# Chemoenzymatic Synthesis of Asymmetric Bisecting Bi-, Tri-, and Tetra-Antennary *N*-Glycans

**DOI:** 10.1101/2025.06.26.661710

**Authors:** Balasaheb K. Ghotekar, Seema K. Bhagwat, Pradeep Chopra, Thomas Buckley, Geert-Jan Boons

**Affiliations:** Complex Carbohydrate Research Center, University of Georgia, Athens, Georgia 30602, United States; Department of Chemistry, University of Georgia, Athens, Georgia 30602, United States; Chemical Biology and Drug Discovery, Utrecht Institute for Pharmaceutical Sciences, and Bijvoet Center for Biomolecular Research, Utrecht University, 3584 CG Utrecht, The Netherlands

## Abstract

*N*-Acetylglucosaminyltransferase-III (GnT-III) is a glycosyltransferase that can install a β1,4-linked *N*-acetylglucosamine (GlcNAc) residue at the central β-mannoside of *N*-glycans. The resulting so-called bisecting GlcNAc is not further extended by glycosyl transferases and has been implicated a wide range of biological processes. The molecular mechanisms by which bisection modulates the biosynthesis of *N*-glycans and influences molecular recognition is not well understood, which is due to a lack of well-defined *N*-glycans with and without bisection. We describe a chemoenzymatic methodology that can readily provide a wide range of asymmetrical bisecting bi-, tri- and tetra-antennary *N*-glycans. It was found GnT-III can act on bi-, tri- and tetra-antennary *N*-glycans and can also accepts *N*-glycans having a β1,2GlcNTFA or GlcN3 moiety at the α1,2Man- or α1,6Man-antenna making it possible to prepare panels of asymmetrical *N*-glycans with and without bisection and having different patterns of sialylation and fucosylation. Kinetic experiments showed GnT-III preferentially modifies bi-antennary glycans. The compounds were printed as a glycan microarray, which was screened for lectin binding. It was found that some lectins preferentially bind to bisecting glycans, whereas others do not tolerate or are not affected by this modification. We investigated receptor specificities of human H1N1 and H2N3 influenza viruses and animal H5N1 viruses that pose a pandemic threat including a virus that has become endemic in cattle. The H1N1 and H2N3 viruses did not tolerate bisection whereas it did not affect H5N1 viruses. A/bovine had the broadest receptor specificity providing a rationale for its wide host range.

## INTRODUCTION

Glycans are the most prominent post-translational modification of proteins in terms of complexity and diversity.^1,2^ These biomolecules encode information by an ability to recruit, in a context dependent manner, regulatory proteins. Glycans are important for protein folding, cell signaling, fertilization, embryogenesis, neuronal development, hormone activity and the proliferation of cells and their organization into specific tissues.^3–6^ Glycans are involved in the etiology of almost every human disease such as pathogen recognition, inflammation, neurological disorders and the development of autoimmune diseases and cancer.^7,8^ Advances in the understanding the biological roles of specific glycans, along with the factors that influence or alter their functions, will provide avenues for the development of therapeutics, diagnostics and nutraceuticals.^9^

The biosynthesis of glycans is a non-template mediated process that occurs in the secretory pathway where glycosyl transferases catalyze the transfer of monosaccharides from sugar-nucleotides to specific hydroxyls of a growing oligosaccharide chain. The complexity of *N*-glycans arises from the modification of a common core pentasaccharide by mannosyl-glycoprotein *N*-acetylglucosaminyltransferases (GnTs) resulting in oligosaccharides that have various numbers and patterns of branching *N*-acetylglucosamine (GlcNAc) moieties (Figure 1A). Galactosyltransferases (GalT) can convert these GlcNAc residues into *N*-acetyl lactosamine (LacNAc) that can then be further modified by other glycosyltransferases into complex epitopes. GnT enzymes have strict substrate requirements. GnT-II and IV require a terminal GlcNAc at the GnT-I position, whereas GnT-V needs a terminal GlcNAc at the GnT-II arm.^10^

*N*-Acetylglucosaminyltransferase-III (GnT-III, EC 2.4.1.144) is the biosynthetic enzyme that installs a β1,4-linked GlcNAc moiety at the core β-mannosyl residue, which has been designated as bisecting GlcNAc (Figure 1A). Unlike other branching GlcNAc moieties, it is not elaborated by glycosyltransferases into complex structures and remains a terminal residue. *N*-glycans modified by bisecting GlcNAc are abundantly observed in brain and kidney tissues, where they modulate various biological functions and disease processes.^11^ Mice deficient in the *Mgat3* gene, which encodes the GnT-III protein, showed improved AD pathology with reduced amyloid-plaque formation.^12,13^ The lack of bisecting GlcNAc caused relocation of the amyloid-producing enzyme, beta-site APP-cleaving enzyme-1 (BACE1) from early endosomes to lysosomes.^14,15^ Bisecting GlcNAc has also been implicated in cancer and mice that lack *Mgat3* display develop faster polyoma middle T (PyMT)-induced mammary tumors, and higher incidence of early metastasis to lung.^16^ Aberrant modification of *N*-glycans with bisecting GlcNAc in cancer makes it a potential biomarker for diagnosis and prognosis.^6^

Tissues of mice lacking a functional *mgat3* gene display *N*-glycans having higher levels of terminal glycan modifications such as sialylation, which indicates that bisecting GlcNAc exerts an influence on other antenna ensuing glycosyltransferases.^17–21^ Upregulation of bisecting GlcNAc reduces galectin binding, probably by inhibition the biosynthesis of poly-*N*-LacNAc residues.^5^ Bisecting GlcNAc has an impact on conformational properties of *N*-glycans and adopt a back-fold conformation in which the α1,6-mannoside is folded to the reducing end.

Despite the importance of bisecting GlcNAc, the biology of this modification at a molecular level is still poorly understood. Bisecting GlcNAc can influence interactions with glycan binding proteins by altering the biosynthesis of terminal epitopes.^21^ It is also possible that a bisecting GlcNAc moiety directly influences binding of glycan binding proteins by altering the glycan conformation or function as a recognition element. Panels of well-defined *N*-glycans with and without bisecting GlcNAc are needed to unravel, at a molecular level, its influence on biological recognition.

The chemical and chemoenzymatic synthesis of bisected *N*-glycans has received little attention. Bisecting *N*-glycans have been prepared by chemical synthesis of a core oligosaccharide followed by enzymatic installation of terminal epitopes.^22–25^ These approaches are very time consuming and have been limited to the preparation of symmetrical bi-antennary *N*-glycans.^26^ Recombinant GnT-III has been employed to introduce bisecting GlcNAc, however, this approach has only be applied to the preparation of biantennary glycans.^27,28^ Currently, no methods have been described to prepare, in a systematic manner, asymmetric multi-antennary bisecting *N*-glycans.

Although bisection interferes with branching enzymes,^17,29–32^ glycomic analysis has shown the presence of bisecting tri- and tetra-antennary glycans indicating that such compounds are biosynthetically feasible. Furthermore, most *N*-glycans have asymmetrical architectures in which the various antennae are modified by unique epitopes. Several studies have indicated that the presentation of an epitope at a specific antenna can influence molecular recognition.^33^ It is thus imperative that methods become available to prepare asymmetrical bisecting multi-antennary *N*-glycans.

Previously, we described a stop-and-go chemoenzymatic strategy for the preparation of asymmetrical *N*-glycans.^34^ It employs a glycopeptide isolated from egg yolk powder that in five chemical and enzymatic steps can be converted into biantennary glycan **1**. Next, recombinant GnT-V and unnatural UDP-2-deoxy-2-trifluoro-*N*-acetamido-glucose (UDP-GlcNTFA, Figure 1B) was employed to transform **1** into a tri-antennary glycan. The trifluoroacetyl (TFA) moiety could easily be removed by aqueous ammonia and the resulting free amine transformed into azide by an azido transfer reaction. Another antenna could be installed employing GnT-IV in the presence of UDP-GlcNTFA and treatment of the resulting product with aqueous ammonia gave a tetra-antennary *N*-glycan. We exploited that GlcNH2 and GlcN3 are resistant to modification by relevant glycosyltransferases, however, at an appropriate stage of synthesis these unnatural monosaccharides can sequentially be converted into natural GlcNAc for selective enzymatic elaboration into complex appendages thereby giving entry into asymmetrical multi-antennary *N*-glycans (Figure 1C). We have also demonstrated that A2-Asn can be further trimmed to a core pentasaccharide (Man3GlcNAc2) that can be modified by GnT-I and GnT-II in the presence of UDP-GlcNTFA or UDP-GlcNAc to give access to asymmetrical biantennary glycans.^35^

Here, we describe a *stop-and-go* methodology for the preparation of asymmetrical bisecting bi-, tri- and tetra-antennary *N*-glycan. We found that recombinant GnT-III can act on bi-, tri- and tetra-antennary *N*-glycans to install a bisecting GlcNAc moiety. On the other hand, a bisecting bi-antennary glycan could not be further branched by GnT-IV and GnT-V, and thus the order of introducing branching points is important for the synthesis of multi-antennary bisecting *N*-glycans. It was also discovered that GnT-III can act on glycans having a β1,2GlcNTFA or GlcN3 moiety at the α1,2Man- or the α1,6Man-antenna opening opportunities to prepare asymmetrical bisecting multi-antennary *N*-glycans. GnT-IV and GnT-V did, however, not tolerate a GlcN3 at α1,2-Man or the α1,6-Man antenna, highlighting that these enzymes have different substrate requirements. Collectively, the finding made it possible to prepare a library of asymmetrical *N*-glycans with and without bisection (Figure 1D). We focused on the preparation of compounds having a terminal α2,3- and α2,6-sialyl and a sialyl Lewis^x^ epitope. The compounds were printed as a glycan microarray, which was screened for lectin binding. It was found that PHA-E preferential binds to bisecting glycans whereas Con A, SNA, MAL-I, GSL-II does not tolerate this moiety. Several lectins including AAL, WGA, ECL were not affected by bisection. We employed the newly developed glycan microarray to investigate receptor specificities of human H1N1 and H2N3 influenza viruses and animal H5N1 viruses that pose a pandemic threat including a virus that has become endemic in cattle in North America.^36^ It was found that the H1N1 and H2N3 viruses did not tolerate bisection. This modification did, however, not affect the H5N1 viruses. Furthermore, it was found that A/bovine/Ohio/B24OSU-432/2024 (H5N1) has broader receptor specificity than the evolutionary earlier A/Vietnam/1203/2004 (H5N1) and requires only one SLe^x^ moiety on an *N*-glycan for potent binding.

**Figure 1.**
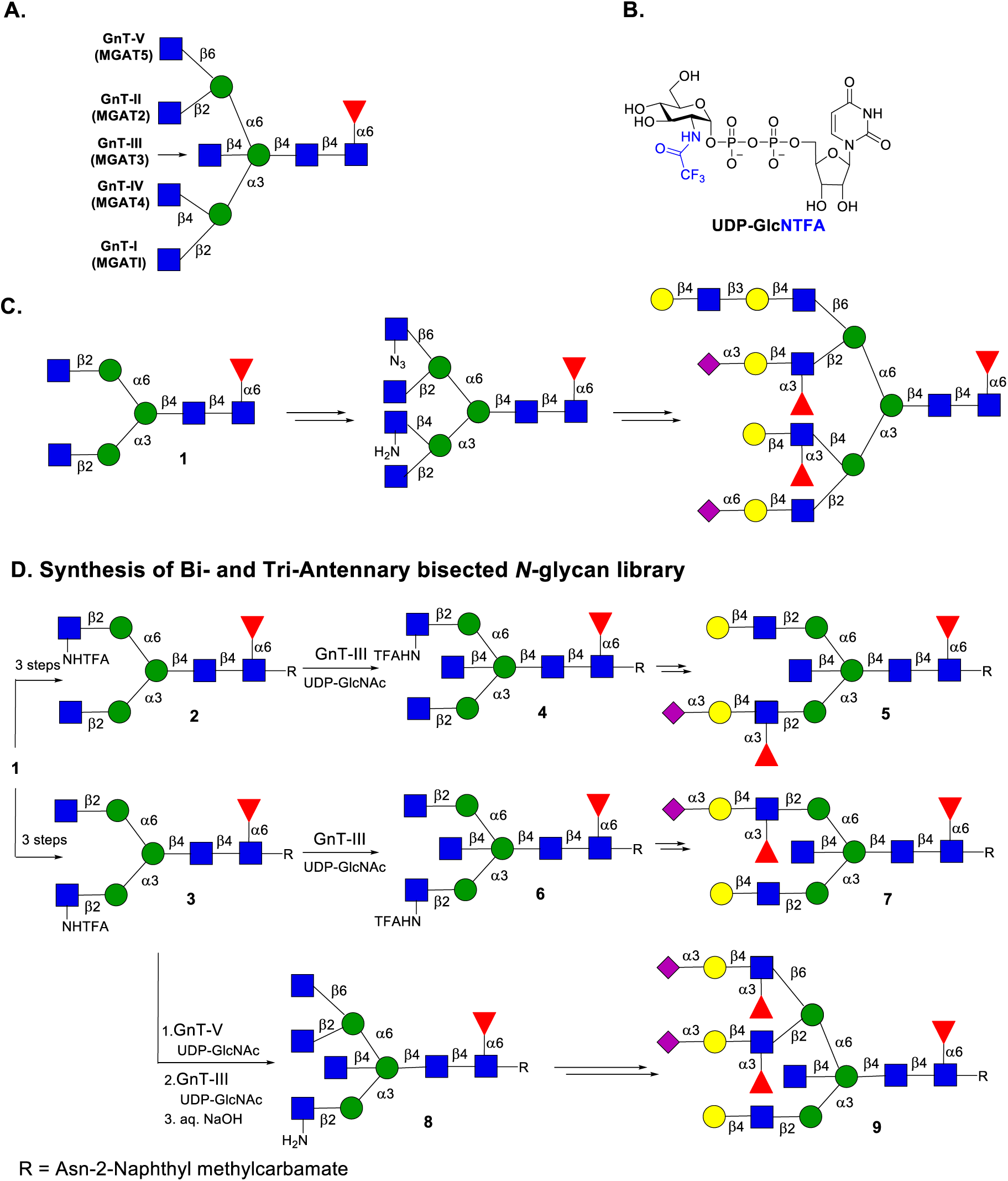
Representative structure of *N*-glycans and stop-and-go strategy for synthesis of multi-antenna and bisected asymmetrical *N*-glycans. A) GnT enzymes responsible for the addition of branching GlcNAc moeities. B) examine of asymmetrical branched B-glycans C) stop and go strategy for the synthesis of asymmetrical *N*-glycans. D) Structure of UDP-GlcTFA. E) Compound **2** and **3** was synthesized from SGP, modified as bisected glycan **4** and **6** with UDP-GlcNAc and GnT-III to access asymmetrical biantennary glycans such as **5** and **7**, whereas **3** with upon addition with GnT-V and GnT-III arm, followed by TFA removal as stop point is appropriate starting material for bisected multiantenna, complex glycans like **9**.

## RESULT AND DISCUSSION

### Substrate Specificity of GnT-III

Several observations have indicated that bisecting GlcNAc can interferes in the activity of the branching enzymes GnT-IV and GnT-V.^17,18,32^ Thus, multi-antennary bisecting *N*-glycans are likely biosynthesized by the action of GnT-IV and GnT-V to give higher branched *N*-glycans that are then modified by GnT-III to install bisection. To examine the substrate tolerance of GnT-III, we prepared bi- (**1**), tri- (**10**, **11**) and tetra- (**12**) antennary *N*-glycans to examine if they can be modified by GnT-III (Scheme 1A). The preparation of these compounds started from glycosylated amino acid **S2** (SI, section 2b) that was readily obtained by subjecting a sialoglycopeptide (SGP) isolated from egg yolk to a three-step procedure entailing hydrolysis of the sialosides and galactosides by the neuraminidase from *Clostridium perfringens* and galactosidase from *Aspergillus niger* followed by pronase treatment to remove the peptide leaving an anomeric asparagine (Asn).^34^ The a-amine of Asn of **S2** was protected as a 2-naphthyl methylcarbamate using 2-naphthyl methyl (NAP) chlorocarbonate in aqueous NaHCO3 and finally a core fucoside was installed using recombinant α-fucosyltransferases 8 (FUT8)^37^ and guanosine 5′-diphospho-β-l-fucose (GDP-fucose) to give **1**. The latter compound was subject to either recombinant GnT-IVB or GnT-V in the presence of uridine 5′-diphospho-*N*-acetyl glucosamine (UDP-GlcNAc) and after an incubation time of 18 h, LC-MS analysis showed complete conversion of the starting material into the expected products. The products were purified by bench top C18 reverse phase (RP) column chromatography using a gradient of water and acetonitrile to give homogenous **10** and **11** in yields of 81% and 91%, respectively. Tetra-antennary *N*-glycan **12** was obtained in a yield of 93% by subjecting **11** to GnT-IVB in the presence UDP-GlcNAc followed by purification by C18 RP column chromatography.

Next, we explored whether **10, 11** and **12** can be converted into the corresponding bisecting structures **13**, **14**, and **15**, respectively by treatment with recombinant GnT-III in the presence of UDP-GlcNAc. Gratifyingly, the transformations proceeded readily and were completed within a period of 18 h providing the target compounds **13**, **14**, and **15** in high yields. All compounds were fully characterized by 1D and 2D NMR and MS.

To validate that GnT-IVB and GnT-V cannot act on bisecting structures, compound **1** was converted into the corresponding bisecting derivative **16** by treatment with GnT-III, which was followed by exposure to GnT-IVB and GnT-V in the presence of UDP-GlcNAc. Even after a prolonged reaction times, LC-MS did not show any product formation highlighting that bisecting multi-antennary glycans are biosynthesized by first branching by GnT-IVB and GnT-V followed by modification by GnT-III (Scheme 1B).

**Scheme 1.**
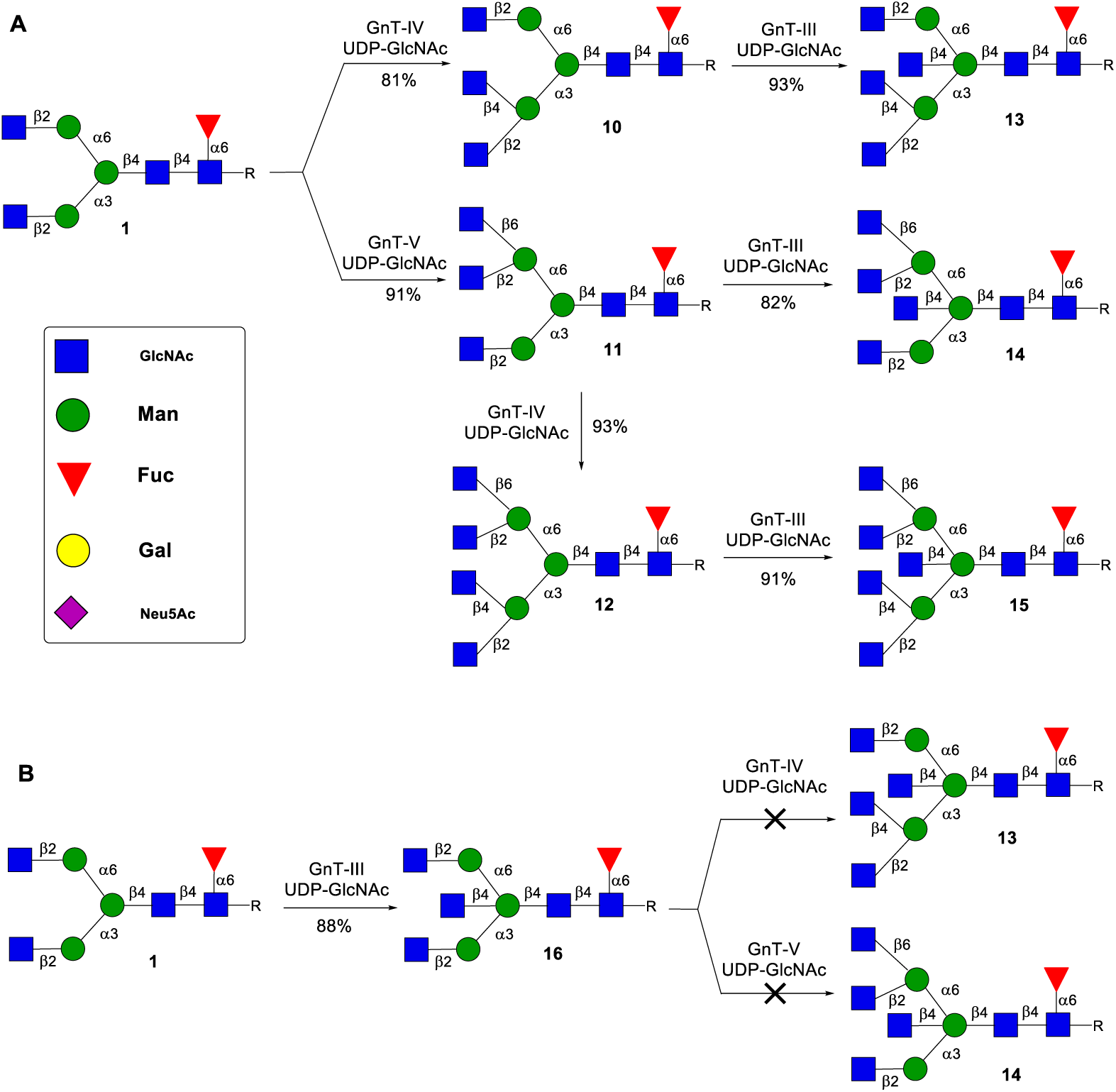
substrate specificity of A) tri- and tetra-antennary *N*-glycans **10**, **11**, and **12** and B) bisecting glycan **16**.

Compounds having bisecting GlcNAc have well separated H-1 at 4.39 ppm for this residue established by ^1^H and ^1^H-^13^C HSQC experiments. All other protons were assigned by using ^1^H-^1^H COSY and ^1^H-^1^H TOCSY experiment. H-4 of bisecting GlcNAc appeared at an upfield region (triplet, δ 3.18-3.16 with *J* = 9.3 Hz) as compared to other protons which was confirmed by COSY and TOCSY 2D experiments. Representative NMR data for compound **16** are shown in Figure 2. The reducing GlcNAc-H1 (GlcNAc-1) could be assigned (δ 4.85) due to a substantial upfield shift of its carbon value (δ 78.1) resulting from the β-linked amide. The signals at δ 4.48 and δ 4.47 integrated to a value of two corresponding to GlcNAc-3 and -5 (GnT-I and GnT-II antennae) as these monosaccharides occupy a similar electronic environment and both have β-anomeric linkages. GlcNAc-2 showed a small characteristic downfield shift at δ 4.54. The characteristic bisecting GlcNAc appeared upfield at δ 4.38. After the assignment of H-1 of GlcNAc-1, a COSY correlation showed the coupling with H-2 at δ 3.61, which in turn correlated with H-3 at δ 3.33. H-4 was confirmed by the TOCSY experiment and appeared at δ 3.19 (for complete characterization and correlation see NMR section of SI). H-4 showed correlations with to H-5 at δ 3.49.

**Figure 2.**
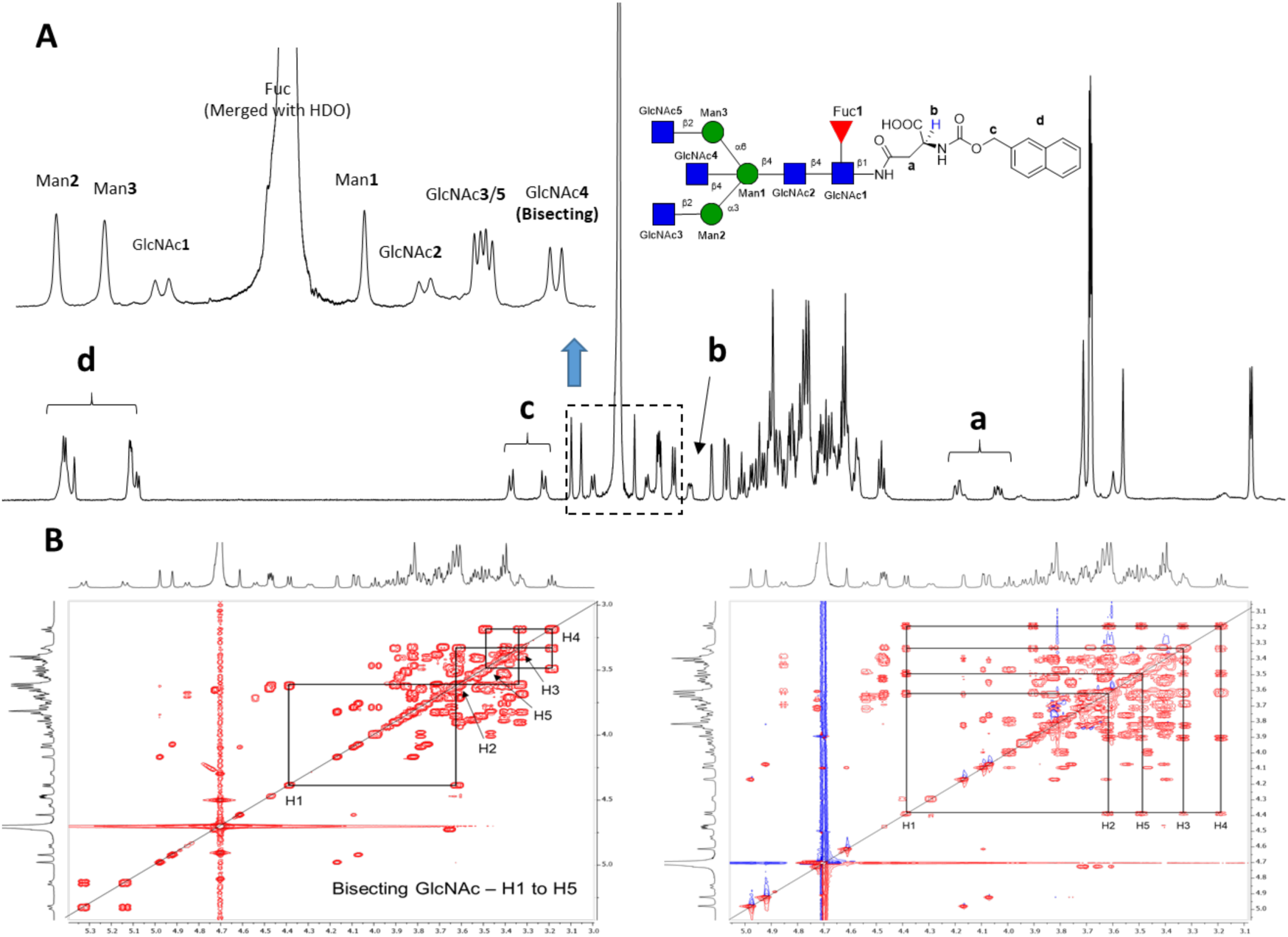
NMR analysis of bisected *N*-glycan **16**. A) Full ^1^H NMR and Anomeric proton signals of **16**. B) Assignment of H-1 to H-5 proton using ^1^H-^1^H COSY and ^1^H-^1^H TOCSY experiments.

### Tolerance of GnT-III to Substrates having a GlcNTFA or GlcN3 Moiety at the α1,3- or 1,6-Antenna

Next, we focused on investigating whether GnT-III can modify *N*-glycans having an unnatural glucosamine moiety at one of the antennae. The stop-and-go strategy^34^ (Scheme 2A) relies on the presence of GlcNH2 or GlcN3 moiety at one of the branching points to temporarily block enzymatic extension by other glycosyl transferases (Figure 1C). Furthermore, we have found that bisection should be the last branching point to be installed. Thus, to extend the stop-and-go strategy to the preparation of bisecting asymmetrical *N*-glycans, it is imperative that GnT-III can accepted *N*-glycans having an unnatural glucosamine at various antennae. Thus, we investigated whether a β1,2-GlcNTFA or β1,2-GlcN3 moiety is tolerated when present at the α1,3- or α1,6-antenna. For this purpose, we prepared compounds **2**, **3**, **17**, and **18** for testing as possible substrates for GnT-III. Thus, compound **1** was treated with β-*N*-acetylglucosaminidase S to give Man3GlcNAcFuc-Asn-NAP that subsequently was treated with GnT-I in the presence of UDP-GlcNTFA and then with GnT-II and UDP-GlcNAc to give **3** (See Scheme S1). The corresponding azido derivative **17** was synthesized by treatment of **3** with aqueous sodium hydroxide (pH = 10) followed by an azido transfer reaction using potassium carbonate (K2CO3) and imidazole-1-sulfonyl azide (ImSO2N3)^38^ (See Scheme S2). Compounds **2** and **18** were prepared by a similar strategy, however, in this case UDP-GlcNAc was employed in combination GnT-I and UDP-GlcNTFA for the GnT-II catalyzed conversion.

Gratifyingly, exposure of TFA containing *N*-glycans **3** and **2** to GnT-III and UDP-GlcNAc resulted in full conversion into the expected products **6** and **4**, respectively. A GlcN3 moiety at the α1,3- or α1,6-antenna was also tolerated by GnT-III allowing the conversion of **17** and **18** into **19** and **20**, respectively. Interestingly, compounds **17** and **18** were not converted by GnT-IVB and GnT-V into the corresponding products highlight different substrate requirements of the different branching enzymes (Scheme S3).

Next, attention was focused on the preparation of bisecting tri-antennary glycans **21** and **22** starting from **2** and **3** employing a GnT enzyme to install an additional branching point. Thus, acceptors **2** and **3** were subjected to GnT-IVB or GnT-V, respectively in the presence of UDP-GlcNAc to furnish tri-antennary glycan intermediates, which were subjected to GnT-III resulting in the formation of bisecting tri-antennary glycans **21** and **22**, respectively (Scheme 2B).

**Scheme 2.**
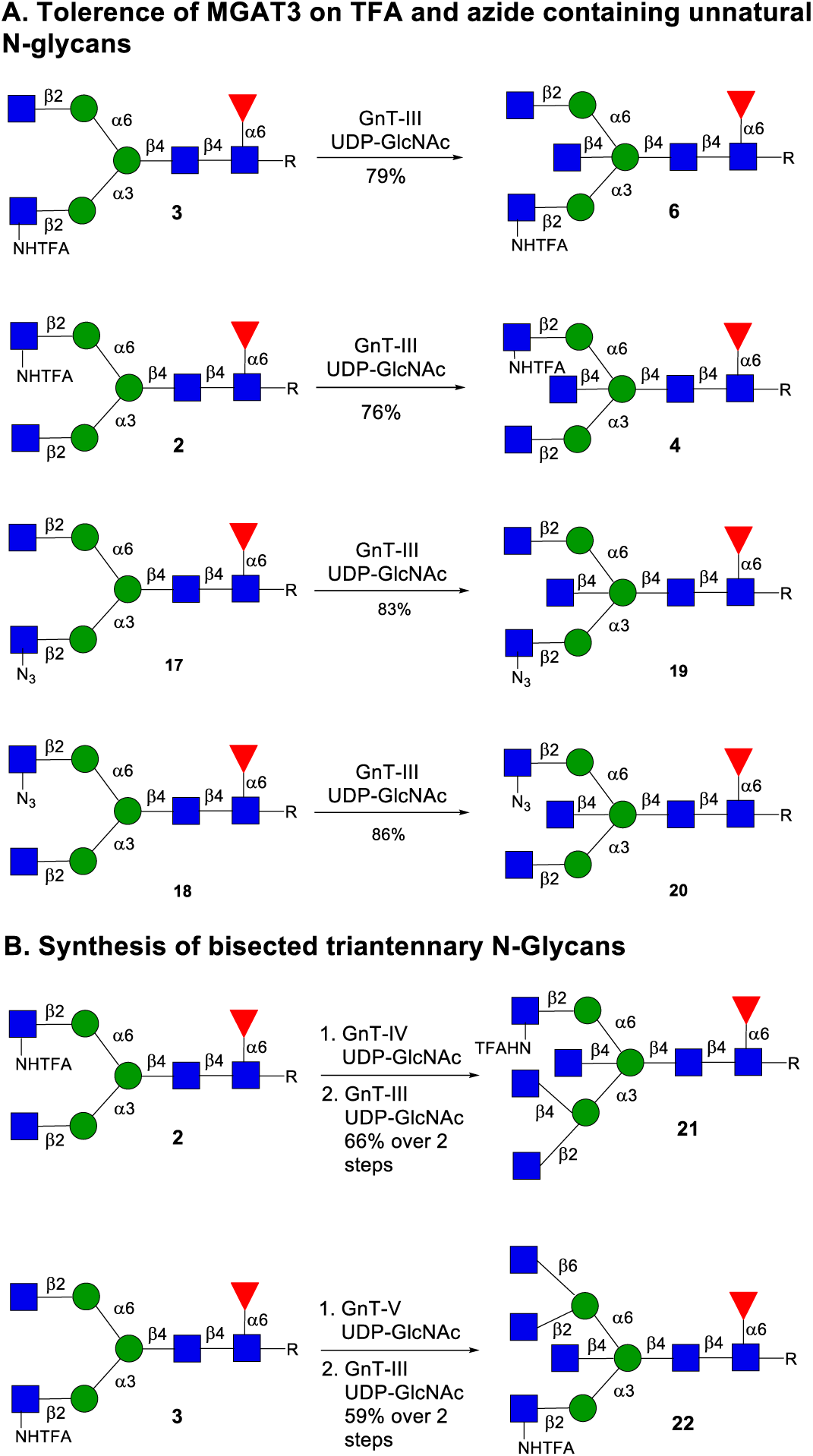
A) Tolerance of GnT-III for unnatural *N*-glycans **2**, **3**, **17**, and **18**. B) Synthesis of bisecting tri-antennary *N*-glycans **21** and **22**.

With compound **6** in hand, we explored the preparation of asymmetrical bisecting *N*-glycans. To block the GnT-I arm from enzymatic conversion, compounds **6** was treated with aq. NaOH at pH = 8 to hydrolyze the TFA moiety to give GlcNH2 containing glycan **23**. The latter compound contains a natural GlcNAc residue at the α1,6-antenna, which was galactosylated with β1,4-galactosyltransferase-1 (B4GalT1) and uridine 5′-diphosphogalactose (UDP-Gal) to install a LacNAc motif to give asymmetrically branched glycan **24**. Next, acetylation of the amine of **24** using Ac2O in the aqueous NaHCO3 gave **25**. The latter compound was sialylated with either ST3 β-galactoside α2,3-sialyltransferase 4 (ST3Gal4) to form α2,3-linked sialoside **26** or ST6 β-galactoside *α*2,6-sialyltransferase 1 (ST6GAL1) to give α2,6-linked sialoside **27**. Compound **26** was further *α*1,3-fucosylated using α-fucosyltransferase 6 (FUT6) to give SLe^x^-containing bisecting *N*-glycan **28**. The GlcNAc moiety at the GnT-I antenna of compounds **26** and **28** were galactosylated by recombinant B4GalT1 and UDP-Gal to produce LacNAc-containing derivatives **29** and **7**, respectively (Scheme 3).

**Scheme 3.**
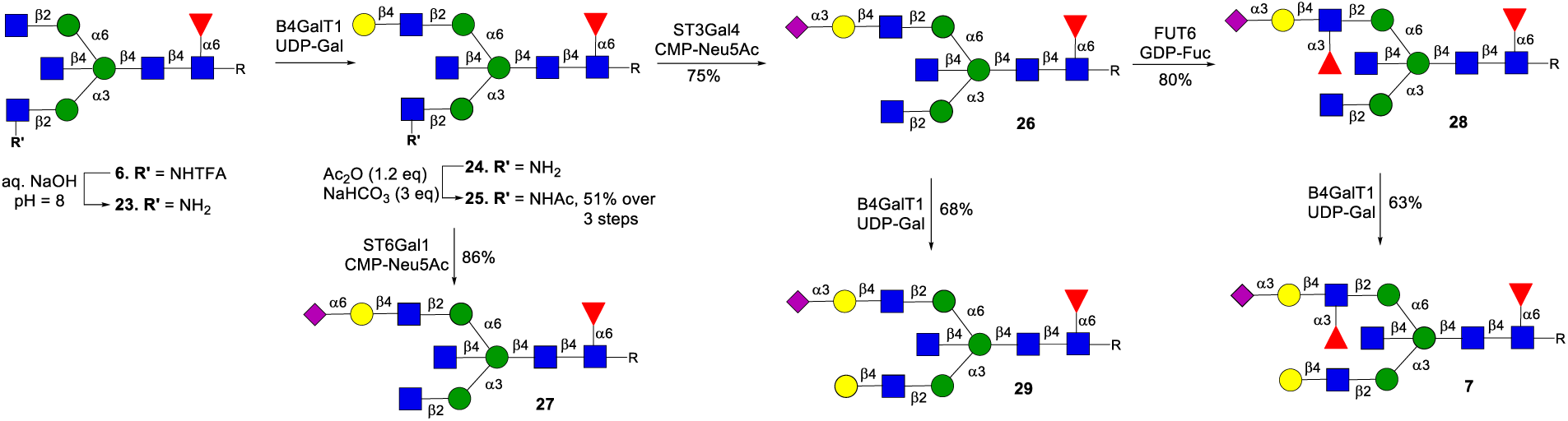
Chemoenzymatic synthesis of core fucosylated asymmetrical bisected *N*-glycans by stopping GnT-I arm.

A similar strategy was employed to prepare a range of isomeric compounds but in this case the synthesis started from **4**, which has a GlcNAc moiety at the α1,3-arm and a GlcNTFA derivative at the α1,6-antenna (Scheme 4). Treatment of **4** with base gave **30** having GlcNH2 at the α1,6-antenna disabling it from enzymatic modification. The natural terminal GlcNAc moiety of **30** was converted into LacNAc by B4GalT1 to give **31** which was *N*-acetylation to provide **32**. The latter compound was converted into **33**, **34**, **35**, **36**, and **5** by similar manipulations as described above.

**Scheme 4.**
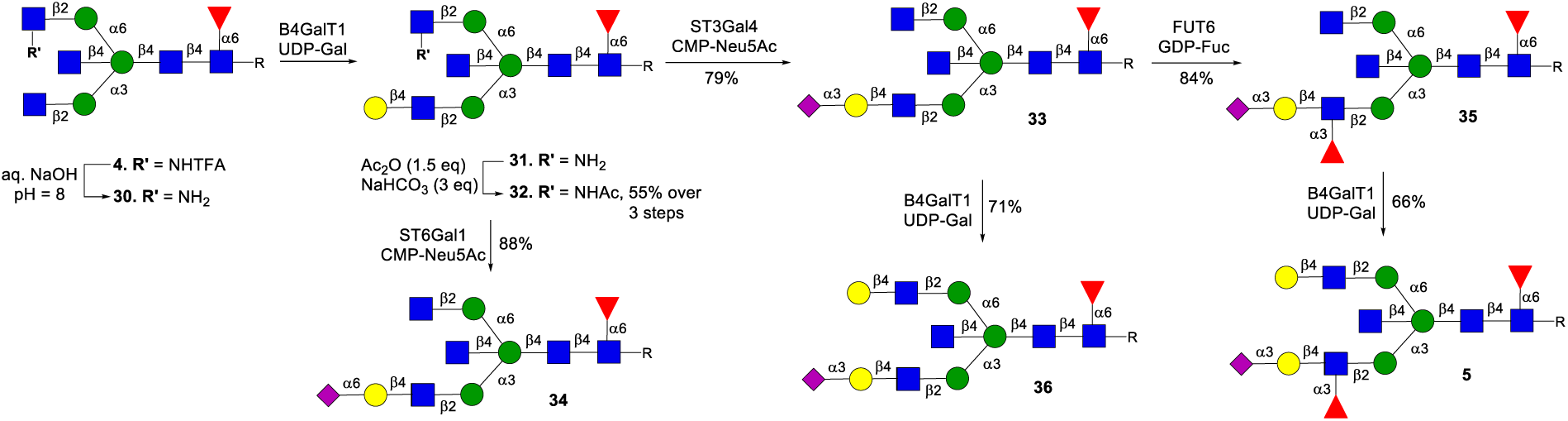
Chemoenzymatic synthesis of core fucosylated asymmetrical bisected *N*-glycans by stopping GnT-II arm.

### Preparation of Bisecting Multi-Antennary *N*-Glycans

We prepared bisecting *N*-glycans **38**-**41** and **9** to demonstrate the versatility of the approach (Scheme 5). Thus, the NHTFA moiety of **22** was saponified by aqueous NaOH, and then the natural GlcNAc residues at the GnT-II and GnT-V arms of the resulting compound **8** were galactosylated with B4GalT1 in the presence of UDP-Gal to provide **37**. Next, the free amine of **37** was acetylated to give **38**. The LacNAc moieties were selectively sialylated with ST3Gal4 providing **39** that was further fucosylated by FUT6 to afford SLe^x^-containing structure **40**. This step exploits that FUT6 acts on LacNAc but not on terminal GlcNAc moieties. The GnT-I arm of **39** and **40** was further extended by a galactoside using B4GalT1 to furnish complex multi-antennary glycans **41** and **9**, respectively.

**Scheme 5.**
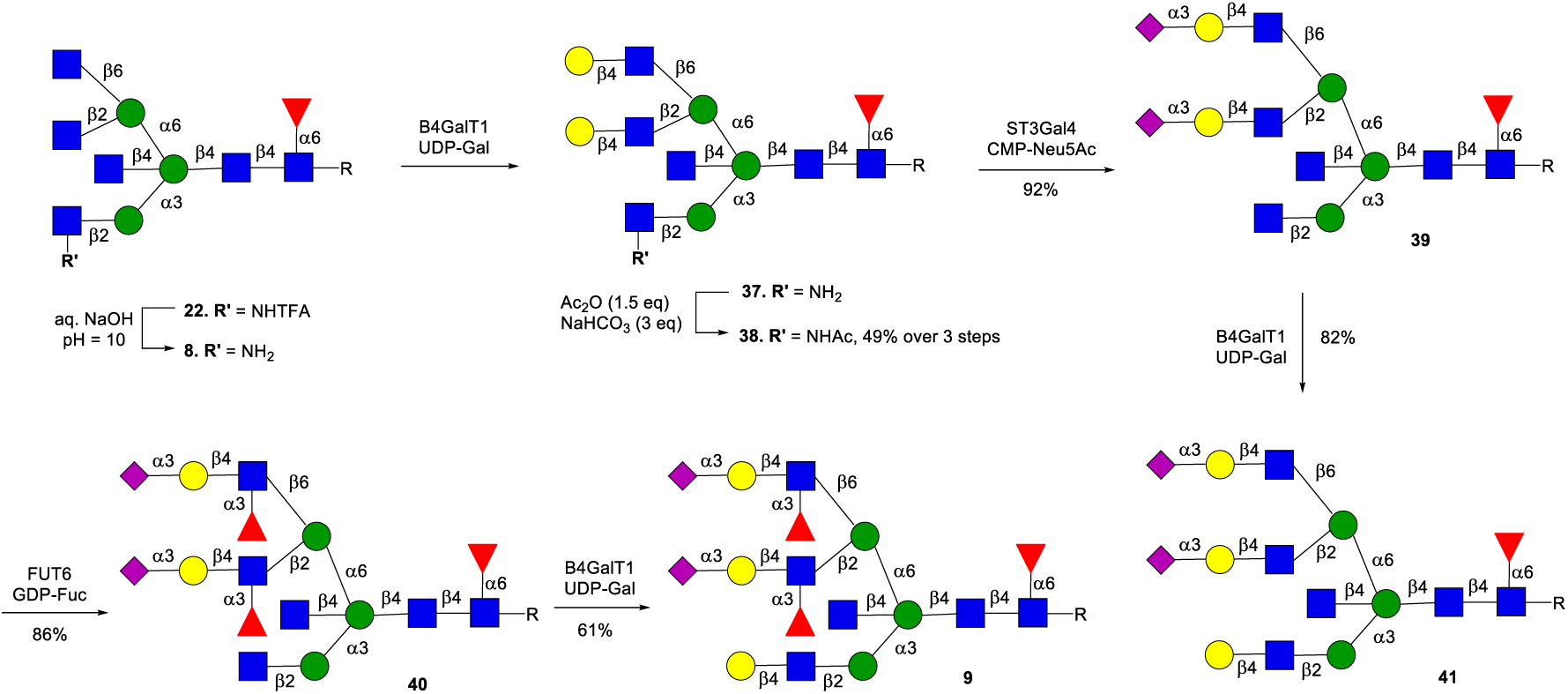
Synthesis of asymmetric branched tri-antennary bisected *N*-glycans.

### Kinetic Parameters of GlcNAc Transfer by GnT-III, GnT-IV and GnT-V for Multi-Antennary *N*-glycan Biosynthesis

Glycomic analyses have shown that bisecting multi-antennary glycans occur naturally.^39,40^ Furthermore, we found that bi-(**1**), tri-(**10**, **11**) and tetra-antennary (**12**) glycans can be modified by GnT-III to give the corresponding bisecting *N*-glycans. To obtain further insight into the biosynthesis of *N*-glycans and establish how readily various glycans are modified by bisection, we determined kinetic parameters of GlcNAc transfer to **1**, **10**, **11** and **12** by GnT-III using a UDP-Glo^®^ assay (Promega). Thus, different concentrations of the synthetic glycans (2000 – 31.25 μM), ultrapure UDP-GlcNAc (1000 μM) and recombinant GnT-III were incubated for 1 h and the formation of UDP was determined by UDP-Glo® assay (see Table 1 and Figure S62).^41^ It was found that bi-antennary glycan **1** is a substantially better substrate than the higher branched structures **10**, **11** and **12**. A GlcNAc installed by GnT-IV appears to be better tolerated than a similar modification by GnT-V. Furthermore, mono-antennary *N*-glycan **S17** is an appropriate substrate for GnT-III whereas isomeric structure **S18** was not converted into the corresponding bisecting *N*-glycan. Also, we found that Man3Glc2 is not a substrate for this enzyme. These results demonstrate that the 1,2-linked GlcNAc installed by GnT-I is critical for GnT-III activity.

Kinetic parameters were also obtained for the donor substrate (UDP-GlcNAc). Different concentrations of UDP-GlcNAc (6000 – 66.25 µM), acceptor (400 µM) and recombinant GnT-III were subjected to the same assay format. It was found that the bi- (**1**), mono- (**S17**), and tetra- (**12**) multi-antennary *N*-glycans have lower Km values for UDP-GlcNAc than that of the tri-antennary *N*-glycans (**10**, and **11**).

Next, we determined kinetic parameters for GlcNAc transfer catalyzed by GnT-IVB and GnT-V to bi-antennary glycan **1**. Due to the reported low affinity of these enzymes for the donor substrate,^42,43^ 7 mM and 10 mM of UDP-GlcNAc were employed, respectively. These glycosyl transferases have substantially higher Km values for the acceptor substrate compared to GnT-III.

GnT-III is compartmentalized in the *trans*-Golgi while GnT-I, II, IV and V are found in the *medial*-golgi.^44^ The low affinity of the *N*-glycans and sugar donors for GnT-IVB and GnT-V coupled with GnT-III being localized in a separate compartment provides an explanation of the higher abundance of bisecting biantennary *N*-glycans.^45^

**Table 1.**
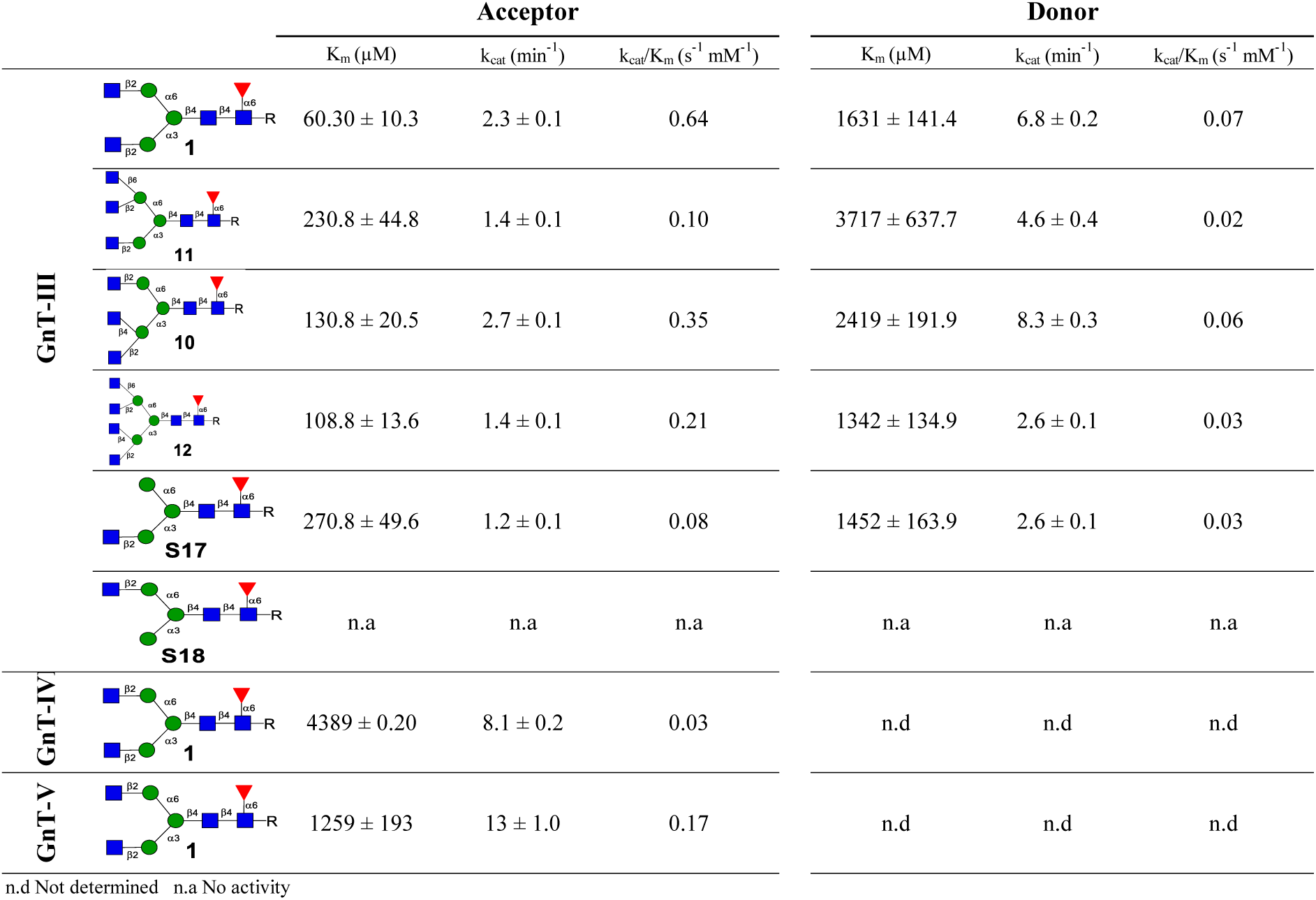
Kinetic parameters of GlcNAc transfer catalyzed by GnT-III, GnT-IV and GnT-V via UDP-Glo assay. Acceptor = **1**, **10**, **11**, **12**, **S17**. Donor = UDP-GlcNAc. Kinetic values Kcat, Km and Vmax were calculated by nonlinear curve fitting in GraphPad Prism 6.

### Glycan Microarray Development

To systematically examine the influence of *N*-glycan bisection on protein binding, we prepared a range of control structures that lack a bisecting GlcNAc moiety following procedures illustrated in Schemes S4 and S5. All synthetic compounds have an anomeric asparagine (Asn) moiety in which the α-amine is protected as a naphthyl methyl (NAP) carbamate. The latter protecting group was readily removed by hydrogenation of **16**, **25**-**29**, **7**, **32**-**36**, **5**, **38**-**41**, and **9** over Pd/C in a mixture of *tert*-butanol/water (*t*BuOH/H2O, 1/1, v/v) to afford compounds **A**-**R** and **S11**, **S12**, **S15**, **S16** to **E’**, **F’**, **K’**, and **L’**, respectively (Figure 3A). The resulting α-amine facilitated printing on *N*-hydroxysuccinimide (NHS) activated glass slides using a non-contact microarray printer. The glycans are arranged in an increasing order of complexity; first biantennary glycans with terminal modifications such as galactosylation, sialylation and fucosylation on the β1,6-antenna (**A**-**F’**), biantennary having a complex extension at the β1,3-antenna (**G**-**L’**), followed by tri-antennary glycans (**M**-**Q**) and finally control structures (**R**-**T**). To confirm the robustness of the platform, we employed *Phaseolus vulgaris* E (PHA-E), which is a prototypical lectin that binds bisecting *N*-glycans having a terminal galactosyl moiety.^46^ Thus, a subarray was incubated with biotinylated lectin in binding buffer at room temperature for 1 h, after which the slide was washed, dried, and then incubated with streptavidin-Alexa Fluor®647 to detect lectin binding. As anticipated, all compounds having bisection and a terminal Gal moiety showed strong responsiveness (Figure 3). Promiscuity for structures with multiple Gal residues (**S** and **T**) was observed. Addition of a fucoside to LacNAc at the α1,6GlcNAc arm (**D** *vs.* **F**) abolished PHA-E binding.^47^ *Concanavalin* A (Con A) is another lectin exhibiting sensitivity to bisection and bound preferentially non-bisecting α-linked mannosyl residues. The presence of bisecting GlcNAc introduces a conformation change which probably is responsible for weak binding (**A** *vs.* **A**’, **C** *vs.* **C**’, **D** *vs.* **D**’ etc).

The fucosyl recognizing lectin, *Aleuria aurantia* lectin (AAL) bound to fucosylated *N*-glycans irrespective of the presence of bisecting GlcNAc. *Sambucus nigra* agglutinin (SNA) lectin preferentially binds α2,6-linked sialosides. It bound strongly to the positive control structure **T** and displayed a clear preference for non-bisecting glycans (**C** *vs.* **C’** and **I** *vs.* **I’**). *Maackia amurensis* lectin I (MAL-I) preferentially bind to α2,3-linked sialosides and responded only to such sialosides lacking bisection (**D** *vs.* **D’** and **J** *vs.* **J’**). Noteworthy, MAL-1 failed to engage SLe^x^ containing structures like **E’**, **F’**, **K’**, and **L’**. *Griffonia simplicifolia* lectin II (GSL-II) and *Datura stramonium* Lectin (DSL) detect glycans having a terminal GlcNAc moiety. It was observed GSL-II binds only strongly to non-bisecting glycans (**A’**, *vs.* **A**, **C’** *vs.* C and **E’** *vs.* **E**) and prefers structures with GlcNAc on the GnT-I arm (**A**’ *vs.* **G’**, **C’** *vs.* **I’**, **E’** *vs.* **K’**). On the other hand, DSL prefers glycans with multiple GlcNAc moieties in the backbone as in **S** and **T** and bound poorly to bi-antennary structures with fewer GlcNAc units. It showed strong binding to bisected tri-antennary glycans (**M**, **N**, **O)**, but their SLe^x^ counterparts **P** and **Q** did not show responsiveness. It is known that *Wisteria floribunda* lectin (WFL) and *Erythrina cristagalli* lectin (ECL) bind to terminal Gal residues.^48^ We observed WFL preferentially binds to glycans having Gal on the β1,3-GlcNAc branching arm and engages better with bisected structures (**A** *vs.* **A’**, **D** *vs.* **D’**, **G** *vs.* **G’**). In contrast, ECL did not differentiate between bisecting and non-bisecting glycans (**D** *vs.* **D’** and **G** *vs.* **G’**) and recognized glycans having a LacNAc moiety on both the GnT-I and GnT-II arm (**D** *vs.* **G**). Finally, to detect the SLe^x^ structures printed on the array, we employed an anti-CD15s monoclonal antibody. It only bound to the tri-antennary glycans **P** and **Q** having at least two SLe^x^ epitopes. The bi-antennary bisecting or non-bisecting glycans having only one SLe^x^ at the GnT-I or GnT-II arm did not show responsiveness.

**Figure 3.**
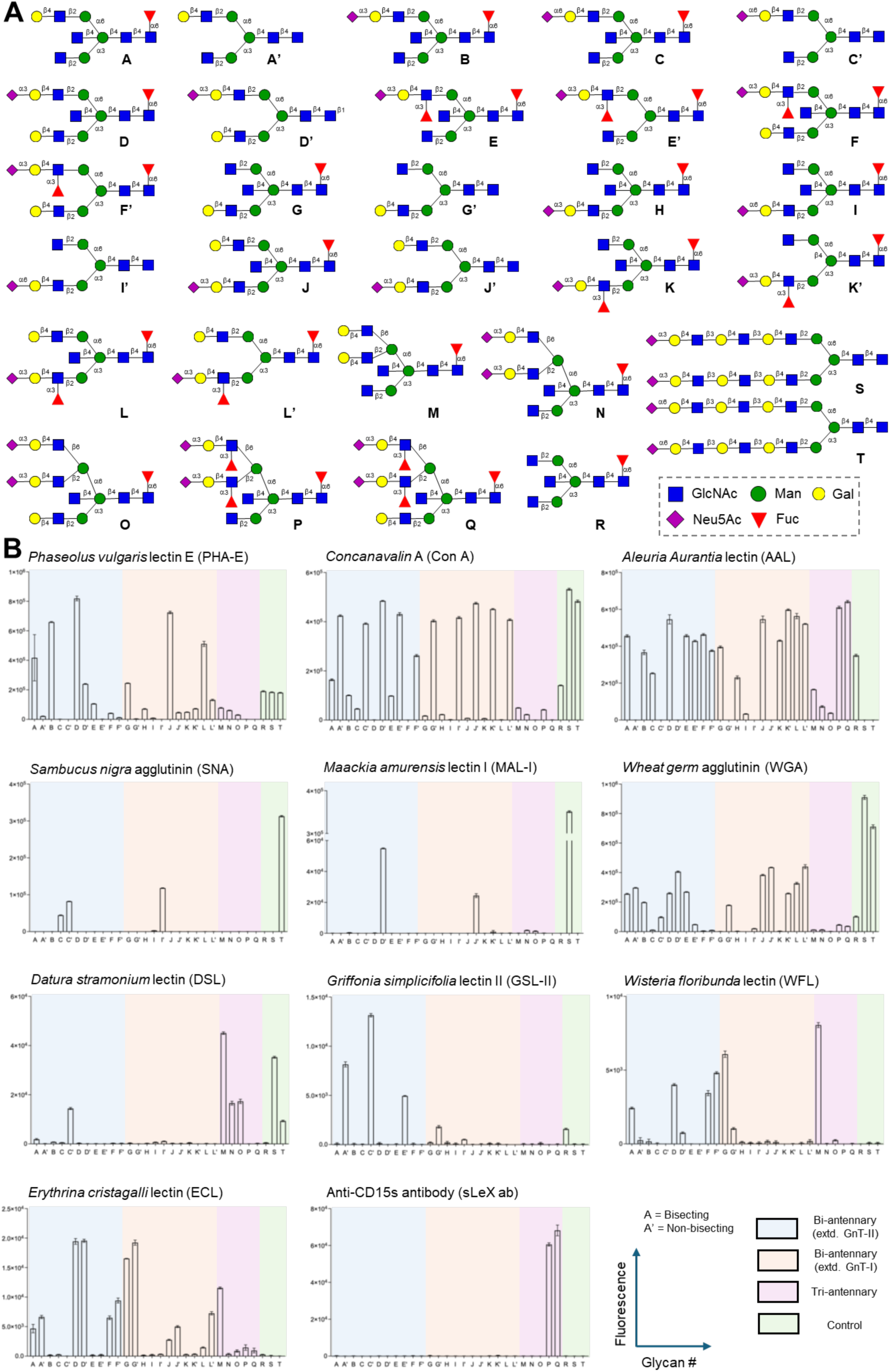
A) Structures of glycans printed on the microarray. All *N*-glycans have an α-amine at the reducing end Asn moiety. B) Binding data for *N*-glycans printed on amine reactive microarray slides for various lectins and CD15s antibody. Experimental details are provided in the SI. Data are presented as mean ± SD (n = 4). Representative data are shown for each lectin, which was repeated at least two times.

### Receptor Specificity of Highly Pathogenic Avian Influenza Viruses

Next, attention was focused on investigating the influence of bisection on influenza A virus (IAVs) receptor specificities. Bisecting glycans have been observed in airway tissues,^49^ and thus it is important to explore whether this modification influences viral binding. Furthermore, there is evidence to support that highly pathogenic avian influenza (HPAI) viruses have broadened their receptor specificity and can bind fucosylated structures.^50,51^ HPAI viruses of the H5Nx A/goose/Guangdong/1/96 lineage have frequent spilled over into mammals.^52^ In 2021, new H5N1 viruses belonging to the 2.3.4.4b hemagglutinin (HA) phylogenetic clade became endemic in wild birds causing widespread infections in poultry. These avian viruses have been detected in several mammalian species including humans, elevating concerns regarding the pandemic potential of these viruses. In early 2024, infections with HPAI H5N1 clade 2.3.4.4b viruses were detected in dairy cattle in Texas (USA) that have spread to multiple farms.^53^ There is evidence for cow-to-cow transmission, interspecies transmissions to birds, domestic cats, and a raccoon and also to dairy cow farmworkers.^54,55^

We and others have found that A/bovine/H5N1 preferentially binds to “avian type” receptors (α2,3-sialosides).^56–58^ Glycan microarray screening indicated that these viruses prefer bi-antennary *N*-glycans having extended LacNAc moieties at both antennae terminating in α2,3-sialosides. Compounds with a sialoside at only one antenna gave low responsiveness. In thes studies, the evolutionary earlier A/Vietnam/1203/2004 (H5N1) virus (BPL-inactivated) gave a similar binding profile for *N*-glycans. The binding selectivity of the two viruses for *O*-glycans differed and A/bovine/OH/B24OSU-432/2024 recognizes *O*-glycans having a sialyl LacNAc moieties modified by an α1,3-fucoside or 6-*O*-sulfate whereas this was not the case for A/Vietnam/1203/2004. Thus, we were compelled to further investigate the binding of these viruses to glycan of the newly developed microarray. We included A/California/04/2009 (pdmH1N1) and A/Perth/16/2009 (H3N2) as examples of human influenza viruses. Thus, subarrays were incubated with α-propiolactone (BPL)-inactivated whole viruses, A/California/04/2009 (pdmH1N1), A/Perth/16/2009 (H3N2), A/Vietnam/1203/2004 (H5N1) and A/Bovine/Ohio/B24OSU-432/2024 (H5N1) at room temperature for 2 h in the presence of a neuraminidase inhibitor (Oseltamivir, 10 μM). After washing and drying, the subarrays were incubated with haemagglutinin anti-stem antibody [1A06 for group-1 (H1N1 and H5N1) and CR8020 for group-2 (H3N2)] and Alexa Fluor®647 anti-human IgG antibody to detect viral binding.

**Figure 4.**
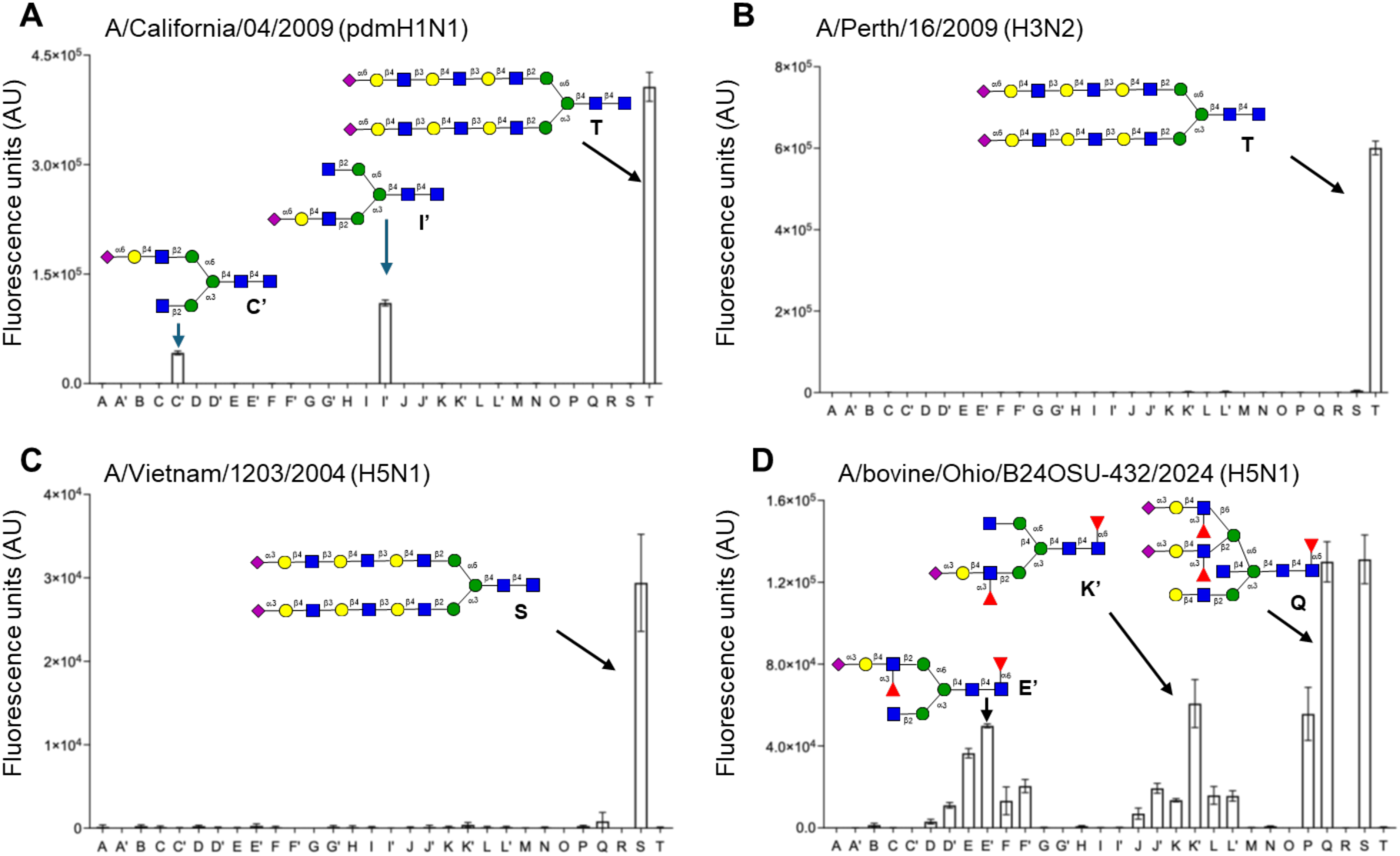
Glycan array binding analysis to determine the receptor-binding specificities of A) A/California/04/2009 (pdmH1N1). B) A/Perth/16/2009 (H3N2). C) A/Vietnam/1203/2004 (H5N1). and D) A/bovine/Ohio/B24OSU-432/2024 (H5N1) viruses. Data are presented as mean ± SD (n = 4). Representative data are shown for each virus, which was repeated at least two times.

A/California/04/2009 (pdmH1N1) bound only to α2,6-sialosides with a strong preference for non-bisecting glycans (**C** *vs.* **C’** and **I** *vs.* **I’** units (Figure 4). A/ a/16/2009 (H3N2) bound to an *N*-glycan having the α2,6-sialosides presented on extended LacNAc moieties (**T**). In agreement with our previous report,^57^ A/bovine/Ohio/B24OSU-432/2024 (H5N1) and A/Vietnam/1203/2004 (H5N1) showed responsiveness to only to α2,3-sialosides. It displays a broad receptor specificity and tolerates both bisecting GlcNAc (*e.g.* **E**, **K**) and 1,3-linked fucosides (*e.g.* **P**, **Q**). The evolutionary earlier avian isolate only bound to bi-antennary glycan **S** having extended LacNAc moiety capped by 2,3-linked sialosides. HPAI viruses of the clade 2.3.4.4 can accommodate fucosylation and sulfation due to K222Q and S227R substitution.^50,51^ These mutations may also be responsible for tolerating bisecting GlcNAc. *N*-glycans that have a SLe^x^ moiety at one of the antennae were well recognized by the virus. In case of sialyl LacNAc moieties, two of these epitopes were required for binding. It is likely that sialyl LacNAc is a sub-optimal ligand and requires a bivalent interaction between two protomers of an HA trimer conferring high avidity of binding.^59^ In the case of SLe^x^, only one epitope at an *N*-glycan is sufficient for robust binding. The data highlight that glycan microarray data can be misinterpreted if it does not display a relevant collection of glycans.

## CONCLUSIONS

We have developed a chemoenzymatic approach for the preparation of asymmetrical bisecting bi-, tri- and tetra-antennary *N*-glycans. It exploits the finding that GnT-III can modify bi-, tri- and tetra-antennary glycans with a bisecting GlcNAc moiety. In addition, it was found that GnT-III tolerates GlcNH2 and GlcN3 at the α1,3- or α1,6-antenna of *N*-glycans making it possible to prepare asymmetrical bisecting *N*-glycans. It exploits that GlcNH2 and GlcN3 temporarily block modifications by glycosyl transferases but at an appropriate point in the synthesis can easily be converted into natural GlcNAc for selective enzymatic elaboration.

The panel of oligosaccharides made it possible to examine kinetic parameters of GlcNAc transfer catalyzed by GnT-III for various acceptors. It was found that a bi-antennary glycan is the most favorable substrate which agrees with the observation that such glycans are most commonly modified by bisection. It was also found that the GlcNAc at the α1,3-antenna is critical for transfer and agrees with the observation that hybrid type glycans can also be modified by bisection. Our studies] confirmed that the branching enzymes GnT-IV and GnT-V cannot modify bisected bi-antennary glycans highlighting bisection controls the branching of *N*-glycans. Furthermore, GnT-IV and GnT-V have much higher Km values than GnT-III and thus when cells express the latter enzyme, the expectation is that the biosynthesis of multi-antennary glycans is inhibited.

Plant lectins are commonly employed to profile glycan compositions of proteins and cells, and thus it is important to have a detailed understanding of their ligand requirements.^60^ Therefore, the collection of glycans with and without bisection was printed as a glycan microarray that was screened for binding for a panel of plant lectins, which validated proper printing. The interplay between the glycan structures on the host cell surface and the glycan-specificity of infecting viruses is an important determinant of host range, transmission, and pathogenesis. It is well established that avian viruses preferentially bind α2,3-linked sialic acids, which are found in the duck enteric and chicken upper respiratory tract.^61,62^ On the other hand, human IAVs recognize α2,6-linked sialic acids, which are predominantly found in the upper respiratory tract of humans. IAVs that circulate in humans are from avian origin, having acquired an ability to recognize α2,6-sialosides through mutations in the receptor binding pocket of HA. In the lower respiratory tract, the “avian-type receptor” is also expressed, and therefore it is possible that avian viruses can infect humans, causing severe disease but without efficient transmission. Currently, there is a worldwide outbreak of H5N1 viruses in many avian and mammalian species, including recent cases in cows, increasing the risk of zoonotic events.^63,64^ We employed the array of glycans to examine receptor spcificities of two human viruses which did not tolerate bisection. A/bovine/Ohio/B24OSU-432/2024 (H5N1) and A/Vietnam/1203/2004 (H5N1) only bound to α2,3-sialosides. The bovine virus displayed, however, a broader receptor specificity. It tolerates bisection and bound to SLe^x^ containing *N*-glycans. Only one SLe^x^ moiety at either the α1,3- or α1,6-antenna was sufficient for binding, whereas for sialyl LacNAc two of such epitopes are needed for recognition. It is likely that for the latter epitope a bivalent interaction is needed between two HA protomers of a trimer to provide a sufficient high avidity of binding.

## ASSOCIATED CONTENT

### Supporting Information

The Supporting Information is available free of charge at https://pubs.cs.org/doi/

Methods, analytical data, additional figures and schemes, and copies of NMR spectra (PDF).

## AUTHOR INFORMATION

### Authors

**Balasaheb K. Ghotekar** - Complex Carbohydrate Research Center, University of Georgia, Athens, Georgia 30602, United States.

**Seema K. Bhagwat** - Complex Carbohydrate Research Center, University of Georgia, Athens, Georgia 30602, United States.

**Pradeep Chopra** - Complex Carbohydrate Research Center, University of Georgia, Athens, Georgia 30602, United States.

**Thomas Buckley** - Complex Carbohydrate Research Center and Chemistry Department, University of Georgia, Athens, Georgia 30602, United States.

### Notes

The authors declare no competing financial interest.

## ACKNOWLEDGMENTS

This research was funded in with Federal funds from the National Institute of Allergy and Infectious Diseases, National Institutes of Health, Department of Health and Human Services, under Award Number R01 AI165692 (G.J.B.) and Contract Numbers 75N93021C00018 (GJB); NIAID Centers for Excellence for Influenza Research and Response (CEIRR). We thank Dr. Mark S. Tompkins and Mr. Sean D. Ray, Center for Vaccine and Immunology (CVI), Center for Influenza Disease and Emergence Response (CIDER) and Department of Infectious Diseases of University of Georgia for sharing the influenza viruses used for receptor specificity evaluation in this study.

## TOC Graphic

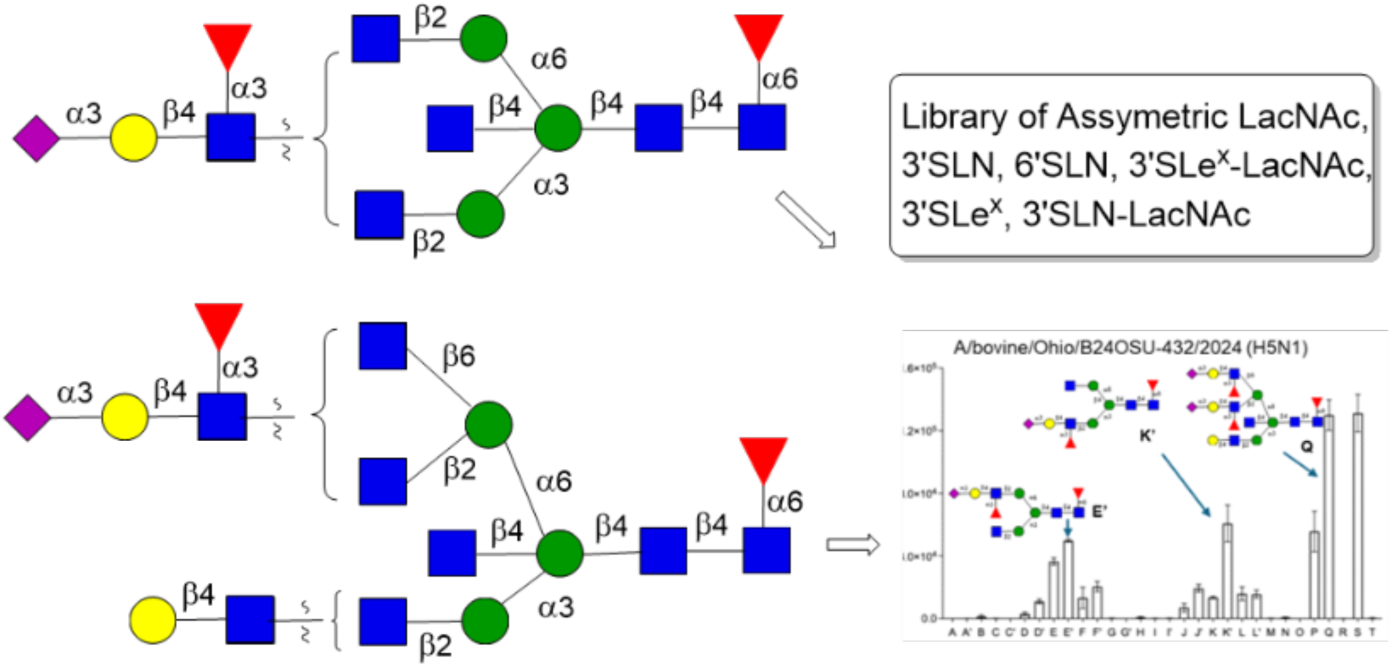

