## Supplementary material for "Chemoenzymatic Synthesis of Asymmetric Bisecting Bi-, Tri-, and Tetra-Antennary *N*-Glycans": SI

### Table of Contents

| | page |
|---|---|
| 1. Materials and Methods | S3 |
| 2. General Procedure for SGP Extraction and Trimming | S5 |
| 2a. Extraction of Sialyl Glycopeptide (SGP) | S5 |
| 2b. Enzymatic Modification of SGP to Prepare Glycosyl Asparagine-2-naphthyl methylcarbamate (**1**) | S6 |
| 3. General Protocols for Enzymatic Modification of Glycosyl Asparagine | S8 |
| 4. General Protocol for Purification | S12 |
| 5. Enzymatic Synthesis | S13 |
| **Scheme S1**. Synthesis of bi-antennary *N*-glycans **2** and **3** | S13 |
| **Scheme S2**. Synthesis of bi-antennary azido derivatives **17** and **18** | S13 |
| **Scheme S3**. Attempt towards the synthesis of **S5**-**S8** using GnT-IVB and GnT-V | S13 |
| **Scheme S4**. Synthesis of **S11** and **S12** from **3** | S14 |
| **Scheme S5**. Synthesis of **S15** and **S16** from **2** | S14 |
| 6. Condition for Liquid Chromatography (LC) Trace Analysis | S15 |
| 7. NMR Nomenclature | S16 |
| 8. Synthesis and Characterization of *N*-Glycan | S17 |
| 9. Kinetic Parameters of GlcNAc transfer GnT-III, GnT-IV and GnT-V for Multi-antennary Glycans (including **Figure S62**) | S109 |
| 10. Glycan Microarray Development | S111 |
| 11. References | S114 |
| 12. NMR Spectra | S115 |

### 1. Materials and Methods

All Chemical reagents were purchased from Sigma-Aldrich (unless noted) and used without further purification. Egg yolk powder (catalogue# 40504) was purchased from Natural Foods Inc, *Clostridium perfringens* Neuraminidase (catalogue# P0720) was obtained from New England Biolabs and Pronase from *Streptomyces griseus* (catalogue# P5147-1G) was purchased from Sigma-Aldrich. Reverse phase column chromatography was performed on Bondapak C-18 (Acetonitile:10mM ammonium bicarbonate or Water as eluent). Size exclusion chromatography was carried out on a Bio-Gel P-2 or G-50 gel column using 100 mM ammonium bicarbonate as eluant. HILIC-HPLC purification of compounds was performed on a Shimadzu 20AD UFLC LCMS-IT-TOF with a Waters XBridge BEH, Amide column, 5 µm, 10 x 250 mm. HPLC grade acetonitrile and water were purchased from Fischer. $^{1}H$ and $^{13}C$ NMR spectra were recorded on Bruker 600 or 900 MHz spectrometer. Signals are reported in terms of chemical shift [δ in parts per million (ppm)] relative to tetramethylsilane (TMS) as the internal standard. NMR data is presented as follows: Chemical shift, multiplicity (s = singlet, br. s = broad singlet, d = doublet, t = triplet, dd = doublet of doublet, m = multiplet and/or multiple resonances), coupling constant in Hertz (Hz), integration. All NMR signals were assigned on the basis of 1D: $^{1}H$ NMR, $^{13}C$ NMR and 2D: COSY, HSQC, TOCSY, HMBC, NOESY experiments. Mass spectras were recorded on Shimadzu LC/MS-IT-TOF mass spectrometer. ‡Carbon assignments were extracted from HSQC experiment.

All enzymatic reactions were performed in aqueous buffers at the appropriate pH for each enzyme. Recombinant human glycosyl transferases including α-1,3-mannosyl-glycoprotein 2-β-*N*-acetylglucosaminyltransferase (GnT-I), α-1,6-mannosyl-glycoprotein 2-β-*N*-acetylglucosaminyltransferase (GnT-II), alpha-1,3-mannosyl-glycoprotein 4-beta-*N*-acetylglucosaminyltransferase B (GnT-IVB), alpha-1,6-mannosylglycoprotein 6-beta-*N*-acetylglucosaminyltransferase (GnT-V), beta-1,4-mannosyl-glycoprotein 4-beta-*N*-acetylglucosaminyltransferase (GnT-III), β-1,4-galactosyltransferase 1 (B4GalT1), β-galactoside-α-2,6-sialyltransferase 1 (ST6Gal1), and β-galactoside-α-2, 3-sialyltransferase 4 (ST3Gal4) Fucosyltransferase 8 (FUT8), Fucosyltransferase 6 (FUT6) were provided by Dr. K. W. Moremen (Complex Carbohydrate Research Center, Athens, GA, USA) which were expressed according to published protocols.[1-4] β-*N*-acetylglucosaminidase S was purchased from New England Biolab

(Catalog # P0744S). Alkaline phosphatase from calf intestine (CIAP) and bovine serum albumin (BSA) were purchased from Sigma-Aldrich. Uridine 5′-diphospho-N-acetylglucosamine (UDP-GlcNAc) was purchased from Sigma-Aldrich. Uridine 5′-diphosphogalactose diphosphate galactose (UDP-Gal) and cytidine 5′monophospho-N-acetylneuraminc acid (CMP-Neu5Ac) were both purchased from Roche. Guanosine. 5’-diphospho-β-L-fucose (GDP-Fuc) was purchased from Carbosynth.Uridine 5′-diphospho-N-trifluoroglucosamine (UDP-GlcNTFA) was synthesized utilizing a one-pot three-enzyme combination as previously reported.[4]

### 2. General Procedure for SGP Extraction and Trimming

#### 2a. Extraction of Sialyl Glycopeptide (SGP)

SGP was obtained from commercially available egg yolk powder following and modifying our previously reported procedure.[5] In brief, an acetone suspension (2.0 L) of egg yolk powder (1.0 kg) was stirred at room temperature to remove lipids and other organic soluble components. After 2 h, the filtrate was discarded, and the insoluble powder was re-suspended in acetone and the same process was repeated again. Next, solid residue obtained was re-suspended with aq. acetone (70% v/v, 2.0 L), stirred for 2 h, filtered again, filtrate was discarded. Next, solid residue obtained was re-suspended with aq. acetone (40% v/v, 2.0 L), stirred for 2 h, and filtered. The insoluble material was discarded, and the filtrate was concentrated under reduced pressure at 40 °C. The collected liquid was concentrated in vacuo, transferred to centrifugal tubes, and kept at 0 °C. After 12 h, tubes were centrifuged (10 min, at 3500 rpm) to remove solid insoluble material. The clear liquid was transferred to fresh centrifugal tubes and diluted with dichloromethane (20 mL, DCM) and centrifuged again (10 min, at 3500 rpm). The aqueous layer was collected, concentrated under vacuum to minimum volume which was loaded on to a Bio-Gel G50 column SEC column. For elution 0.1 M ammonium bicarbonat e was employed and the desired fractions were pooled and concentrated in vacuo to yield pure SGP as a fluffy, white powder (0.9 g SGP/kg of egg yolk powder). The purity of SGP was determined by liquid chromatography-mass spectrometry (LC-MS) analysis on a Shimadzu LC/MS-IT-TOF mass spectrometer using a Xbridge amide analytical column (3.5 µm, 2.1 x 150 mm) with linear gradient of 80-50% solvent B (100% acetonitrile) in A (100% water) over a 60 min period at a flow rate of 0.25 mL/min.

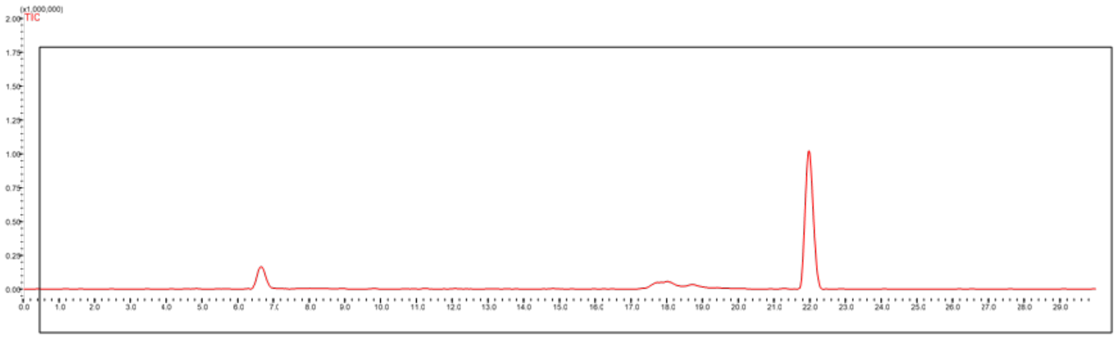

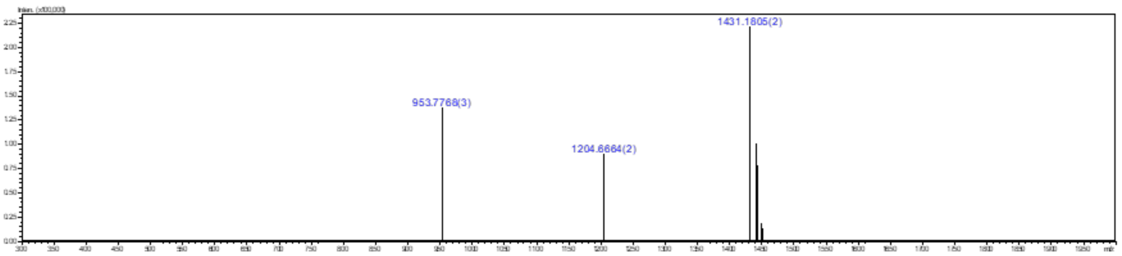

**Figure S1.** Liquid chromatography (LC) trace and mass (ESI-MS) spectrum (peak at retention time = 22.0 min) of sialylglycopeptide (SGP).

**2b. Enzymatic Modification of SGP to Prepare Glycosyl Asparagine-2-naphthyl methylcarbamate (1)**

To a solution of a sialoglycopeptide (SGP, 10 mM, 1.1 g) in sodium acetate buffer (50 mM, pH 5.5) containing $CaCl_2$ (5 mM) was added neuraminidase from *Clostridium perfringens* ((New England Biolabs # P0720L, 160 μL, 2000 units), and the mixture was incubated at 37 °C for 18 h. once ESI-MS shows completion of reaction, the pH of the reaction mixture was adjusted to 5 with acetic acid after which, β-galactosidase (800 μL, 800 units) from *Aspergillus niger* (Megazyme # E-BGLAN) were added. The reaction was incubated at 37 °C with shaking overnight, after which another 200 μL of *β*-galactosidase were added. There are no any substrates in reaction solution and full galactose removal was monitored by ESI-MS, after which an equal volume of cooled alcohol was added and kept at 0 °C to precipitate protein residuals. The supernatant liquid was concentrated in vacuum and purified by size-exclusion chromatography using G-50 BioGel eluting with a 100 mM ammonium bicarbonate solution. Product **S1** containing fractions were lyophilized to obtain the desired product in the form of a white, amorphous solid.

Peptide attached *N*-glycan **S1** (500 mg) was dissolved in 2.5 mL of Tris buffer (100 mM, pH 8.0) containing 5 mM $CaCl_2$ with final reaction concentration of 10 mM. Pronase from *Streptomyces griseus* (SigmaAldrich # P5147-1G, 200 mg) was added, and the reaction was incubated for 3 days at 50 °C. Following an 3 days incubation period at 50 °C, portion of 120 mg Pronase was added. The reaction was monitored by ESI-MS, until there is no starting material remains. Equal amount of Ethanol was added into the reaction mixture, shake, and kept at 0 °C to precipitate protein residuals. Thereafter, supernatant liquid concentrated under reduced pressure at 40 °C and purified by size-exclusion chromatography using P2 BioGel eluting with a 100 mM ammonium bicarbonate solution. Product **S2** containing fractions were combined and lyophilized to afford the desired product **S2** as a white amorphous solid. Next, compound **S2** (0.2 g, 0.139 mmol) was dissolved in 1,4-Dioxane: $H_2O$ (1:1, 14 mL) and 2-naphthylmethyl (NAP) chlorocarbonate (92 mg, 0.419 mmol), Sodium bicarbonate (0.12 g, 1.39 mmol) were added at room temparature for 16 h. After complete consumption of starting material (Monitored by TLC and LC-MS), concentrated and purified by size-exclusion chromatography using P2 BioGel eluting with a 100 mM ammonium bicarbonate solution to afford **S3** as white amorphous solid. The fractions containing the trimmed Compound **S3** were pooled, lyophilized, and dissolved in 10 mL of Tris buffer (100 mM, pH 7.5). To this mixture BSA (1 mg), calf intestine alkaline phosphatase (CIAP, 100 µL, 2kU/mL), $MgCl_2$ (10mM), GDP-Fucose (0.2 g), and FUT8 (200 µL, 1 mg/mL) were added and the reaction was incubated overnight at 37 °C with shaking. The reaction was lyophilized and purified by size-exclusion chromatography using P-2 BioGel eluting with a 0.1 M ammonium bicarbonate solution. The fractions containing **1** were pooled, lyophilized, and subjected to C18 followed by P-4 (extrafine, <45 µm) Biogel size-exclusion column chromatography purification according to the general protocol (0.125 g, 52% over 2 steps).

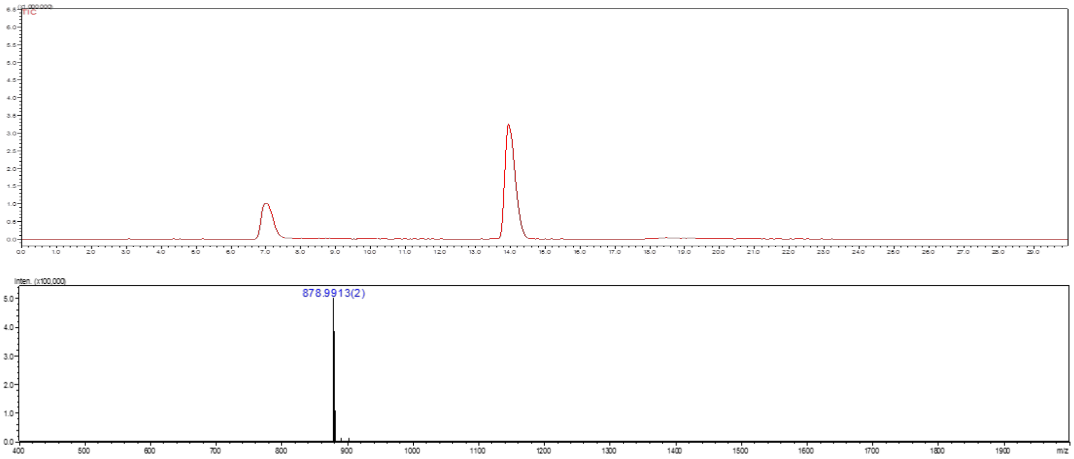

**Figure S2.** Liquid chromatography (LC) trace and mass (ESI-MS) spectrum (peak at retention time = 13.5 min) of **1**.

### 3. General Protocols for Enzymatic Modification of Glycosyl Asparagine

#### 3a. General Procedure for the Installation of β1,2-GlcNAc or β1,2-GlcNTFA using GnT-I

Asparagine acceptor (1.0 equiv) and UDP-GlcNAc or UDP-GlcNTFA (1.5 equiv) were dissolved in a MES buffer (100 mM, pH 6.5) containing $MnCl_2$ (10 mM) keeping a final acceptor concentration of 10 mM. Recombinant GnT-I (50 µg/µmol acceptor) were successively added, and the reaction mixture was incubated overnight at 37 °C with shaking. Reaction progress was monitored by LC-ESI-MS and another portion of GnT-I was added if starting material remained after 18 h. The product was purified by C18 followed by P-4 (extrafine, <45 µm) Biogel size-exclusion column chromatography according to the general protocol.

#### 3b. General Procedure for the Installation of β1,2-GlcNAc or β1,2-GlcNTFA using GnT-II

Acceptor (1.0 equiv) and UDP-GlcNAc or UDP-GlcNTFA (1.5 equiv) were dissolved in a MES buffer (100 mM, pH 6.5) containing $MnCl_2$ (10 mM) to provide a final acceptor concentration of 10 mM. GnT-II (50 µg/µmol acceptor) were successively added, and the mixture solution was incubated overnight at 37 °C with shaking. Reaction progress was monitored by ESI-MS and if starting material remained after 18 h another portion of GnT-II was added until no starting material could be detected. The product was purified by C18 followed by P-4 (extrafine, <45 µm) Biogel size-exclusion column chromatography according to the general protocol.

#### 3c. General Procedure for the Installation of β1,2-GlcNAc using GnT-III

Asparagine acceptor (1.0 equiv) and UDP-GlcNAc (1.5 equiv) were dissolved in a MES buffer (100 mM, pH 6.5) containing $MnCl_2$ (2 mM), and BSA (1% total volume, stock solution = 10 mg/mL) to provide a final acceptor concentration of 10 mM. GnT-III (50 µg/µmol acceptor) were successively added, and the mixture solution was incubated overnight at 37 °C with shaking. Reaction progress was monitored by ESI-MS and if starting material remained after 18 h another portion of GnT-III was added until no starting material could be detected. The product was purified by C18 followed by P-4 (extrafine, <45 µm) Biogel size-exclusion column chromatography according to the general protocol.

#### 3d. General Procedure for the Installation of β1,2-GlcNAc using GnT-IVB

Acceptor (1.0 equiv) and UDP-GlcNAc (1.5 equiv) were dissolved in a Tris buffer (100 mM, pH 7.5) containing $MnCl_2$ (5 mM), and BSA (1% total volume, stock solution = 10 mg/mL) to provide a final acceptor concentration of 10 mM. CIAP (1% total volume, stock solution = 1 kU/mL) and GnT-IVB (50 µg/µmol acceptor) were successively added, and the mixture solution was incubated overnight at 37 °C with shaking. Reaction progress was monitored by ESI-MS and if starting material remained after 18 h another portion of GnT-IVB was added until no starting material could be detected. The product was purified by C18 followed by P-4 (extrafine, <45 µm) Biogel size-exclusion column chromatography according to the general protocol.

#### 3e. General Procedure for the Installation of β1,2-GlcNAc or β1,2-GlcNTFA using GnT-V

Asparagine acceptor (1.0 equiv) and UDP-GlcNAc or UDP-GlcNTFA (1.5 equiv) were dissolved in a sodium cacodylate buffer (100 mM, pH 6.5) containing MnCl2 (2 mM), and BSA (1% total volume, stock solution = 10 mg/mL) to provide a final acceptor concentration of 10 mM. CIAP (1% total volume, stock solution = 1 kU/mL) and GnT-V (50 µg/µmol acceptor) were successively added, and the mixture solution was incubated overnight at 37 °C with shaking. Reaction progress was monitored by ESI-MS and if starting material remained after 18 h another portion of GnT-V was added until no starting material could be detected. The product was purified by C18 followed by P-4 (extrafine, <45 µm) Biogel size-exclusion column chromatography according to the general protocol.

#### 3f. General Procedure for the Installation of β1,4 Gal using B4GalT1

Glycosyl asparagine acceptor (1 eq) and UDP-Gal (1.5 eq per Gal to be added) were dissolved to prepare an reaction concentration of 5 mM in a Tris buffer (100 mM, pH 7.5) containing $MnCl_2$ (10 mM) and BSA (1% total volume, stock solution = 10 mg/mL). CIAP (1% total volume, stock solution = 1 kU/mL) and B4GalT1 (0.5% wt/wt relative to acceptor substrate) were added, and the reaction mixture was incubated overnight at 37 °C with gentle shaking. Reaction progress was monitored by LC-ESI-MS, and if starting material remained after 18 h another portion of B4GalT1 was added until no starting material could be detected. The product was purified by C18 followed by P-4 (extrafine, <45 µm) Biogel size-exclusion column chromatography according to the general protocol.

#### 3g. General Procedure for the Installation of α2,3 Neu5Ac using ST3Gal4 or ST6Gal1

Asparagine acceptor (1 eq) and CMP-Neu5Ac (1.5 eq per Neu5Ac to be added) were dissolved at a final acceptor concentration of 5 mM in a sodium cacodylate buffer (50 mM, pH 7.2) containing BSA (1% total volume, stock solution = 10 mg/mL). CIAP (1% total volume, stock solution = 1 kU/mL) and ST3Gal4 or ST6Gal1 (1% wt/wt relative to acceptor substrate) were added, and the reaction mixture was incubated overnight at 37 °C with gentle shaking. Reaction progress was monitored by ESI-TOF MS, and if starting material remained after 18 h another portion of ST3Gal4 or ST6Gal1 was added until no starting material could be detected. The product was purified by C18 followed by P-4 (extrafine, <45 μm) Biogel size-exclusion column chromatography according to the general protocol.

#### 3h. General Procedure for the Installation of α1,3 Fuc using FUT6

Acceptor (1 eq) and GDP-Fuc (1.5 eq per Fuc to be added) were dissolved at a final acceptor concentration of 5 mM in a Tris buffer (50 mM, pH 7.3) containing $MnCl_2$ (10 mM). CIAP (1% total volume, stock solution = 1 kU/mL) and FUT6 (1% wt/wt) were added, and the reaction mixture was incubated overnight at 37 °C with gentle shaking. Reaction progress was monitored by ESI-TOF MS, and if starting material remained after 18 h another portion of FUT6 was added until no starting material could be detected. The product was purified by C18 followed by P-4 (extrafine, <45 μm) Biogel size-exclusion column chromatography according to the general protocol.

#### 3i. General Procedure for Removal of TFA Group of *N*-Glycans

*N*-Glycan was dissolved in deionized water with the final reaction concentration of 5 mM. The pH of the reaction solution was adjusted to 10 using NaOH (1 M), and the mixture solution was incubated for 3-5 h at 37 °C with shaking. When complete, the reaction was neutralized by aqueous acetic acid (1 M). The product was first purified by C18 biogel [Eluent – 10mM ammonium bicarbonate and water:acetonitrile (8:2)] followed by P-4 (extrafine, <45 μm) Biogel size-exclusion column chromatography.

#### 3j. General Procedure for Amine Acetylation of *N*-Glycans

The amine containing *N*-glycans were dissolved (1 equiv) in 50 mM of water. Solids of $NaHCO_3$ (10 equiv, pH 8.0) and $Ac_2O$ (1.2 equiv) were added to convert the $NH_2$ moiety into NHAc moiety and the reaction mixture was gently mixed until all solids were dissolved and incubated for 1–2 h at 37 °C. The reaction progress was monitored by LC-ESI-MS and additional $NaHCO_3$ (depending upon pH) and $Ac_2O$ was added if the starting material was remaining. The product was first purified by C18 biogel [Eluent - water and 10mM ammonium bicarbonate:acetonitrile (8:2)] followed by P-4 (extrafine, <45 μm) Biogel size-exclusion column chromatography.

#### 3k. General Procedure for Removal of 2-Naphthyl methylcarbamate Group

*N*-Glycans were dissolved in a 50% tert-butanol/water solution (1 mg/mL substrate concentration). To this solution Pd/C (1 mg/2 mg of starting material) was added. The reaction was stirred vigorously under an atmosphere of hydrogen (1 atm) and monitored by LC-ESI-MS. Once the reaction had gone to completion, it was filtered through a Whatman syringe filter (0.2 μm) to remove the catalyst, and the filtrate was lyophilized to provide the final product.

#### 3l. General Procedure for the Conversion of $GlcNH_2$ into $GlcN_3$

Intermediate $GlcNH_2$ glycans (5 μmol, 1 eq) was dissolved in water (2 mL) and to this solution was added imidazole-1-sulfonyl azide hydrogen sulfate (50 μmol), $K_2CO_3$ (50 μmol) and catalytic $CuSO_4.5H_2O$. The reaction mixture was incubated overnight at 37 °C with gentle shaking. Reaction progress was monitored by MALDI-TOF MS. The reaction mixture was lyophilized and the salts were removed by C18 biogel [Eluent - water and water:acetonitrile (8:2)] followed by P-4 (extrafine, <45 μm) Biogel size-exclusion column chromatography.

### 4. General Protocol for Purification

The reaction mixture was centrifuged over a Nanosep® Omega ultrafiltration device (10 kDa MWCO) to remove proteins. The decanted solution was first loaded on C18 Biogel and washed with 8-10 mL $H_2O$ to remove excess sugar nucleotides and further product was eluted using 10 mM ammonium bicarbonate (or $H_2O$ for base sensitive TFA containing compound) : acetonitrile (8:2). Fractions containing product were collected and lyophilized. The lyophilized product was dissolved and loaded on P-4 (extrafine, <45 µm) Biogel size-exclusion column chromatography using 100 mM ammonium bicarbonate as eluent.

### 5. Enzymatic Synthesis

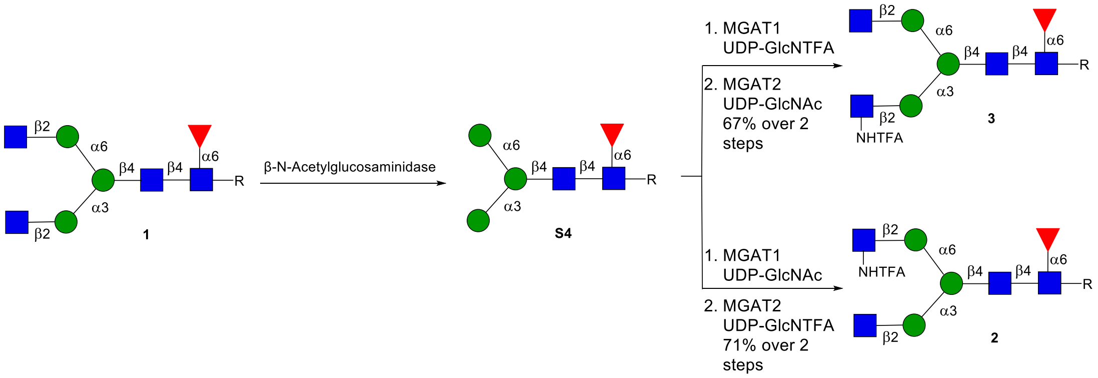

**Scheme S1**. Synthesis of bi-antennary *N*-glycans **2** and **3**.

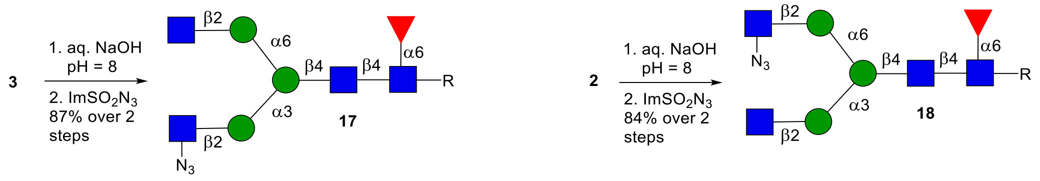

**Scheme S2**. Synthesis of bi-antennary azido derivatives **17** and **18**.

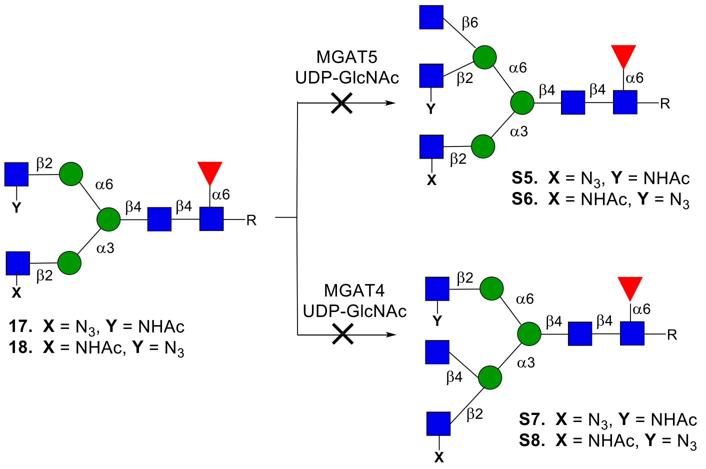

**Scheme S3**. Attempt towards the synthesis of **S5-S8** using GnT-IVB and GnT-V.

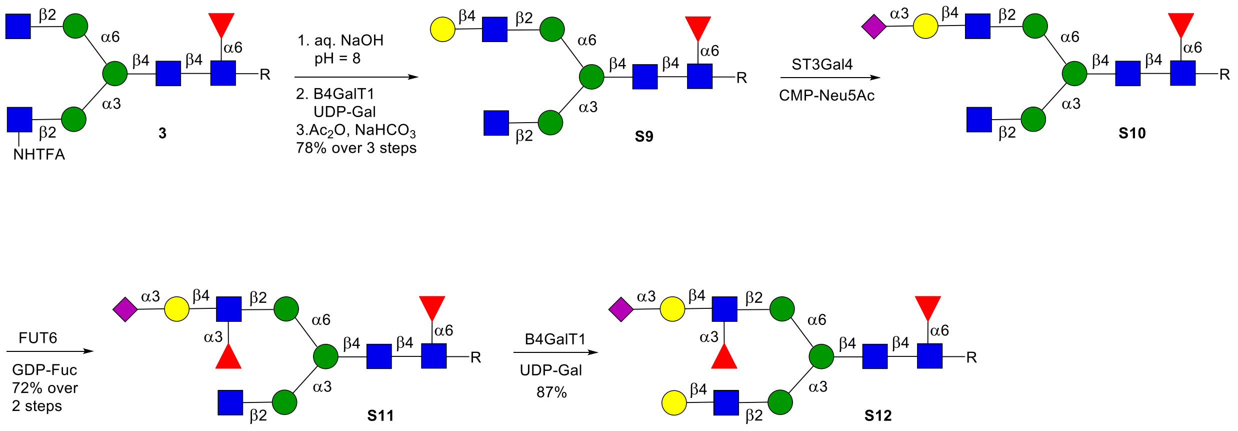

**Scheme S4**. Synthesis of asymmetric bi-antennary glycosyl asparagine **S11** and **S12** from **3**.

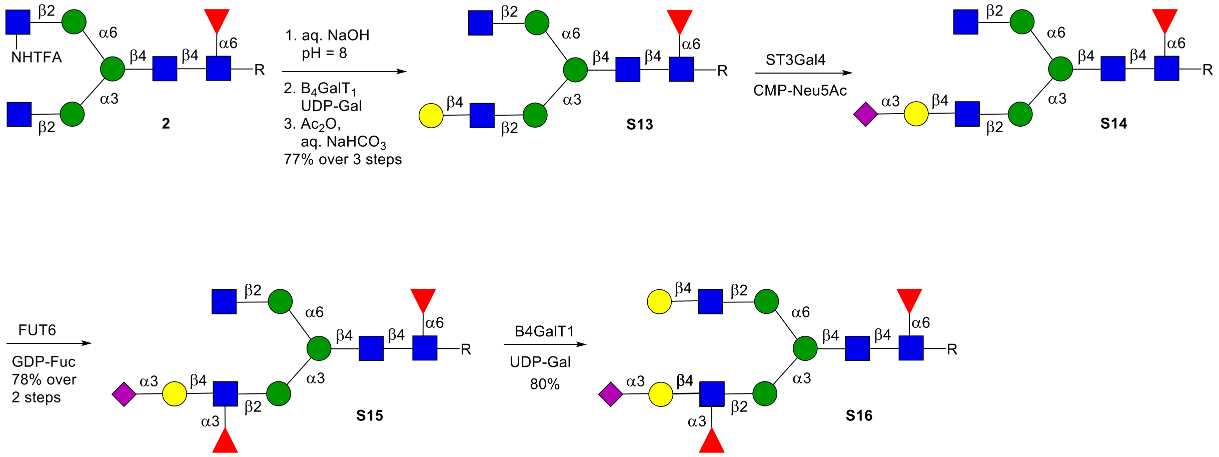

**Scheme S5**. Synthesis of asymmetric bi-antennary glycosyl asparagine **S15** and **S16** from **2**.

### 6. Condition for Liquid Chromatography (LC) Trace Analysis

Liquid Chromatography (LC) was performed on a Shimadzu LC-ESI-IT-TOF with a Waters XBridge BEH, Amide column, 3.5 μm, 2.1 x 150 mm (Semi-preparative HILIC) at a flow rate of 0.16 mL/min, injection volume of 100-500 μL, with 1% of the flow is diverted to the ESI-MS detector using a splitter. Mobile phase A was 10 mM ammonium formate in water, adjusted to pH 3.5 with formic acid; mobile phase B was acetonitrile. The general condition using a linear gradient is as follows:

| Time (Min) | B % | A% |
|---|---|---|
| 0.01 | 80 | 20 |
| 18 | 50 | 50 |
| 20 | 25 | 75 |
| 24 | 25 | 75 |
| 26 | 80 | 20 |
| 30 | 80 | 20 |

The common peak in LC trace at retention time = 6 to 7 min is Salt peak from sodium and ammonium formate.

### 7. NMR Nomenclature

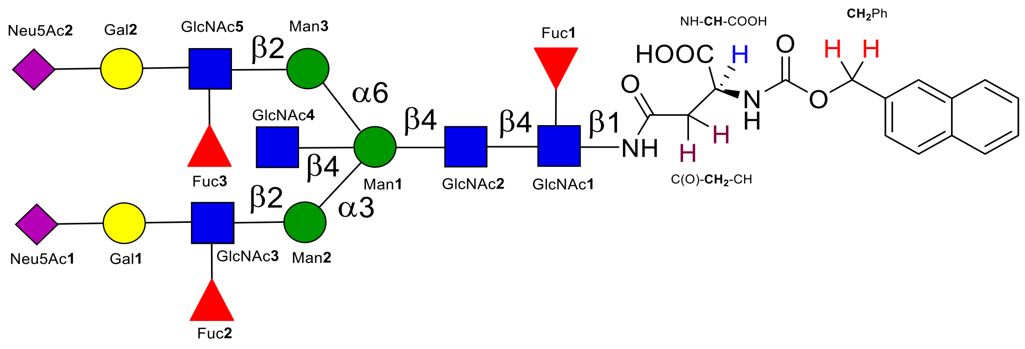

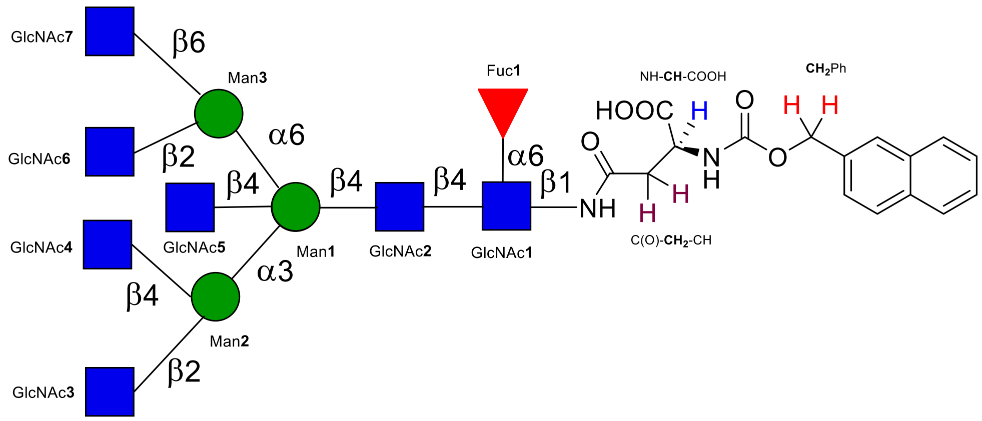

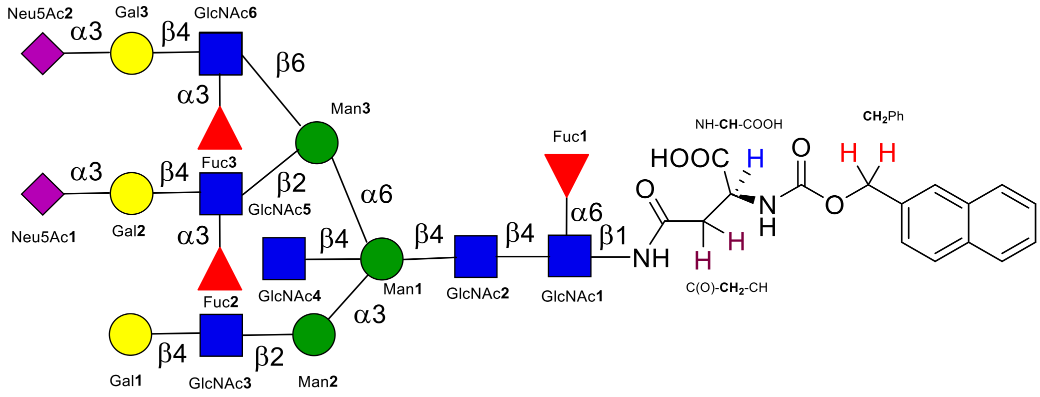

### 8. Synthesis and Characterization of *N*-Glycan

#### Compound 1

**1** was synthesized from SGP (1.1 g, 401.2 μmol) following the general protocol **2b**.

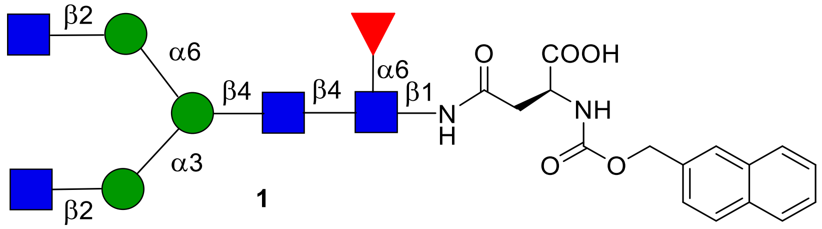

**1**

**$^{1}$H NMR (600 MHz, $D_2O$): δ (ppm)**

| | H1 | H2 | H3 | H4 | H5 | H6 | NHAc |
|---|---|---|---|---|---|---|---|
| GlcNAc**1** | 4.85 | 3.69 | 3.44 | 3.38 | N/R[a] | N/R | 2.02-1.79 (12H) |
| GlcNAc**2** | 4.55 | 3.70 | 3.50 | 3.65 | N/R | N/R | 2.02-1.79 (12H) |
| GlcNAc**3** | 4.47 | 3.63 | 3.47 | 3.38 | N/R | N/R | 2.02-1.79 (12H) |
| GlcNAc**4** | 4.47 | 3.63 | 3.47 | 3.36 | N/R | N/R | 2.02-1.79 (12H) |
| Man**1** | 4.70 | 4.17 | 3.69 | 3.54 | N/R | N/R | - |
| Man**2** | 5.03 | 4.10 | 3.82 | 3.42 | 3.54 | N/R | - |
| Man**3** | 4.84 | 4.03 | 3.81 | 3.42 | 3.54 | N/R | - |
| Fuc**1** | 4.72 | 3.65 | 3.72 | 3.60 | 3.94 | 1.05 | - |

| Signal | Proton | Carbon |
|---|---|---|
| **Aromatic** | 7.92-7.84 (m, 4H)<br>7.53-7.47 (m, 3H) | 128.2-125.8 |
| **$CH_2$**Ph | 5.32 (d, $J$ = 12.7 Hz)<br>5.14 (d, $J$ = 12.6 Hz) | 66.7 |
| NH-**CH**-COOH | 4.29 (dd, $J$ = 9.5, 3.6 Hz) | 53.0 |
| C(O)-**$CH_2$**-CH | 2.75 (dd, $J$ = 14.8, 2.9 Hz)<br>2.51 (dd, $J$ = 15.0, 9.9 Hz) | 38.6 |

**$^{13}C$ NMR (150 MHz, $D_2O$): δ (ppm)**

| | **C1** |
|---|---|
| GlcNAc**1** | 78.2 |
| GlcNAc**2** | 100.8 |
| GlcNAc**3** | 99.6 |
| GlcNAc**4** | 99.6 |
| Man**1** | 100.5 |
| Man**2** | 99.5 |
| Man**3** | 97.0 |
| Fuc**1** | 99.2 |

[a] Not reported

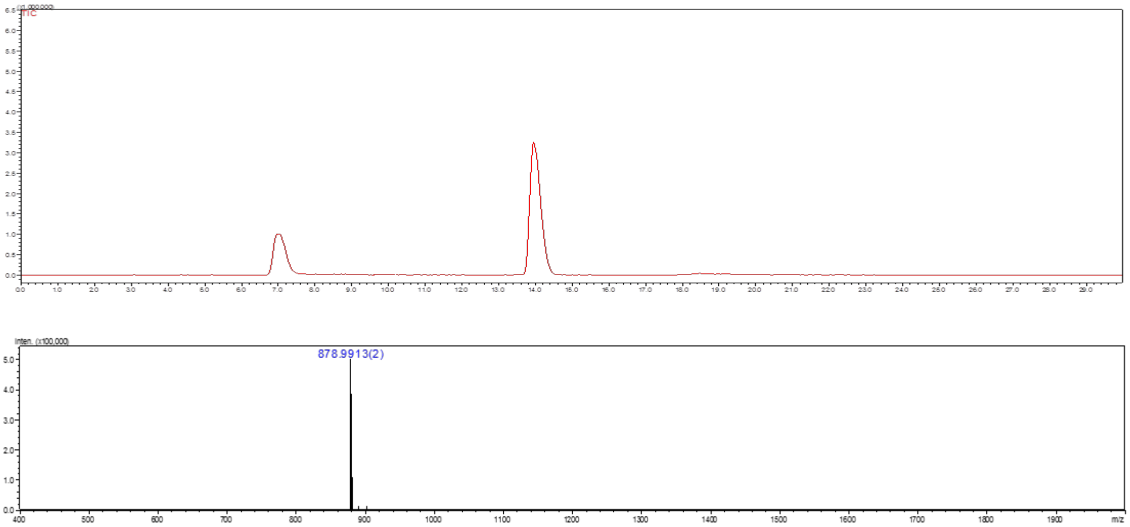

**Figure S6.** Liquid chromatography (LC) trace and mass (ESI-MS) spectrum (peak at retention time = 14.0 min) of **1**.

**Compound 10**

**10** was synthesized from **1** (2 mg, 1.13 μmol) following the general protocol **3d** for the installation of GlcNAc moiety using UDP-GlcNAc. The product **10** was obtained as a white fluffy solid (1.8 mg, 81%).

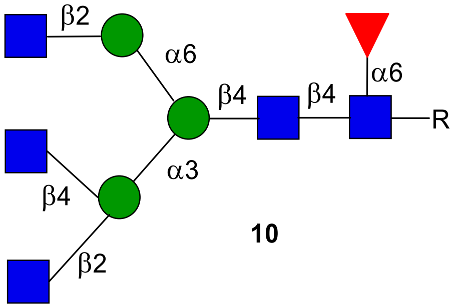

**10**

**$^1$H NMR (600 MHz, $D_2O$): δ (ppm)**

| | H1 | H2 | H3 | H4 | H5 | H6 | NHAc |
|---|---|---|---|---|---|---|---|
| GlcNAc**1** | 4.86 | 3.70 | 3.44 | 3.39 | N/R[a] | N/R | 2.03-1.80 (15H) |
| GlcNAc**2** | 4.55 | 3.71 | 3.50 | 3.65 | N/R | N/R | 2.03-1.80 (15H) |
| GlcNAc**3** | 4.45 | 3.68 | 3.44 | 3.41 | N/R | N/R | 2.03-1.80 (15H) |
| GlcNAc**4** | 4.48 | 3.63 | 3.47 | 3.37 | N/R | N/R | 2.03-1.80 (15H) |
| GlcNAc**5** | 4.48 | 3.63 | 3.47 | 3.37 | N/R | N/R | 2.03-1.80 (15H) |
| Man**1** | 4.68 | 4.14 | 3.69 | 3.55 | N/R | N/R | - |
| Man**2** | 5.04 | 4.14 | 3.97 | 3.55 | 3.52 | N/R | - |
| Man**3** | 4.84 | 4.04 | 3.83 | 3.54 | 3.43 | N/R | - |
| Fuc**1** | 4.72 | 3.67 | 3.73 | 3.61 | 3.95 | 1.08 | - |

| Signal | Proton | Carbon |
|---|---|---|
| **Aromatic** | 7.93-7.85 (m, 4H)<br>7.54-7.48 (m, 3H) | 128.4-125.7 |
| **$CH_2$**Ph | 5.34 (d, $J$ = 12.8 Hz)<br>5.14 (d, $J$ = 12.7 Hz) | 66.9 |
| NH-**CH**-COOH | 4.30 (dd, $J$ = 9.5, 3.6 Hz) | 53.1 |
| C(O)-**$CH_2$**-CH | 2.75 (dd, $J$ = 14.8, 2.9 Hz) | 38.6 |

| | |
|---|---|
| | 2.51 (dd, $J$ = 15.0, 9.9 Hz) |

**$^{13}C$ NMR (150 MHz, $D_2O$): δ (ppm)**

| | **C1** |
|---|---|
| GlcNAc**1** | 78.2 |
| GlcNAc**2** | 100.8 |
| GlcNAc**3** | 99.6 |
| GlcNAc**4** | 101.7 |
| GlcNAc**5** | 99.6 |
| Man**1** | 100.3 |
| Man**2** | 99.0 |
| Man**3** | 97.0 |
| Fuc**1** | 99.2 |

[a] Not reported

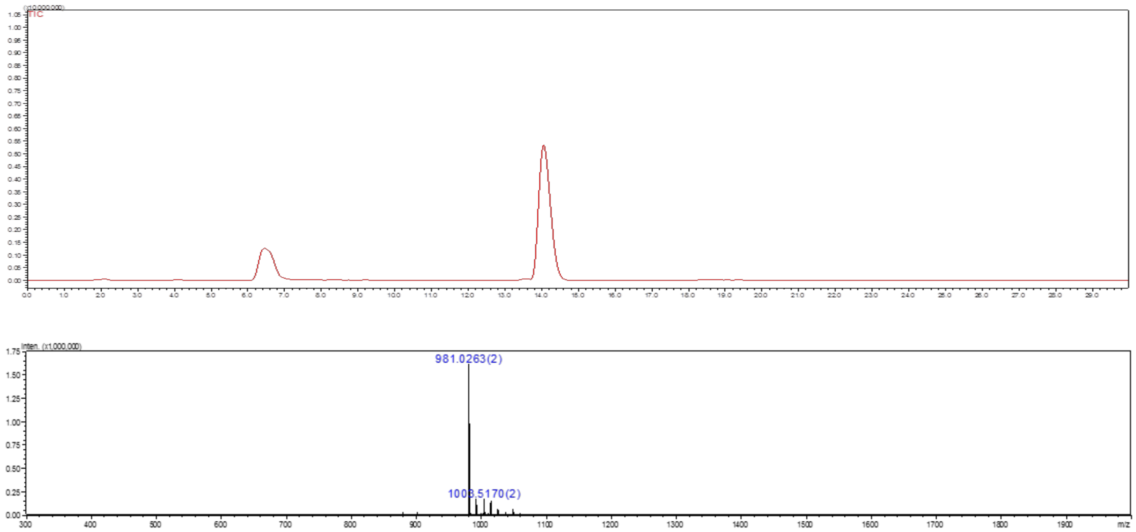

**Figure S7.** Liquid chromatography (LC) trace and mass (ESI-MS) spectrum (peak at retention time = 14.2 min) of **10**.

**Compound 11**

**11** was synthesized from **1** (10 mg, 5.67 µmol) following the general protocol **3e** for the installation of GlcNAc moiety using UDP-GlcNAc. The product **11** was obtained as a white fluffy solid (10.1 mg, 91%).

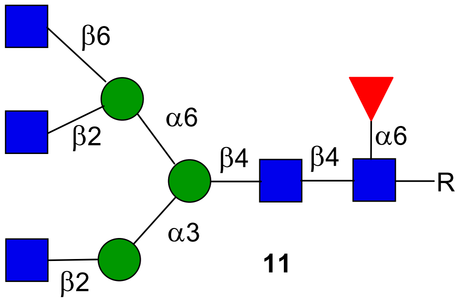

**11**

**$^1$H NMR (900 MHz, $D_2O$): δ (ppm)**

| | **H1** | **H2** | **H3** | **H4** | **H5** | **H6** | **NHAc** |
|---|---|---|---|---|---|---|---|
| GlcNAc**1** | 4.86 | 3.70 | 3.45 | 3.38 | N/R[a] | N/R | 2.02-1.79 (15H) |
| GlcNAc**2** | 4.54 | 3.69 | 3.50 | 3.65 | N/R | N/R | 2.02-1.79 (15H) |
| GlcNAc**3** | 4.48 | 3.59/3.64[b] | 3.45/3.47[b] | 3.36/3.38[b] | N/R | N/R | 2.02-1.79 (15H) |
| GlcNAc**4** | 4.48 | 3.59/3.64[b] | 3.45/3.47[b] | 3.36/3.38[b] | N/R | N/R | 2.02-1.79 (15H) |
| GlcNAc**5** | 4.46 | 3.64 | 3.49 | 3.39 | N/R | N/R | 2.02-1.79 (15H) |
| Man**1** | 4.69 | 4.17 | 3.69 | 3.54 | N/R | N/R | - |
| Man**2** | 5.05 | 4.12 | 3.83 | 3.67 | N/R | N/R | - |
| Man**3** | 4.79 | 4.02 | 3.80 | 3.66 | 3.34 | N/R | - |
| Fuc**1** | 4.73 | 3.66 | 3.71 | 3.61 | 3.93 | 1.06 | - |

| **Signal** | **Proton** | **Carbon** |
|---|---|---|
| **Aromatic** | 7.91-7.85 (m, 4H)<br>7.53-7.47 (m, 3H) | 128.3-125.7 |
| **$CH_2$**Ph | 5.33 (d, $J$ = 12.8 Hz)<br>5.14 (d, $J$ = 12.7 Hz) | 66.8 |
| NH-**CH**-COOH | 4.29 (dd, $J$ = 9.5, 3.6 Hz) | 53.1 |
| C(O)-**$CH_2$**-CH | 2.75 (dd, $J$ = 14.8, 2.9 Hz) | 38.6 |

| | |
|---|---|
| | 2.51 (dd, $J$ = 15.0, 9.9 Hz) |

**$^{13}$C NMR (225 MHz, $D_2O$): δ (ppm)**

| | **C1** |
|---|---|
| GlcNAc**1** | 78.2 |
| GlcNAc**2** | 100.8 |
| GlcNAc**3** | 99.7 |
| GlcNAc**4** | 99.6 |
| GlcNAc**5** | 101.6 |
| Man**1** | 100.4 |
| Man**2** | 99.5 |
| Man**3** | 97.1 |
| Fuc**1** | 99.2 |

[a] Not reported
[b] Could not be determined

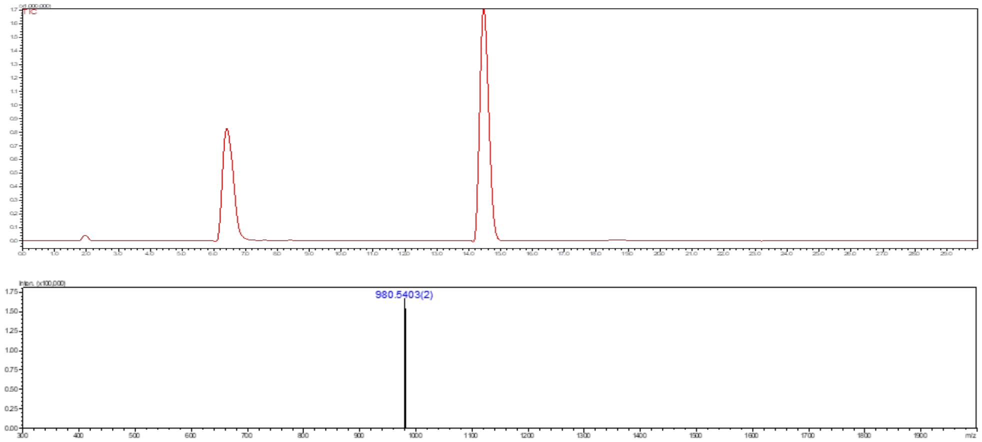

**Figure S8.** Liquid chromatography (LC) trace and mass (ESI-MS) spectrum (peak at retention time = 14.5 min) of **11**.

**Compound 12**

**12** was synthesized from **11** (7 mg, 3.56 µmol) following the general protocol **3d** for the installation of GlcNAc moiety using UDP-GlcNAc. The product **12** was obtained as a white fluffy solid (7.7 mg, 93%).

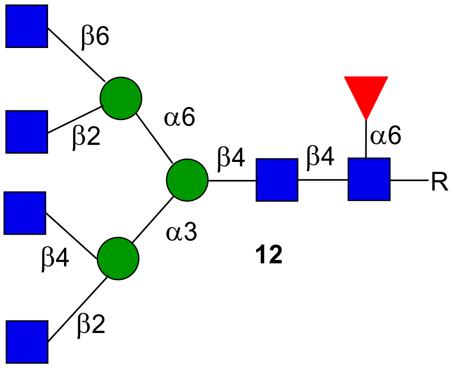

**12**

**$^1$H NMR (600 MHz, $D_2O$): δ (ppm)**

| | **H1** | **H2** | **H3** | **H4** | **H5** | **H6** | **NHAc** |
|---|---|---|---|---|---|---|---|
| GlcNAc**1** | 4.84 | 3.69 | 3.50 | 3.36 | N/R[a] | N/R | 2.01-1.78 (18H) |
| GlcNAc**2** | 4.53 | 3.69 | 3.49 | 3.66 | N/R | N/R | 2.01-1.78 (18H) |
| GlcNAc**3** | 4.48/4.47[b] | 3.59/3.62[b] | 3.48/3.49[b] | 3.36/3.38[b] | N/R | N/R | 2.01-1.78 (18H) |
| GlcNAc**4** | 4.46 | 3.66/3.64[b] | 3.50/3.49[b] | 3.42 | N/R | N/R | 2.01-1.78 (18H) |
| GlcNAc**5** | 4.48/4.47[b] | 3.59/3.62[b] | 3.48/3.49[b] | 3.36/3.38[b] | N/R | N/R | 2.01-1.78 (18H) |
| GlcNAc**6** | 4.44 | 3.66/3.64[b] | 3.50/3.49[b] | 3.38 | N/R | N/R | 2.01-1.78 (18H) |
| Man**1** | 4.69 | 4.15 | 3.70 | 3.41 | N/R | N/R | - |
| Man**2** | 5.02 | 4.13 | 3.96 | 3.53 | 3.72 | 3.50 | - |
| Man**3** | 4.77 | 4.00 | 3.78 | 3.66 | N/R | N/R | - |
| Fuc**1** | 4.71 | 4.65 | 3.70 | 3.60 | 3.94 | 1.05 | - |

| **Signal** | **Proton** | **Carbon** |
|---|---|---|
| **Aromatic** | 7.90-7.83 (m, 4H)<br>7.52-7.46 (m, 3H) | 128.4-125.7 |
| **$CH_2$**Ph | 5.31 (d, $J$ = 12.8 Hz)<br>5.12 (d, $J$ = 12.7 Hz) | 66.8 |

| | | |
|---|---|---|
| NH-**CH**-COOH | 4.28 (dd, *J* = 9.5, 3.6 Hz) | 53.1 |
| C(O)-**$CH_2$**-CH | 2.74 (dd, *J* = 14.8, 2.9 Hz)<br>2.50 (dd, *J* = 15.0, 9.9 Hz) | 38.7 |

**$^{13}C$ NMR (150 MHz, $D_2O$): δ (ppm)**

| | **C1** |
|---|---|
| GlcNAc**1** | 78.2 |
| GlcNAc**2** | 100.9 |
| GlcNAc**3** | 99.7 |
| GlcNAc**4** | 101.6 |
| GlcNAc**5** | 99.7 |
| GlcNAc**6** | 101.6 |
| Man**1** | 100.3 |
| Man**2** | 99.1 |
| Man**3** | 97.2 |
| Fuc**1** | 99.2 |

[a] Not reported

[b] Could not be determined

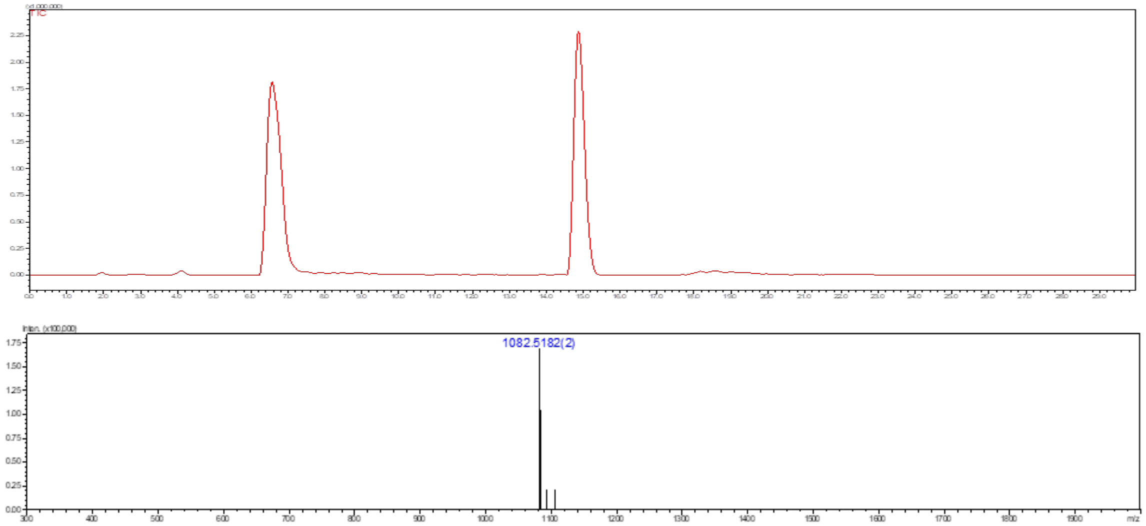

**Figure S11.** Liquid chromatography (LC) trace and mass (ESI-MS) spectrum (peak at retention time = 14.9 min) of **12**.

**Compound 13**

**13** was synthesized from **10** (3 mg, 1.52 µmol) following the general protocol **3c** for the installation of GlcNAc moiety using UDP-GlcNAc. The product **13** was obtained as a white fluffy solid (3.1 mg, 93%).

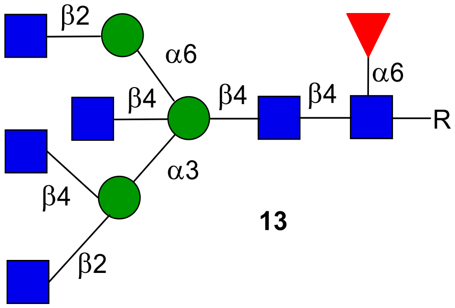

**13**

**$^1$H NMR (900 MHz, $D_2O$): δ (ppm)**

| | H1 | H2 | H3 | H4 | H5 | H6 | NHAc |
|---|---|---|---|---|---|---|---|
| GlcNAc**1** | 4.83 | 3.68 | 3.43 | 3.36 | N/R[a] | N/R | 2.01-1.78 (18H) |
| GlcNAc**2** | 4.52 | 3.70 | 3.46 | 3.62 | N/R | N/R | 2.01-1.78 (18H) |
| GlcNAc**3** | 4.45 | 3.62 | 3.49 | 3.38 | N/R | N/R | 2.01-1.78 (18H) |
| GlcNAc**4** | 4.43 | 3.67 | 3.46 | 3.39 | N/R | N/R | 2.01-1.78 (18H) |
| GlcNAc**5** | 4.37 | 3.60 | 3.31 | 3.18 | 3.48 | N/R | 2.01-1.78 (18H) |
| GlcNAc**6** | 4.45 | 3.62 | 3.49 | 3.38 | N/R | N/R | 2.01-1.78 (18H) |
| Man**1** | 4.59 | 4.06 | 3.76 | N/R | N/R | N/R | - |
| Man**2** | 4.96 | 4.19 | 3.96 | 3.49 | N/R | N/R | - |
| Man**3** | 4.90 | 4.06 | 3.75 | 3.46 | N/R | N/R | - |
| Fuc**1** | 4.71 | 3.65 | 3.70 | 3.59 | 3.92 | 1.04 | - |

| Signal | Proton | Carbon |
|---|---|---|
| **Aromatic** | 7.90-7.84 (m, 4H)<br>7.51-7.46 (m, 3H) | 128.4-125.7 |
| **$CH_2$**Ph | 5.31 (d, $J$ = 12.8 Hz)<br>5.12 (d, $J$ = 12.7 Hz) | 66.8 |
| NH-**CH**-COOH | 4.28 (dd, $J$ = 9.5, 3.6 Hz) | 53.1 |

| | | |
|---|---|---|
| C(O)-**CH$_2$**-CH | 2.74 (dd, *J* = 14.8, 2.9 Hz)<br>2.50 (dd, *J* = 15.0, 9.9 Hz) | 38.6 |

**$^{13}$C NMR (225 MHz, D$_2$O): δ (ppm)**

| | **C1** |
|---|---|
| GlcNAc**1** | 78.2 |
| GlcNAc**2** | 100.9 |
| GlcNAc**3** | 99.8 |
| GlcNAc**4** | 101.6 |
| GlcNAc**5** | 100.5 |
| GlcNAc**6** | 99.8 |
| Man**1** | 100.1 |
| Man**2** | 99.4 |
| Man**3** | 97.6 |
| Fuc**1** | 99.1 |

[a] Not reported

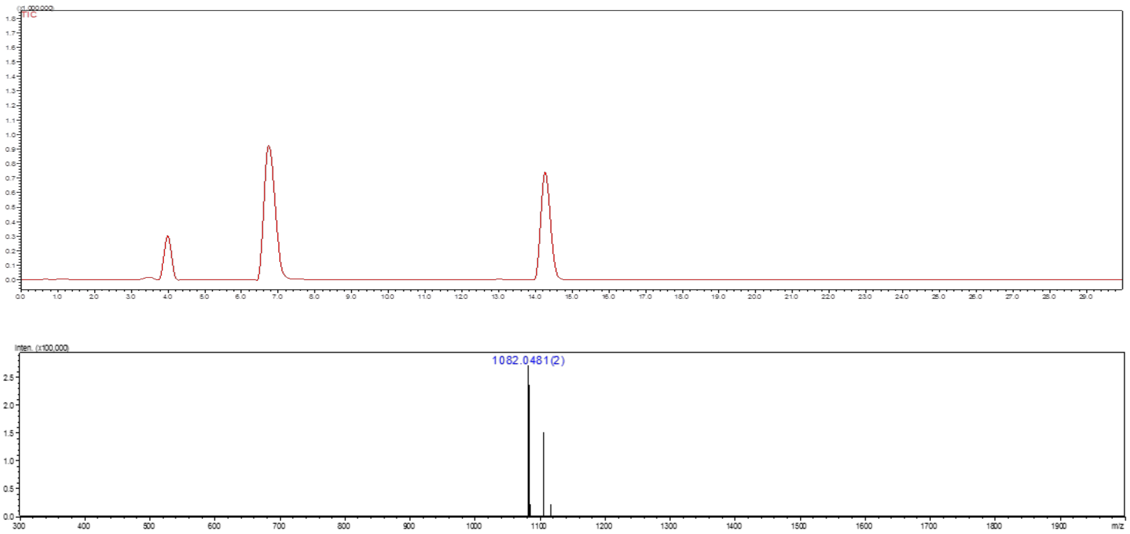

**Figure S9.** Liquid chromatography (LC) trace and mass (ESI-MS) spectrum (peak at retention time = 14.3 min) of **13**.

**Compound 5**

**14** was synthesized from **11** (3 mg, 1.52 μmol) following the general protocol **3c** for the installation of GlcNAc moiety using UDP-GlcNAc. The product **14** was obtained as a white fluffy solid (2.7 mg, 82%).

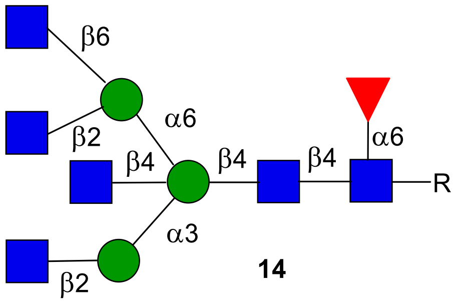

**14**

**$^1$H NMR (900 MHz, $D_2O$): δ (ppm)**

| | **H1** | **H2** | **H3** | **H4** | **H5** | **H6** | **NHAc** |
|---|---|---|---|---|---|---|---|
| GlcNAc**1** | 4.91 | 3.75 | 3.50 | 3.44 | N/R[a] | N/R | 2.08-1.85 (18H) |
| GlcNAc**2** | 4.59 | 3.76 | 3.54 | 3.71 | N/R | N/R | 2.08-1.85 (18H) |
| GlcNAc**3** | 4.56 | 3.71 | 3.48 | N/R | N/R | N/R | 2.08-1.85 (18H) |
| GlcNAc**4** | 4.47 | 3.69 | 3.37 | 3.26 | 3.59 | N/R | 2.08-1.85 (18H) |
| GlcNAc**5** | 4.54 | 3.65 | 3.44 | 3.36 | N/R | N/R | 2.08-1.85 (18H) |
| GlcNAc**6** | 4.50 | 3.72 | 3.56 | 3.44 | N/R | N/R | 2.08-1.85 (18H) |
| Man**1** | 4.68 | 4.15 | 3.84 | 3.51 | N/R | N/R | - |
| Man**2** | 5.04 | 4.22 | 3.88 | 3.46 | 3.57 | N/R | - |
| Man**3** | 4.89 | 4.11 | 3.79 | 3.41 | 3.53 | N/R | - |
| Fuc**1** | 4.79 | 3.71 | 3.76 | 3.66 | 4.00 | 1.10 | - |

| **Signal** | **Proton** | **Carbon** |
|---|---|---|
| **Aromatic** | 7.91-7.85 (m, 4H)<br>7.53-7.47 (m, 3H) | 128.2-125.5 |
| **$CH_2$**Ph | 5.33 (d, *J* = 12.8 Hz)<br>5.14 (d, *J* = 12.7 Hz) | 66.9 |
| NH-**CH**-COOH | 4.29 (dd, *J* = 9.5, 3.6 Hz) | 53.1 |

| | | |
|---|---|---|
| C(O)-**$CH_2$**-CH | 2.75 (dd, *J* = 14.8, 2.9 Hz)<br>2.51 (dd, *J* = 15.0, 9.9 Hz) | 38.9 |

**$^{13}$C NMR (225 MHz, $D_2O$): δ (ppm)**

| | **C1** |
|---|---|
| GlcNAc**1** | 78.2 |
| GlcNAc**2** | 100.9 |
| GlcNAc**3** | 99.8 |
| GlcNAc**4** | 99.9 |
| GlcNAc**5** | 99.8 |
| GlcNAc**6** | 101.7 |
| Man**1** | 99.9 |
| Man**2** | 99.8 |
| Man**3** | 97.9 |
| Fuc**1** | 99.2 |

[a] Not reported

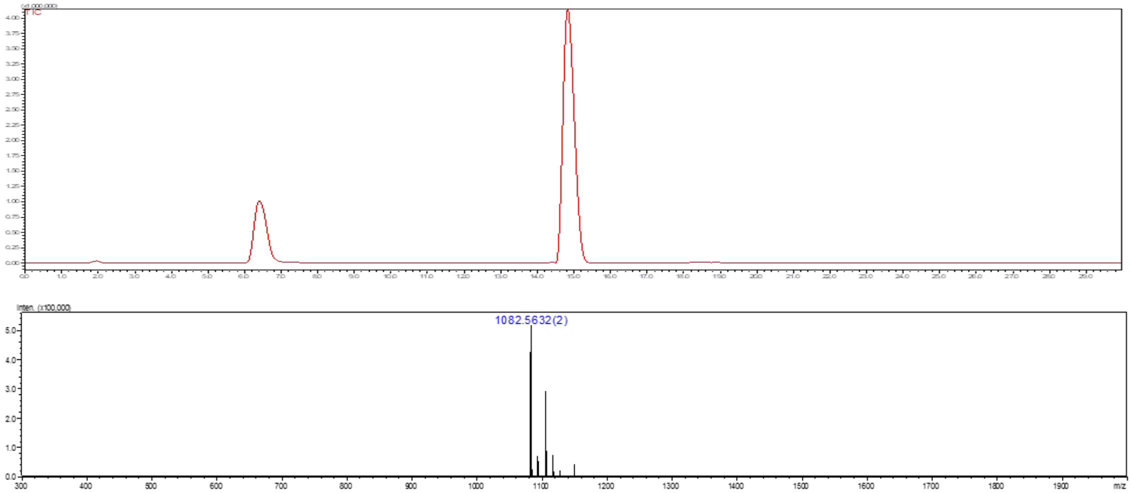

**Figure S10.** Liquid chromatography (LC) trace and mass (ESI-MS) spectrum (peak at retention time = 14.9 min) of **14**.

**Compound 15**

**15** was synthesized from **12** (4 mg, 1.84 µmol) following the general protocol **3c** for the installation of GlcNAc moiety using UDP-GlcNAc. The product **15** was obtained as a white fluffy solid (3.9 mg, 91%).

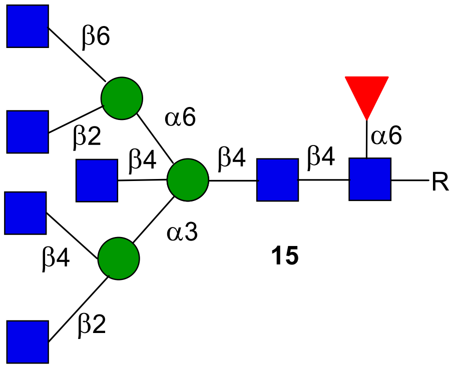

**15**

**$^1$H NMR (600 MHz, $D_2O$): δ (ppm)**

| | H1 | H2 | H3 | H4 | H5 | H6 | NHAc |
|---|---|---|---|---|---|---|---|
| GlcNAc**1** | 4.68 | 3.70 | 3.50 | 3.38 | N/R[a] | N/R | 2.02-1.79 (21H) |
| GlcNAc**2** | 3.53 | 3.72 | 3.48 | 3.65 | N/R | N/R | 2.02-1.79 (21H) |
| GlcNAc**3** | 4.50 | 3.64 | 3.52 | 3.41 | N/R | N/R | 2.02-1.79 (21H) |
| GlcNAc**4** | 4.44 | 3.66 | 3.50 | 3.42 | N/R | N/R | 2.02-1.79 (21H) |
| GlcNAc**5** | 4.41 | 3.61 | 3.31 | 3.21 | 3.53 | N/R | 2.02-1.79 (21H) |
| GlcNAc**6** | 4.48 | 3.59 | 3.38 | 3.31 | N/R | N/R | 2.02-1.79 (21H) |
| GlcNAc**7** | 4.43 | 3.68 | 3.51 | 3.39 | N/R | N/R | 2.02-1.79 (21H) |
| Man**1** | 4.61 | 4.06 | 3.79 | 4.08 | N/R | N/R | - |
| Man**2** | 4.98 | 4.20 | 3.97 | 3.51 | N/R | N/R | - |
| Man**3** | 4.82 | 4.04 | 3.73 | 3.48 | N/R | N/R | - |
| Fuc**1** | 4.73 | 3.67 | 3.72 | 3.62 | 3.93 | 1.06 | - |

| Signal | Proton | Carbon |
|---|---|---|
| **Aromatic** | 7.91-7.85 (m, 4H)<br>7.53-7.47 (m, 3H) | 128.4-125.7 |
| **CH$_2$**Ph | 5.32 (d, $J$ = 12.8 Hz) | 66.8 |

| | | |
|---|---|---|
| | 5.14 (d, $J$ = 12.7 Hz) | |
| NH-**CH**-COOH | 4.29 (dd, $J$ = 9.5, 3.6 Hz) | 53.1 |
| C(O)-**$CH_2$**-CH | 2.75 (dd, $J$ = 14.8, 2.9 Hz)<br>2.51 (dd, $J$ = 15.0, 9.9 Hz) | 38.6 |

**$^{13}$C NMR (150 MHz, $D_2O$): δ (ppm)**

| | **C1** |
|---|---|
| GlcNAc**1** | 78.2 |
| GlcNAc**2** | 100.9 |
| GlcNAc**3** | 99.9 |
| GlcNAc**4** | 101.7 |
| GlcNAc**5** | 99.8 |
| GlcNAc**6** | 99.9 |
| GlcNAc**7** | 101.7 |
| Man**1** | 99.8 |
| Man**2** | 99.5 |
| Man**3** | 98.0 |
| Fuc**1** | 99.2 |

[a] Not reported

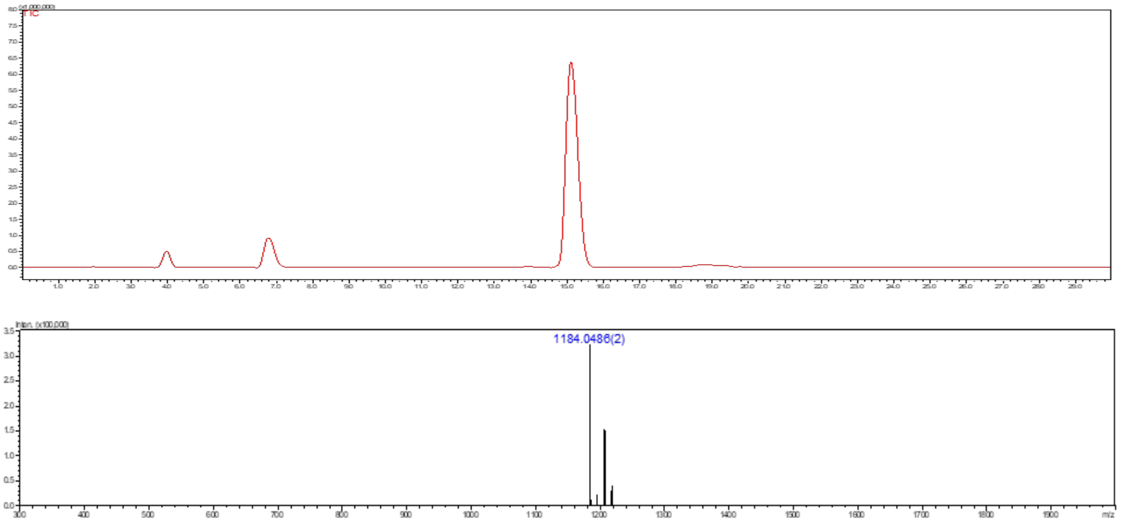

**Figure S12.** Liquid chromatography (LC) trace and mass (ESI-MS) spectrum (peak at retention time = 15.2 min) of **15**.

**Compound 16**

Compound **16** was synthesized from **1** (6 mg, 3.40 μmol) following the general protocol **3c** for the installation of GlcNAc moiety using UDP-GlcNAc. The product **16** was obtained as a white fluffy solid (5.6 mg, 88%).

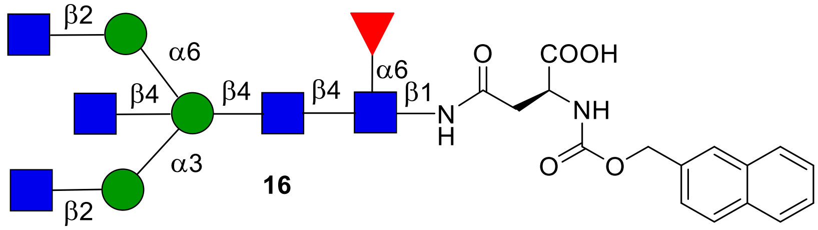

**16**

**$^{1}$H NMR (600 MHz, $D_2O$): δ (ppm)**

| | **H1** | **H2** | **H3** | **H4** | **H5** | **H6** | **NHAc** |
|---|---|---|---|---|---|---|---|
| GlcNAc**1** | 4.85 | 3.69 | 3.44 | 3.38 | N/R[a] | N/R | 2.02-1.79 (12H) |
| GlcNAc**2** | 4.54 | 3.72 | 3.48 | 3.66 | N/R | N/R | 2.02-1.79 (12H) |
| GlcNAc**3** | 4.48 | 3.65 | 3.50 | 3.41 | N/R | N/R | 2.02-1.79 (12H) |
| GlcNAc**4** | 4.39 | 3.62 | 3.33 | 3.19 | 3.49 | 3.62 | 2.02-1.79 (12H) |
| GlcNAc**5** | 4.47 | 3.63 | 3.50 | 3.41 | N/R | N/R | 2.02-1.79 (12H) |
| Man**1** | 4.61 | 4.10 | 3.79 | 4.00 | 3.46 | N/R | - |
| Man**2** | 4.98 | 4.17 | 3.82 | 3.40 | 3.53 | 3.61 | - |
| Man**3** | 4.92 | 4.07 | 3.75 | 3.41 | 3.55 | 3.56 | - |
| Fuc**1** | 4.72 | 3.65 | 3.71 | 3.61 | 3.93 | 1.05 | - |

| **Signal** | **Proton** | **Carbon** |
|---|---|---|
| **Aromatic** | 7.95-7.82 (m, 4H)<br>7.54-7.47 (m, 3H) | 128.4-125.7 |
| **$CH_2$**Ph | 5.33 (d, $J$ = 12.7 Hz)<br>5.14 (d, $J$ = 12.6 Hz) | 66.9 |
| NH-**CH**-COOH | 4.29 (dd, $J$ = 9.5, 3.6 Hz) | 53.1 |
| C(O)-**$CH_2$**-CH | 2.73 (dd, $J$ = 14.8, 2.9 Hz)<br>2.51 (dd, $J$ = 15.0, 9.9 Hz) | 38.7 |

**$^{13}$C NMR (150 MHz, $D_2O$): δ (ppm)**

| | **C1** |
|---|---|
| GlcNAc**1** | 78.2 |
| GlcNAc**2** | 100.8 |
| GlcNAc**3** | 99.9 |
| GlcNAc**4** | 100.6 |
| GlcNAc**5** | 99.5 |
| Man**1** | 100.1 |
| Man**2** | 99.9 |
| Man**3** | 97.7 |
| Fuc**1** | 99.2 |

[a] Not reported

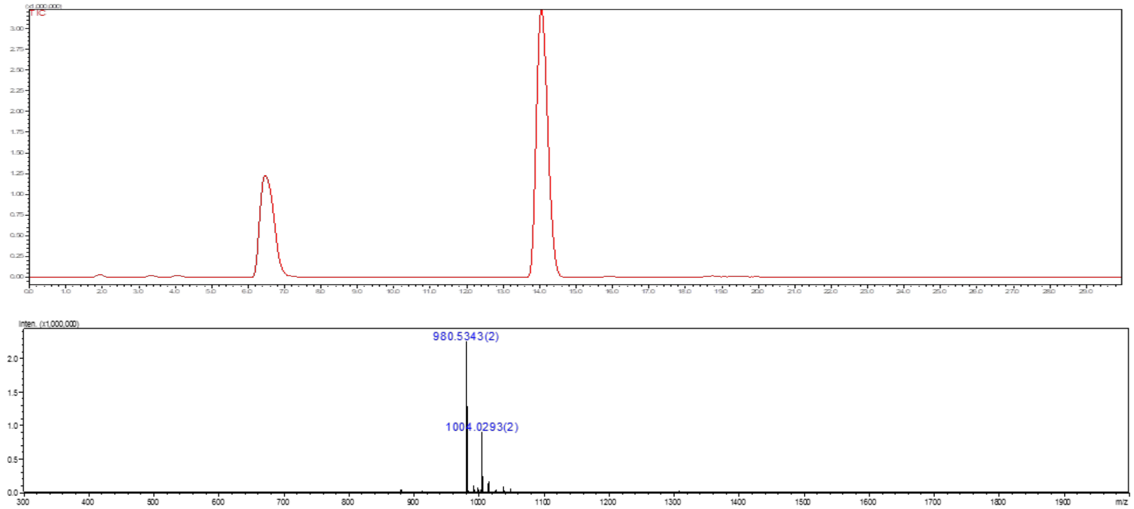

**Figure S13.** Liquid chromatography (LC) trace and mass (ESI-MS) spectrum (peak at retention time = 14.1 min) of **16**.

### Compound R

The amine of *N*-glycan compound **16** was subjected for the hydrogenation using a general protocol **3k** to obtain **R** as a white solid.

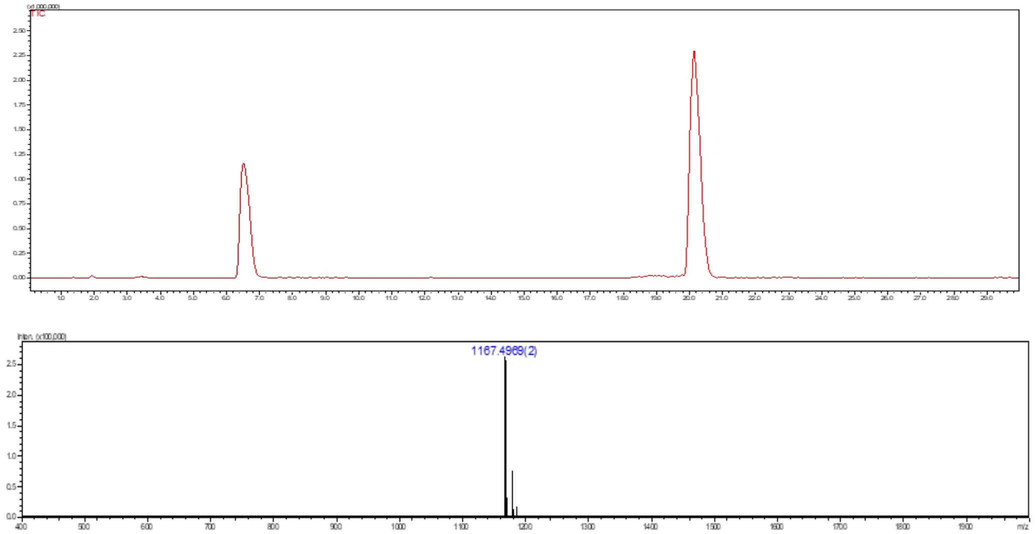

**Figure S29.** Liquid chromatography (LC) trace and mass (ESI-MS) spectrum (peak at retention time = 20.2 min) of **R**.

### Compound 6

Compound **6** was synthesized from **3** (18 mg, 1.65 μmol) following the general protocol **3c** for the installation of GlcNAc moiety using UDP-GlcNAc. The product **6** was obtained as a white fluffy solid (19.9 mg, 79%).

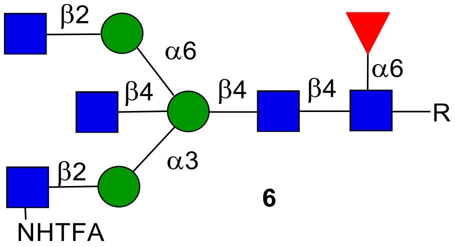

**6**

**$^1$H NMR (600 MHz, $D_2O$): δ (ppm)**

| | H1 | H2 | H3 | H4 | H5 | H6 | NHAc |
|---|---|---|---|---|---|---|---|
| GlcNAc**1** | 4.85 | 3.70 | 3.44 | 3.39 | 3.61 | N/R[a] | 2.02-1.79 (12H) |

| | | | | | | | |
|---|---|---|---|---|---|---|---|
| GlcNAc**2** | 4.54 | 3.71 | 3.49 | 3.64 | N/R | N/R | 2.02-1.79 (12H) |
| GlcNTFA | 4.46 | 3.63 | 3.43 | N/R | N/R | N/R | - |
| GlcNAc**3** | 4.39 | 3.62 | 3.34 | 3.19 | 3.49 | N/R | 2.02-1.79 (12H) |
| GlcNAc**4** | 4.61 | 3.74 | 3.41 | 3.32 | N/R | N/R | 2.02-1.79 (12H) |
| Man**1** | 4.61 | 4.09 | 3.76 | 3.98 | 3.47 | 3.82 | - |
| Man**2** | 4.93 | 1.19 | 3.83 | 3.36 | 3.47 | N/R | - |
| Man**3** | 4.92 | 4.07 | 3.76 | 3.42 | 3.55 | N/R | - |
| Fuc**1** | 4.73 | 3.66 | 3.72 | 3.62 | 3.95 | 1.07 | - |

| **Signal** | **Proton** | **Carbon** |
|---|---|---|
| **Aromatic** | 7.91-7.84 (m, 4H)<br>7.53-7.48 (m, 3H) | 128.3-125.7 |
| **$CH_2$**Ph | 5.33 (d, *J* = 12.8 Hz)<br>5.14 (d, *J* = 12.7 Hz) | 66.9 |
| NH-**CH**-COOH | 4.29 (dd, *J* = 9.5, 3.6 Hz) | 53.1 |
| C(O)-**$CH_2$**-CH | 2.75 (dd, *J* = 14.8, 2.9 Hz)<br>2.51 (dd, *J* = 15.0, 9.9 Hz) | 38.6 |

**$^{13}$C NMR (150 MHz, $D_2O$): δ (ppm)**

| | **C1** |
|---|---|
| GlcNAc**1** | 78.2 |
| GlcNAc**2** | 100.8 |
| GlcNTFA | 98.9 |
| GlcNAc**3** | 100.5 |
| GlcNAc**4** | 99.4 |
| Man**1** | 100.1 |
| Man**2** | 99.8 |
| Man**3** | 97.6 |
| Fuc**1** | 99.1 |

[a] Not reported

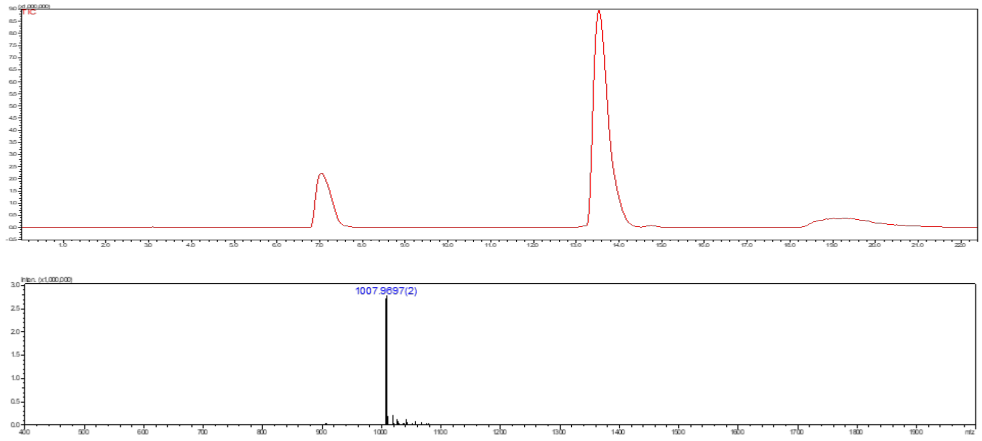

**Figure S14.** Liquid chromatography (LC) trace and mass (ESI-MS) spectrum (peak at retention time = 13.7 min) of **6**.

**Compound 4**

**4** was synthesized from **2** (3 mg, 1.65 µmol) following the general protocol **3c** for the installation of GlcNAc moiety using UDP-GlcNAc. The product **4** was obtained as a white fluffy solid (2.3 mg, 76%).

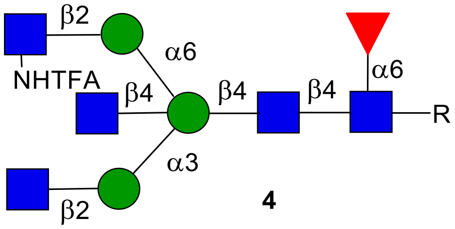

**4**

**$^1$H NMR (600 MHz, $D_2O$): δ (ppm)**

| | H1 | H2 | H3 | H4 | H5 | H6 | NHAc |
|---|---|---|---|---|---|---|---|
| GlcNAc**1** | 4.85 | 3.69 | 3.44 | 3.39 | 3.61 | N/R[a] | 2.01-1.79 (12H) |
| GlcNAc**2** | 4.54 | 3.71 | 3.49 | 3.66 | N/R | N/R | 2.01-1.79 (12H) |
| GlcNAc**3** | 4.47 | 3.63 | 3.52 | 3.38 | N/R | N/R | 2.01-1.79 (12H) |
| GlcNAc**4** | 4.37 | 3.62 | 3.35 | 3.18 | 3.49 | N/R | 2.01-1.79 (12H) |
| GlcNTFA | 4.60 | 3.73 | 3.54 | 3.44 | 3.38 | N/R | - |

| | | | | | | | |
|---|---|---|---|---|---|---|---|
| Man**1** | 4.60 | 4.09 | 3.78 | 3.97 | 3.46 | 3.79 | - |
| Man**2** | 4.98 | 4.17 | 3.82 | 3.41 | 3.62 | N/R | - |
| Man**3** | 4.89 | 4.11 | 3.77 | 3.35 | 3.53 | N/R | - |
| Fuc**1** | 4.72 | 3.66 | 3.72 | 3.60 | 3.94 | 1.05 | - |

| Signal | Proton | Carbon |
|---|---|---|
| **Aromatic** | 7.91-7.84 (m, 4H)<br>7.53-7.47 (m, 3H) | 128.4-125.7 |
| $\mathbf{CH_2}$Ph | 5.33 (d, $J$ = 12.8 Hz)<br>5.14 (d, $J$ = 12.7 Hz) | 66.9 |
| NH-**CH**-COOH | 4.29 (dd, $J$ = 9.5, 3.6 Hz) | 53.1 |
| C(O)-$\mathbf{CH_2}$-CH | 2.75 (dd, $J$ = 14.8, 2.9 Hz)<br>2.51 (dd, $J$ = 15.0, 9.9 Hz) | 38.6 |

**$^{13}$C NMR (150 MHz, $D_2O$): δ (ppm)**

| | C1 |
|---|---|
| GlcNAc**1** | 78.2 |
| GlcNAc**2** | 100.9 |
| GlcNAc**3** | 99.9 |
| GlcNAc**4** | 100.6 |
| GlcNTFA | 98.7 |
| Man**1** | 100.2 |
| Man**2** | 99.9 |
| Man**3** | 97.6 |
| Fuc**1** | 99.2 |

[a] Not reported

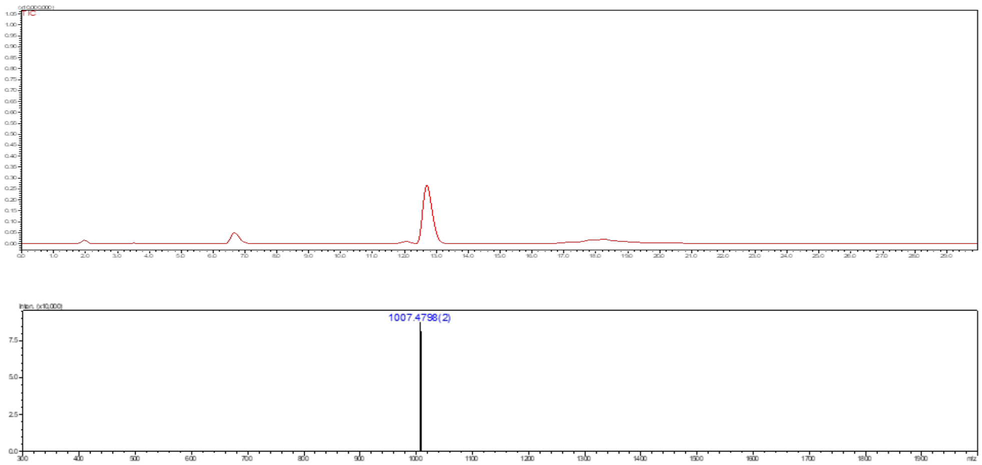

**Figure S15.** Liquid chromatography (LC) trace and mass (ESI-MS) spectrum (peak at retention time = 12.8 min) of **4**.

**Compound 17**

**17** was prepared starting from **3** (2.9 mg, 1.65 μmol) by first converting GlcNHTFA to $GlcNH_2$ following general procedure **3i** to get amine intermediate. To the resulting intermediate was subjected for the conversion of $GlcNH_2$ to $GlcN_3$ using the general procedure **3l** to obtain **17** as a white solid (2.4 mg, 87% over 2 steps).

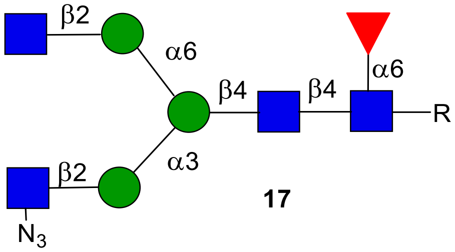

**$^1$H NMR (600 MHz, $D_2O$): δ (ppm)**

| | **H1** | **H2** | **H3** | **H4** | **H5** | **H6** | **NHAc** |
|---|---|---|---|---|---|---|---|
| GlcNAc**1** | 4.86 | 3.70 | 3.44 | 3.38 | 3.61 | N/R[a] | 2.02-1.79 (9H) |
| GlcNAc**2** | 4.55 | 3.71 | 3.51 | N/R | N/R | N/R | 2.02-1.79 (9H) |
| $GlcN_3$ | 4.55 | 3.31 | 3.35 | N/R | N/R | N/R | - |
| GlcNAc**3** | 4.46 | 3.63 | 3.48 | 3.36 | N/R | N/R | 2.02-1.79 (9H) |

| | | | | | | | |
|---|---|---|---|---|---|---|---|
| Man**1** | 4.71 | 4.17 | 3.70 | 3.57 | N/R | N/R | - |
| Man**2** | 5.05 | 4.11 | 3.82 | 3.36 | 3.54 | N/R | - |
| Man**3** | 5.04 | 4.14 | 3.89 | 3.57 | N/R | N/R | - |
| Fuc**1** | 4.72 | 3.66 | 3.71 | 3.60 | 3.93 | 1.04 | - |

| **Signal** | **Proton** | **Carbon** |
|---|---|---|
| **Aromatic** | 7.91-7.84 (m, 4H)<br>7.53-7.48 (m, 3H) | 128.3-125.7 |
| **$CH_2$**Ph | 5.33 (d, *J* = 12.8 Hz)<br>5.14 (d, *J* = 12.7 Hz) | 66.8 |
| NH-**CH**-COOH | 4.29 (dd, *J* = 9.5, 3.6 Hz) | 53.1 |
| C(O)-**$CH_2$**-CH | 2.75 (dd, *J* = 14.8, 2.9 Hz)<br>2.51 (dd, *J* = 15.0, 9.9 Hz) | 38.6 |

**$^{13}$C NMR (150 MHz, $D_2O$): δ (ppm)**

| | **C1** |
|---|---|
| GlcNAc**1** | 78.0 |
| GlcNAc**2** | 100.8 |
| $GlcN_3$ | 99.6 |
| GlcNAc**3** | 99.7 |
| Man**1** | 100.5 |
| Man**2** | 99.2 |
| Man**3** | 96.6 |
| Fuc**1** | 99.1 |

[a] Not reported

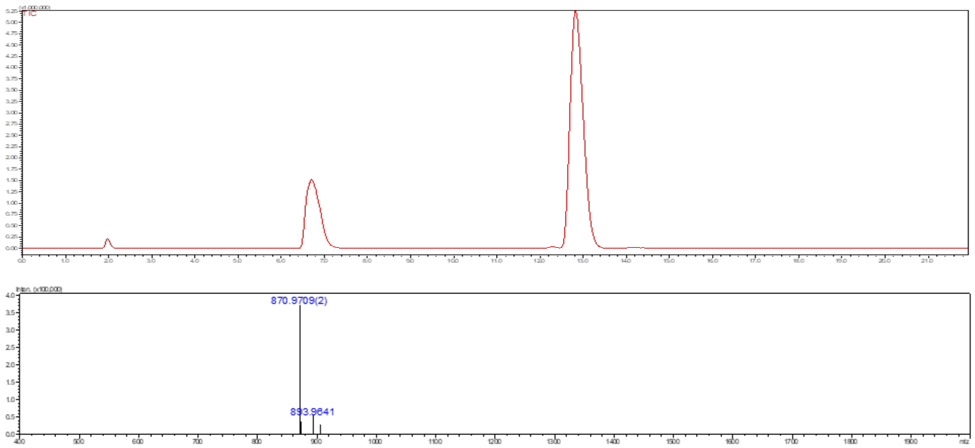

**Figure S16.** Liquid chromatography (LC) trace and mass (ESI-MS) spectrum (peak at retention time = 12.9 min) of **17**.

**Compound 18**

**18** was prepared starting from **2** (2.6 mg, 1.49 μmol) by first converting GlcNHTFA to $GlcNH_2$ following general procedure **3i** to get amine intermediate. To the resulting intermediate was subjected for the conversion of $GlcNH_2$ to $GlcN_3$ using the general procedure **3l** to obtain **18** as a white solid (2.1 mg, 84% over 2 steps).

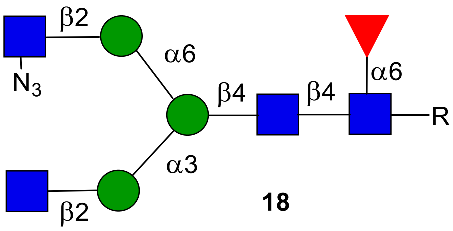

**18**

**$^1$H NMR (600 MHz, $D_2O$): δ (ppm)**

| | H1 | H2 | H3 | H4 | H5 | H6 | NHAc |
|---|---|---|---|---|---|---|---|
| GlcNAc**1** | 4.87 | 3.75 | 3.45 | 3.38 | 3.61 | N/R[a] | 2.01-1.79 (9H) |
| GlcNAc**2** | 4.56 | 3.71 | 3.50 | N/R | N/R | N/R | 2.01-1.79 (9H) |
| GlcNAc**3** | 4.48 | 3.63 | 3.48 | 3.36 | N/R | N/R | 2.01-1.79 (9H) |
| $GlcN_3$ | 4.56 | 3.31 | 3.33 | N/R | N/R | N/R | - |

| | | | | | | | |
|---|---|---|---|---|---|---|---|
| Man**1** | 4.71 | 4.20 | 3.72 | 3.56 | N/R | N/R | **-** |
| Man**2** | 5.22 | 4.23 | 3.89 | 3.57 | N/R | N/R | **-** |
| Man**3** | 4.84 | 4.04 | 3.82 | 3.44 | 3.54 | N/R | **-** |
| Fuc**1** | 4.73 | 3.67 | 3.71 | 3.60 | 3.94 | 1.05 | **-** |

| **Signal** | **Proton** | **Carbon** |
|---|---|---|
| **Aromatic** | 7.91-7.84 (m, 4H)<br>7.53-7.47 (m, 3H) | 128.4-125.7 |
| $\mathbf{CH_2}$Ph | 5.33 (d, *J* = 12.8 Hz)<br>5.14 (d, *J* = 12.7 Hz) | 66.9 |
| NH-**CH**-COOH | 4.29 (dd, *J* = 9.5, 3.6 Hz) | 53.1 |
| C(O)-$\mathbf{CH_2}$-CH | 2.75 (dd, *J* = 14.8, 2.9 Hz)<br>2.51 (dd, *J* = 15.0, 9.9 Hz) | 38.6 |

**$^{13}$C NMR (150 MHz, $D_2O$): δ (ppm)**

| | **C1** |
|---|---|
| GlcNAc**1** | 78.2 |
| GlcNAc**2** | 100.9 |
| GlcNAc**3** | 100.1 |
| $GlcN_3$ | 99.6 |
| Man**1** | 100.4 |
| Man**2** | 99.6 |
| Man**3** | 97.1 |
| Fuc**1** | 99.2 |

[a] Not reported

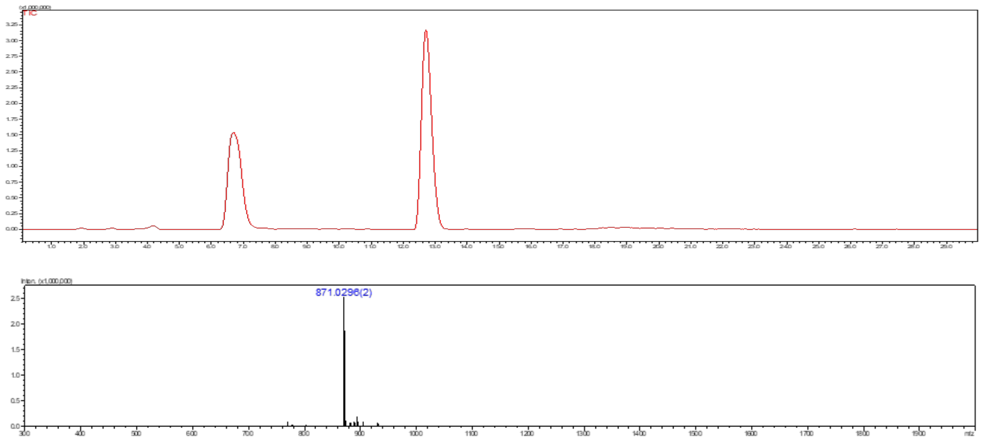

**Figure S17.** Liquid chromatography (LC) trace and mass (ESI-MS) spectrum (peak at retention time = 12.8 min) of **18**.

### Compound 21

**21** was prepared starting from **2** (3 mg, 1.48 μmol) by first installing a β1,4 GlcNAc using the general procedure **3d** for GnT-IVB. The reaction was monitored by ESI-MS and after complete conversion to MGAT4 arm installed intermediate [confirmed by LC-ESI-MS, peak at retention time = 12.5 min, m/z = 1007.5259(1)]. To the resulting compound was subjected for the installation of a β1,4 GlcNAc using the general procedure **3c** to obtain **21** as a white solid (2.2 mg, 66% over 2 steps).

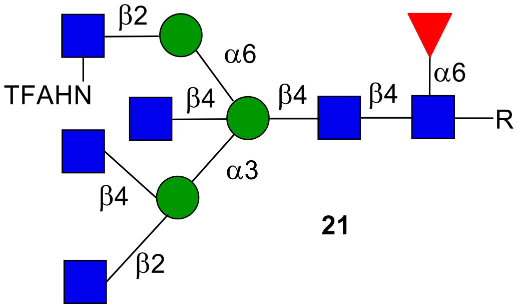

**21**

**$^1$H NMR (600 MHz, $D_2O$): δ (ppm)**

| | H1 | H2 | H3 | H4 | H5 | H6 | NHAc |
|---|---|---|---|---|---|---|---|
| GlcNAc**1** | 4.86 | 3.69 | 3.44 | 3.39 | N/R[a] | N/R | 2.01-1.79 (15H) |
| GlcNAc**2** | 4.54 | 3.71 | 3.48 | 3.63 | N/R | N/R | 2.01-1.79 (15H) |

| | | | | | | | |
|---|---|---|---|---|---|---|---|
| GlcNAc**3** | 4.46 | 3.65 | 3.51 | 3.38 | N/R | N/R | 2.01-1.79 (15H) |
| GlcNAc**4** | 4.43 | 3.68 | 3.49 | 3.43 | N/R | N/R | 2.01-1.79 (15H) |
| GlcNAc**5** | 4.37 | 3.62 | 3.33 | 3.18 | 3.49 | N/R | 2.01-1.79 (15H) |
| GlcNTFA | 4.60 | 3.74 | 3.54 | 3.44 | N/R | N/R | - |
| Man**1** | 4.59 | 4.07 | 3.77 | 3.97 | 3.52 | N/R | - |
| Man**2** | 4.97 | 4.21 | 3.97 | 3.52 | 3.65 | 3.46 | - |
| Man**3** | 4.89 | 4.10 | 3.77 | 3.35 | 3.53 | N/R | - |
| Fuc**1** | 4.73 | 3.66 | 3.73 | 3.61 | 3.94 | 1.06 | - |

| **Signal** | **Proton** | **Carbon** |
|---|---|---|
| **Aromatic** | 7.91-7.85 (m, 4H)<br>7.53-7.48 (m, 3H) | 128.4-125.9 |
| **CH$_2$**Ph | 5.33 (d, *J* = 12.8 Hz)<br>5.14 (d, *J* = 12.7 Hz) | 66.8 |
| NH-**CH**-COOH | 4.29 (dd, *J* = 9.5, 3.6 Hz) | 53.2 |
| C(O)-**CH$_2$**-CH | 2.75 (dd, *J* = 14.8, 2.9 Hz)<br>2.51 (dd, *J* = 15.0, 9.9 Hz) | 38.9 |

**$^{13}$C NMR (150 MHz, D$_2$O): δ (ppm)**

| | **C1** |
|---|---|
| GlcNAc**1** | 78.2 |
| GlcNAc**2** | 100.9 |
| GlcNAc**3** | 99.9 |
| GlcNAc**4** | 101.8 |
| GlcNAc**4** | 100.6 |
| GlcNTFA | 98.8 |
| Man**1** | 100.1 |
| Man**2** | 99.6 |
| Man**3** | 97.5 |
| Fuc**1** | 99.2 |

[a] Not reported

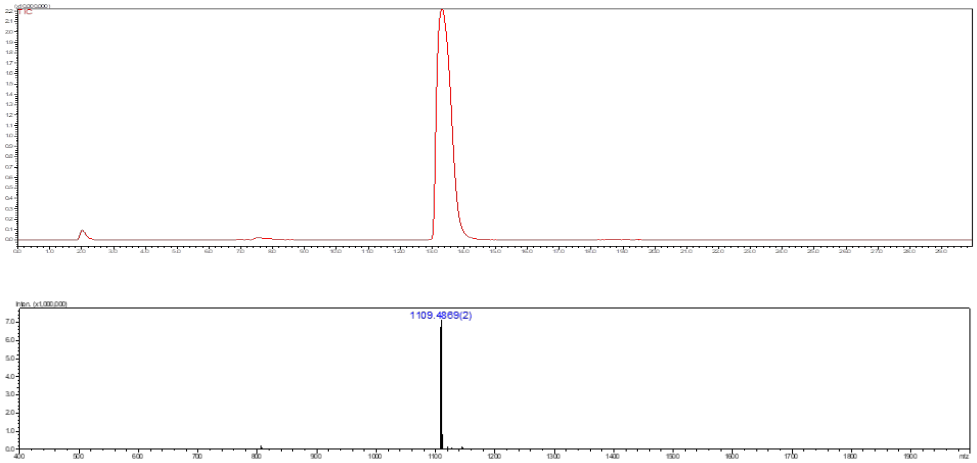

**Figure S18.** Liquid chromatography (LC) trace and mass (ESI-MS) spectrum (peak at retention time = 13.5 min) of **21**.

#### Compound 22

**22** was prepared starting from **3** (14.2 mg, 7.82 μmol) by first installing a β1,4 GlcNAc using the general procedure **3e** for GnT-V. The reaction was monitored by ESI-MS and after complete conversion to MGAT5 installed intermediate [confirmed by LC-ESI-MS, peak at retention time = 12.8 min, m/z = 1007.5259(1)], then fraction containing compound was lyophilized. To the resulting compound was subjected for the installation of a β1,4 GlcNAc using the general procedure **3c** to obtain **22** as a white solid (10.2 mg, 59% over 2 steps).

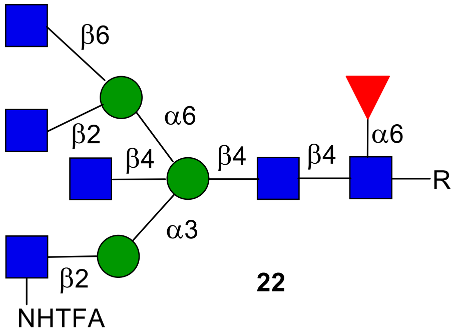

**22**

**$^{1}$H NMR (600 MHz, $D_2O$): δ (ppm)**

| | H1 | H2 | H3 | H4 | H5 | H6 | NHAc |
|---|---|---|---|---|---|---|---|
| GlcNAc**1** | 4.86 | 3.69 | 3.45 | 3.39 | N/R[a] | N/R | 2.02-1.79 (15H) |
| GlcNAc**2** | 4.53 | 3.73 | 3.48 | 3.65 | N/R | N/R | 2.02-1.79 (15H) |

| | | | | | | | |
|---|---|---|---|---|---|---|---|
| GlcNTFA | 4.66 | 3.75 | 3.47 | N/R | N/R | N/R | - |
| GlcNAc**3** | 4.42 | 3.63 | 3.32 | 3.21 | 3.54 | N/R | 2.02-1.79 (15H) |
| GlcNAc**4** | 4.49 | 3.60 | 3.39 | 3.32 | N/R | N/R | 2.02-1.79 (15H) |
| GlcNAc**5** | 4.45 | 3.67 | 3.51 | 3.41 | N/R | N/R | 2.02-1.79 (15H) |
| Man**1** | 4.62 | 4.10 | 3.77 | 4.05 | 3.45 | N/R | - |
| Man**2** | 4.93 | 4.18 | 3.84 | 3.35 | N/R | N/R | - |
| Man**3** | 4.83 | 4.05 | 3.73 | 3.35 | N/R | N/R | - |
| Fuc**1** | 4.73 | 3.67 | 3.72 | 3.61 | 3.94 | 1.06 | - |

| **Signal** | **Proton** | **Carbon** |
|---|---|---|
| **Aromatic** | 7.91-7.84 (m, 4H)<br>7.53-7.47 (m, 3H) | 128.4-125.7 |
| **$CH_2$**Ph | 5.33 (d, *J* = 12.8 Hz)<br>5.14 (d, *J* = 12.7 Hz) | 66.8 |
| NH-**CH**-COOH | 4.29 (dd, *J* = 9.5, 3.6 Hz) | 53.0 |
| C(O)-**$CH_2$**-CH | 2.75 (dd, *J* = 14.8, 2.9 Hz)<br>2.51 (dd, *J* = 15.0, 9.9 Hz) | 38.6 |

**$^{13}C$ NMR (150 MHz, $D_2O$): δ (ppm)**

| | **C1** |
|---|---|
| GlcNAc**1** | 78.2 |
| GlcNAc**2** | 101.0 |
| GlcNTFA | 98.9 |
| GlcNAc**3** | 99.7 |
| GlcNAc**4** | 99.7 |
| GlcNAc**5** | 101.7 |
| Man**1** | 99.9 |
| Man**2** | 99.9 |
| Man**3** | 98.0 |
| Fuc**1** | 99.2 |

[a] Not reported

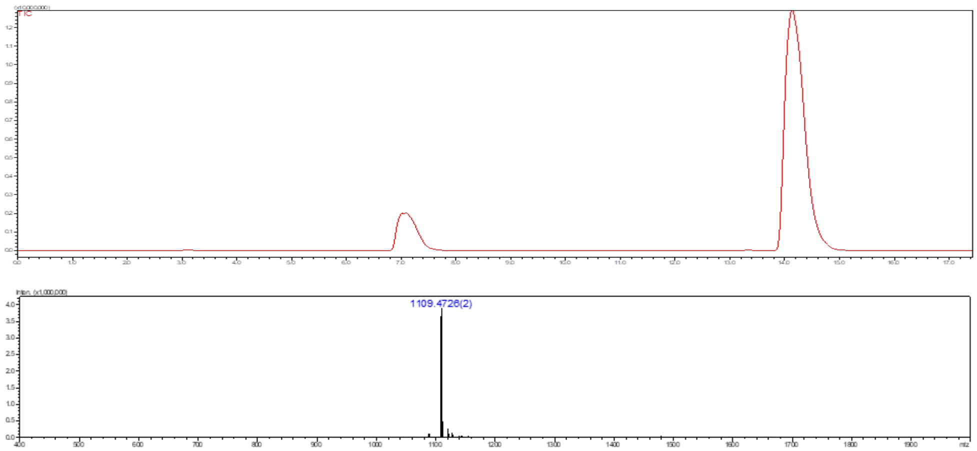

**Figure S19.** Liquid chromatography (LC) trace and mass (ESI-MS) spectrum (peak at retention time = 14.3 min) of **22**.

#### Compound 25

*N*-Glycan **6** (10.1 mg, 5.10 µmol) was dissolved in deionized water with the final reaction concentration of 5 mM. The pH of the reaction solution was adjusted to 10 using NaOH (1 M), and the mixture solution was incubated for 3-5 h at 37 °C with shaking. After complete conversion of starting material to amine intermediate **23** [confirmed by LC-ESI-MS, peak at retention time = 15.8 min, m/z = 959.5598(1)]. Then, reaction was neutralized by aqueous acetic acid (1 M). The product was purified by C18 biogel [Eluent - water and water:acetonitrile (8:2)]. Next, compound **23** was subjected for the installation of a β1,4 Gal using the general procedure **3f** to obtained **24** as a white solid. The amine of *N*-glycan compound **24** was subjected for the acetylation using a general protocol **3j** to obtain **25** as a white solid (5.4 mg, 51% over 3 steps).

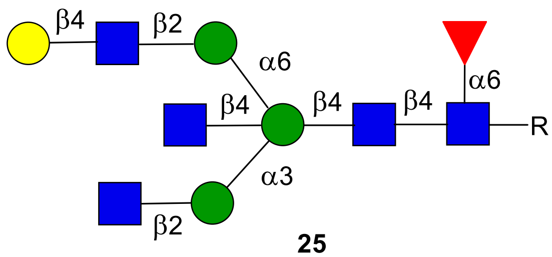

**25**

**$^1$H NMR (900 MHz, $D_2O$): δ (ppm)**

| | **H1** | **H2** | **H3** | **H4** | **H5** | **H6** | **NHAc** |
|---|---|---|---|---|---|---|---|
| GlcNAc**1** | 4.85 | 3.69 | 3.60 | 3.51 | N/R[a] | N/R | 2.02-1.79 (15H) |
| GlcNAc**2** | 4.53 | 3.72 | 3.64 | 3.67 | 3.49 | N/R | 2.02-1.79 (15H) |
| GlcNAc**3** | 4.50 | 3.67 | 3.56 | N/R | N/R | N/R | 2.02-1.79 (15H) |
| GlcNAc**4** | 4.39 | 3.61 | 3.33 | 3.19 | 3.49 | N/R | 2.02-1.79 (15H) |
| GlcNAc**5** | 4.48 | 3.64 | 3.53 | 3.41 | N/R | N/R | 2.02-1.79 (15H) |
| Man**1** | 4.60 | 4.09 | 3.78 | 4.00 | 3.46 | N/R | - |
| Man**2** | 4.98 | 4.17 | 3.83 | 3.63 | 3.85 | 3.49 | - |
| Man**3** | 4.93 | 4.06 | 3.75 | 3.55 | 3.40 | N/R | - |
| Fuc**1** | 4.73 | 3.66 | 3.72 | 3.62 | 3.94 | 1.05 | - |
| Gal**1** | 4.40 | 3.47 | 3.66 | 3.85 | N/R | N/R | - |

| **Signal** | **Proton** | **Carbon** |
|---|---|---|
| **Aromatic** | 7.92-7.85 (m, 4H)<br>7.53-7.48 (m, 3H) | 128.3-125.7 |
| **$CH_2$**Ph | 5.32 (d, *J* = 12.8 Hz)<br>5.14 (d, *J* = 12.7 Hz) | 66.9 |
| NH-**CH**-COOH | 4.29 (dd, *J* = 9.5, 3.6 Hz) | 53.1 |
| C(O)-**$CH_2$**-CH | 2.75 (dd, *J* = 14.8, 2.9 Hz)<br>2.51 (dd, *J* = 15.0, 9.9 Hz) | 38.7 |

**$^{13}$C NMR (225 MHz, $D_2O$): δ (ppm)**

| | **C1** |
|---|---|
| GlcNAc**1** | 78.1 |
| GlcNAc**2** | 100.9 |
| GlcNAc**3** | 99.4 |
| GlcNAc**4** | 100.6 |
| GlcNAc**5** | 99.8 |

| | |
|---|---|
| Man**1** | 100.0 |
| Man**2** | 99.9 |
| Man**3** | 97.7 |
| Fuc**1** | 99.2 |
| Gal**1** | 102.9 |

[a] Not reported

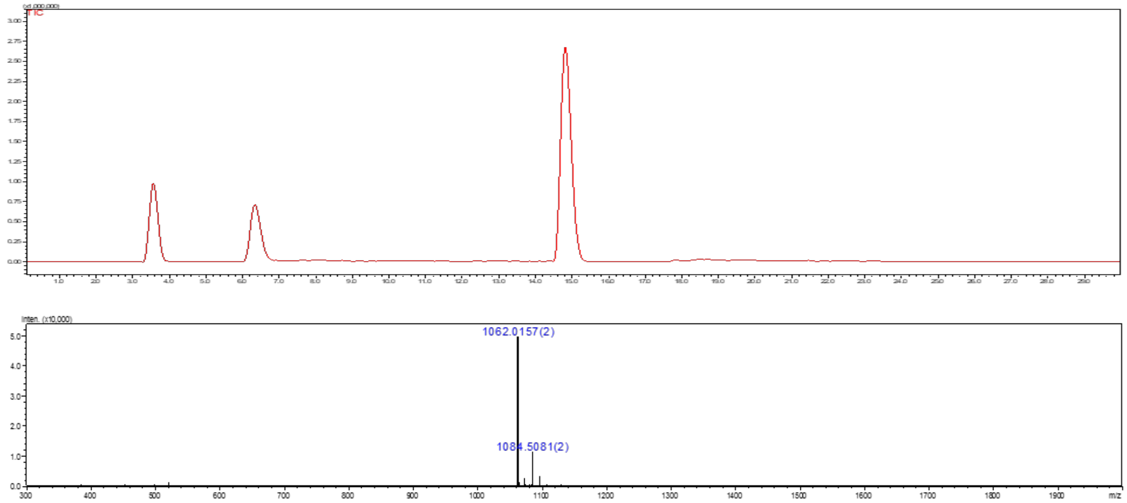

**Figure S20.** Liquid chromatography (LC) trace and mass (ESI-MS) spectrum (peak at retention time = 14.9 min) of **25.**

### Compound A

The amine of *N*-glycan compound **25** was subjected for the hydrogenation using a general protocol **3k** to obtain **A** as a white solid.

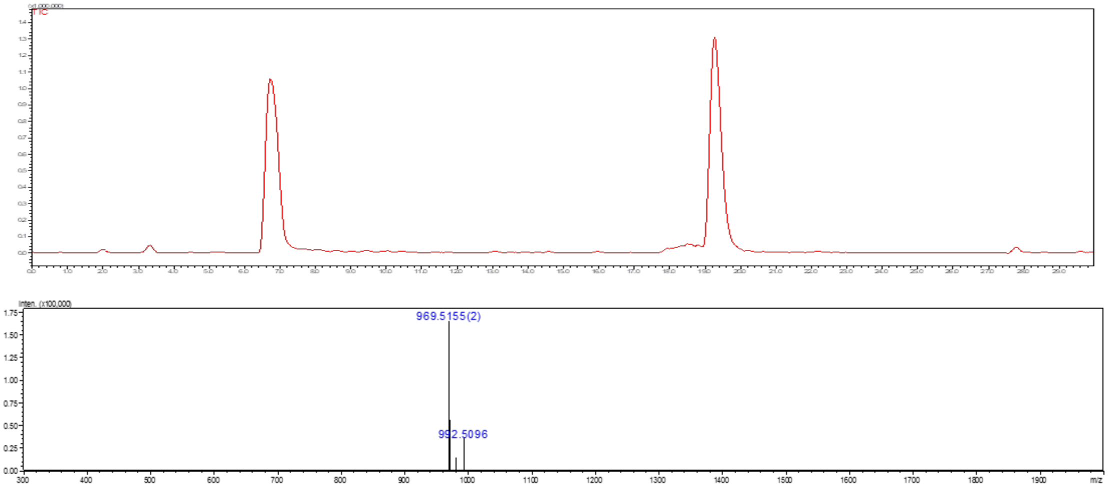

**Figure S21.** Liquid chromatography (LC) trace and mass (ESI-MS) spectrum (peak at retention time = 19.4 min) of **A**.

### Compound 26

**26** was synthesized from **25** (12 mg, 5.64 µmol) following the general protocol **3g** for the installation of Neu5Ac moiety using ST3Gal4. The product **26** was obtained as a white fluffy solid (10.2 mg, 75%).

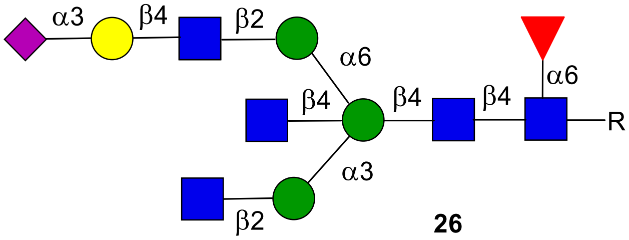

**26**

**$^1$H NMR (600 MHz, $D_2O$): δ (ppm)**

| | H1 | H2 | H3 | H4 | H5 | H6 | NHAc |
|---|---|---|---|---|---|---|---|
| GlcNAc**1** | 4.85 | 3.68 | 3.59 | 3.45 | N/R[a] | N/R | 2.02-1.79 (15H) |
| GlcNAc**2** | 4.53 | 3.71 | 3.63 | 3.66 | 3.48 | N/R | 2.02-1.79 (15H) |
| GlcNAc**3** | 4.49 | 3.67 | 3.56 | N/R | N/R | N/R | 2.02-1.79 (15H) |
| GlcNAc**4** | 4.38 | 3.62 | 3.32 | 3.20 | 3.49 | N/R | 2.02-1.79 (15H) |
| GlcNAc**5** | 4.47 | 3.65 | 3.51 | 3.40 | N/R | N/R | 2.02-1.79 (15H) |
| Man**1** | 4.60 | 4.09 | 3.80 | 4.00 | 3.45 | N/R | **-** |
| Man**2** | 4.98 | 4.16 | 3.83 | 3.62 | 3.84 | N/R | **-** |
| Man**3** | 4.93 | 4.07 | 3.76 | 3.54 | 3.40 | N/R | **-** |
| Fuc**1** | 4.73 | 3.67 | 3.71 | 3.61 | 3.94 | 1.07 | **-** |
| Gal**1** | 4.47 | 3.51 | 4.04 | 3.88 | N/R | N/R | **-** |

| | H1 | H2 | H3 | H4 | H5 | H6 | H7 | H8 | H9 |
|---|---|---|---|---|---|---|---|---|---|
| Neu5Ac9 | - | - | 2.68<br>1.71 | 3.61 | 3.77 | N/R | N/R | N/R | N/R |

| Signal | Proton | Carbon |
|---|---|---|
| **Aromatic** | 7.91-7.85 (m, 4H)<br>7.53-7.48 (m, 3H) | 128.3-125.7 |
| **$CH_2$**Ph | 5.33 (d, *J* = 12.8 Hz)<br>5.13 (d, *J* = 12.7 Hz) | 66.9 |
| NH-**CH**-COOH | 4.29 (dd, *J* = 9.5, 3.6 Hz) | 53.0 |
| C(O)-**$CH_2$**-CH | 2.75 (dd, *J* = 14.8, 2.9 Hz)<br>2.51 (dd, *J* = 15.0, 9.9 Hz) | 38.7 |

**$^{13}$C NMR (150 MHz, $D_2O$): δ (ppm)**

| | C1 |
|---|---|
| GlcNAc**1** | 78.2 |

| | |
|---|---|
| GlcNAc**2** | 100.9 |
| GlcNAc**3** | 99.4 |
| GlcNAc**4** | 100.6 |
| GlcNAc**5** | 99.8 |
| Man**1** | 100.0 |
| Man**2** | 99.9 |
| Man**3** | 97.8 |
| Fuc**1** | 99.1 |
| Gal**1** | 102.7 |

[a] Not reported

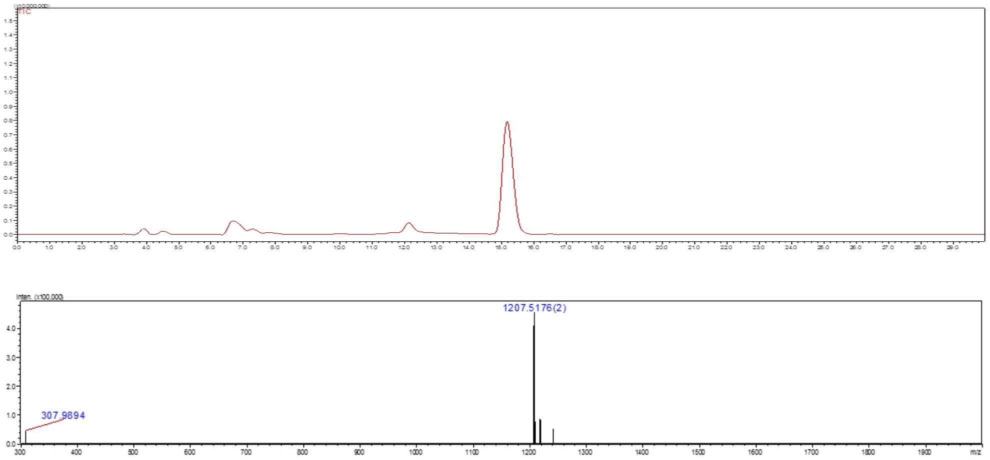

**Figure S22.** Liquid chromatography (LC) trace and mass (ESI-MS) spectrum (peak at retention time = 15.2 min) of **26**.

### Compound B

The amine of *N*-glycan compound **26** was subjected for the hydrogenation using a general protocol **3k** to obtain **B** as a white solid.

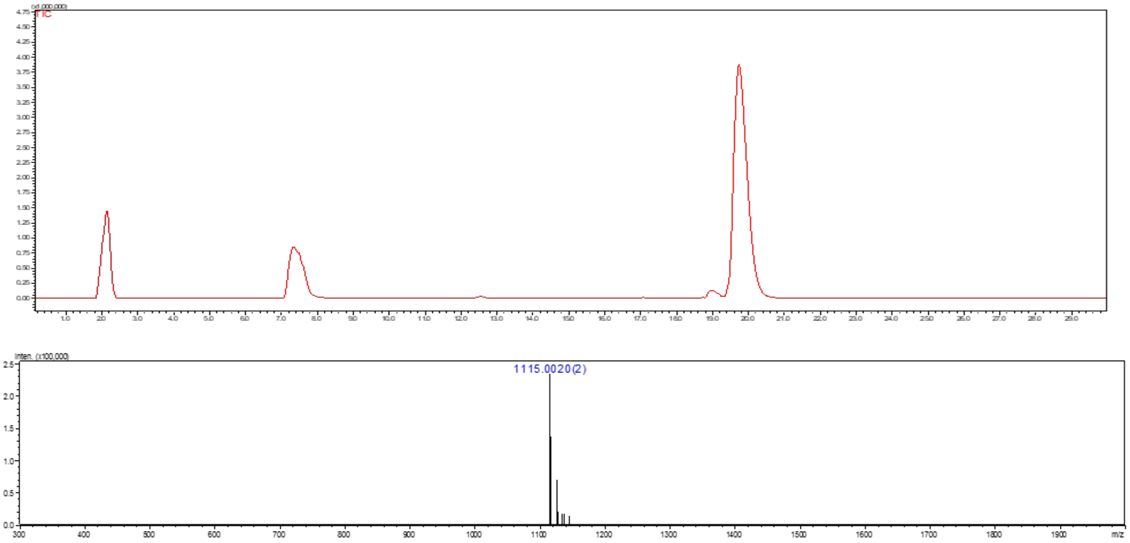

**Figure S23.** Liquid chromatography (LC) trace and mass (ESI-MS) spectrum (peak at retention time = 19.2 min) of **B**.

### Compound 27

**27** was synthesized from **25** (3 mg, 1.40 µmol) following the general protocol **3g** for the installation of Neu5Ac moiety using ST6Gal1. The product **27** was obtained as a white fluffy solid (2.9 mg, 86%).

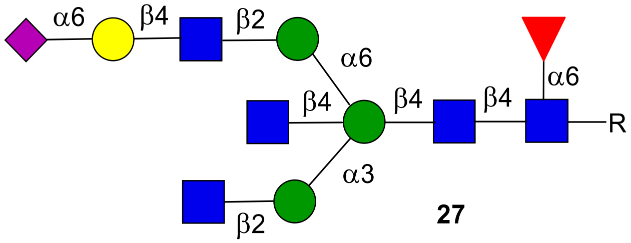

**27**

**$^{1}$H NMR (900 MHz, $D_2O$): δ (ppm)**

| | H1 | H2 | H3 | H4 | H5 | H6 | NHAc |
|---|---|---|---|---|---|---|---|
| GlcNAc**1** | 4.87 | 3.70 | 3.59 | 3.44 | 3.39 | N/R[a] | 2.02-1.79 (15H) |
| GlcNAc**2** | 4.54 | 3.70 | 3.64 | 3.65 | 3.49 | N/R | 2.02-1.79 (15H) |
| GlcNAc**3** | 4.53 | 3.67 | 3.60 | N/R | N/R | N/R | 2.02-1.79 (15H) |
| GlcNAc**4** | 4.40 | 3.63 | 3.35 | 3.19 | 3.50 | N/R | 2.02-1.79 (15H) |
| GlcNAc**5** | 4.49 | 3.65 | 3.53 | 3.40 | N/R | N/R | 2.02-1.79 (15H) |
| Man**1** | 4.61 | 4.11 | 3.81 | 4.09 | 3.49 | N/R | - |
| Man**2** | 4.99 | 4.18 | 3.83 | 3.63 | 3.41 | N/R | - |
| Man**3** | 4.94 | 4.08 | 3.78 | 3.54 | 3.43 | N/R | - |
| Fuc**1** | 4.75 | 3.67 | 3.71 | 3.61 | 3.95 | 1.07 | - |
| Gal**1** | 4.38 | 3.47 | 3.59 | 3.87 | N/R | N/R | - |

| | H1 | H2 | H3 | H4 | H5 | H6 | H7 | H8 | H9 |
|---|---|---|---|---|---|---|---|---|---|
| Neu5Ac9 | - | - | 2.60<br>1.65 | 3.58 | 3.73 | N/R | N/R | N/R | N/R |

| Signal | Proton | Carbon |
|---|---|---|
| **Aromatic** | 7.92-7.85 (m, 4H)<br>7.53-7.48 (m, 3H) | 128.4-125.7 |
| **$CH_2$**Ph | 5.33 (d, *J* = 12.8 Hz)<br>5.14 (d, *J* = 12.7 Hz) | 66.8 |
| NH-**CH**-COOH | 4.29 (dd, *J* = 9.5, 3.6 Hz) | 53.0 |
| C(O)-**$CH_2$**-CH | 2.75 (dd, *J* = 14.8, 2.9 Hz)<br>2.51 (dd, *J* = 15.0, 9.9 Hz) | 38.7 |

**$^{13}$C NMR (225 MHz, $D_2O$): δ (ppm)**

| | **C1** |
|---|---|
| GlcNAc**1** | 78.2 |
| GlcNAc**2** | 100.9 |
| GlcNAc**3** | 99.2 |
| GlcNAc**4** | 100.6 |
| GlcNAc**5** | 100.1 |
| Man**1** | 100.2 |
| Man**2** | 100.0 |
| Man**3** | 97.5 |
| Fuc**1** | 99.3 |
| Gal**1** | 103.6 |

[a] Not reported

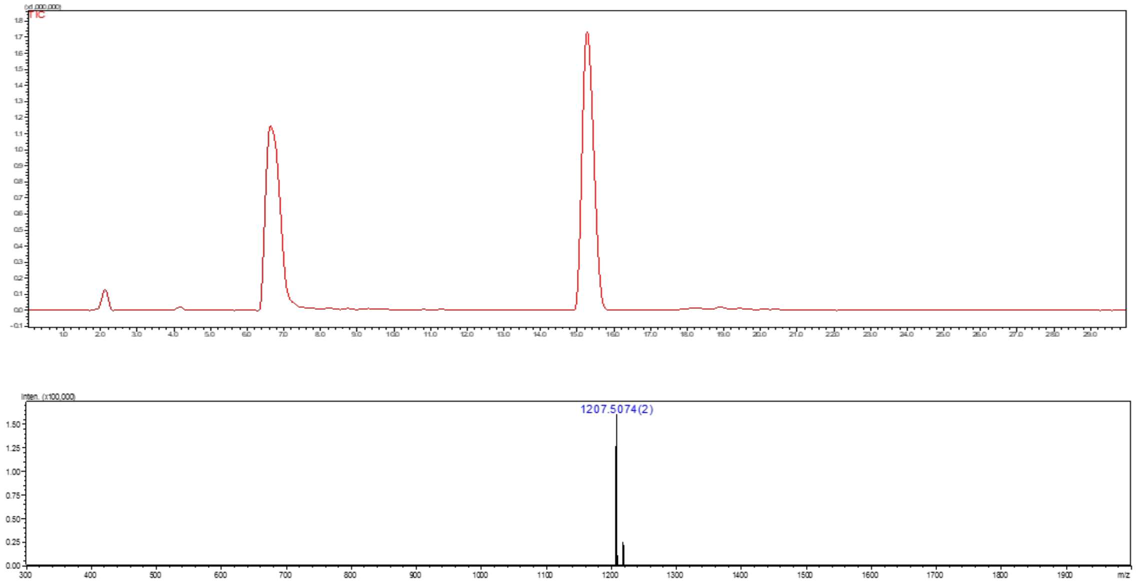

**Figure S24.** Liquid chromatography (LC) trace and mass (ESI-MS) spectrum (peak at retention time = 15.4 min) of **27**.

**Compound C**

The amine of *N*-glycan compound **27** was subjected for the hydrogenation using a general protocol **3k** to obtain **C** as a white solid.

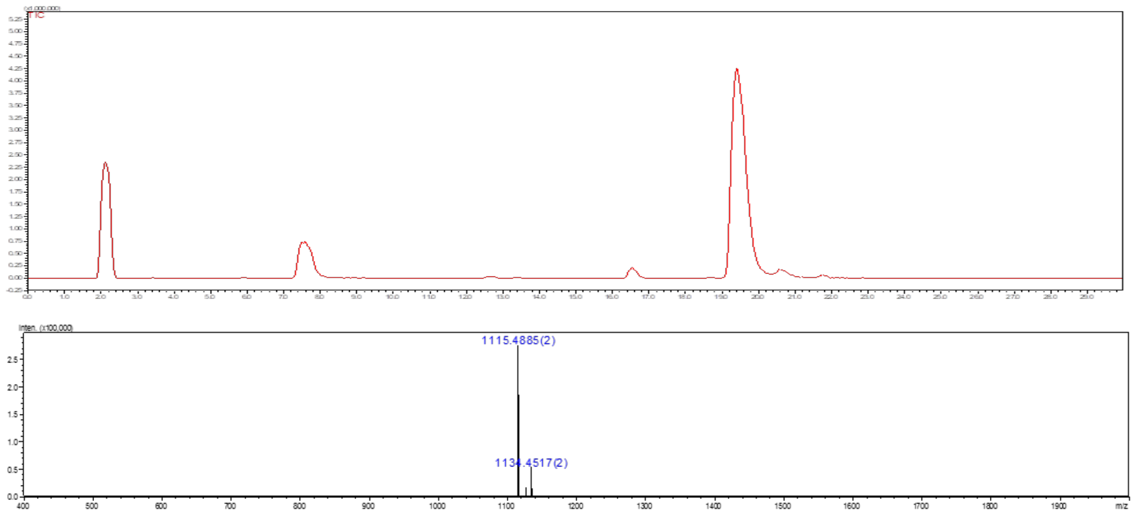

**Figure S25.** Liquid chromatography (LC) trace and mass (ESI-MS) spectrum (peak at retention time = 19.5 min) of **C**

**Compound 28**

**28** was synthesized from **26** (1 mg, 0.41 μmol) following the general protocol **3h** for the installation of Fuc moiety using FUT6. The product **28** was obtained as a white fluffy solid (0.85 mg, 80%).

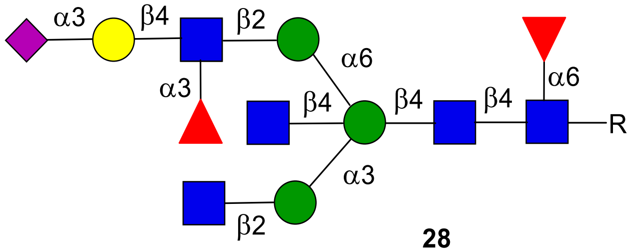

**28**

**$^1$H NMR (600 MHz, $D_2O$): δ (ppm)**

| | **H1** | **H2** | **H3** | **H4** | **H5** | **H6** | **NHAc** |
|---|---|---|---|---|---|---|---|
| GlcNAc**1** | 4.88 | 3.70 | 3.62 | 3.44 | N/R[a] | N/R | 2.02-1.79 (15H) |
| GlcNAc**2** | 4.54 | 3.73 | 3.65 | 3.15 | N/R | N/R | 2.02-1.79 (15H) |
| GlcNAc**3** | 4.50 | 3.85 | 3.50 | N/R | N/R | N/R | 2.02-1.79 (15H) |
| GlcNAc**4** | 4.39 | 3.62 | 3.35 | 3.19 | 3.49 | N/R | 2.02-1.79 (15H) |
| GlcNAc**5** | 4.48 | 3.65 | 3.52 | 3.40 | N/R | N/R | 2.02-1.79 (15H) |
| Man**1** | 4.62 | 4.11 | 3.79 | 3.98 | 3.50 | N/R | **-** |
| Man**2** | 4.98 | 4.17 | 3.83 | 3.64 | 3.40 | N/R | **-** |
| Man**3** | 4.94 | 4.06 | 3.78 | 3.54 | 3.40 | N/R | **-** |
| Fuc**1** | 4.74 | 3.65 | 3.72 | 3.61 | 3.95 | 1.07 | **-** |
| Gal**1** | 4.44 | 3.44 | 4.00 | 3.85 | N/R | N/R | **-** |
| Fuc**2** | 5.05 | 3.61 | 3.84 | 3.71 | 4.75 | 1.10 | **-** |

| | **H1** | **H2** | **H3** | **H4** | **H5** | **H6** | **H7** | **H8** | **H9** |
|---|---|---|---|---|---|---|---|---|---|
| Neu5Ac9 | - | - | 2.68<br>1.72 | 3.61 | 3.76 | N/R | N/R | N/R | N/R |

| **Signal** | **Proton** | **Carbon** |
|---|---|---|
| **Aromatic** | 7.91-7.85 (m, 4H)<br>7.53-7.47 (m, 3H) | 128.4-125.7 |
| **$CH_2$**Ph | 5.32 (d, *J* = 12.8 Hz)<br>5.16 (d, *J* = 12.7 Hz) | 66.9 |
| NH-**CH**-COOH | 4.38 (dd, *J* = 9.5, 3.6 Hz) | 53.1 |
| C(O)-**$CH_2$**-CH | 2.77 (dd, *J* = 14.8, 2.9 Hz)<br>2.59 (dd, *J* = 15.0, 9.9 Hz) | 38.6 |

**$^{13}$C NMR (150 MHz, $D_2O$): δ (ppm)**

| | **C1** |
|---|---|
| GlcNAc**1** | 78.2 |
| GlcNAc**2** | 100.9 |
| GlcNAc**3** | 98.9 |
| GlcNAc**4** | 100.4 |
| GlcNAc**5** | 99.9 |
| Man**1** | 100.2 |
| Man**2** | 99.9 |
| Man**3** | 97.3 |
| Fuc**1** | 99.2 |
| Gal**1** | 101.6 |
| Fuc**2** | 98.4 |

[a] Not reported

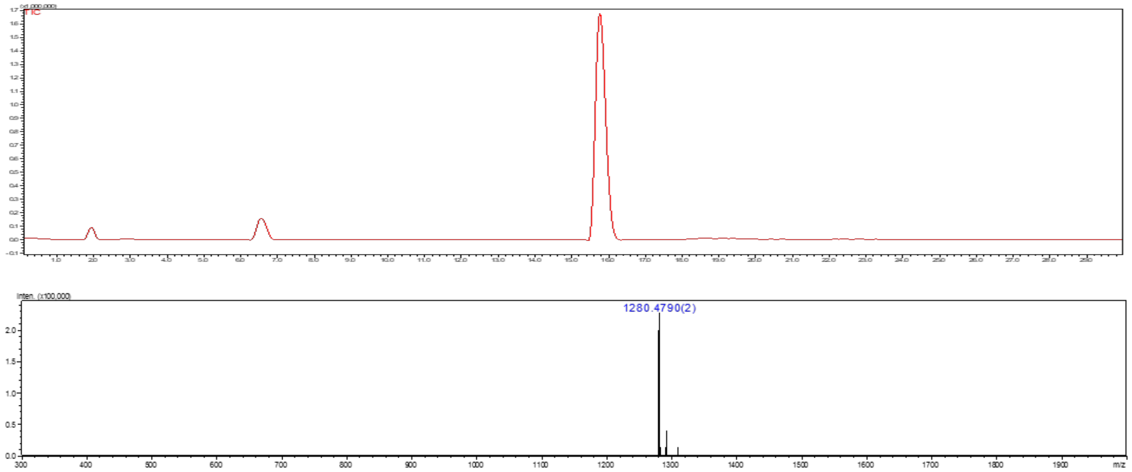

**Figure S26.** Liquid chromatography (LC) trace and mass (ESI-MS) spectrum (peak at retention time = 15.9 min) of **28**.

**Compound E**

The amine of *N*-glycan compound **28** was subjected for the hydrogenation using a general protocol **3k** to obtain **E** as a white solid.

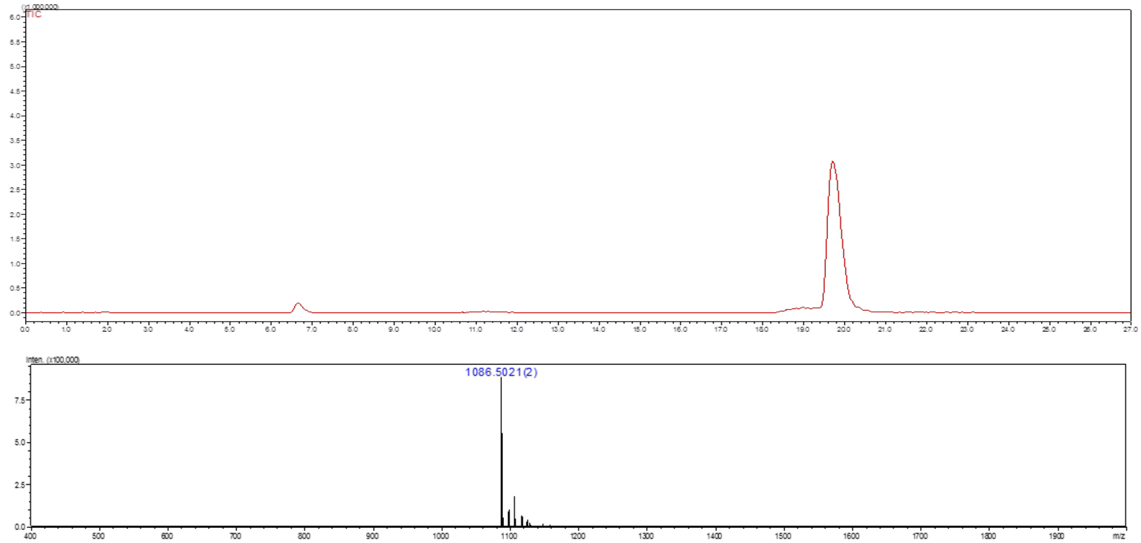

**Figure S27.** Liquid chromatography (LC) trace and mass (ESI-MS) spectrum (peak at retention time = 19.8 min) of **E**.

**Compound 29**

**29** was synthesized from **26** (2 mg, 0.82 μmol) following the general protocol **3f** for the installation of Gal moiety using B4GalT1. The product **29** was obtained as a white fluffy solid (1.45 mg, 68%).

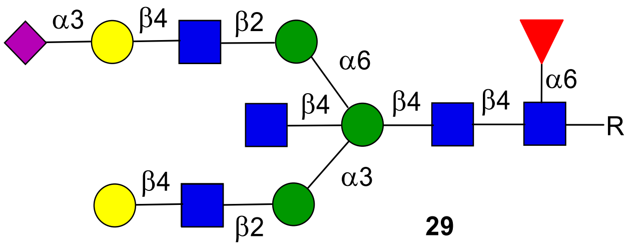

**29**

**$^{1}$H NMR (600 MHz, $D_2O$): δ (ppm)**

| | **H1** | **H2** | **H3** | **H4** | **H5** | **H6** | **NHAc** |
|---|---|---|---|---|---|---|---|
| GlcNAc**1** | 4.85 | 3.69 | 3.59 | N/R[a] | N/R | N/R | 2.03-1.79 (15H) |
| GlcNAc**2** | 4.53 | 3.73 | 3.64 | 3.66 | 3.48 | N/R | 2.03-1.79 (15H) |
| GlcNAc**3** | 4.51 | 3.69 | 3.55 | N/R | N/R | N/R | 2.03-1.79 (15H) |
| GlcNAc**4** | 4.38 | 3.61 | 3.32 | 3.19 | 3.39 | N/R | 2.03-1.79 (15H) |
| GlcNAc**5** | 4.49 | 3.67 | 3.55 | 3.46 | N/R | N/R | 2.03-1.79 (15H) |
| Man**1** | 4.60 | 4.08 | 3.79 | 4.00 | 3.45 | N/R | **-** |
| Man**2** | 4.98 | 4.20 | 3.83 | 3.62 | N/R | N/R | **-** |
| Man**3** | 4.94 | 4.07 | 3.74 | 3.54 | 3.40 | N/R | **-** |
| Fuc**1** | 4.73 | 3.66 | 3.71 | 3.62 | 3.93 | 1.07 | **-** |
| Gal**1** | 4.38 | 3.47 | N/R | 3.84 | N/R | N/R | **-** |
| Gal**2** | 4.48 | 3.51 | 4.04 | 3.89 | N/R | N/R | **-** |

| | **H1** | **H2** | **H3** | **H4** | **H5** | **H6** | **H7** | **H8** | **H9** |
|---|---|---|---|---|---|---|---|---|---|
| Neu5Ac9 | **-** | **-** | 2.68<br>1.73 | 3.60 | 3.77 | N/R | N/R | N/R | N/R |

| **Signal** | **Proton** | **Carbon** |
|---|---|---|
| **Aromatic** | 7.92-7.85 (m, 4H)<br>7.53-7.48 (m, 3H) | 128.3-125.7 |
| **$CH_2$**Ph | 5.33 (d, *J* = 12.8 Hz)<br>5.14 (d, *J* = 12.7 Hz) | 66.8 |
| NH-**CH**-COOH | 4.29 (dd, *J* = 9.5, 3.6 Hz) | 53.0 |
| C(O)-**$CH_2$**-CH | 2.75 (dd, *J* = 14.8, 2.9 Hz)<br>2.51 (dd, *J* = 15.0, 9.9 Hz) | 38.6 |

**$^{13}C$ NMR (150 MHz, $D_2O$): δ (ppm)**

| | **C1** |
|---|---|
| GlcNAc**1** | 78.2 |
| GlcNAc**2** | 100.9 |
| GlcNAc**3** | 99.7 |
| GlcNAc**4** | 100.6 |
| GlcNAc**5** | 99.4 |
| Man**1** | 100.0 |
| Man**2** | 99.9 |
| Man**3** | 97.7 |
| Fuc**1** | 99.2 |
| Gal**1** | 102.9 |
| Gal**2** | 102.7 |

[a] Not reported

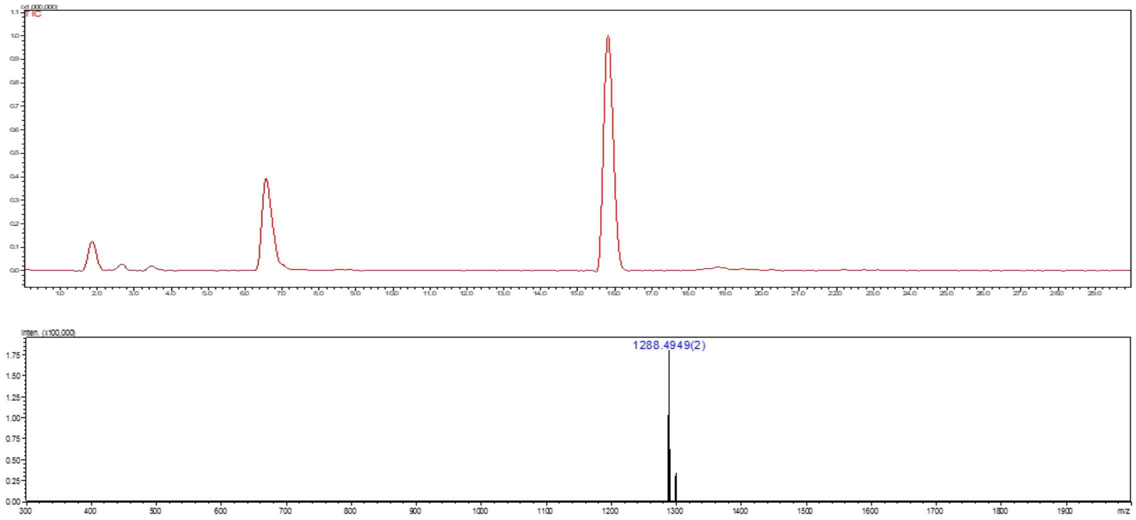

**Figure S28.** Liquid chromatography (LC) trace and mass (ESI-MS) spectrum (peak at retention time = 15.9 min) of **29**.

**Compound D**

The amine of *N*-glycan compound **29** was subjected for the hydrogenation using a general protocol **3k** to obtain **D** as a white solid.

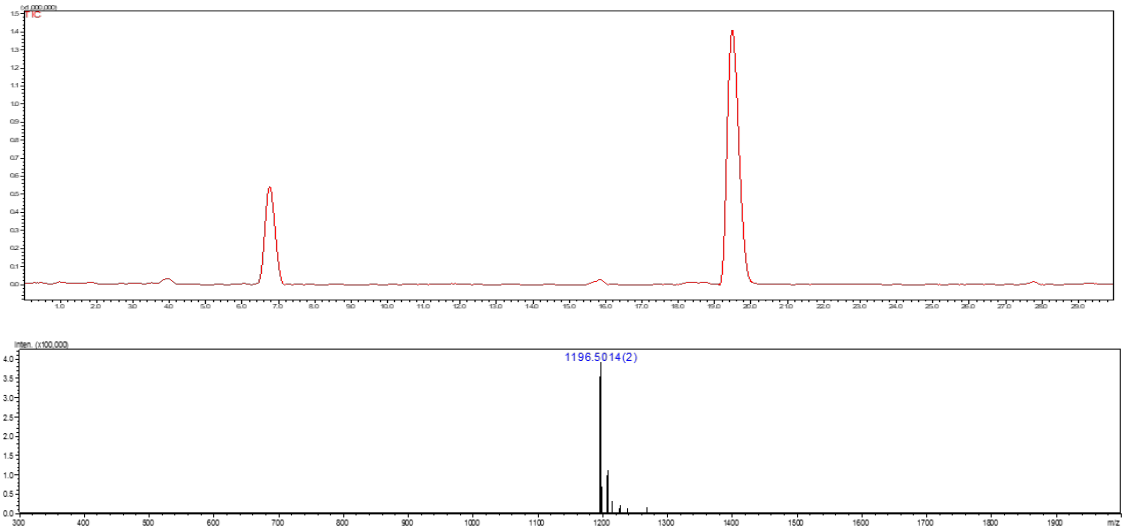

**Figure S29.** Liquid chromatography (LC) trace and mass (ESI-MS) spectrum (peak at retention time = 19.6 min) of **D**.

**Compound 7**

**7** was synthesized from **28** (1.5 mg, 0.58 μmol) following the general protocol **3f** for the installation of Gal moiety using B4GalT1. The product **7** was obtained as a white fluffy solid (0.99 mg, 63%).

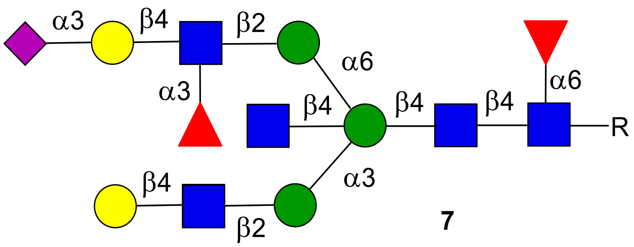

**7**

**$^1$H NMR (900 MHz, $D_2O$): δ (ppm)**

| | H1 | H2 | H3 | H4 | H5 | H6 | NHAc |
|---|---|---|---|---|---|---|---|
| GlcNAc**1** | 4.84 | 3.69 | 3.59 | N/R[a] | N/R | N/R | 2.03-1.79 (15H) |
| GlcNAc**2** | 4.52 | 3.73 | 3.65 | N/R | N/R | N/R | 2.03-1.79 (15H) |
| GlcNAc**3** | 4.50 | 3.69 | 3.50 | N/R | N/R | N/R | 2.03-1.79 (15H) |
| GlcNAc**4** | 4.39 | 3.62 | 3.33 | 3.18 | 3.49 | N/R | 2.03-1.79 (15H) |
| GlcNAc**5** | 4.49 | 3.68 | N/R | N/R | N/R | N/R | 2.03-1.79 (15H) |
| Man**1** | 4.61 | 4.10 | 3.78 | 3.98 | 3.50 | N/R | - |
| Man**2** | 4.97 | 4.17 | 3.83 | N/R | N/R | N/R | - |
| Man**3** | 4.94 | 4.05 | 3.76 | 3.54 | 3.40 | N/R | - |
| Fuc**1** | 4.72 | 3.65 | 3.71 | 3.62 | 3.93 | 1.06 | - |
| Fuc**2** | 5.04 | 3.62 | 3.84 | 3.65 | 3.75 | 1.10 | - |
| Gal**1** | 4.39 | 3.46 | 3.94 | N/R | N/R | N/R | - |
| Gal**2** | 4.45 | 3.43 | 4.00 | N/R | N/R | N/R | - |

| | H1 | H2 | H3 | H4 | H5 | H6 | H7 | H8 | H9 |
|---|---|---|---|---|---|---|---|---|---|
| Neu5Ac9 | - | - | 2.68<br>171 | 3.60 | 3.76 | N/R | N/R | N/R | N/R |

| Signal | Proton | Carbon |
|---|---|---|
| **Aromatic** | 7.92-7.85 (m, 4H)<br>7.52-7.47 (m, 3H) | 128.4-125.7 |
| **CH$_2$**Ph | 5.32 (d, *J* = 12.8 Hz)<br>5.14 (d, *J* = 12.7 Hz) | 66.8 |
| NH-**CH**-COOH | 4.29 (dd, *J* = 9.5, 3.6 Hz) | 53.1 |
| C(O)-**CH$_2$**-CH | 2.75 (dd, *J* = 14.8, 2.9 Hz)<br>2.51 (dd, *J* = 15.0, 9.9 Hz) | 38.8 |

**$^{13}C$ NMR (225 MHz, $D_2O$): δ (ppm)**

| | **C1** |
|---|---|
| GlcNAc**1** | 78.1 |
| GlcNAc**2** | 100.9 |
| GlcNAc**3** | 99.7 |
| GlcNAc**4** | 100.4 |
| GlcNAc**5** | 99.7 |
| Man**1** | 100.4 |
| Man**2** | 99.9 |
| Man**3** | 97.2 |
| Fuc**1** | 99.2 |
| Gal**1** | 102.8 |
| Gal**2** | 101.5 |
| Fuc**2** | 98.4 |

[a] Not reported

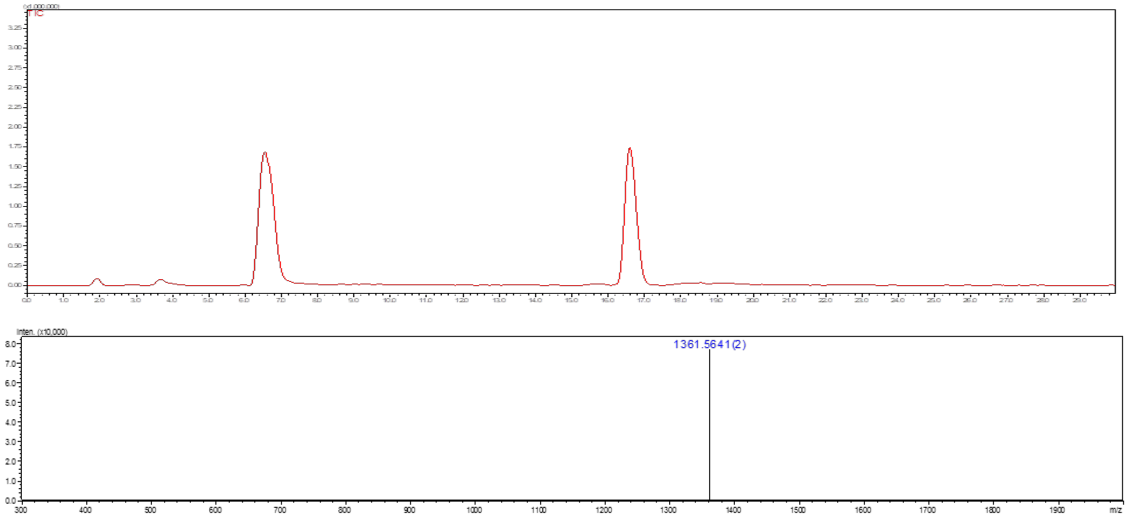

**Figure S30.** Liquid chromatography (LC) trace and mass (ESI-MS) spectrum (peak at retention time = 16.6 min) of **7**.

**Compound F**

The amine of *N*-glycan compound **7** was subjected for the hydrogenation using a general protocol **3k** to obtain **F** as a white solid.

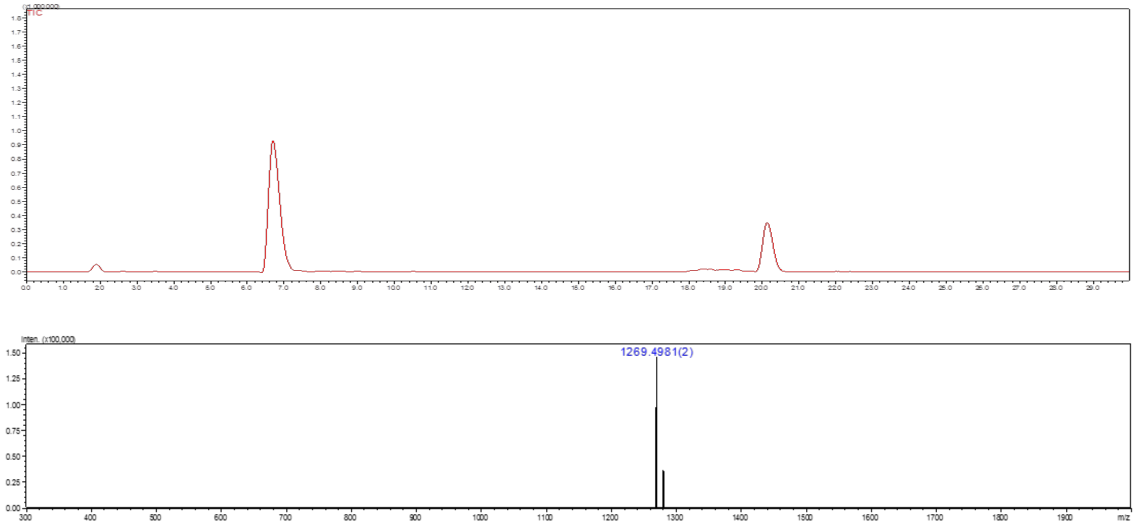

**Figure S31.** Liquid chromatography (LC) trace and mass (ESI-MS) spectrum (peak at retention time = 20.2 min) of **F**.

**Compound 30**

*N*-Glycan **4** (22 mg, 10.90 µmol) was dissolved in deionized water with the final reaction concentration of 5 mM. The pH of the reaction solution was adjusted to 10 using NaOH (1 M), and the mixture solution was incubated for 3-5 h at 37 °C with shaking. After complete conversion of starting material to amine intermediate **30** [confirmed by LC-ESI-MS, peak at retention time = 15.8 min, m/z = 959.5598(1)]. Then, reaction was neutralized by aqueous acetic acid (1 M). The product was purified by C18 biogel [Eluent - water and water:acetonitrile (8:2)]. Next, compound **30** was subjected for the installation of a β1,4 Gal using the general procedure **3f** to obtained **31** as a white solid which was subjected for the acetylation using a general protocol **3j** to obtain **32** as a white solid (11.9 mg, 55% over 3 steps).

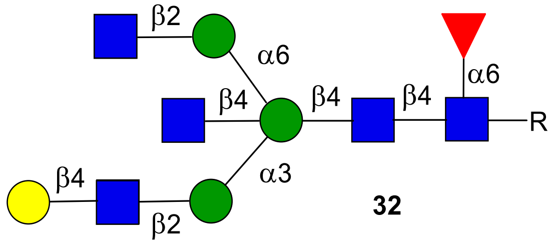

**32**

$^{1}$**H NMR (600 MHz, $D_2O$): δ (ppm)**

| | **H1** | **H2** | **H3** | **H4** | **H5** | **H6** | **NHAc** |
|---|---|---|---|---|---|---|---|
| GlcNAc**1** | 4.86 | 3.70 | 3.60 | 3.44 | 3.38 | N/R[a] | 2.02-1.79 (15H) |
| GlcNAc**2** | 4.54 | 3.73 | 3.65 | 3.64 | N/R | N/R | 2.02-1.79 (15H) |
| GlcNAc**3** | 4.50 | 3.69 | 3.54 | N/R | N/R | N/R | 2.02-1.79 (15H) |
| GlcNAc**4** | 4.39 | 3.63 | 3.33 | 3.19 | 3.49 | N/R | 2.02-1.79 (15H) |
| GlcNAc**5** | 4.48 | 3.64 | 3.41 | N/R | N/R | N/R | 2.02-1.79 (15H) |
| Man**1** | 4.61 | 4.11 | 3.80 | 3.66 | N/R | N/R | - |
| Man**2** | 4.98 | 4.18 | 3.83 | 3.63 | N/R | 3.52 | - |
| Man**3** | 4.93 | 4.07 | 3.76 | 3.55 | 3.40 | N/R | - |
| Fuc**1** | 4.73 | 3.67 | 3.72 | 3.62 | 3.94 | 1.06 | - |
| Gal**1** | 4.40 | 3.48 | 3.59 | 3.85 | N/R | N/R | - |

| **Signal** | **Proton** | **Carbon** |
|---|---|---|
| **Aromatic** | 7.92-7.84 (m, 4H)<br>7.53-7.48 (m, 3H) | 128.3-125.7 |
| **CH$_2$**Ph | 5.33 (d, $J$ = 12.8 Hz)<br>5.14 (d, $J$ = 12.7 Hz) | 66.8 |
| NH-**CH**-COOH | 4.29 (dd, $J$ = 9.5, 3.6 Hz) | 53.1 |
| C(O)-**CH$_2$**-CH | 2.75 (dd, $J$ = 14.8, 2.9 Hz)<br>2.51 (dd, $J$ = 15.0, 9.9 Hz) | 38.6 |

**$^{13}$C NMR (150 MHz, $D_2O$): δ (ppm)**

| | **C1** |
|---|---|
| GlcNAc**1** | 78.2 |
| GlcNAc**2** | 100.9 |
| GlcNAc**3** | 99.3 |
| GlcNAc**4** | 100.5 |
| GlcNAc**5** | 99.9 |
| Man**1** | 100.0 |
| Man**2** | 99.8 |
| Man**3** | 97.7 |
| Fuc**1** | 99.2 |
| Gal**1** | 103.0 |

[a] Not reported

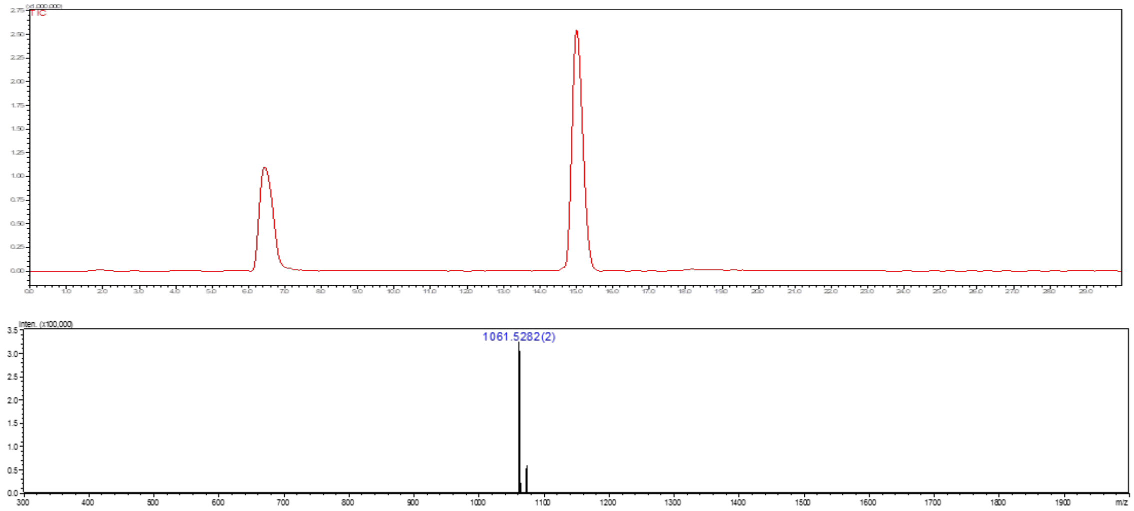

**Figure S32.** Liquid chromatography (LC) trace and mass (ESI-MS) spectrum (peak at retention time = 15.0 min) of **32**.

### Compound G

The amine of *N*-glycan compound **32** was subjected for the hydrogenation using a general protocol **3k** to obtain **G** as a white solid.

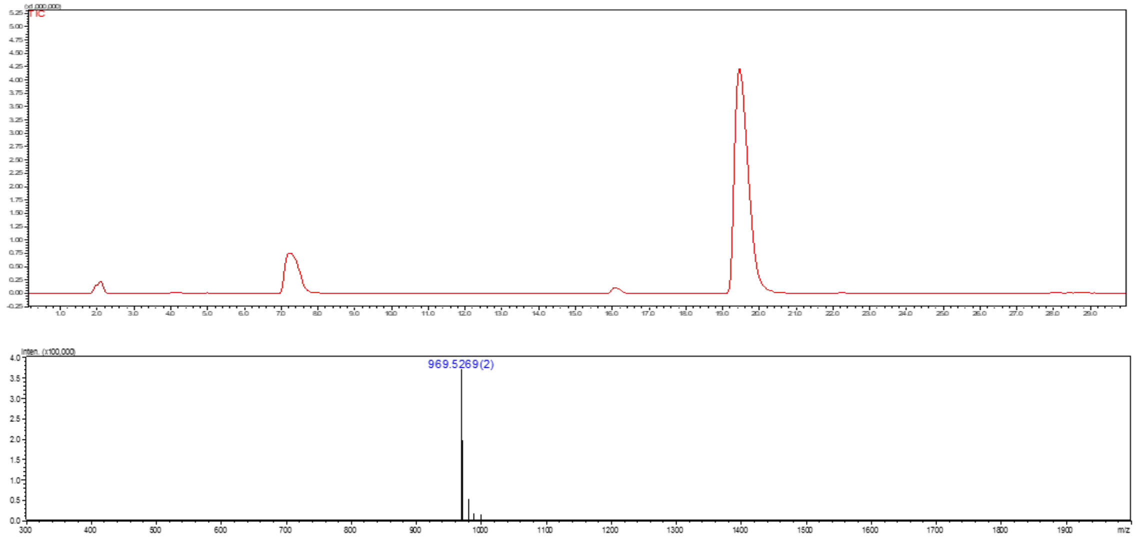

**Figure S33.** Liquid chromatography (LC) trace and mass (ESI-MS) spectrum (peak at retention time = 19.7 min) of **G**.

### Compound 33

**33** was synthesized from **32** (10 mg, 4.69 μmol) following the general protocol **3g** for the installation of Neu5Ac moiety using ST3Gal4. The product **33** was obtained as a white fluffy solid (9.0 mg, 79%).

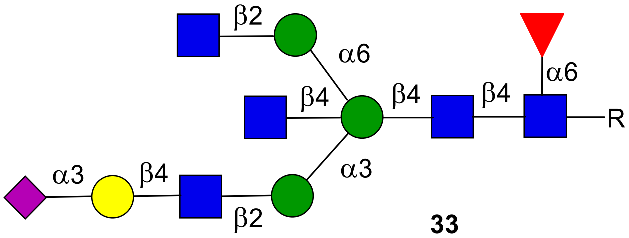

**33**

**$^1$H NMR (600 MHz, $D_2O$): δ (ppm)**

| | H1 | H2 | H3 | H4 | H5 | H6 | NHAc |
|---|---|---|---|---|---|---|---|
| GlcNAc**1** | 4.85 | 3.70 | 3.60 | 3.44 | N/R[a] | N/R | 2.02-1.79 (15H) |
| GlcNAc**2** | 3.54 | 3.72 | 3.64 | 3.64 | 3.48 | N/R | 2.02-1.79 (15H) |
| GlcNAc**3** | 4.50 | 3.68 | 3.49 | N/R | N/R | N/R | 2.02-1.79 (15H) |
| GlcNAc**4** | 4.38 | 3.62 | 3.34 | 3.18 | 3.50 | N/R | 2.02-1.79 (15H) |
| GlcNAc**5** | 4.47 | 3.64 | 3.54 | N/R | N/R | N/R | 2.02-1.79 (15H) |
| Man**1** | 4.62 | 4.10 | 3.80 | 3.98 | 3.47 | N/R | - |
| Man**2** | 4.98 | 4.18 | 3.83 | 3.62 | N/R | N/R | - |
| Man**3** | 4.92 | 4.08 | 3.78 | 3.54 | 3.41 | N/R | - |
| Fuc**1** | 4.73 | 3.66 | 3.72 | 3.61 | 3.94 | 1.05 | - |
| Gal**1** | 4.47 | 3.49 | 4.04 | 3.89 | N/R | N/R | - |

| | H1 | H2 | H3 | H4 | H5 | H6 | H7 | H8 | H9 |
|---|---|---|---|---|---|---|---|---|---|
| Neu5Ac9 | - | - | 2.68<br>1.73 | 3.61 | 3.77 | N/R | N/R | N/R | N/R |

| Signal | Proton | Carbon |
|---|---|---|
| **Aromatic** | 7.92-7.85 (m, 4H)<br>7.52-7.48 (m, 3H) | 128.4-125.8 |
| **$CH_2$**Ph | 5.33 (d, *J* = 12.8 Hz)<br>5.14 (d, *J* = 12.7 Hz) | 66.7 |
| NH-**CH**-COOH | 4.29 (dd, *J* = 9.5, 3.6 Hz) | 53.1 |
| C(O)-**$CH_2$**-CH | 2.75 (dd, *J* = 14.8, 2.9 Hz)<br>2.51 (dd, *J* = 15.0, 9.9 Hz) | 38.7 |

**$^{13}$C NMR (150 MHz, $D_2O$): δ (ppm)**

| | **C1** |
|---|---|
| GlcNAc**1** | 78.0 |
| GlcNAc**2** | 100.8 |
| GlcNAc**3** | 99.7 |
| GlcNAc**4** | 100.6 |
| GlcNAc**5** | 99.5 |
| Man**1** | 100.1 |
| Man**2** | 99.9 |
| Man**3** | 97.6 |
| Fuc**1** | 99.2 |
| Gal**1** | 102.6 |

[a] Not reported

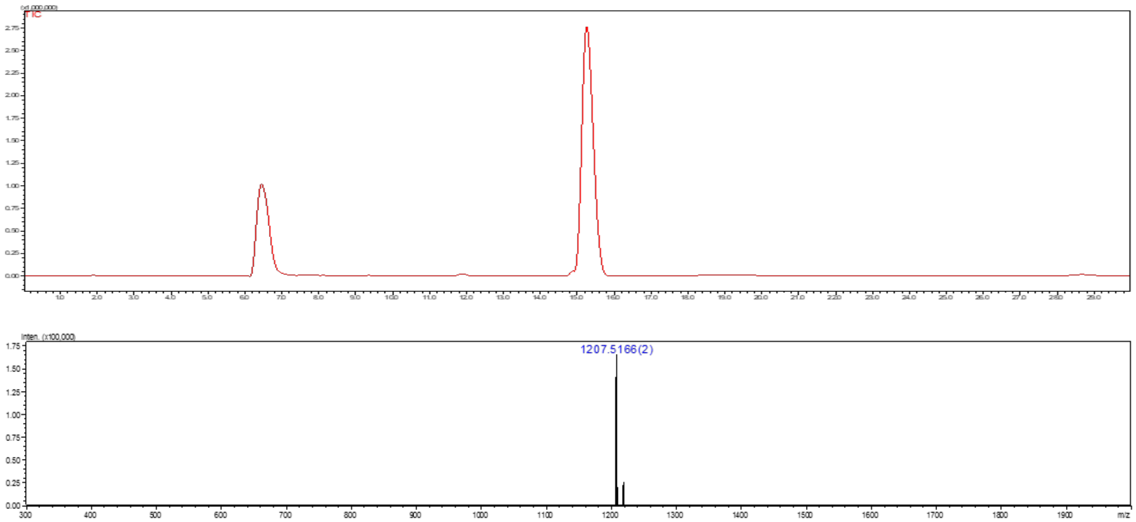

**Figure S34.** Liquid chromatography (LC) trace and mass (ESI-MS) spectrum (peak at retention time = 15.4 min) of **33**.

**Compound H**

The amine of *N*-glycan compound **33** was subjected for the hydrogenation using a general protocol **3k** to obtain **H** as a white solid.

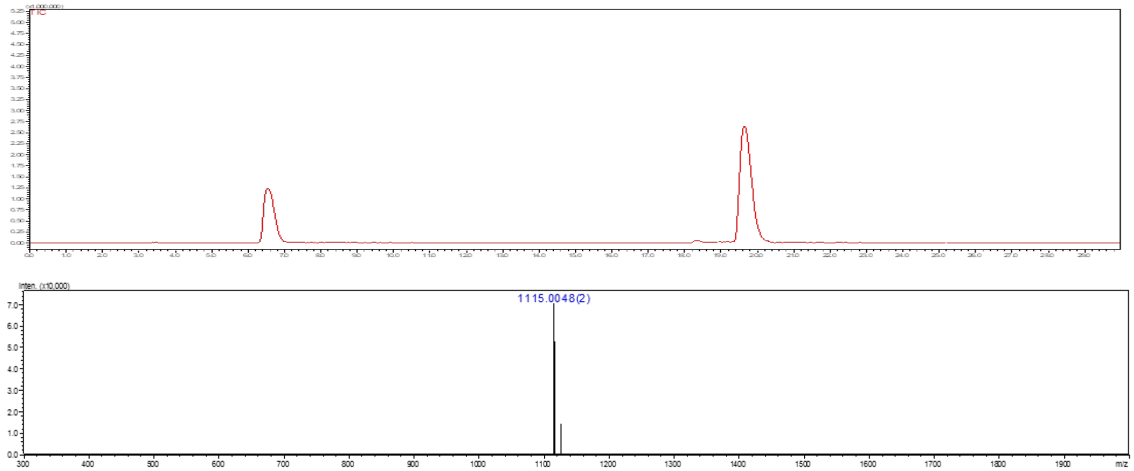

**Figure S35.** Liquid chromatography (LC) trace and mass (ESI-MS) spectrum (peak at retention time = 19.3 min) of **H**.

**Compound 34**

**34** was synthesized from **32** (3 mg, 1.40 µmol) following the general protocol **3g** for the installation of Neu5Ac moiety using ST6Gal1. The product **34** was obtained as a white fluffy solid (3.0 mg, 88%).

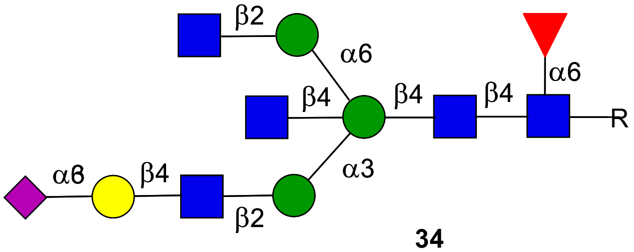

**34**

**$^1$H NMR (600 MHz, $D_2O$): δ (ppm)**

| | H1 | H2 | H3 | H4 | H5 | H6 | NHAc |
|---|---|---|---|---|---|---|---|
| GlcNAc**1** | 4.85 | 3.69 | 3.60 | 3.43 | N/R[a] | N/R | 2.02-1.79 (15H) |
| GlcNAc**2** | 4.54 | 3.73 | 3.62 | N/R | 3.48 | N/R | 2.02-1.79 (15H) |
| GlcNAc**3** | 4.54 | 3.69 | 3.49 | N/R | N/R | N/R | 2.02-1.79 (15H) |
| GlcNAc**4** | 4.39 | 3.61 | 3.34 | 3.19 | 3.49 | N/R | 2.02-1.79 (15H) |
| GlcNAc**5** | 4.47 | 3.63 | 3.41 | 3.34 | N/R | N/R | 2.02-1.79 (15H) |
| Man**1** | 4.62 | 4.10 | 3.80 | 3.99 | 3.48 | N/R | **-** |
| Man**2** | 5.00 | 4.19 | 3.83 | 3.63 | N/R | N/R | **-** |
| Man**3** | 4.92 | 4.08 | 3.77 | 3.54 | 3.40 | N/R | **-** |
| Fuc**1** | 4.73 | 3.66 | 3.71 | 3.60 | 3.95 | 1.06 | **-** |
| Gal**1** | 4.37 | 3.45 | N/R | 3.86 | N/R | N/R | **-** |

| | H1 | H2 | H3 | H4 | H5 | H6 | H7 | H8 | H9 |
|---|---|---|---|---|---|---|---|---|---|
| Neu5Ac9 | **-** | **-** | 2.59<br>1.65 | 3.58 | 3.73 | N/R | N/R | N/R | N/R |

| Signal | Proton | Carbon |
|---|---|---|
| **Aromatic** | 7.92-7.85 (m, 4H)<br>7.52-7.48 (m, 3H) | 128.4-125.7 |
| **$CH_2$**Ph | 5.33 (d, *J* = 12.8 Hz)<br>5.14 (d, *J* = 12.7 Hz) | 66.9 |
| NH-**CH**-COOH | 4.29 (dd, *J* = 9.5, 3.6 Hz) | 53.1 |
| C(O)-**$CH_2$**-CH | 2.75 (dd, *J* = 14.8, 2.9 Hz)<br>2.51 (dd, *J* = 15.0, 9.9 Hz) | 38.6 |

**$^{13}$C NMR (150 MHz, $D_2O$): δ (ppm)**

| | **C1** |
|---|---|
| GlcNAc**1** | 78.2 |
| GlcNAc**2** | 100.9 |
| GlcNAc**3** | 99.4 |
| GlcNAc**4** | 99.4 |
| GlcNAc**5** | 100.6 |
| Man**1** | 100.1 |
| Man**2** | 99.8 |
| Man**3** | 97.7 |
| Fuc**1** | 99.2 |
| Gal**1** | 103.4 |

[a] Not reported

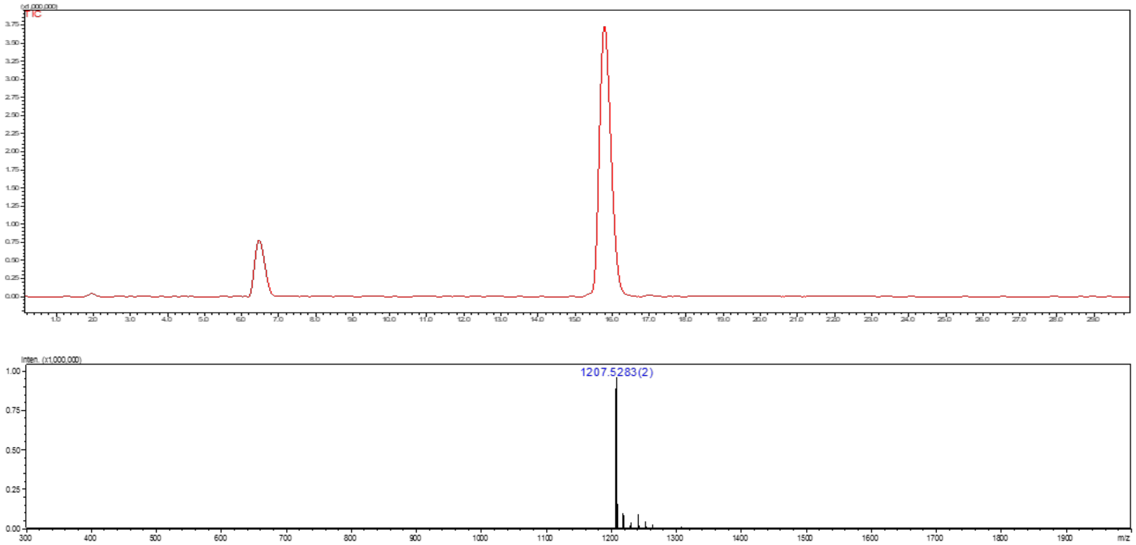

**Figure S36.** Liquid chromatography (LC) trace and mass (ESI-MS) spectrum (peak at retention time = 15.9 min) of **34**.

**Compound I**

The amine of *N*-glycan compound **34** was subjected for the hydrogenation using a general protocol **3k** to obtain **I** as a white solid.

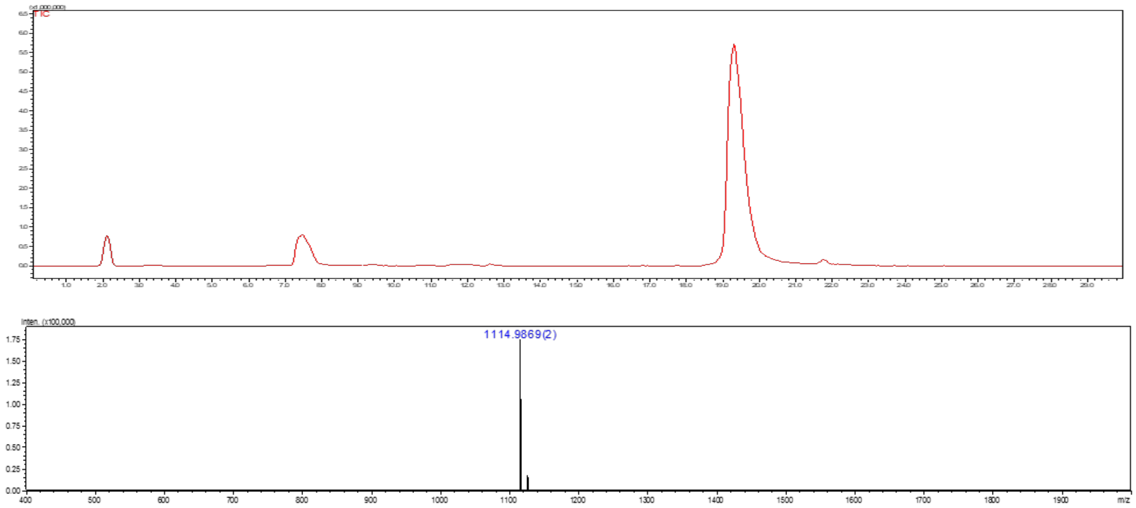

**Figure S37.** Liquid chromatography (LC) trace and mass (ESI-MS) spectrum (peak at retention time = 19.4 min) of **I**.

**Compound 35**

**35** was synthesized from **33** (4 mg, 1.65 μmol) following the general protocol **3h** for the installation of Fuc moiety using FUT6. The product **35** was obtained as a white fluffy solid (3.5 mg, 84%).

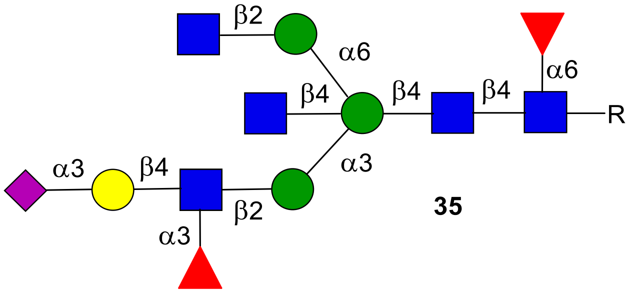

**35**

**$^{1}$H NMR (600 MHz, $D_2O$): δ (ppm)**

| | **H1** | **H2** | **H3** | **H4** | **H5** | **H6** | **NHAc** |
|---|---|---|---|---|---|---|---|
| GlcNAc**1** | 4.86 | 3.70 | 3.60 | 3.45 | N/R[a] | N/R | 2.02-1.79 (15H) |
| GlcNAc**2** | 4.55 | 3.72 | 3.65 | 3.65 | 3.49 | N/R | 2.02-1.79 (15H) |
| GlcNAc**3** | 4.51 | 3.87 | 3.52 | N/R | N/R | N/R | 2.02-1.79 (15H) |
| GlcNAc**4** | 4.38 | 3.63 | 3.36 | 3.18 | 3.19 | N/R | 2.02-1.79 (15H) |
| GlcNAc**5** | 4.47 | 3.64 | 3.41 | 3.40 | N/R | N/R | 2.02-1.79 (15H) |
| Man**1** | 4.61 | 4.10 | 3.79 | 3.97 | 3.48 | N/R | - |
| Man**2** | 4.96 | 4.17 | 3.82 | 3.62 | 3.38 | N/R | - |
| Man**3** | 4.92 | 4.08 | 3.78 | 3.55 | 3.41 | N/R | - |
| Fuc**1** | 4.73 | 3.66 | 3.72 | 3.62 | 3.94 | 1.09 | - |
| Gal**1** | 4.45 | 3.46 | 4.03 | 3.88 | N/R | N/R | - |
| Fuc**2** | 5.06 | 3.61 | 3.89 | 3.69 | 4.75 | 1.09 | - |

| | **H1** | **H2** | **H3** | **H4** | **H5** | **H6** | **H7** | **H8** | **H9** |
|---|---|---|---|---|---|---|---|---|---|
| Neu5Ac9 | - | - | 2.68<br>1.71 | 3.60 | 3.77 | N/R | N/R | N/R | N/R |

| **Signal** | **Proton** | **Carbon** |
|---|---|---|
| **Aromatic** | 7.92-7.85 (m, 4H)<br>7.52-7.47 (m, 3H) | 128.4-125.7 |
| **$CH_2$**Ph | 5.32 (d, *J* = 12.8 Hz)<br>5.13 (d, *J* = 12.7 Hz) | 66.8 |
| NH-**CH**-COOH | 4.29 (dd, *J* = 9.5, 3.6 Hz) | 53.0 |
| C(O)-**$CH_2$**-CH | 2.75 (dd, *J* = 14.8, 2.9 Hz)<br>2.51 (dd, *J* = 15.0, 9.9 Hz) | 38.7 |

**$^{13}C$ NMR (150 MHz, $D_2O$): δ (ppm)**

| | **C1** |
|---|---|
| GlcNAc**1** | 78.2 |
| GlcNAc**2** | 100.9 |
| GlcNAc**3** | 99.4 |
| GlcNAc**4** | 100.7 |
| GlcNAc**5** | 99.5 |
| Man**1** | 100.2 |
| Man**2** | 99.7 |
| Man**3** | 97.7 |
| Fuc**1** | 99.2 |
| Gal**1** | 101.7 |
| Fuc**2** | 98.5 |

[a] Not reported

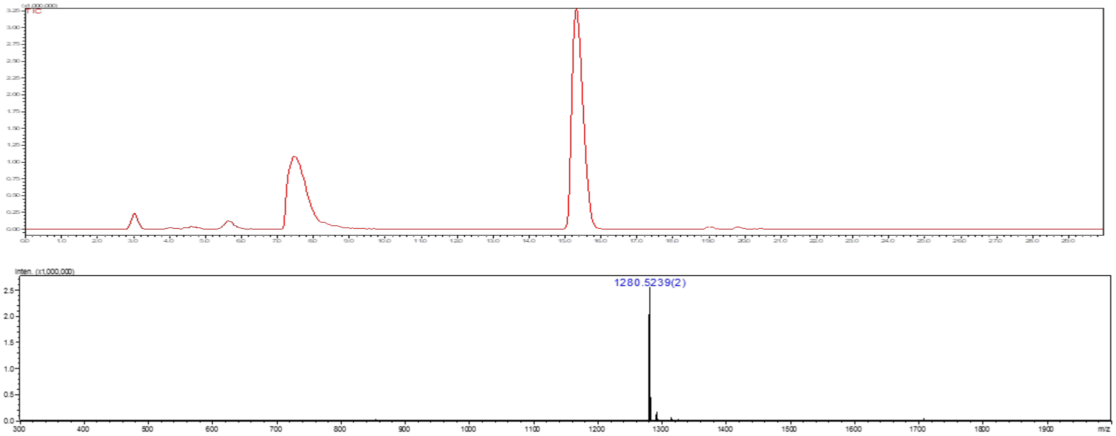

**Figure S38.** Liquid chromatography (LC) trace and mass (ESI-MS) spectrum (peak at retention time = 15.5 min) of **35**.

**Compound K**

The amine of *N*-glycan compound **35** was subjected for the hydrogenation using a general protocol **3k** to obtain **K** as a white solid.

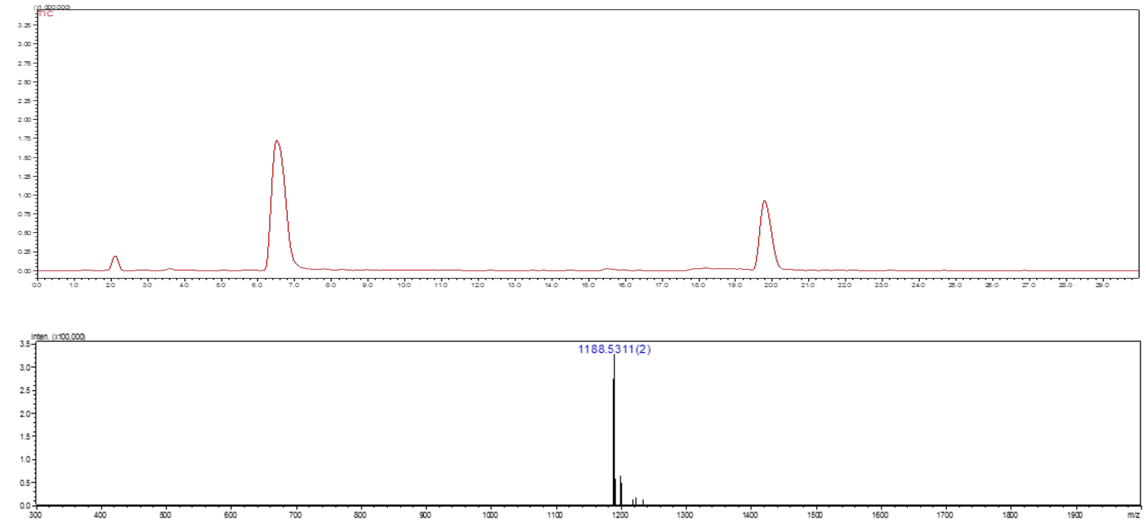

**Figure S39.** Liquid chromatography (LC) trace and mass (ESI-MS) spectrum (peak at retention time = 19.8 min) of **K**.

**Compound 39**

**36** was synthesized from **33** (3 mg, 1.24 µmol) following the general protocol **3f** for the installation of Gal moiety using B4GalT1. The product **36** was obtained as a white fluffy solid (2.30 mg, 71%).

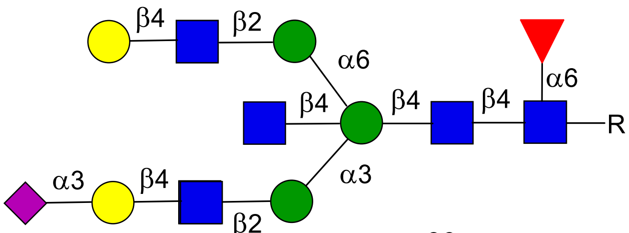

**36**

**$^1$H NMR (900 MHz, $D_2O$): δ (ppm)**

| | **H1** | **H2** | **H3** | **H4** | **H5** | **H6** | **NHAc** |
|---|---|---|---|---|---|---|---|
| GlcNAc**1** | 4.87 | 3.70 | 3.60 | N/R[a] | N/R | N/R | 2.02-1.79 (15H) |
| GlcNAc**2** | 4.55 | 3.73 | 3.62 | 3.66 | 3.48 | N/R | 2.02-1.79 (15H) |
| GlcNAc**3** | 4.52 | 3.70 | 3.53 | N/R | N/R | N/R | 2.02-1.79 (15H) |
| GlcNAc**4** | 4.38 | 3.62 | 3.33 | 3.19 | 3.49 | N/R | 2.02-1.79 (15H) |
| GlcNAc**5** | 4.51 | 3.68 | 3.56 | 3.46 | N/R | N/R | 2.02-1.79 (15H) |
| Man**1** | 4.61 | 4.10 | 3.80 | 4.01 | 3.46 | N/R | **-** |
| Man**2** | 4.99 | 4.18 | 3.83 | 3.63 | N/R | N/R | **-** |
| Man**3** | 4.94 | 4.07 | 3.78 | 3.55 | 3.40 | N/R | **-** |
| Fuc**1** | 4.74 | 3.67 | 3.72 | 3.61 | 3.95 | 1.07 | **-** |
| Gal**1** | 4.47 | 3.49 | 4.03 | 3.88 | N/R | N/R | **-** |
| Gal**2** | 4.39 | 3.48 | N/R | 3.87 | N/R | N/R | **-** |

| | **H1** | **H2** | **H3** | **H4** | **H5** | **H6** | **H7** | **H8** | **H9** |
|---|---|---|---|---|---|---|---|---|---|
| Neu5Ac9 | **-** | **-** | 2.68<br>1.72 | 3.60 | 3.77 | N/R | N/R | N/R | N/R |

| **Signal** | **Proton** | **Carbon** |
|---|---|---|
| **Aromatic** | 7.92-7.85 (m, 4H)<br>7.52-7.48 (m, 3H) | 128.4-125.7 |
| **$CH_2$**Ph | 5.32 (d, *J* = 12.8 Hz)<br>5.14 (d, *J* = 12.7 Hz) | 66.7 |
| NH-**CH**-COOH | 4.32 (dd, *J* = 9.5, 3.6 Hz) | 53.0 |
| C(O)-**$CH_2$**-CH | 2.75 (dd, *J* = 14.8, 2.9 Hz)<br>2.54 (dd, *J* = 15.0, 9.9 Hz) | 38.6 |

**$^{13}C$ NMR (225 MHz, $D_2O$): δ (ppm)**

| | **C1** |
|---|---|
| GlcNAc**1** | 78.2 |
| GlcNAc**2** | 100.9 |
| GlcNAc**3** | 99.7 |
| GlcNAc**4** | 100.6 |
| GlcNAc**5** | 99.4 |
| Man**1** | 100.0 |
| Man**2** | 99.9 |
| Man**3** | 97.6 |
| Fuc**1** | 99.2 |
| Gal**1** | 102.9 |
| Gal**2** | 102.6 |

[a] Not reported

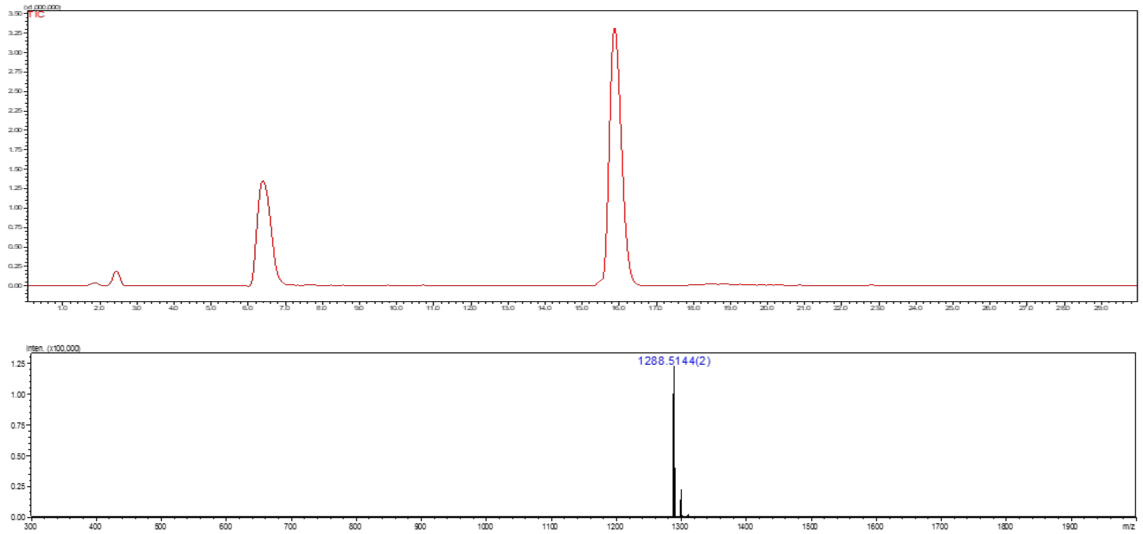

**Figure S40.** Liquid chromatography (LC) trace and mass (ESI-MS) spectrum (peak at retention time = 15.9 min) of **36**.

**Compound J**

The amine of *N*-glycan compound **36** was subjected for the hydrogenation using a general protocol **3k** to obtain **J** as a white solid.

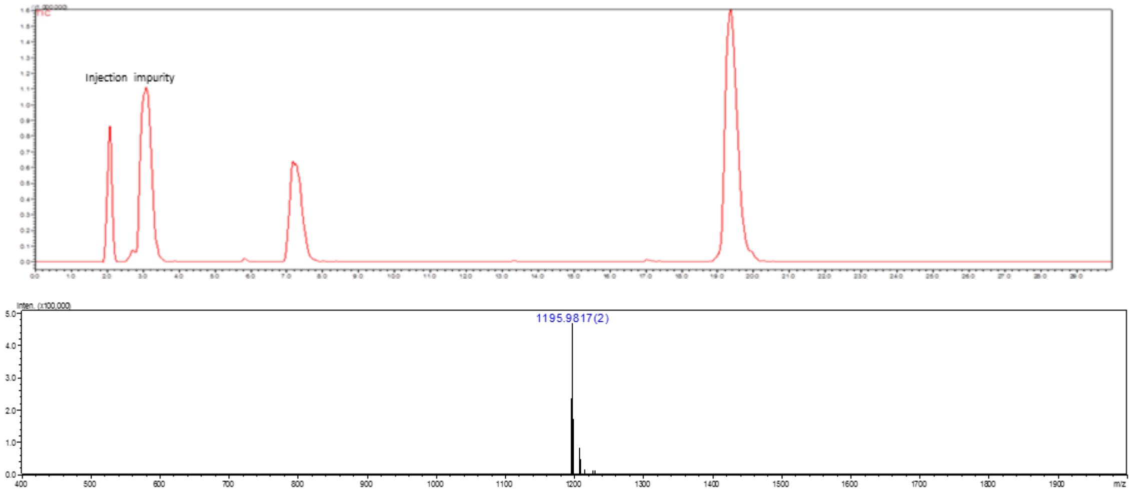

**Figure S41.** Liquid chromatography (LC) trace and mass (ESI-MS) spectrum (peak at retention time = 19.5 min) of **J**.

**Compound 5**

**5** was synthesized from **35** (3 mg, 1.16 µmol) following the general protocol **3f** for the installation of Gal moiety using B4GalT1. The product **5** was obtained as a white fluffy solid (2.1 mg, 66%).

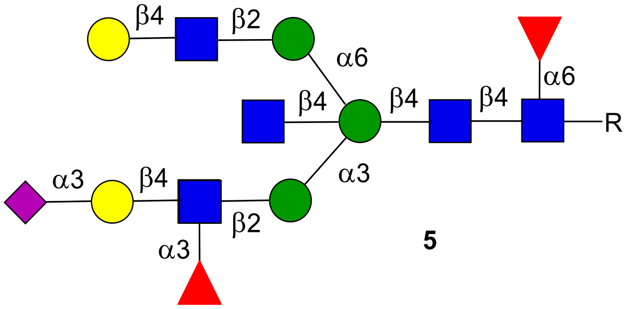

**5**

**$^{1}$H NMR (600 MHz, $D_2O$): δ (ppm)**

| | **H1** | **H2** | **H3** | **H4** | **H5** | **H6** | **NHAc** |
|---|---|---|---|---|---|---|---|
| GlcNAc**1** | 4.86 | 3.70 | 3.60 | N/R[a] | N/R | N/R | 2.02-1.80 (15H) |
| GlcNAc**2** | 4.54 | 3.73 | 3.62 | 3.66 | 3.48 | N/R | 2.02-1.80 (15H) |
| GlcNAc**3** | 4.51 | 3.86 | 3.54 | N/R | N/R | N/R | 2.02-1.80 (15H) |
| GlcNAc**4** | 4.38 | 3.62 | 3.34 | 3.18 | 3.50 | N/R | 2.02-1.80 (15H) |
| GlcNAc**5** | 4.50 | 3.67 | 3.57 | 3.46 | N/R | N/R | 2.02-1.80 (15H) |
| Man**1** | 4.61 | 4.10 | 3.80 | 3.97 | 3.47 | N/R | **-** |
| Man**2** | 4.96 | 4.18 | 3.83 | 3.63 | N/R | N/R | **-** |
| Man**3** | 4.94 | 4.06 | 3.77 | 3.55 | 3.41 | N/R | **-** |
| Fuc**1** | 4.73 | 3.68 | 3.72 | 3.60 | 3.94 | 1.06 | **-** |
| Fuc**2** | 5.06 | 3.61 | N/R | 3.67 | 4.74 | 1.09 | **-** |
| Gal**1** | 4.39 | 3.47 | N/R | 3.85 | N/R | N/R | **-** |
| Gal**2** | 4.45 | 3.46 | 4.01 | 3.87 | N/R | N/R | **-** |

| | **H1** | **H2** | **H3** | **H4** | **H5** | **H6** | **H7** | **H8** | **H9** |
|---|---|---|---|---|---|---|---|---|---|
| Neu5Ac9 | **-** | **-** | 2.67<br>1.71 | 3.60 | 3.77 | N/R | N/R | N/R | N/R |

| **Signal** | **Proton** | **Carbon** |
|---|---|---|
| **Aromatic** | 7.91-7.85 (m, 4H)<br>7.53-7.48 (m, 3H) | 128.3-125.6 |
| **CH$_2$**Ph | 5.35 (d, $J$ = 12.8 Hz)<br>5.15 (d, $J$ = 12.7 Hz) | 66.8 |
| NH-**CH**-COOH | 4.35 (dd, $J$ = 9.5, 3.6 Hz) | 52.6 |
| C(O)-**CH$_2$**-CH | 2.76 (dd, $J$ = 14.8, 2.9 Hz)<br>2.55 (dd, $J$ = 15.0, 9.9 Hz) | 38.4 |

**$^{13}$C NMR (150 MHz, $D_2O$): δ (ppm)**

| | **C1** |
|---|---|
| GlcNAc**1** | 78.2 |
| GlcNAc**2** | 100.8 |
| GlcNAc**3** | 99.4 |
| GlcNAc**4** | 100.6 |
| GlcNAc**5** | 99.4 |
| Man**1** | 100.0 |
| Man**2** | 99.8 |
| Man**3** | 97.6 |
| Fuc**1** | 99.1 |
| Gal**1** | 102.9 |
| Gal**2** | 101.7 |
| Fuc**2** | 98.5 |

[a] Not reported

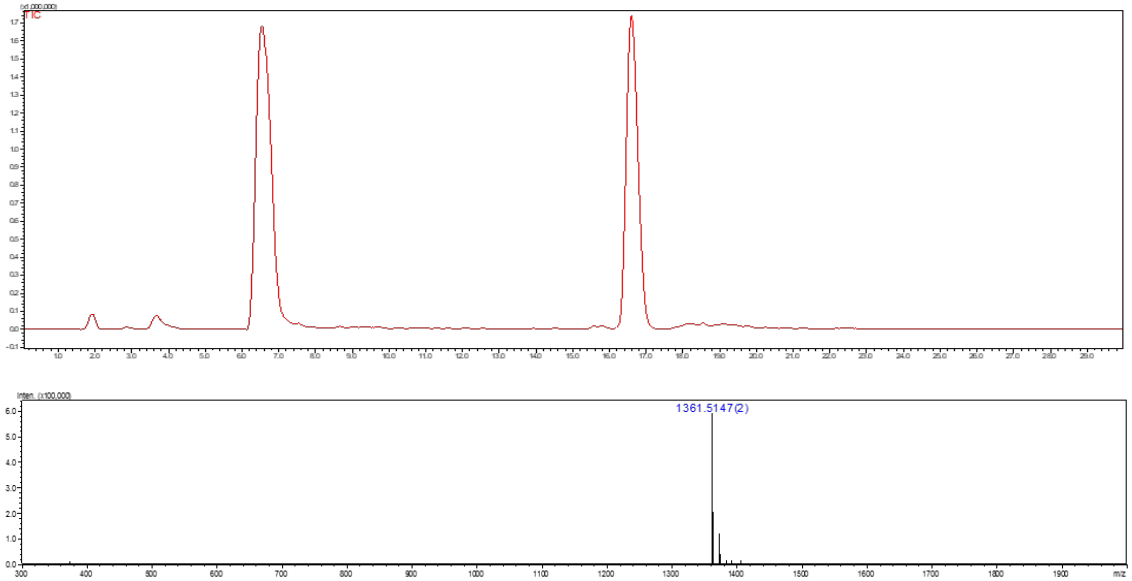

**Figure S42.** Liquid chromatography (LC) trace and mass (ESI-MS) spectrum (peak at retention time = 16.6 min) of **5**.

**Compound L**

The amine of *N*-glycan compound **5** was subjected for the hydrogenation using a general protocol **3k** to obtain **L** as a white solid.

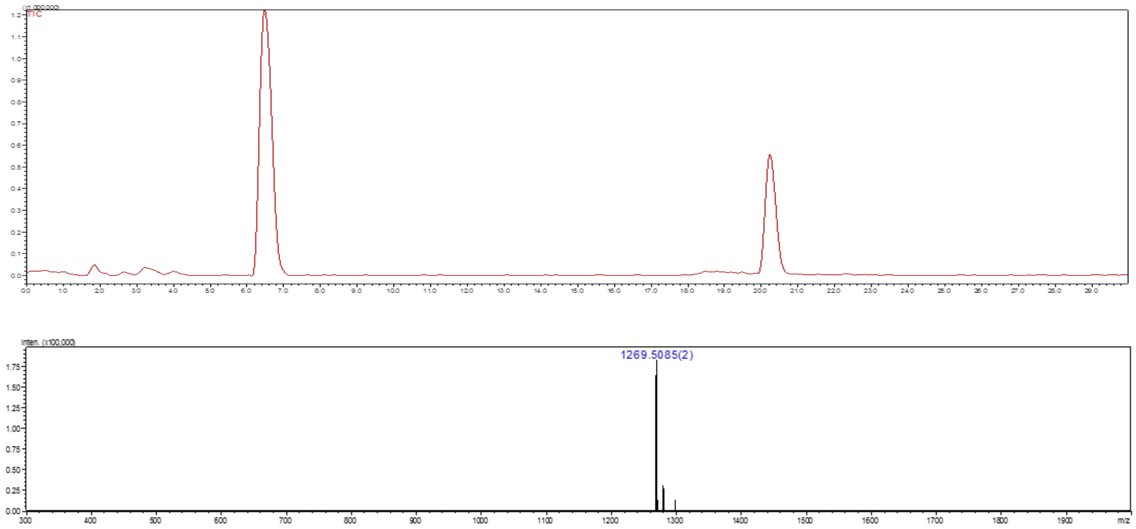

**Figure S43.** Liquid chromatography (LC) trace and mass (ESI-MS) spectrum (peak at retention time = 20.3 min) of **L.**

**Compound 38**

*N*-Glycan **22** (17 mg, 7.99 µmol) was dissolved in deionized water with the final reaction concentration of 5 mM. The pH of the reaction solution was adjusted to 10 using NaOH (1 M), and the mixture solution was incubated for 3-5 h at 37 °C with shaking. After complete conversion of starting material to amine intermediate **8** [confirmed by LC-ESI-MS]. Then, reaction was neutralized by aqueous acetic acid (1 M). The product was purified by C18 biogel [Eluent - water and water:acetonitrile (8:2)]. Next, compound **8** was subjected for the installation of a β1,4 Gal using the general procedure **3f** to obtained **37** as a white solid which was subjected for the acetylation using a general protocol **3j** to obtain **38** as a white solid (9.5 mg, 49% over 3 steps).

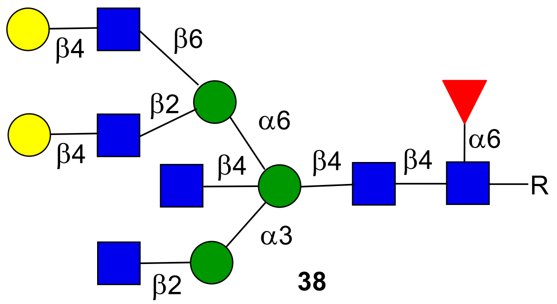

**38**

**[1]H NMR (900 MHz, $D_2O$): δ (ppm)**

| | H1 | H2 | H3 | H4 | H5 | H6 | NHAc |
|---|---|---|---|---|---|---|---|
| GlcNAc**1** | 4.86 | 3.69 | 3.45 | 3.35 | N/R[a] | N/R | 2.02-1.79 (18H) |
| GlcNAc**2** | 4.53 | 3.71 | 3.43 | N/R | N/R | N/R | 2.02-1.79 (18H) |
| GlcNAc**3** | 4.54 | 3.64 | 3.56 | 3.44 | N/R | N/R | 2.02-1.79 (18H) |
| GlcNAc**4** | 4.42 | 3.62 | 3.30 | 3.20 | 3.53 | 3.40 | 2.02-1.79 (18H) |
| GlcNAc**5** | 4.51 | 3.64 | 3.54 | 3.42 | N/R | N/R | 2.02-1.79 (18H) |
| GlcNAc**6** | 4.46 | 3.71 | 3.65 | 3.54 | N/R | N/R | 2.02-1.79 (18H) |
| Man**1** | 4.62 | 4.09 | 3.80 | 3.43 | N/R | N/R | - |
| Man**2** | 4.99 | 4.16 | 3.82 | 3.40 | 3.52 | N/R | - |
| Man**3** | 4.83 | 4.04 | 3.72 | 3.34 | 3.61 | N/R | - |
| Fuc**1** | 4.74 | 3.65 | 3.71 | 3.60 | 3.93 | 1.05 | - |
| Gal**1** | 4.40 | 3.47 | 3.60 | 3.85 | N/R | N/R | - |
| Gal**2** | 4.41 | 3.48 | 3.61 | 3.85 | N/R | N/R | - |

| Signal | Proton | Carbon |
|---|---|---|
| **Aromatic** | 7.92-7.84 (m, 4H)<br>7.52-7.47 (m, 3H) | 128.4-125.7 |
| **$CH_2$**Ph | 5.32 (d, $J$ = 12.8 Hz)<br>5.14 (d, $J$ = 12.7 Hz) | 66.7 |
| NH-**CH**-COOH | 4.29 (dd, $J$ = 9.5, 3.6 Hz) | 53.0 |
| C(O)-**$CH_2$**-CH | 2.74 (dd, $J$ = 14.8, 2.9 Hz)<br>2.51 (dd, $J$ = 15.0, 9.9 Hz) | 38.6 |

**$^{13}$C NMR (225 MHz, $D_2O$): δ (ppm)**

| | **C1** |
|---|---|
| GlcNAc**1** | 78.1 |
| GlcNAc**2** | 101.0 |
| GlcNAc**3** | 99.6 |
| GlcNAc**4** | 99.8 |
| GlcNAc**5** | 99.9 |
| GlcNAc**6** | 101.6 |
| Man**1** | 99.8 |
| Man**2** | 99.9 |
| Man**3** | 97.8 |
| Fuc**1** | 99.2 |
| Gal**1** | 103.0 |
| Gal**2** | 103.0 |

[a] Not reported

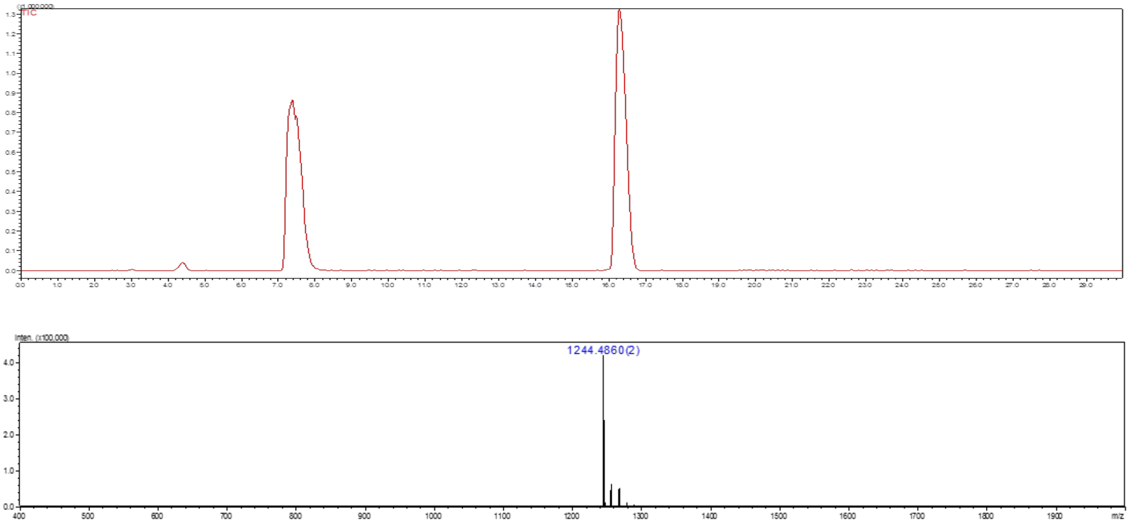

**Figure S44.** Liquid chromatography (LC) trace and mass (ESI-MS) spectrum (peak at retention time = 16.4 min) of **38**.

#### Compound M

The amine of *N*-glycan compound **38** was subjected for the hydrogenation using a general protocol **3k** to obtain **M** as a white solid.

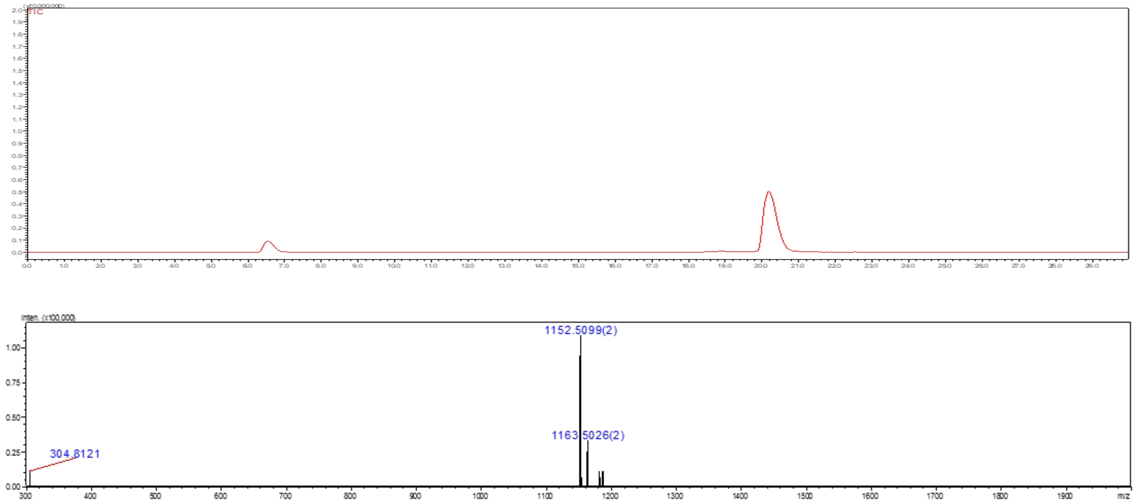

**Figure S45.** Liquid chromatography (LC) trace and mass (ESI-MS) spectrum (peak at retention time = 20.6 min) of **M**.

#### Compound 39

**39** was synthesized from **38** (6.8 mg, 2.74 μmol) following the general protocol **3g** for the installation of Neu5Ac moiety using ST3Gal4. The product **39** was obtained as a white fluffy solid (7.7 mg, 92%).

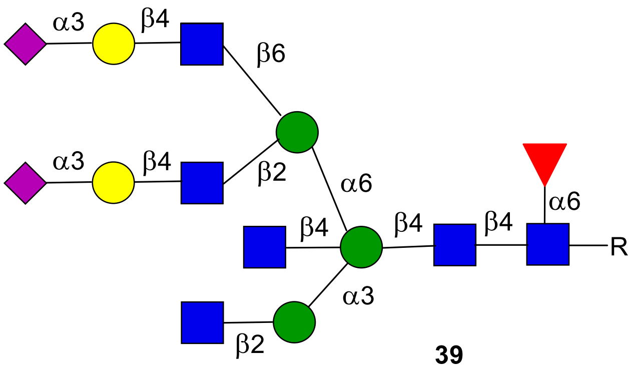

**39**

**$^1$H NMR (600 MHz, $D_2O$): δ (ppm)**

| | H1 | H2 | H3 | H4 | H5 | H6 | NHAc |
|---|---|---|---|---|---|---|---|
| GlcNAc**1** | 4.87 | 3.71 | 3.48 | 3.41 | N/R[a] | N/R | 2.02-1.79 (18H) |
| GlcNAc**2** | 4.52 | 3.73 | 3.49 | N/R | N/R | N/R | 2.02-1.79 (18H) |
| GlcNAc**3** | 4.52 | 3.66 | 3.55 | 3.43 | N/R | N/R | 2.02-1.79 (18H) |
| GlcNAc**4** | 4.42 | 3.63 | 3.30 | 3.20 | 3.54 | N/R | 2.02-1.79 (18H) |
| GlcNAc**5** | 4.52 | 3.66 | 3.55 | 3.43 | N/R | N/R | 2.02-1.79 (18H) |
| GlcNAc**6** | 4.46 | 3.73 | 3.66 | 3.54 | N/R | N/R | 2.02-1.79 (18H) |
| Man**1** | 4.63 | 4.10 | 3.79 | 3.43 | N/R | N/R | - |
| Man**2** | 4.99 | 4.17 | 3.83 | 3.39 | 3.62 | N/R | - |
| Man**3** | 4.83 | 4.05 | 3.72 | 3.34 | N/R | N/R | - |
| Fuc**1** | 4.75 | 3.67 | 3.71 | 3.60 | 3.94 | 1.06 | - |
| Gal**1** | 4.47 | 3.50 | 4.05 | 3.89 | N/R | N/R | - |
| Gal**2** | 4.48 | 3.51 | 4.05 | 3.89 | N/R | N/R | - |

| | H1 | H2 | H3 | H4 | H5 | H6 | H7 | H8 | H9 |
|---|---|---|---|---|---|---|---|---|---|
| Neu5Ac9-**1** | - | - | 2.68<br>1.71 | 3.61 | 3.77 | N/R | N/R | N/R | N/R |
| Neu5Ac9-**2** | - | - | 2.68<br>1.71 | 3.61 | 3.77 | N/R | N/R | N/R | N/R |

| Signal | Proton | Carbon |
|---|---|---|
| **Aromatic** | 7.92-7.85 (m, 4H)<br>7.53-7.48 (m, 3H) | 128.4-125.7 |
| **$CH_2$**Ph | 5.33 (d, *J* = 12.8 Hz)<br>5.13 (d, *J* = 12.7 Hz) | 66.9 |
| NH-**CH**-COOH | 4.29 (dd, *J* = 9.5, 3.6 Hz) | 52.4 |
| C(O)-**$CH_2$**-CH | 2.75 (dd, *J* = 14.8, 2.9 Hz)<br>2.51 (dd, *J* = 15.0, 9.9 Hz) | 38.4 |

**$^{13}$C NMR (150 MHz, $D_2O$): δ (ppm)**

| | **C1** |
|---|---|
| GlcNAc**1** | 78.2 |
| GlcNAc**2** | 101.1 |
| GlcNAc**3** | 99.8 |
| GlcNAc**4** | 99.8 |
| GlcNAc**5** | 99.9 |
| GlcNAc**6** | 101.6 |
| Man**1** | 99.7 |
| Man**2** | 99.9 |
| Man**3** | 98.0 |
| Fuc**1** | 99.2 |
| Gal**1** | 102.6 |
| Gal**2** | 102.6 |

[a] Not reported

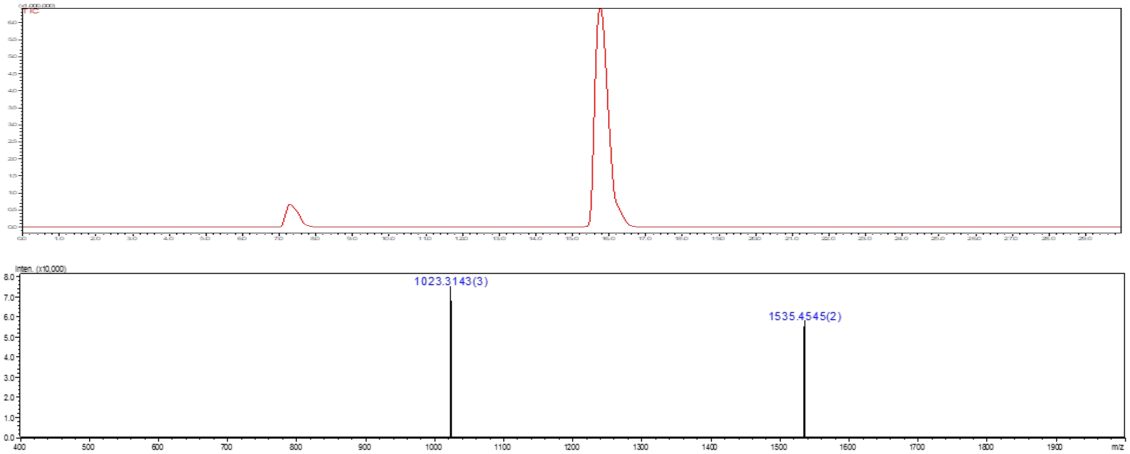

**Figure S46.** Liquid chromatography (LC) trace and mass (ESI-MS) spectrum (peak at retention time = 15.9 min) of **39**.

**Compound N**

The amine of *N*-glycan compound **39** was subjected for the hydrogenation using a general protocol **3k** to obtain **N** as a white solid.

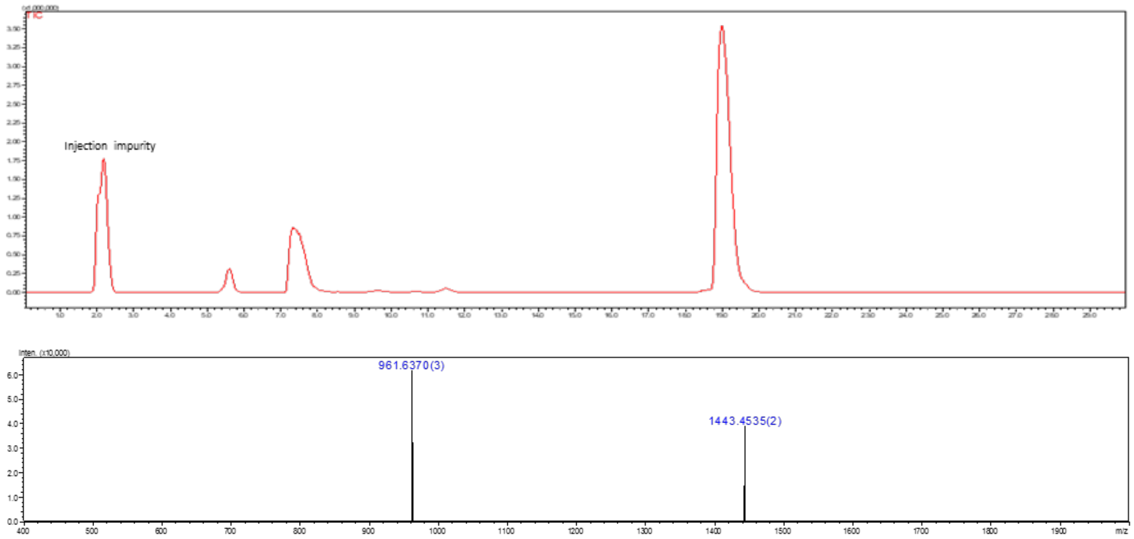

**Figure S47.** Liquid chromatography (LC) trace and mass (ESI-MS) spectrum (peak at retention time = 19.2 min) of **N**.

**Compound 40**

**40** was synthesized from **39** (5.8 mg, 1.88 μmol) following the general protocol **3h** for the installation of Fuc moiety using FUT6. The product **40** was obtained as a white fluffy solid (5.4 mg, 86%).

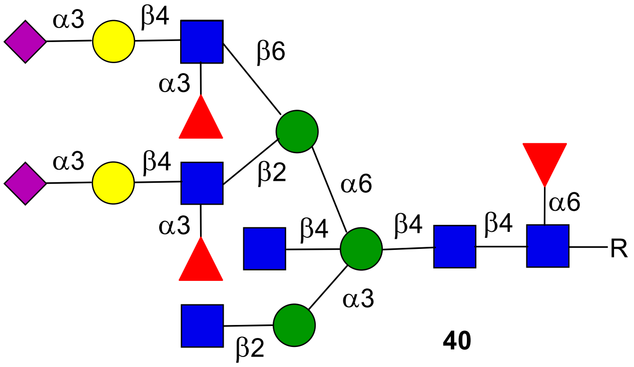

**40**

**$^1$H NMR (900 MHz, $D_2O$): δ (ppm)**

| | H1 | H2 | H3 | H4 | H5 | H6 | NHAc |
|---|---|---|---|---|---|---|---|
| GlcNAc**1** | 4.86 | 3.70 | 3.45 | 3.40 | N/R[a] | N/R | 2.03-1.79 (18H) |
| GlcNAc**2** | 4.53 | 3.72 | 3.49 | N/R | N/R | N/R | 2.03-1.79 (18H) |
| GlcNAc**3** | 4.53 | 3.64 | 3.54 | 3.44 | N/R | N/R | 2.03-1.79 (18H) |
| GlcNAc**4** | 4.46 | 3.63 | 3.34 | 3.22 | 3.51 | N/R | 2.03-1.79 (18H) |
| GlcNAc**5** | 4.53 | 3.64 | 3.54 | 3.41 | N/R | N/R | 2.03-1.79 (18H) |
| GlcNAc**6** | 4.46 | 3.85 | 3.63 | N/R | N/R | N/R | 2.03-1.79 (18H) |
| Man**1** | 4.62 | 4.08 | 3.77 | 3.52 | N/R | N/R | - |
| Man**2** | 4.96 | 4.15 | 3.84 | 3.35 | 3.60 | N/R | - |
| Man**3** | 4.83 | 4.06 | 3.74 | 3.33 | N/R | N/R | - |
| Fuc**1** | 4.73 | 3.66 | 3.71 | 3.61 | 3.94 | 1.06 | - |
| Gal**1** | 4.45 | 3.45 | 4.01 | 3.85 | N/R | N/R | - |
| Gal**2** | 4.45 | 3.45 | 4.01 | 3.85 | N/R | N/R | - |
| Fuc**2** | 5.04 | 3.62 | 3.71 | 3.61 | 4.76 | 1.11 | - |
| Fuc**3** | 5.08 | 3.62 | 3.71 | 3.61 | 4.76 | 1.11 | - |

| | H1 | H2 | H3 | H4 | H5 | H6 | H7 | H8 | H9 |
|---|---|---|---|---|---|---|---|---|---|
| Neu5Ac9-**1** | - | - | 2.68<br>1.72 | 3.60 | 1.73 | N/R | N/R | N/R | N/R |
| Neu5Ac9-**2** | - | - | 2.68<br>1.72 | 3.60 | 1.73 | N/R | N/R | N/R | N/R |

| Signal | Proton | Carbon |
|---|---|---|
| **Aromatic** | 7.91-7.85 (m, 4H)<br>7.53-7.48 (m, 3H) | 128.3-125.7 |
| **$CH_2$**Ph | 5.33 (d, *J* = 12.8 Hz)<br>5.14 (d, *J* = 12.7 Hz) | 66.8 |
| NH-**CH**-COOH | 4.29 (dd, *J* = 9.5, 3.6 Hz) | 53.0 |
| C(O)-**$CH_2$**-CH | 2.76 (dd, *J* = 14.8, 2.9 Hz) | 38.7 |

| | |
|---|---|
| | 2.51 (dd, $J$ = 15.0, 9.9 Hz) |

**$^{13}$C NMR (225 MHz, $D_2O$): δ (ppm)**

| | **C1** |
|---|---|
| GlcNAc**1** | 78.1 |
| GlcNAc**2** | 101.0 |
| GlcNAc**3** | 99.8 |
| GlcNAc**4** | 99.2 |
| GlcNAc**5** | 99.9 |
| GlcNAc**6** | 101.0 |
| Man**1** | 100.1 |
| Man**2** | 99.9 |
| Man**3** | 97.7 |
| Fuc**1** | 99.2 |
| Gal**1** | 101.5 |
| Gal**2** | 101.5 |

[a] Not reported

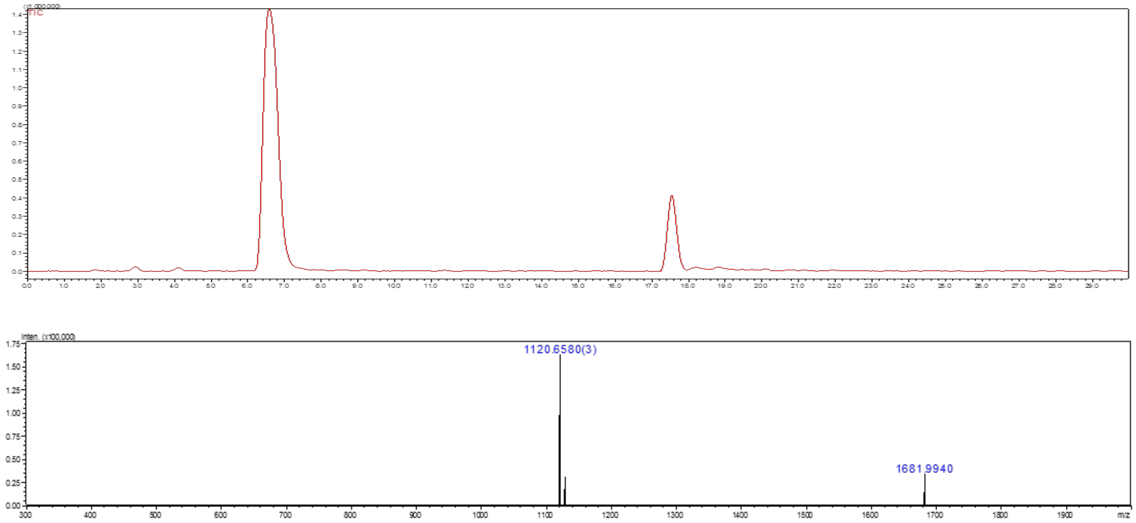

**Figure S48.** Liquid chromatography (LC) trace and mass (ESI-MS) spectrum (peak at retention time = 17.6 min) of **40**.

### Compound P

The amine of *N*-glycan compound **40** was subjected for the hydrogenation using a general protocol **3k** to obtain **P** as a white solid.

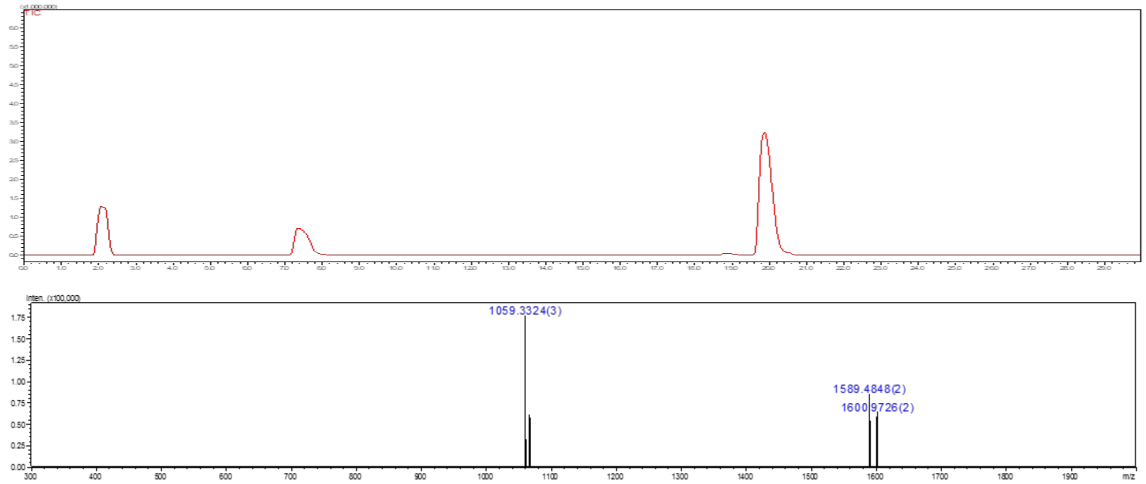

**Figure S49.** Liquid chromatography (LC) trace and mass (ESI-MS) spectrum (peak at retention time = 20.0 min) of **P**.

### Compound 41

**41** was synthesized from **39** (2.5 mg, 0.81 µmol) following the general protocol **3f** for the installation of Gal moiety using B4GalT1. The product **41** was obtained as a white fluffy solid (2.14 mg, 82%).

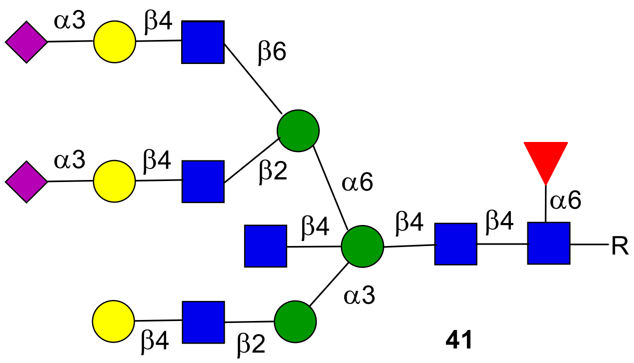

**41**

**$^{1}$H NMR (900 MHz, $D_2O$): δ (ppm)**

| | **H1** | **H2** | **H3** | **H4** | **H5** | **H6** | **NHAc** |
|---|---|---|---|---|---|---|---|
| GlcNAc**1** | 4.85 | 3.70 | 3.46 | 3.39 | N/R[a] | N/R | 2.03-1.79 (18H) |
| GlcNAc**2** | 4.55 | 3.69 | 3.58 | N/R | N/R | N/R | 2.03-1.79 (18H) |
| GlcNAc**3** | 4.52 | 3.71 | 3.54 | 3.43 | N/R | N/R | 2.03-1.79 (18H) |
| GlcNAc**4** | 4.42 | 3.62 | 3.29 | 3.20 | 3.54 | N/R | 2.03-1.79 (18H) |
| GlcNAc**5** | 4.52 | 3.61 | 3.55 | 3.48 | N/R | N/R | 2.03-1.79 (18H) |
| GlcNAc**6** | 4.46 | 3.72 | 3.66 | 3.53 | N/R | N/R | 2.03-1.79 (18H) |
| Man**1** | 4.61 | 4.09 | 3.78 | 3.44 | N/R | N/R | - |
| Man**2** | 4.97 | 4.17 | 3.83 | 3.39 | 3.54 | N/R | - |
| Man**3** | 4.82 | 4.03 | 3.71 | 3.34 | N/R | N/R | - |
| Fuc**1** | 4.74 | 3.66 | 3.71 | 3.61 | 3.93 | 1.06 | - |
| Gal**1** | 4.38 | 3.46 | 3.58 | 3.84 | N/R | N/R | - |
| Gal**2** | 4.47 | 3.50 | 4.05 | 3.88 | N/R | N/R | - |
| Gal**3** | 4.48 | 3.51 | 4.05 | 3.88 | N/R | N/R | - |

| | **H1** | **H2** | **H3** | **H4** | **H5** | **H6** | **H7** | **H8** | **H9** |
|---|---|---|---|---|---|---|---|---|---|
| Neu5Ac9-**1** | - | - | 2.68<br>1.72 | 3.61 | 3.77 | N/R | N/R | N/R | N/R |
| Neu5Ac9-**2** | - | - | 2.68<br>1.72 | 3.61 | 3.77 | N/R | N/R | N/R | N/R |

| **Signal** | **Proton** | **Carbon** |
|---|---|---|
| **Aromatic** | 7.91-7.84 (m, 4H)<br>7.52-7.47 (m, 3H) | 128.3-125.6 |
| **$CH_2$**Ph | 5.33 (d, *J* = 12.8 Hz)<br>5.14 (d, *J* = 12.7 Hz) | 66.8 |
| NH-**CH**-COOH | 4.32 (dd, *J* = 9.5, 3.6 Hz) | 52.8 |
| C(O)-**$CH_2$**-CH | 2.76 (dd, *J* = 14.8, 2.9 Hz)<br>2.53 (dd, *J* = 15.0, 9.9 Hz) | 38.7 |

**$^{13}C$ NMR (225 MHz, $D_2O$): δ (ppm)**

| | **C1** |
|---|---|
| GlcNAc**1** | 78.2 |
| GlcNAc**2** | 101.1 |
| GlcNAc**3** | 99.6 |
| GlcNAc**4** | 99.7 |
| GlcNAc**5** | 99.6 |
| GlcNAc**6** | 101.7 |
| Man**1** | 99.8 |
| Man**2** | 99.8 |
| Man**3** | 97.8 |
| Fuc**1** | 99.2 |
| Gal**1** | 102.8 |
| Gal**2** | 102.6 |
| Gal**3** | 102.6 |

[a] Not reported

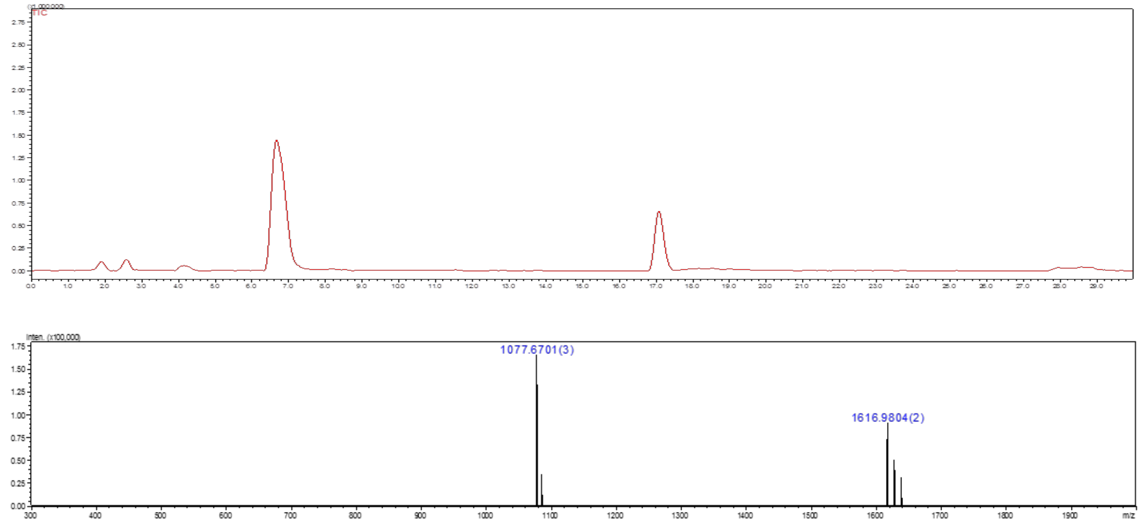

**Figure S50.** Liquid chromatography (LC) trace and mass (ESI-MS) spectrum (peak at retention time = 17.1 min) of **41**.

### Compound O

The amine of *N*-glycan compound **41** was subjected for the hydrogenation using a general protocol **3k** to obtain **O** as a white solid.

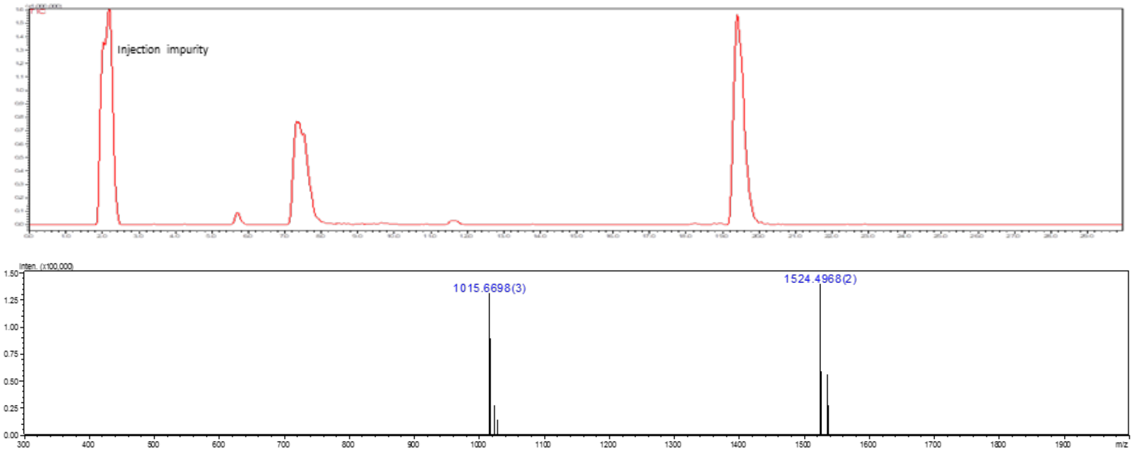

**Figure S51.** Liquid chromatography (LC) trace and mass (ESI-MS) spectrum (peak at retention time = 19.5 min) of **O**.

### Compound 9

**9** was synthesized from **40** (5 mg, 1.48 μmol) following the general protocol **3f** for the installation of Gal moiety using B4GalT1. The product **9** was obtained as a white fluffy solid (3.2 mg, 61%).

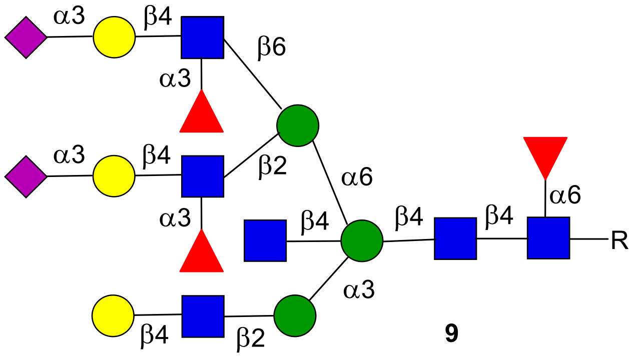

**9**

**$^1$H NMR (600 MHz, $D_2O$) δ (ppm)**

| | H1 | H2 | H3 | H4 | H5 | H6 | NHAc |
|---|---|---|---|---|---|---|---|
| GlcNAc**1** | 4.86 | 3.69 | 3.46 | 3.38 | N/R[a] | N/R | 2.03-1.95 (18H) |
| GlcNAc**2** | 4.53 | 3.72 | 3.48 | N/R | N/R | N/R | 2.03-1.95 (18H) |
| GlcNAc**3** | 4.57 | 3.64 | 3.53 | N/R | N/R | N/R | 2.03-1.95 (18H) |
| GlcNAc**4** | 4.45 | 3.63 | 3.35 | 3.21 | 3.58 | N/R | 2.03-1.95 (18H) |
| GlcNAc**5** | 4.49 | 3.64 | 3.53 | N/R | N/R | N/R | 2.03-1.95 (18H) |
| GlcNAc**6** | 4.46 | 3.85 | 3.63 | N/R | N/R | N/R | 2.03-1.95 (18H) |
| Man**1** | 4.62 | 4.09 | 3.74 | N/R | N/R | N/R | - |
| Man**2** | 4.96 | 4.16 | 3.85 | 3.35 | N/R | N/R | - |
| Man**3** | 4.83 | 4.06 | 3.74 | N/R | N/R | N/R | - |
| Fuc**1** | 4.72 | 3.67 | 3.70 | 3.62 | 3.94 | 1.08 | - |
| Fuc**2** | 5.04 | 3.63 | 3.72 | 3.61 | 4.75 | 1.10 | - |
| Fuc**3** | 5.09 | 3.63 | 3.72 | 3.61 | 4.75 | 1.10 | - |
| Gal**1** | 4.38 | 3.47 | N/R | N/R | N/R | N/R | - |
| Gal**2** | 4.45 | 3.44 | 4.01 | 3.86 | N/R | N/R | - |
| Gal**3** | 4.45 | 3.44 | 4.01 | 3.86 | N/R | N/R | - |

| | H1 | H2 | H3 | H4 | H5 | H6 | H7 | H8 | H9 |
|---|---|---|---|---|---|---|---|---|---|
| Neu5Ac9-**1** | - | - | 2.68<br>1.72 | 3.60 | 3.72 | - | - | - | - |
| Neu5Ac9-**2** | - | - | 2.68<br>1.72 | 3.60 | 3.72 | - | - | - | - |

| **Signal** | **Proton** | **Carbon** |
|---|---|---|
| **Aromatic** | 7.92-7.86 (m, 4H)<br>7.53-7.48 (m, 3H) | 128.4-125.8 |
| **CH$_2$**Ph | 5.33 (d, *J* = 12.8 Hz, 1H)<br>5.14 (d, *J* = 12.8 Hz, 1H) | 66.8 |
| NH-**CH**-COOH | 4.29 (dd, *J* = 9.5, 3.6 Hz 1H) | 52.8 |
| C(O)-**CH**-COOH | 2.75 (dd, *J* = 14.8, 2.9, 1H)<br>2.51 (dd, *J* = 15.0, 9.9, 1H) | 38.7 |

[a] Not reported

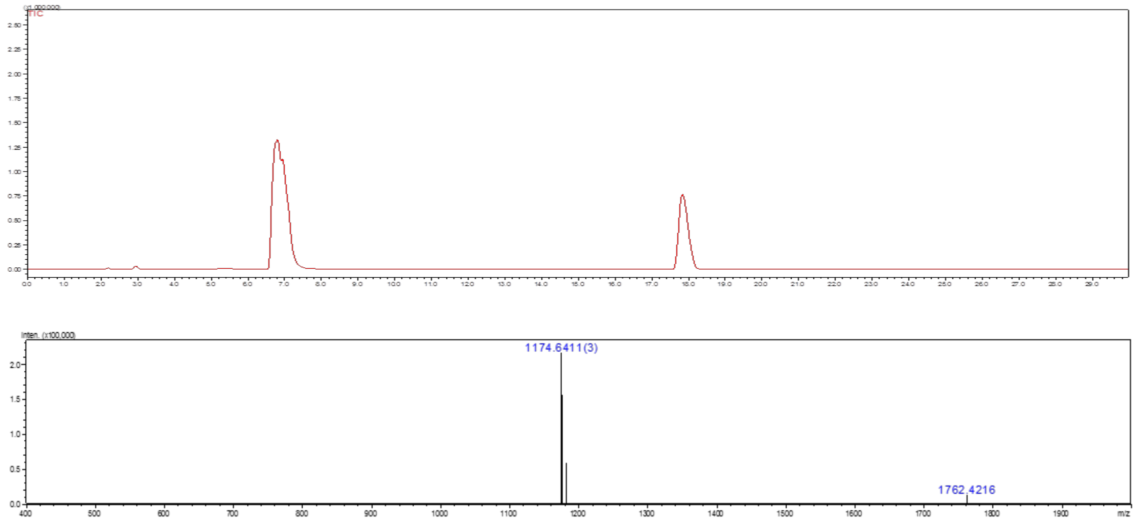

**Figure S52.** Liquid chromatography (LC) trace and mass (ESI-MS) spectrum (peak at retention time = 17.9 min) of **9**.

**Compound Q**

The amine of *N*-glycan compound **9** was subjected for the hydrogenation using a general protocol **3k** to obtain **Q** as a white solid.

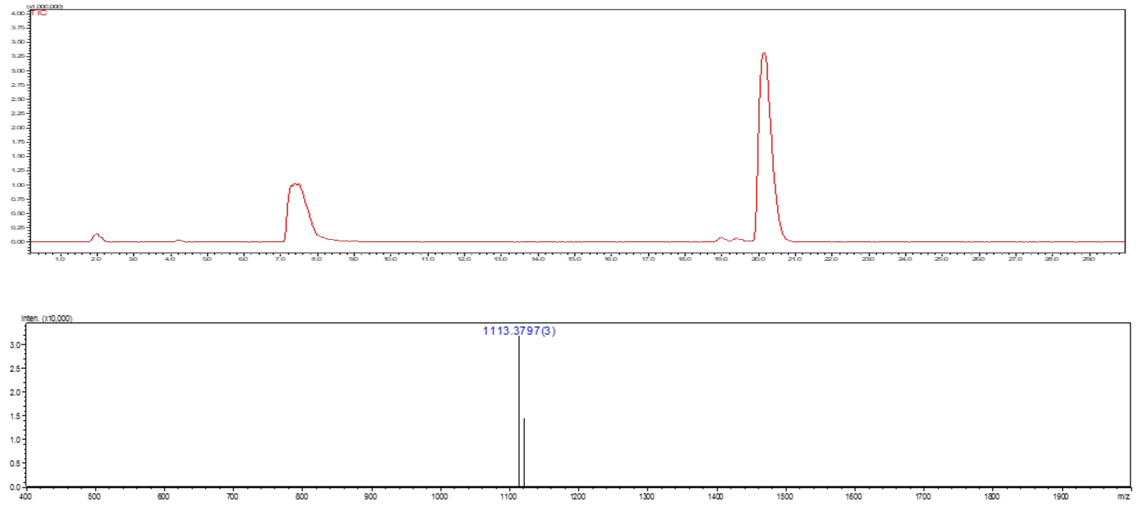

**Figure S53.** Liquid chromatography (LC) trace and mass (ESI-MS) spectrum (peak at retention time = 20.3 min) of **Q**.

**Compound S11**

*N*-Glycan **3** (10 mg, 6.41 µmol) was dissolved in deionized water with the final reaction concentration of 5 mM. The pH of the reaction solution was adjusted to 10 using NaOH (1 M), and the mixture solution was incubated for 3-5 h at 37 °C with shaking. After complete conversion of starting material to amine intermediate reaction was neutralized by aqueous acetic acid (1 M). The product was purified by C18 biogel [Eluent - water and water:acetonitrile (8:2)]. Next, resulting compound was subjected for the installation of a β1,4 Gal using the general procedure **3f** followed by acetylation using a general protocol **3j** to obtain **S9** as a white solid (8.8 mg, 78%).
**S9** (6.1 mg, 3.4 µmol) was subjected to installation of 2, 3-Sialic acid following the general protocol **3g** for the installation of Neu5Ac moiety using ST3Gal4 to get product **S10** as a white fluffy solid. **S11** was synthesized from **S10** following the general protocol **3h** for the installation of Fuc moiety using FUT6. The product **S11** was obtained as a white fluffy solid (5.9 mg, 72%).

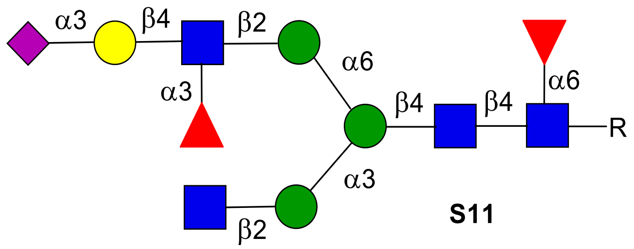

**S11**

**$^1$H NMR (600 MHz, $D_2O$) δ (ppm)**

| | **H1** | **H2** | **H3** | **H4** | **H5** | **H6** | **NHAc** |
|---|---|---|---|---|---|---|---|
| GlcNAc**1** | 4.86 | 3.69 | 3.44 | 3.37 | N/R[a] | N/R | 1.98-1.79 (12H) |
| GlcNAc**2** | 4.54 | 3.70 | 3.51 | 3.65 | N/R | N/R | 1.98-1.79 (12H) |
| GlcNAc**3** | 4.50 | 3.84 | N/R | N/R | N/R | N/R | 1.98-1.79 (12H) |
| GlcNAc**4** | 4.47 | 3.63 | 3.47 | 3.37 | N/R | N/R | 1.98-1.79 (12H) |
| Man**1** | 4.70 | 3.79 | N/R | N/R | N/R | N/R | - |
| Man**2** | 5.03 | 4.11 | 3.82 | 3.42 | N/R | N/R | - |
| Man**3** | 4.83 | 4.02 | 3.86 | 3.41 | N/R | N/R | - |
| Fuc**1** | 4.72 | 3.68 | 3.70 | 3.60 | 3.94 | 1.05 | - |
| Gal**1** | 4.44 | 3.45 | 4.01 | 3.85 | N/R | N/R | - |
| Fuc**2** | 5.05 | 3.60 | 3.83 | 3.71 | 3.74 | 1.10 | - |

| | **H1** | **H2** | **H3** | **H4** | **H5** | **H6** | **H7** | **H8** | **H9** |
|---|---|---|---|---|---|---|---|---|---|
| Neu5Ac9 | - | - | 2.68<br>1.72 | 3.60 | 3.72 | - | - | - | - |

| **Signal** | **Proton** | **Carbon** |
|---|---|---|
| **Aromatic** | 7.91-7.84 (m, 4H)<br>7.52-7.47 (m, 3H) | 128.3-125.6 |
| **$CH_2$**Ph | 5.33 (d, $J$ = 12.8 Hz)<br>5.14 (d, $J$ = 12.7 Hz) | 66.8 |
| NH-**CH**-COOH | 4.32 (dd, $J$ = 9.5, 3.6 Hz) | 52.8 |

| | | |
|---|---|---|
| C(O)-**CH**-COOH | 2.76 (dd, $J$ = 14.8, 2.9 Hz)<br>2.53 (dd, $J$ = 15.0, 9.9 Hz) | 38.7 |

| | **C1** |
|---|---|
| GlcNAc**1** | 78.3 |
| GlcNAc**2** | 100.8 |
| GlcNAc**3** | 99.2 |
| GlcNAc**4** | 99.6 |
| Man**1** | 100.5 |
| Man**2** | 99.5 |
| Man**3** | 96.9 |
| Fuc**1** | 99.2 |
| Gal**1** | 101.4 |
| Fuc**2** | 98.4 |

[a] Not reported

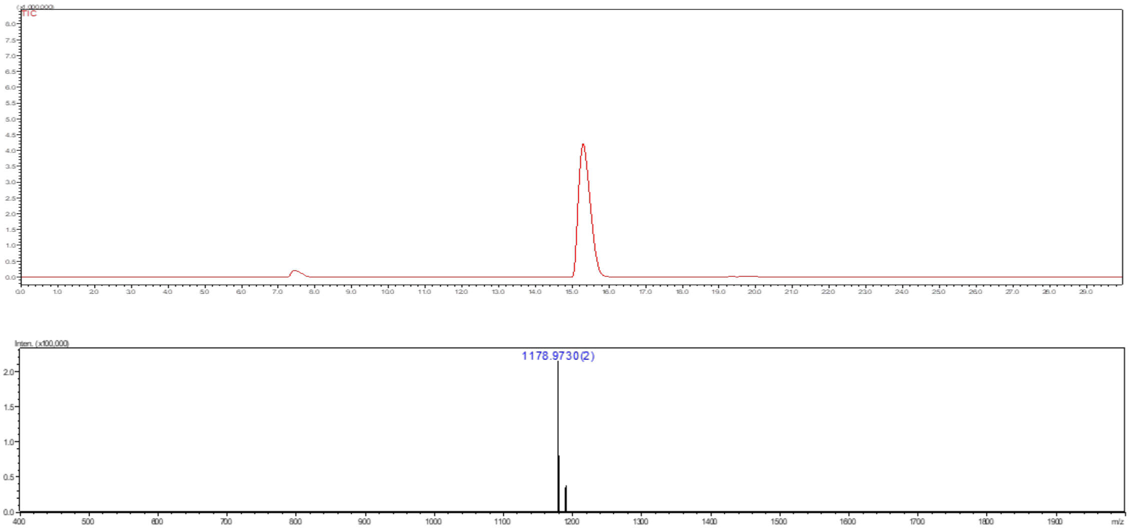

**Figure S54.** Liquid chromatography (LC) trace and mass (ESI-MS) spectrum (peak at retention time = 15.4 min) of **S11**.

**Compound E′**

The amine of *N*-glycan compound **S11** was subjected for the hydrogenation using a general protocol **3k** to obtain **E′** as a white solid.

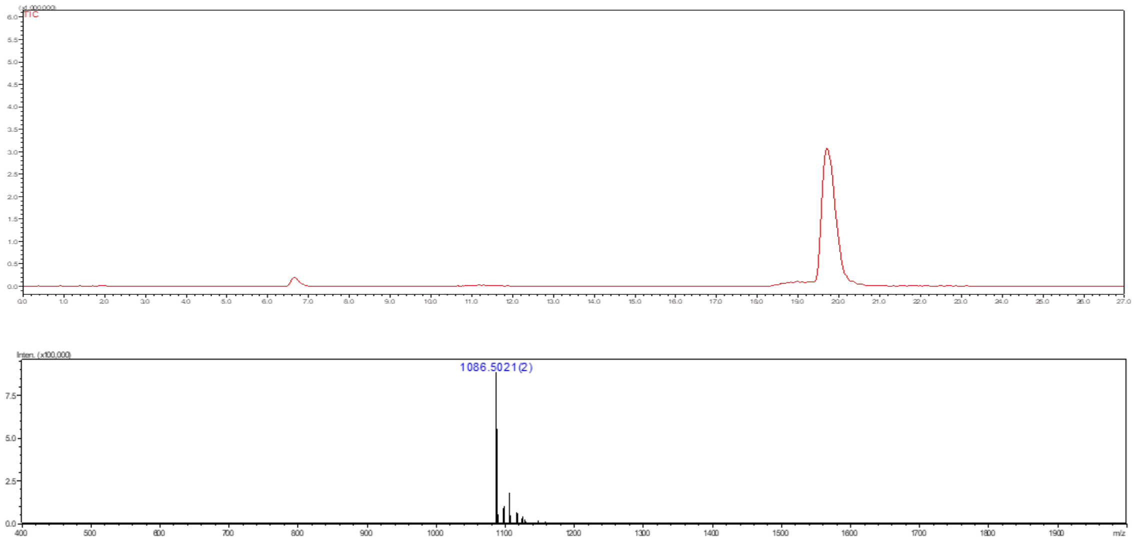

**Figure S55.** Liquid chromatography (LC) trace and mass (ESI-MS) spectrum (peak at retention time = 19.8 min) of **E′**.

**Compound S12**

**S12** was synthesized from **S11** (3 mg, 1.27 μmol) following the general protocol **3f** for the installation of Gal moiety using B4GalT1. The product **S12** was obtained as a white fluffy solid (2.8 mg, 87%).

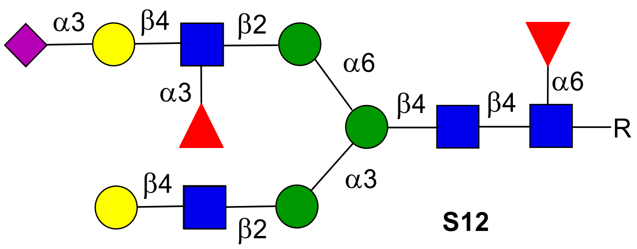

**S12**

**$^{1}$H NMR (600 MHz, $D_2O$) δ (ppm)**

| | **H1** | **H2** | **H3** | **H4** | **H5** | **H6** | **NHAc** |
|---|---|---|---|---|---|---|---|
| GlcNAc**1** | 4.85 | 3.70 | 3.61 | N/R[a] | N/R | N/R | 1.98-1.79 (12H) |
| GlcNAc**2** | 4.54 | 3.70 | 3.65 | N/R | N/R | N/R | 1.98-1.79 (12H) |
| GlcNAc**3** | 4.50 | 3.66 | 3.67 | N/R | N/R | N/R | 1.98-1.79 (12H) |
| GlcNAc**4** | 4.50 | 3.66 | 3.67 | N/R | N/R | N/R | 1.98-1.79 (12H) |
| Man**1** | 4.69 | 4.17 | 3.68 | 3.85 | N/R | N/R | - |
| Man**2** | 5.04 | 4.11 | 3.82 | 3.70 | N/R | N/R | - |
| Man**3** | 4.83 | 4.02 | 3.81 | N/R | N/R | N/R | - |
| Fuc**1** | 4.72 | 3.67 | 3.71 | 3.60 | 3.93 | 1.05 | - |
| Fuc**2** | 5.05 | 3.60 | 3.70 | 3.62 | 4.75 | 1.09 | - |
| Gal**1** | 4.39 | 3.46 | 3.86 | N/R | N/R | N/R | - |
| Gal**2** | 4.44 | 3.45 | 4.01 | 3.86 | N/R | N/R | - |

| | **H1** | **H2** | **H3** | **H4** | **H5** | **H6** | **H7** | **H8** | **H9** |
|---|---|---|---|---|---|---|---|---|---|
| Neu5Ac9 | **-** | **-** | 2.69<br>1.72 | 3.60 | 3.77 | **-** | **-** | **-** | **-** |

| **Signal** | **Proton** | |
|---|---|---|
| **Aromatic** | 7.92-7.85 (m, 4H)<br>7.53-7.48 (m, 3H) | 128.4-125.8 |
| **$CH_2$**Ph | 5.33 (d, *J* = 12.8 Hz, 1H)<br>5.14 (d, *J* = 12.8 Hz, 1H) | 66.8 |
| NH-**CH**-COOH | 4.29 (dd, *J* = 9.5, 3.6 Hz 1H) | 52.8 |
| C(O)-**CH**-COOH | 2.75 (dd, *J* = 14.8, 2.9, 1H)<br>2.51 (dd, *J* = 15.0, 9.9, 1H) | 39.8 |

| | **C1** |
|---|---|
| GlcNAc**1** | 78.2 |
| GlcNAc**2** | 100.9 |
| GlcNAc**3** | 99.4 |
| GlcNAc**4** | 99.4 |
| Man**1** | 100.5 |
| Man**2** | 99.6 |
| Man**3** | 96.8 |
| Fuc**1** | 99.3 |
| Fuc**2** | 98.5 |
| Gal**1** | 103.0 |
| Gal2 | 101.7 |

[a] Not reported

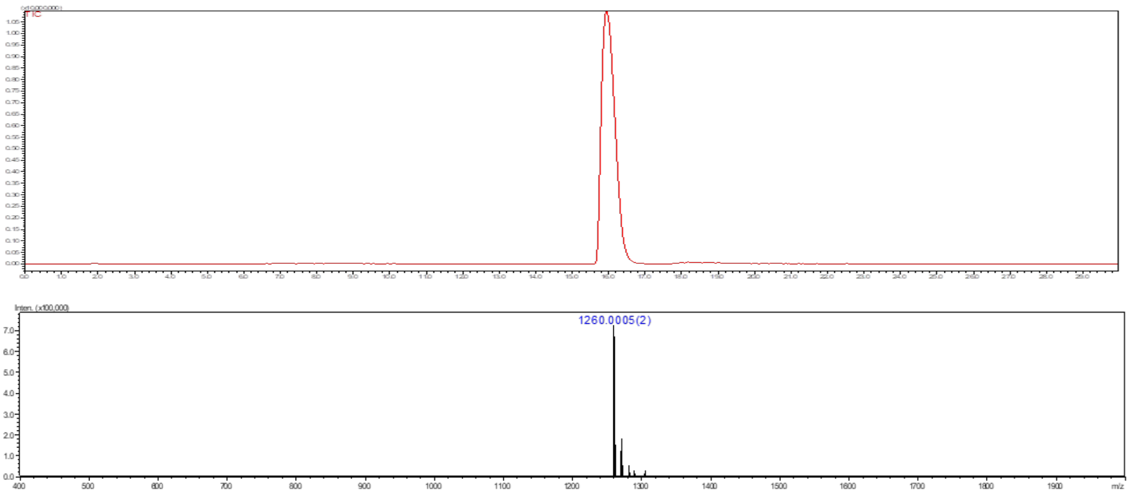

**Figure S56.** Liquid chromatography (LC) trace and mass (ESI-MS) spectrum (peak at retention time = 16.1 min) of **S12**.

### Compound F′

The amine of *N*-glycan compound **S12** was subjected for the hydrogenation using a general protocol **3k** to obtain **F′** as a white solid.

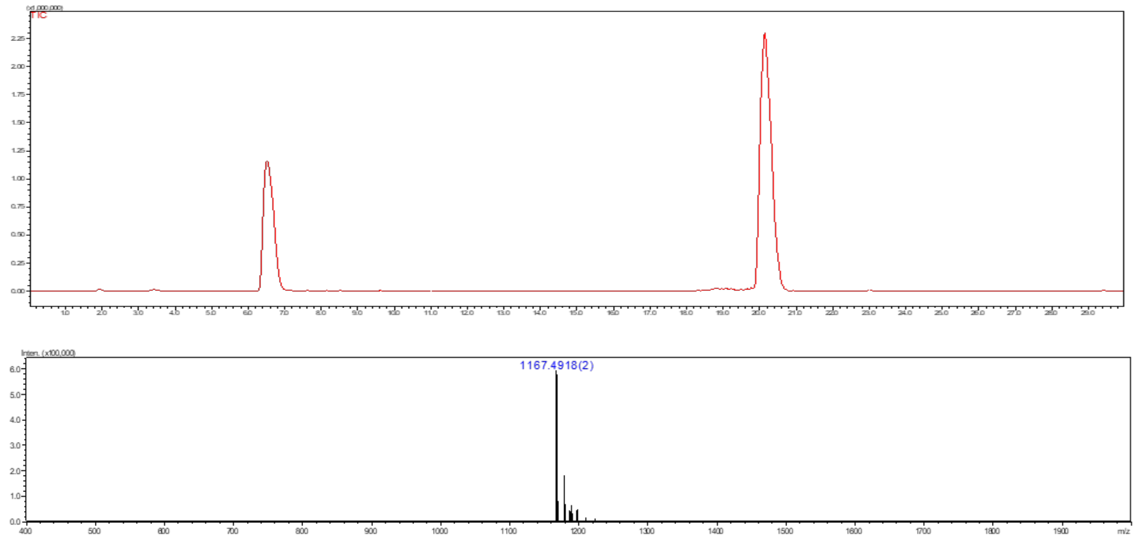

**Figure S57.** Liquid chromatography (LC) trace and mass (ESI-MS) spectrum (peak at retention time = 20.2 min) of **F′**.

### Compound S15

*N*-Glycan **2** (12 mg, 10.90 μmol) was dissolved in deionized water with the final reaction concentration of 5 mM. The pH of the reaction solution was adjusted to 10 using NaOH (1 M), and the mixture solution was incubated for 3-5 h at 37 °C with shaking. The complete conversion of starting material to amine intermediate was confirmed by LC-MS. Then, reaction was neutralized by aqueous acetic acid (1 M). The product was purified by C18 biogel [Eluent - water and water:acetonitrile (8:2)]. Next, resulting compound was subjected for the installation of a β1,4 Gal using the general procedure **3f** to obtained followed by acetylation using a general protocol **3j** to obtain **S13** as a white solid (10.5 mg, 77%).

**S13** (6 mg, 3.2 μmol) was subjected to installation of 2, 3-Sialic acid following the general protocol **3g** for the installation of Neu5Ac moiety using ST3Gal4 to get product **S14** as a white fluffy solid.

**S15** was synthesized from **S14** following the general protocol **3h** for the installation of Fuc moiety using FUT6. The product **S15** was obtained as a white fluffy solid (6.2 mg, 78%).

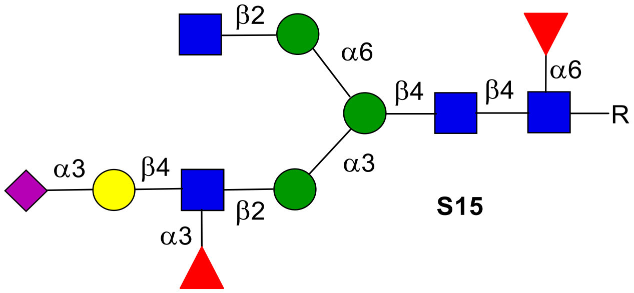

**S15**

**$^1$H NMR (600 MHz, $D_2O$) δ (ppm)**

| | **H1** | **H2** | **H3** | **H4** | **H5** | **H6** | **NHAc** |
|---|---|---|---|---|---|---|---|
| GlcNAc**1** | 4.86 | 3.69 | 3.60 | 3.45 | N/R[a] | N/R | 1.97-1.79 (12H) |
| GlcNAc**2** | 4.54 | 3.70 | 3.64 | N/R | N/R | N/R | 1.97-1.79 (12H) |
| GlcNAc**3** | 4.50 | 3.63 | 3.48 | 3.37 | N/R | N/R | 1.97-1.79 (12H) |
| GlcNAc**4** | 4.48 | 3.63 | 3.48 | 3.37 | N/R | N/R | 1.97-1.79 (12H) |
| Man**1** | 4.69 | 4.18 | 3.70 | N/R | N/R | N/R | - |
| Man**2** | 5.03 | 4.12 | 3.82 | 3.67 | 3.40 | N/R | - |
| Man**3** | 4.84 | 4.04 | 3.81 | 3.55 | N/R | N/R | - |
| Fuc**1** | 4.72 | 3.66 | 3.72 | 3.60 | 3.95 | 1.06 | - |
| Gal**1** | 4.43 | 3.44 | 4.01 | 3.86 | N/R | N/R | - |
| Fuc**2** | 5.04 | 3.61 | 3.83 | 3.70 | 4.74 | 1.09 | - |

| | **H1** | **H2** | **H3** | **H4** | **H5** | **H6** | **H7** | **H8** | **H9** |
|---|---|---|---|---|---|---|---|---|---|
| Neu5Ac9 | - | - | 2.69<br>1.72 | 3.60 | 3.77 | - | - | - | - |

| **Signal** | **Proton** | |
|---|---|---|
| **Aromatic** | 7.92-7.85 (m, 4H)<br>7.53-7.48 (m, 3H) | 128.4-125.8 |
| **$CH_2$**Ph | 5.33 (d, *J* = 12.8 Hz, 1H) | 66.8 |

| | | |
|---|---|---|
| | 5.14 (d, $J$ = 12.8 Hz, 1H) | |
| NH-**CH**-COOH | 4.29 (dd, $J$ = 9.5, 3.6 Hz 1H) | 52.8 |
| C(O)-**CH**-COOH | 2.75 (dd, $J$ = 14.8, 2.9, 1H)<br>2.51 (dd, $J$ = 15.0, 9.9, 1H) | 39.8 |

| | **C1** |
|---|---|
| GlcNAc**1** | 78.2 |
| GlcNAc**2** | 100.7 |
| GlcNAc**3** | 99.2 |
| GlcNAc**4** | 99.5 |
| Man**1** | 100.4 |
| Man**2** | 99.5 |
| Man**3** | 97.0 |
| Fuc**1** | 99.1 |
| Gal**1** | 101.6 |
| Fuc**2** | 98.5 |

[a] Not reported

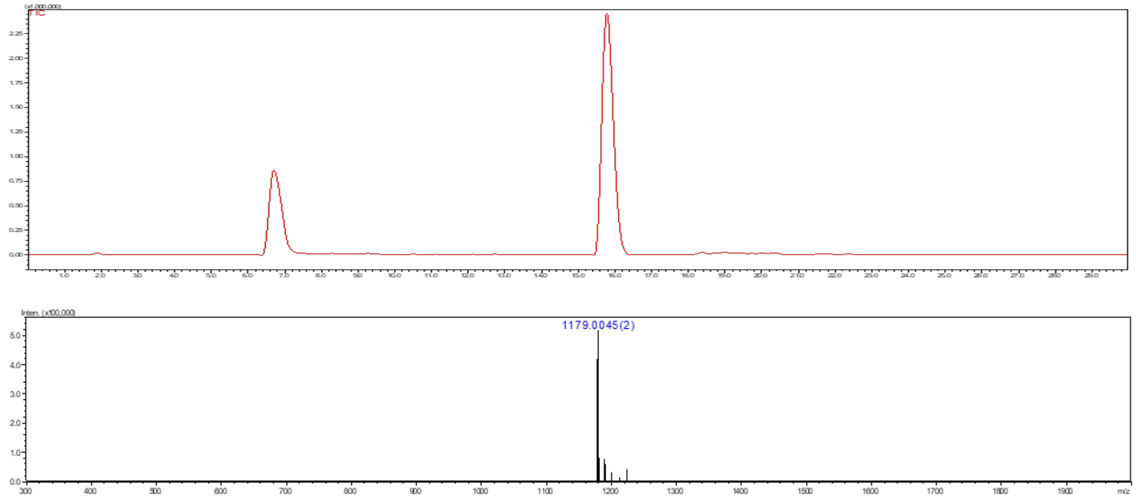

**Figure S58.** Liquid chromatography (LC) trace and mass (ESI-MS) spectrum (peak at retention time = 15.9 min) of **S15**.

### Compound K′

The amine of *N*-glycan compound **S15** was subjected for the hydrogenation using a general protocol **3k** to obtain **K′** as a white solid.

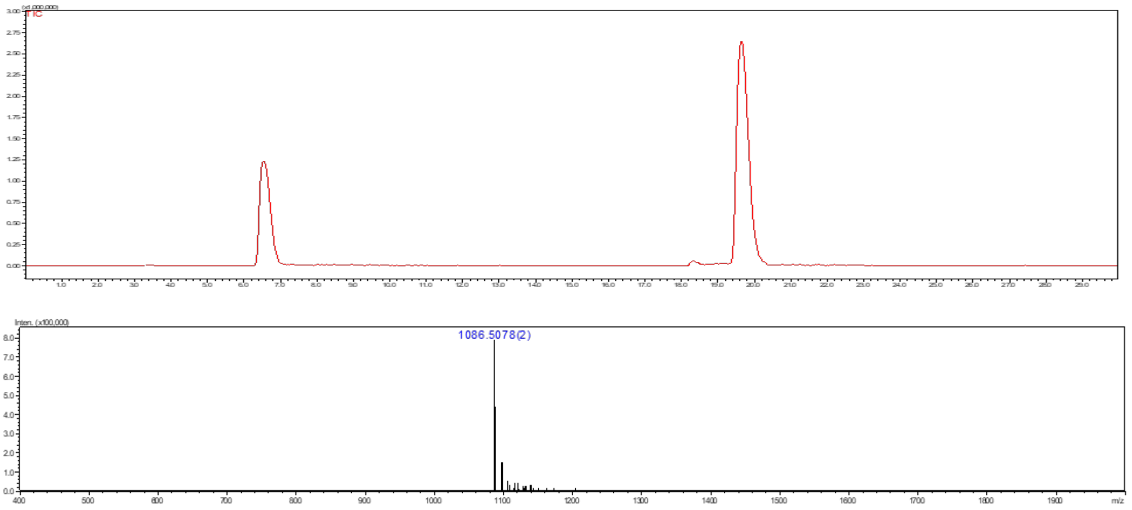

**Figure S59.** Liquid chromatography (LC) trace and mass (ESI-MS) spectrum (peak at retention time = 19.9 min) of **K′**.

### Compound S16

**S16** was synthesized from **S15** (2.5 mg, 0.81 μmol) following the general protocol **3f** for the installation of Gal moiety using B4GalT1. The product **S16** was obtained as a white fluffy solid (2.14 mg, 80%).

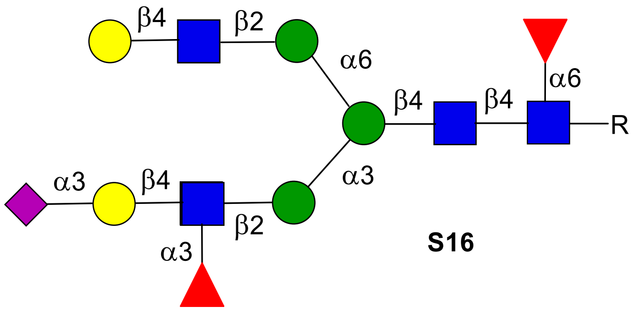

**S16**

**$^1$H NMR (600 MHz, $D_2O$) δ (ppm)**

| | H1 | H2 | H3 | H4 | H5 | H6 | NHAc |
|---|---|---|---|---|---|---|---|
| GlcNAc**1** | 4.86 | 3.70 | 3.60 | N/R[a] | N/R | N/R | 1.98-3.80 (12H) |
| GlcNAc**2** | 4.55 | 3.71 | 3.65 | 3.65 | 3.49 | N/R | 1.98-3.80 (12H) |
| GlcNAc**3** | 4.50 | 3.84 | N/R | N/R | N/R | N/R | 1.98-3.80 (12H) |
| GlcNAc**4** | 4.50 | 3.68 | 3.60 | 3.49 | N/R | N/R | 1.98-3.80 (12H) |
| Man**1** | 4.69 | 4.17 | 3.70 | 3.50 | N/R | N/R | - |
| Man**2** | 5.03 | 4.11 | 3.82 | 3.66 | N/R | N/R | - |
| Man**3** | 4.85 | 4.02 | 3.83 | 3.54 | N/R | N/R | - |
| Fuc**1** | 4.72 | 3.67 | 3.72 | 3.60 | 3.94 | 1.07 | - |
| Fuc**2** | 5.04 | 3.60 | 3.83 | 3.67 | 4.75 | 1.10 | - |
| Gal**1** | 4.43 | 3.45 | 4.00 | 3.85 | N/R | N/R | - |
| Gal**2** | 4.39 | 3.46 | N/R | 3.85 | N/R | N/R | - |

| | H1 | H2 | H3 | H4 | H5 | H6 | H7 | H8 | H9 |
|---|---|---|---|---|---|---|---|---|---|
| Neu5Ac9 | **-** | **-** | 2.69<br>1.72 | 3.60 | 3.77 | **-** | **-** | **-** | **-** |

| Signal | Proton | Carbon |
|---|---|---|
| **Aromatic** | 7.92-7.85 (m, 4H)<br>7.53-7.48 (m, 3H) | 128.4-<br>125.8 |
| **$CH_2$**Ph | 5.33 (d, *J* = 12.8 Hz, 1H)<br>5.14 (d, *J* = 12.8 Hz, 1H) | 66.8 |
| NH-**CH**-COOH | 4.29 (dd, *J* = 9.5, 3.6 Hz 1H) | 52.8 |
| C(O)-**CH**-COOH | 2.75 (dd, *J* = 14.8, 2.9, 1H)<br>2.51 (dd, *J* = 15.0, 9.9, 1H) | 39.8 |

| | **13C** |
|---|---|
| GlcNAc**1** | 78.0 |
| GlcNAc**2** | 100.8 |
| GlcNAc**3** | 99.1 |
| GlcNAc**4** | 99.1 |
| Man**1** | 100.3 |
| Man**2** | 99.5 |
| Man**3** | 96.9 |
| Fuc**1** | 99.4 |
| Fuc**2** | 98.5 |
| Gal**1** | 101.7 |
| Gal**2** | 103.1 |

[a] Not reported

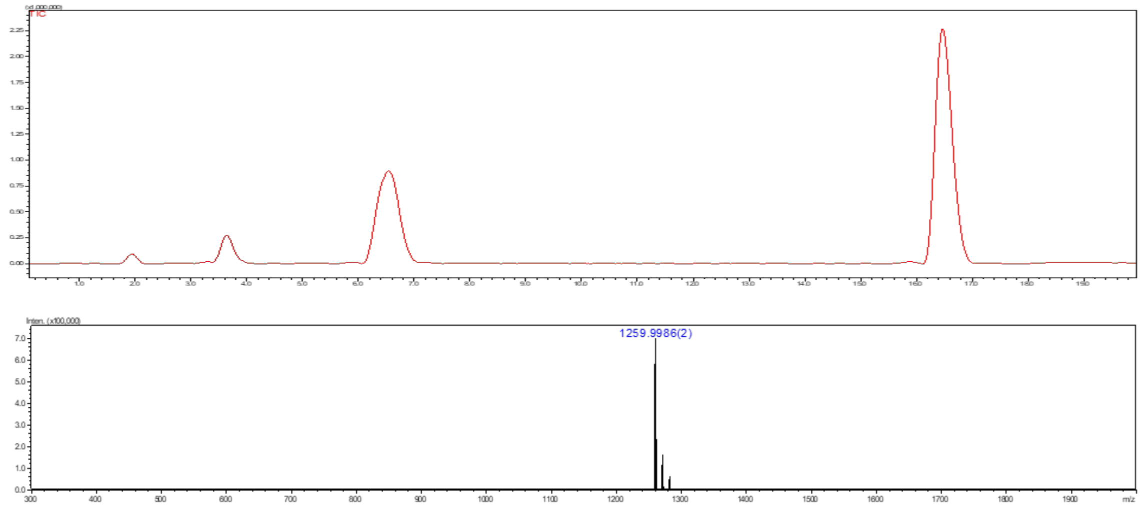

**Figure S60.** Liquid chromatography (LC) trace and mass (ESI-MS) spectrum (peak at retention time = 16.6 min) of **S16**.

**Compound L′**

The amine of *N*-glycan compound **47** was subjected for the hydrogenation using a general protocol **3k** to obtain **L′** as a white solid.

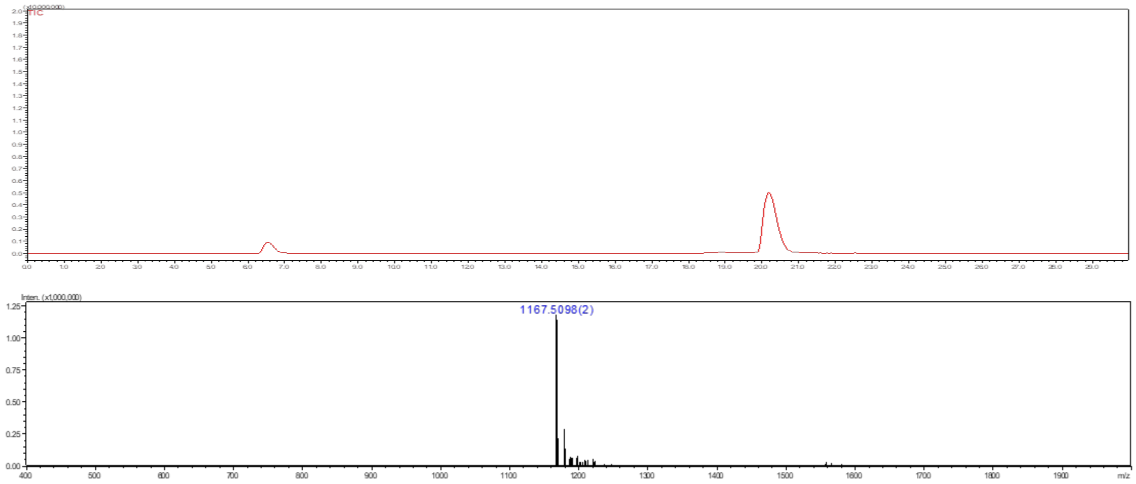

**Figure S61.** Liquid chromatography (LC) trace and mass (ESI-MS) spectrum (peak at retention time = 20.4 min) of **L′**.

### 9. Kinetic Parameters of GlcNAc Transfer GnT-III, GnT-IV and GnT-V for Multi-antennary Glycans

Following established protocol,[6] determination of GnT-III, GnT-IVB, and GnT-V steady state kinetic parameters for sugar donor and acceptor substrates was achieved in 5 µL reactions with 16 ng GnT-III, 50 ng GnT-IVB, and 35 ng GnT-V in Corning® Low Volume 384-well White Flat Bottom Polystyrene NBS Microplates (Catalog #3824). GnT-III and GnT-V were performed using a 50 mM MES, pH 6.5 buffer supplemented with 5 mM $MnCl_2$. GnT-IVB was performed using a 50 mM Tris-HCl, pH 7.5 supplemented with 5 mM $MnCl_2$. For substrate kinetics, acceptors were serially diluted from 2 mM to 0.03 mM or 10 mM to 0.16 mM for GnT-III/GnT-V and GnT-IVB respectively, with a constant donor concentration of 1 mM, 7 mM, and 10 mM for GnT-III, GnT-IVB, and GnT-V respectively. For GnT-III donor kinetics, UDP-GlcNAc was serially diluted from 6 mM to 0.09 mM with a constant acceptor concentration of 0.4 mM. Reactions were incubated for 1 h at 37°C. UDP formation was detected using a UDP-Glo assay following the manufacturer's procedure (Catalog #V6971). Luminescence values were associated with a UDP standard curve for quantification of UDP generated and Michaelis-Menten kinetic $K_{cat}$, $K_m$, and $V_{max}$ were calculated through nonlinear curve fitting in GraphPad Prism 6.

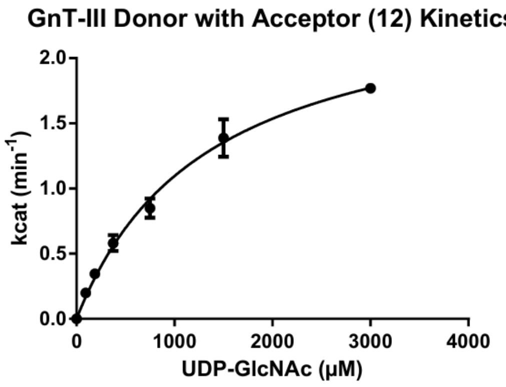

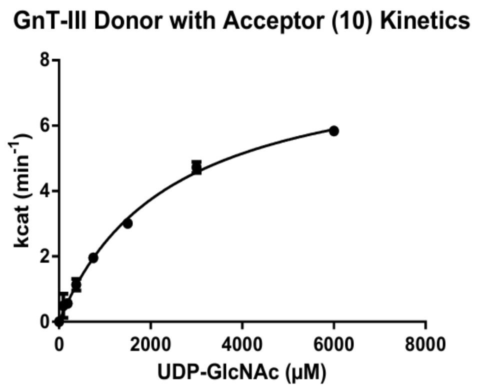

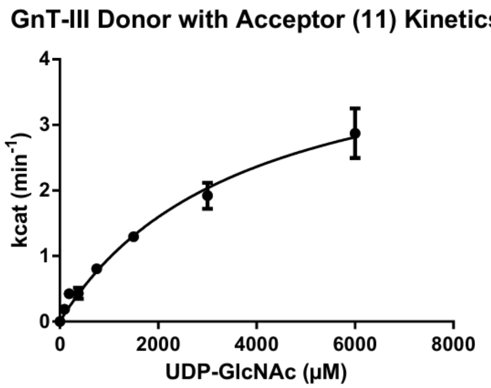

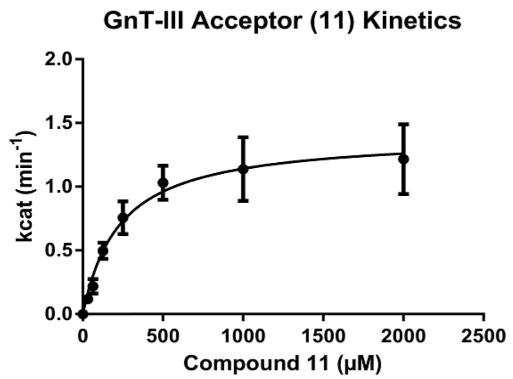

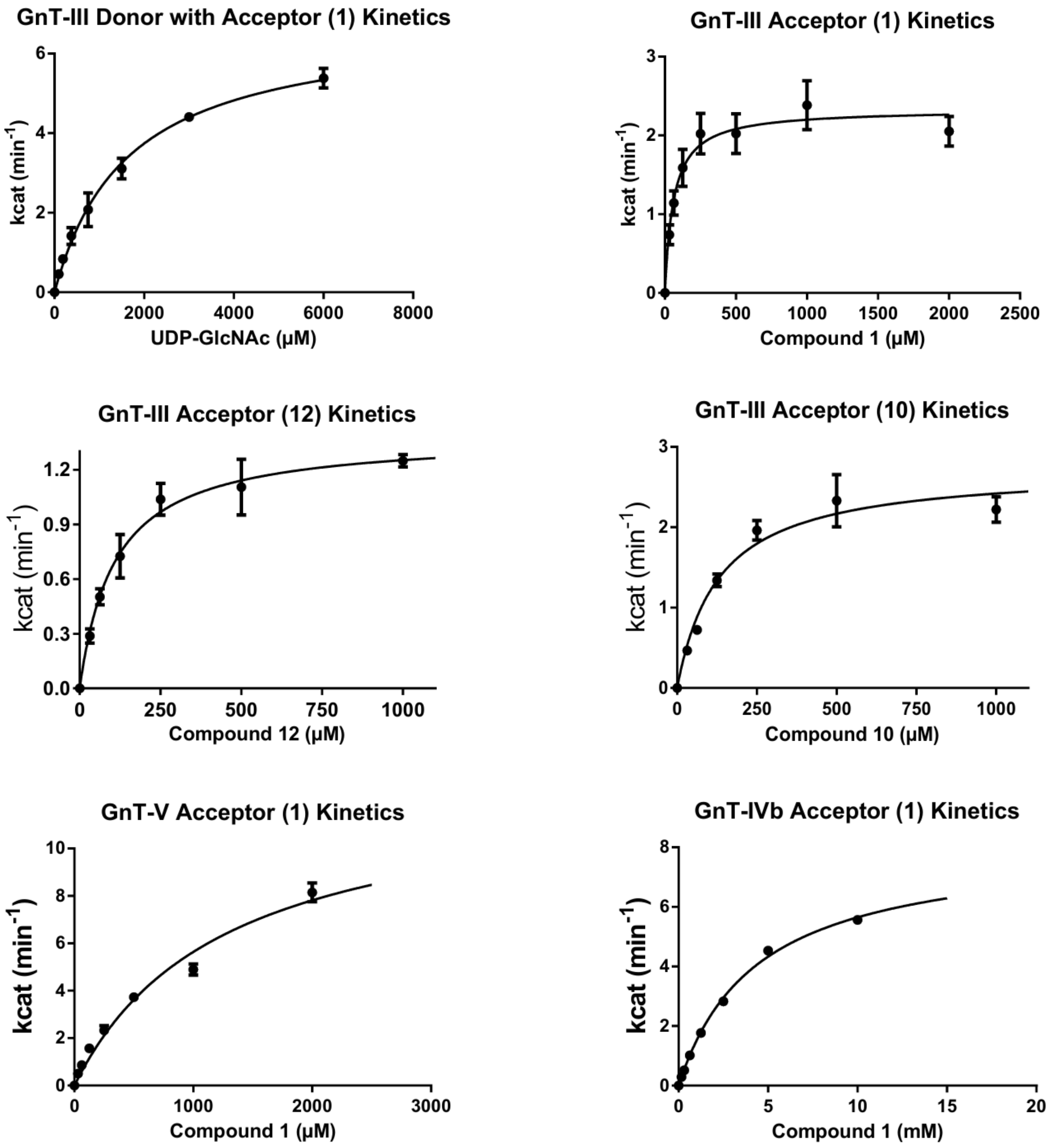

**Figure S62.** Steady state kinetics of GnT-III, GnT-IVB, and GnT-V as determined by UDP-Glo assay. Acceptors: **1**, **10**, **11**, **12**, **S17**, and **S18**. Donor: UDP-GlcNAc. All reactions performed in triplicate with kinetic parameters determined via nonlinear curve fitting in GraphPad Prism 6.

### 10. Glycan Microarray Development

The *N*-glycans containing free α-amine of Asn moiety were printed on NHS-ester activated glass slides (NEXTERION® Slide H, Schott Inc.) using a Scienion sciFLEXARRAYER S3 non-contact microarray equipped with a Scienion PDC80 nozzle (Scienion Inc.). All samples at a concentration of 100 μM in sodium phosphate buffer (0.225 M, pH 8.5) were printed in replicates of 6 with spot volume ~ 400 pL, at 20 °C and 50% relative humidity (each slide has 24 subarrays in a 3x8 layout). After printing, slides were incubated overnight in a saturated NaCl chamber (affording a 75% relative humidity environment), the remaining activated esters were then quenched with 5 mM ethanolamine in a Tris buffer (pH 9.0, 50 mM) at 50 °C for 1 h. Blocked slides were rinsed with DI water, spun dry, and kept in a desiccator at room temperature for future use. Printed glass slide was incubated with protein of interest. After 1 h, the slide was sequentially washed by dipping in TSM wash buffer (2 min, containing 0.05 % Tween 20), TSM buffer (2 min) and, water (2 x 2 min), spun dry. The slide was incubated with fluorescent detection antibody (for details see Tables S1 and S2, in case of anti-CD15s premixed primary antibody and secondary fluorescent antibody were employed) for 1 h and same washing sequence was repeated. The slides were scanned using a GenePix 4000B microarray scanner (Molecular Devices) at the appropriate excitation wavelength with a resolution of 5 μm. Various gains and PMT values were employed in the scanning to ensure that all the signals were within the linear range of the scanner's detector and there was no saturation of signals. The images were analyzed using GenePix Pro 7 software (version 7.2.29.2, Molecular Devices). The data was analyzed with our home written Excel macro to provide the results. The highest and the lowest value of the total fluorescence intensity of the six replicates spots were removed, and the four values in the middle were used to provide the mean value and standard deviation. The data were fitted using Prism software 8 (GraphPad Software, Inc.), bar represents the mean ± SD for each compound. All experiments were performed three times at the minimum.

**Table S1.** List of protein, protein source (type of fusion tag, if present), primary antibodies and secondary antibodies (type of fluorophore, if present) screened against the *N*-glycans-microarray.

| Sr. # | Protein | Source | Tag | Detection | Source |
|---|---|---|---|---|---|
| 1 | *Phaseolus vulgaris* E lectin (PHA-E) | Vector Labs Cat # B-1125-2 | Biotin | Streptavidin-Alexa Fluor® 647 | Thermo, Cat # S32364 |
| 2 | *Concanavalin* A (ConA) | Vector Labs Cat # B-1005-5 | Biotin | Streptavidin-Alexa Fluor® 647 | Thermo, Cat # S32364 |
| 3 | *Aleuria aurantia* lectin (AAL) | Vector Labs Cat # B-1395-1 | Biotin | Streptavidin-Alexa Fluor® 647 | Thermo, Cat # S32364 |
| 4 | *Sambucus nigra* agglutinin (SNA) | Vector Labs Cat # B-1305-2 | Biotin | Streptavidin-Alexa Fluor® 647 | Thermo, Cat # S32364 |
| 5 | *Maackia amurensis* lectin I (MAL-I) | Vector Labs Cat # B-1315-2 | Biotin | Streptavidin-Alexa Fluor® 647 | Thermo, Cat # S32364 |
| 6 | *Wheat Germ* Agglutinin (WGA) | Vector Labs Cat # B-1025-5 | Biotin | Streptavidin-Alexa Fluor® 647 | Thermo, Cat # S32364 |
| 7 | *Datura stramonium* lectin (DSL) | Vector Labs Cat # B-1185-2 | Biotin | Streptavidin-Alexa Fluor® 647 | Thermo, Cat # S32364 |
| 8 | *Griffonia simplicifolia* lectin II (GSL-II) | Vector Labs Cat # B-1215-2 | Biotin | Streptavidin-Alexa Fluor® 647 | Thermo, Cat # S32364 |
| 9 | *Wisteria floribunda* lectin (WFL) | Vector Labs | Biotin | Streptavidin-Alexa Fluor® 647 | Thermo, Cat # S32364 |

| | | | | | |
|---|---|---|---|---|---|
| | | Cat # B-1355-2 | | | |
| 10 | *Erythrina cristagalli* lectin (ECL) | Vector Labs Cat # B-1145-5 | Biotin | Streptavidin-Alexa Fluor® 647 | Thermo, Cat # S32364 |
| 11 | Anti-CD15s antibody | BD Pharmigen Cat # 551344 | untagged | Alexa Fluor® 647 anti-mouse IgG antibody | Jackson Immuno Res |

**Table S2.** List of viruses, haemagglutinin anti-stem antibody, detecting antibody, and source screened against *N*-glycans-microarray.

| Sr. # | Virus | Haemagglutinin anti-stem antibody | Detecting antibody | Source |
|---|---|---|---|---|
| A | A/California/07/2009 (pdmH1N1) | 1A06[7] | Alexa Fluor® 647 anti-human IgG antibody | Jackson Immuno Res |
| B | A/Perth/16/2009 (H3N2) | CR8020 | Alexa Fluor® 647 anti-human IgG antibody | Jackson Immuno Res |
| C | A/Vietnam/1203/2004 (H5N1) | 1A06[7] | Alexa Fluor® 647 anti-human IgG antibody | Jackson Immuno Res |
| D | A/bovine/Ohio/B24OSU-432/2024 (H5N1) | 1A06[7] | Alexa Fluor® 647 anti-human IgG antibody | Jackson Immuno Res |

### 12. NMR Spectra

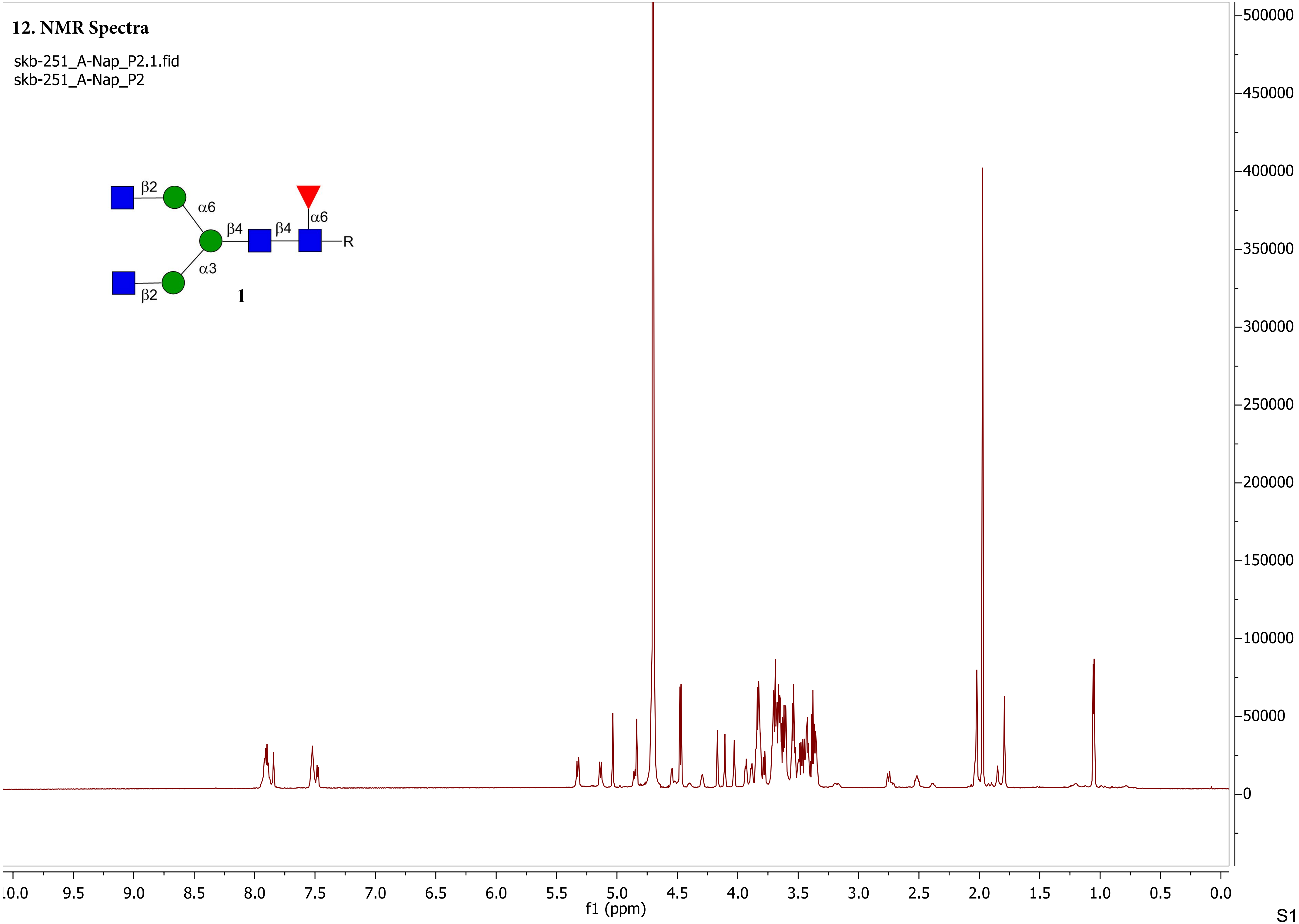

skb-251_A-Nap_P2.2.fid
skb-251_A-Nap_P2

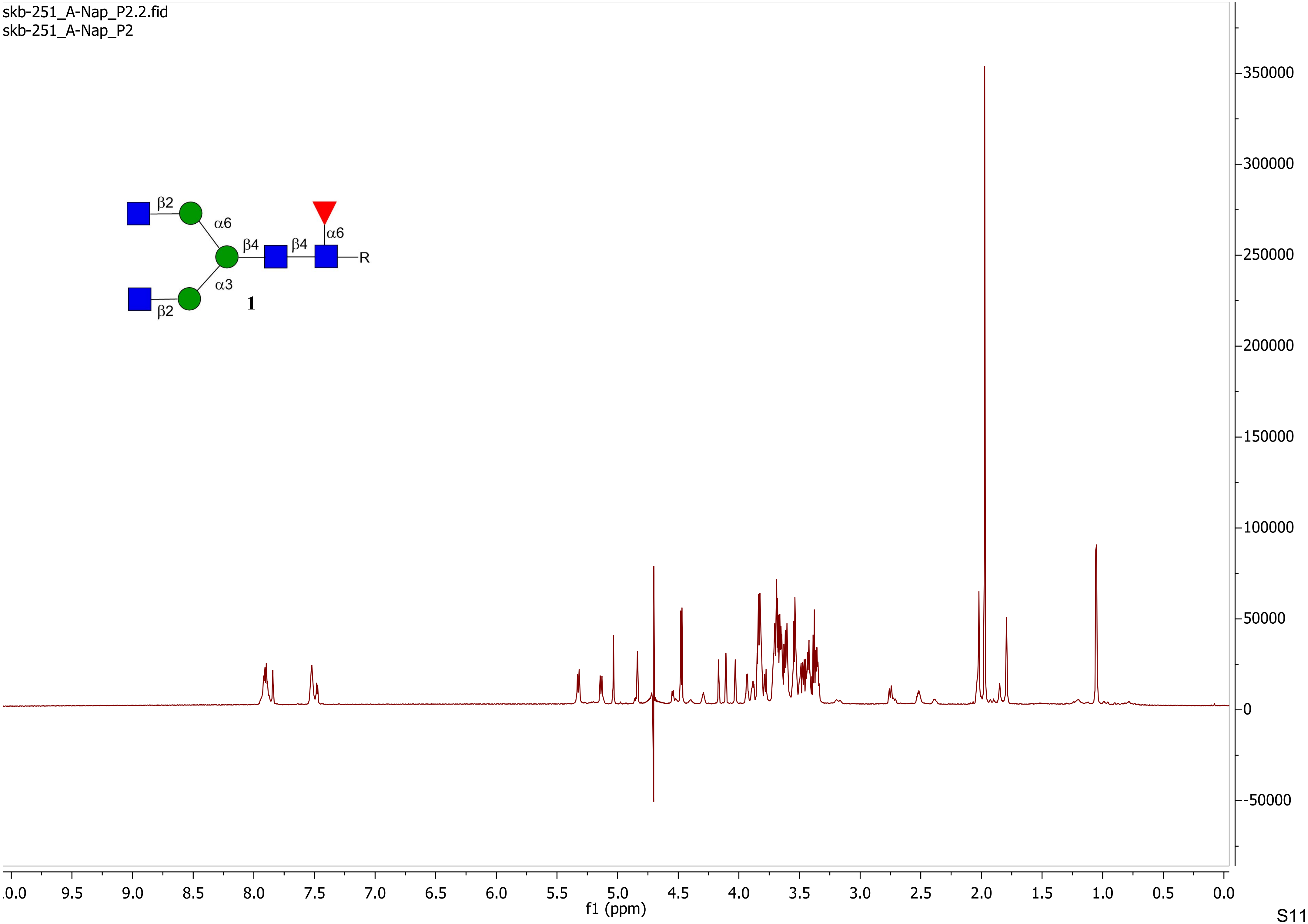

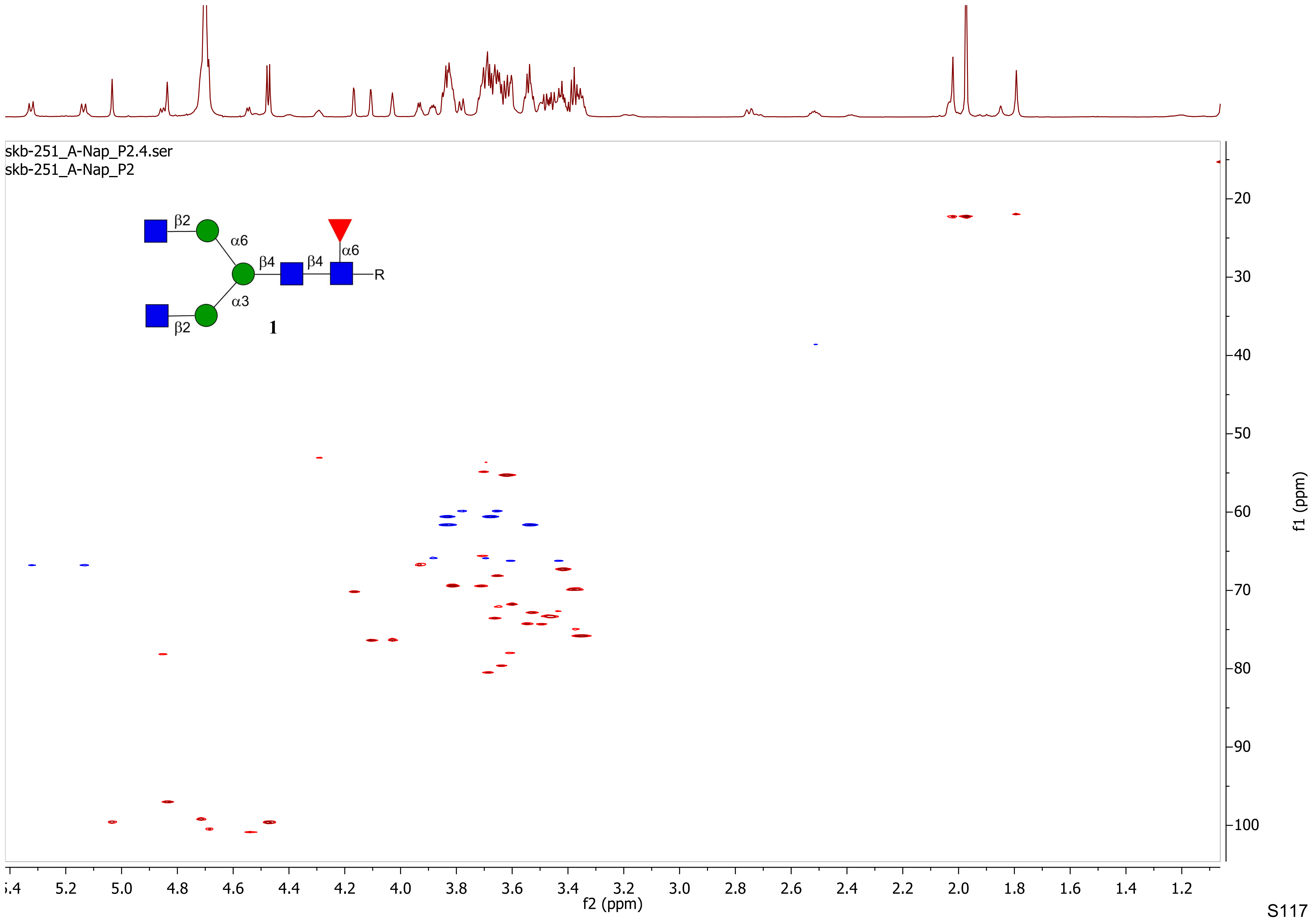

skb-251_A-Nap_P2.4.ser
skb-251_A-Nap_P2
β2
α6
β4
β4
α6
R
α3
β2
1
f1 (ppm)
f2 (ppm)

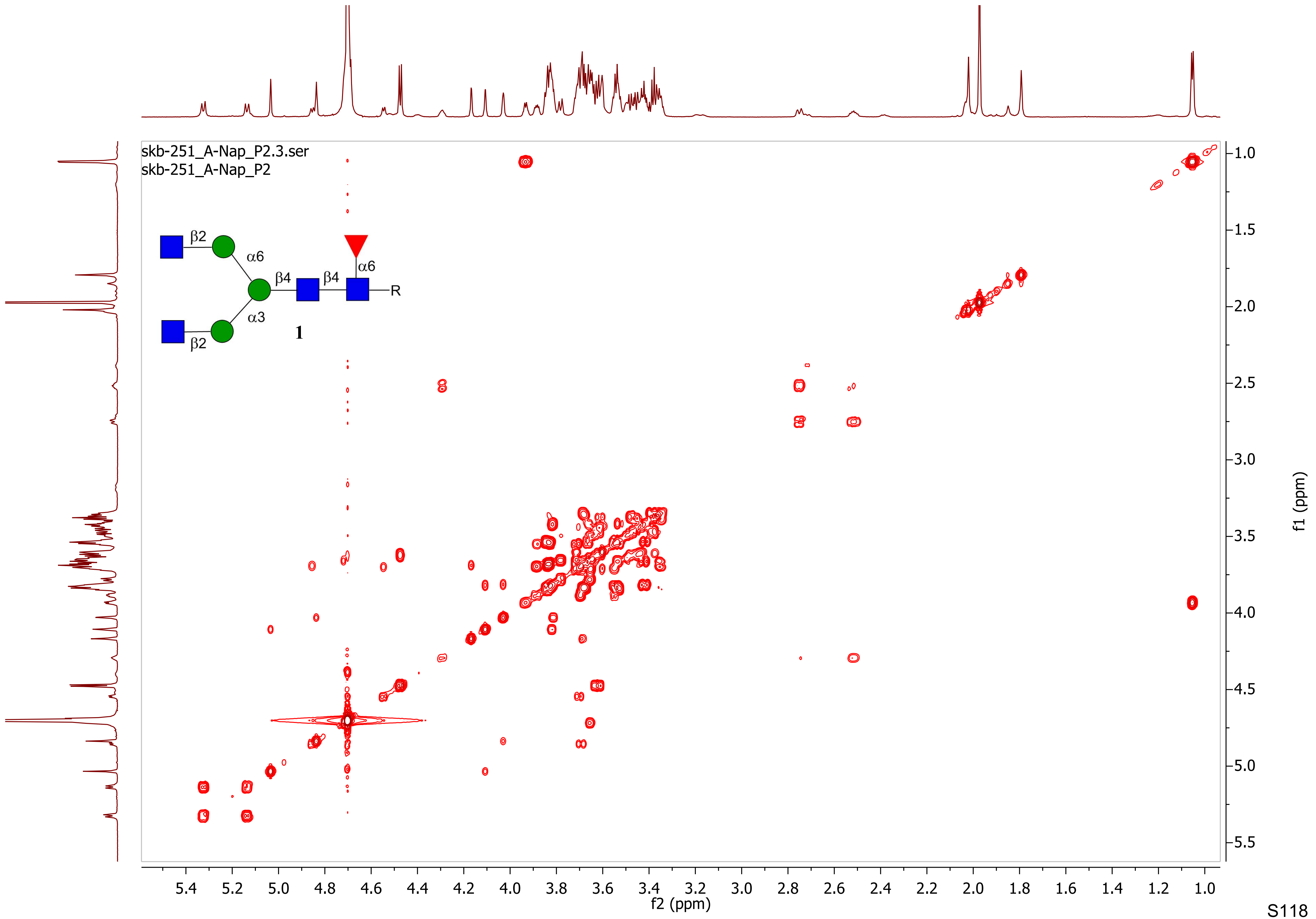

skb-251_A-Nap_P2.3.ser
skb-251_A-Nap_P2
β2
α6
β4
β4
α6
R
α3
β2
1
f2 (ppm)
f1 (ppm)

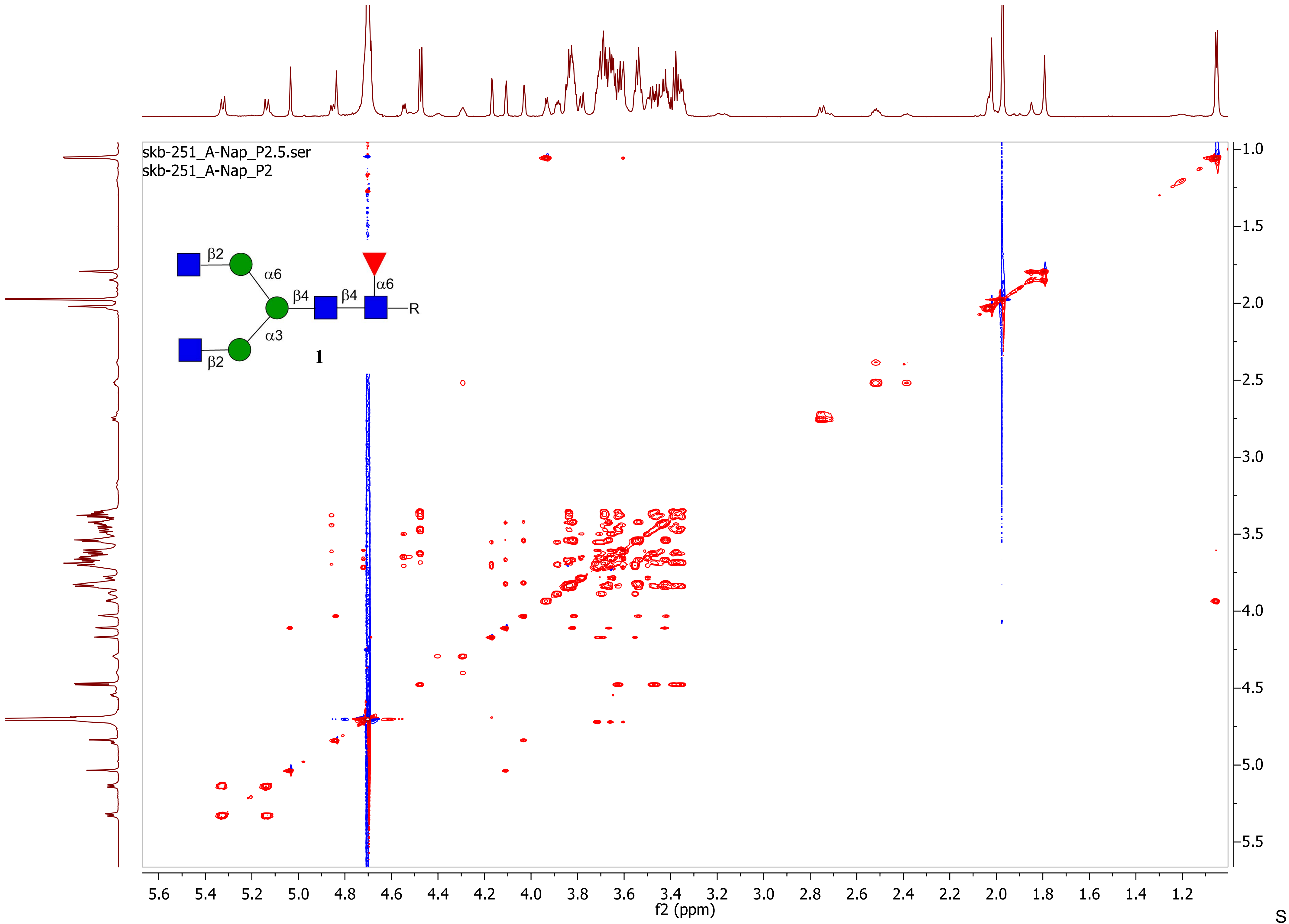

skb-251_A-Nap_P2.5.ser
skb-251_A-Nap_P2
β2
α6
β4
β4
α6
R
α3
β2
1
f2 (ppm)
f1 (ppm)

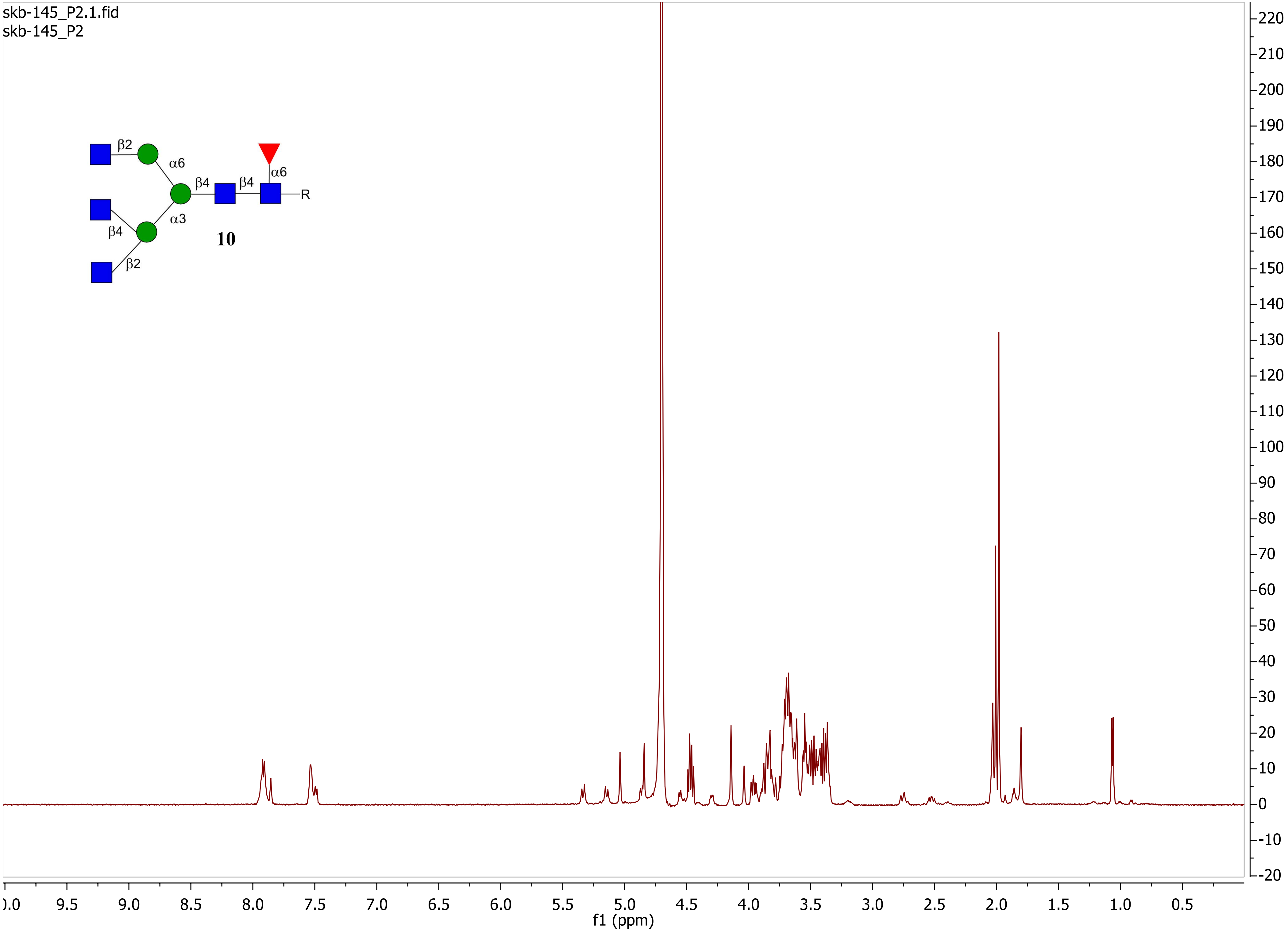

skb-145_P2.1.fid
skb-145_P2
β2
α6
β4
β4
α6
R
β4
α3
10
β2
f1 (ppm)

skb-145_P2.2.fid
skb-145_P2

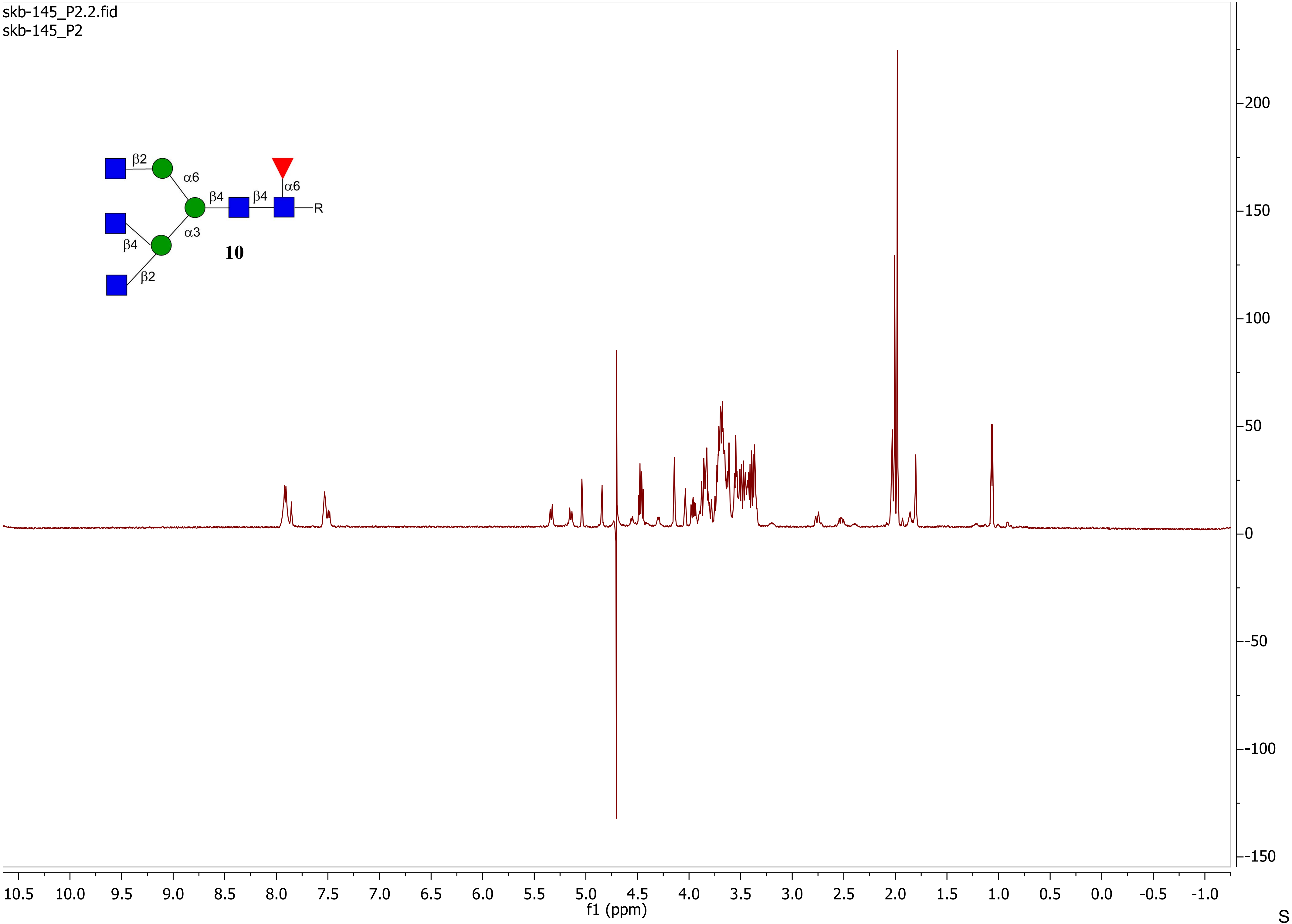

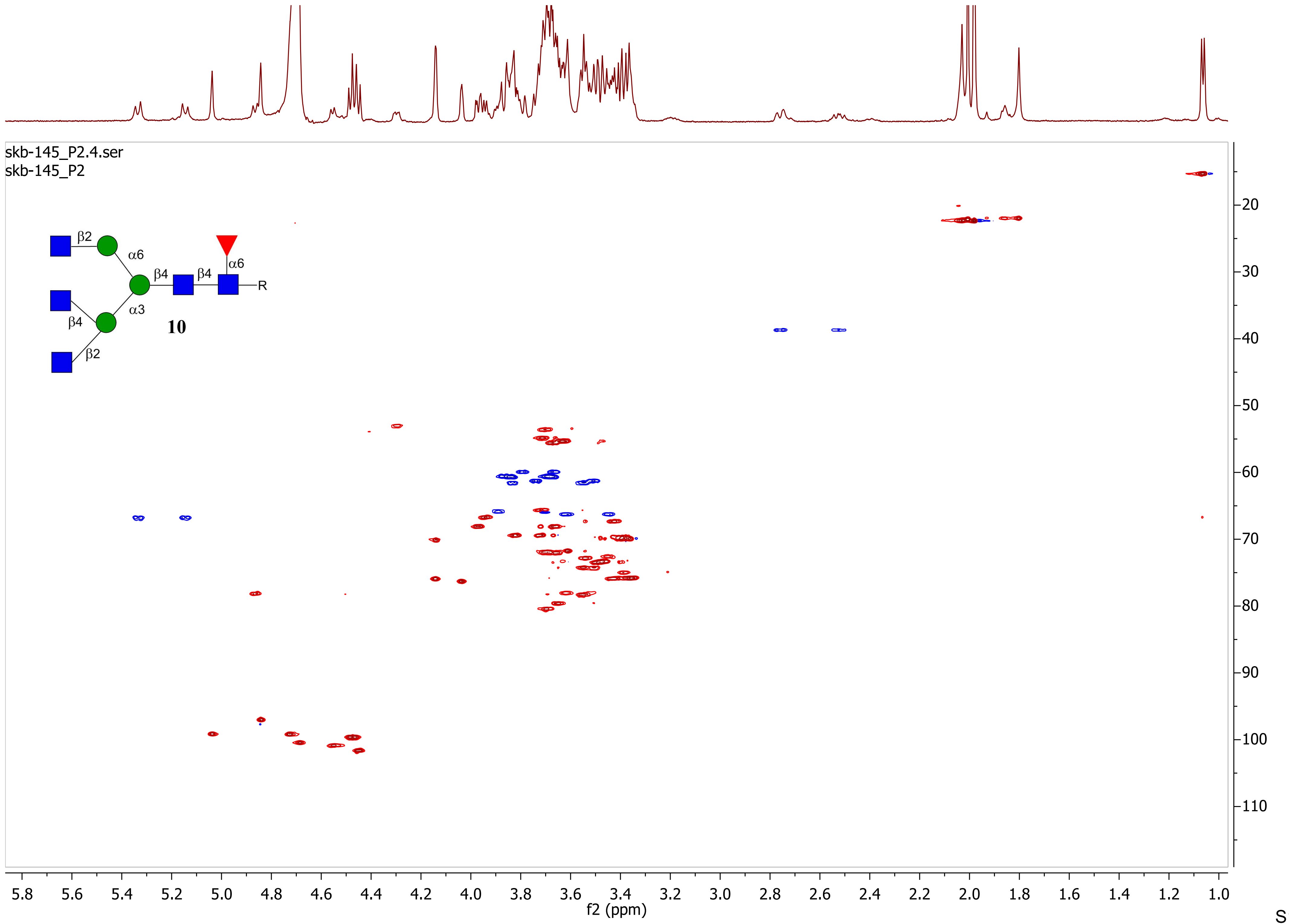
skb-145_P2.4.ser
skb-145_P2
β2
α6
β4
β4
α6
R
α3
β4
β2
10
f2 (ppm)
f1 (ppm)

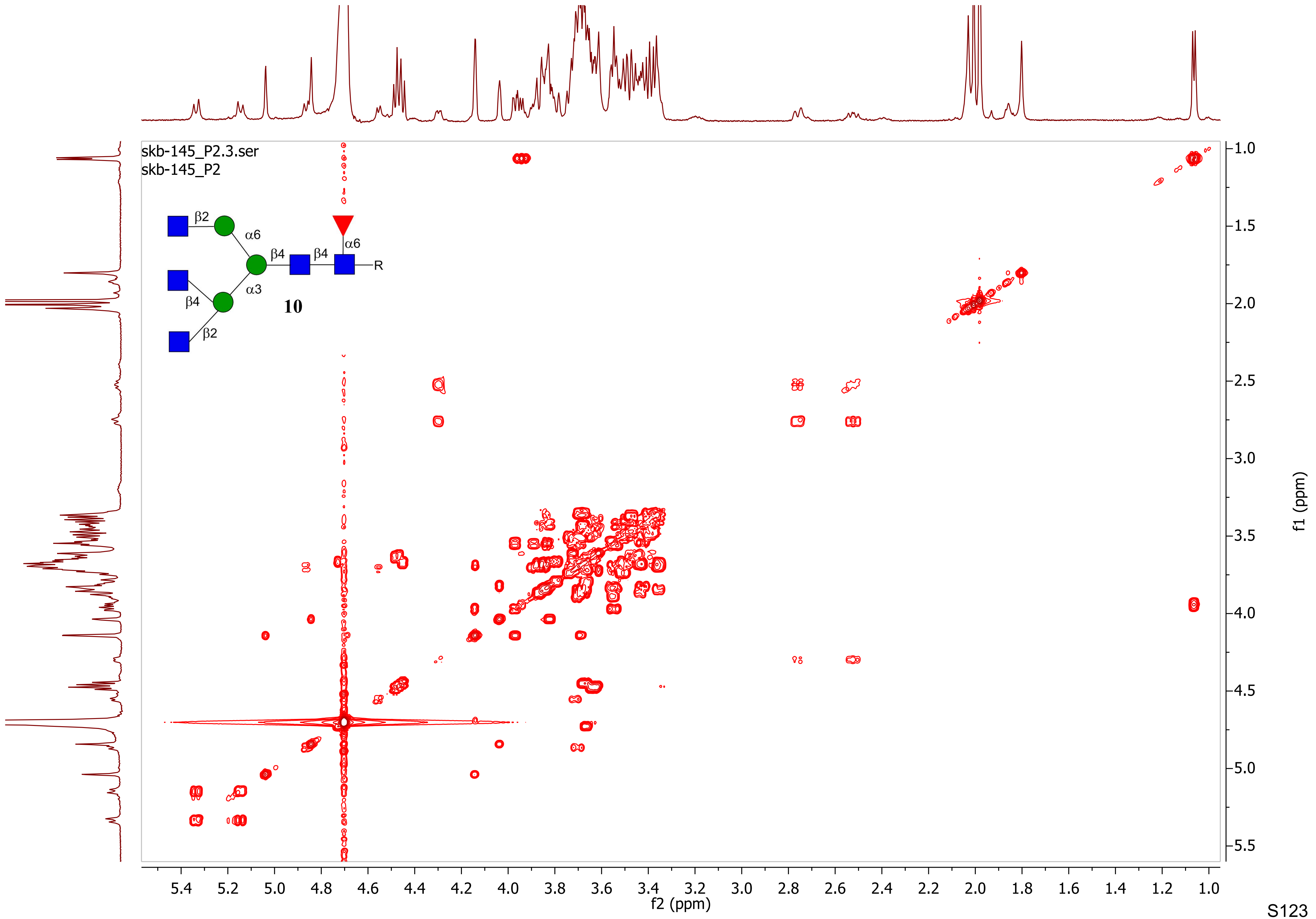

skb-145_P2.3.ser
skb-145_P2
β2
α6
β4
β4
α6
R
α3
β4
β2
10
f1 (ppm)
f2 (ppm)

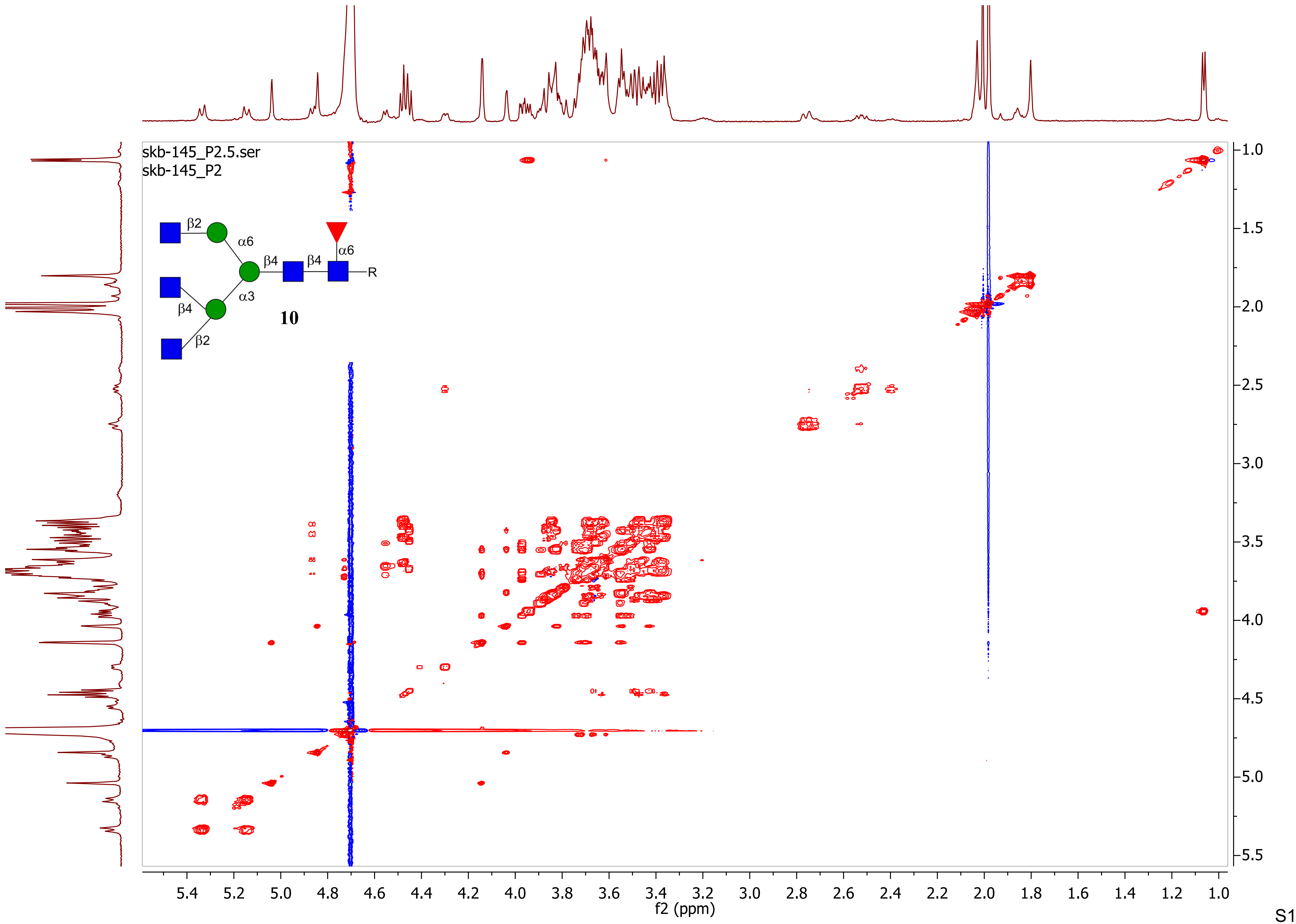

skb-145_P2.5.ser
skb-145_P2
β2
α6
β4
β4
α6
R
β4
α3
β2
10
f2 (ppm)
f1 (ppm)

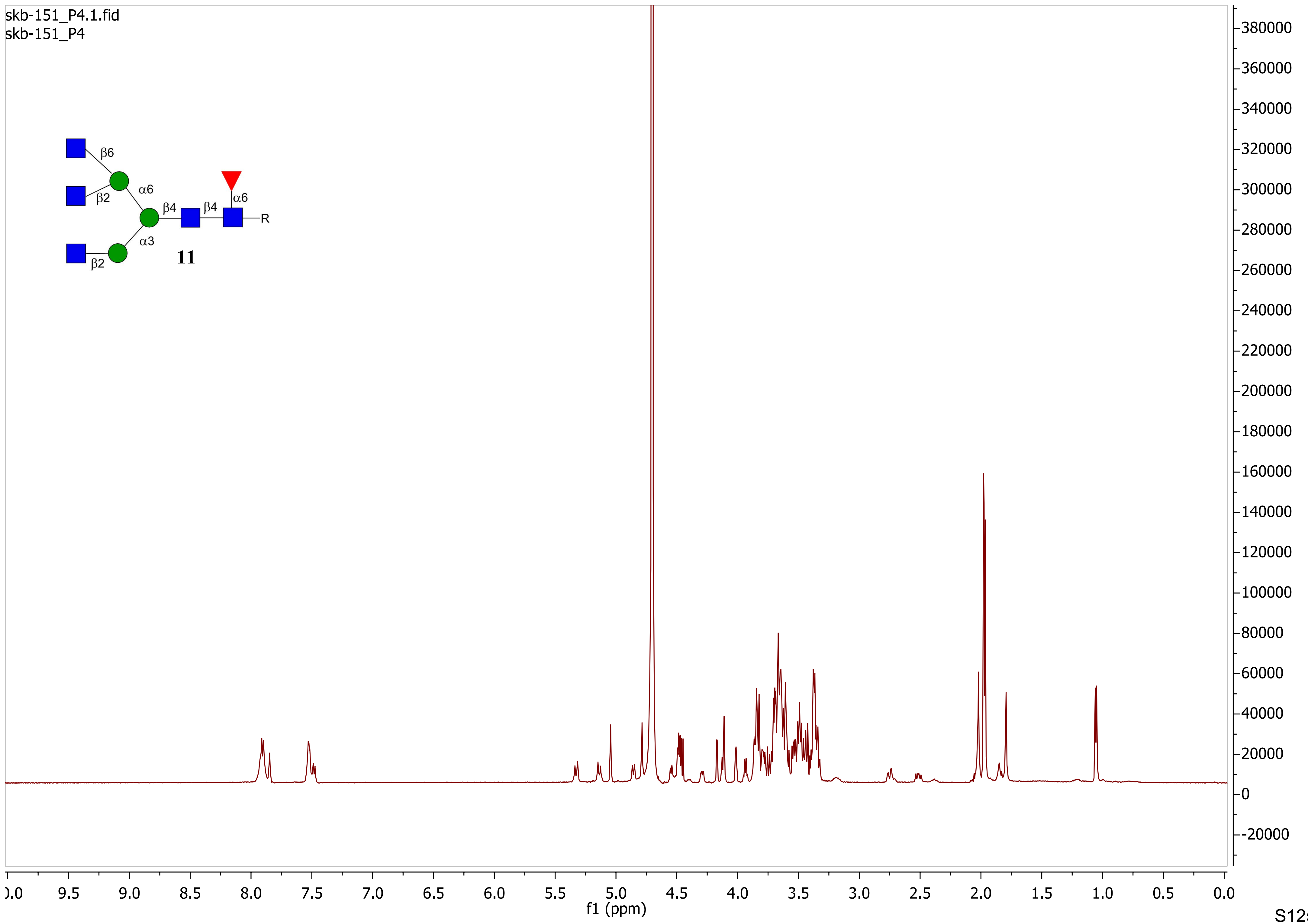

skb-151_P4.1.fid
skb-151_P4
β6
β2
α6
β4
β4
α6
R
α3
β2
11
f1 (ppm)

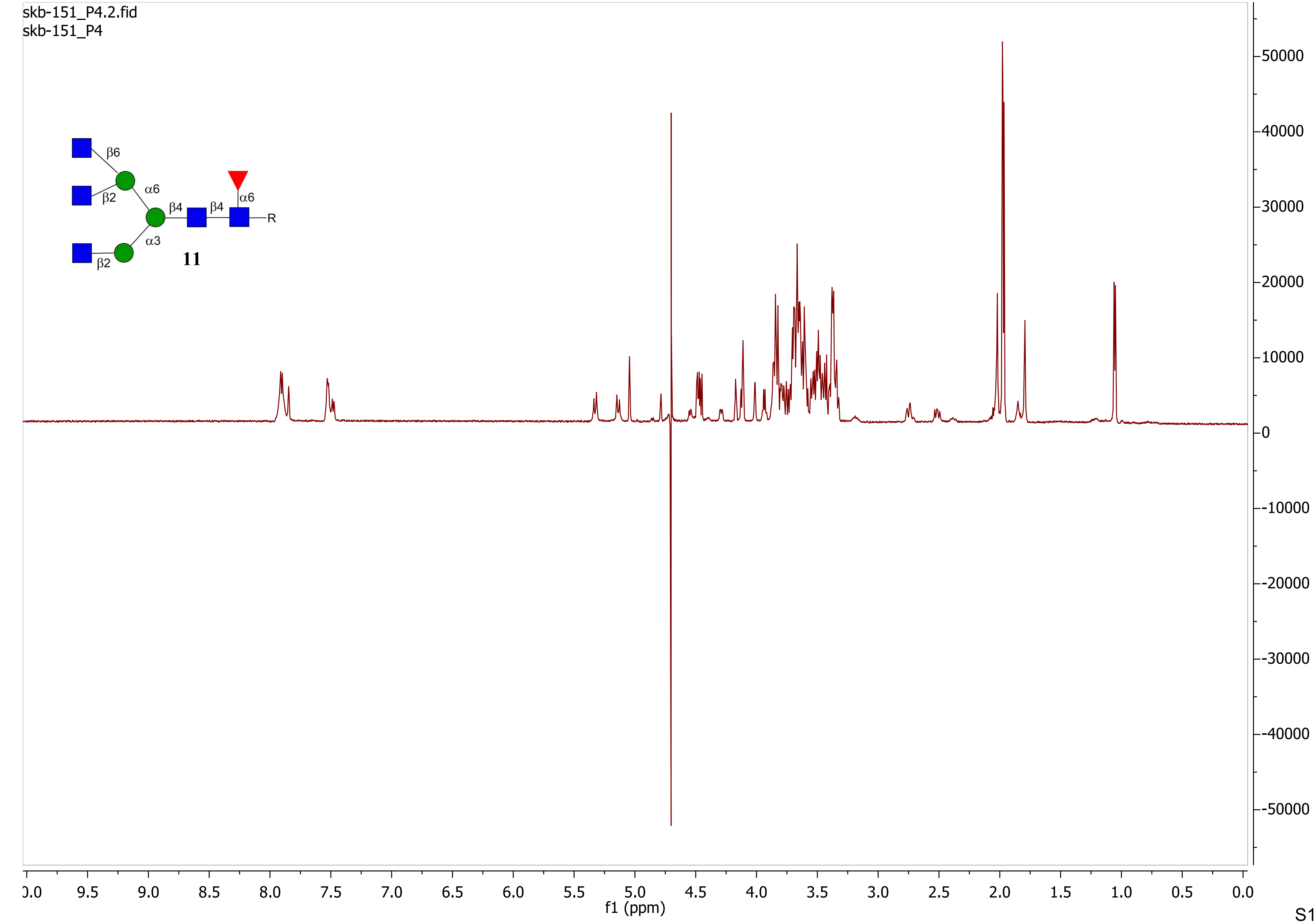
skb-151_P4.2.fid
skb-151_P4
β6
β2
α6
β4
β4
α6
R
α3
β2
11
f1 (ppm)

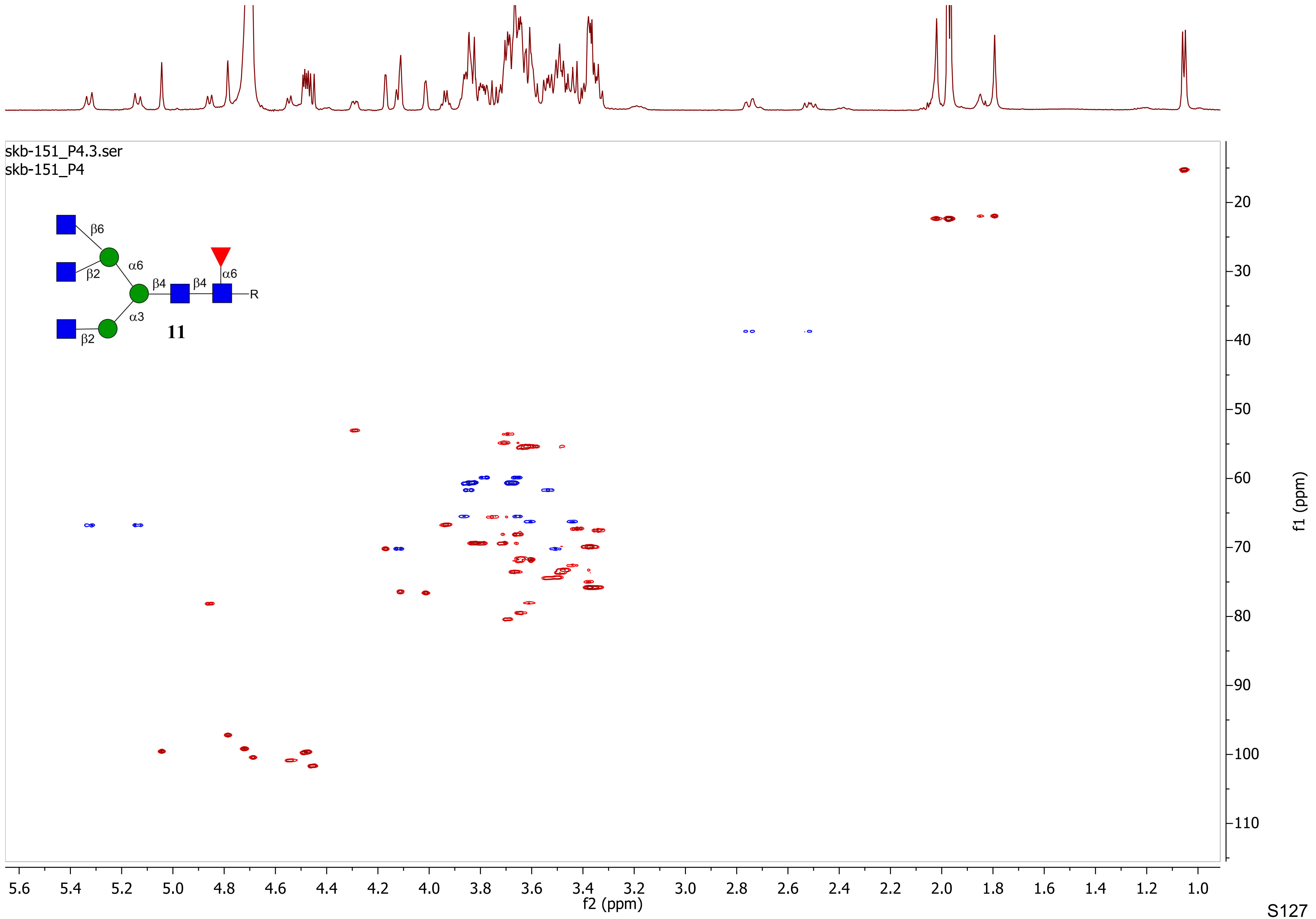
skb-151_P4.3.ser
skb-151_P4
β6
β2
α6
β4
β4
α6
R
α3
β2
11
f2 (ppm)
f1 (ppm)

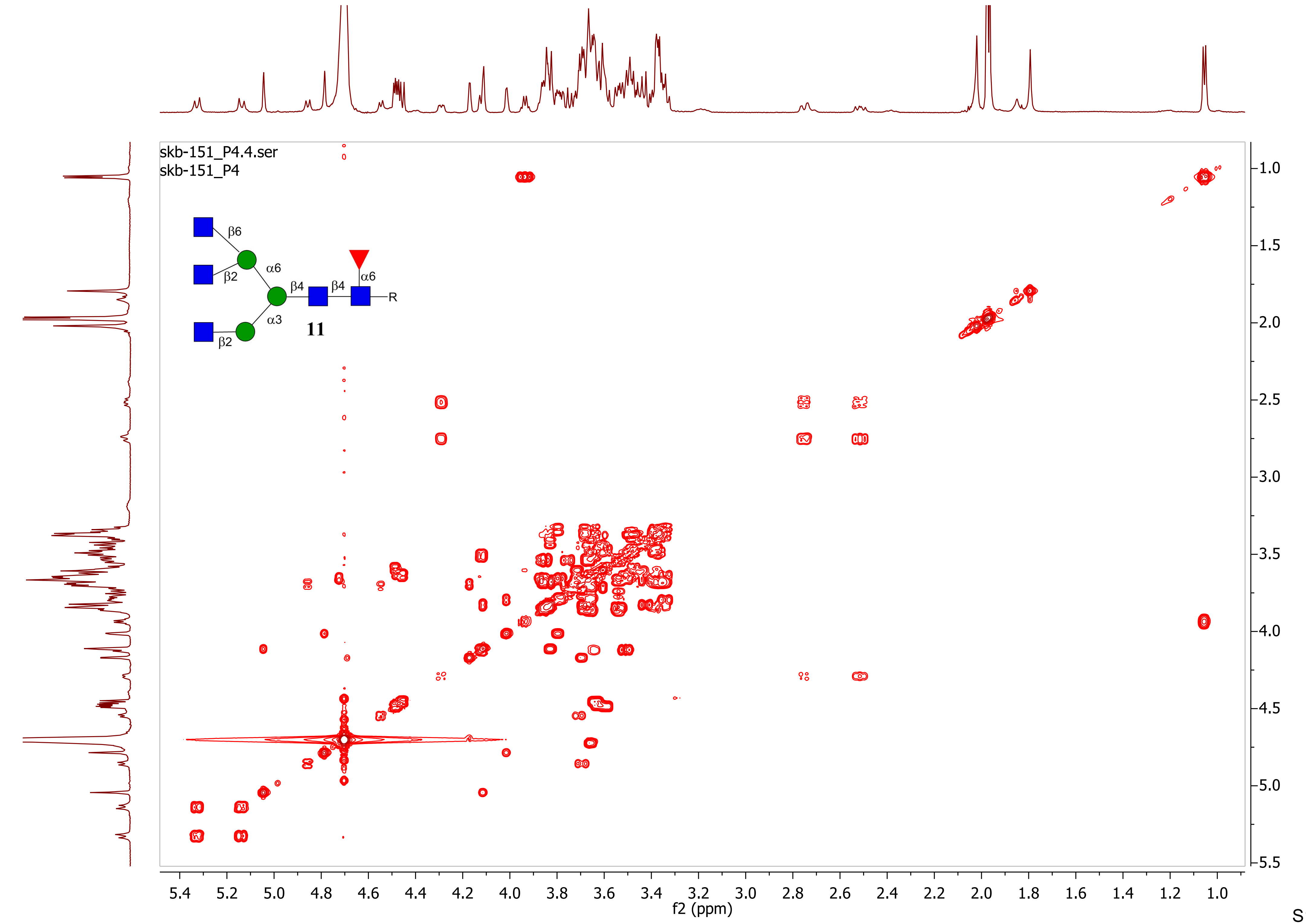
skb-151_P4.4.ser
skb-151_P4
β6
α6
β2
β4
β4
α6
R
α3
β2
11
f2 (ppm)
f1 (ppm)

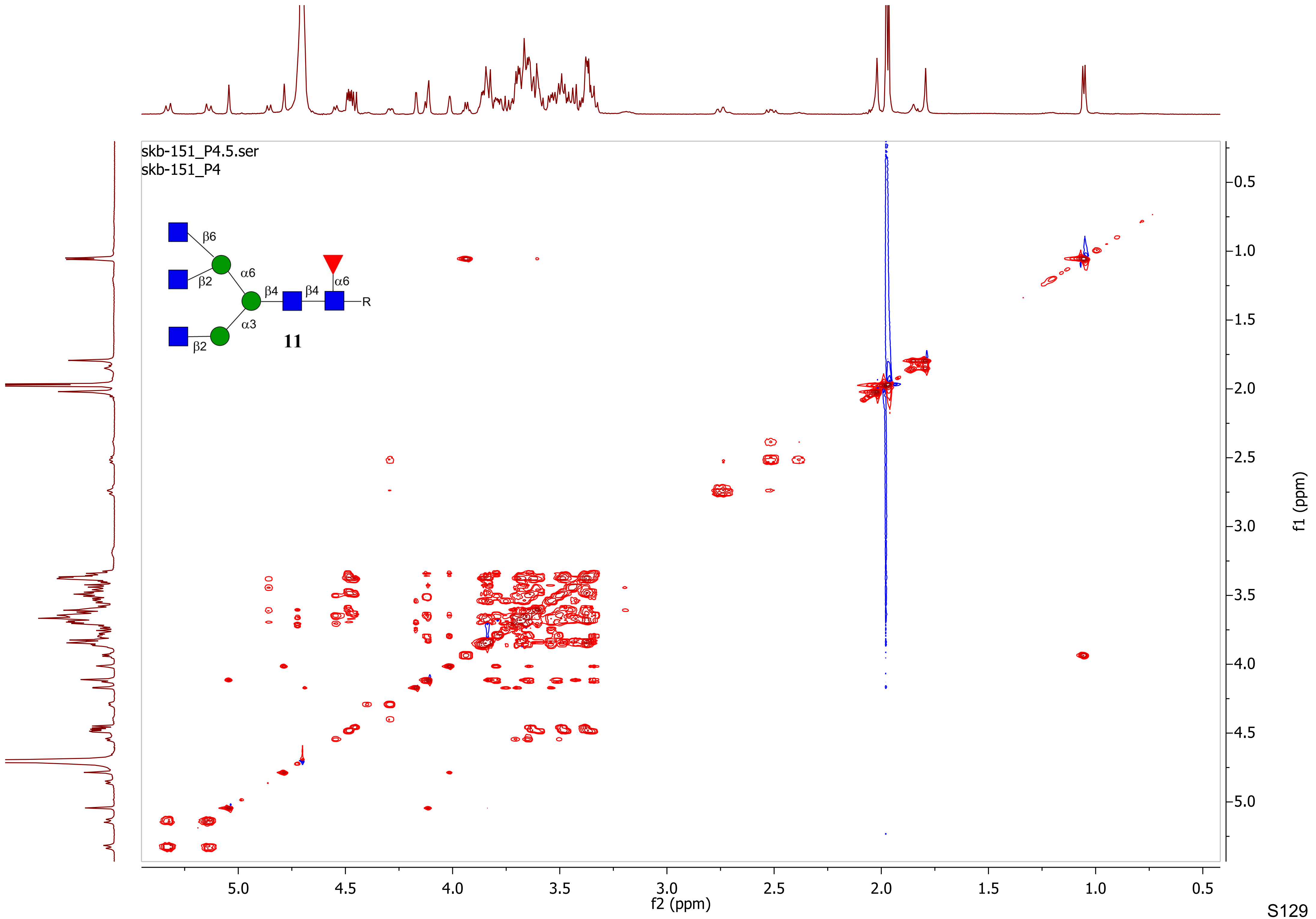
skb-151_P4.5.ser
skb-151_P4
β6
α6
β2
β4
β4
α6
R
α3
β2
11
f2 (ppm)
f1 (ppm)

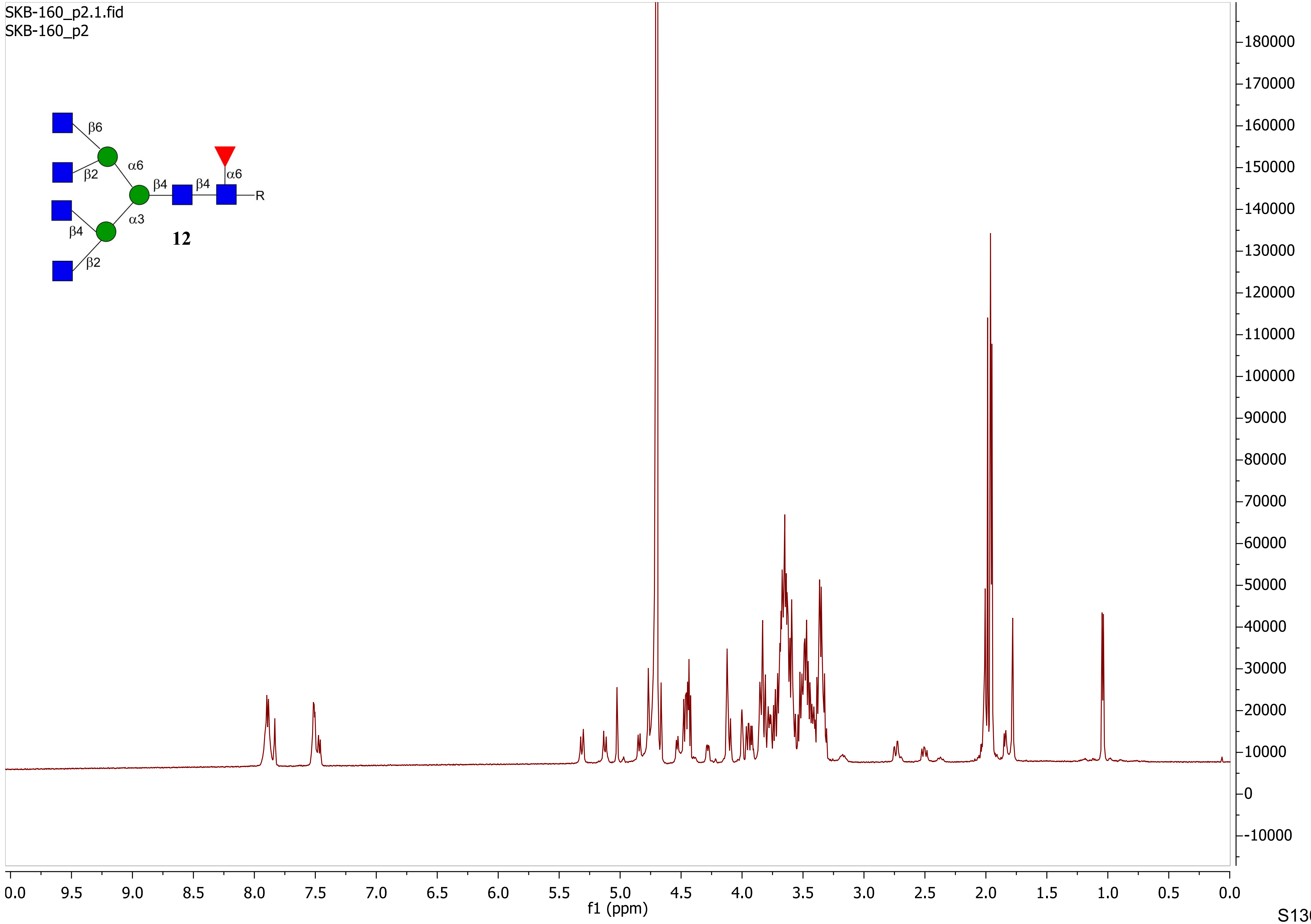
SKB-160_p2.1.fid
SKB-160_p2
β6
β2
α6
β4
β4
α6
R
β4
α3
β2
12
f1 (ppm)
0.0
9.5
9.0
8.5
8.0
7.5
7.0
6.5
6.0
5.5
5.0
4.5
4.0
3.5
3.0
2.5
2.0
1.5
1.0
0.5
0.0
180000
170000
160000
150000
140000
130000
120000
110000
100000
90000
80000
70000
60000
50000
40000
30000
20000
10000
0
-10000

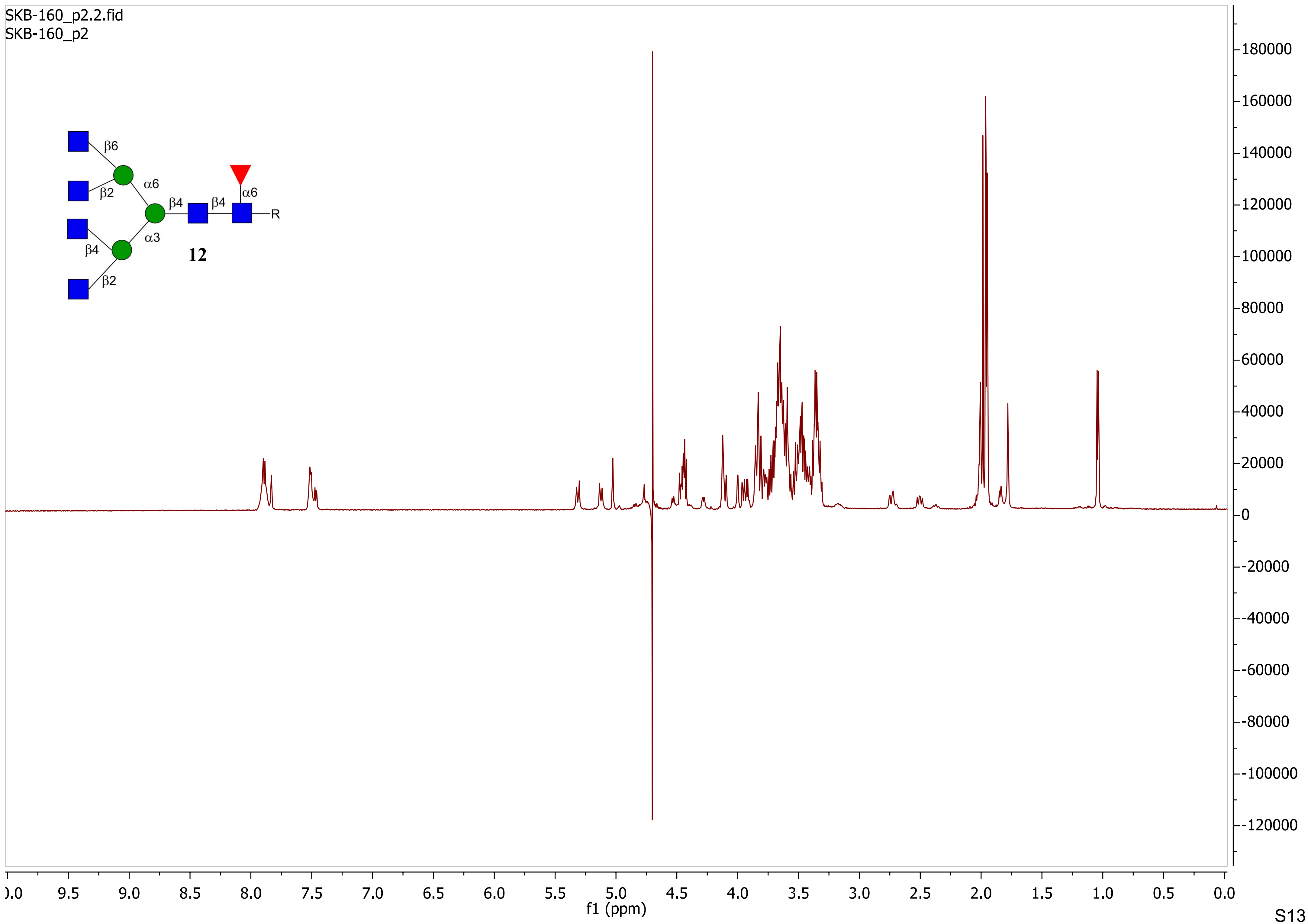
SKB-160_p2.2.fid
SKB-160_p2
β6
β2
α6
β4
β4
α6
R
β4
α3
β2
12
180000
160000
140000
120000
100000
80000
60000
40000
20000
0
-20000
-40000
-60000
-80000
-100000
-120000
9.5
9.0
8.5
8.0
7.5
7.0
6.5
6.0
5.5
5.0
4.5
4.0
3.5
3.0
2.5
2.0
1.5
1.0
0.5
0.0
f1 (ppm)

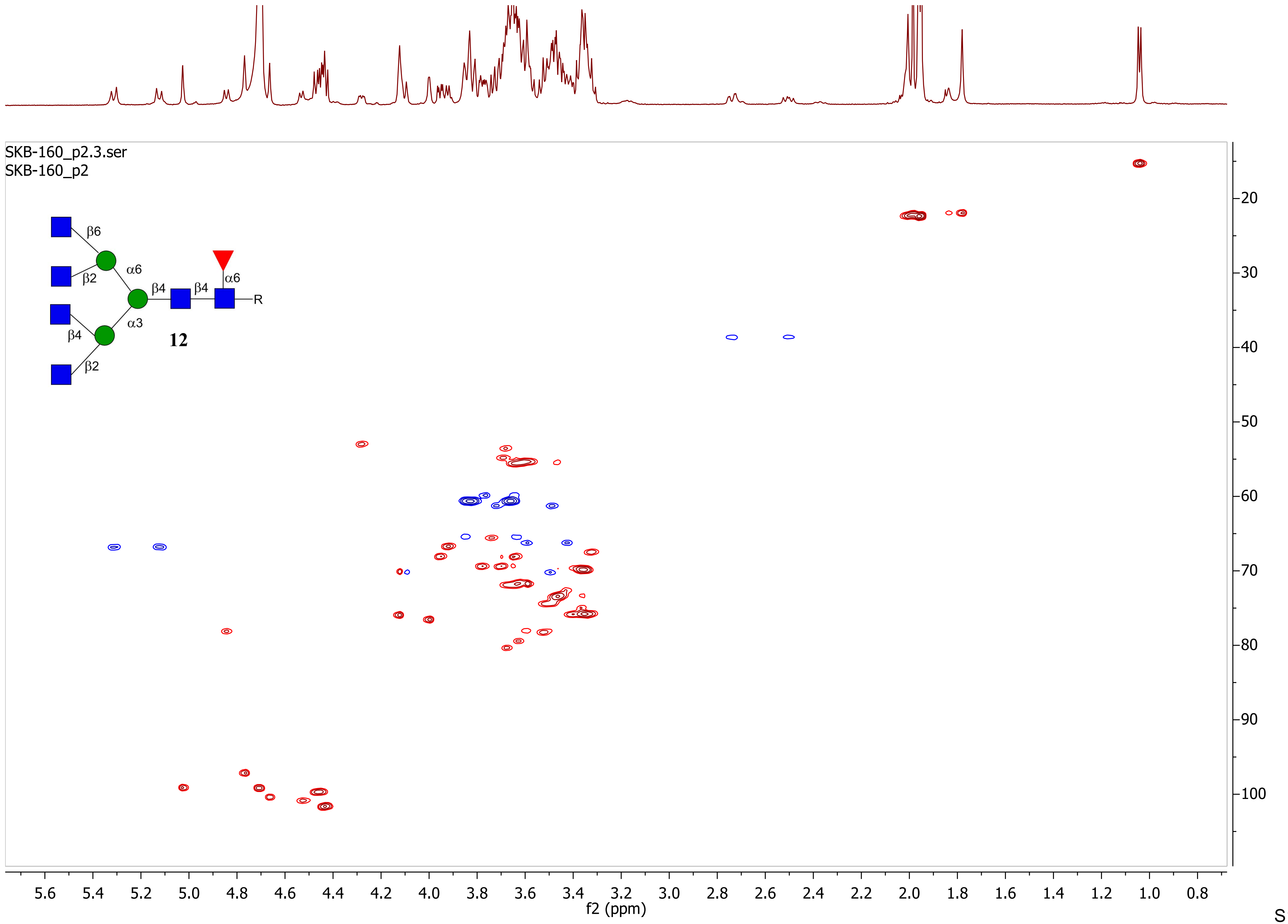

SKB-160_p2.3.ser
SKB-160_p2
β6
β2
α6
β4
β4
α6
R
β4
α3
β2
12
5.6 5.4 5.2 5.0 4.8 4.6 4.4 4.2 4.0 3.8 3.6 3.4 3.2 3.0 2.8 2.6 2.4 2.2 2.0 1.8 1.6 1.4 1.2 1.0 0.8
f2 (ppm)
20 30 40 50 60 70 80 90 100
f1 (ppm)

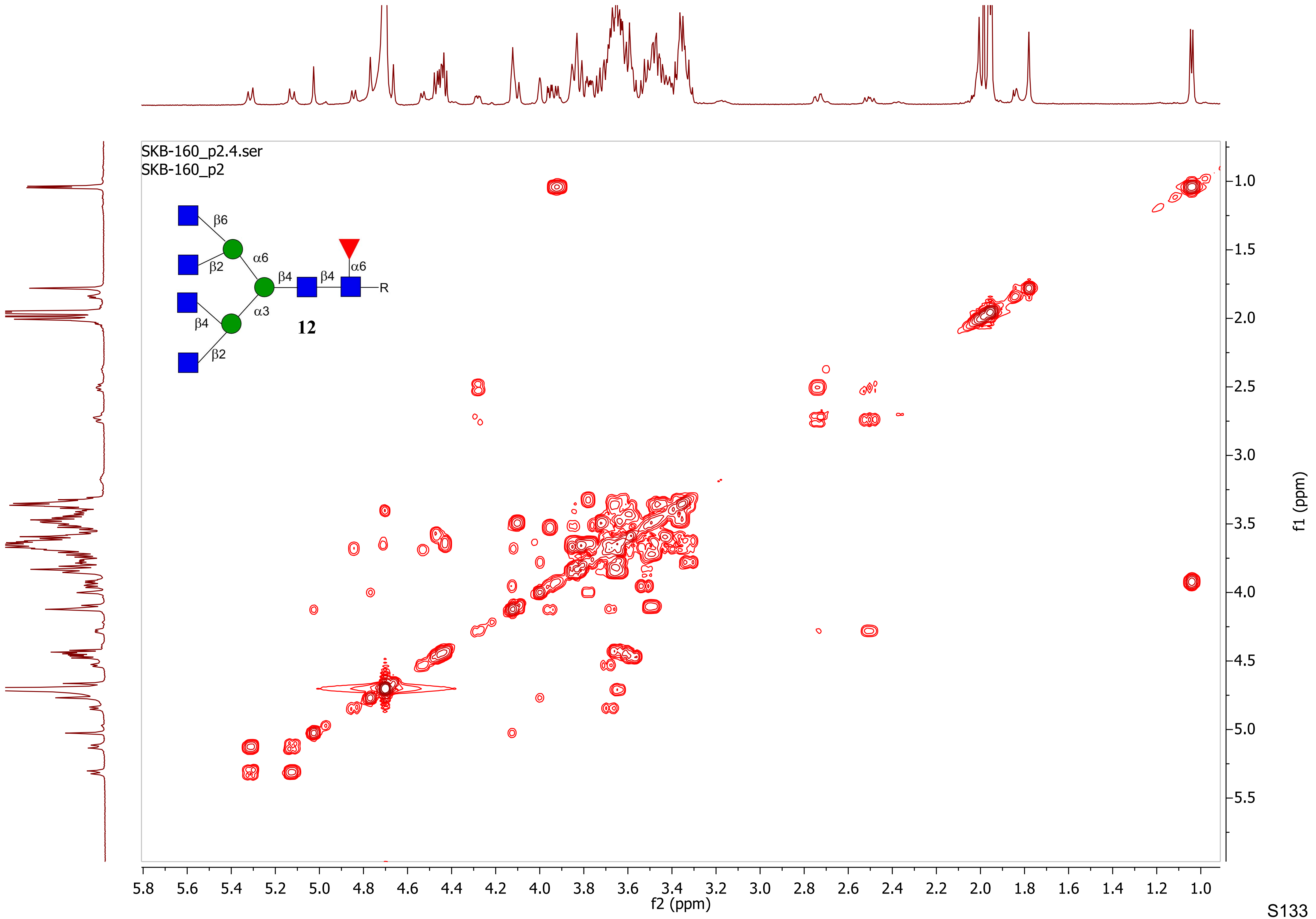
SKB-160_p2.4.ser
SKB-160_p2
β6
β2
α6
β4
β4
α6
R
β4
α3
12
β2
f2 (ppm)
f1 (ppm)

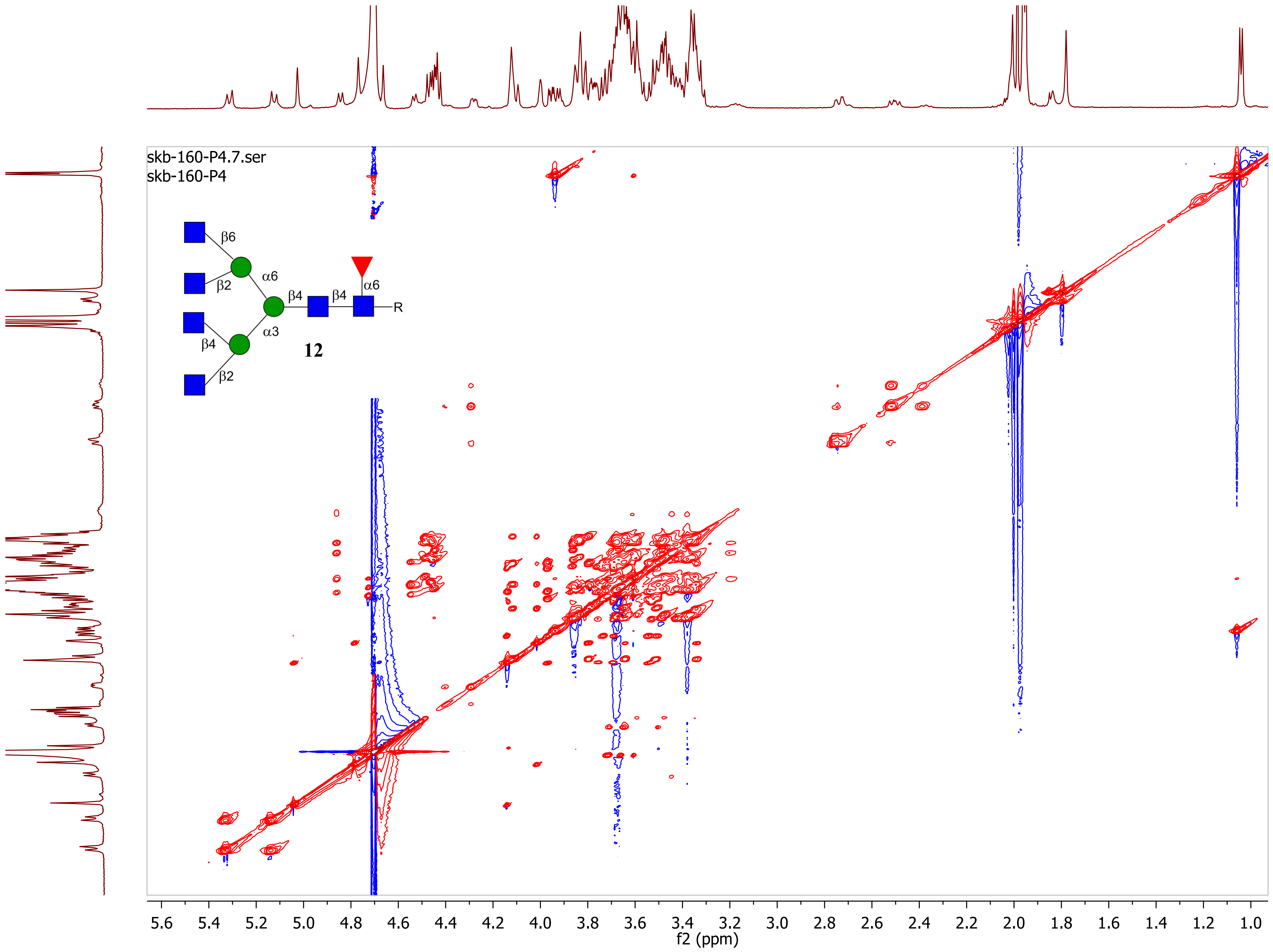
skb-160-P4.7.ser
skb-160-P4
β6
α6
β2
β4
β4
α6
R
β4
α3
12
β2
f2 (ppm)
f1 (ppm)
5.6
5.4
5.2
5.0
4.8
4.6
4.4
4.2
4.0
3.8
3.6
3.4
3.2
3.0
2.8
2.6
2.4
2.2
2.0
1.8
1.6
1.4
1.2
1.0
1.5
2.0
2.5
3.0
3.5
4.0
4.5
5.0
5.5

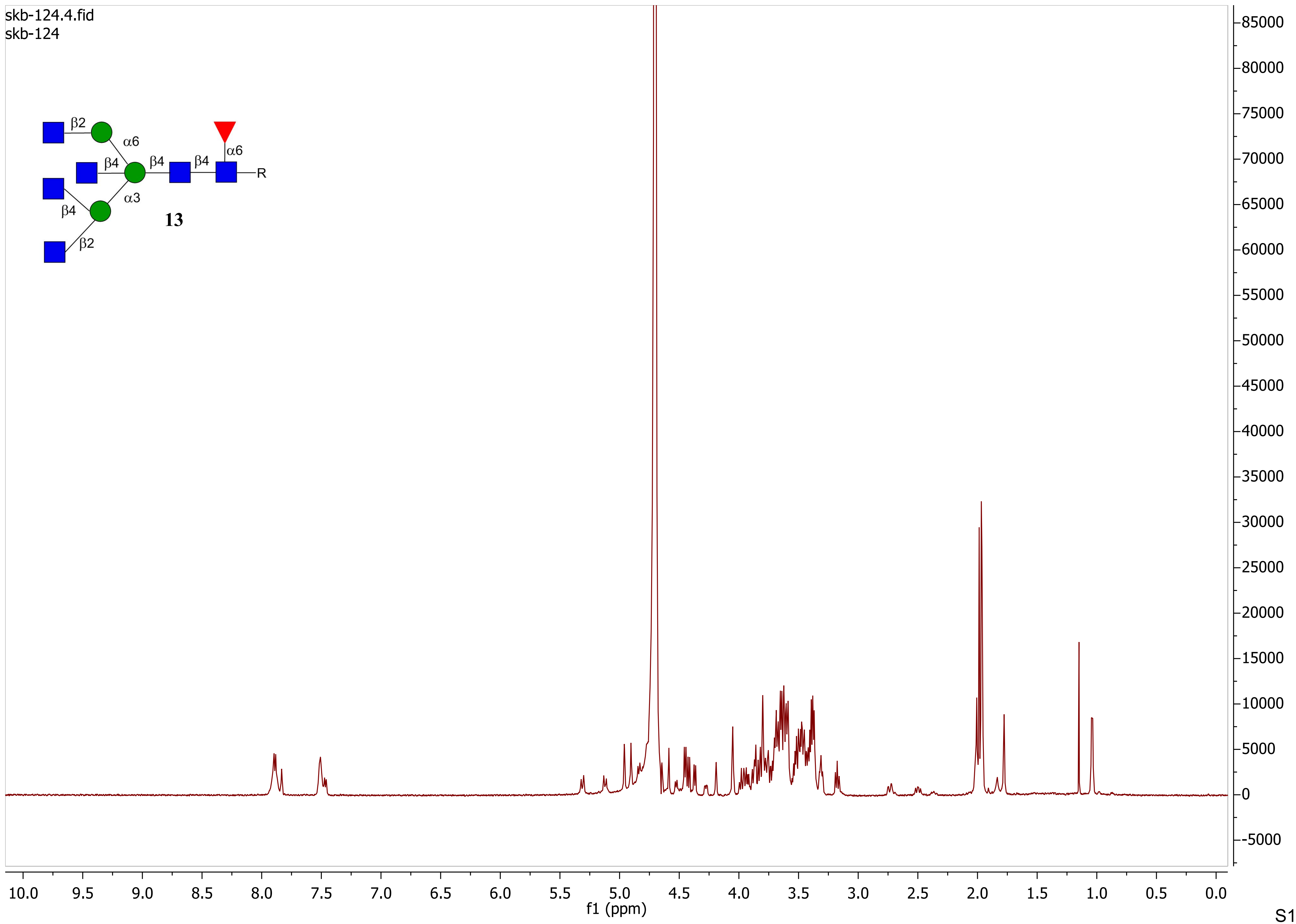

skb-124.4.fid
skb-124
β2
α6
β4
β4
β4
α6
R
α3
β4
β2
13
85000
80000
75000
70000
65000
60000
55000
50000
45000
40000
35000
30000
25000
20000
15000
10000
5000
0
-5000
10.0
9.5
9.0
8.5
8.0
7.5
7.0
6.5
6.0
5.5
5.0
4.5
4.0
3.5
3.0
2.5
2.0
1.5
1.0
0.5
0.0
f1 (ppm)

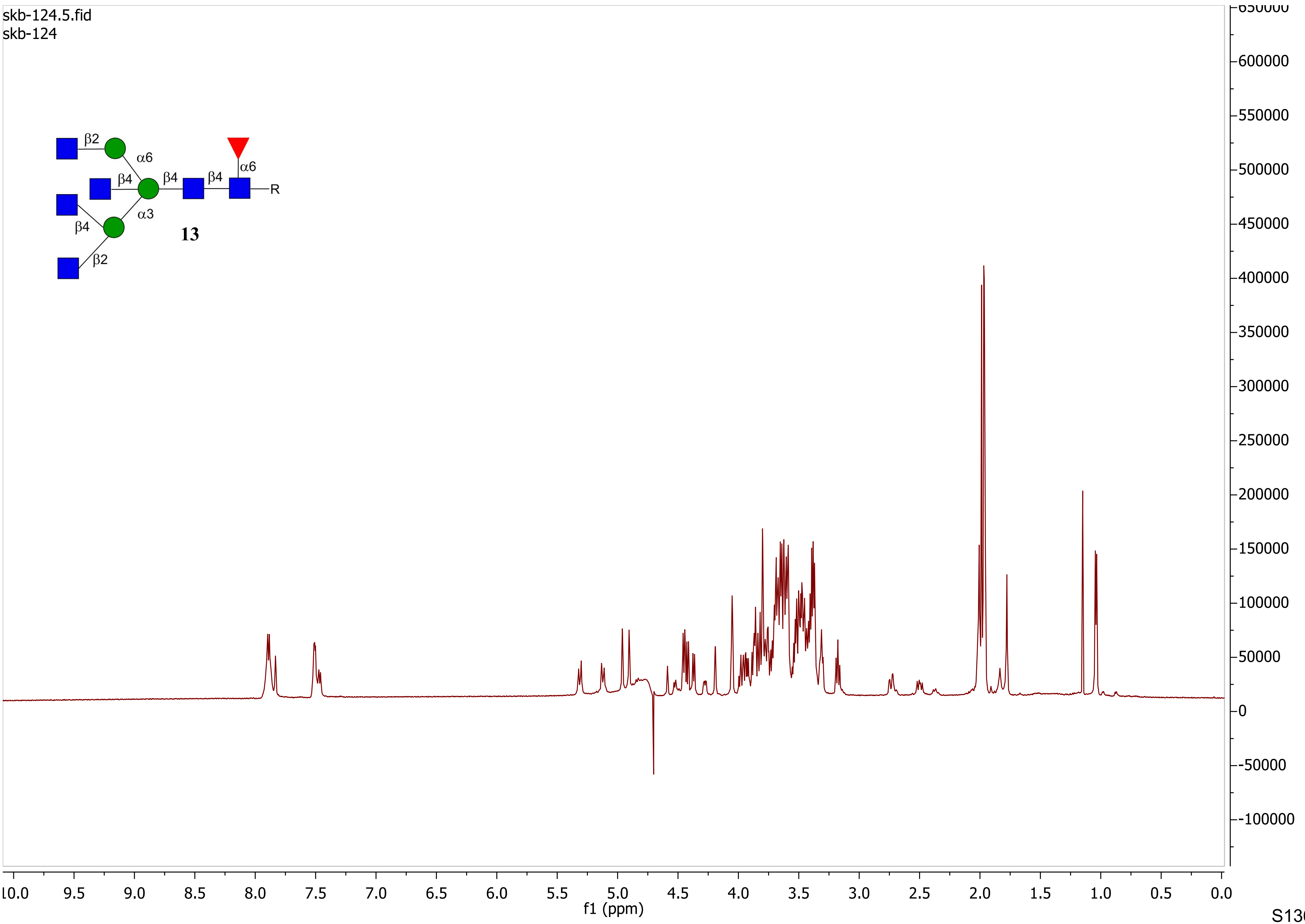
skb-124.5.fid
skb-124
β2
α6
β4
β4
β4
α6
R
α3
β4
β2
13
f1 (ppm)

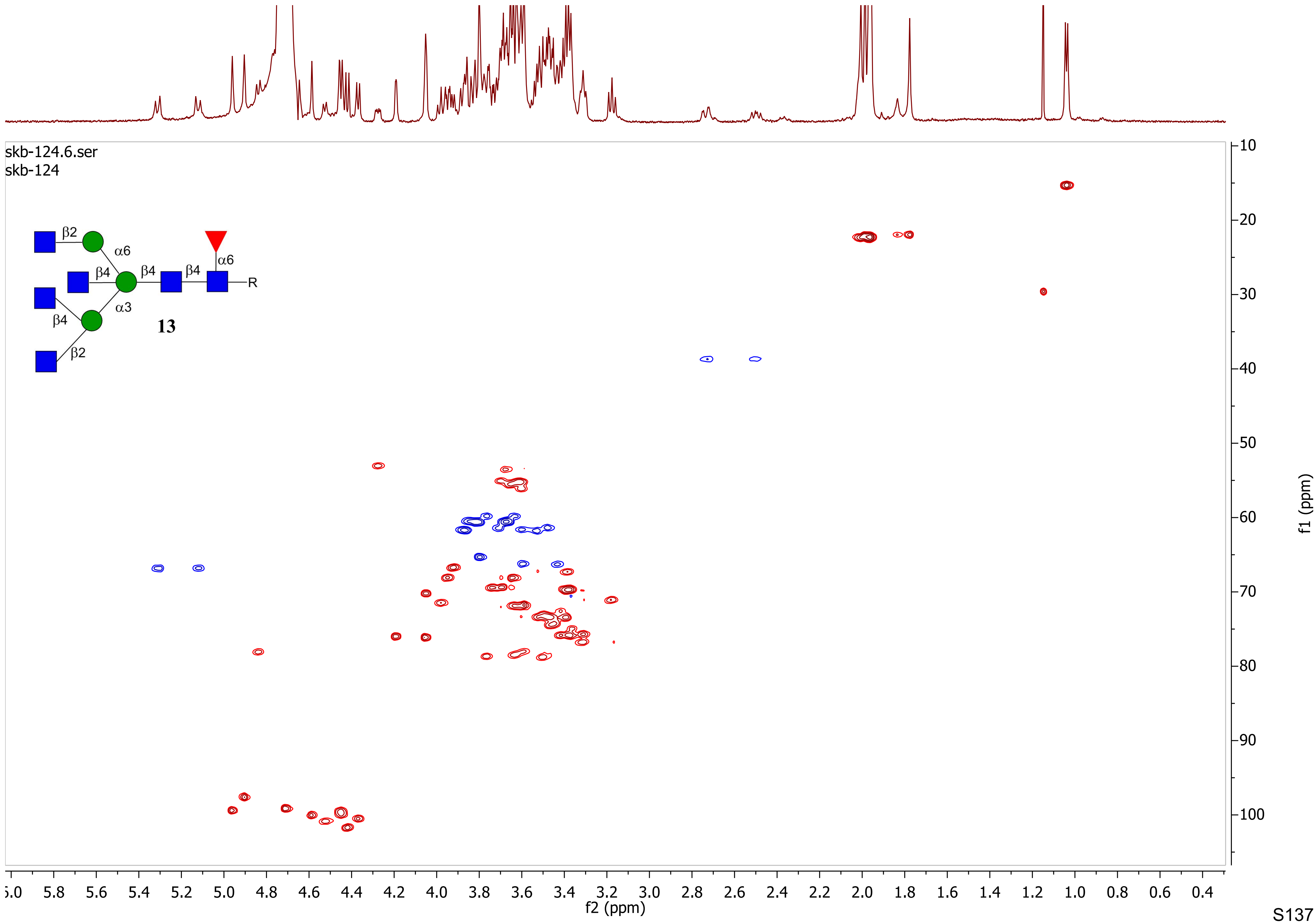
skb-124.6.ser
skb-124
β2
α6
β4
β4
β4
α6
R
α3
β4
β2
13
f2 (ppm)
f1 (ppm)

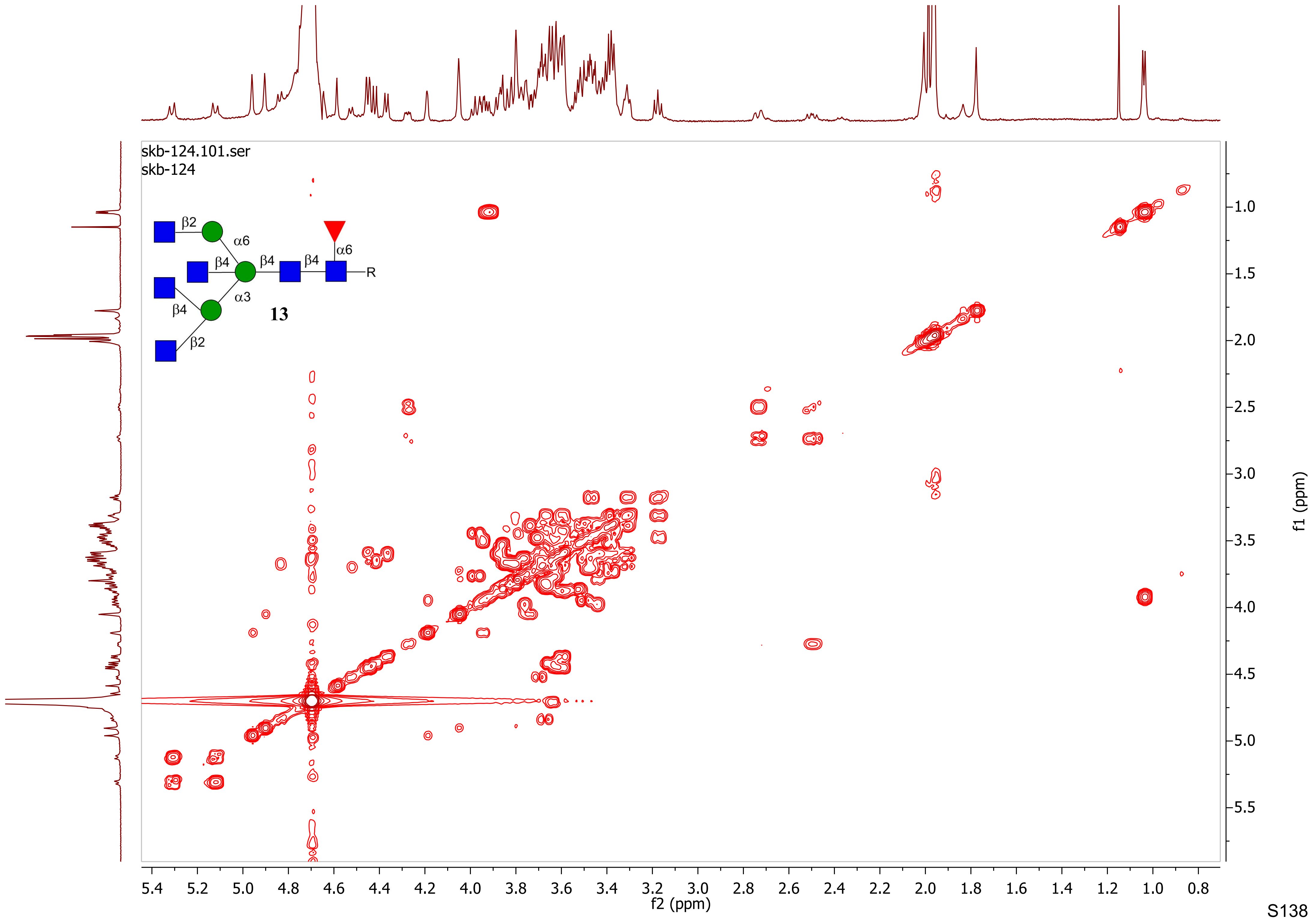
skb-124.101.ser
skb-124
β2
α6
β4
β4
β4
α6
R
α3
β4
β2
13
f2 (ppm)
f1 (ppm)

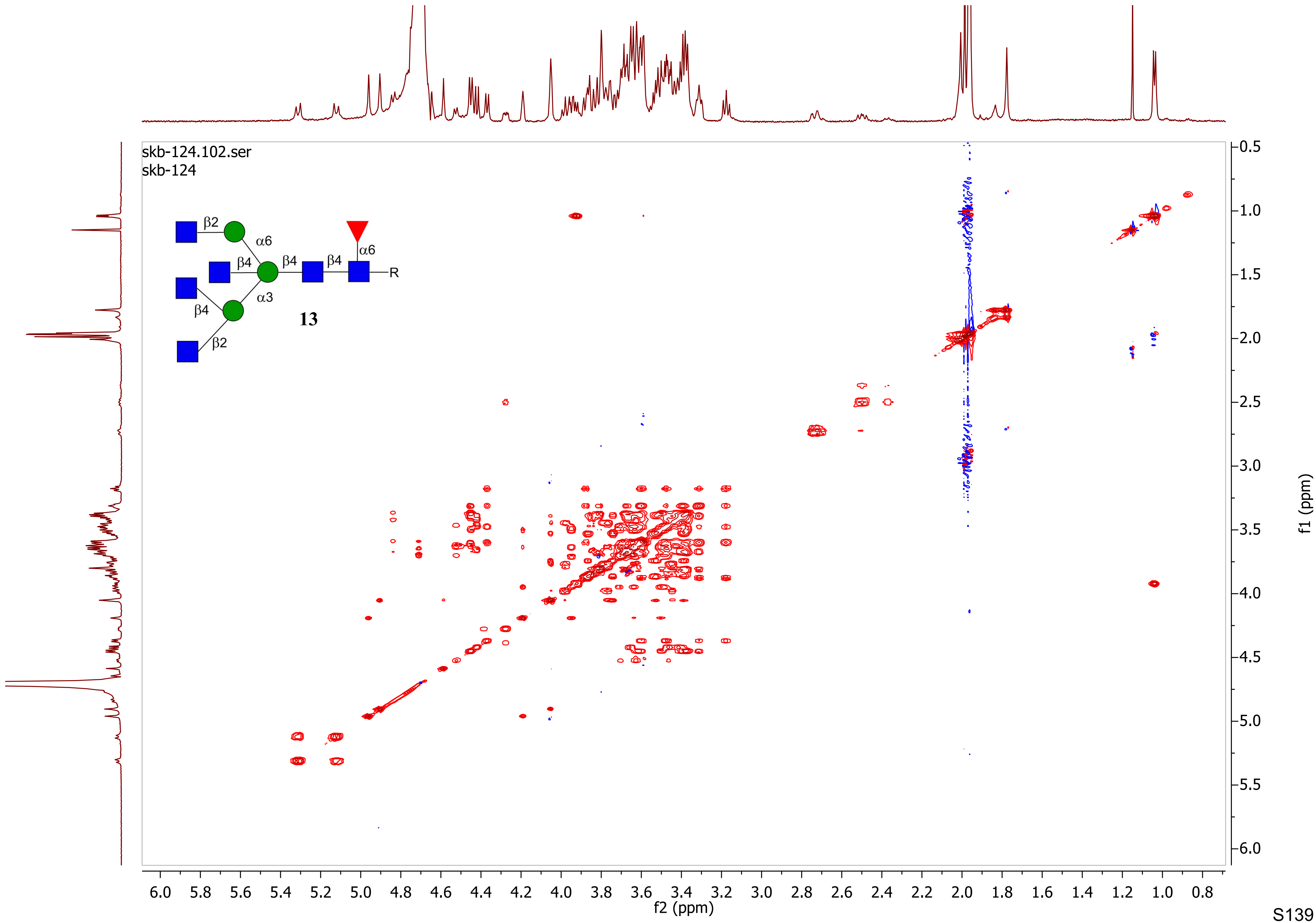

skb-124.102.ser
skb-124
β2
α6
β4
β4
β4
α6
R
β4
α3
β2
13
f2 (ppm)
f1 (ppm)

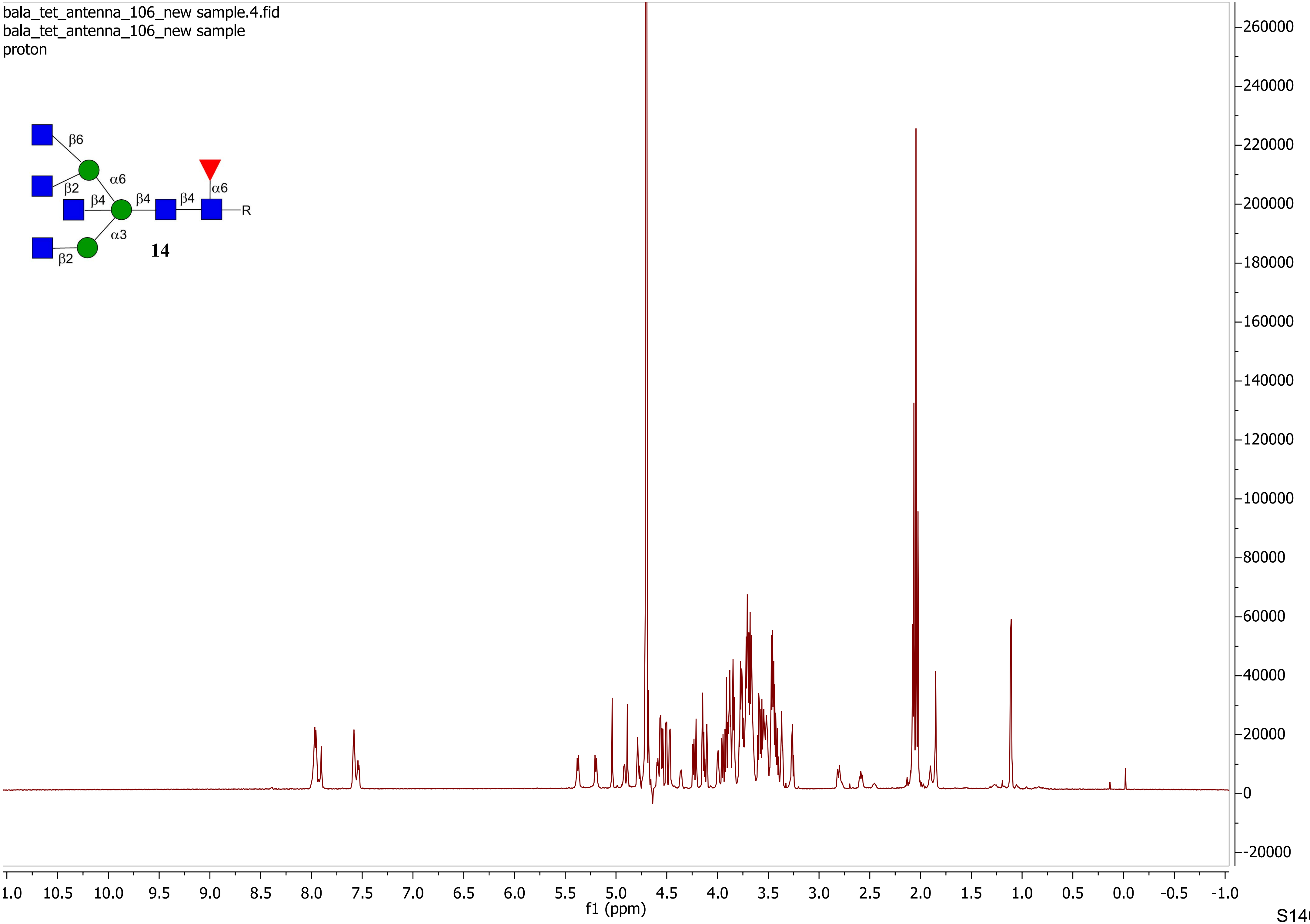
bala_tet_antenna_106_new sample.4.fid
bala_tet_antenna_106_new sample
proton
β6
β2
α6
β4
β4
β4
α6
R
α3
β2
14
f1 (ppm)

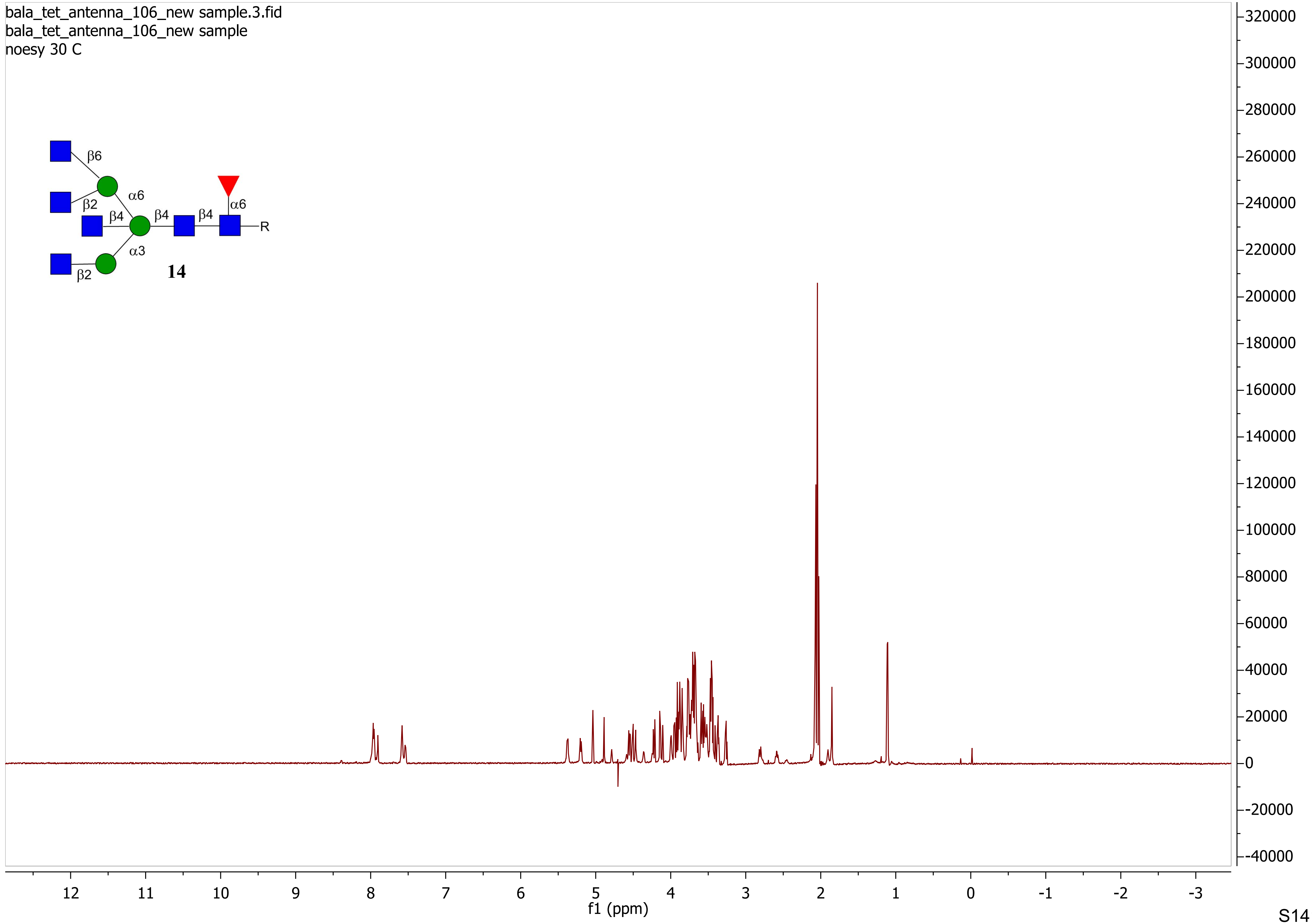

bala_tet_antenna_106_new sample.3.fid
bala_tet_antenna_106_new sample
noesy 30 C
β6
α6
β2
β4
β4
β4
α6
R
α3
β2
14
f1 (ppm)

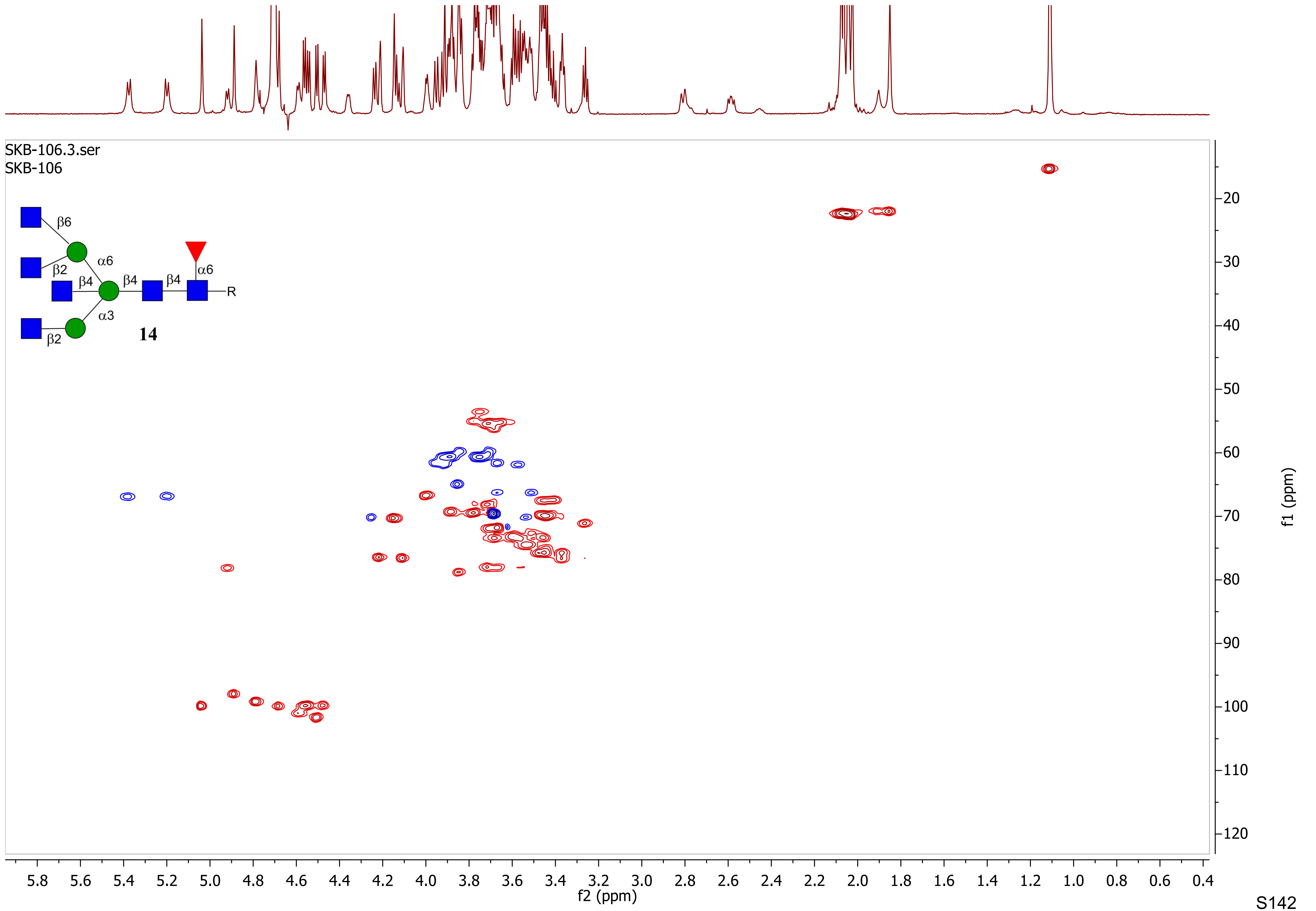
SKB-106.3.ser
SKB-106
β6
α6
β2
β4
β4
β4
α6
R
α3
β2
14
f1 (ppm)
f2 (ppm)

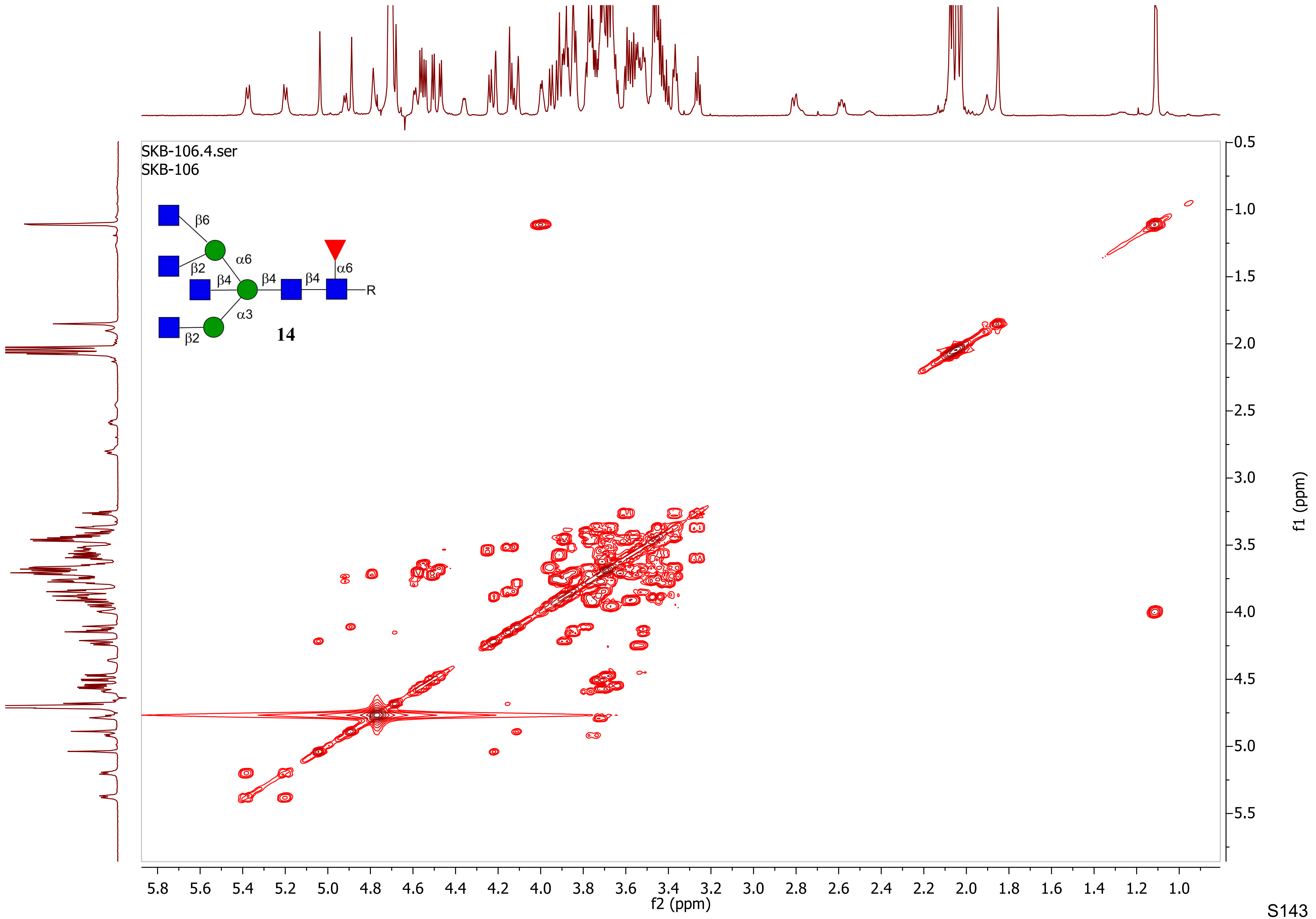

SKB-106.4.ser
SKB-106
β6
α6
β2
β4
β4
β4
α6
R
α3
β2
14
f2 (ppm)
f1 (ppm)
5.8 5.6 5.4 5.2 5.0 4.8 4.6 4.4 4.2 4.0 3.8 3.6 3.4 3.2 3.0 2.8 2.6 2.4 2.2 2.0 1.8 1.6 1.4 1.2 1.0
0.5 1.0 1.5 2.0 2.5 3.0 3.5 4.0 4.5 5.0 5.5

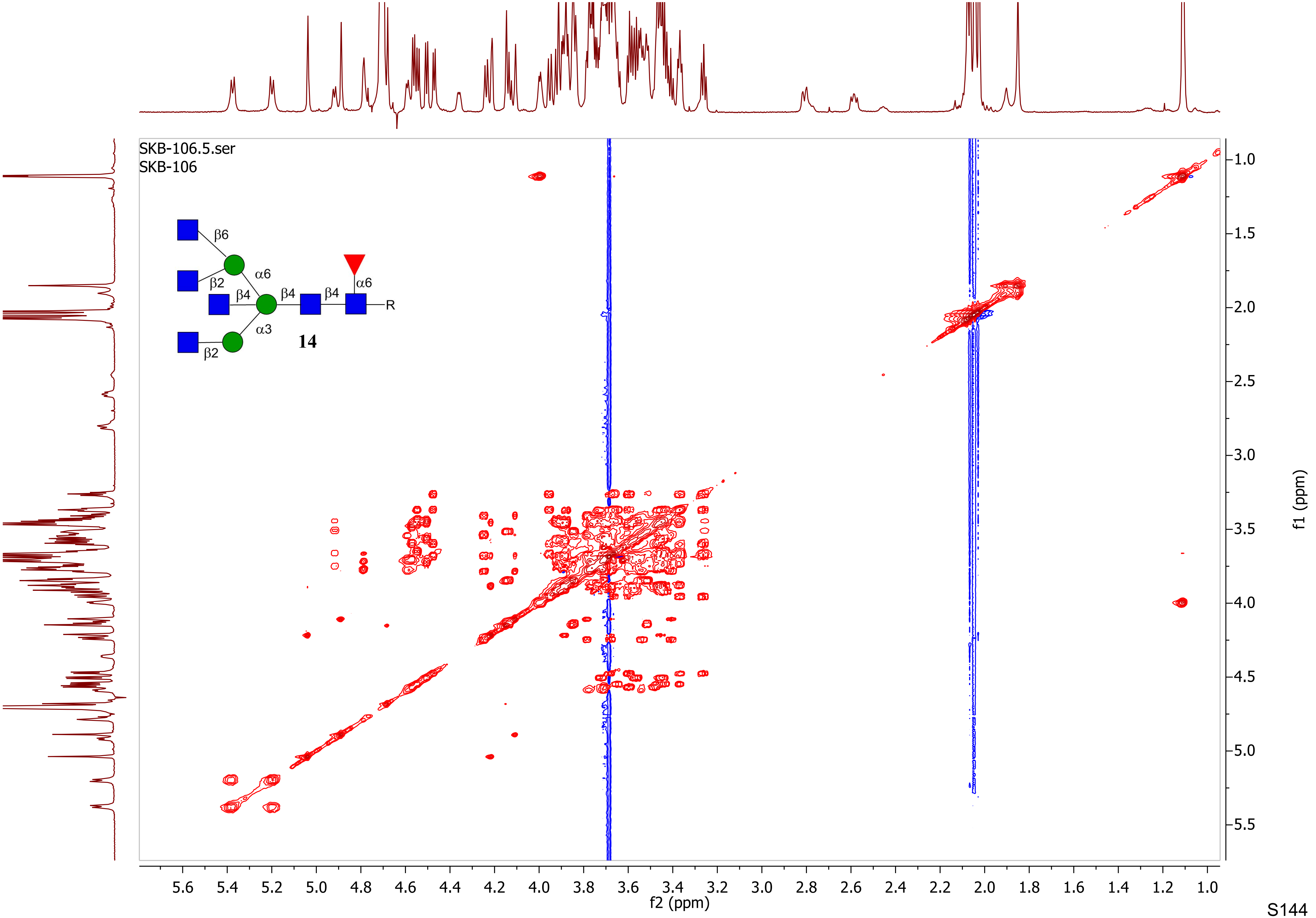

SKB-106.5.ser
SKB-106
β6
β2
α6
β4
β4
β4
α6
R
α3
β2
14
f2 (ppm)
f1 (ppm)

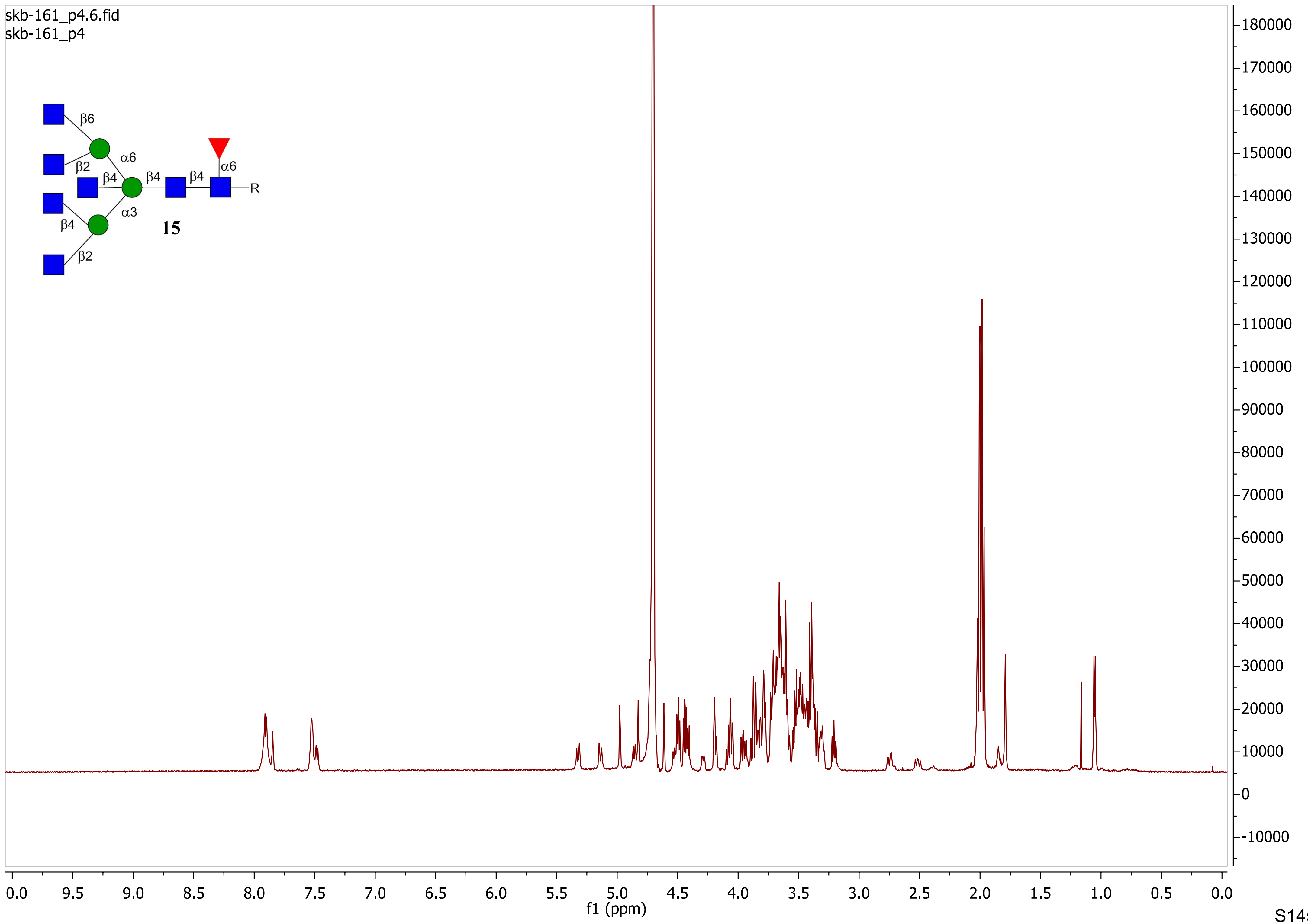

skb-161_p4.6.fid
skb-161_p4
β6
β2
α6
β4
β4
β4
α6
R
β4
α3
β2
15
f1 (ppm)

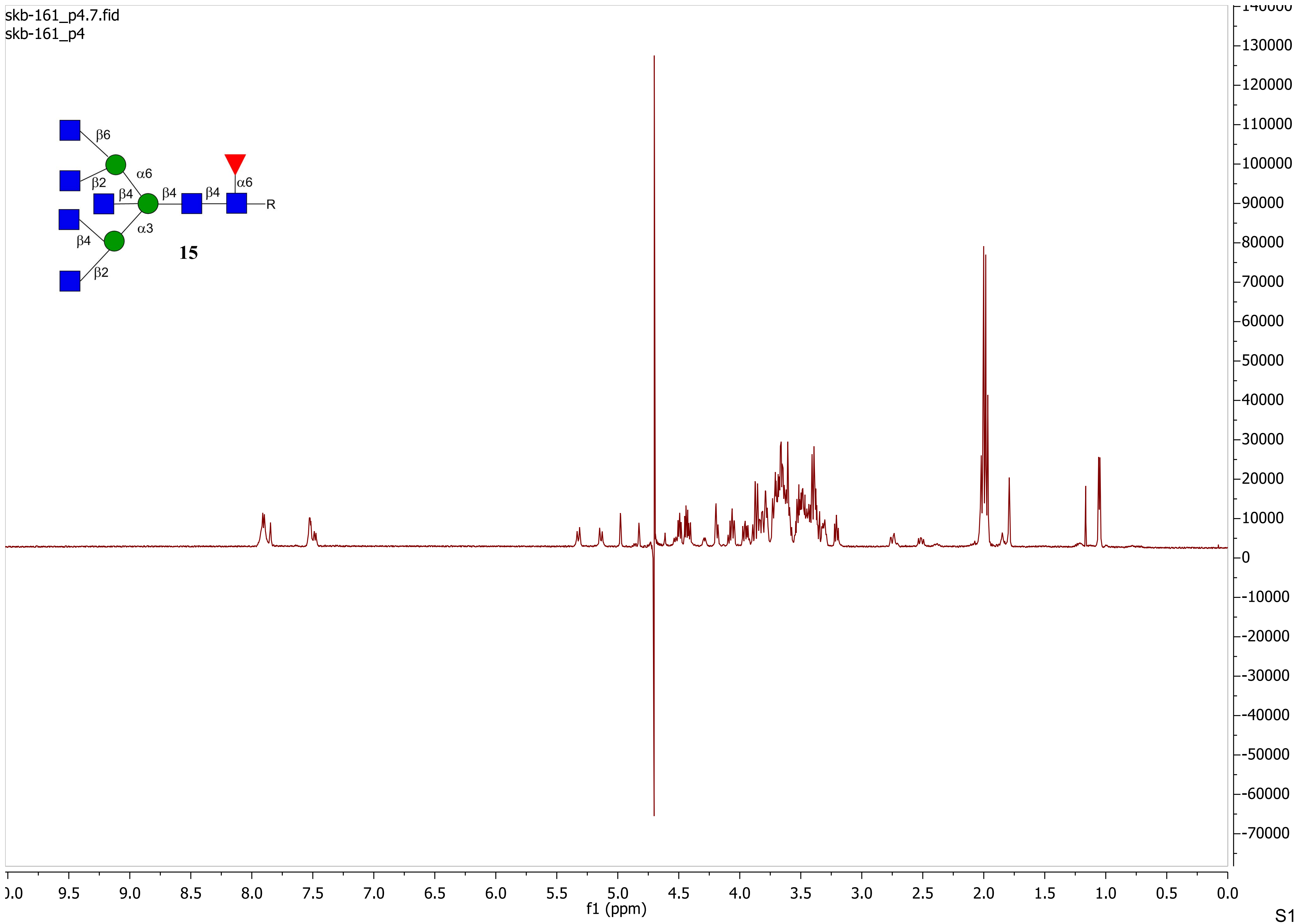
skb-161_p4.7.fid
skb-161_p4
β6
α6
β2
β4
β4
β4
α6
R
α3
β4
15
β2
f1 (ppm)

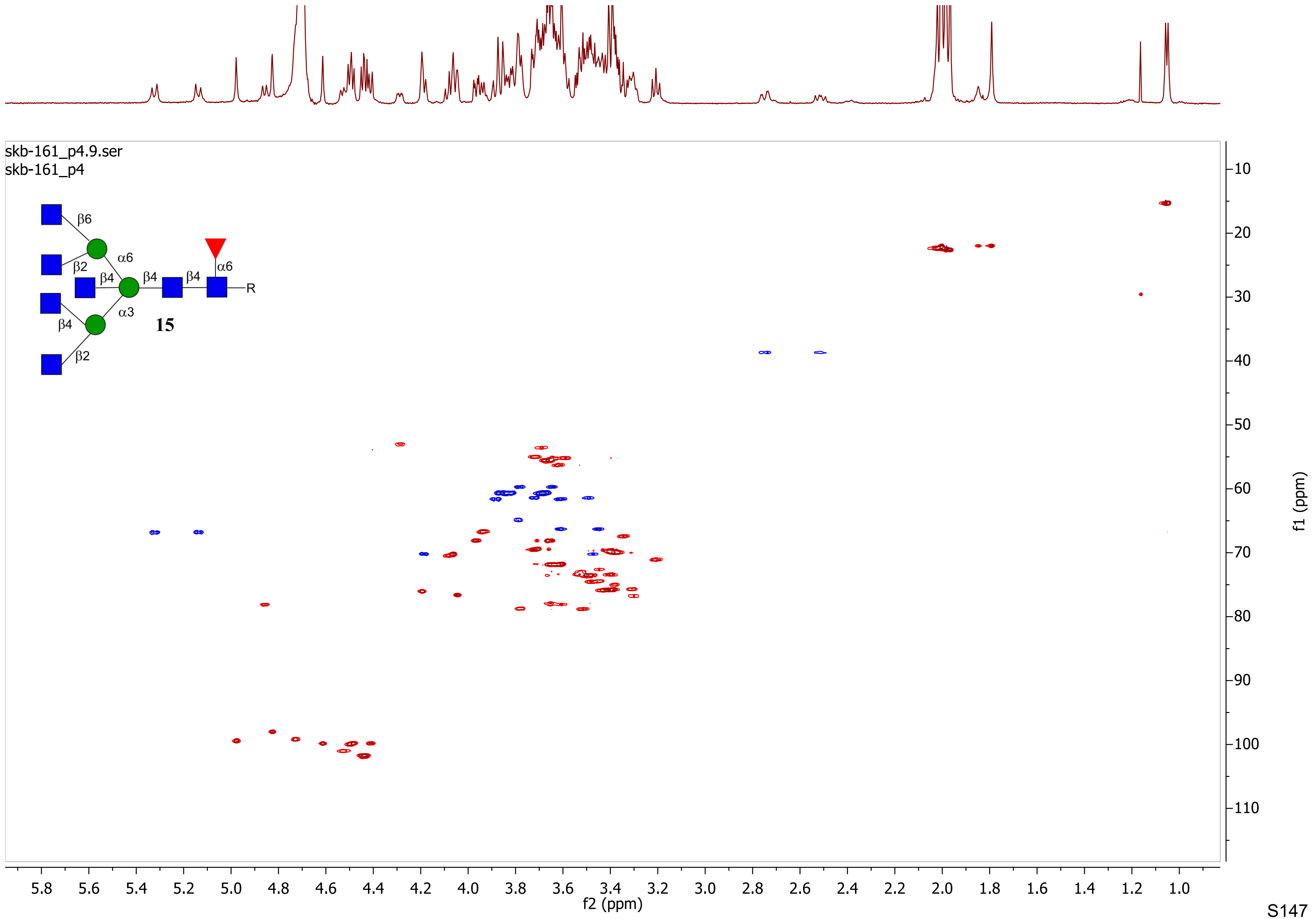
skb-161_p4.9.ser
skb-161_p4
β6
α6
β2
β4
β4
β4
α6
R
α3
β4
15
β2
f2 (ppm)
f1 (ppm)

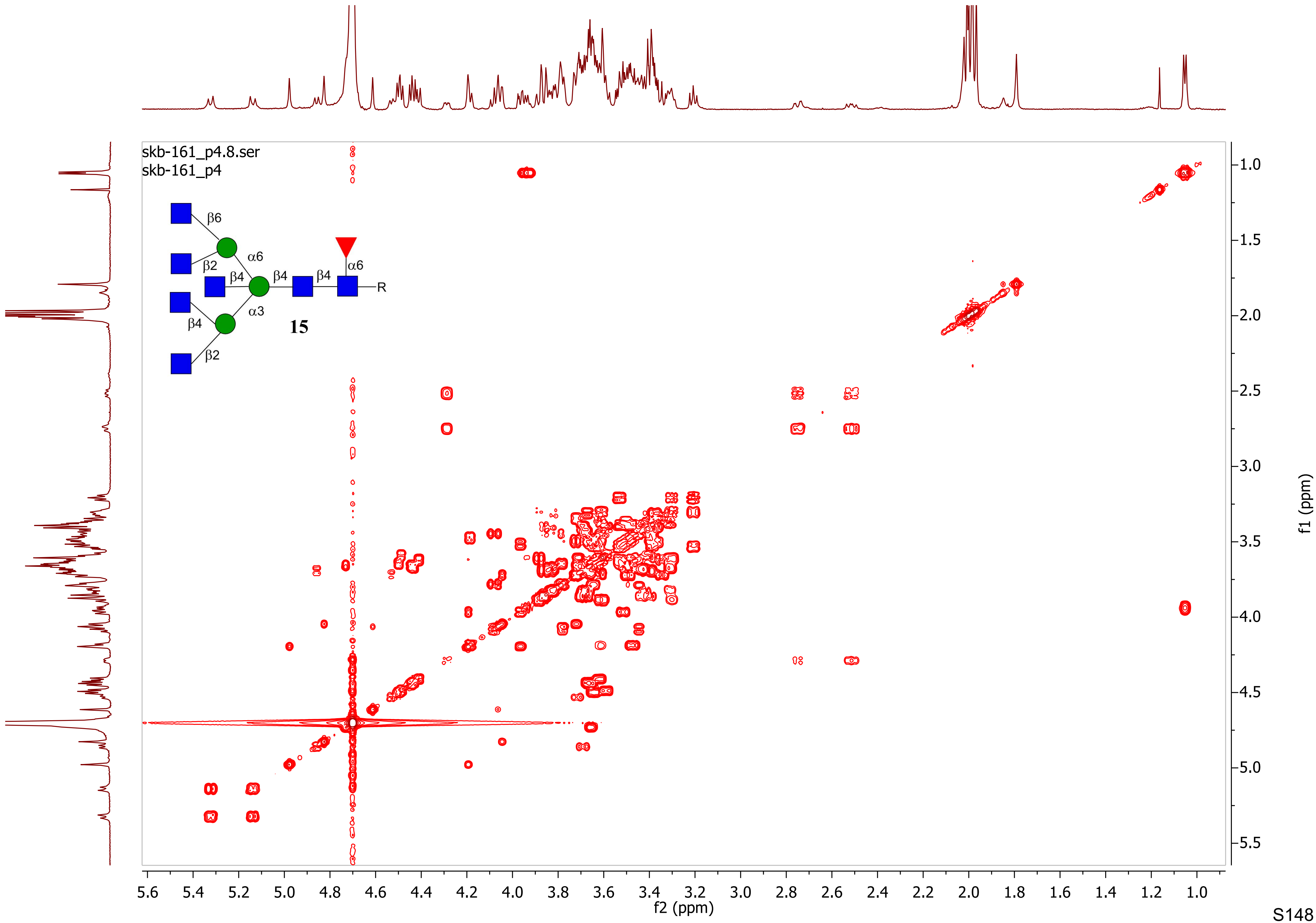

skb-161_p4.8.ser
skb-161_p4
β6
β2
α6
β4
β4
β4
α6
R
β4
α3
15
β2
f2 (ppm)
f1 (ppm)

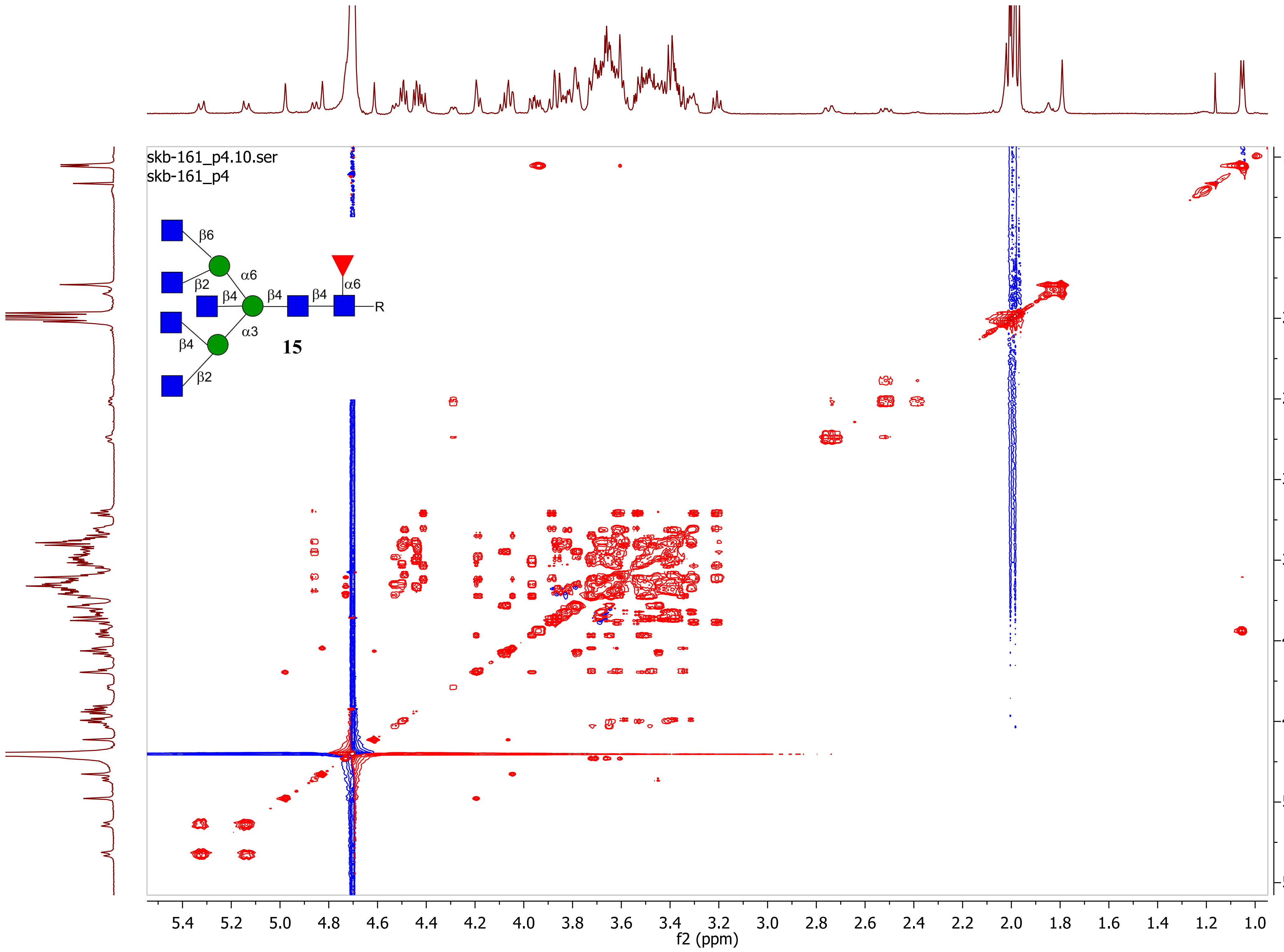
skb-161_p4.10.ser
skb-161_p4
β6
β2
α6
β4
β4
β4
α6
R
α3
15
β4
β2
f2 (ppm)
f1 (ppm)

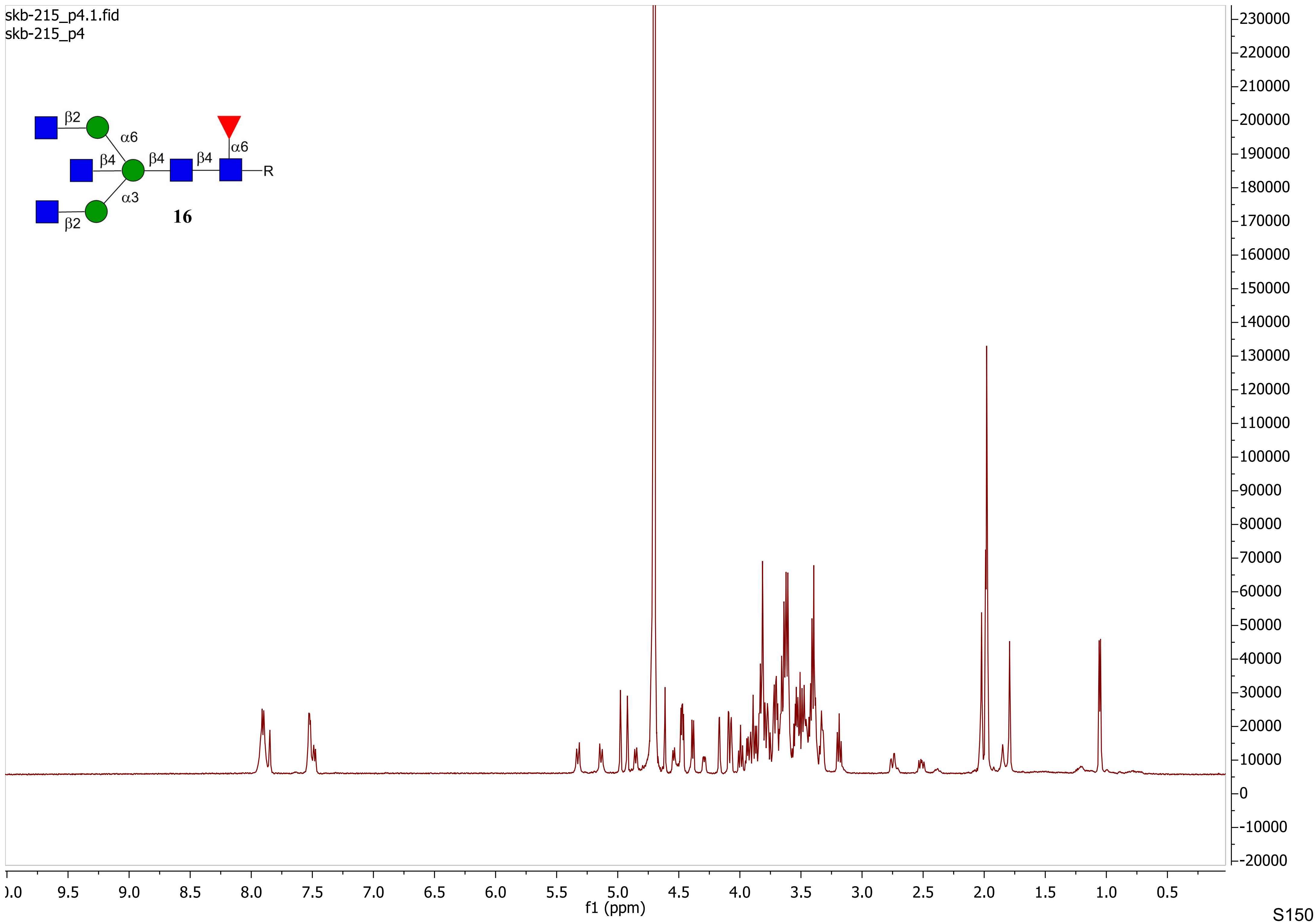

skb-215_p4.1.fid
skb-215_p4
β2
α6
β4
β4
β4
α6
R
α3
β2
16
f1 (ppm)

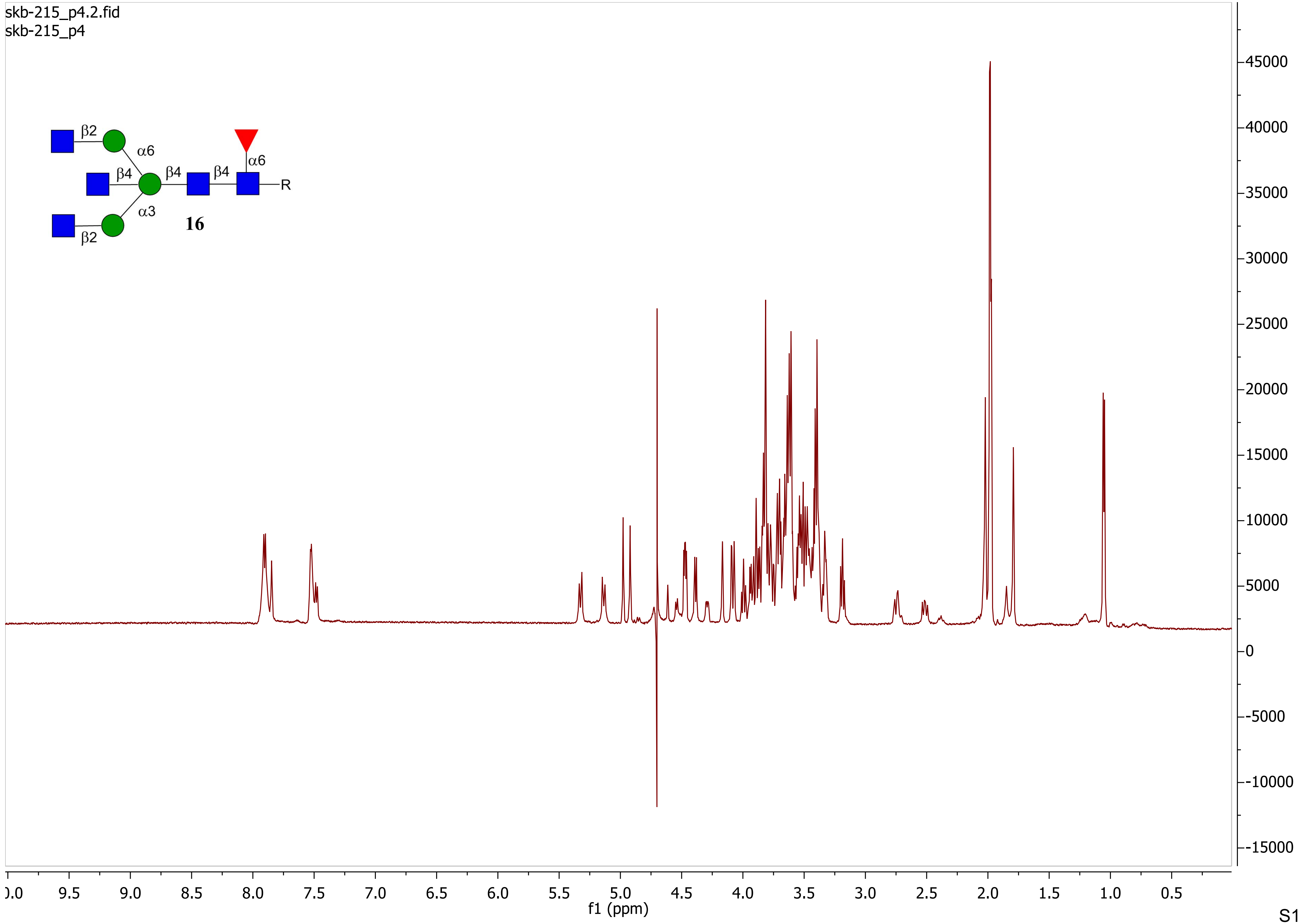

skb-215_p4.2.fid
skb-215_p4
β2
α6
β4
β4
β4
α6
R
α3
16
β2
45000
40000
35000
30000
25000
20000
15000
10000
5000
0
-5000
-10000
-15000
9.5
9.0
8.5
8.0
7.5
7.0
6.5
6.0
5.5
5.0
4.5
4.0
3.5
3.0
2.5
2.0
1.5
1.0
0.5
f1 (ppm)

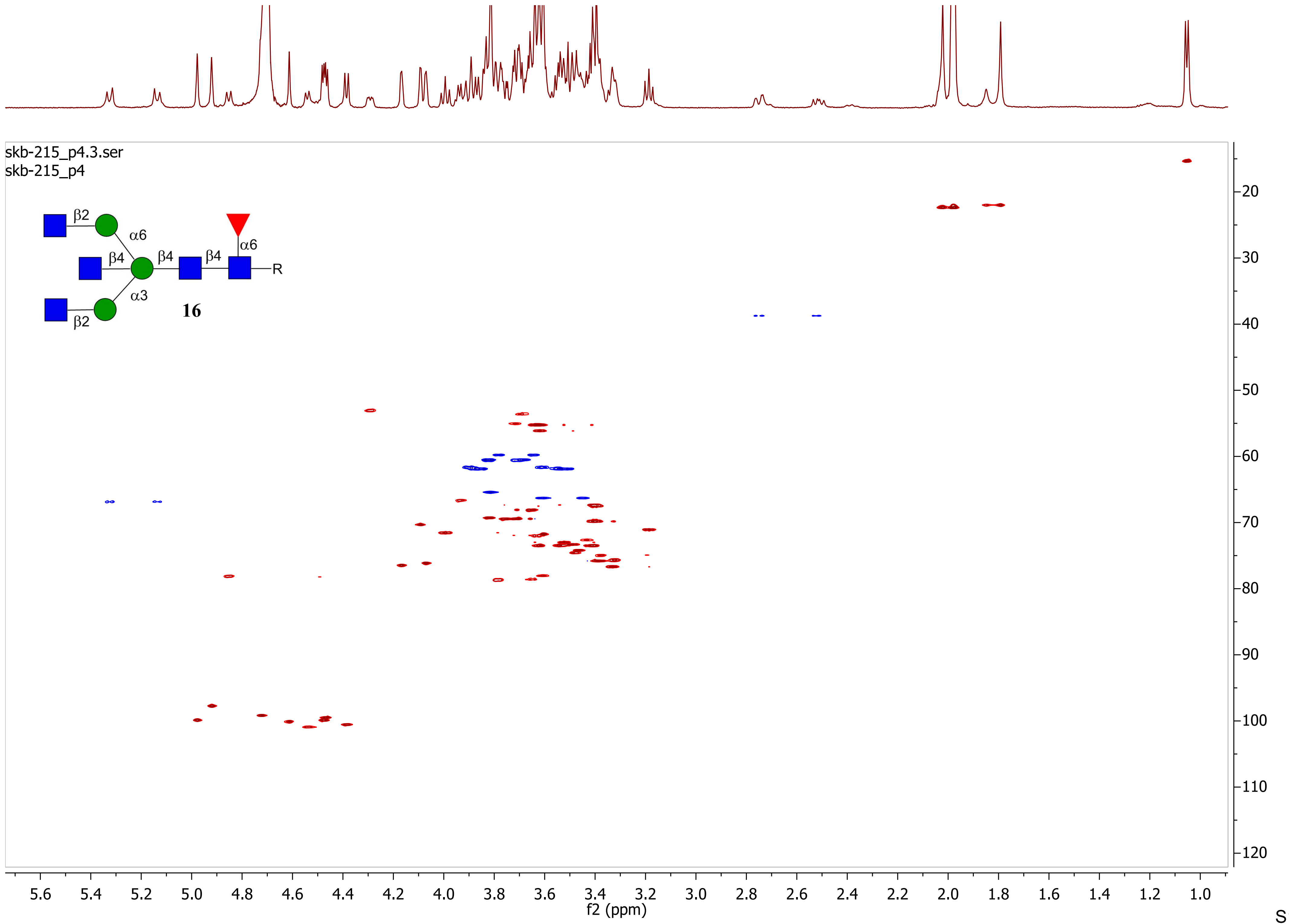

skb-215_p4.3.ser
skb-215_p4
β2
α6
β4
β4
β4
α6
R
α3
16
β2
f2 (ppm)
f1 (ppm)

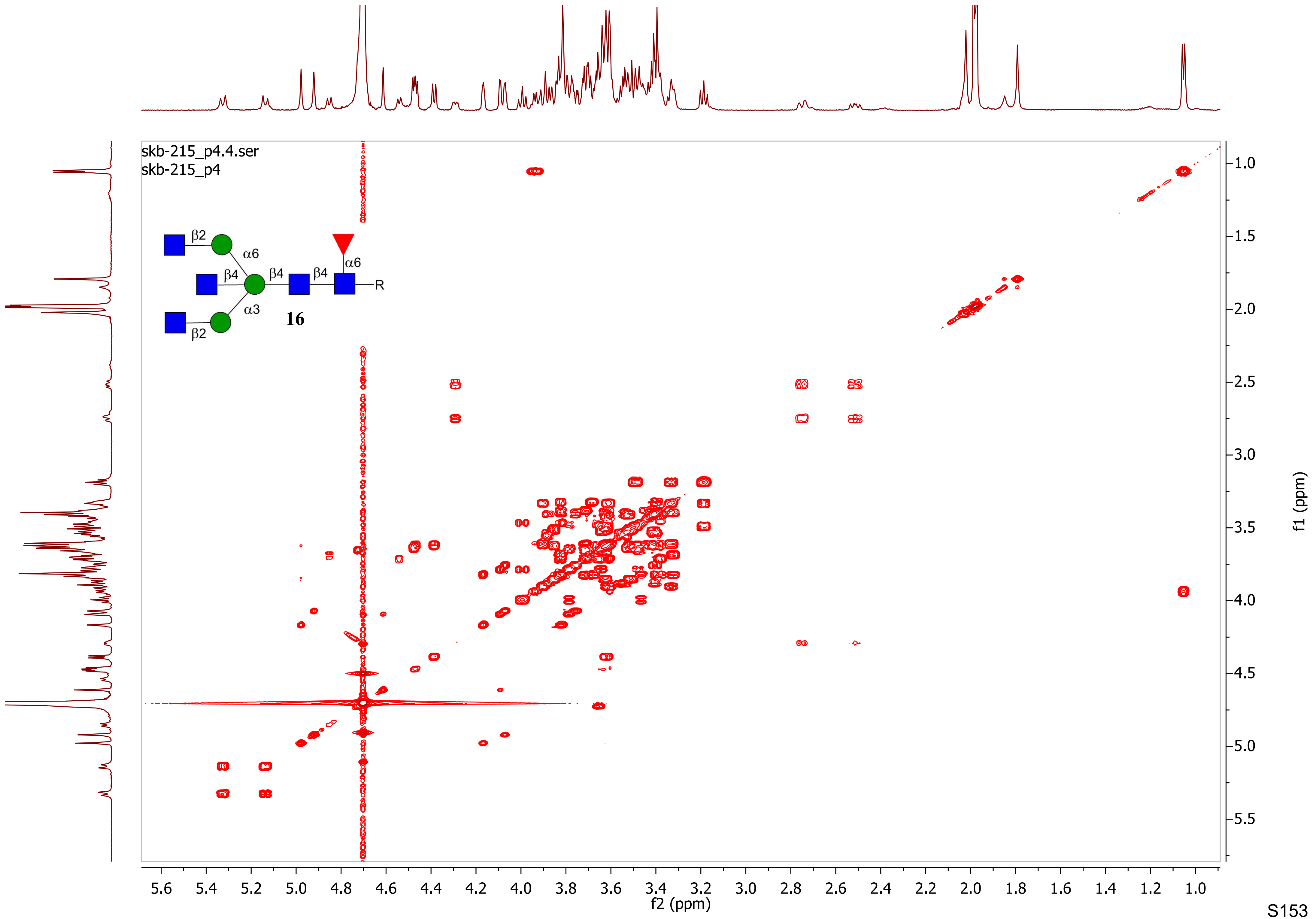
skb-215_p4.4.ser
skb-215_p4
β2
α6
β4
β4
β4
α6
R
α3
16
β2
f2 (ppm)
f1 (ppm)

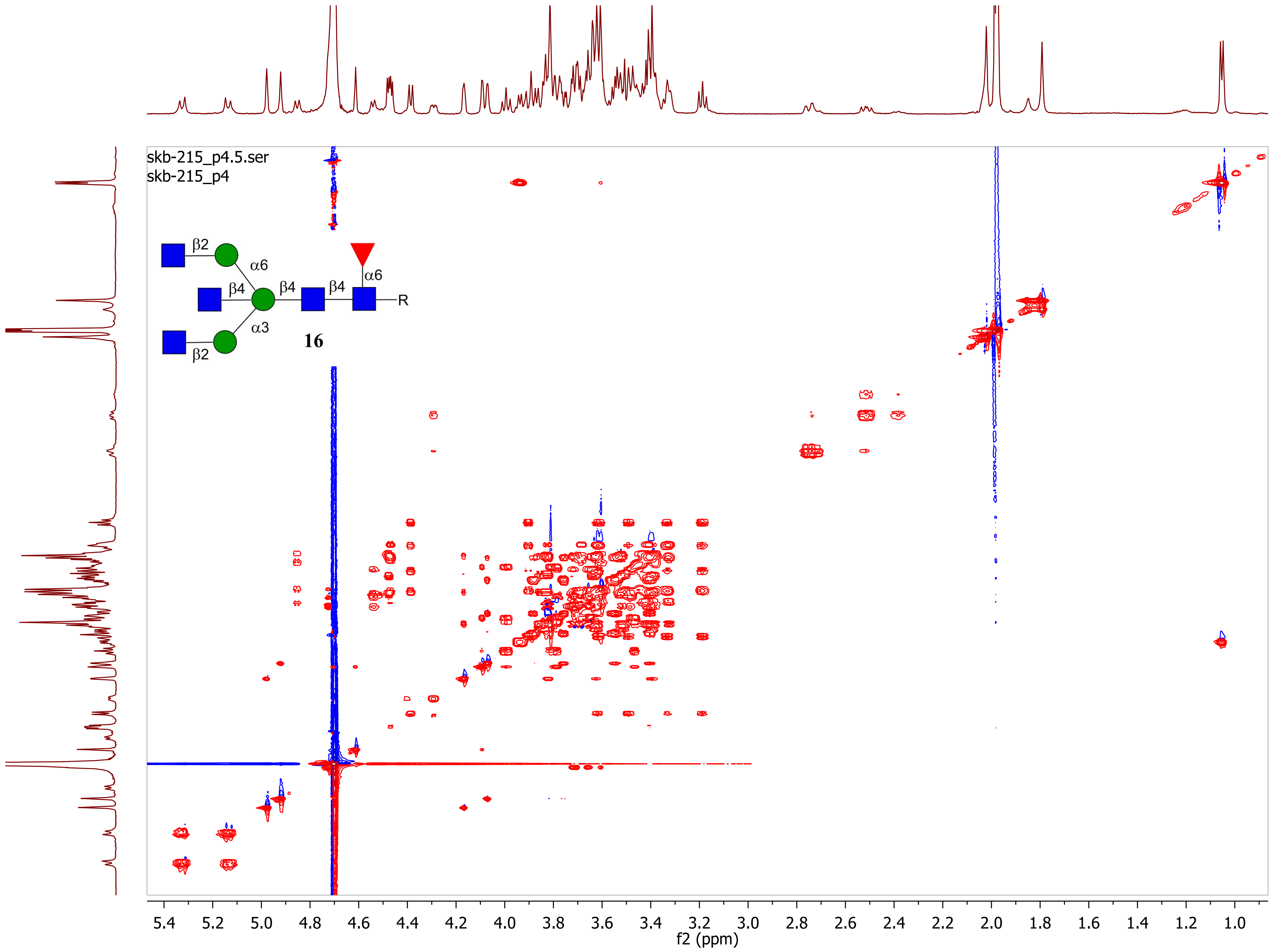
skb-215_p4.5.ser
skb-215_p4
β2
α6
β4
β4
β4
α6
R
α3
β2
16
f2 (ppm)
f1 (ppm)

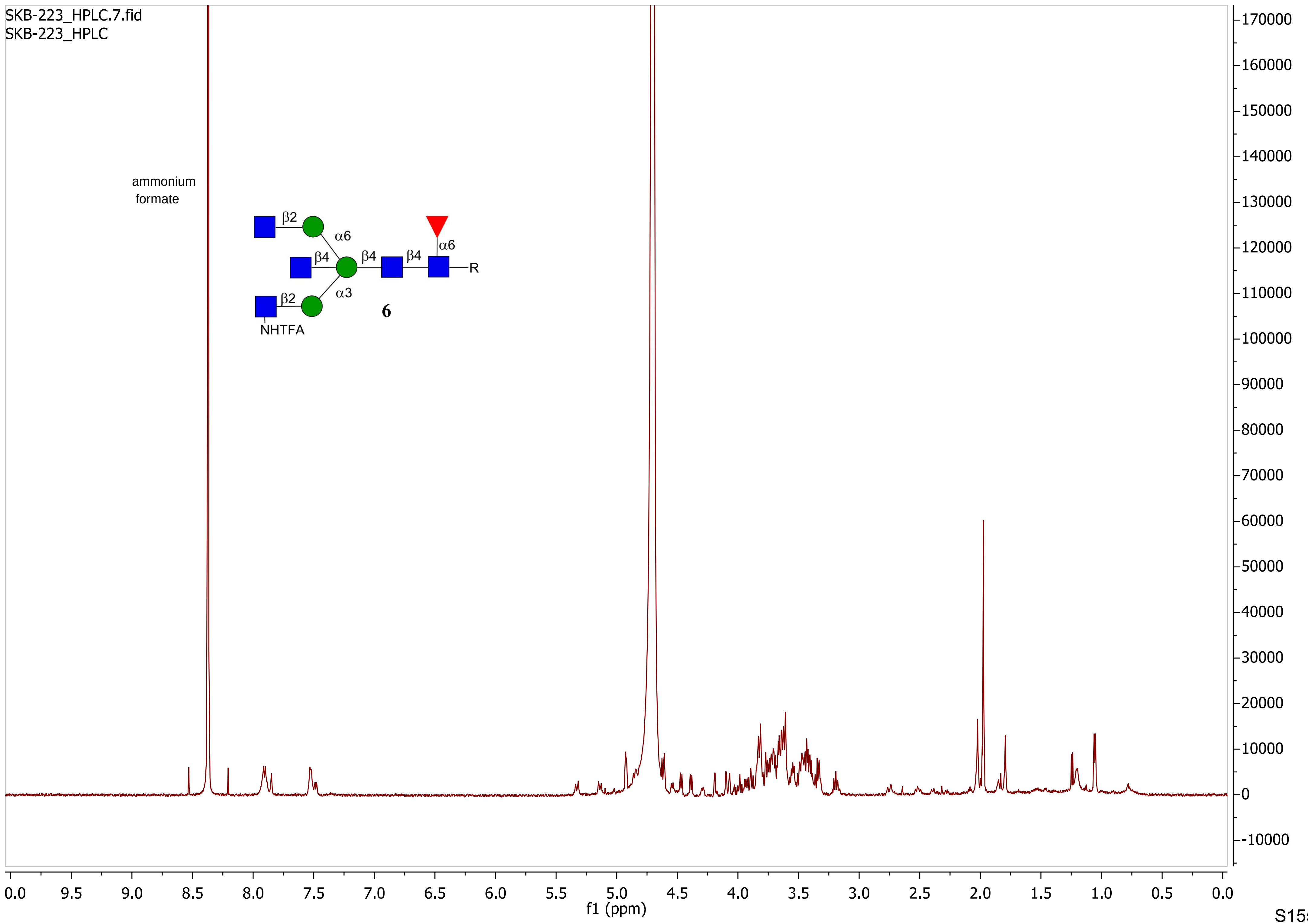

SKB-223_HPLC.7.fid
SKB-223_HPLC
ammonium formate
β2
α6
β4
β4
β4
α6
R
α3
β2
NHTFA
6
f1 (ppm)

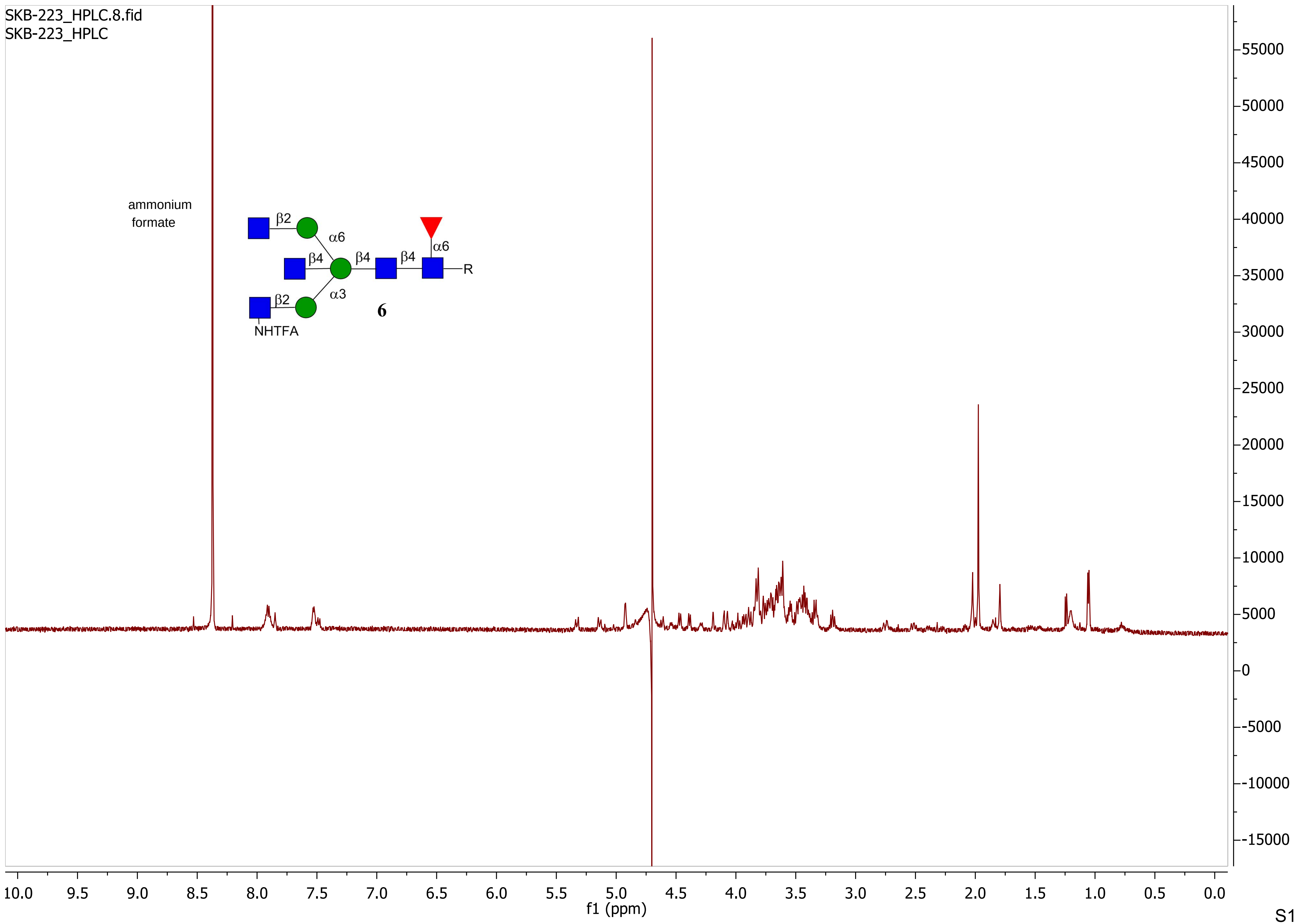

SKB-223_HPLC.8.fid
SKB-223_HPLC
ammonium formate
β2
α6
β4
β4
β4
α6
R
α3
β2
NHTFA
6
f1 (ppm)

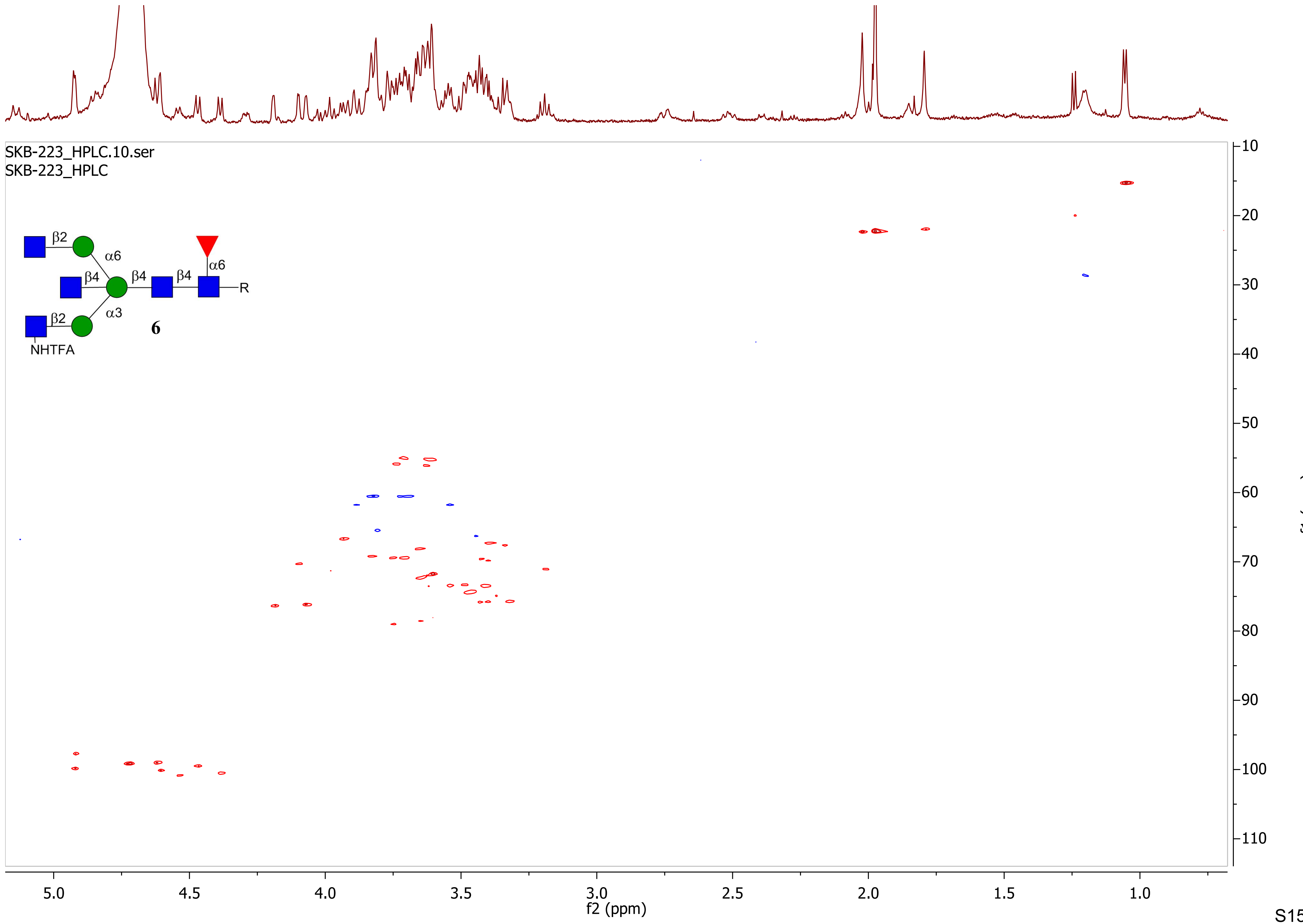
SKB-223_HPLC.10.ser
SKB-223_HPLC
β2
α6
β4
β4
β4
α6
R
α3
β2
6
NHTFA
f2 (ppm)
f1 (ppm)

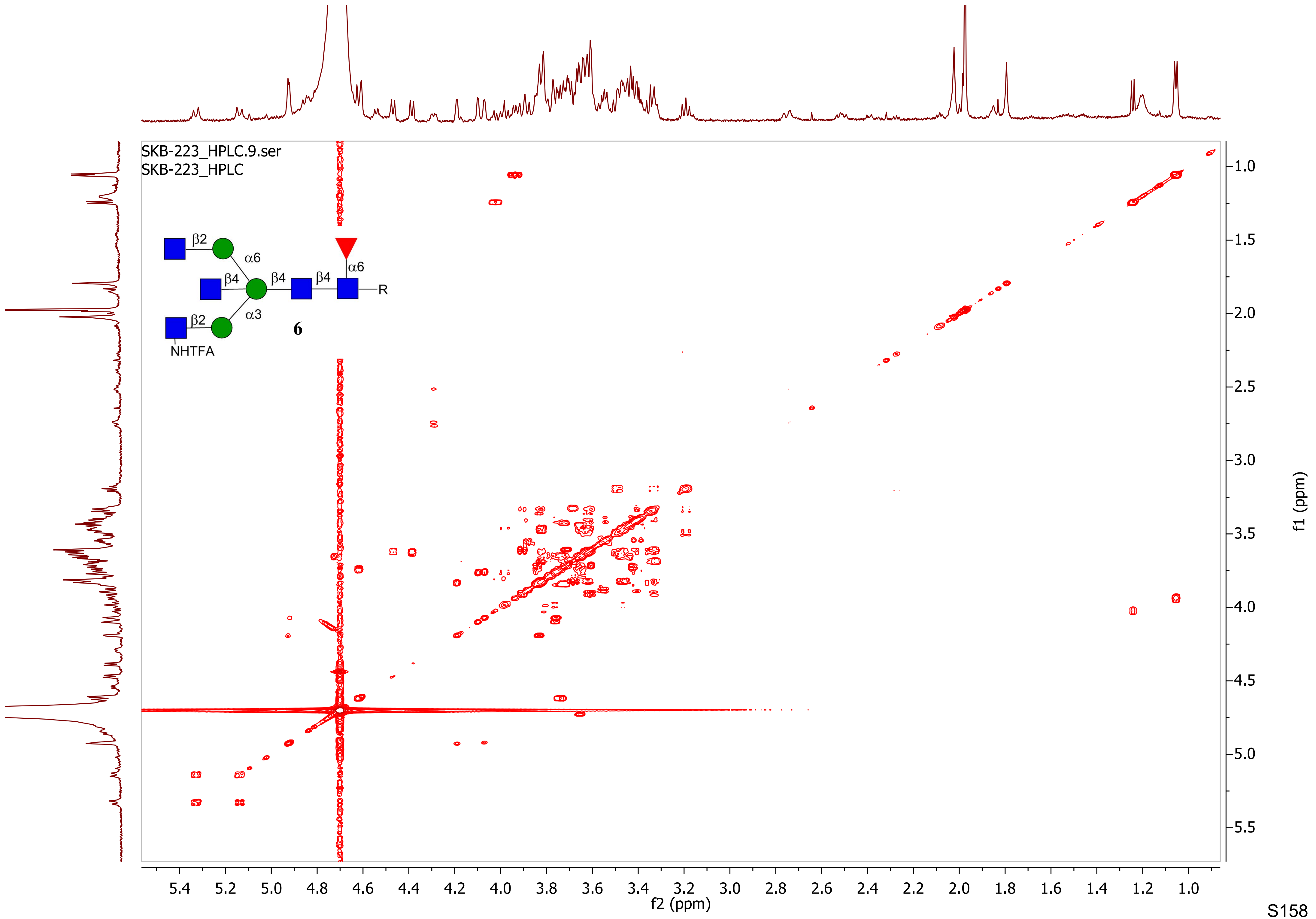
SKB-223_HPLC.9.ser
SKB-223_HPLC
β2
α6
β4
β4
β4
α6
R
β2
α3
NHTFA
6
f2 (ppm)
f1 (ppm)

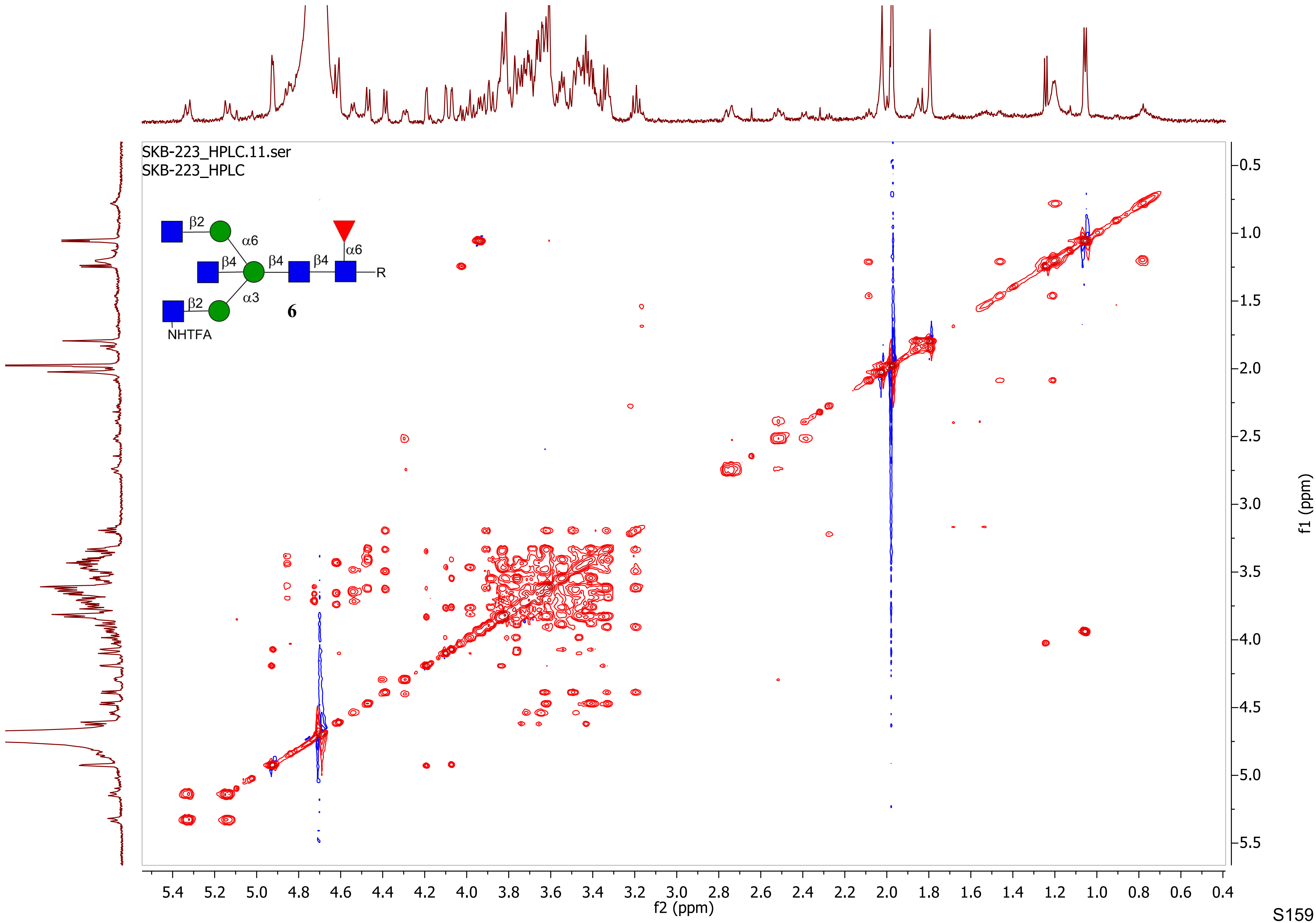
SKB-223_HPLC.11.ser
SKB-223_HPLC
β2
α6
β4
β4
β4
α6
R
β2
α3
6
NHTFA
f2 (ppm)
f1 (ppm)

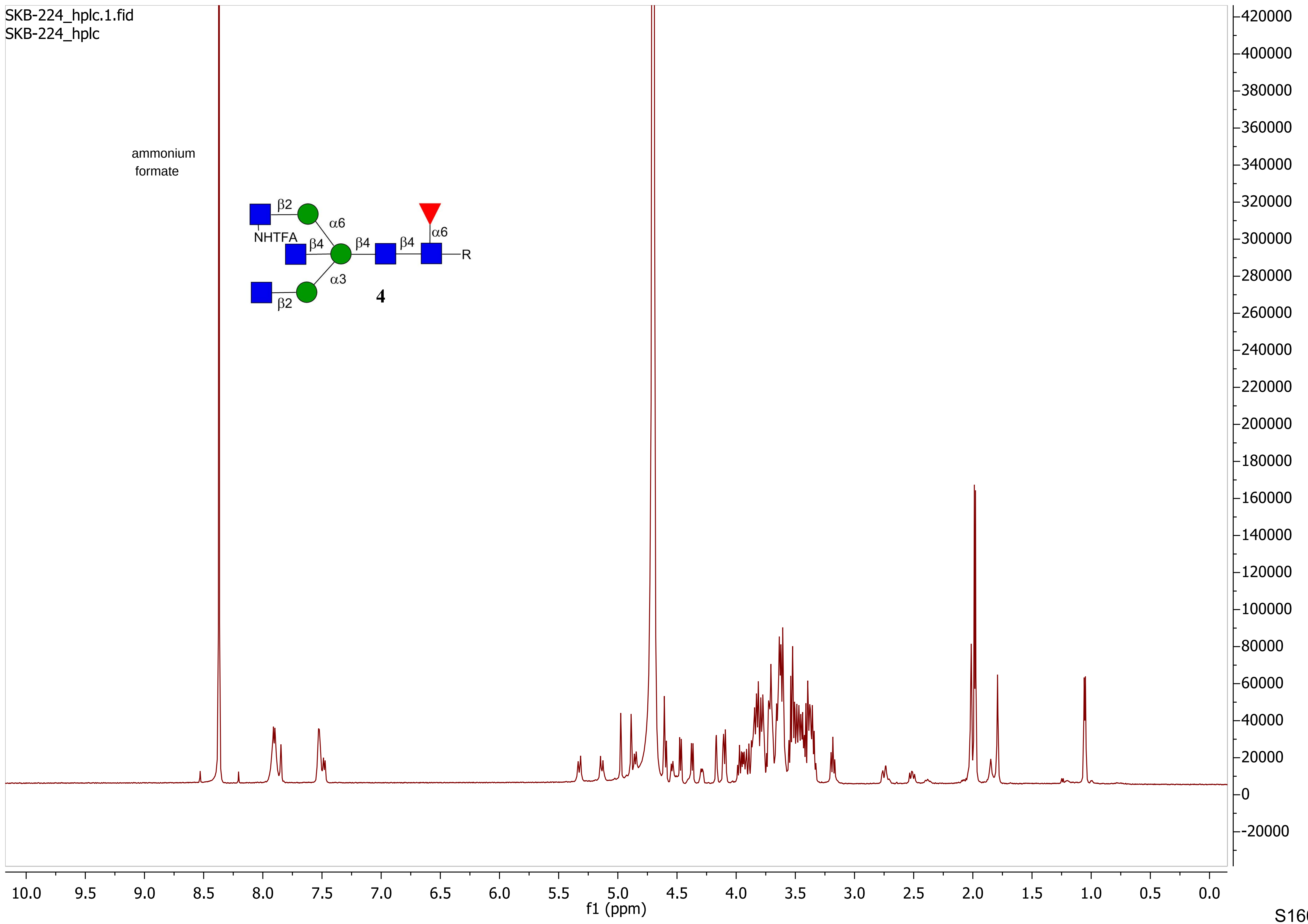

SKB-224_hplc.1.fid
SKB-224_hplc
ammonium formate
β2
α6
NHTFA
β4
β4
β4
α6
R
α3
β2
4
f1 (ppm)

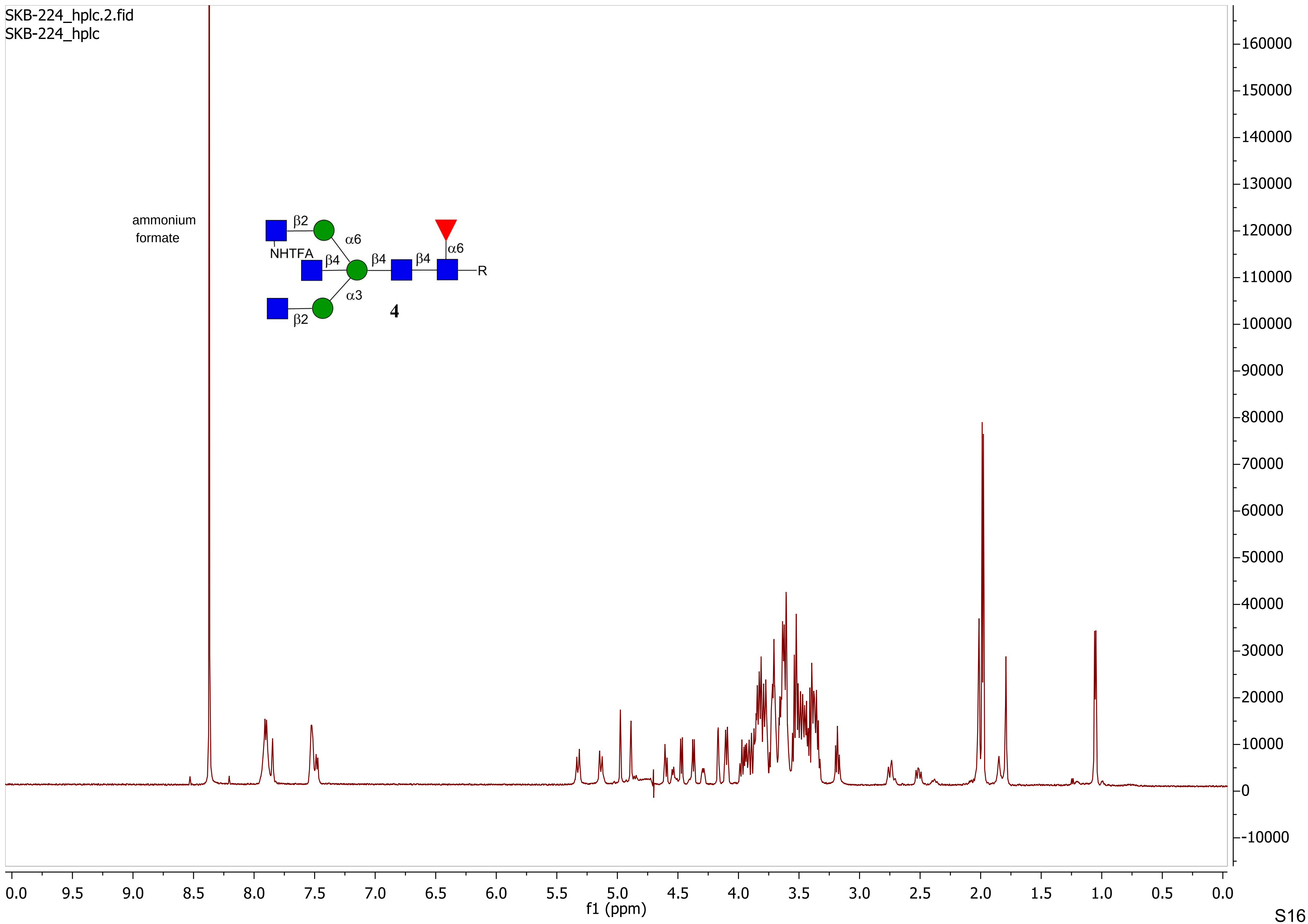
SKB-224_hplc.2.fid
SKB-224_hplc
ammonium
formate
β2
α6
NHTFA
β4
β4
β4
α6
R
α3
β2
4
f1 (ppm)

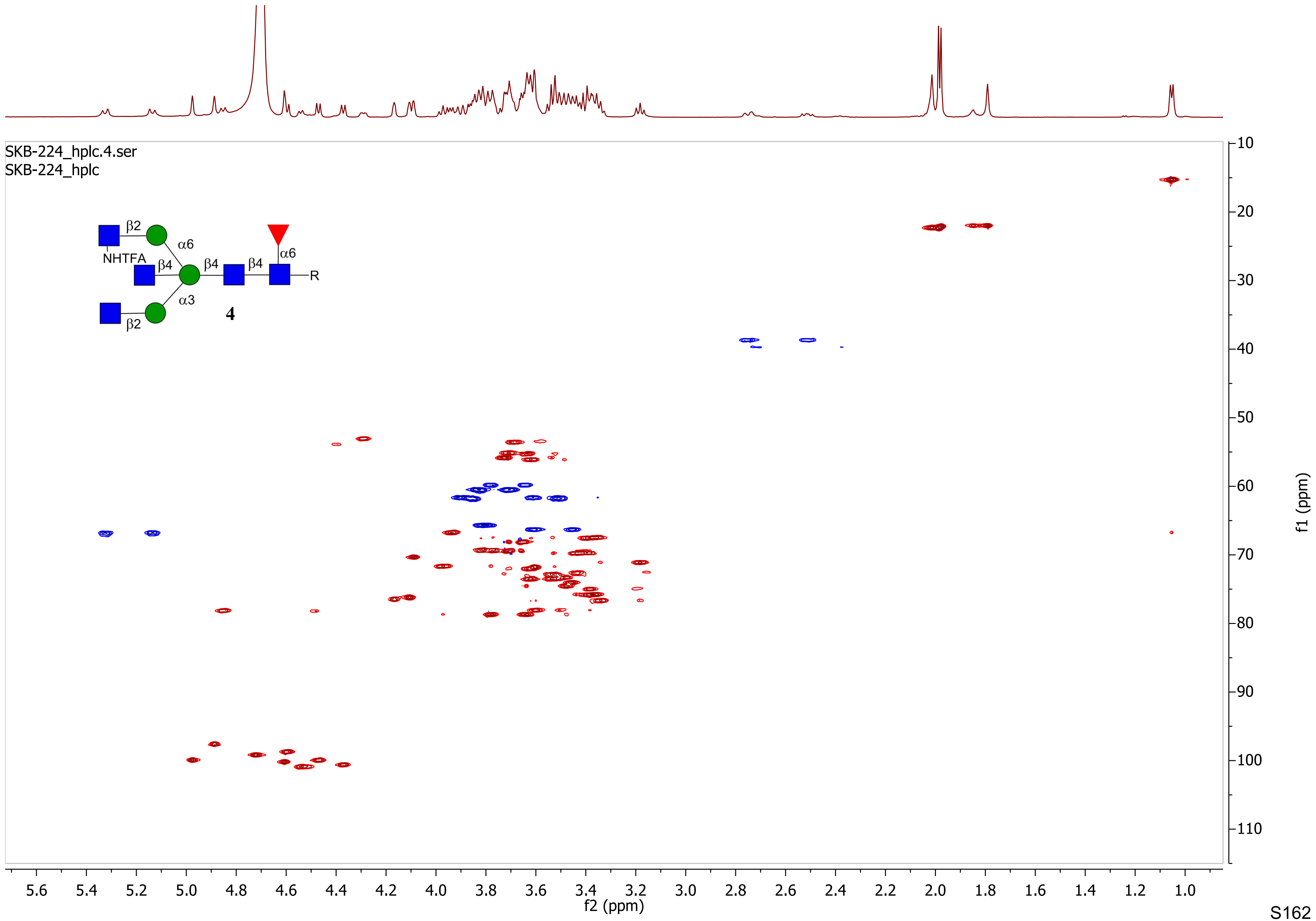
SKB-224_hplc.4.ser
SKB-224_hplc
β2
α6
NHTFA
β4
β4
β4
α6
R
α3
4
β2
f2 (ppm)
f1 (ppm)

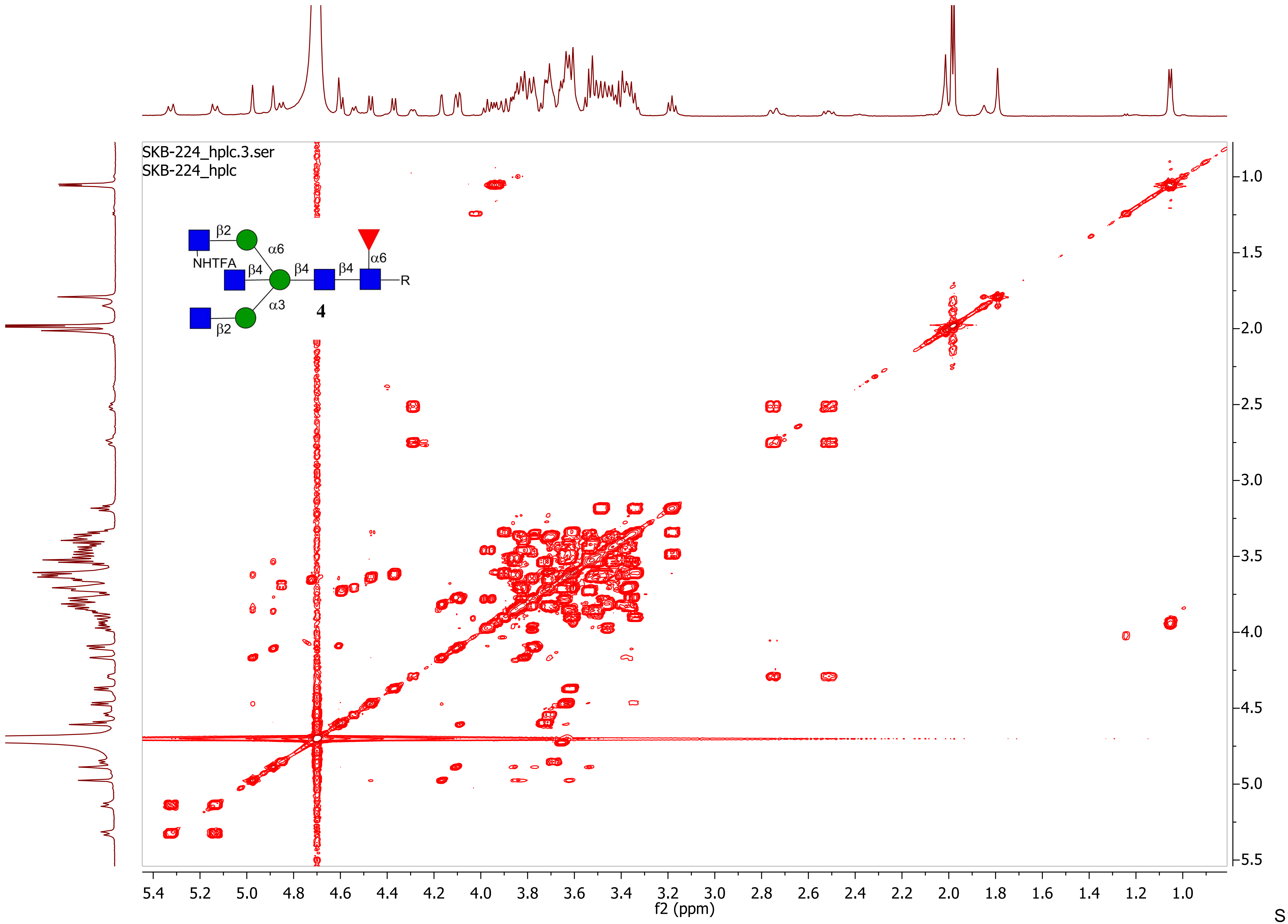

SKB-224_hplc.3.ser
SKB-224_hplc
β2
α6
NHTFA
β4
β4
β4
α6
R
α3
β2
4
f2 (ppm)
f1 (ppm)

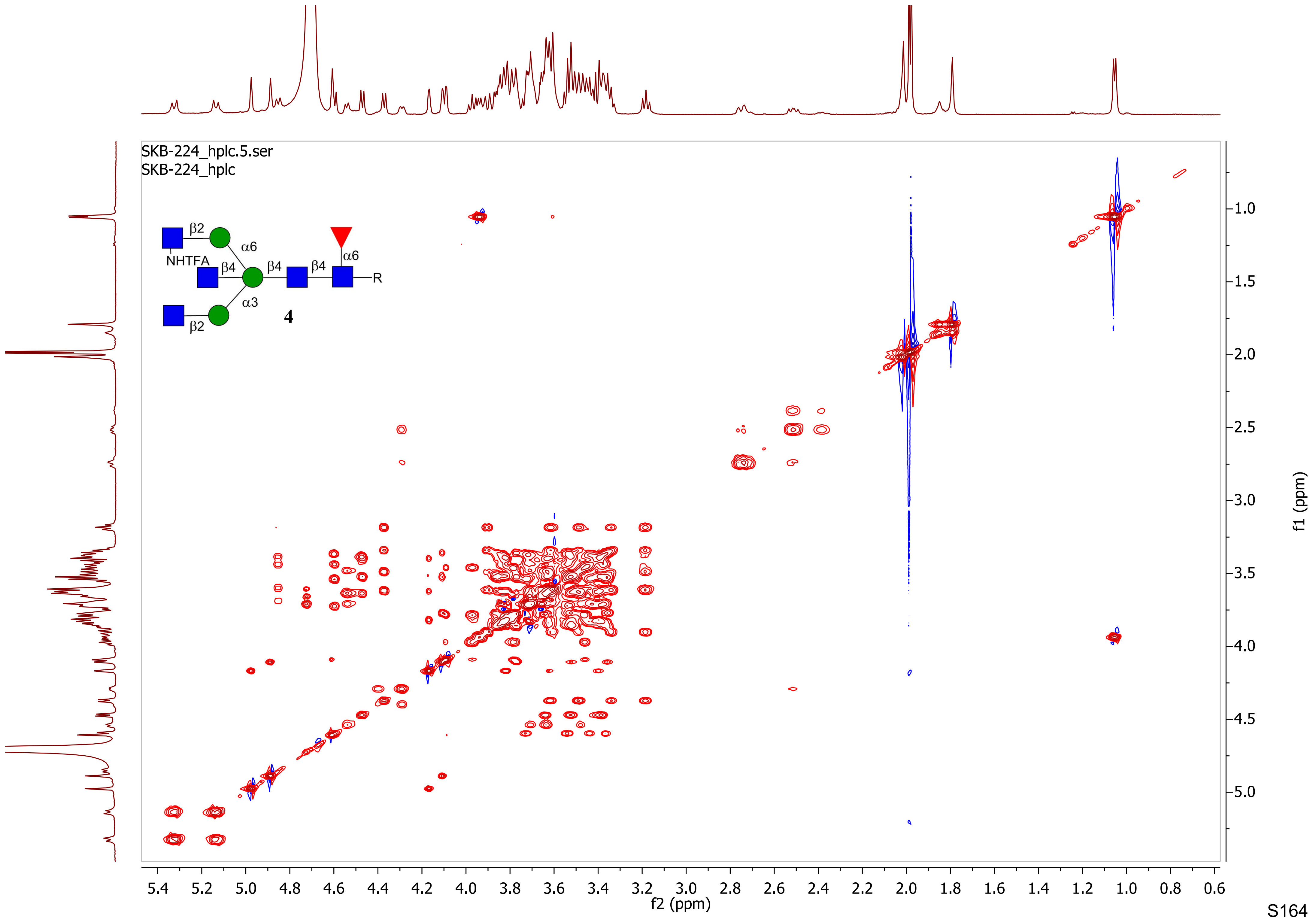
SKB-224_hplc.5.ser
SKB-224_hplc
β2
α6
NHTFA
β4
β4
β4
α6
R
α3
β2
4
f2 (ppm)
f1 (ppm)

skb-225_HPLC.1.fid
skb-225_HPLC

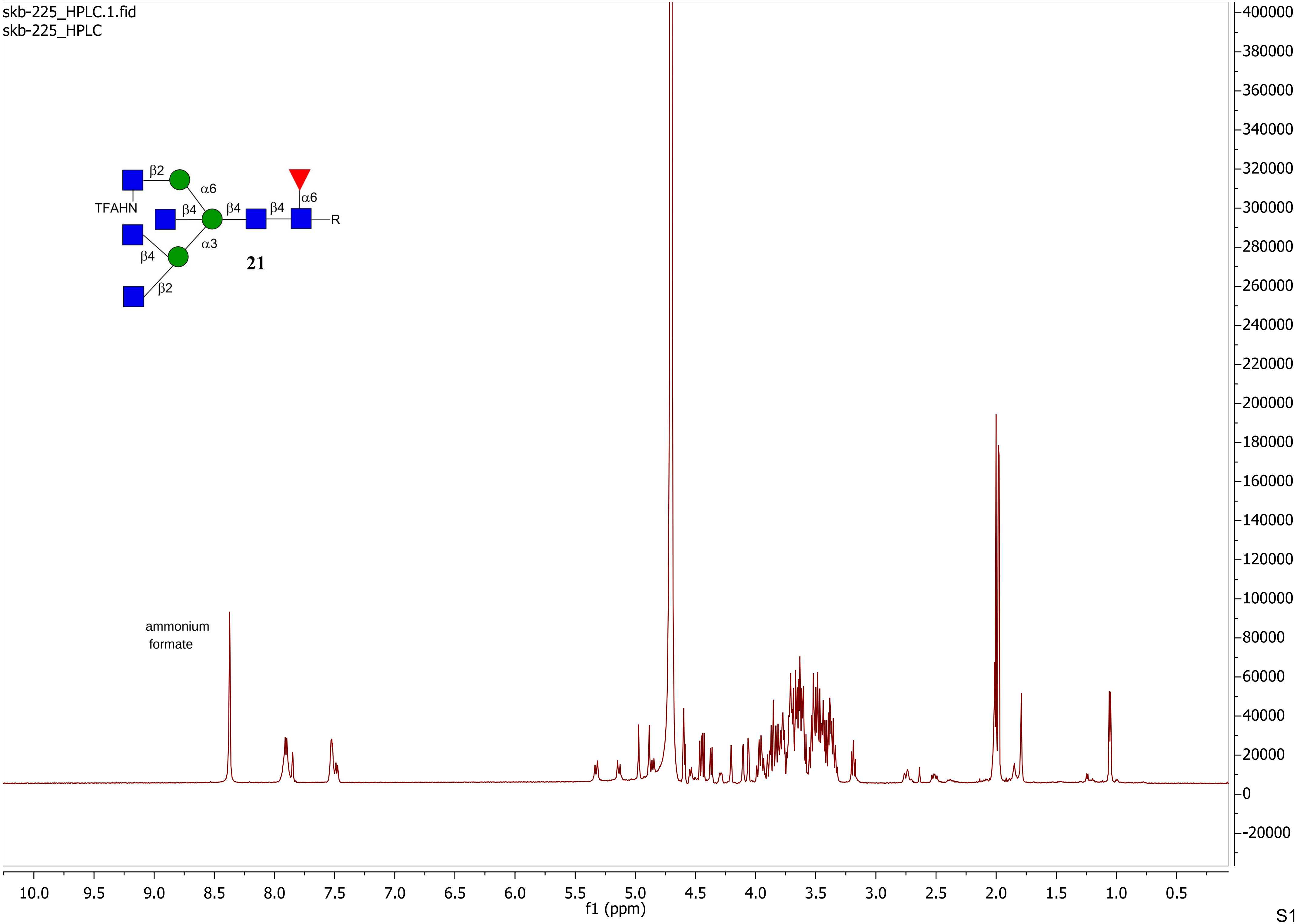

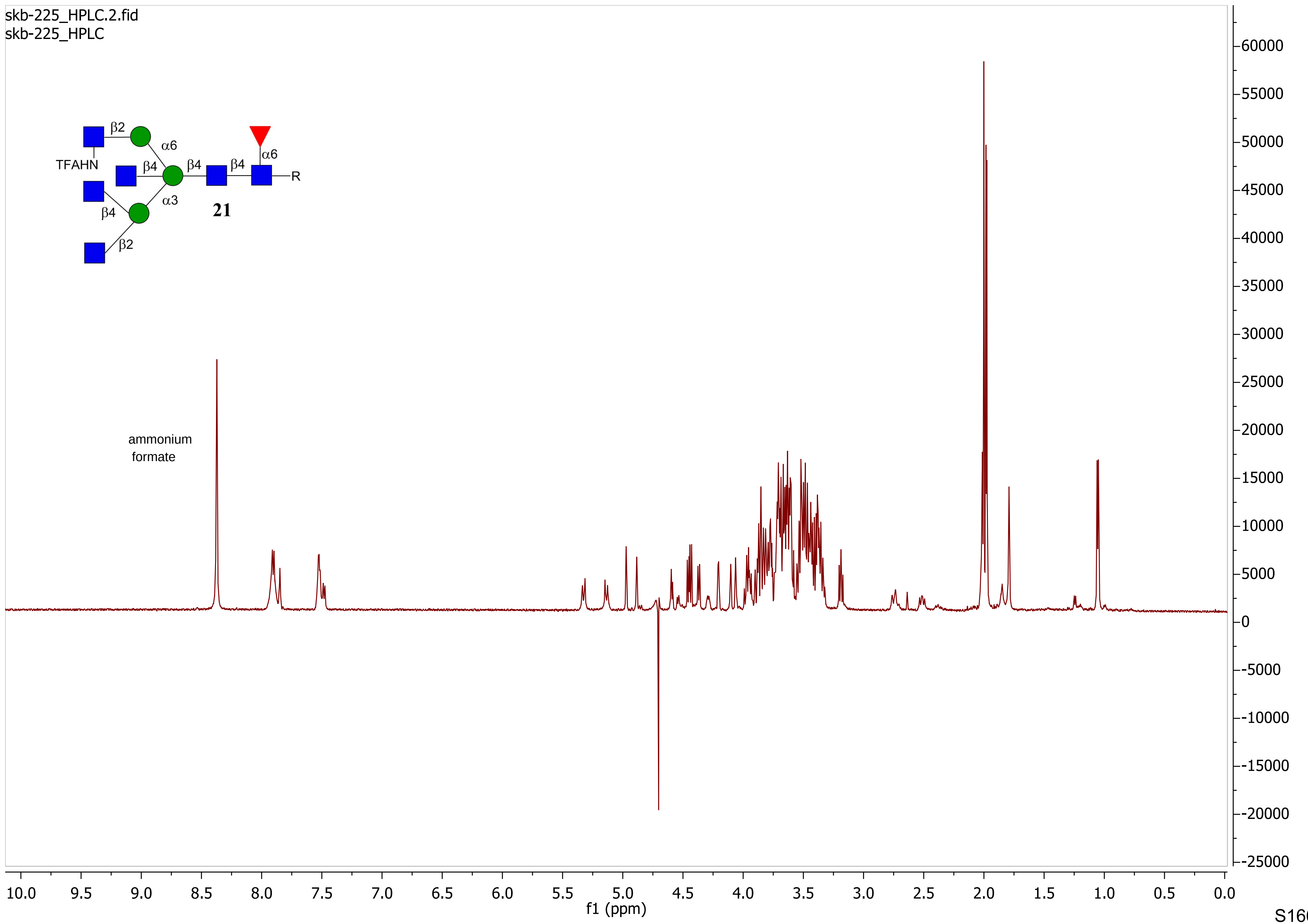
skb-225_HPLC.2.fid
skb-225_HPLC
β2
α6
TFAHN
β4
β4
β4
α6
R
β4
α3
β2
21
ammonium formate
f1 (ppm)

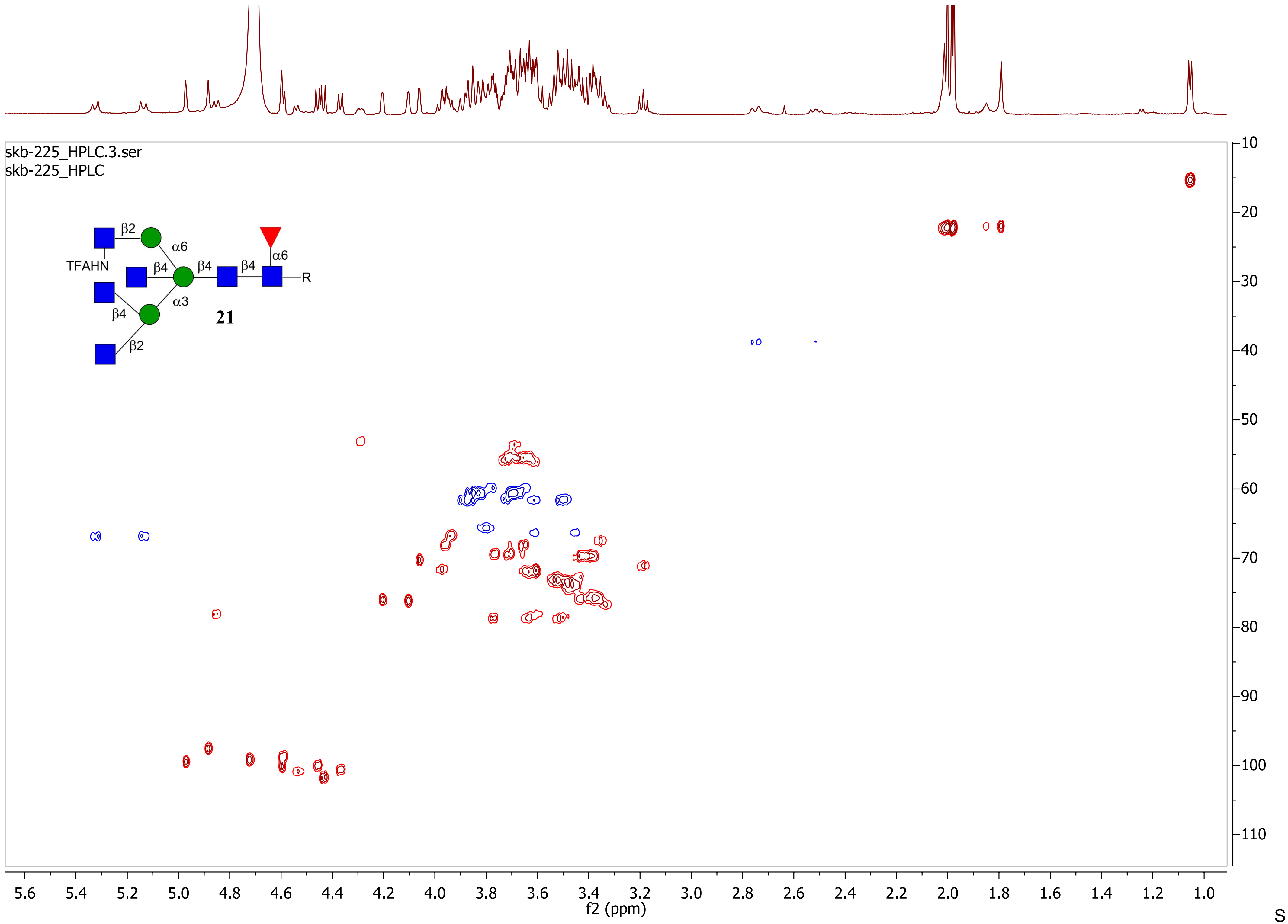

skb-225_HPLC.3.ser
skb-225_HPLC
β2
α6
TFAHN
β4
β4
β4
α6
R
β4
α3
21
β2
f1 (ppm)
f2 (ppm)

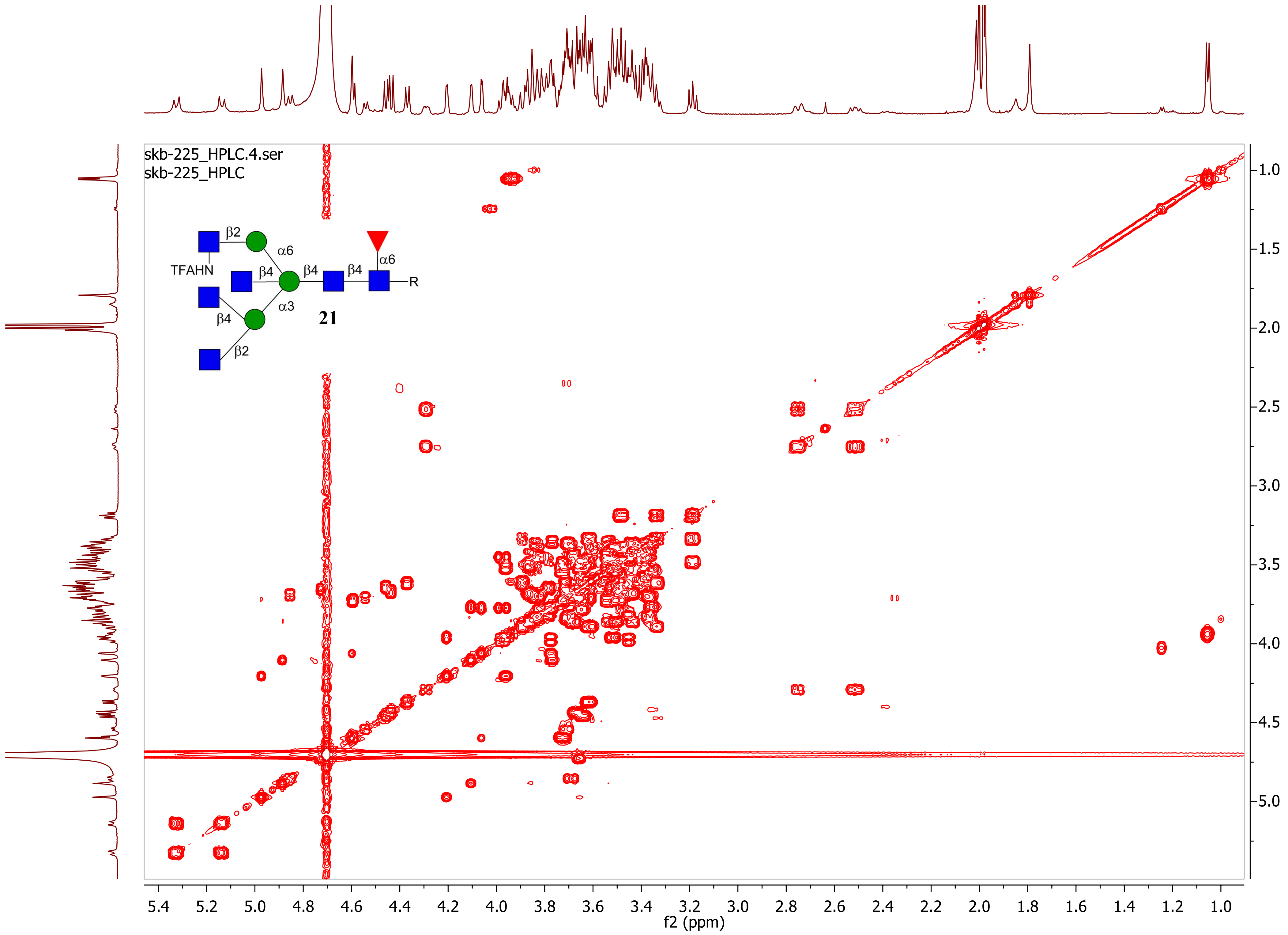
skb-225_HPLC.4.ser
skb-225_HPLC
β2
α6
TFAHN
β4
β4
β4
α6
R
β4
α3
21
β2
f2 (ppm)
f1 (ppm)

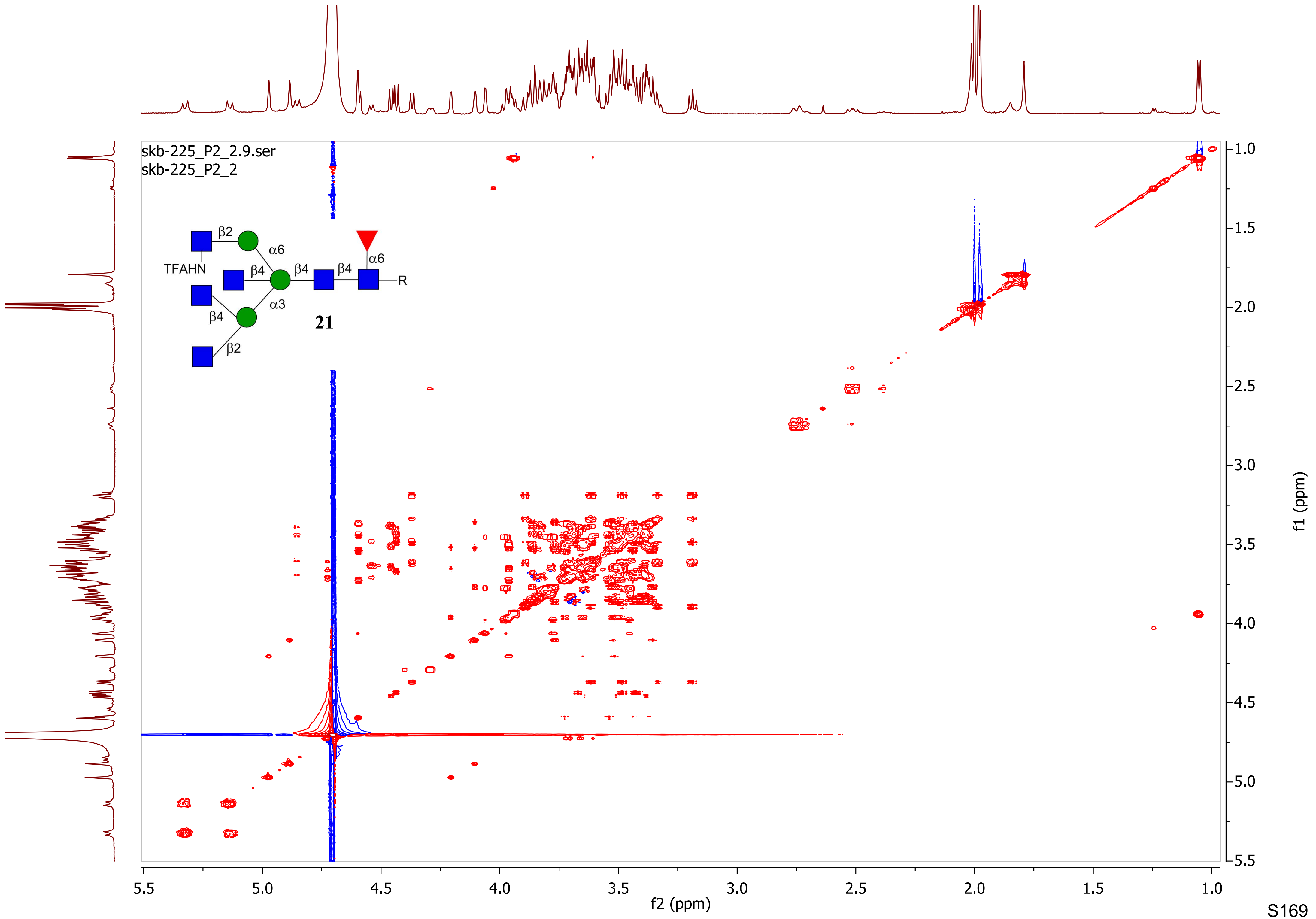

skb-225_P2_2.9.ser
skb-225_P2_2
TFAHN
β2
α6
β4
β4
β4
α6
R
α3
β4
β2
21
f2 (ppm)
f1 (ppm)

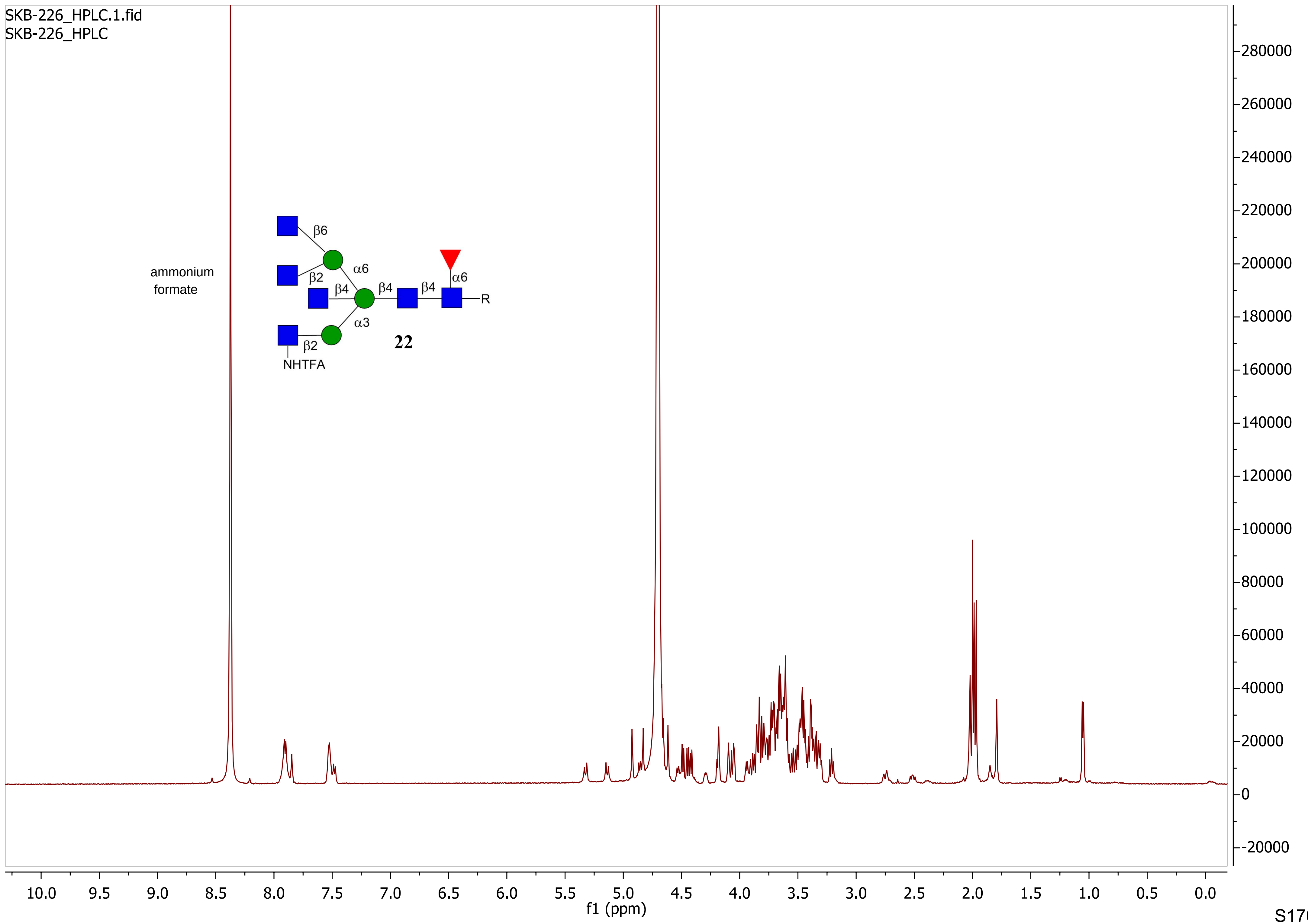
SKB-226_HPLC.1.fid
SKB-226_HPLC
ammonium formate
β6
β2
α6
β4
β4
β4
α6
R
α3
β2
NHTFA
22
f1 (ppm)

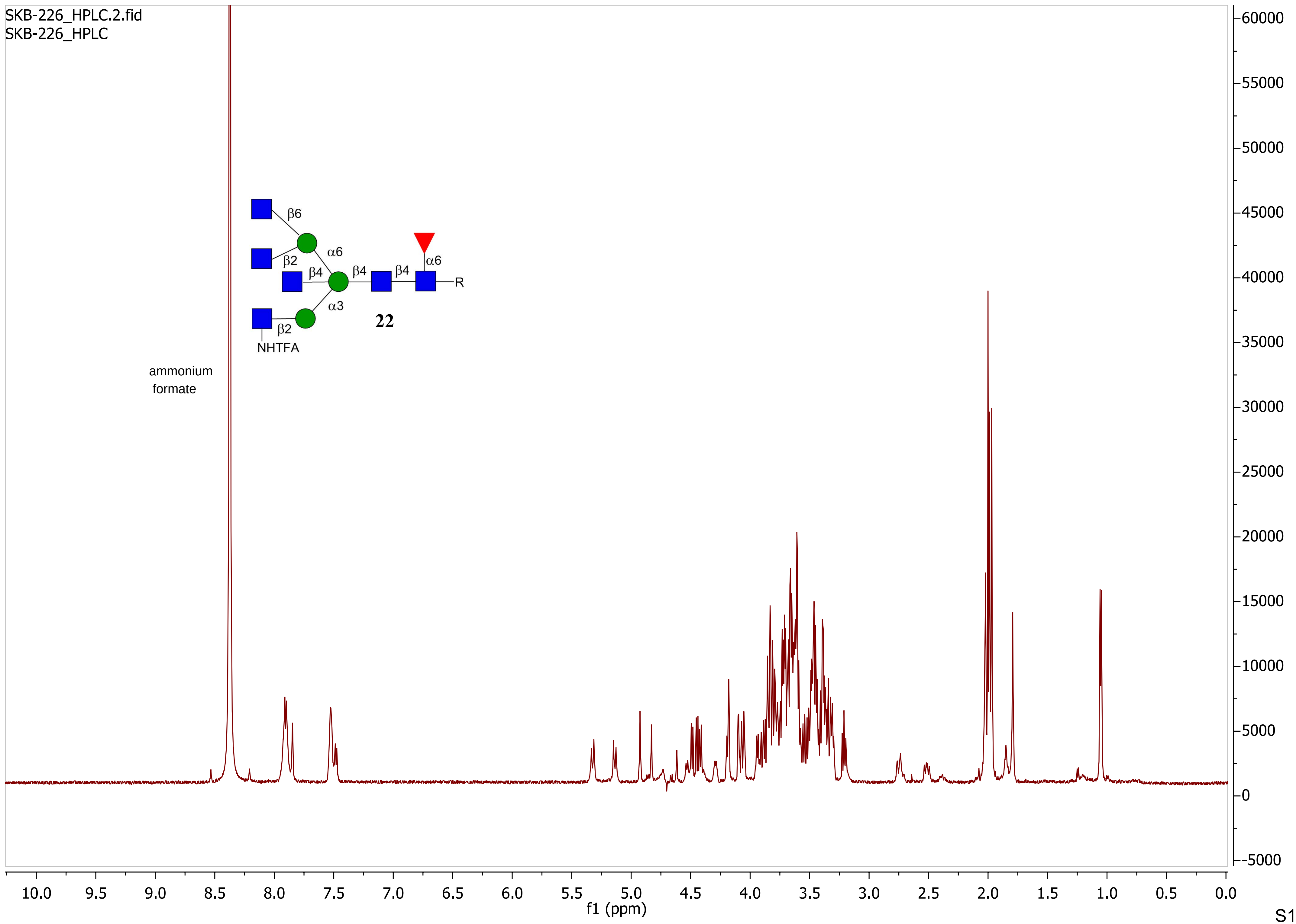

SKB-226_HPLC.2.fid
SKB-226_HPLC
β6
β2
α6
β4
β4
β4
α6
R
α3
β2
NHTFA
22
ammonium
formate
f1 (ppm)

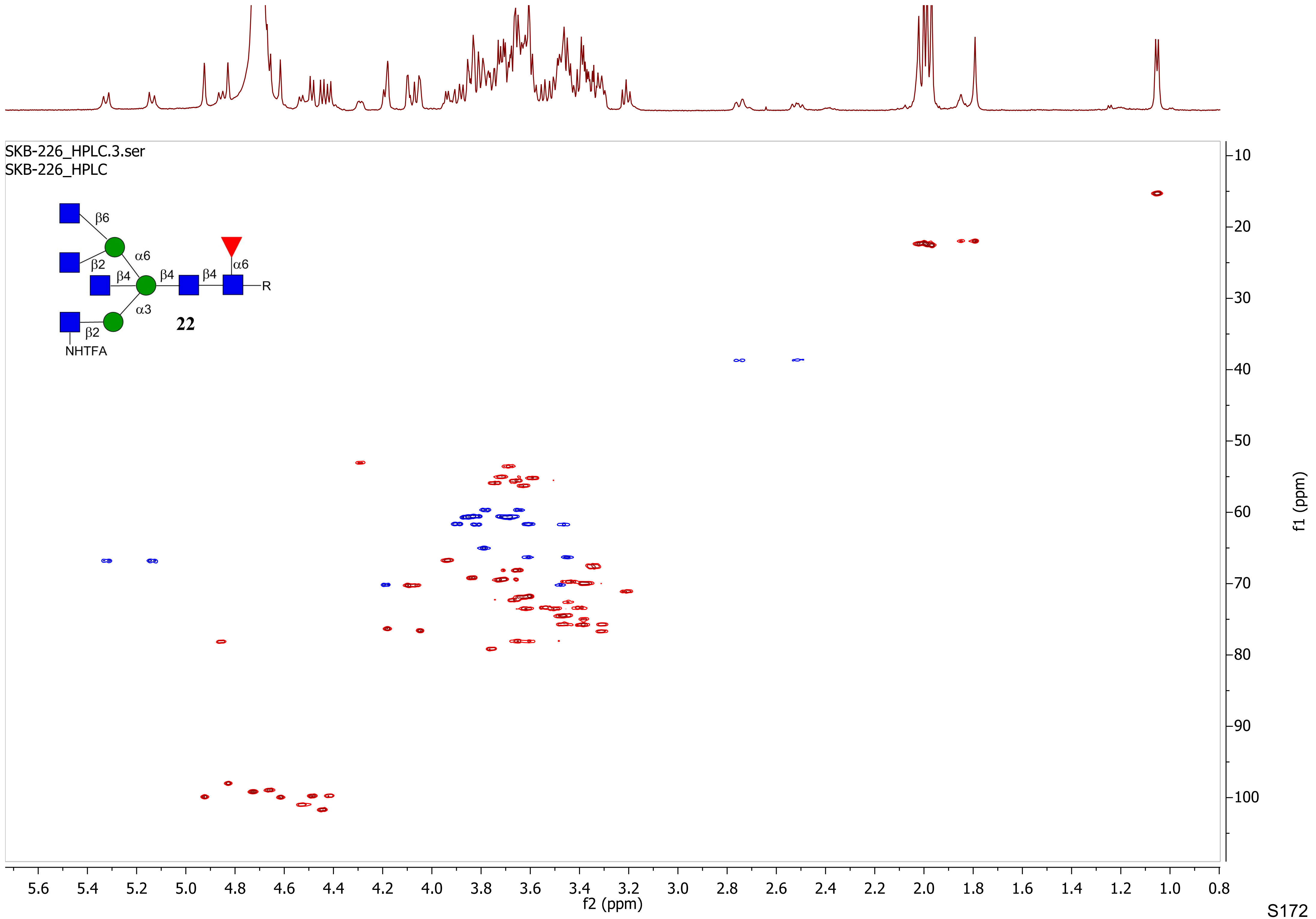
SKB-226_HPLC.3.ser
SKB-226_HPLC
β6
α6
β2
β4
β4
β4
α6
R
α3
22
β2
NHTFA
f1 (ppm)
f2 (ppm)

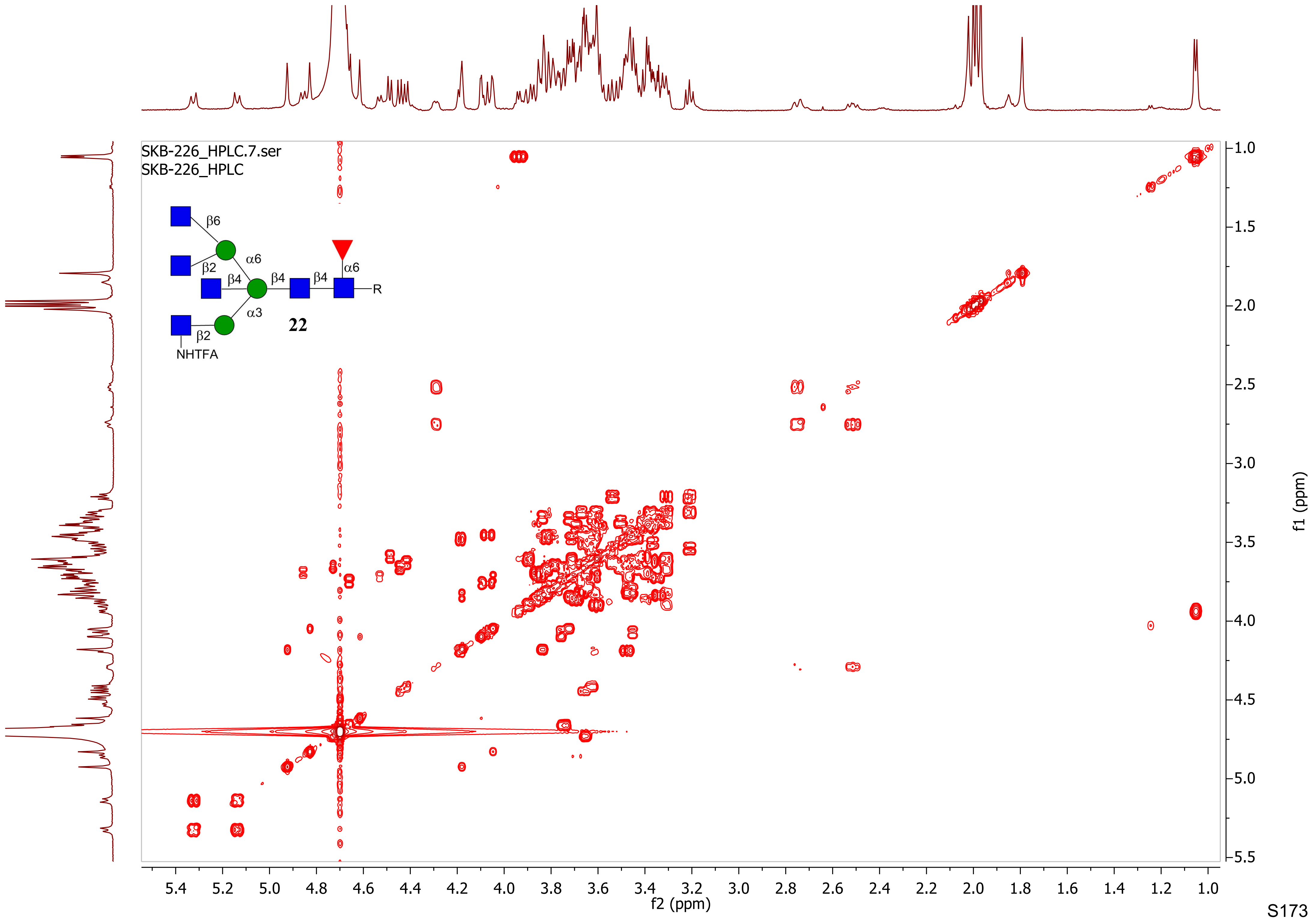

SKB-226_HPLC.7.ser
SKB-226_HPLC
β6
α6
β2
β4
β4
β4
α6
R
α3
22
β2
NHTFA
f1 (ppm)
f2 (ppm)

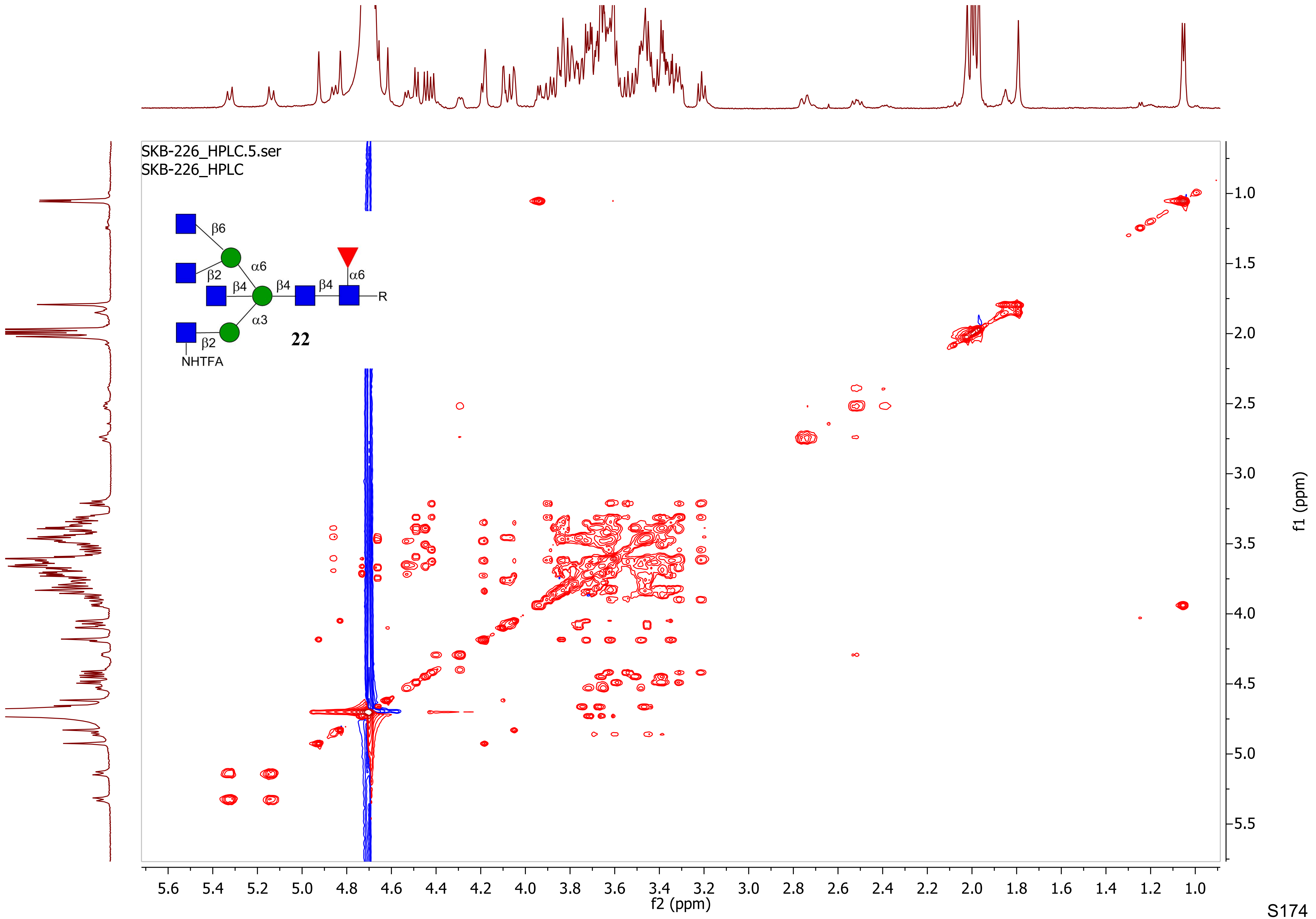

SKB-226_HPLC.5.ser
SKB-226_HPLC
β6
α6
β2
β4
β4
β4
α6
R
α3
β2
NHTFA
22
f2 (ppm)
f1 (ppm)

SKB-MGAT_1_Azide

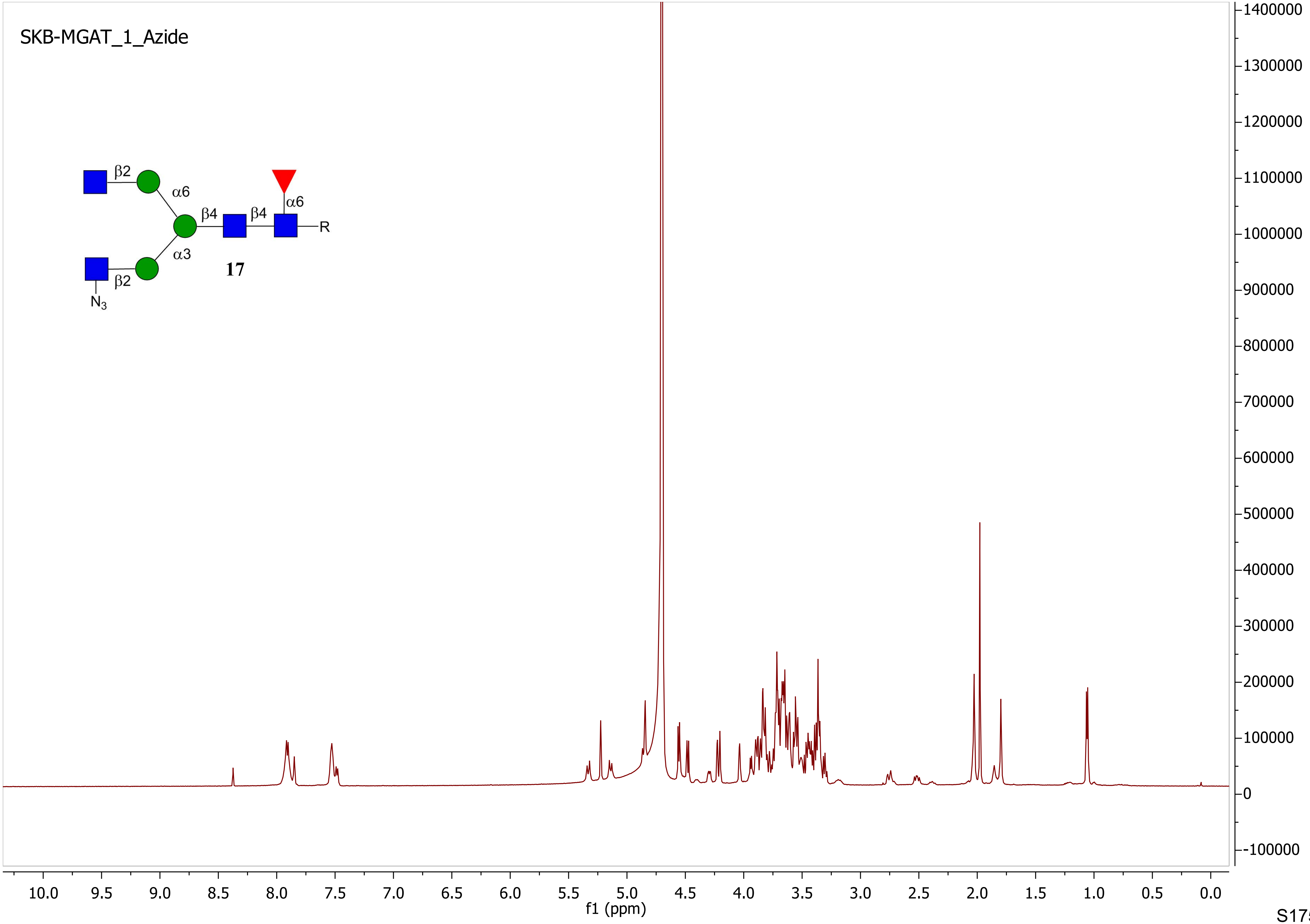

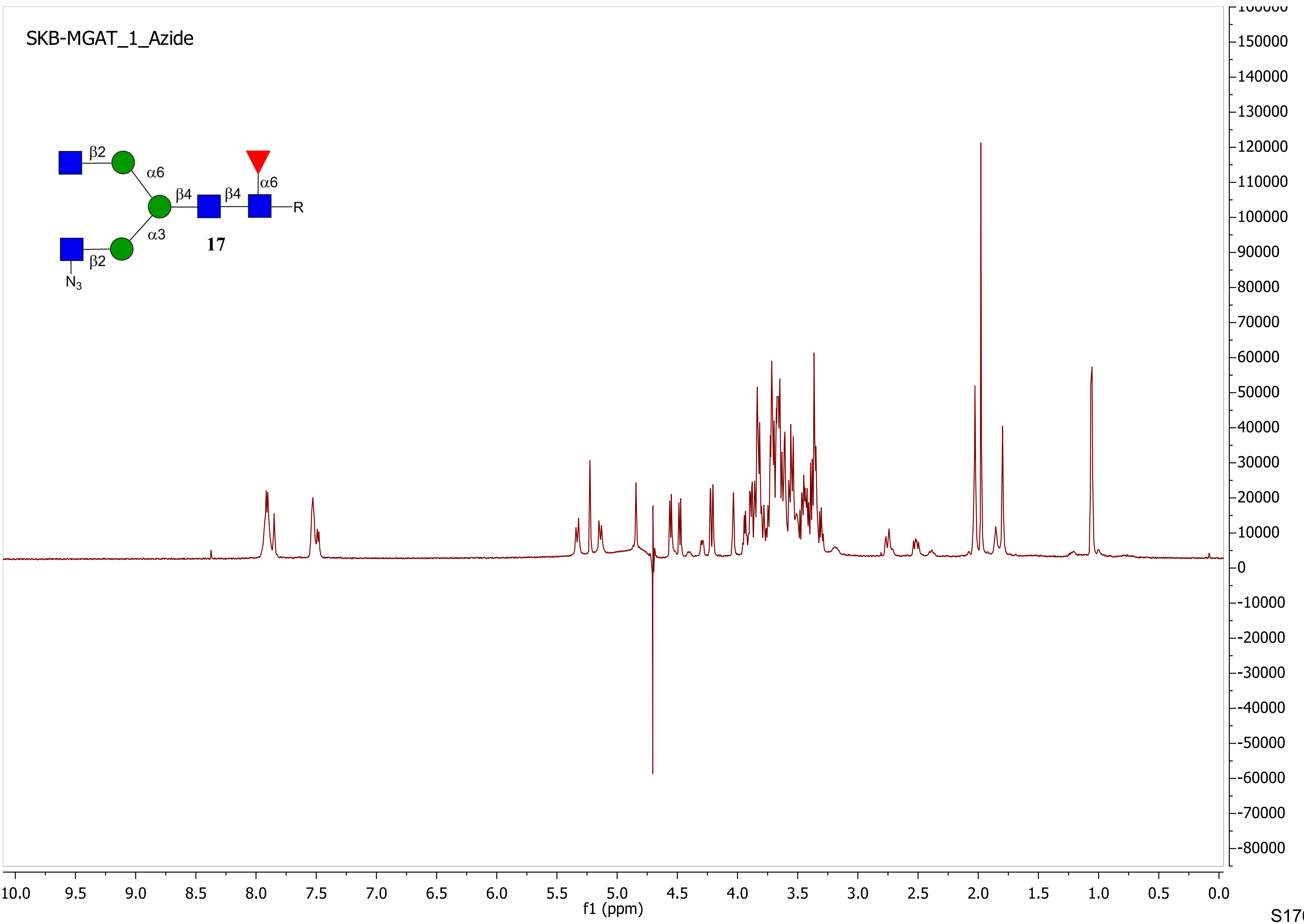
SKB-MGAT_1_Azide
β2
α6
β4
β4
α6
R
α3
17
β2
N3
f1 (ppm)
10.0
9.5
9.0
8.5
8.0
7.5
7.0
6.5
6.0
5.5
5.0
4.5
4.0
3.5
3.0
2.5
2.0
1.5
1.0
0.5
0.0
150000
140000
130000
120000
110000
100000
90000
80000
70000
60000
50000
40000
30000
20000
10000
0
-10000
-20000
-30000
-40000
-50000
-60000
-70000
-80000

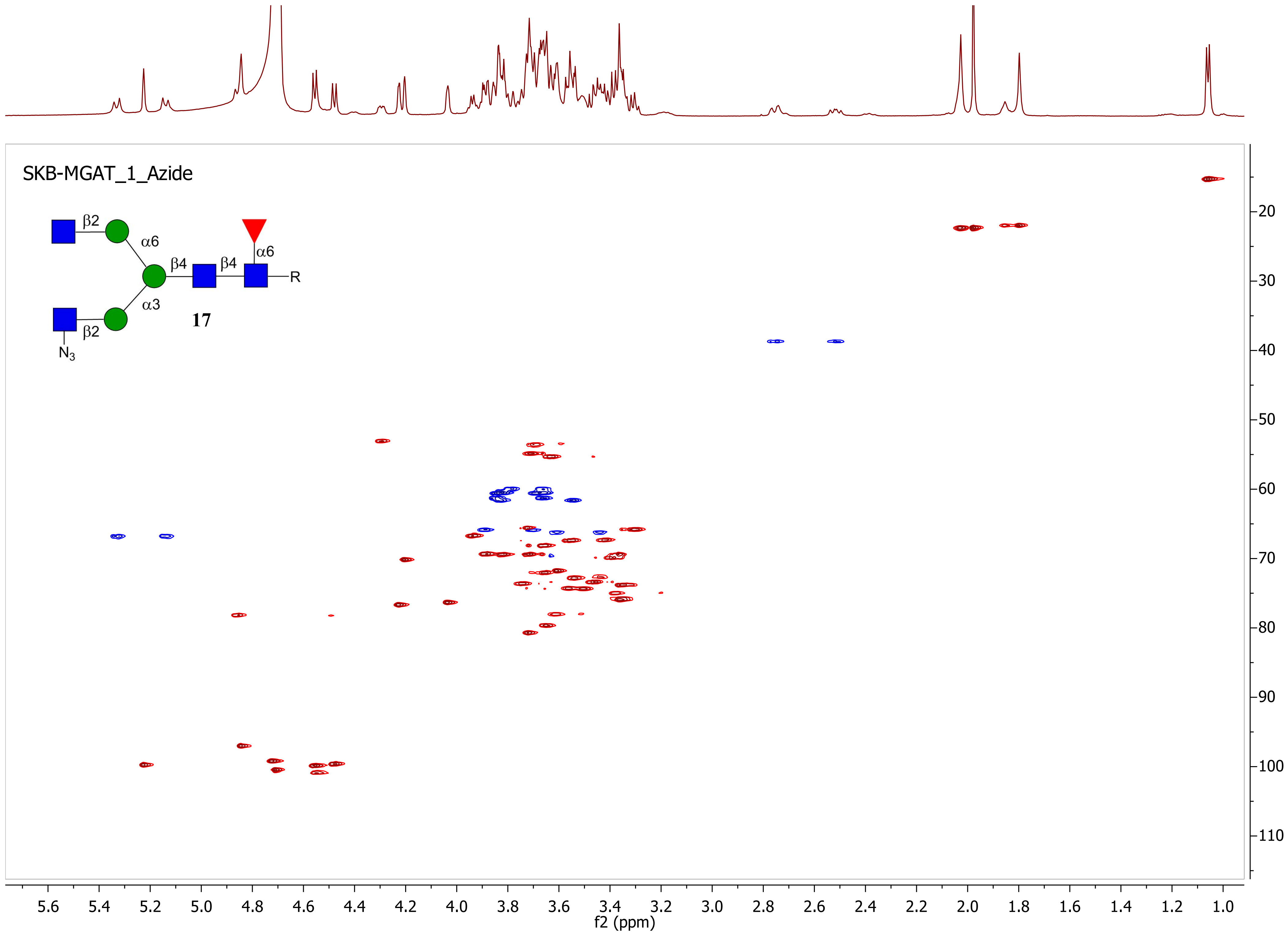
SKB-MGAT_1_Azide
β2
α6
β4
β4
α6
R
α3
17
β2
N3
5.6
5.4
5.2
5.0
4.8
4.6
4.4
4.2
4.0
3.8
3.6
3.4
3.2
3.0
2.8
2.6
2.4
2.2
2.0
1.8
1.6
1.4
1.2
1.0
f2 (ppm)
20
30
40
50
60
70
80
90
100
110
f1 (ppm)

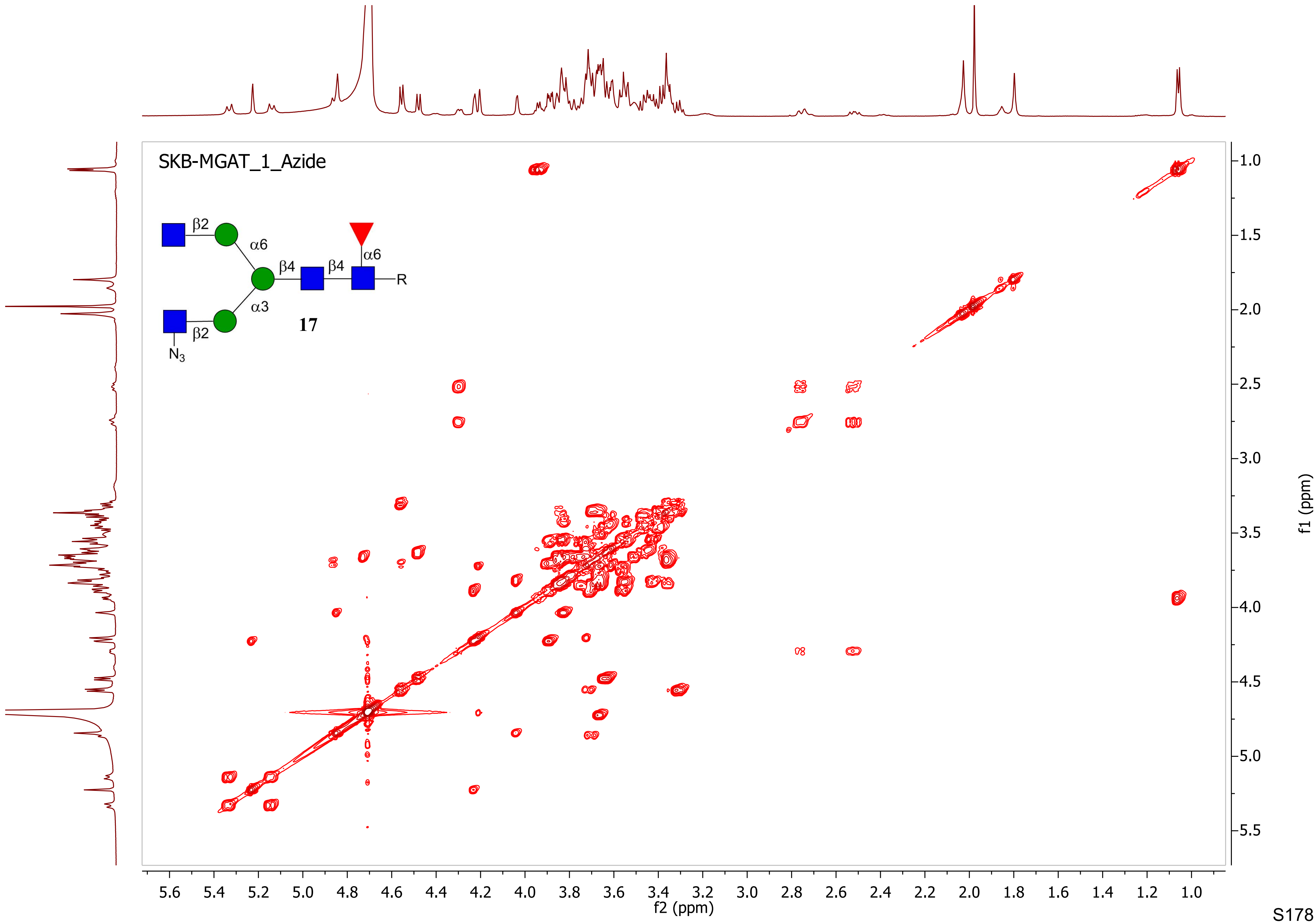
SKB-MGAT_1_Azide
β2
α6
β4
β4
α6
R
α3
β2
$N_3$
17
f2 (ppm)
f1 (ppm)
5.6
5.4
5.2
5.0
4.8
4.6
4.4
4.2
4.0
3.8
3.6
3.4
3.2
3.0
2.8
2.6
2.4
2.2
2.0
1.8
1.6
1.4
1.2
1.0
1.5
2.5
3.5
4.5
5.5

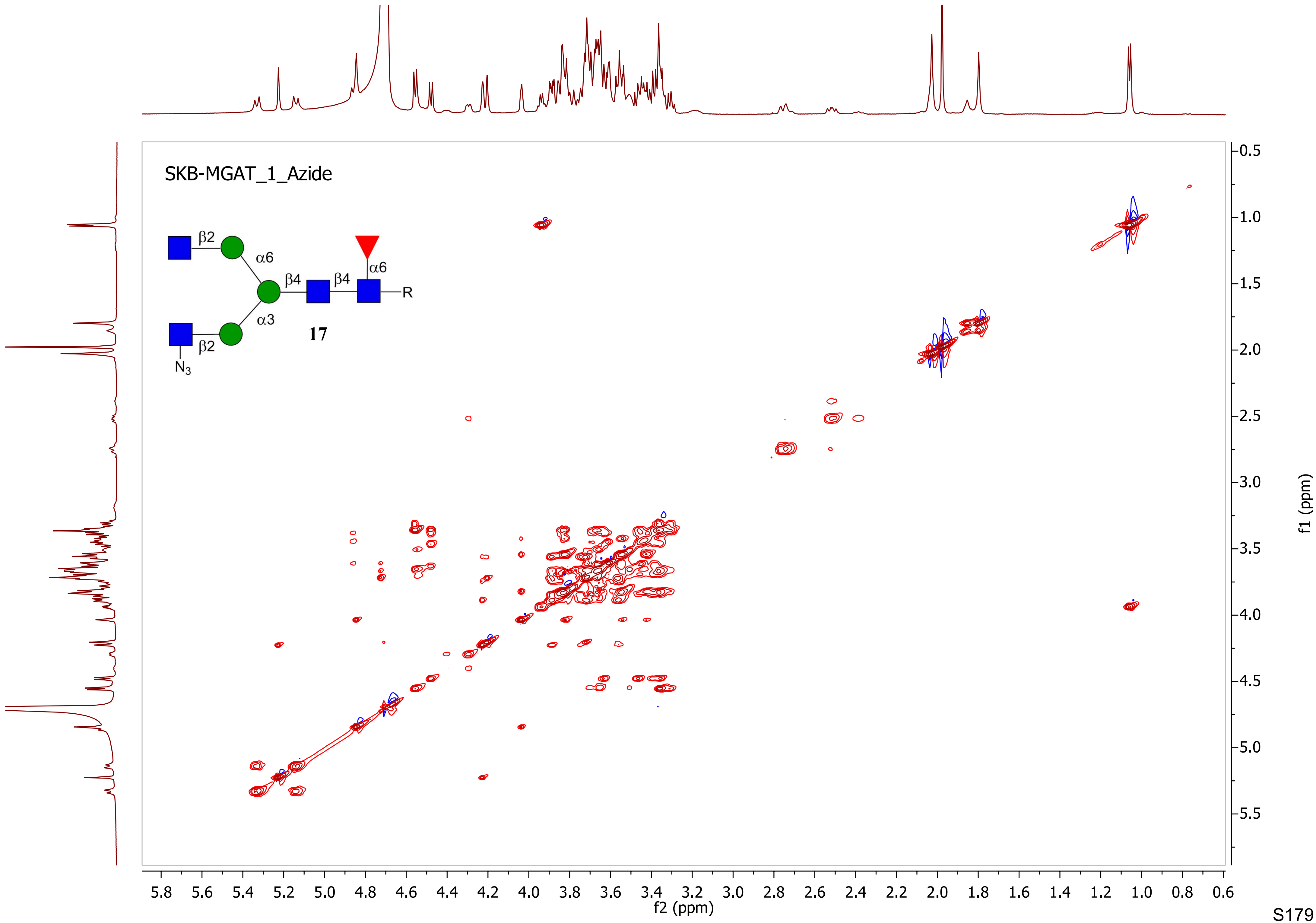

SKB-MGAT_1_Azide
β2
α6
β4
β4
α6
R
α3
17
β2
N3
f2 (ppm)
f1 (ppm)

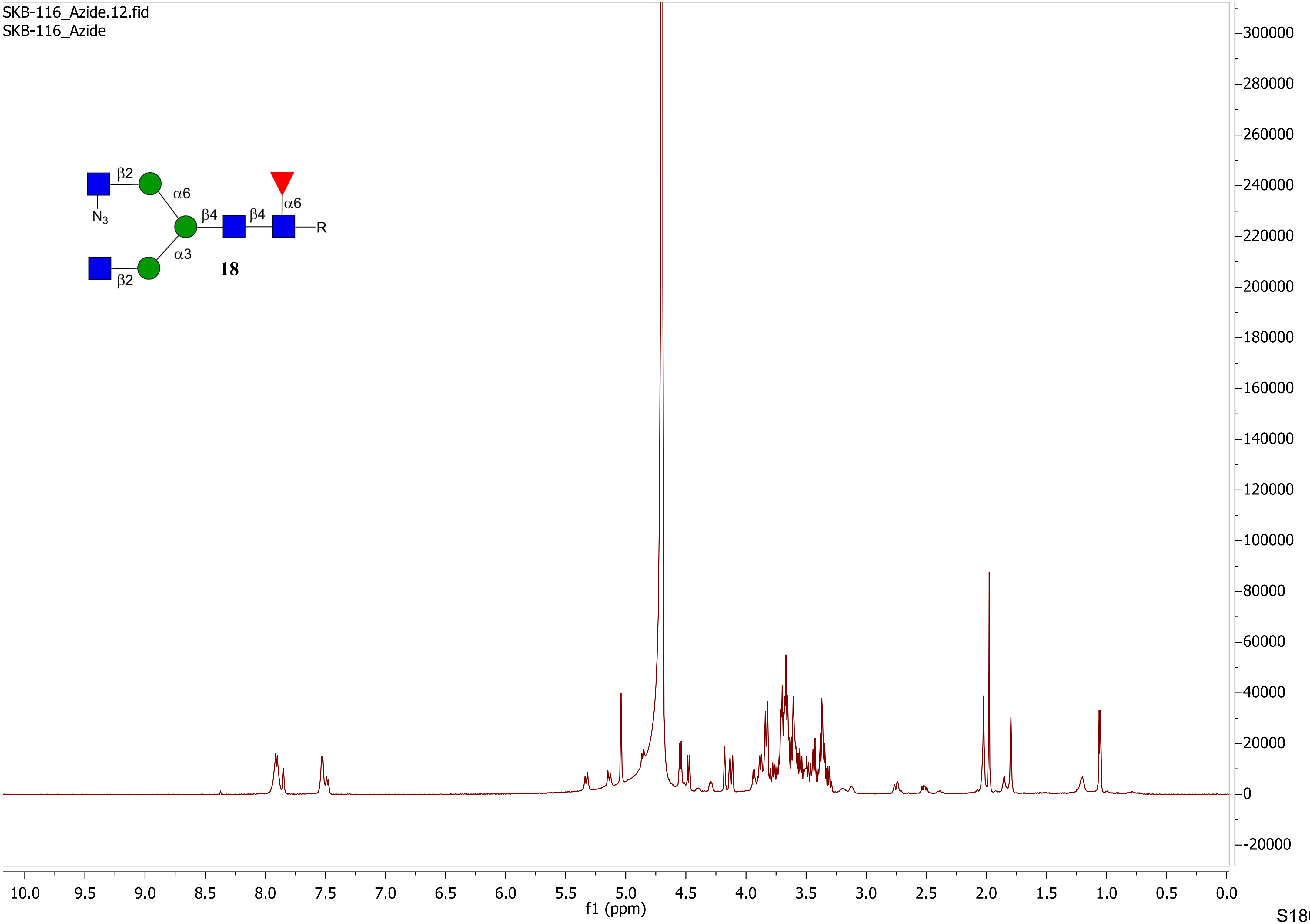
SKB-116_Azide.12.fid
SKB-116_Azide
β2
α6
N3
β4
β4
α6
R
α3
β2
18
f1 (ppm)

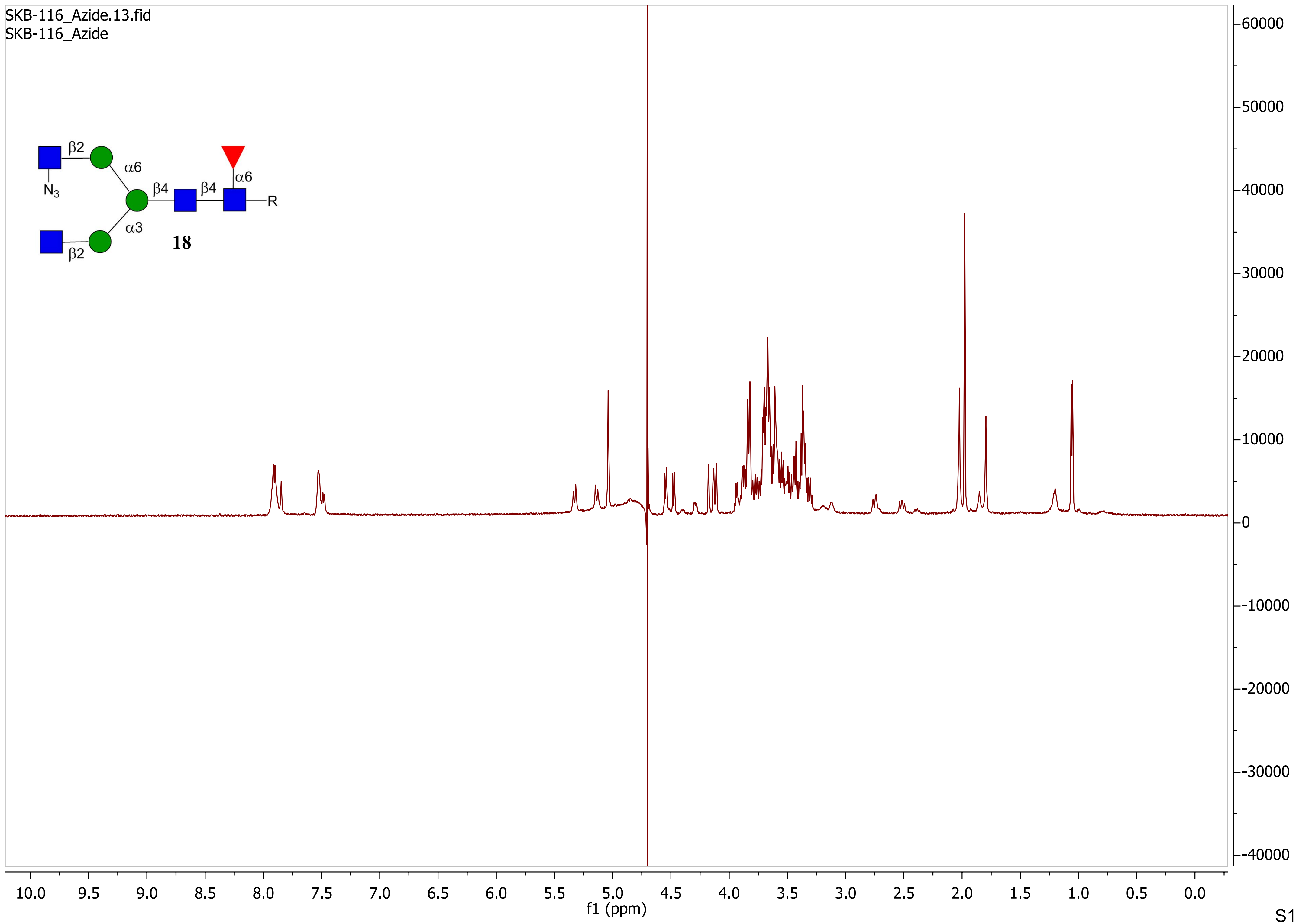

SKB-116_Azide.13.fid
SKB-116_Azide
β2
α6
N3
β4
β4
α6
R
α3
18
β2
60000
50000
40000
30000
20000
10000
0
-10000
-20000
-30000
-40000
10.0 9.5 9.0 8.5 8.0 7.5 7.0 6.5 6.0 5.5 5.0 4.5 4.0 3.5 3.0 2.5 2.0 1.5 1.0 0.5 0.0
f1 (ppm)

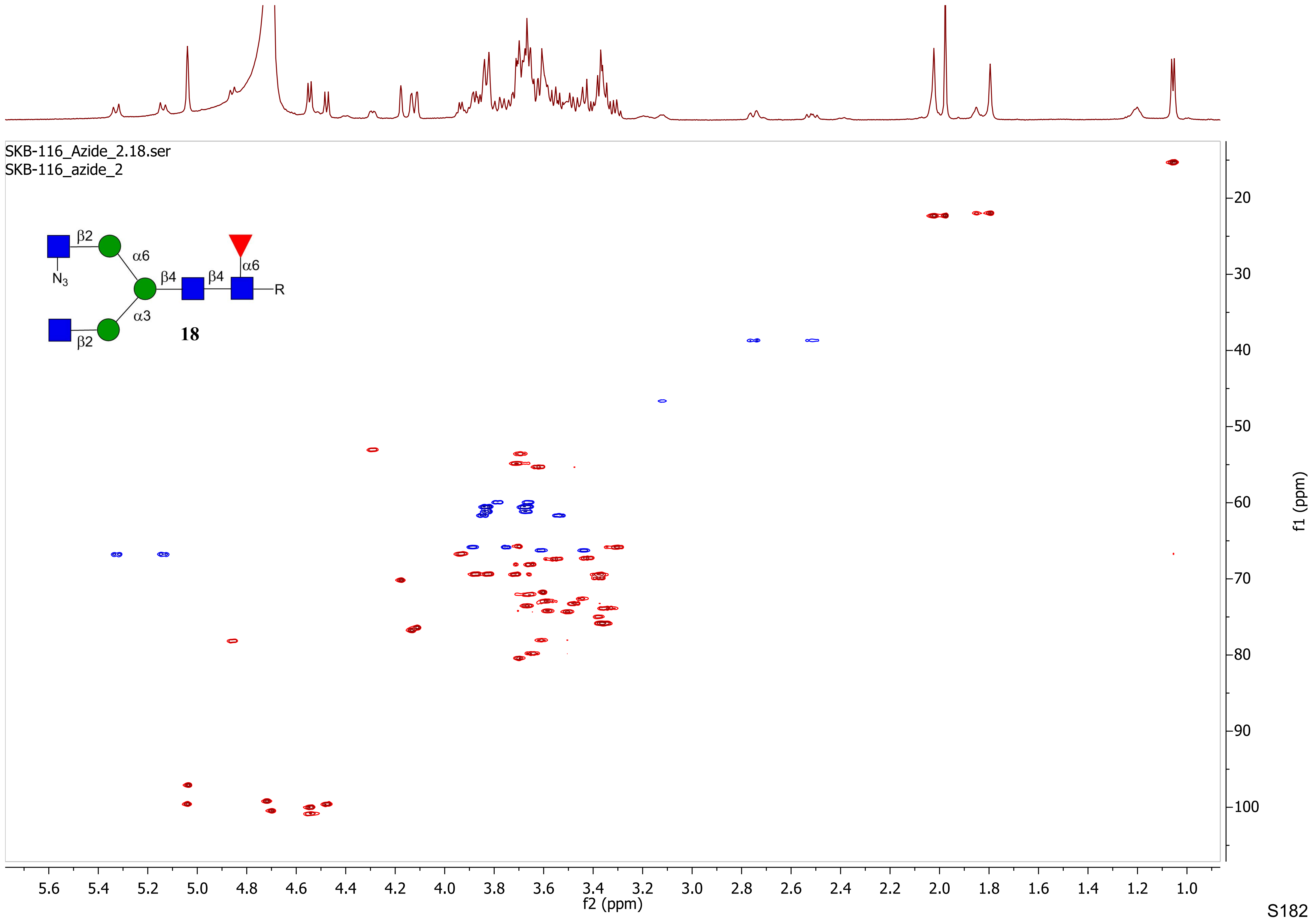
SKB-116_Azide_2.18.ser
SKB-116_azide_2
β2
α6
N3
β4
β4
α6
R
α3
β2
18
f1 (ppm)
f2 (ppm)

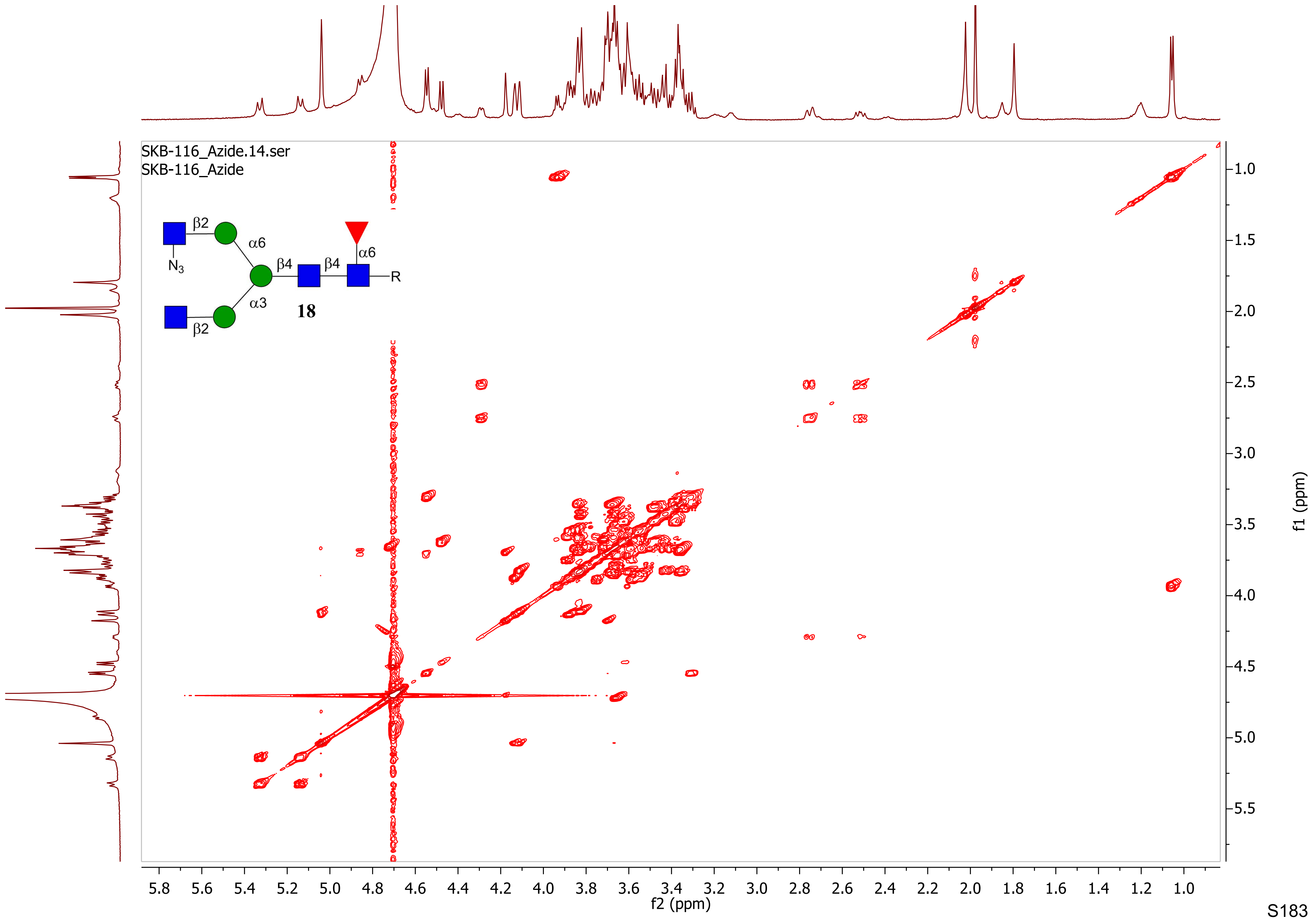

SKB-116_Azide.14.ser
SKB-116_Azide
β2
α6
N3
β4
β4
α6
R
α3
18
β2
f2 (ppm)
f1 (ppm)

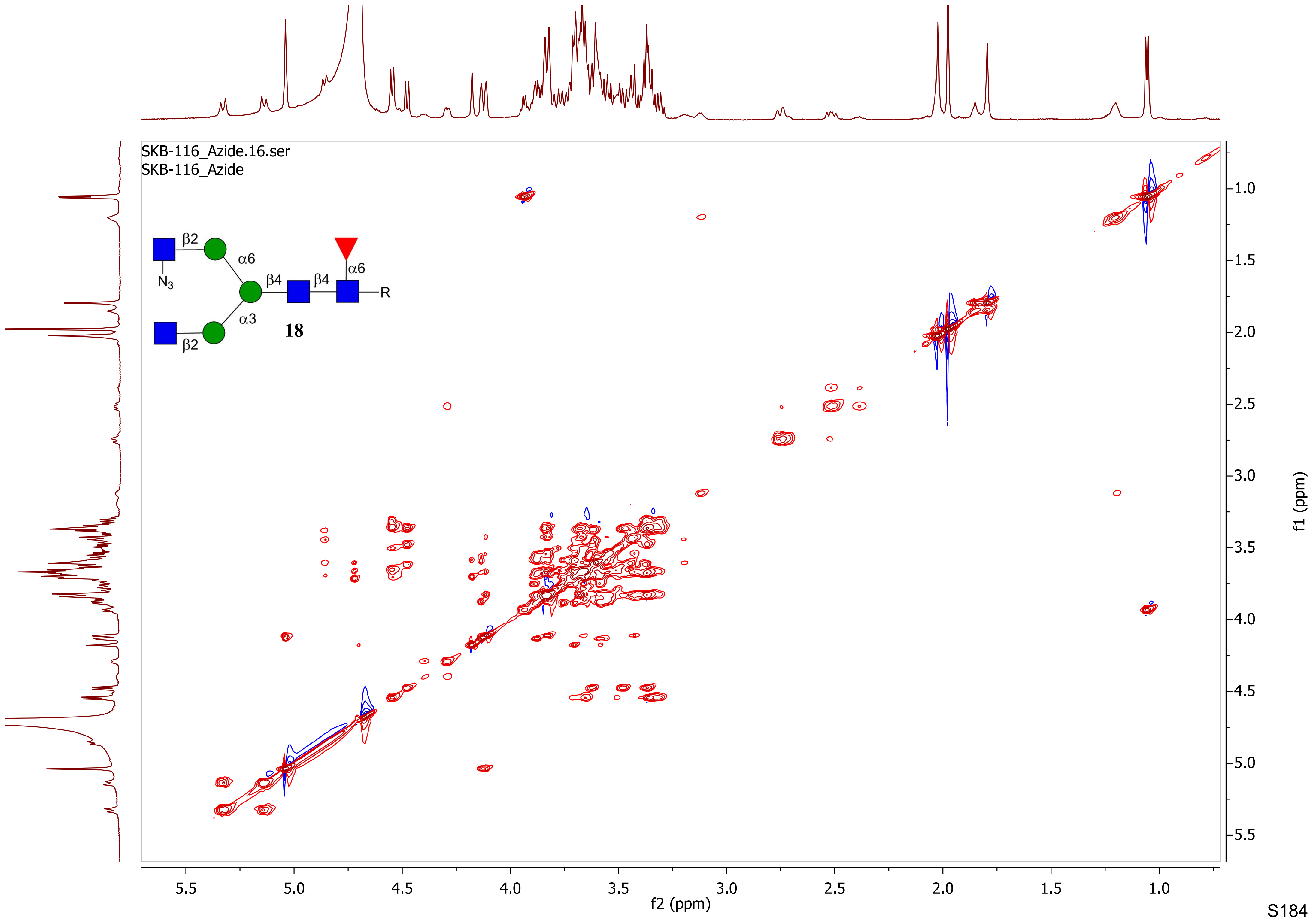
SKB-116_Azide.16.ser
SKB-116_Azide
β2
α6
N3
β4
β4
α6
R
α3
18
β2
f2 (ppm)
f1 (ppm)

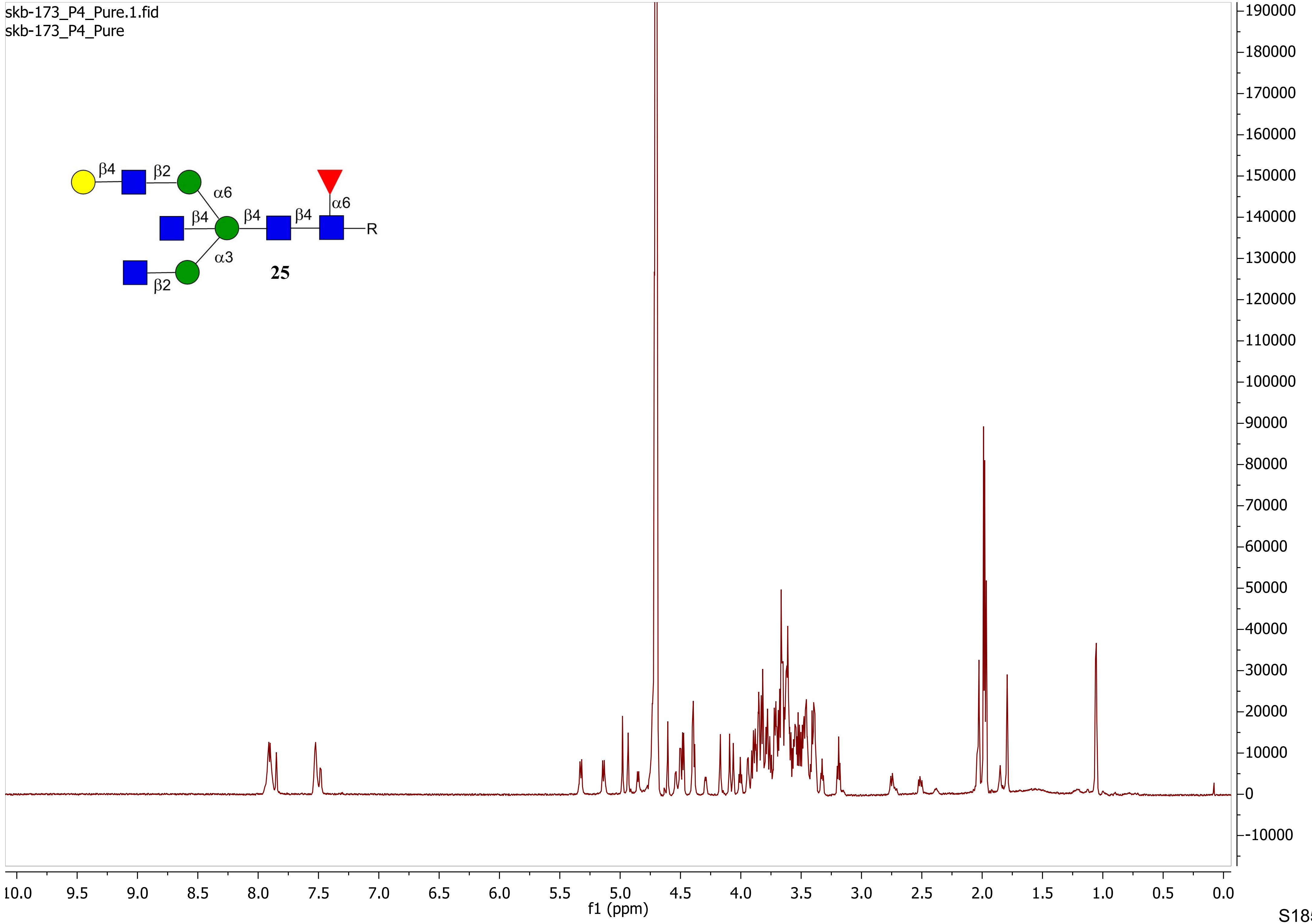
skb-173_P4_Pure.1.fid
skb-173_P4_Pure
β4
β2
α6
β4
β4
β4
α6
R
α3
β2
25
f1 (ppm)

skb-173_P4_Pure.2.fid
skb-173_P4_Pure

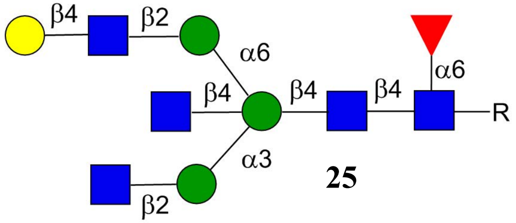

**25**

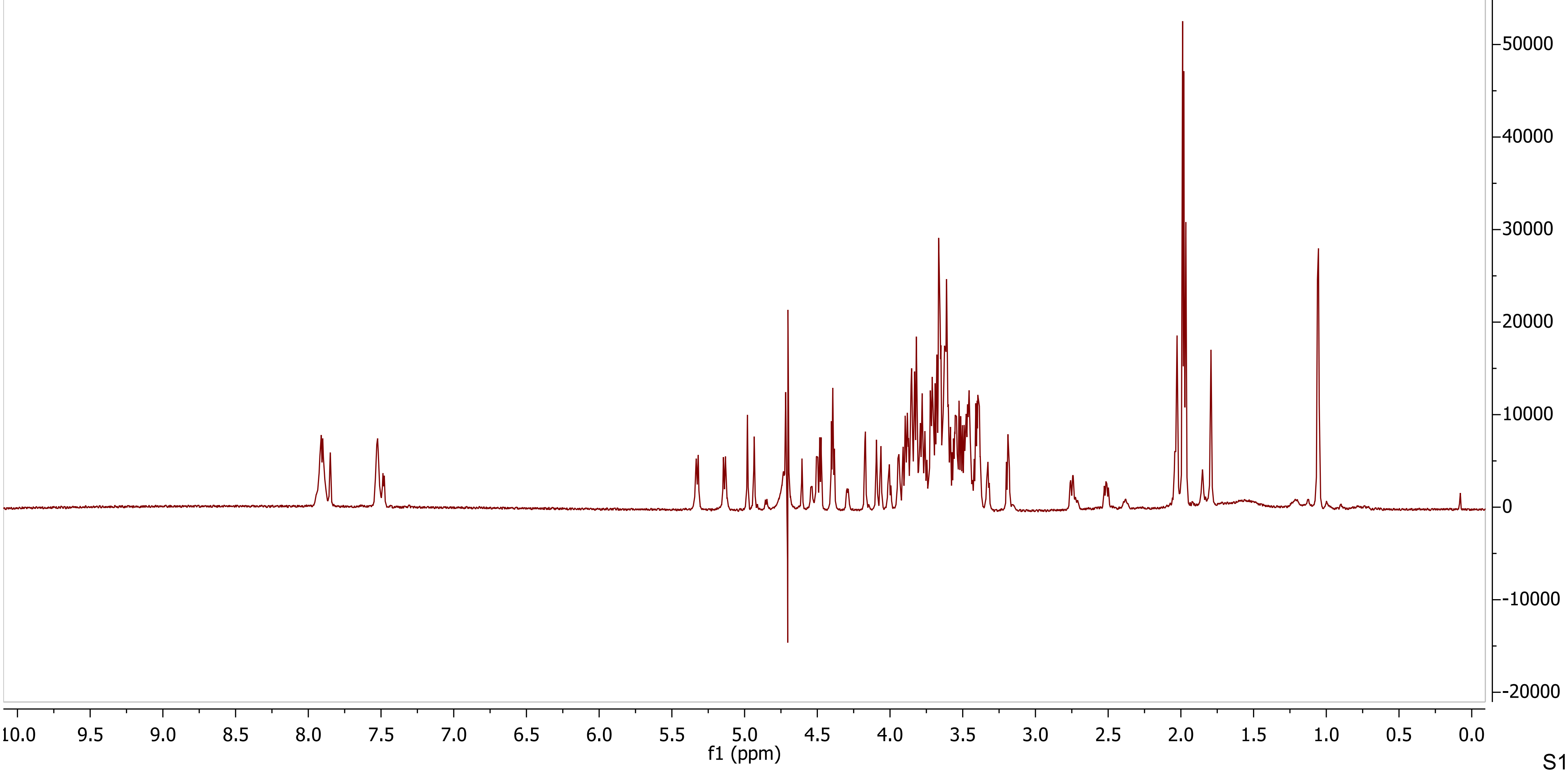

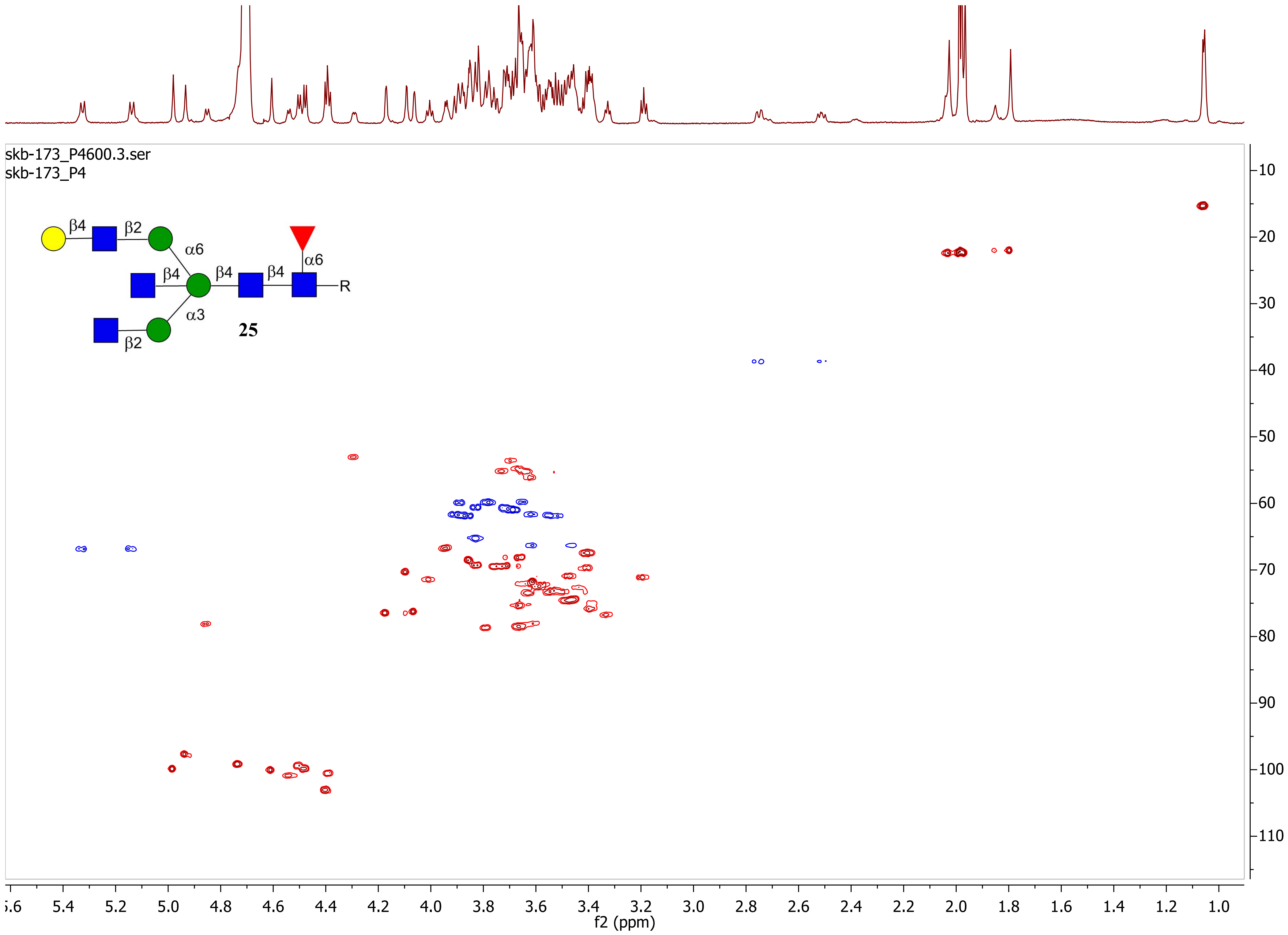
skb-173_P4600.3.ser
skb-173_P4
β4
β2
α6
β4
β4
β4
α6
R
α3
β2
25
f2 (ppm)
f1 (ppm)

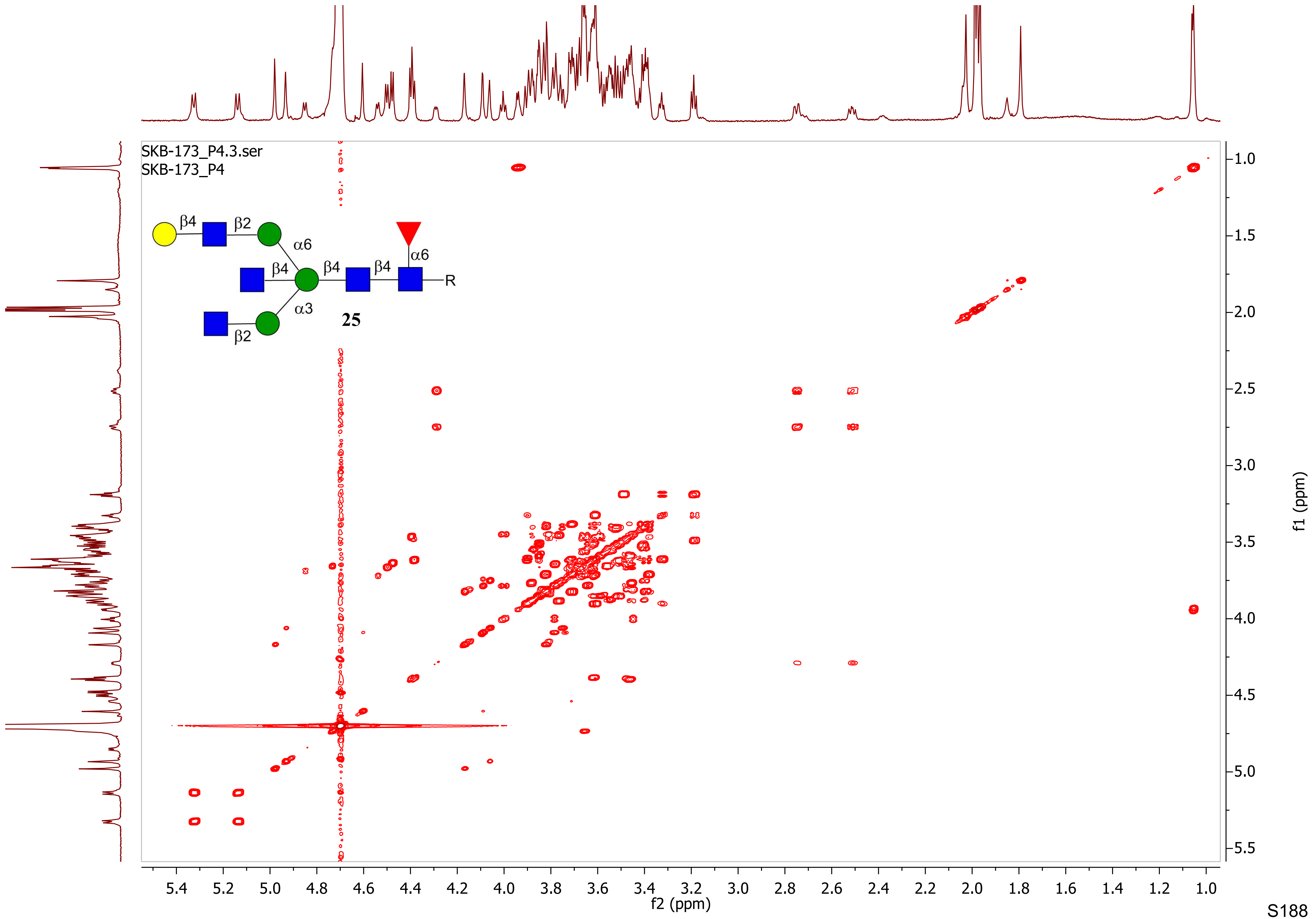

SKB-173_P4.3.ser
SKB-173_P4
β4
β2
α6
β4
β4
β4
α6
R
α3
25
β2
f1 (ppm)
f2 (ppm)

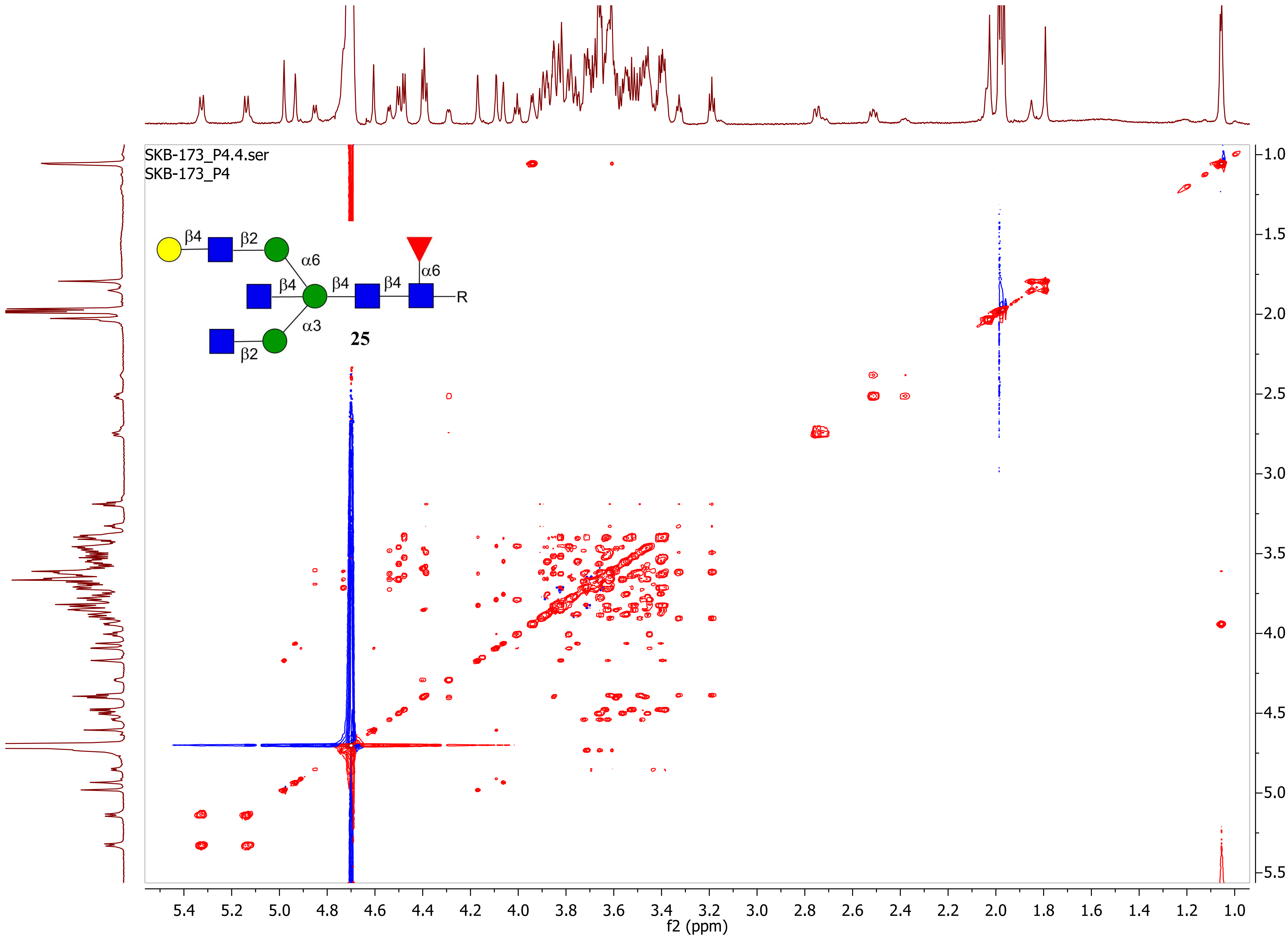
SKB-173_P4.4.ser
SKB-173_P4
β4
β2
α6
β4
β4
β4
α6
R
α3
β2
25
f2 (ppm)
f1 (ppm)

skb-154_P4_2.1.fid
skb-154_P4_2

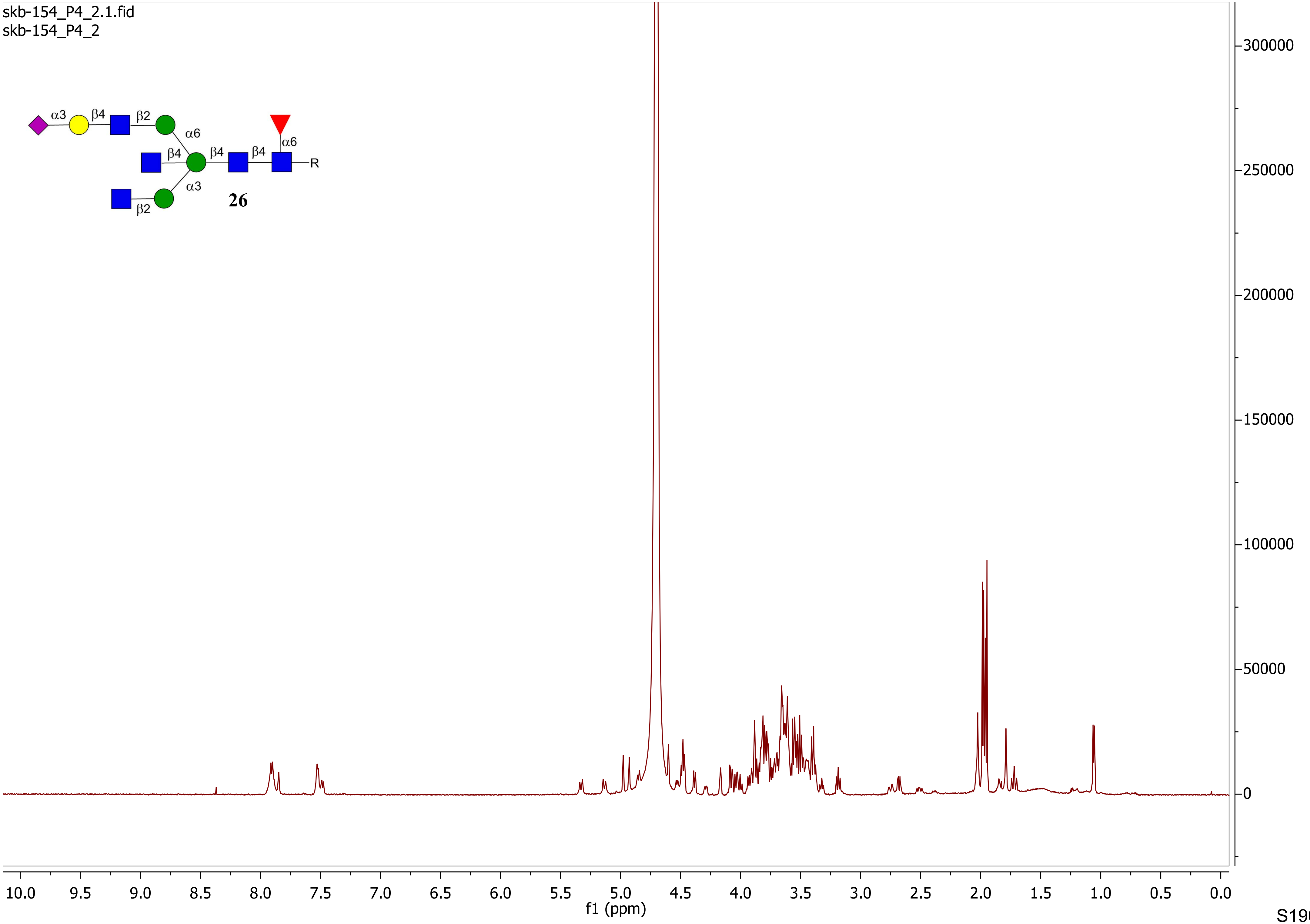

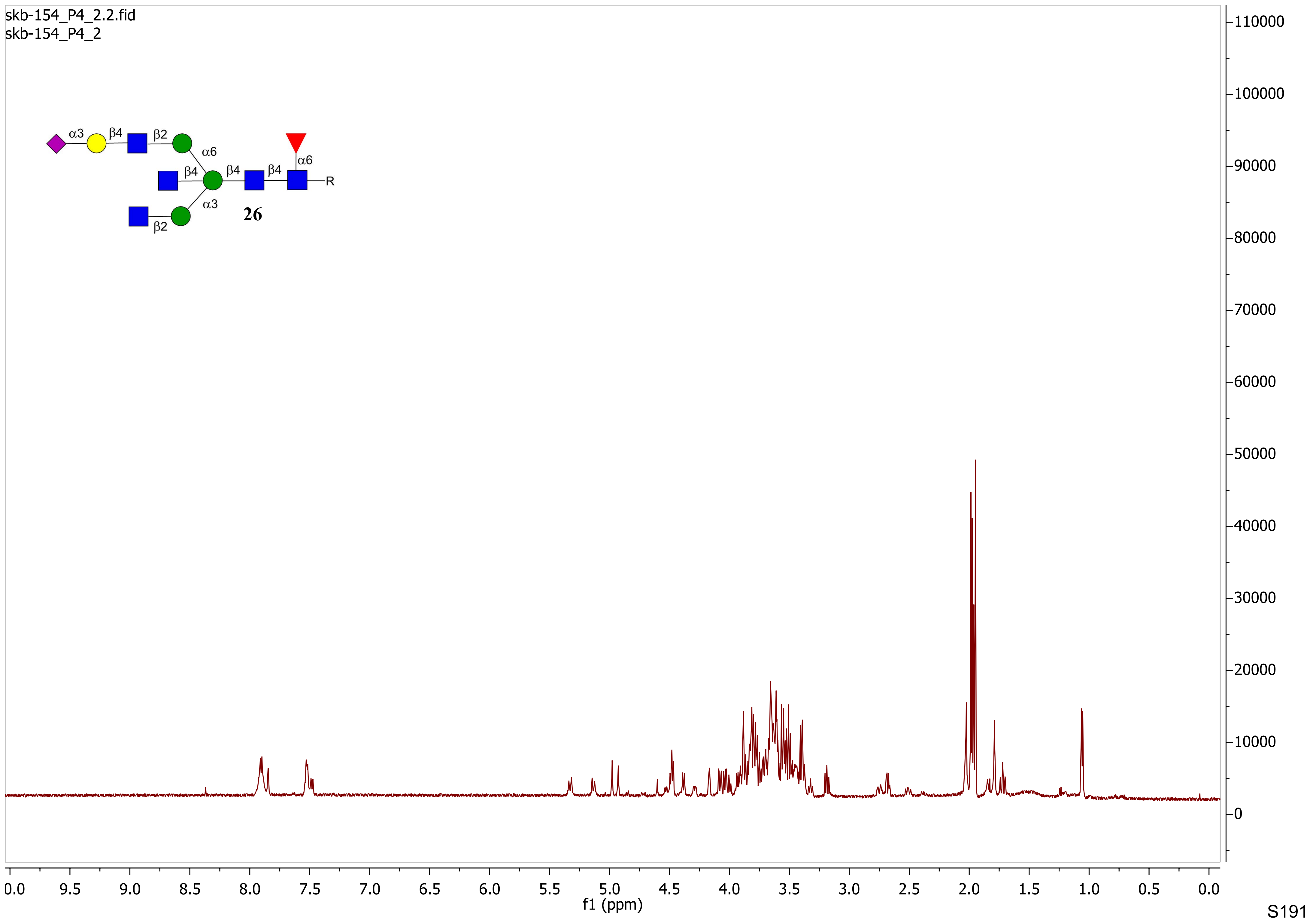
skb-154_P4_2.2.fid
skb-154_P4_2
α3
β4
β2
α6
β4
β4
β4
α6
R
α3
β2
26
110000
100000
90000
80000
70000
60000
50000
40000
30000
20000
10000
0
0.0
9.5
9.0
8.5
8.0
7.5
7.0
6.5
6.0
5.5
5.0
4.5
4.0
3.5
3.0
2.5
2.0
1.5
1.0
0.5
0.0
f1 (ppm)

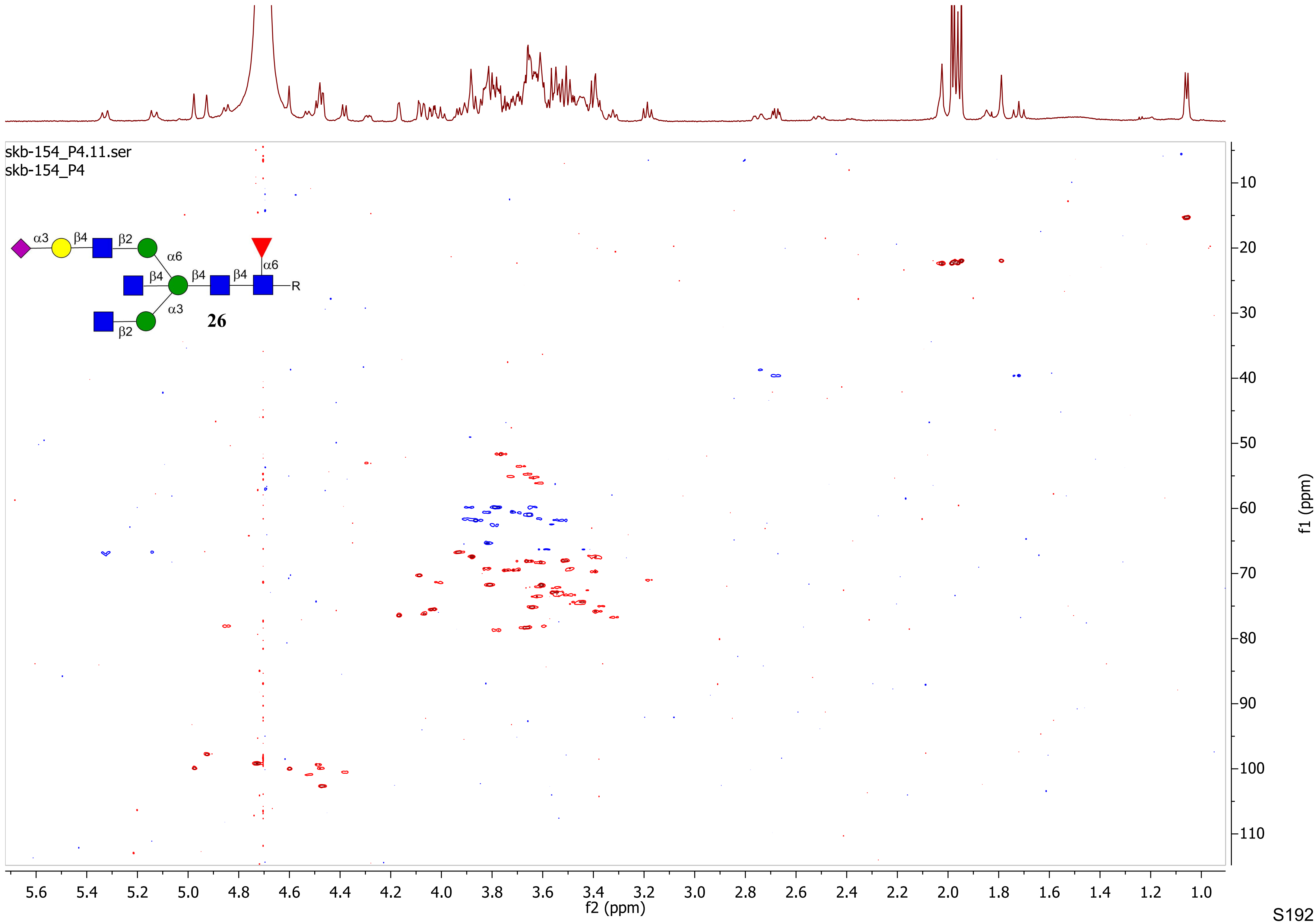
skb-154_P4.11.ser
skb-154_P4
α3
β4
β2
α6
β4
β4
β4
α6
R
α3
β2
26
f2 (ppm)
f1 (ppm)

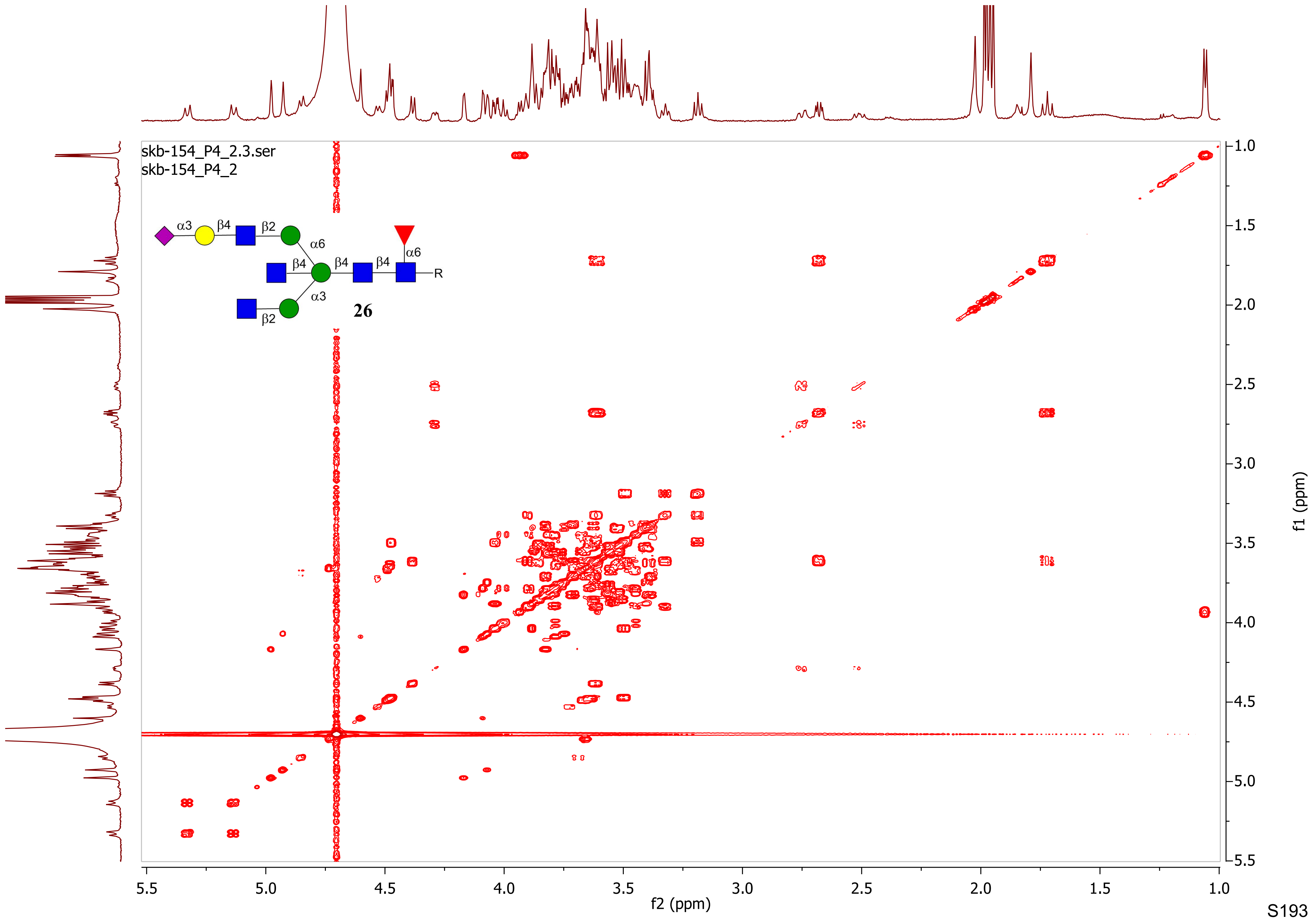
skb-154_P4_2.3.ser
skb-154_P4_2
α3
β4
β2
α6
β4
β4
β4
α6
R
α3
β2
26
f2 (ppm)
f1 (ppm)

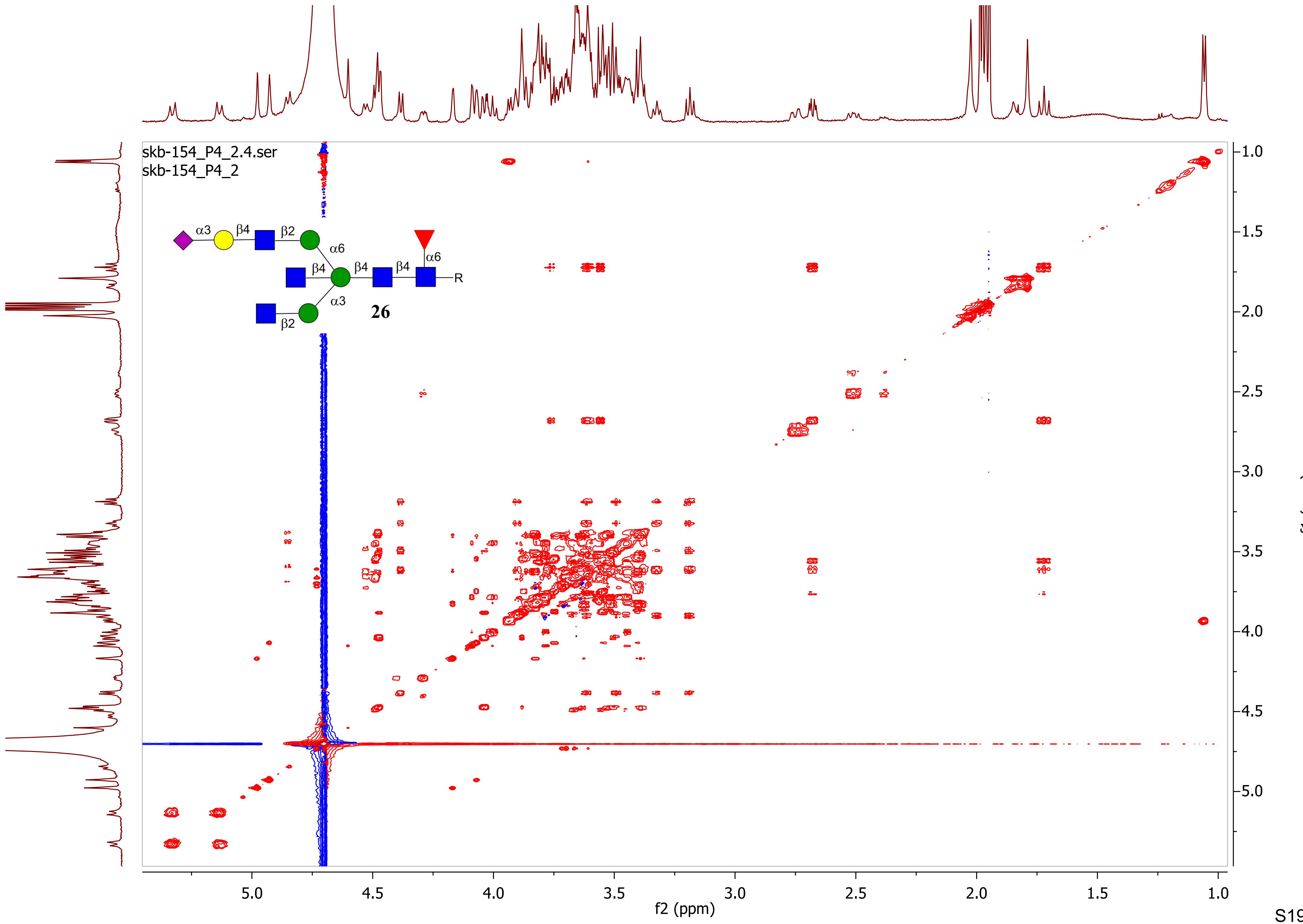
skb-154_P4_2.4.ser
skb-154_P4_2
α3
β4
β2
α6
β4
β4
β4
α6
R
α3
β2
26
f2 (ppm)
f1 (ppm)

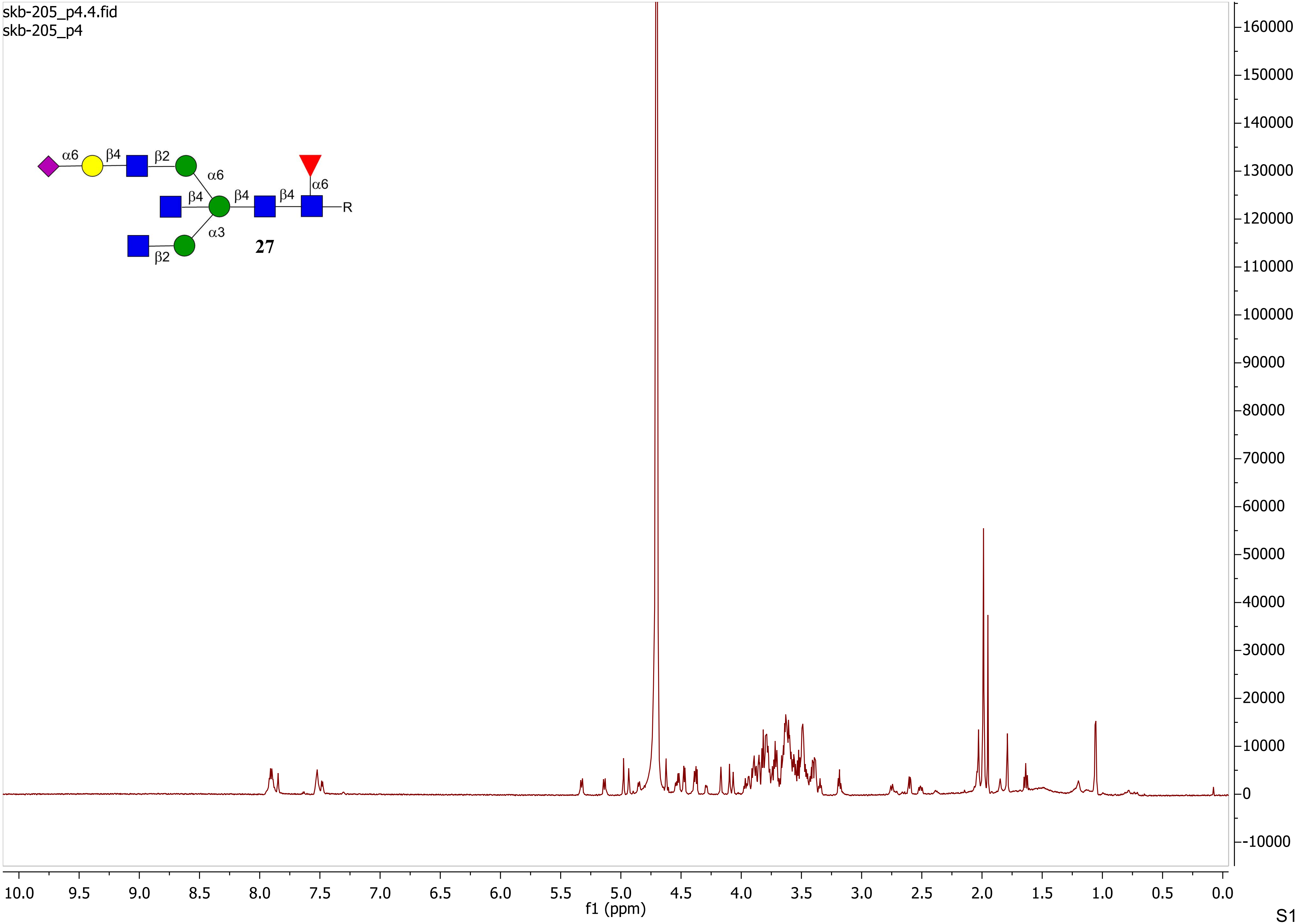
skb-205_p4.4.fid
skb-205_p4
α6
β4
β2
α6
β4
β4
β4
α6
R
α3
β2
27
160000
150000
140000
130000
120000
110000
100000
90000
80000
70000
60000
50000
40000
30000
20000
10000
0
-10000
10.0 9.5 9.0 8.5 8.0 7.5 7.0 6.5 6.0 5.5 5.0 4.5 4.0 3.5 3.0 2.5 2.0 1.5 1.0 0.5 0.0
f1 (ppm)

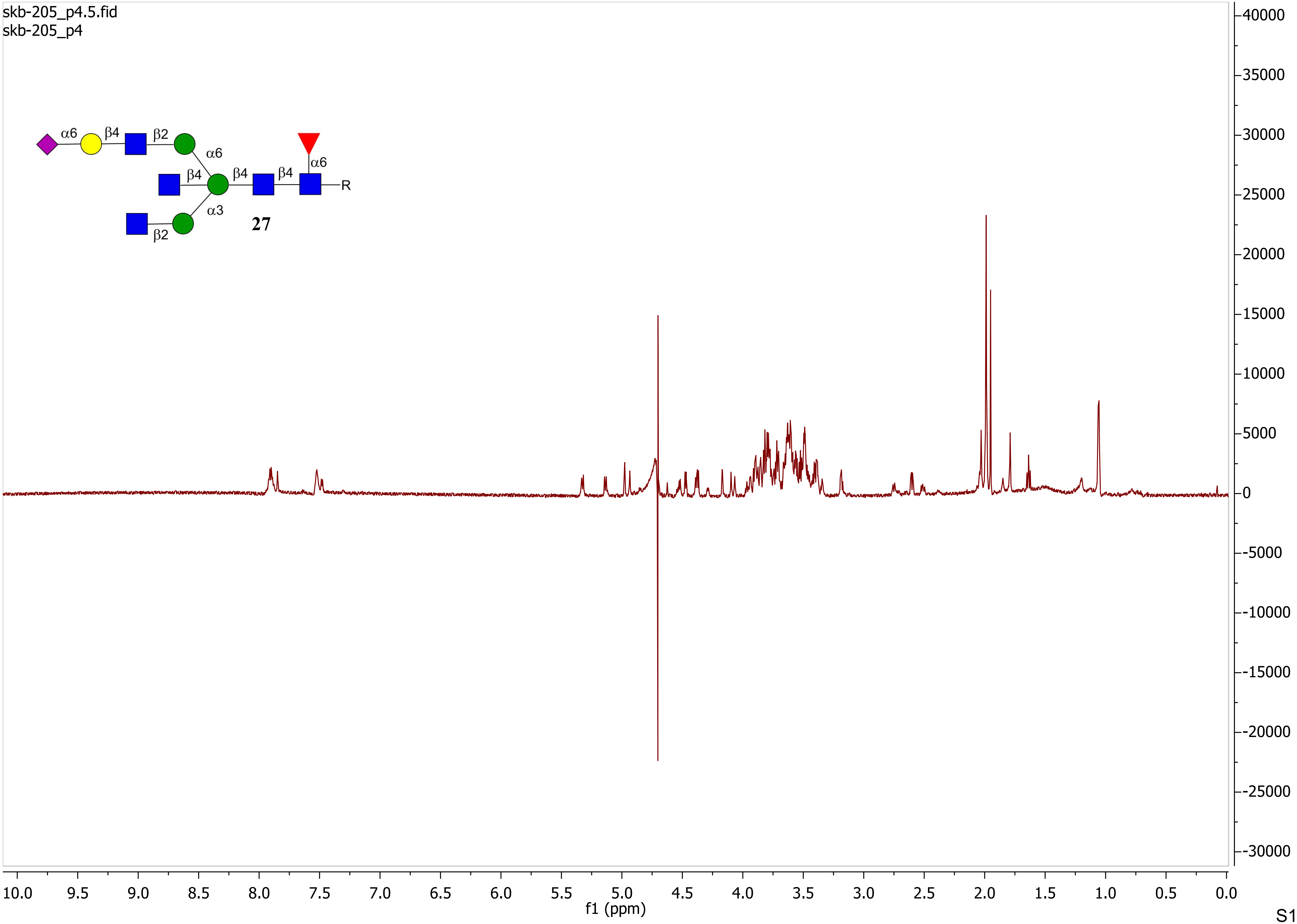
skb-205_p4.5.fid
skb-205_p4
α6
β4
β2
α6
β4
β4
β4
α6
R
α3
β2
27
f1 (ppm)

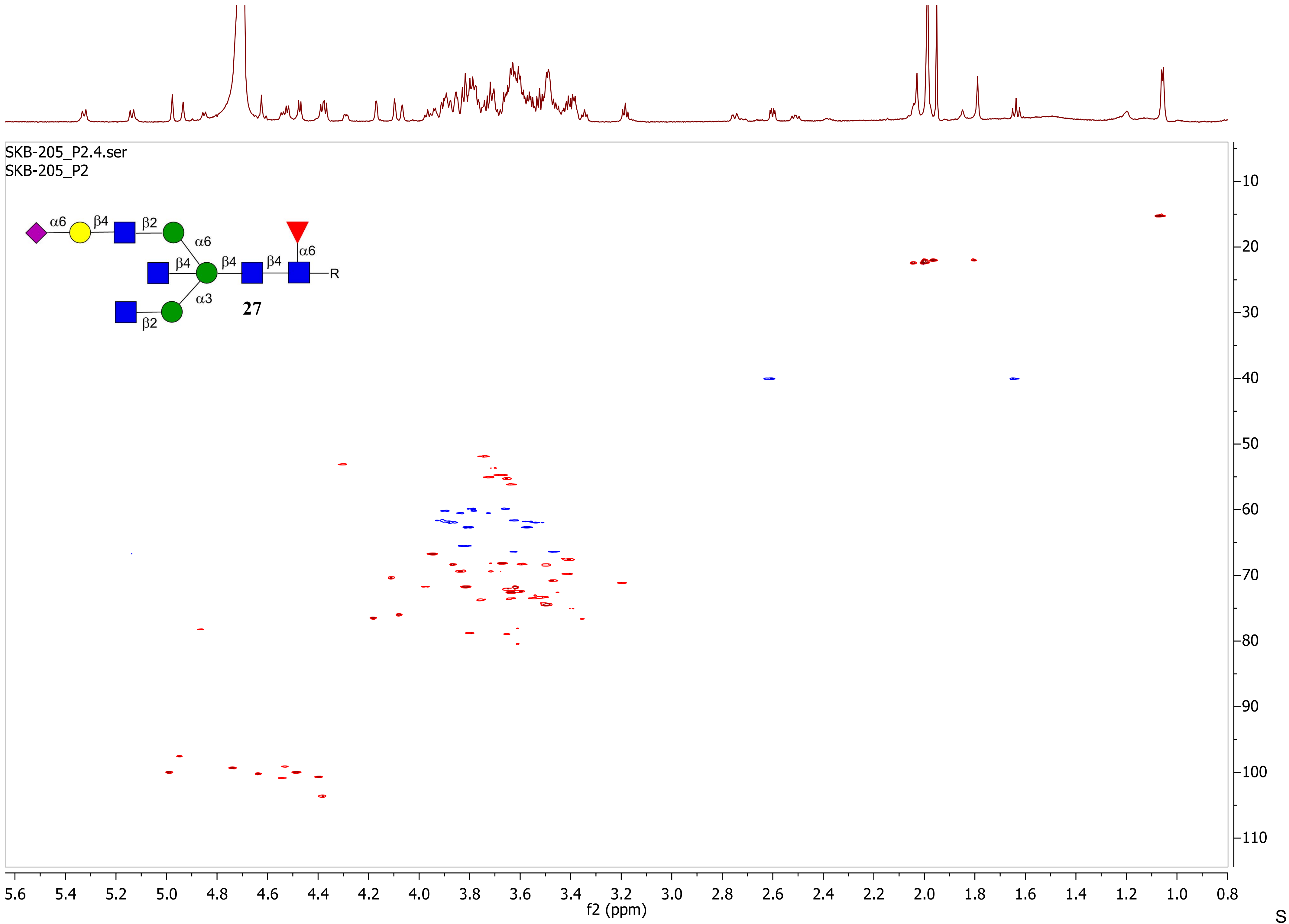

SKB-205_P2.4.ser
SKB-205_P2
α6
β4
β2
α6
β4
β4
β4
α6
R
α3
27
β2
f2 (ppm)
f1 (ppm)

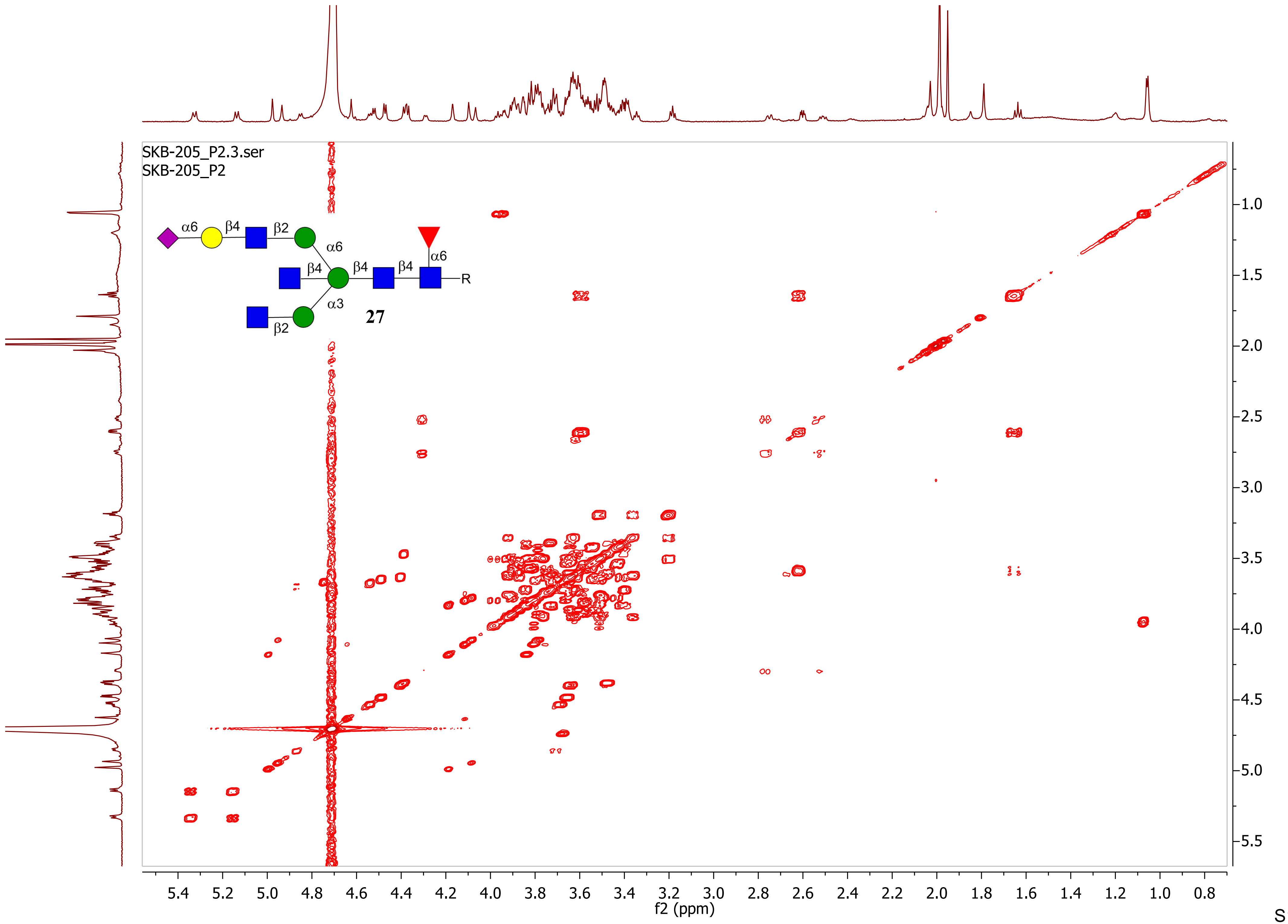

SKB-205_P2.3.ser
SKB-205_P2
α6
β4
β2
α6
β4
β4
β4
α6
R
α3
β2
27
f2 (ppm)
f1 (ppm)

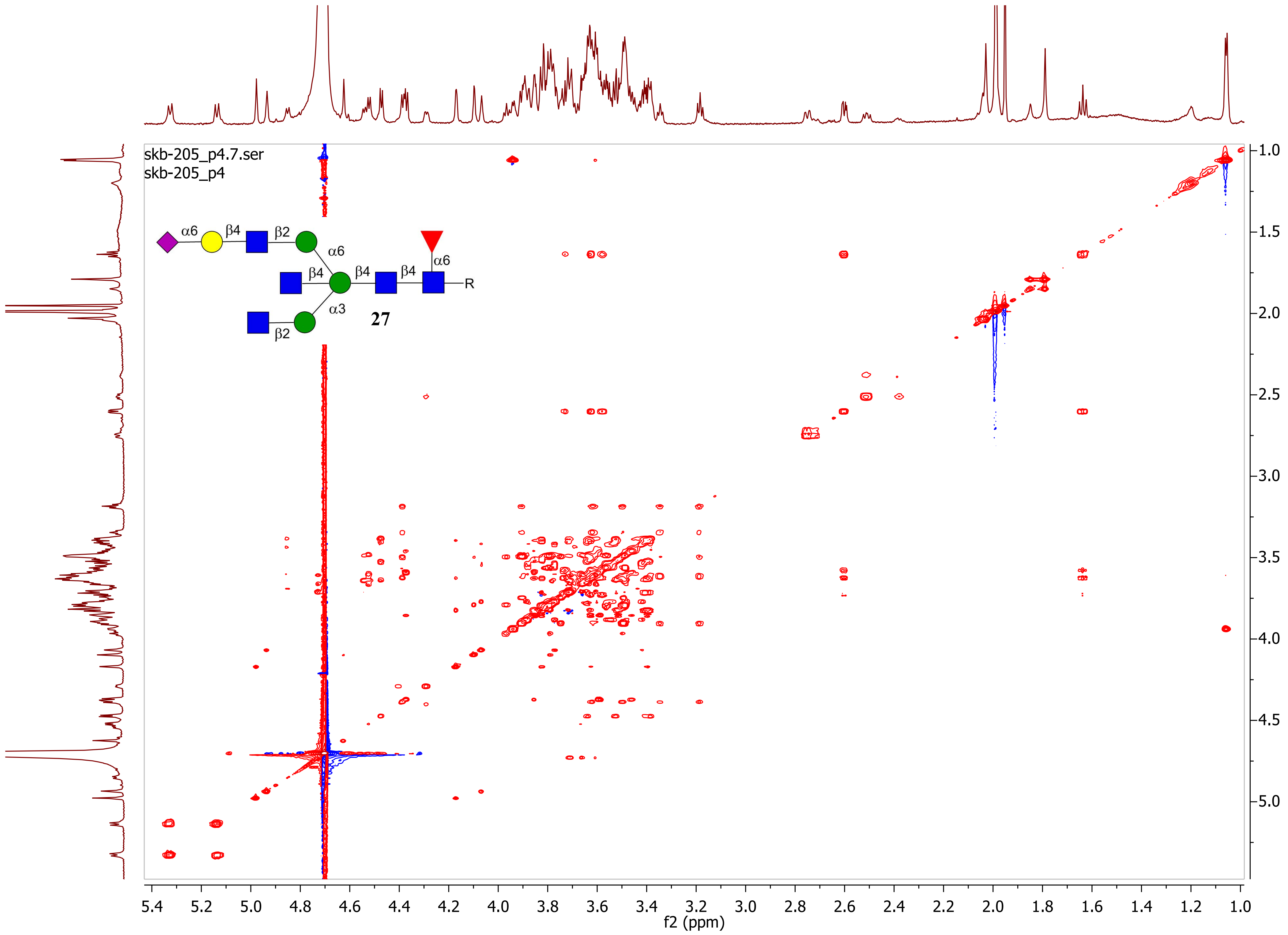
skb-205_p4.7.ser
skb-205_p4
α6
β4
β2
α6
β4
β4
β4
α6
R
α3
β2
27
f2 (ppm)
f1 (ppm)

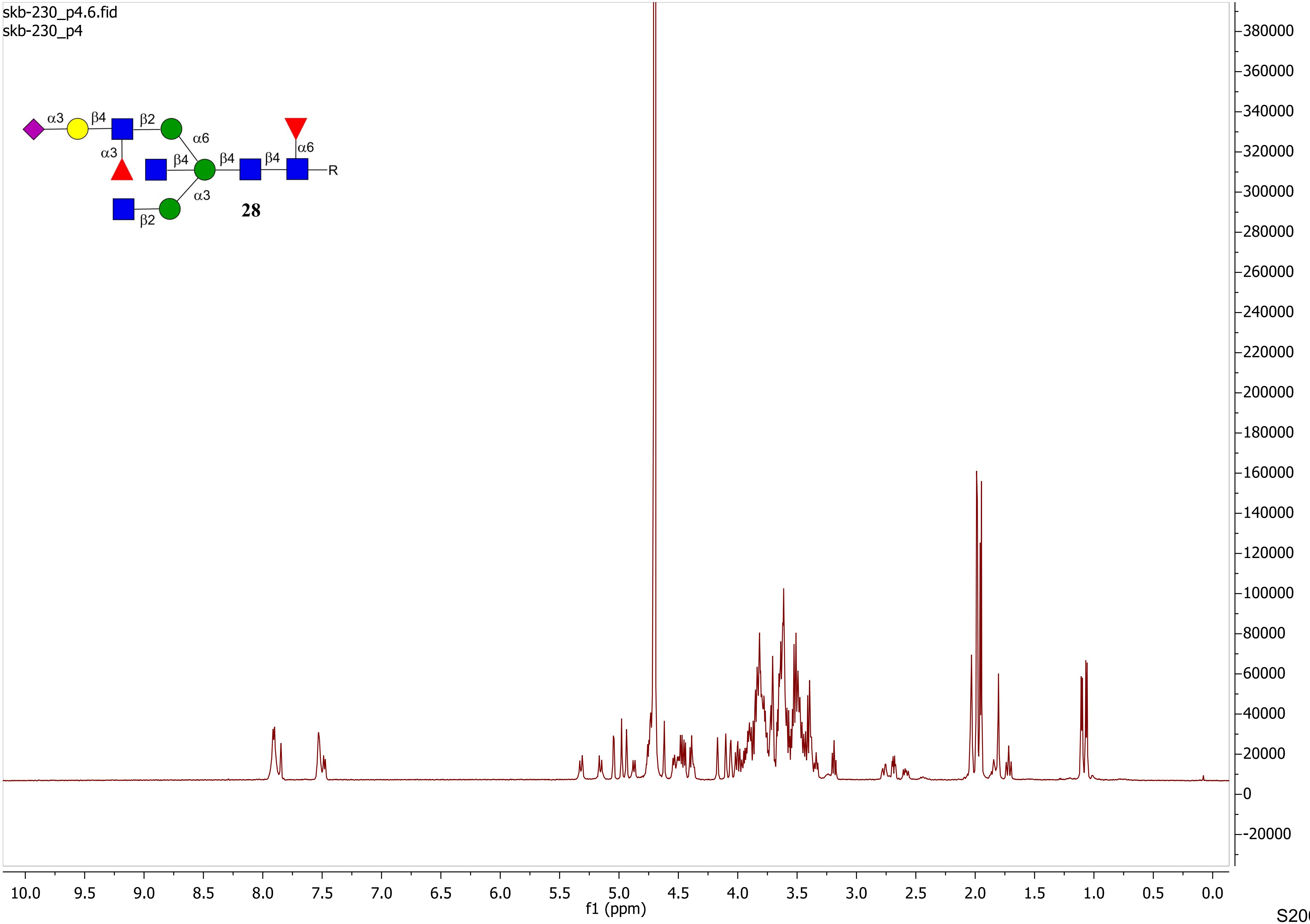

skb-230_p4.6.fid
skb-230_p4
α3
β4
β2
α6
α3
β4
β4
β4
α6
R
α3
β2
28
f1 (ppm)

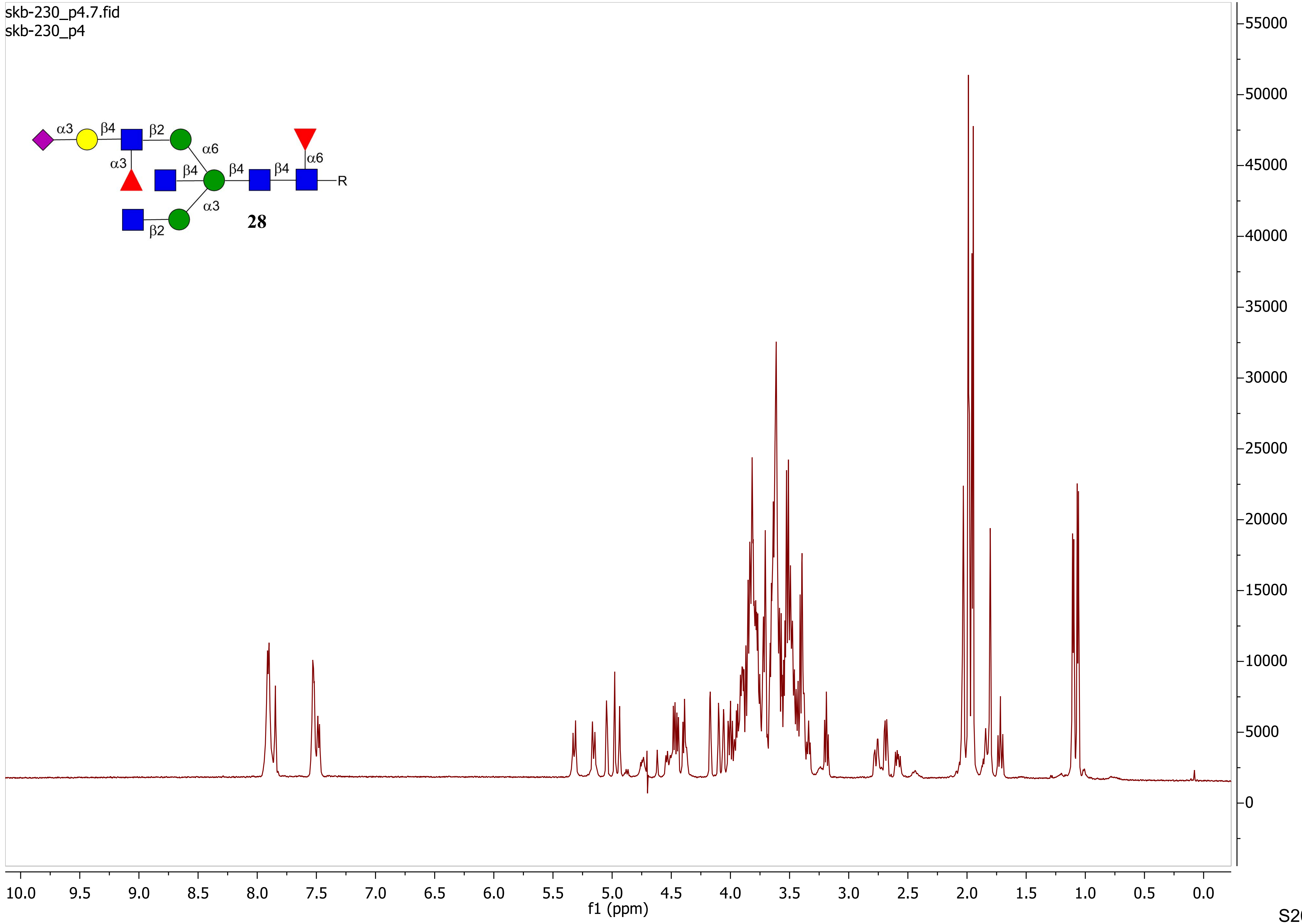
skb-230_p4.7.fid
skb-230_p4
α3
β4
β2
α6
α3
β4
β4
β4
α6
R
α3
β2
28
55000
50000
45000
40000
35000
30000
25000
20000
15000
10000
5000
0
10.0
9.5
9.0
8.5
8.0
7.5
7.0
6.5
6.0
5.5
5.0
4.5
4.0
3.5
3.0
2.5
2.0
1.5
1.0
0.5
0.0
f1 (ppm)

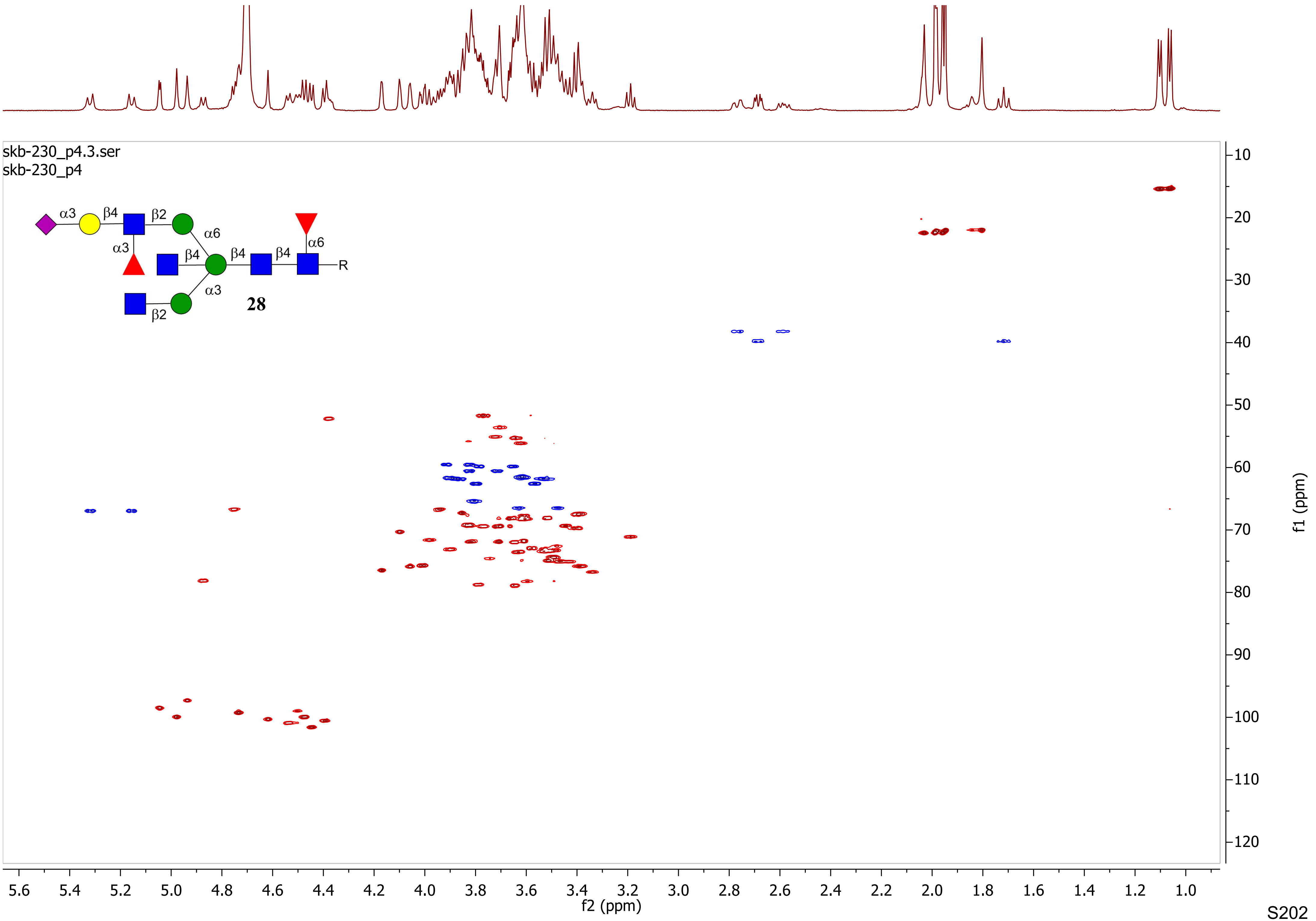
skb-230_p4.3.ser
skb-230_p4
α3
β4
β2
α6
α3
β4
β4
β4
α6
R
α3
β2
28
f2 (ppm)
f1 (ppm)

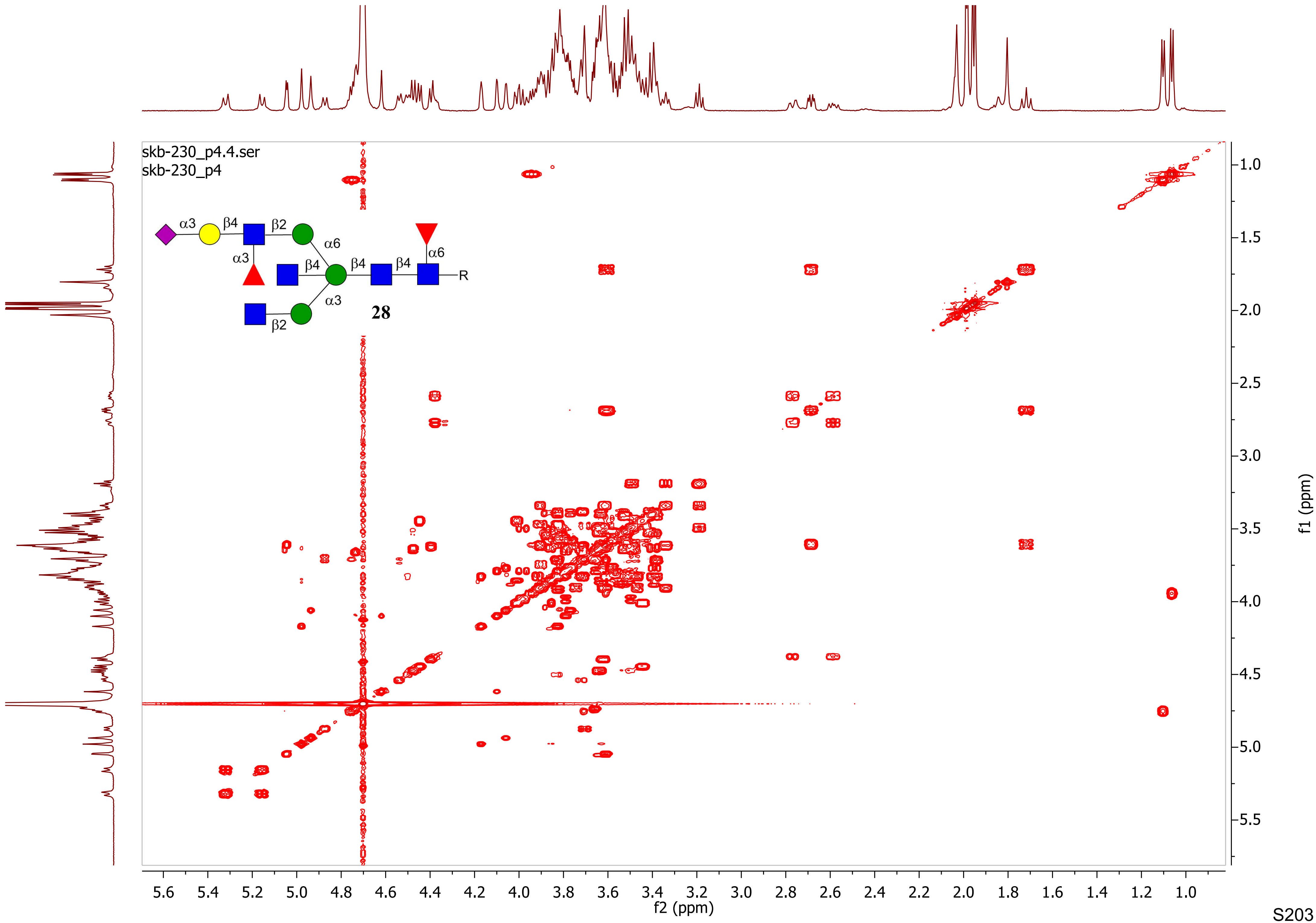
skb-230_p4.4.ser
skb-230_p4
α3
β4
β2
α6
α3
β4
β4
β4
α6
R
α3
β2
28
f2 (ppm)
f1 (ppm)

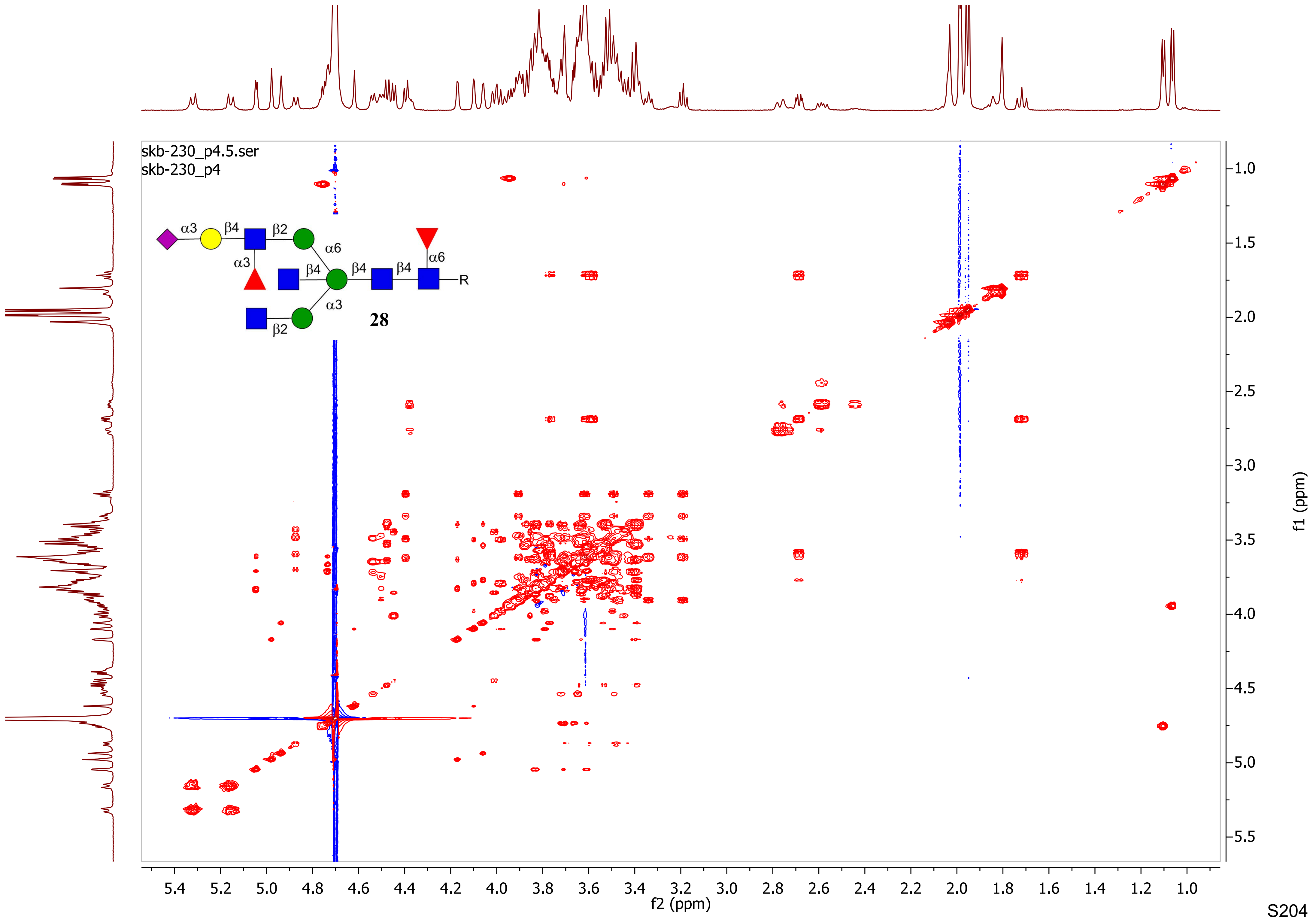
skb-230_p4.5.ser
skb-230_p4
α3
β4
β2
α6
α3
β4
β4
β4
α6
R
α3
β2
28
f2 (ppm)
f1 (ppm)

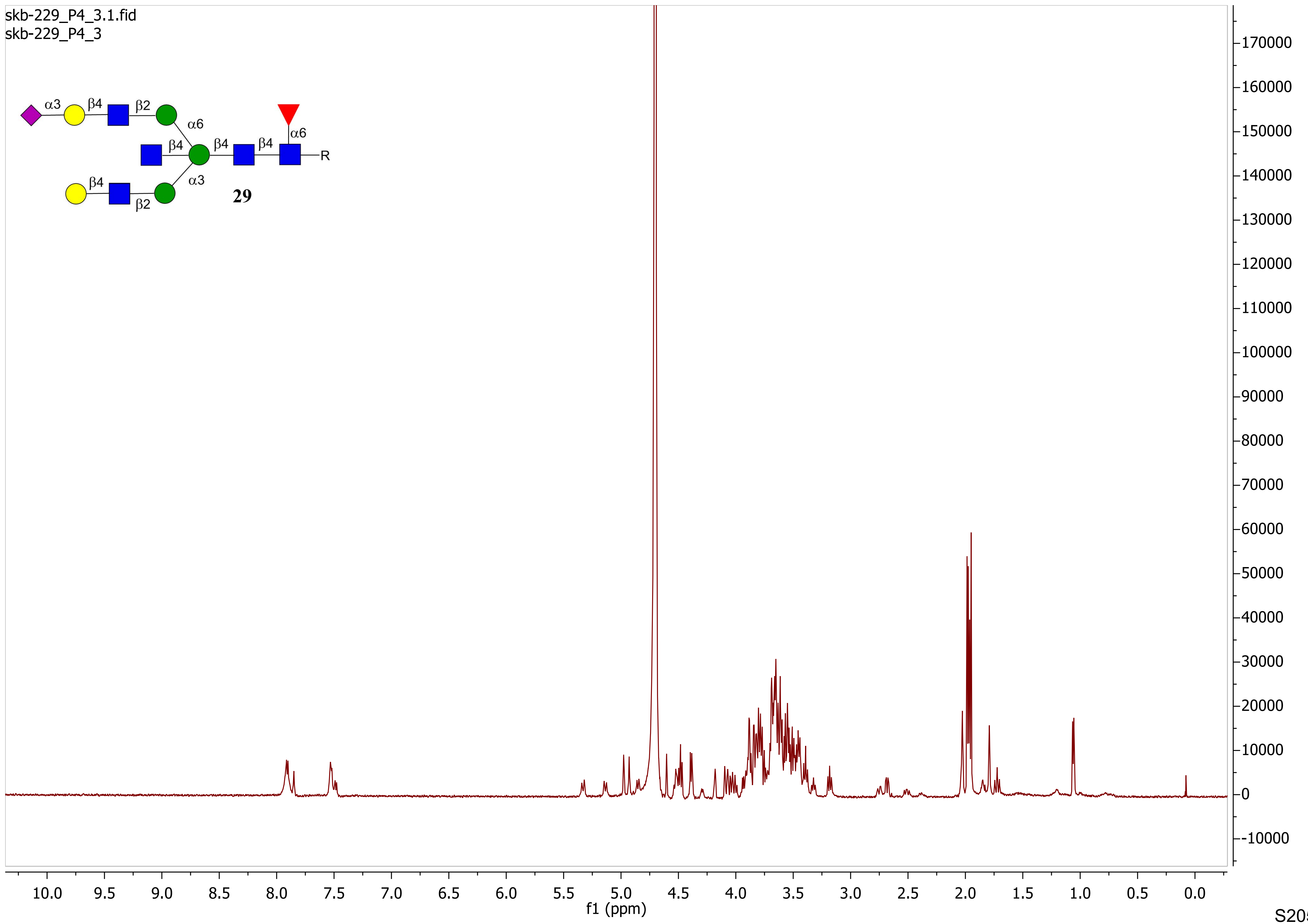

skb-229_P4_3.1.fid
skb-229_P4_3
α3
β4
β2
α6
β4
β4
β4
α6
R
α3
β4
β2
29
170000
160000
150000
140000
130000
120000
110000
100000
90000
80000
70000
60000
50000
40000
30000
20000
10000
0
-10000
10.0 9.5 9.0 8.5 8.0 7.5 7.0 6.5 6.0 5.5 5.0 4.5 4.0 3.5 3.0 2.5 2.0 1.5 1.0 0.5 0.0
f1 (ppm)

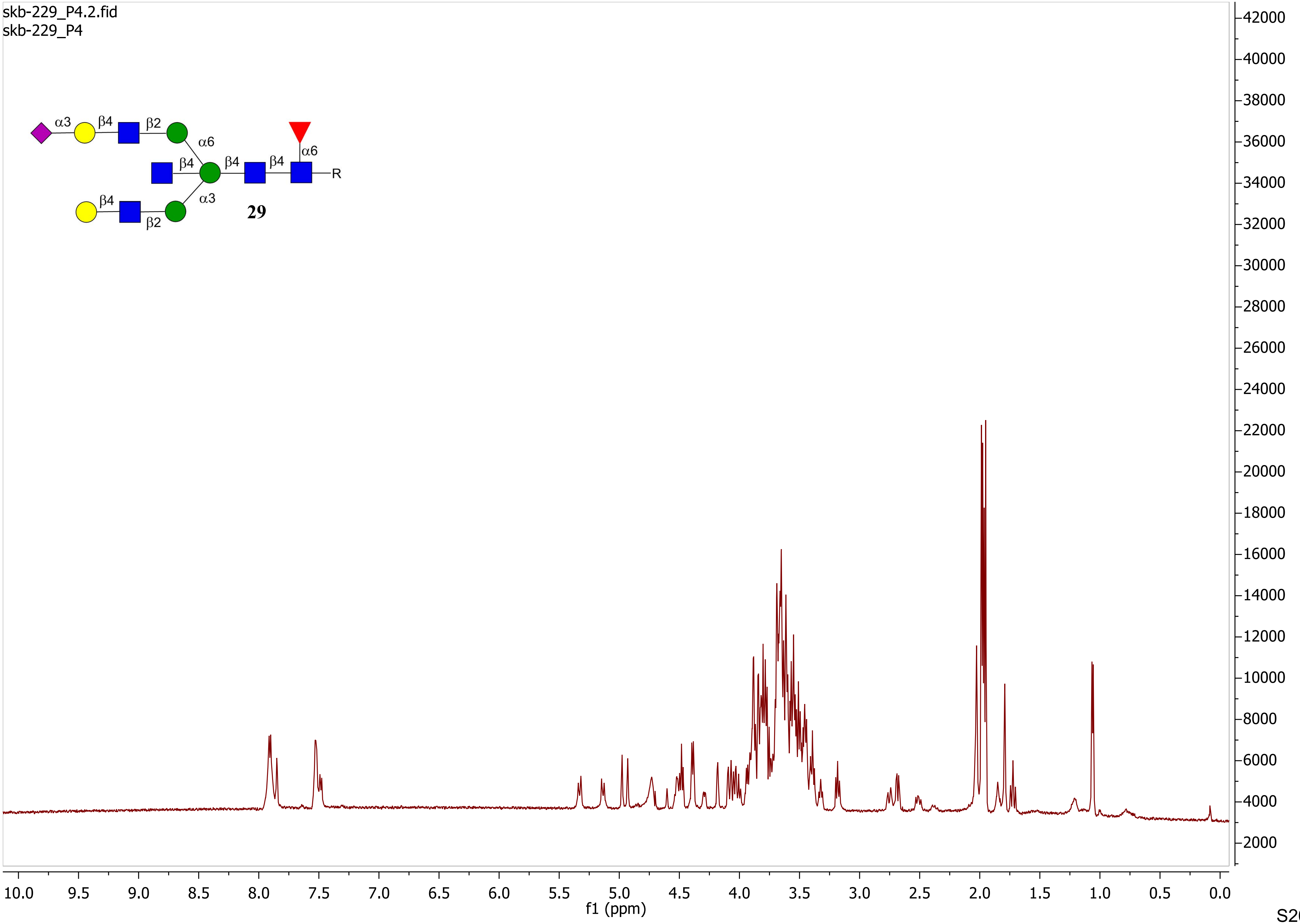
skb-229_P4.2.fid
skb-229_P4
α3
β4
β2
α6
β4
β4
β4
α6
R
β4
β2
α3
29
42000
40000
38000
36000
34000
32000
30000
28000
26000
24000
22000
20000
18000
16000
14000
12000
10000
8000
6000
4000
2000
10.0
9.5
9.0
8.5
8.0
7.5
7.0
6.5
6.0
5.5
5.0
4.5
4.0
3.5
3.0
2.5
2.0
1.5
1.0
0.5
0.0
f1 (ppm)

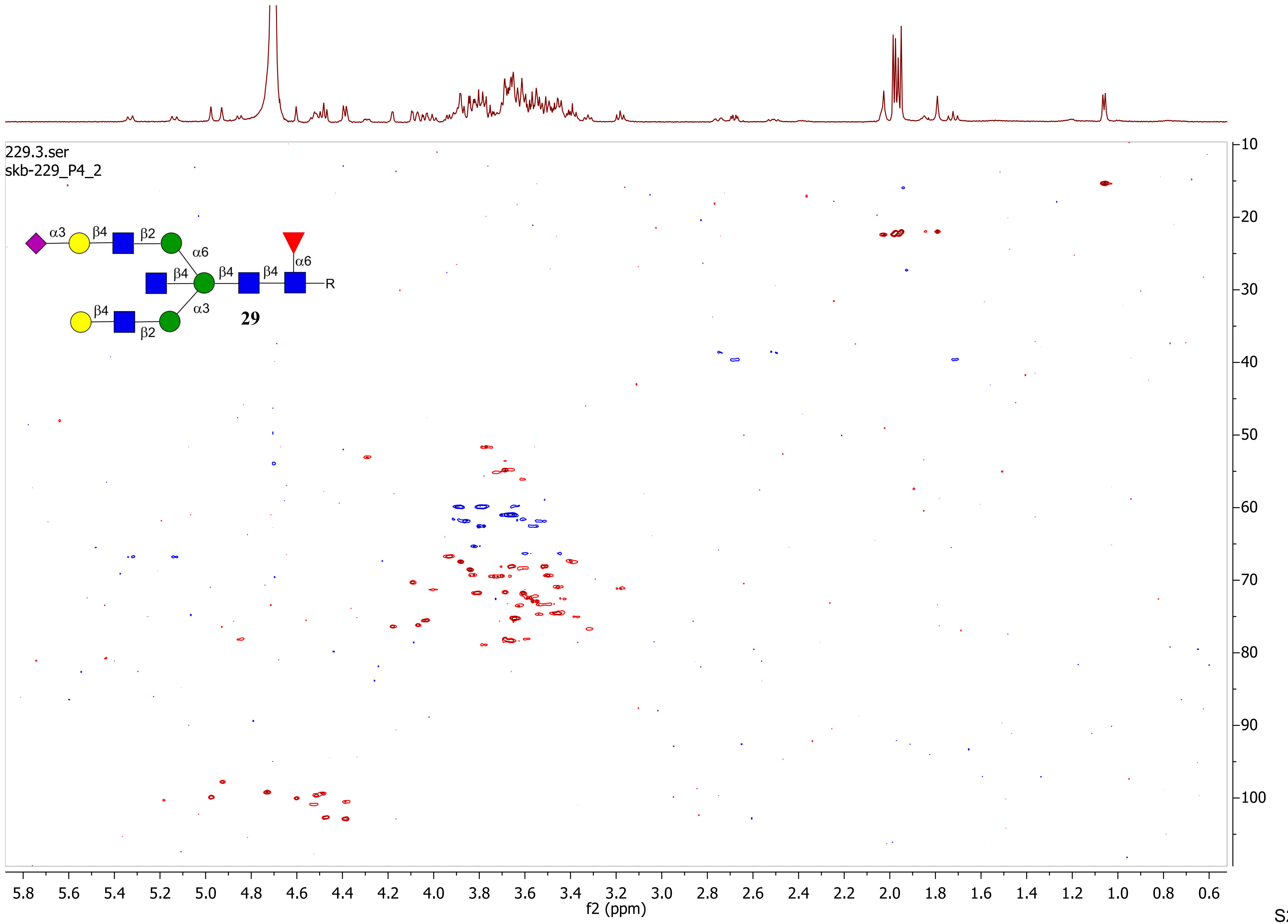

229.3.ser
skb-229_P4_2
α3
β4
β2
α6
β4
β4
β4
α6
R
α3
β4
β2
29
f2 (ppm)
f1 (ppm)

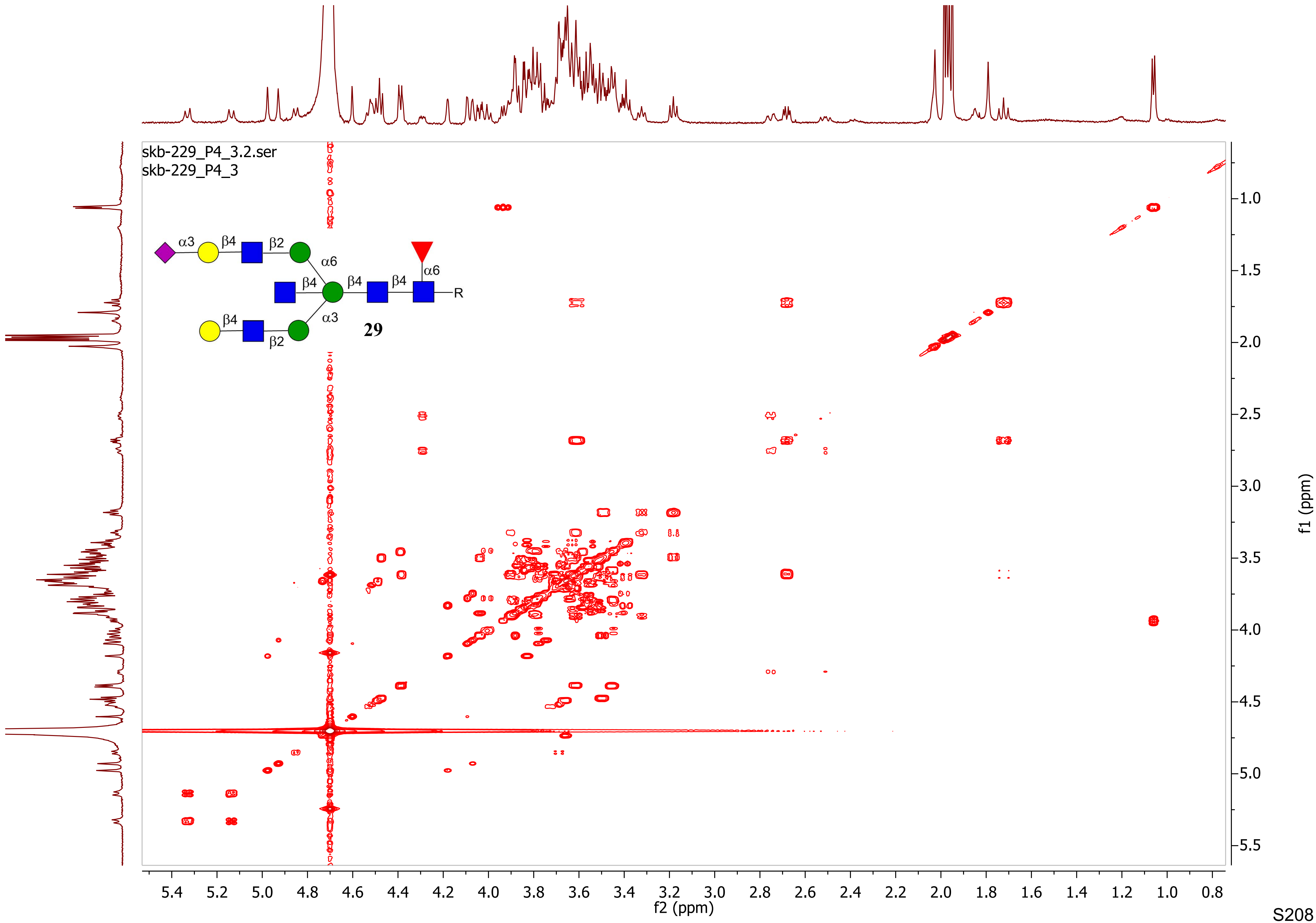
skb-229_P4_3.2.ser
skb-229_P4_3
α3
β4
β2
α6
β4
β4
β4
α6
R
β4
α3
β2
29
f2 (ppm)
f1 (ppm)

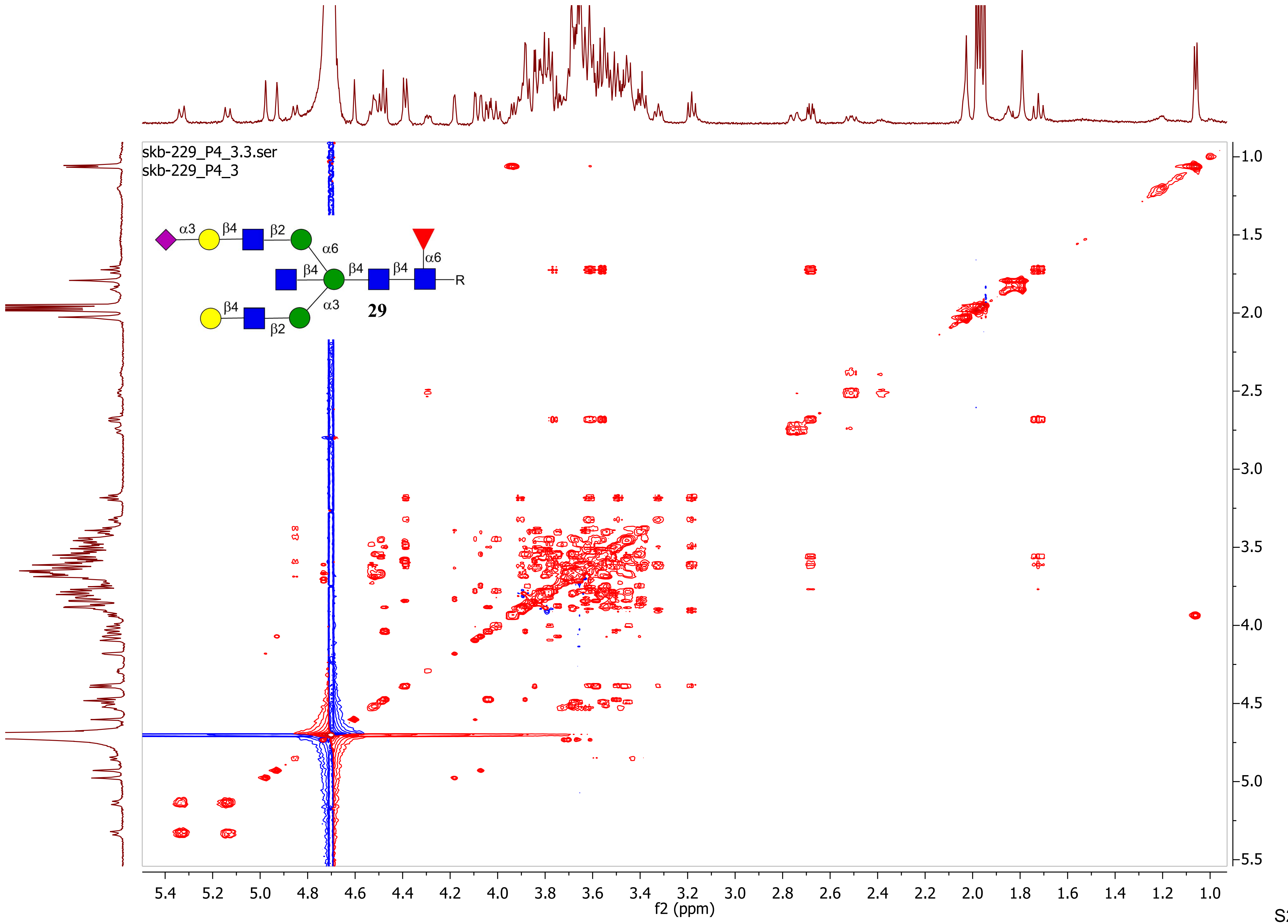
skb-229_P4_3.3.ser
skb-229_P4_3
α3
β4
β2
α6
β4
β4
β4
α6
R
β4
β2
α3
29
f2 (ppm)
f1 (ppm)

skb-176_p4.1.fid
skb-176_p4

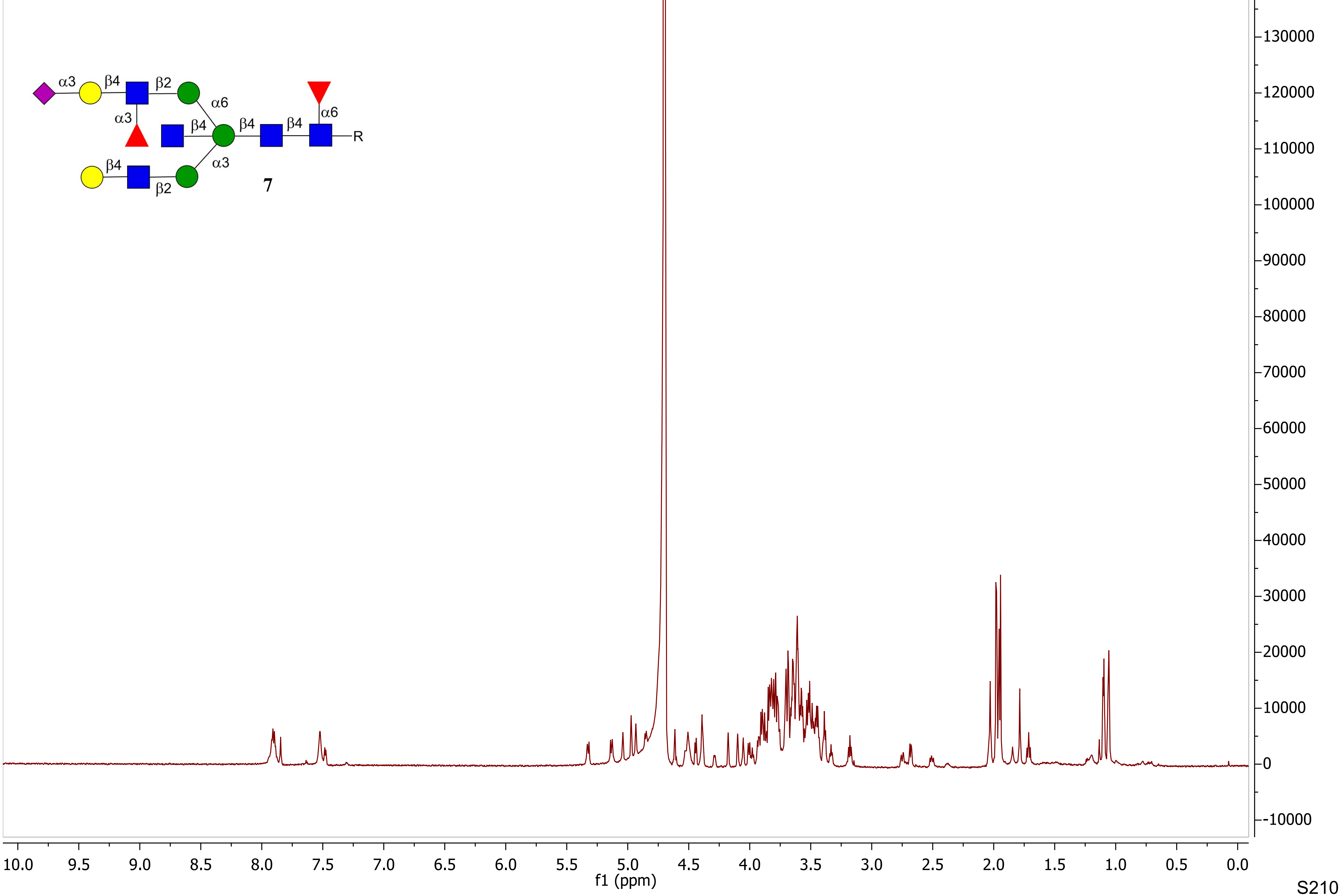

skb-176_p4.2.fid
skb-176_p4

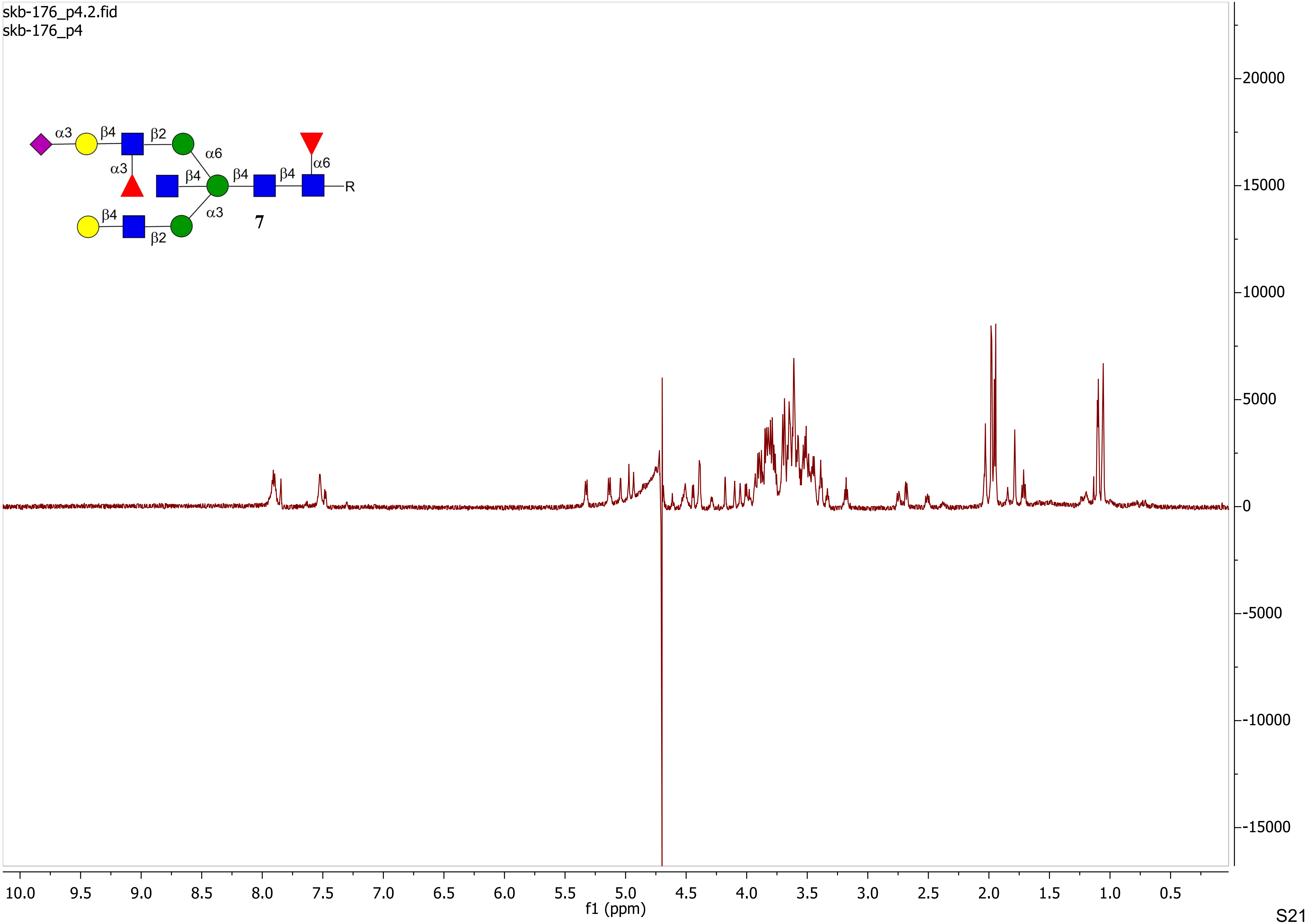

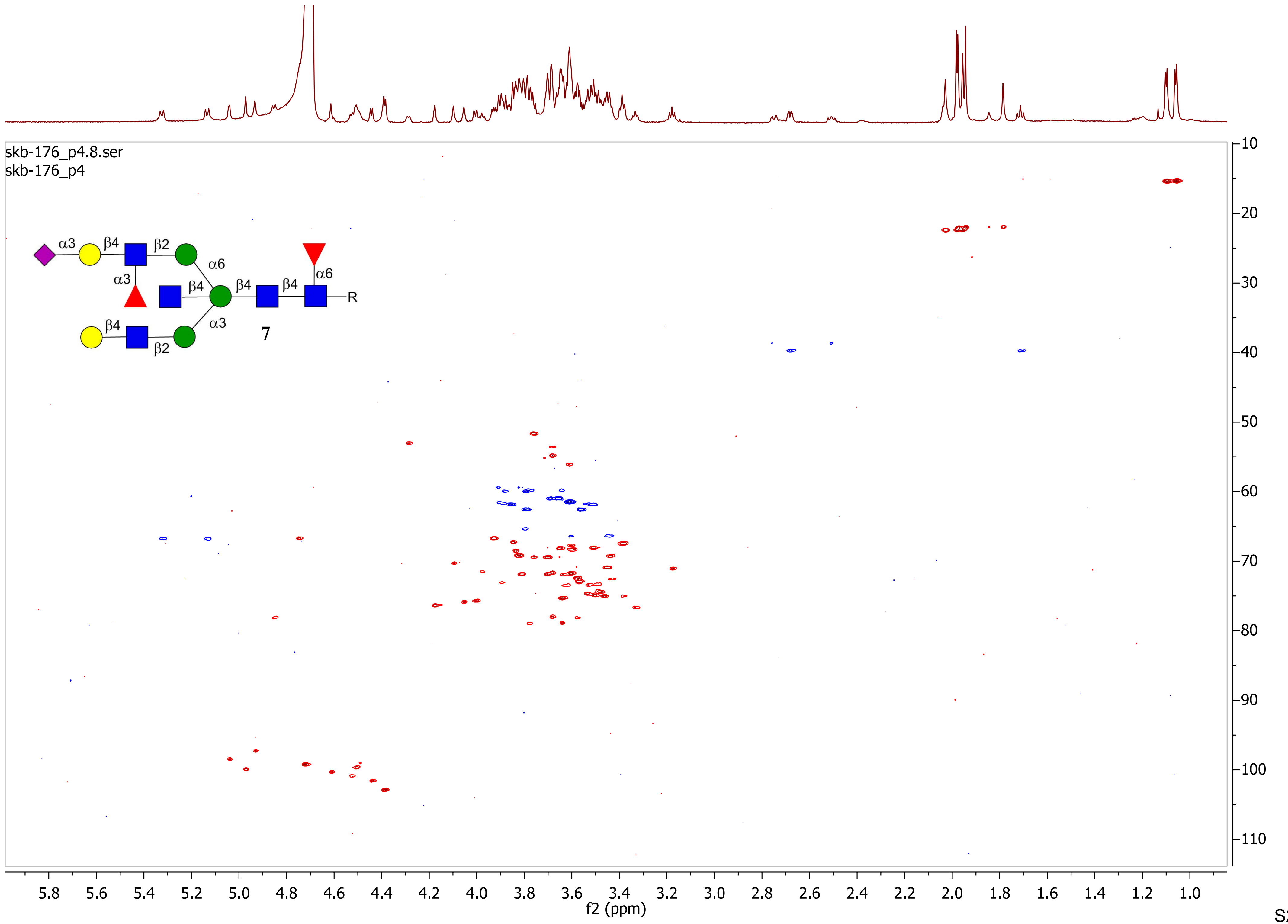

skb-176_p4.8.ser
skb-176_p4
α3
β4
β2
α6
α3
β4
β4
β4
α6
R
α3
β4
β2
7
f2 (ppm)
f1 (ppm)

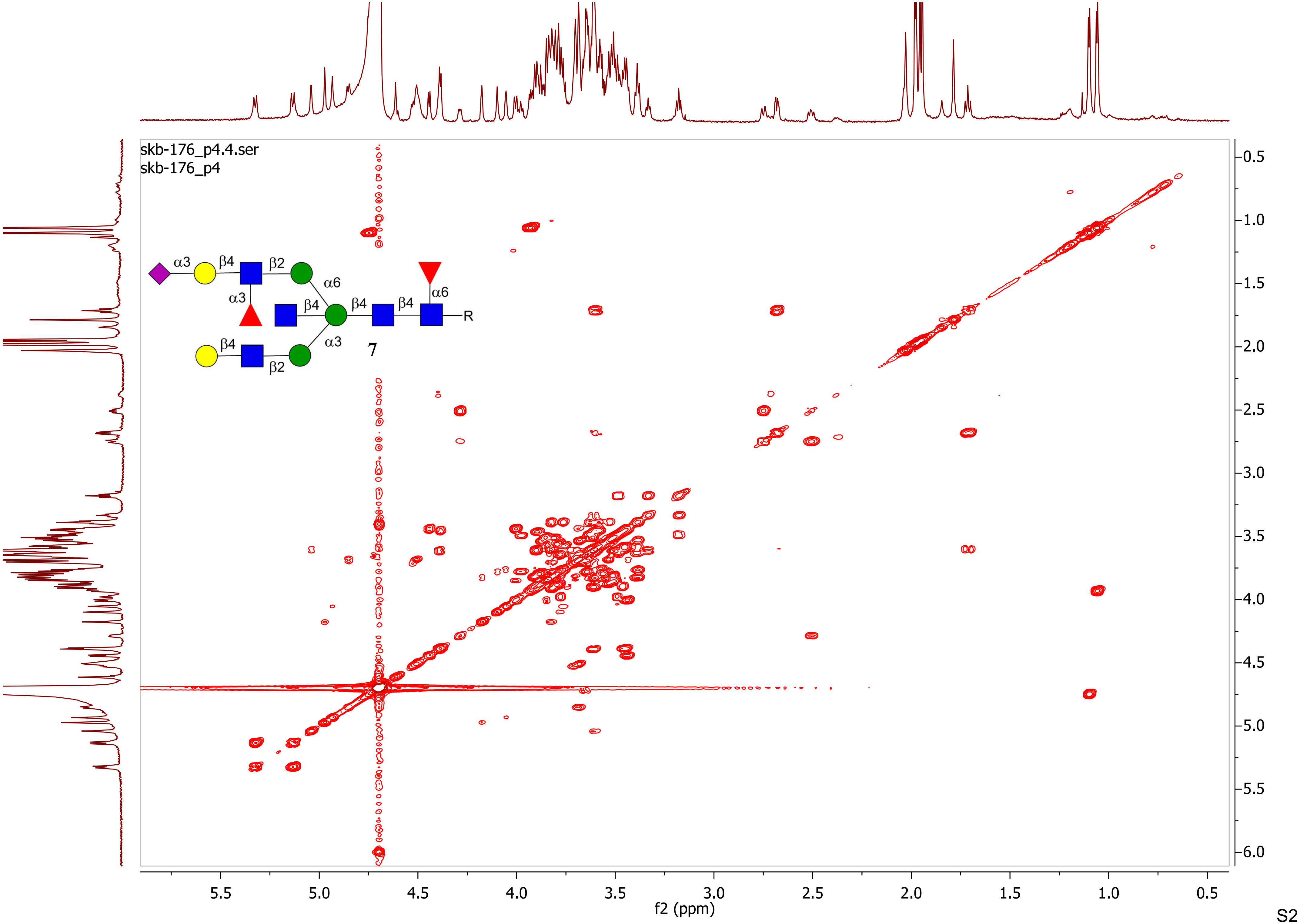
skb-176_p4.4.ser
skb-176_p4
α3
β4
β2
α6
α3
β4
β4
β4
α6
R
β4
α3
β2
7
f2 (ppm)
f1 (ppm)

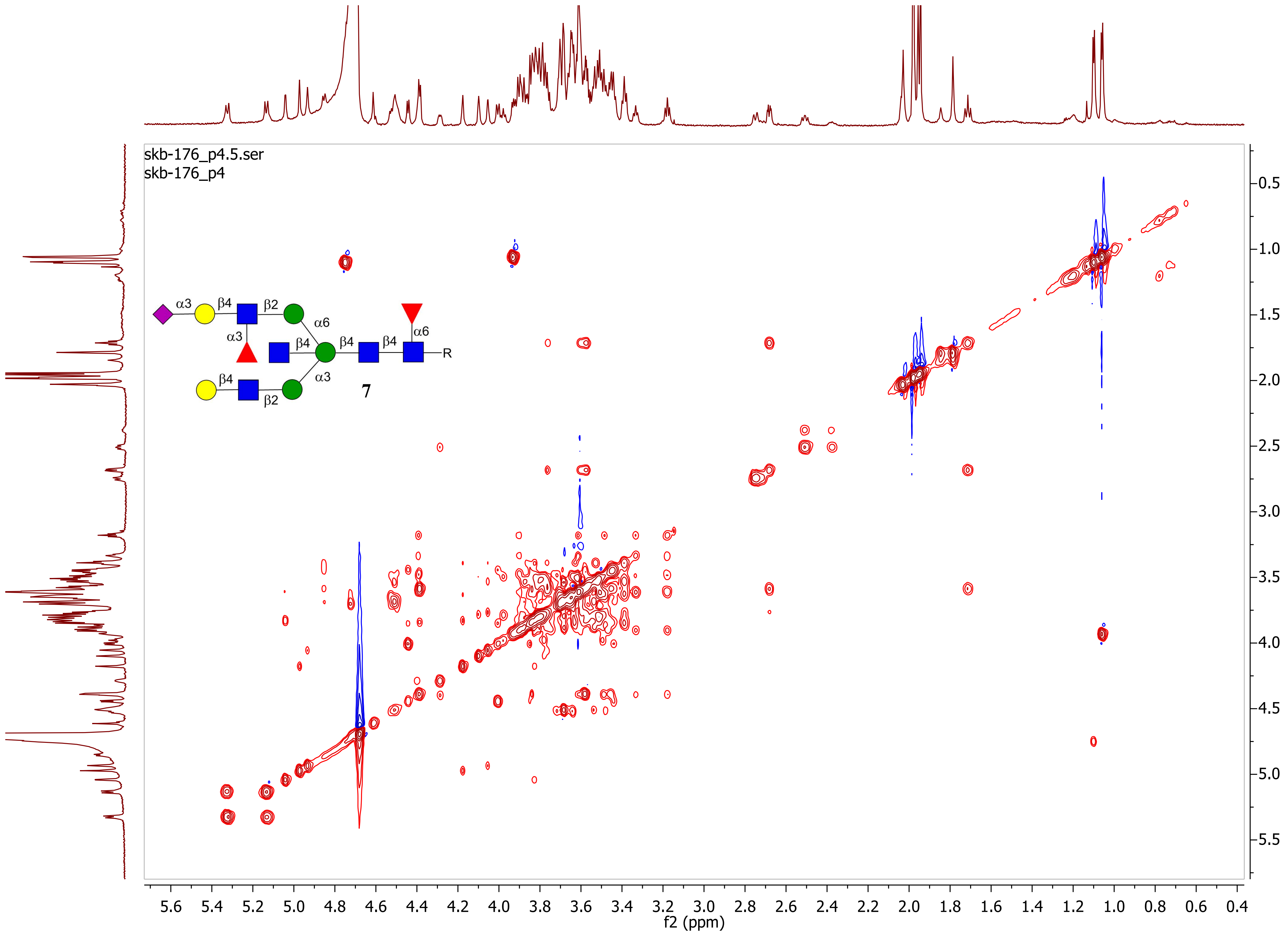
skb-176_p4.5.ser
skb-176_p4
α3
β4
β2
α6
α3
β4
β4
β4
α6
R
β4
α3
β2
7
f2 (ppm)
f1 (ppm)

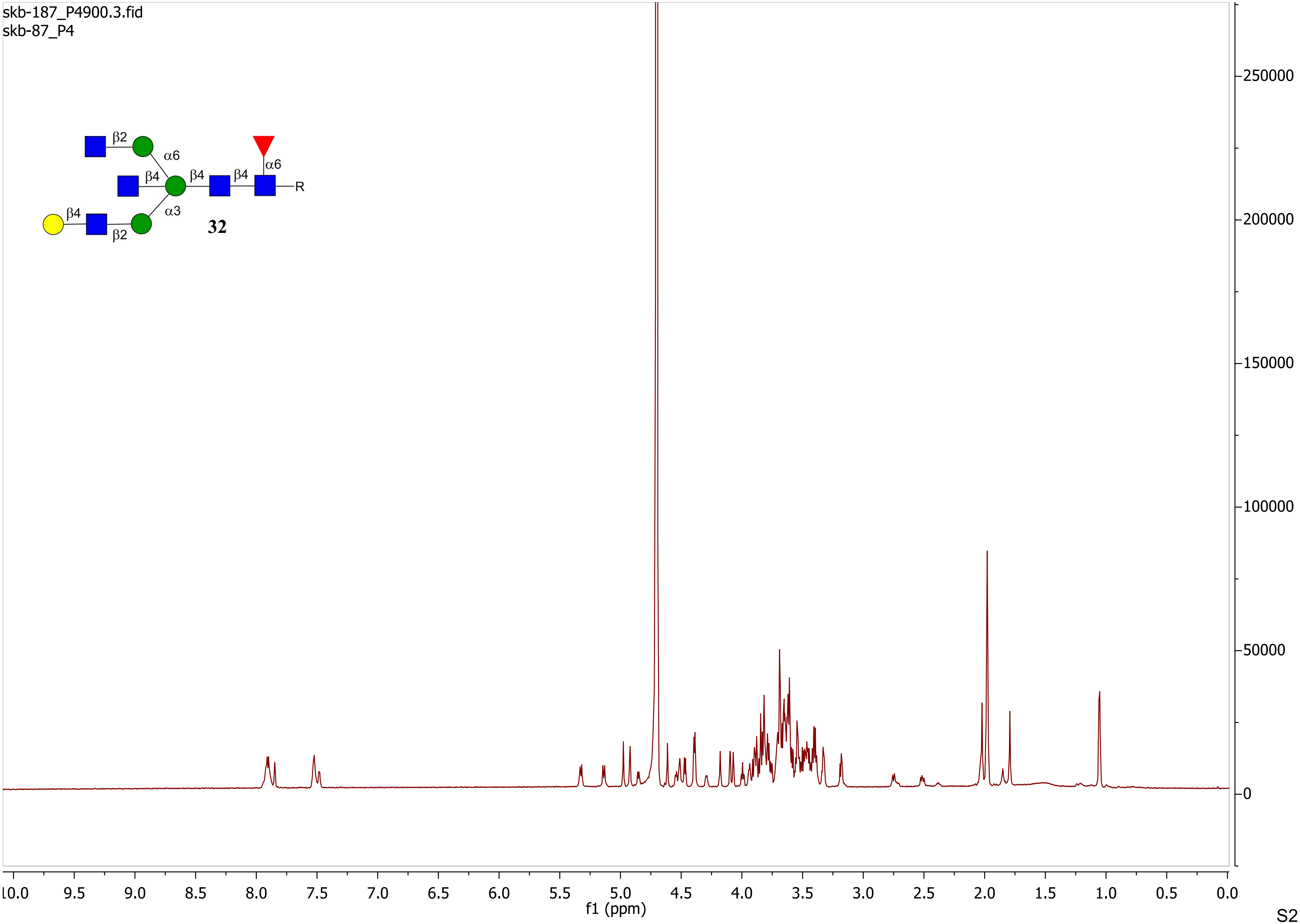

skb-187_P4900.3.fid
skb-87_P4
β2
α6
β4
β4
β4
α6
R
β4
α3
β2
32
250000
200000
150000
100000
50000
0
10.0
9.5
9.0
8.5
8.0
7.5
7.0
6.5
6.0
5.5
5.0
4.5
4.0
3.5
3.0
2.5
2.0
1.5
1.0
0.5
0.0
f1 (ppm)

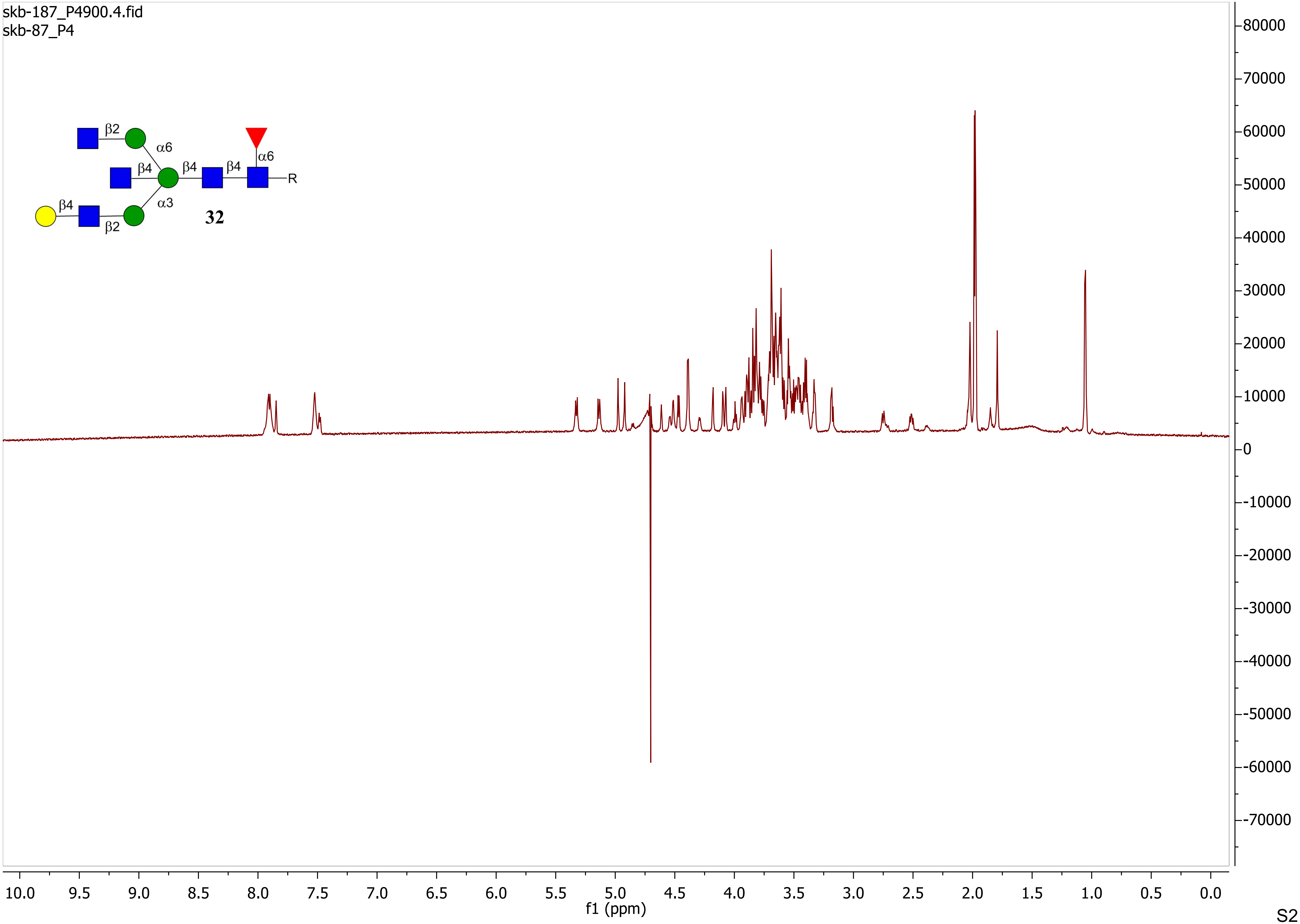
skb-187_P4900.4.fid
skb-87_P4
β2
α6
β4
β4
β4
α6
R
β4
α3
β2
32
f1 (ppm)

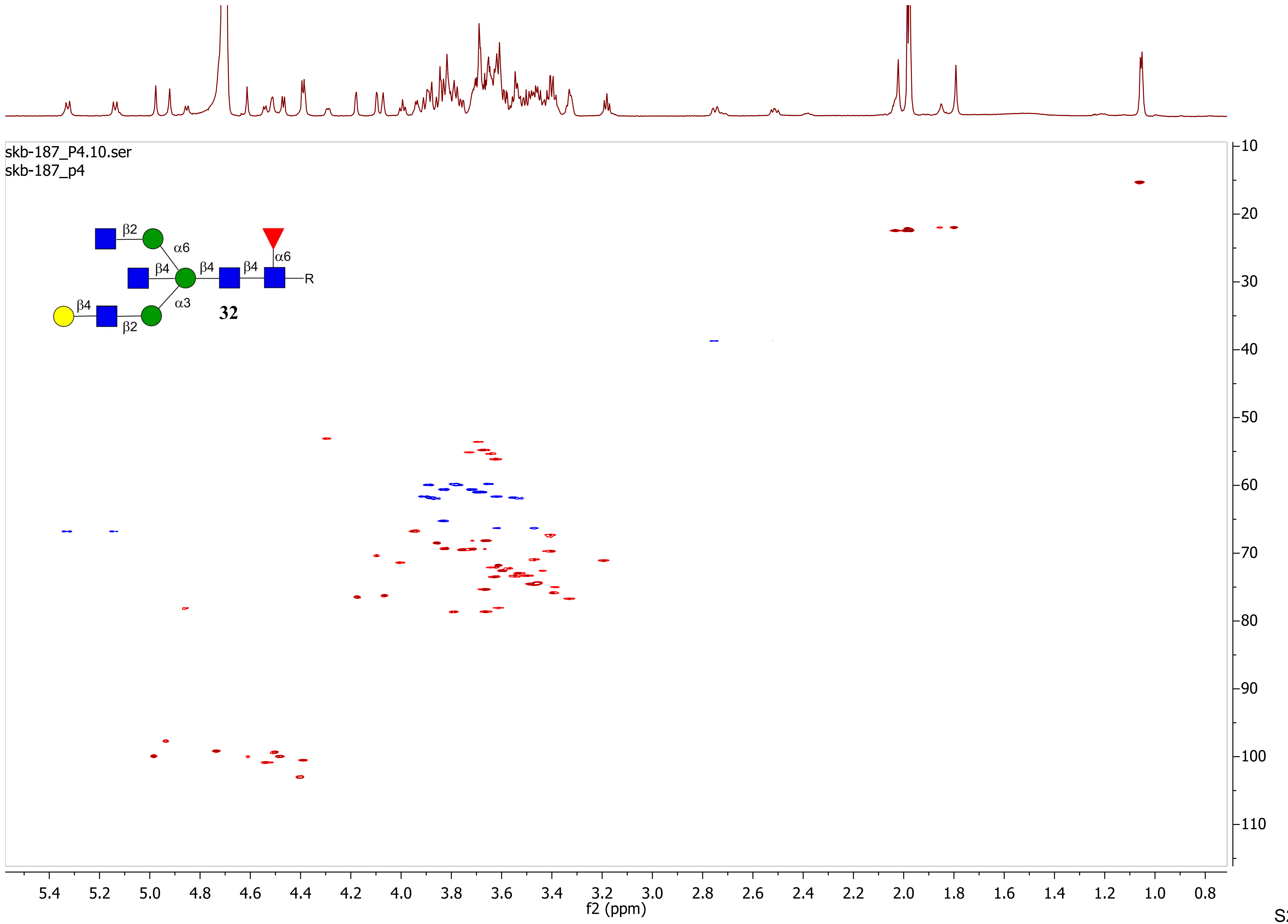
skb-187_P4.10.ser
skb-187_p4
β2
α6
β4
β4
β4
α6
R
β4
α3
β2
32
f2 (ppm)
f1 (ppm)

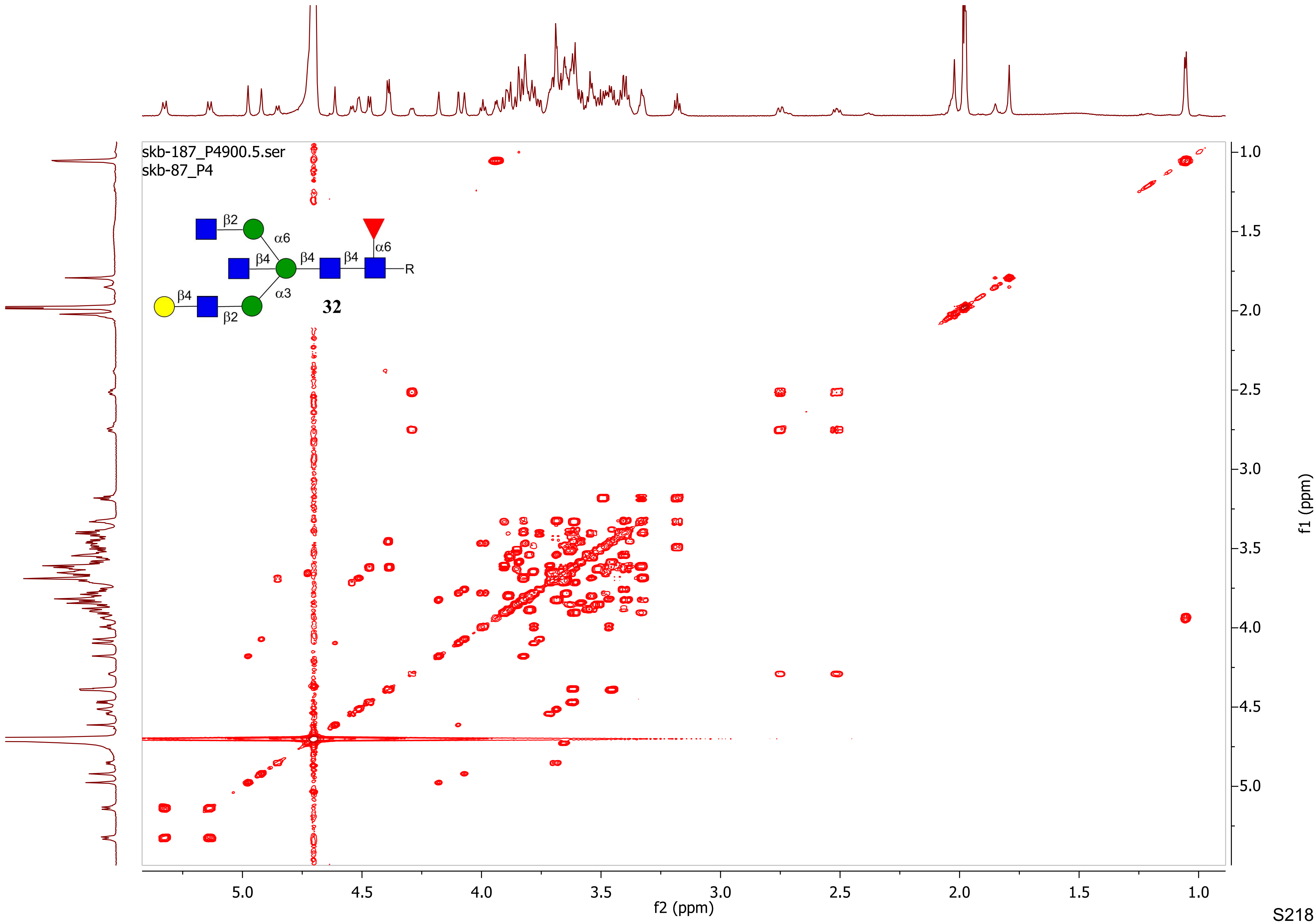

skb-187_P4900.5.ser
skb-87_P4
β2
α6
β4
β4
β4
α6
R
β4
α3
β2
32
5.0
4.5
4.0
3.5
3.0
2.5
2.0
1.5
1.0
f2 (ppm)
f1 (ppm)

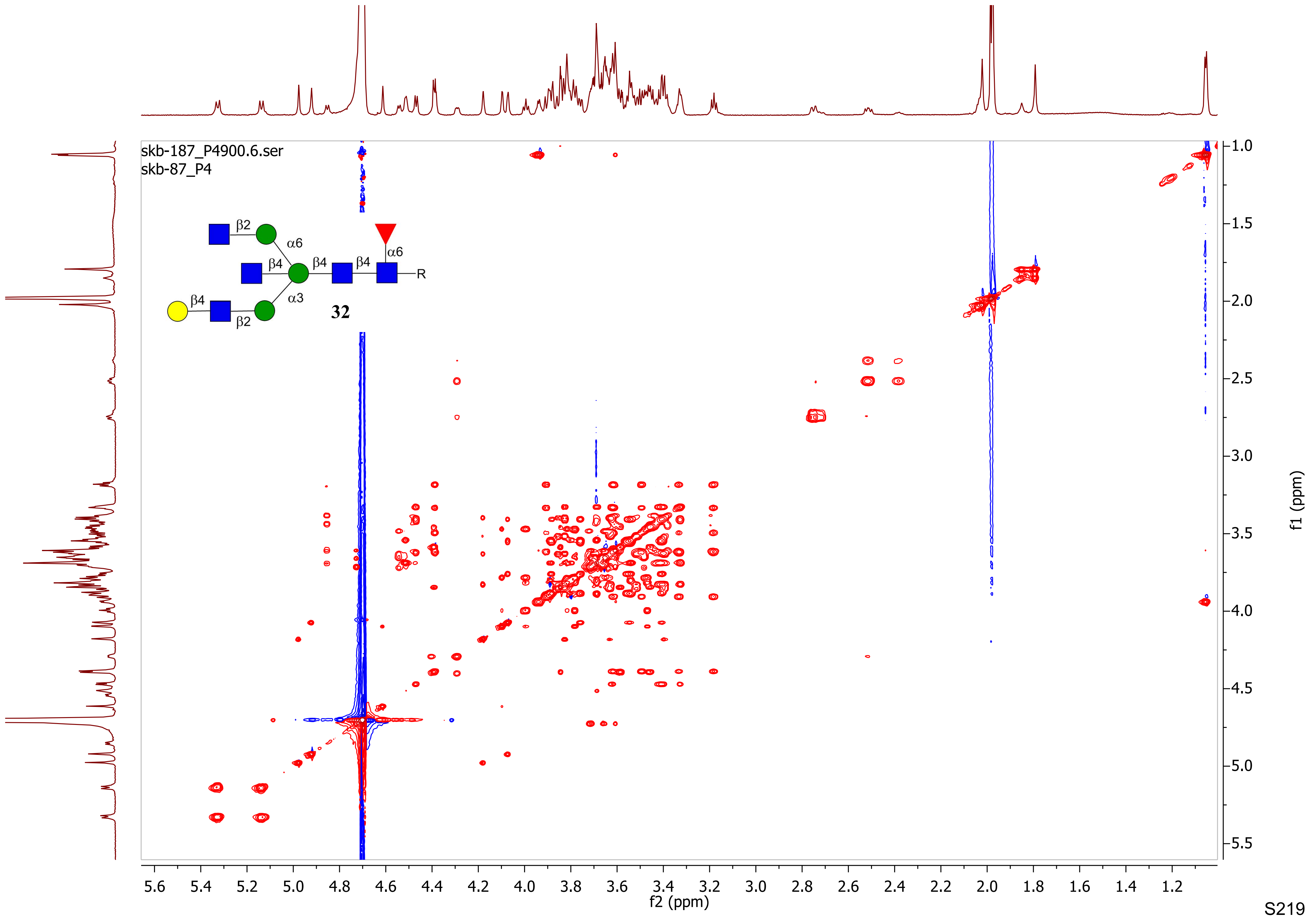
skb-187_P4900.6.ser
skb-87_P4
β2
α6
β4
β4
β4
α6
R
β4
α3
β2
32
f2 (ppm)
f1 (ppm)

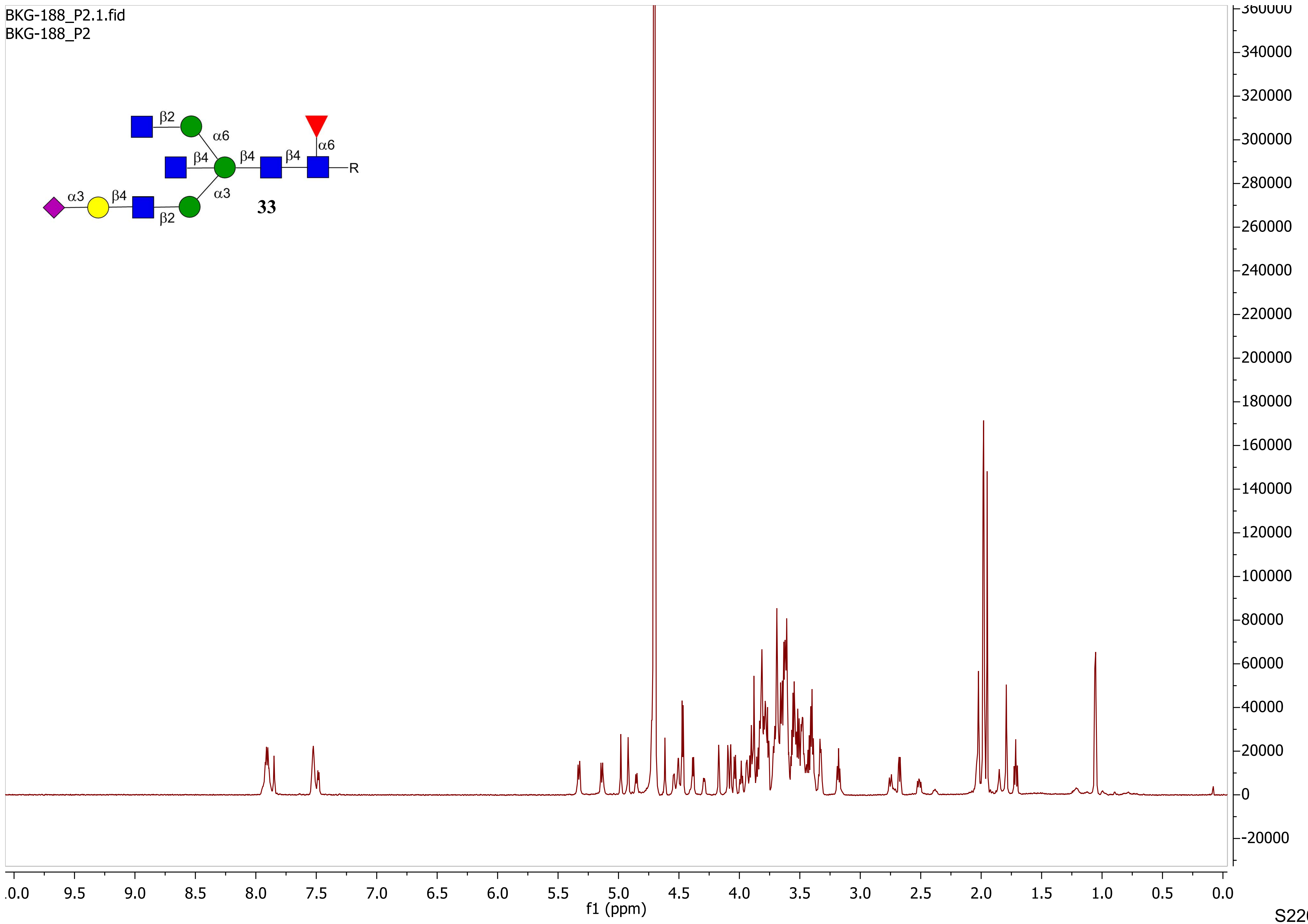
BKG-188_P2.1.fid
BKG-188_P2
β2
α6
β4
β4
β4
α6
R
α3
β4
α3
β2
33
f1 (ppm)

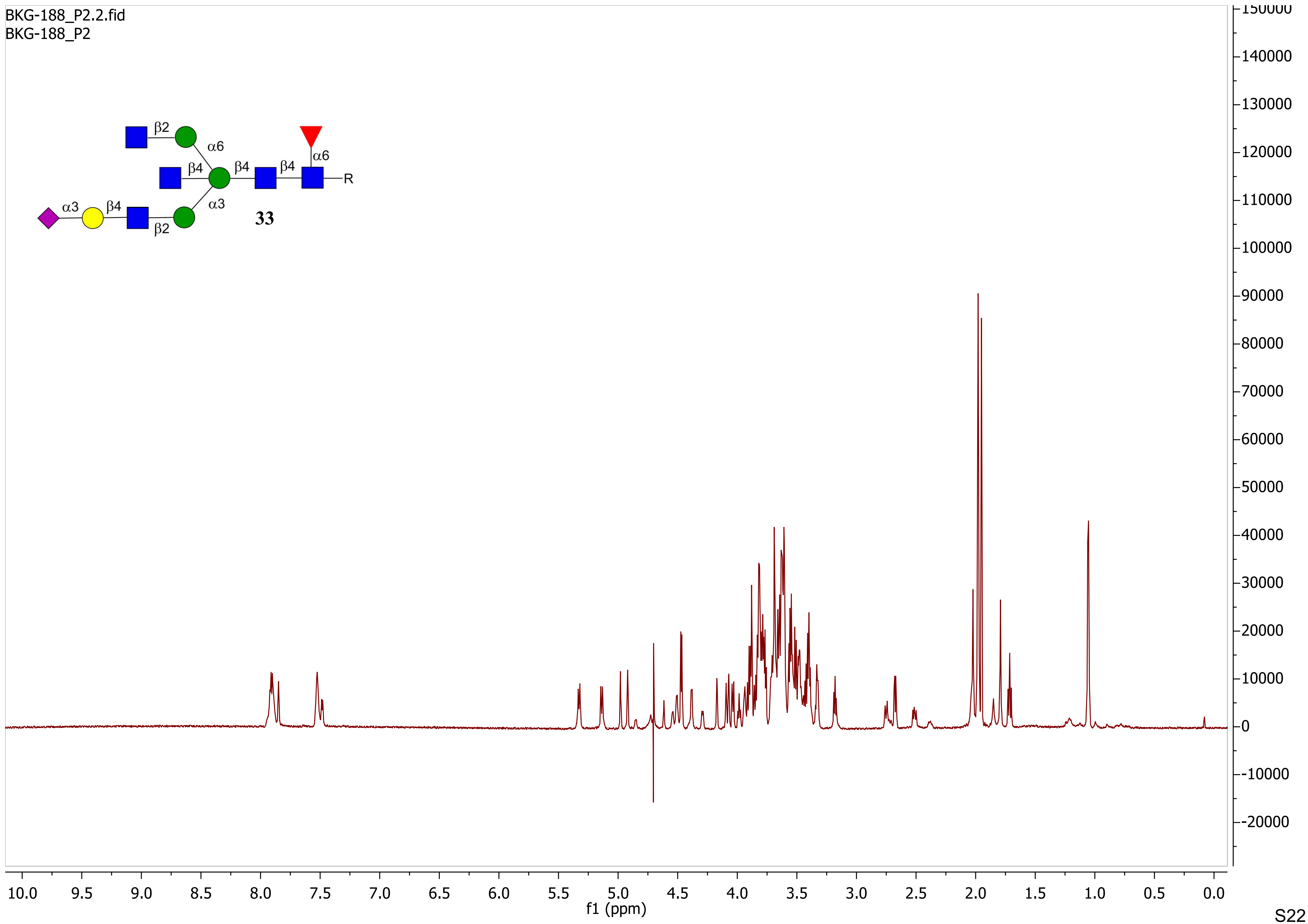
BKG-188_P2.2.fid
BKG-188_P2
β2
α6
β4
β4
β4
α6
R
α3
β4
α3
β2
33
f1 (ppm)

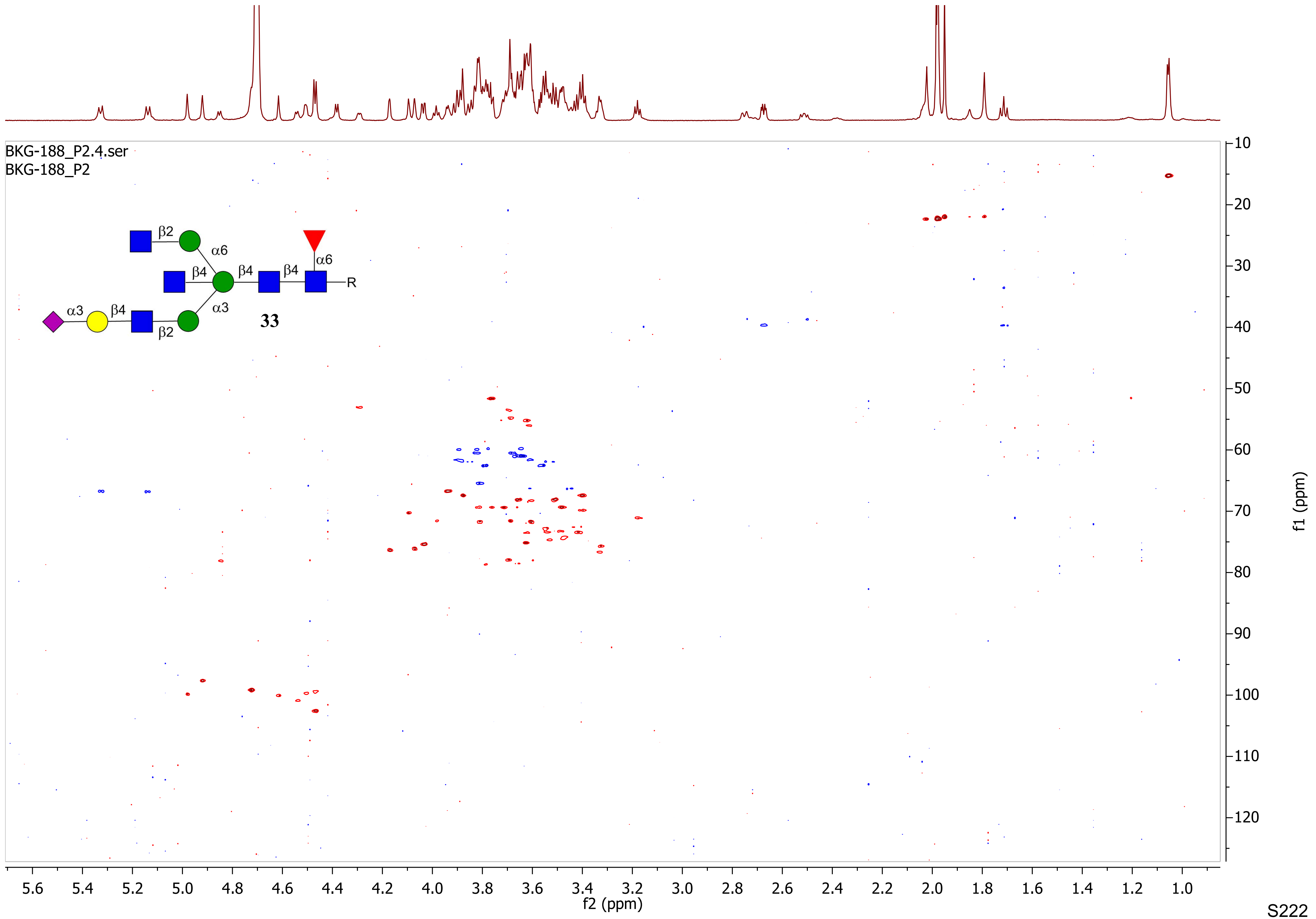
BKG-188_P2.4.ser
BKG-188_P2
β2
α6
β4
β4
β4
α6
R
α3
β4
α3
β2
33
f2 (ppm)
f1 (ppm)

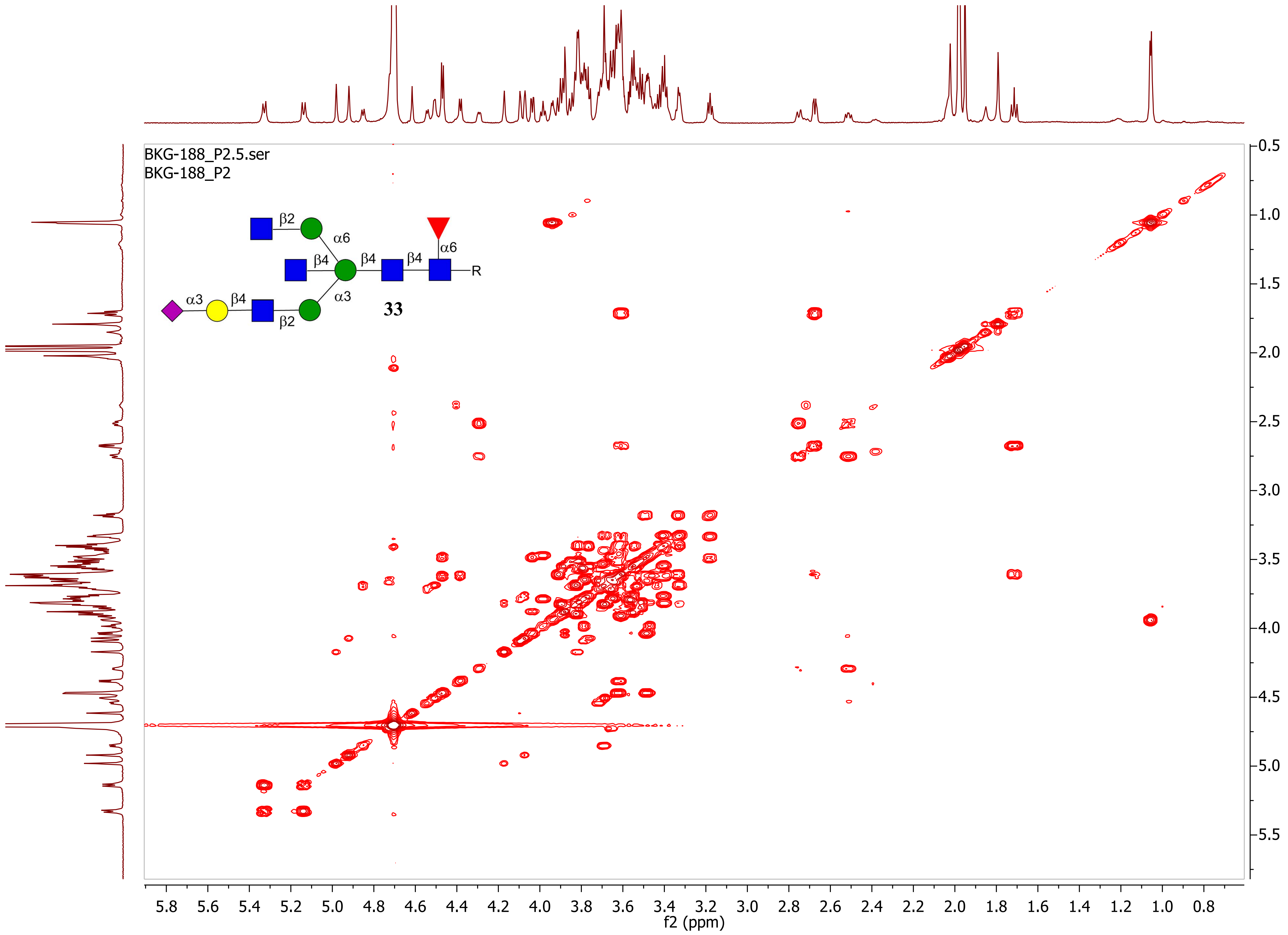
BKG-188_P2.5.ser
BKG-188_P2
β2
α6
β4
β4
β4
α6
R
α3
β4
α3
β2
33
f2 (ppm)
f1 (ppm)

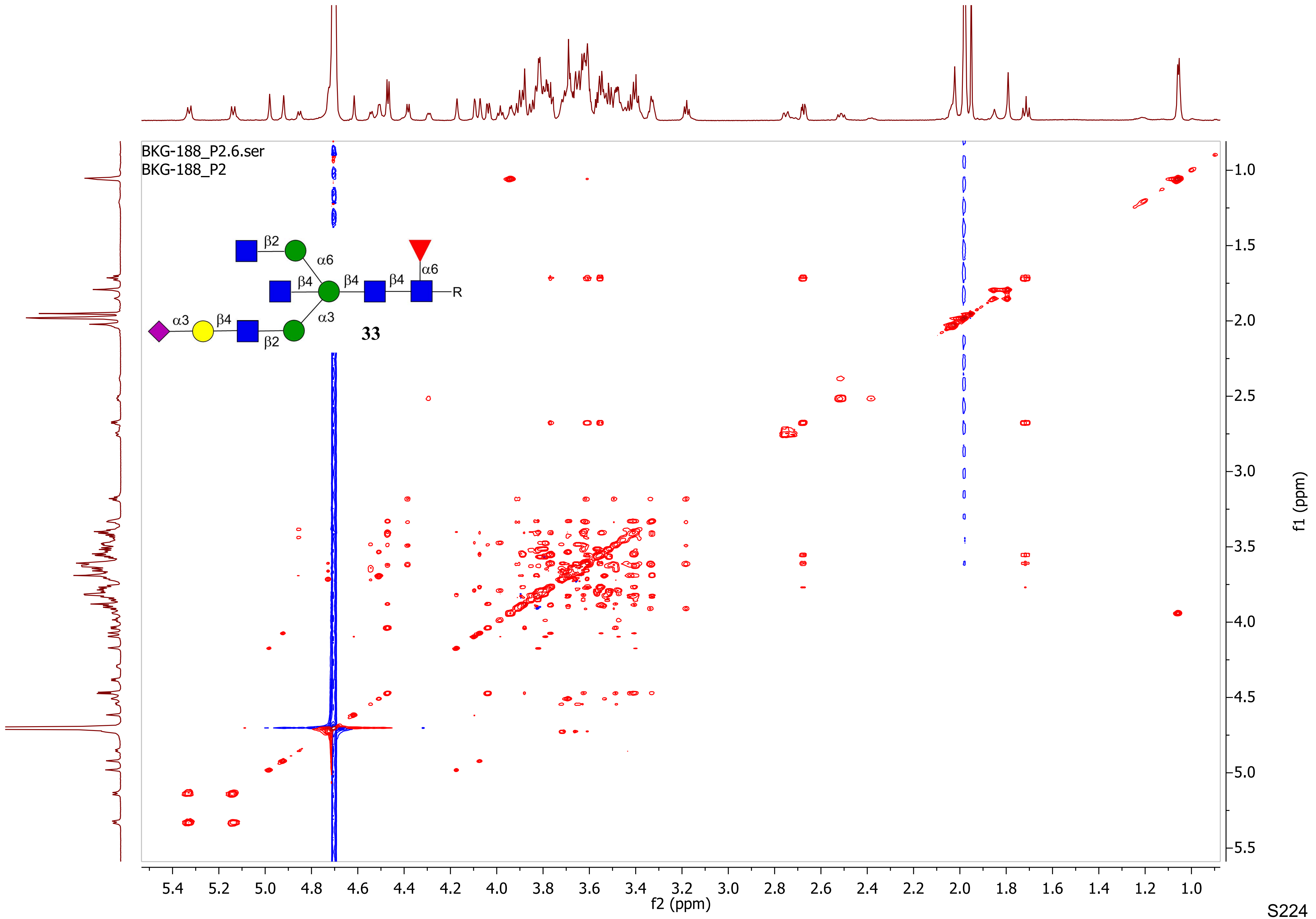

BKG-188_P2.6.ser
BKG-188_P2
β2
α6
β4
β4
β4
α6
R
α3
β4
α3
β2
33
f2 (ppm)
f1 (ppm)

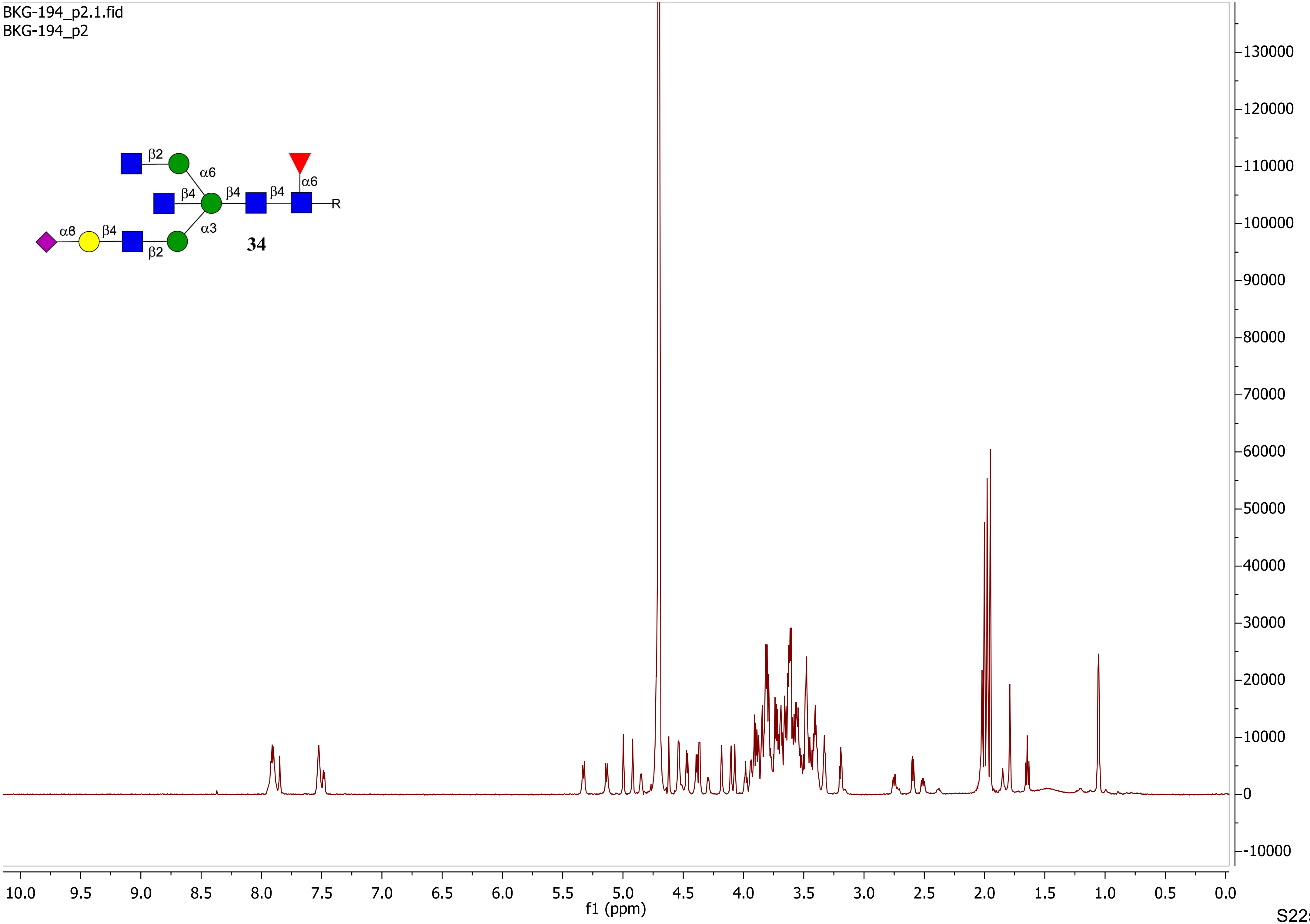
BKG-194_p2.1.fid
BKG-194_p2
β2
α6
β4
β4
β4
α6
R
α3
α8
β4
β2
34
f1 (ppm)
10.0 9.5 9.0 8.5 8.0 7.5 7.0 6.5 6.0 5.5 5.0 4.5 4.0 3.5 3.0 2.5 2.0 1.5 1.0 0.5 0.0
130000 120000 110000 100000 90000 80000 70000 60000 50000 40000 30000 20000 10000 0 -10000

BKG-194_p2.2.fid
BKG-194_p2

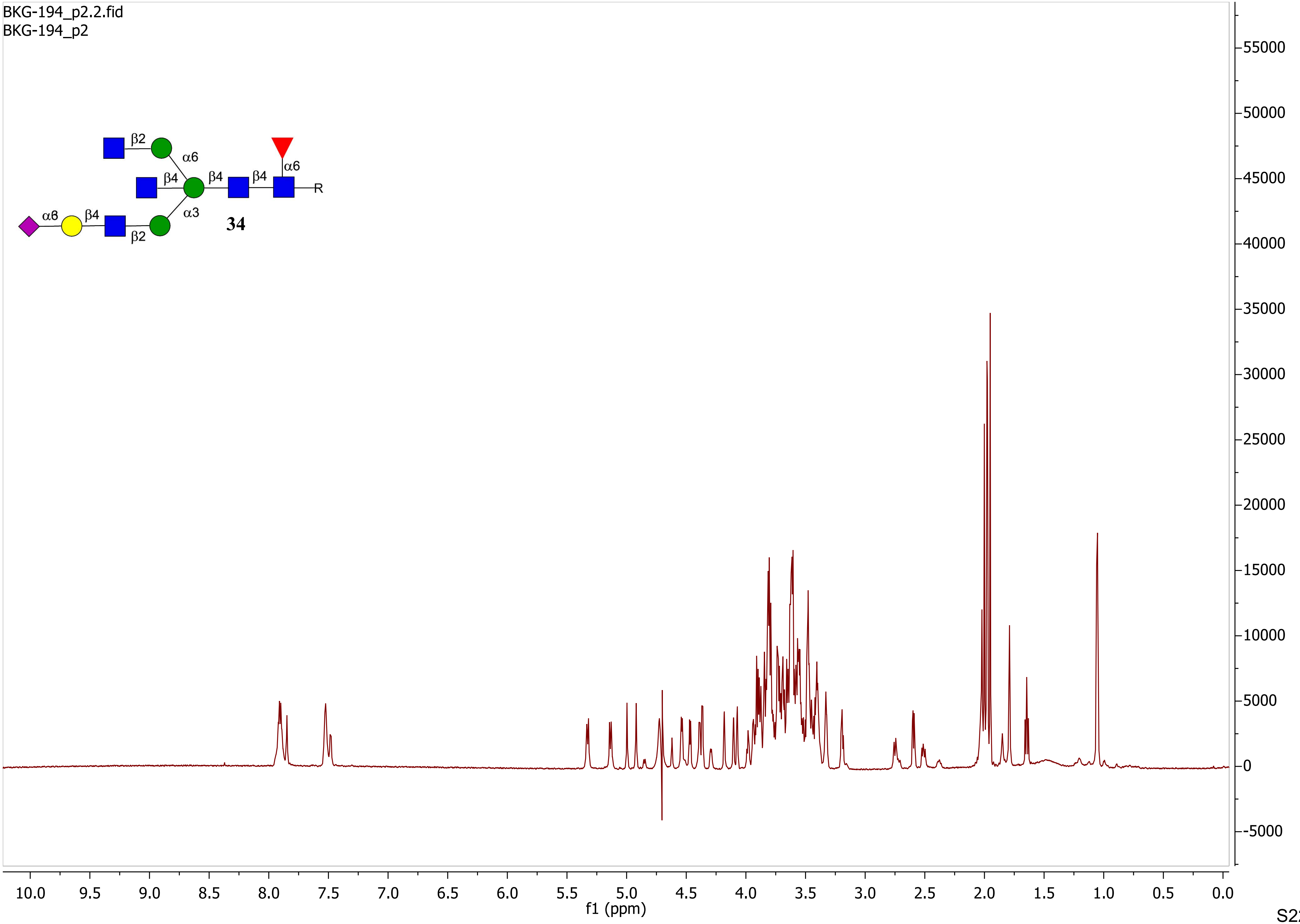

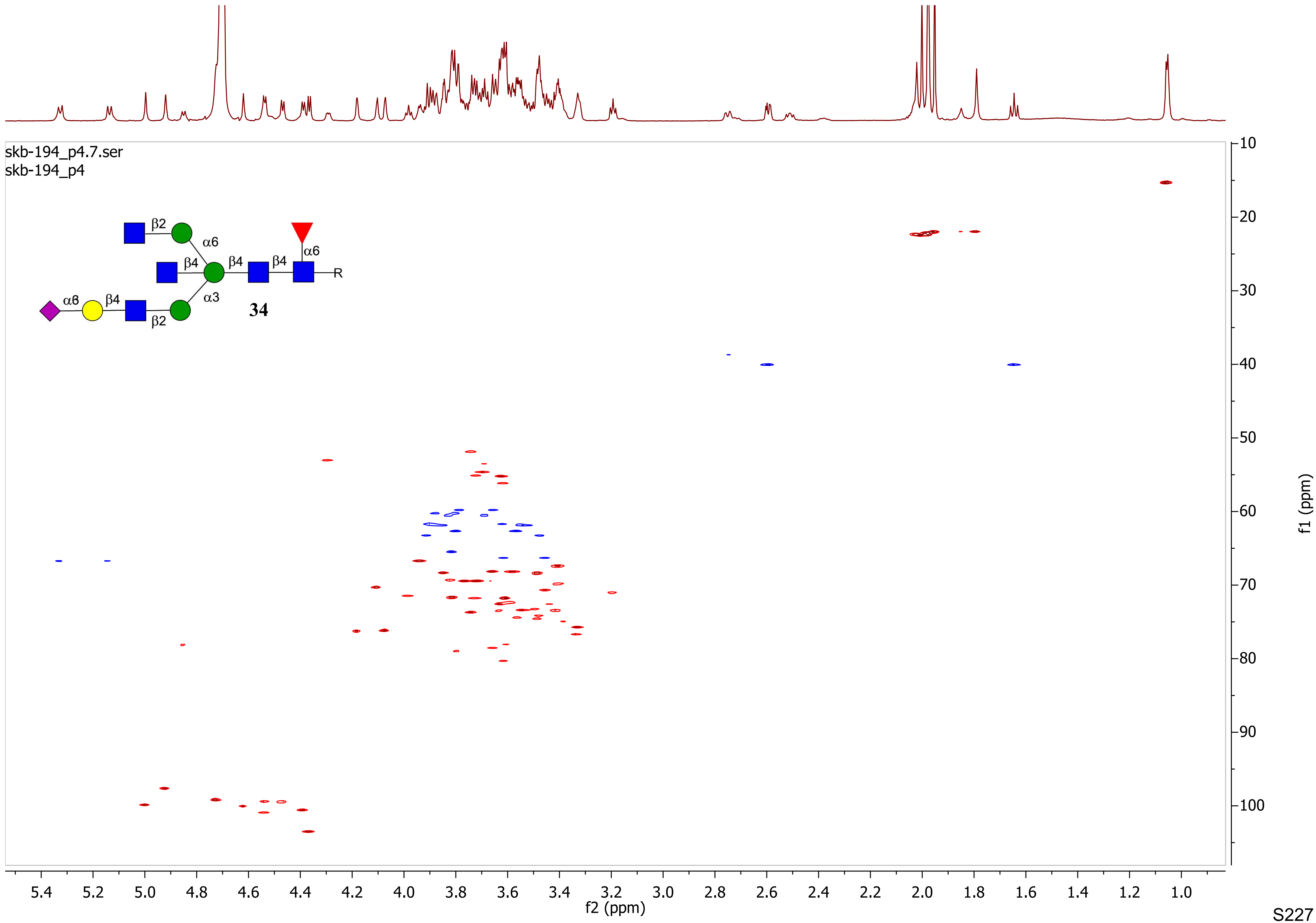

skb-194_p4.7.ser
skb-194_p4
34
f2 (ppm)
f1 (ppm)

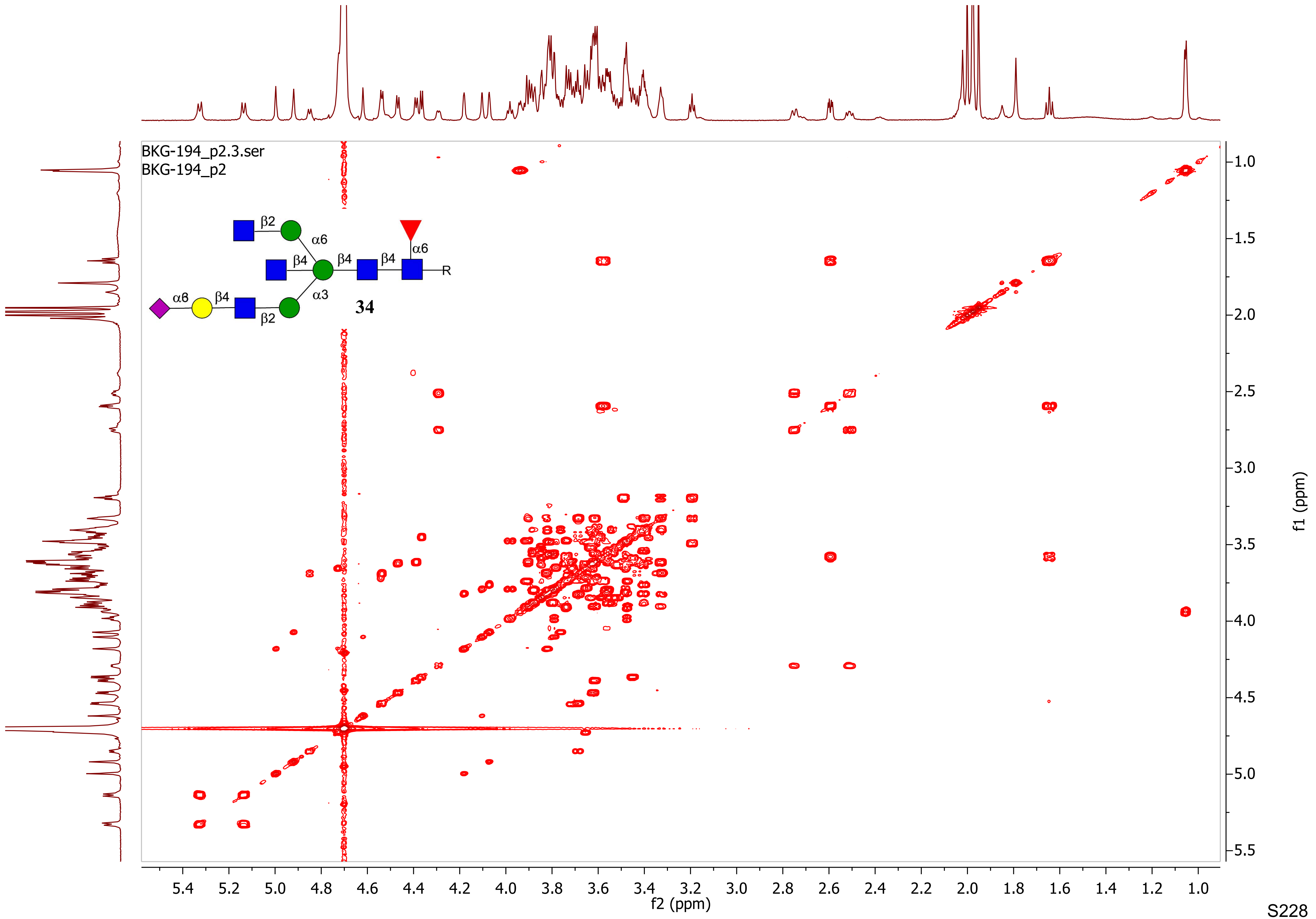

BKG-194_p2.3.ser
BKG-194_p2
β2
α6
β4
β4
β4
α6
R
α6
β4
α3
β2
34
f2 (ppm)
f1 (ppm)

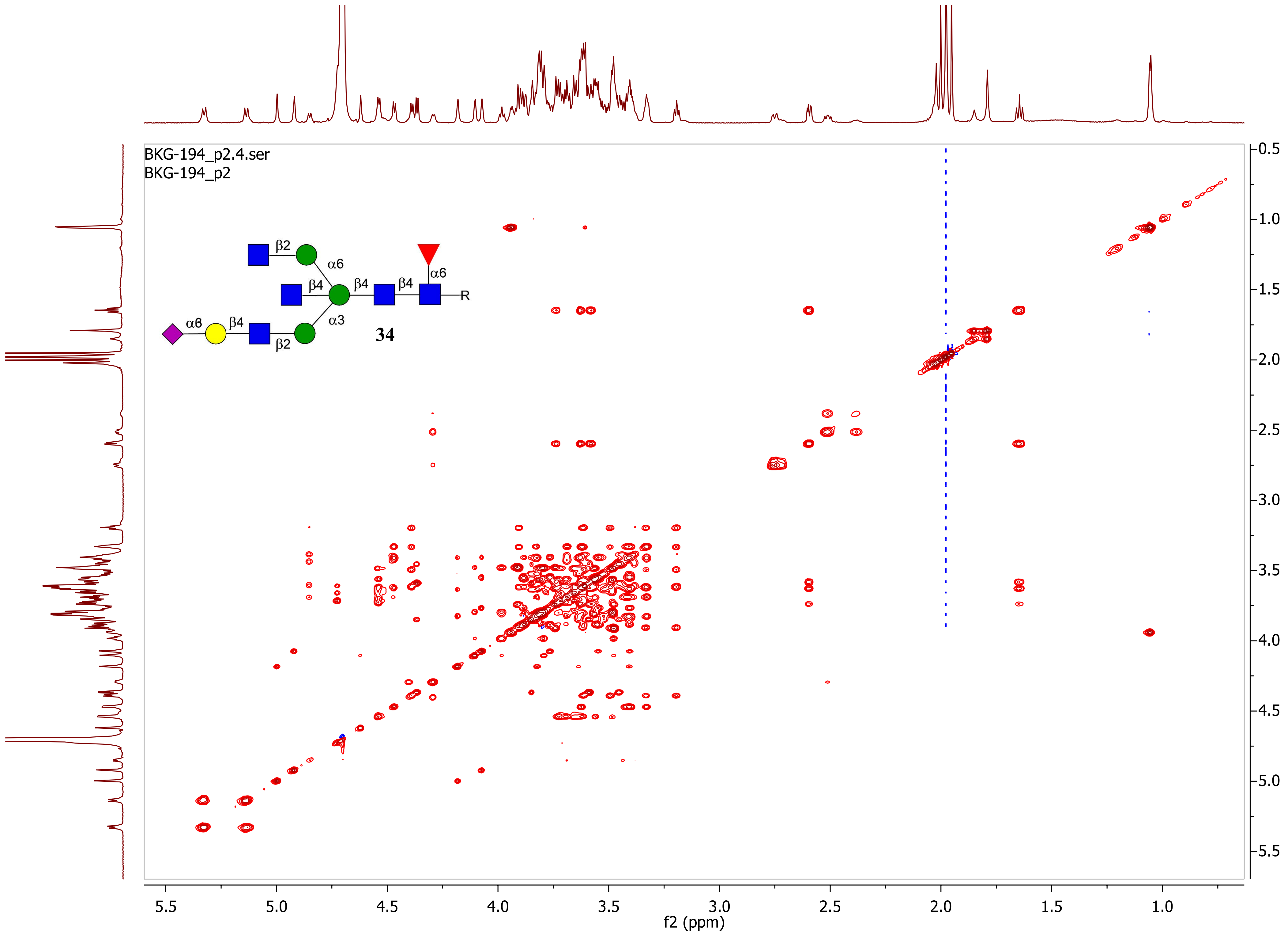

BKG-194_p2.4.ser
BKG-194_p2
β2
α6
β4
β4
β4
α6
R
α6
β4
β2
α3
34
f2 (ppm)
f1 (ppm)

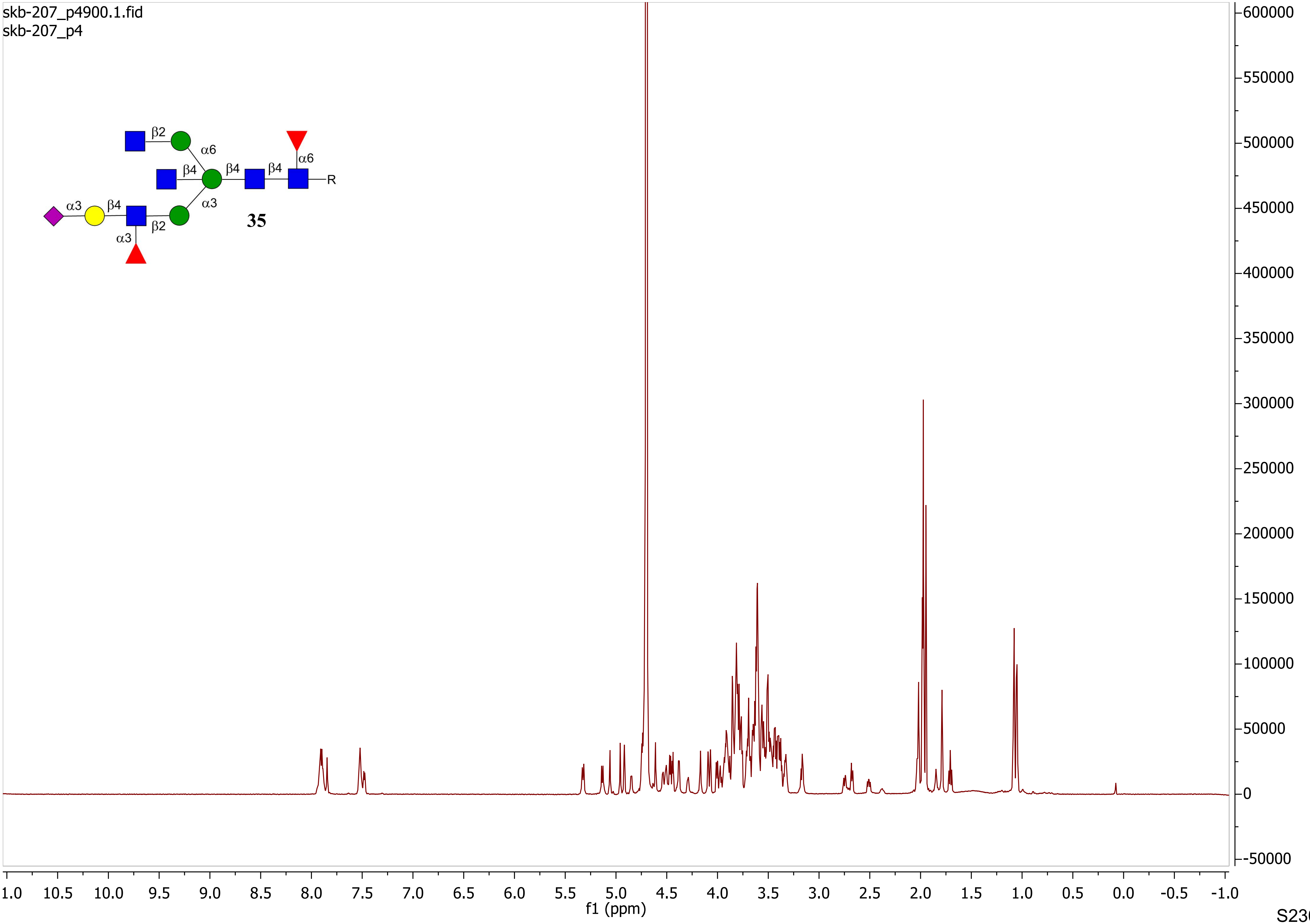
skb-207_p4900.1.fid
skb-207_p4
β2
α6
β4
β4
β4
α6
R
α3
β4
α3
β2
α3
35
f1 (ppm)

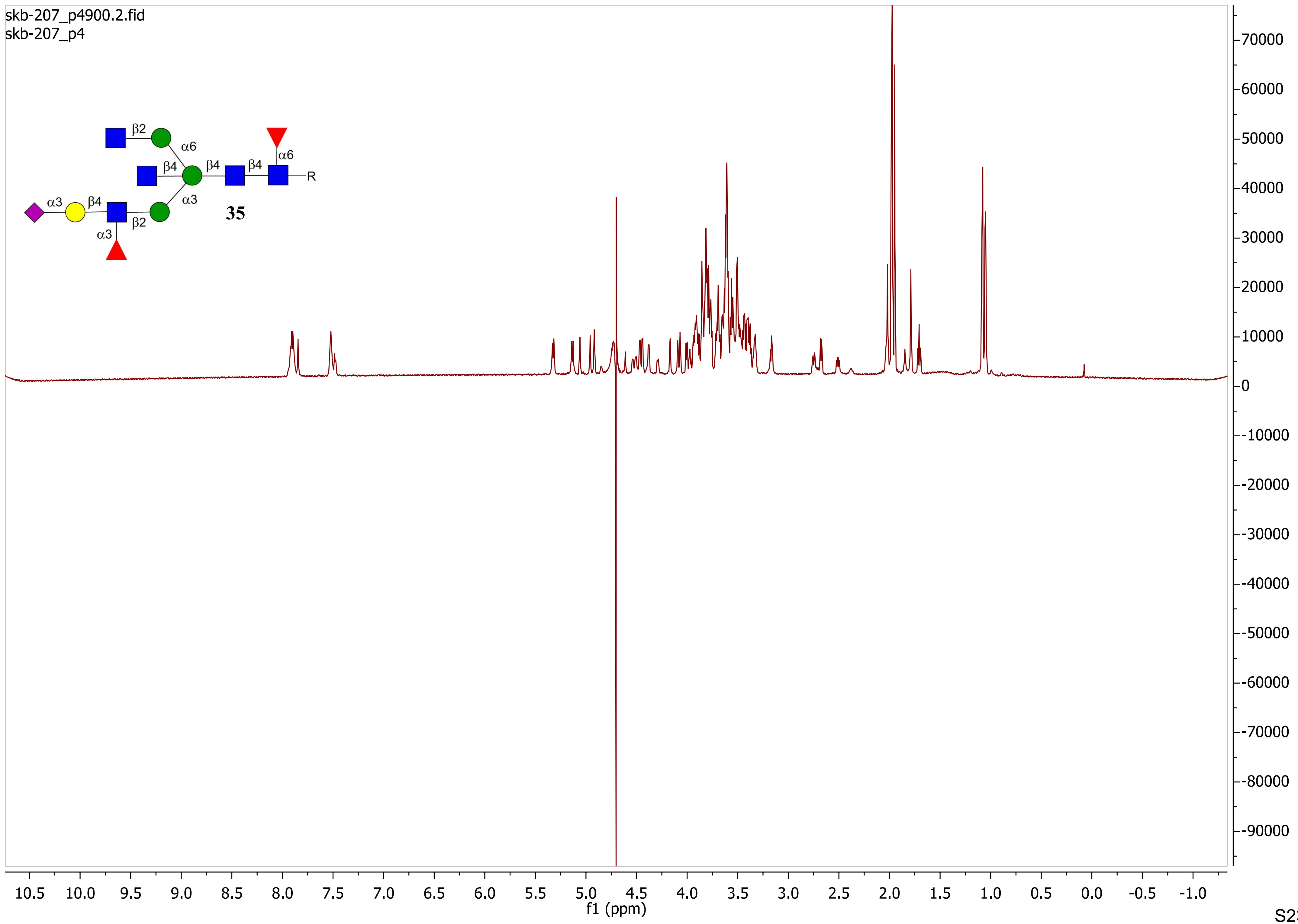
skb-207_p4900.2.fid
skb-207_p4
β2
α6
β4
β4
β4
α6
R
α3
β4
α3
β2
α3
35
f1 (ppm)

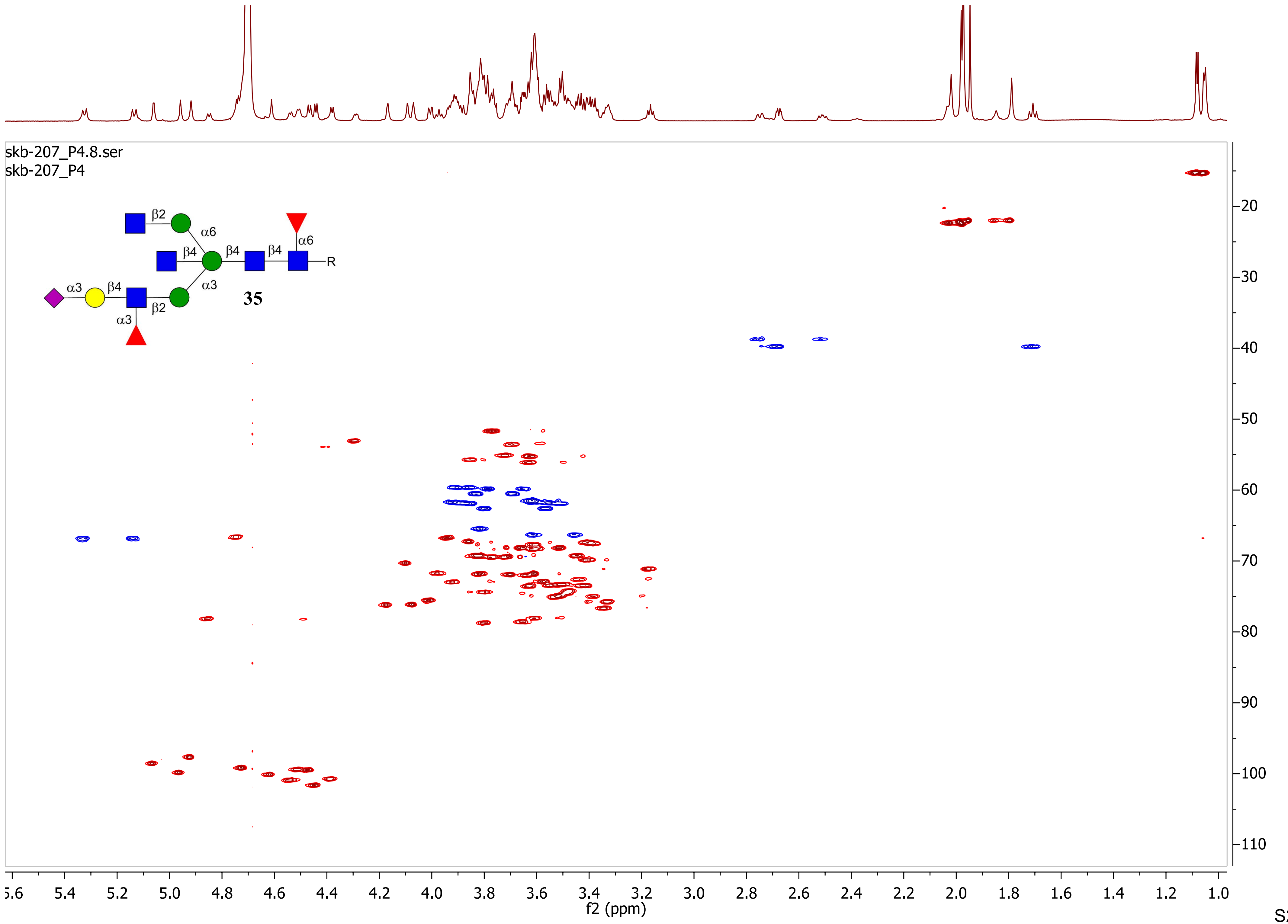

skb-207_P4.8.ser
skb-207_P4
β2
α6
β4
β4
β4
α6
R
α3
β4
α3
β2
α3
35
f1 (ppm)
f2 (ppm)

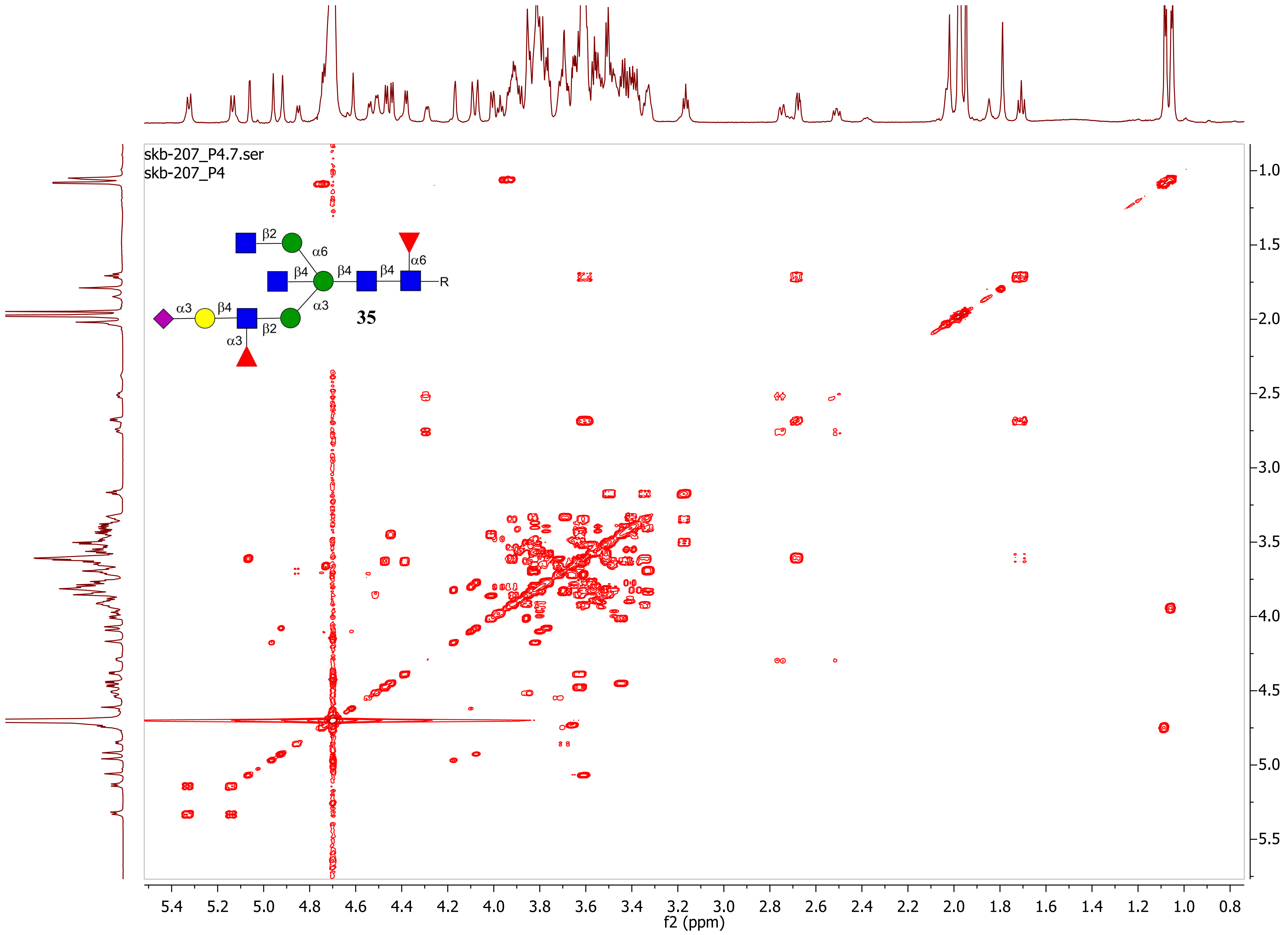

skb-207_P4.7.ser
skb-207_P4
β2
α6
β4
β4
β4
α6
R
α3
β4
α3
β2
α3
35
f2 (ppm)
f1 (ppm)

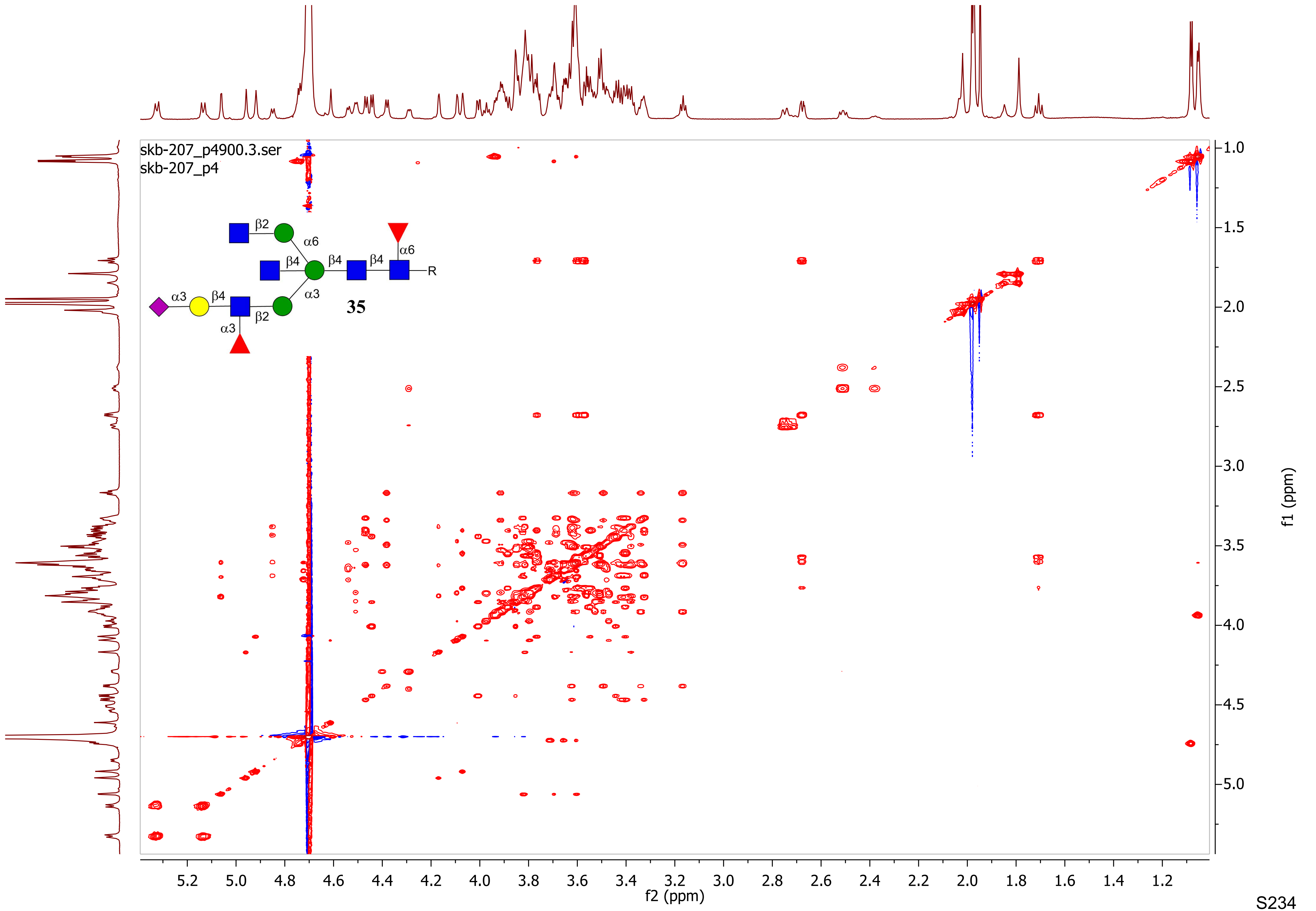

skb-207_p4900.3.ser
skb-207_p4
β2
α6
β4
β4
β4
α6
R
α3
β4
α3
β2
α3
35
f2 (ppm)
f1 (ppm)

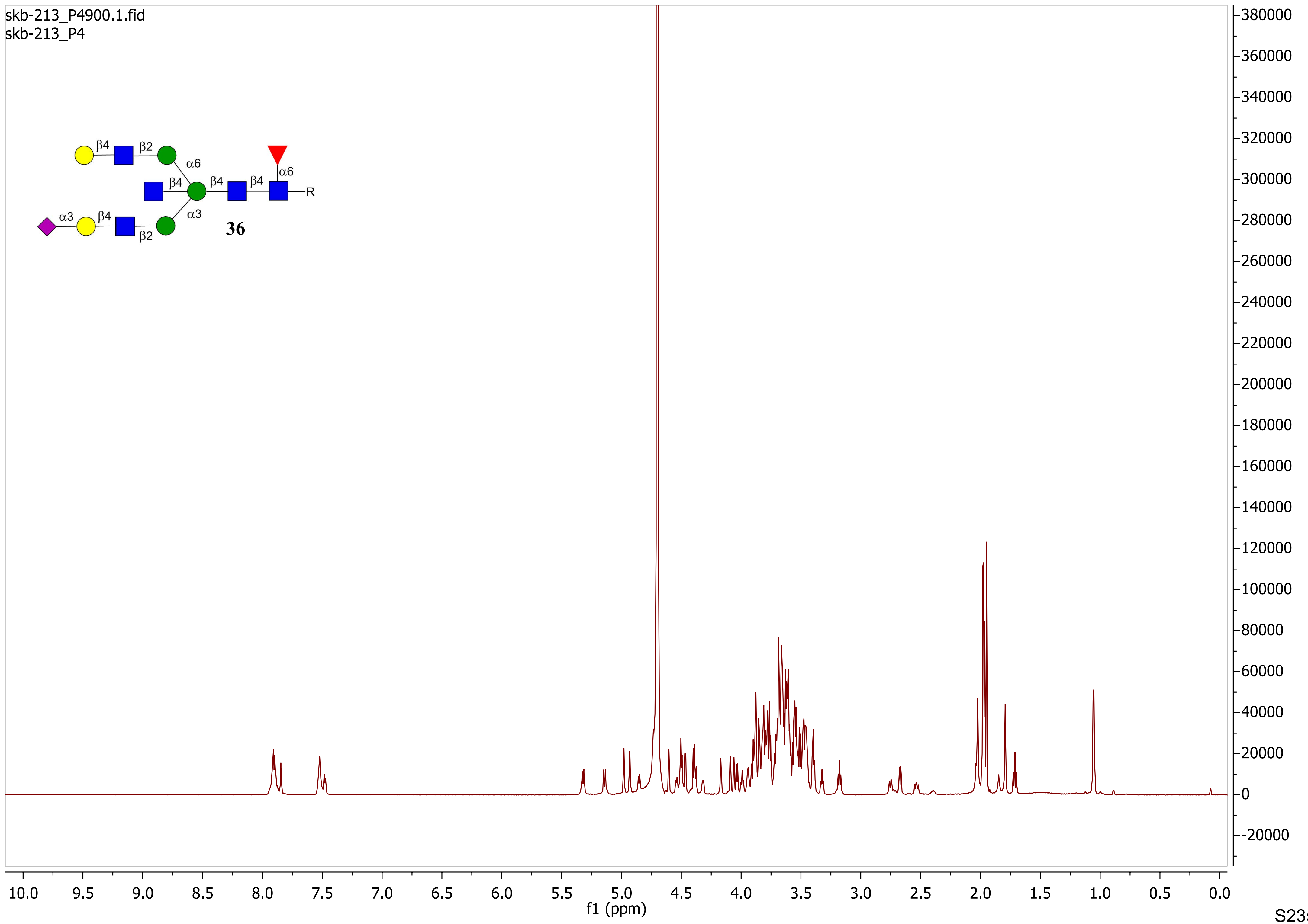
skb-213_P4900.1.fid
skb-213_P4
β4
β2
α6
β4
β4
β4
α6
R
α3
β4
α3
β2
36
f1 (ppm)
10.0 9.5 9.0 8.5 8.0 7.5 7.0 6.5 6.0 5.5 5.0 4.5 4.0 3.5 3.0 2.5 2.0 1.5 1.0 0.5 0.0
380000 360000 340000 320000 300000 280000 260000 240000 220000 200000 180000 160000 140000 120000 100000 80000 60000 40000 20000 0 -20000

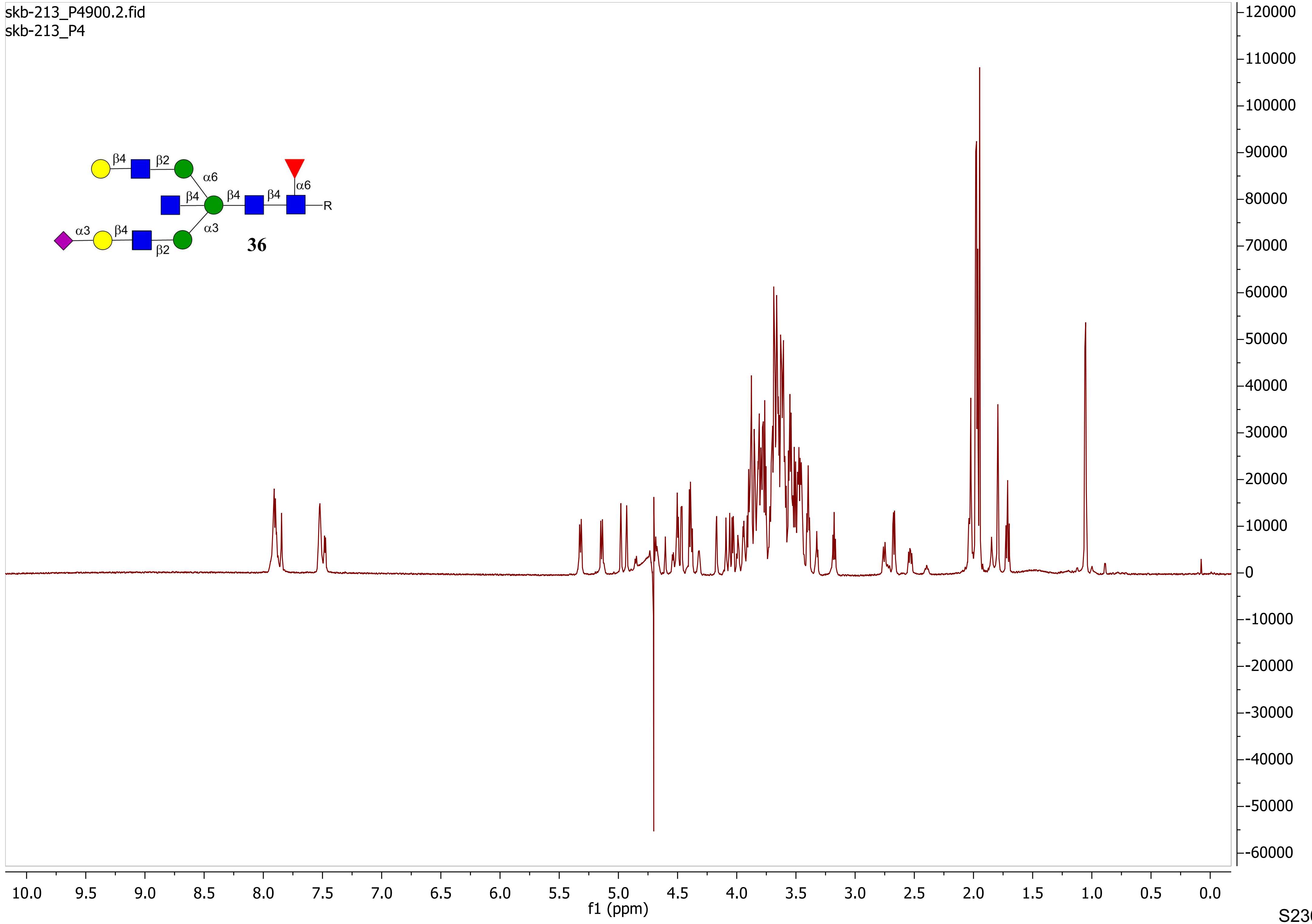

skb-213_P4900.2.fid
skb-213_P4
β4
β2
α6
β4
β4
β4
α6
R
α3
β4
α3
β2
36
f1 (ppm)

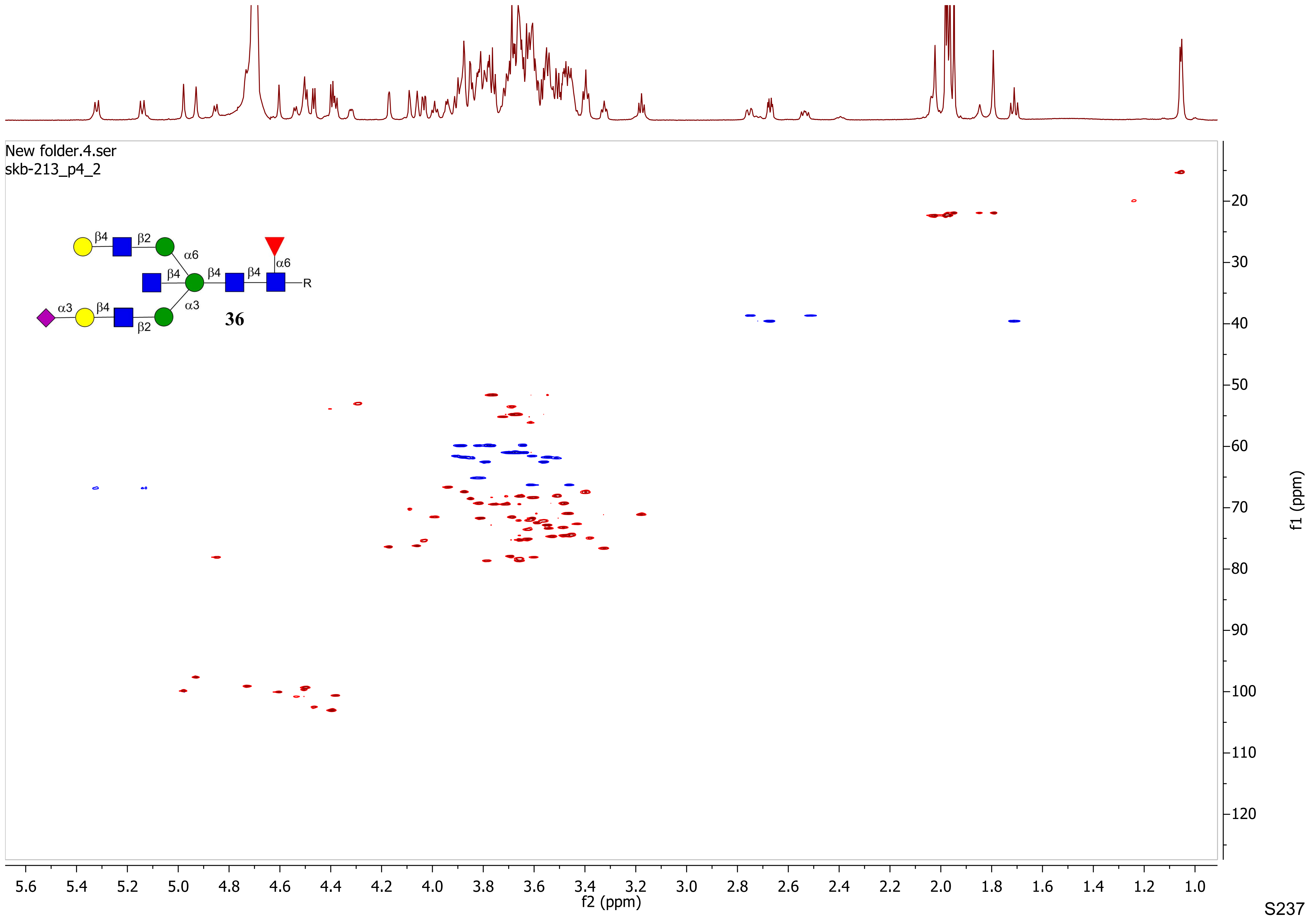
New folder.4.ser
skb-213_p4_2
β4
β2
α6
β4
β4
β4
α6
R
α3
β4
α3
β2
36
f2 (ppm)
f1 (ppm)

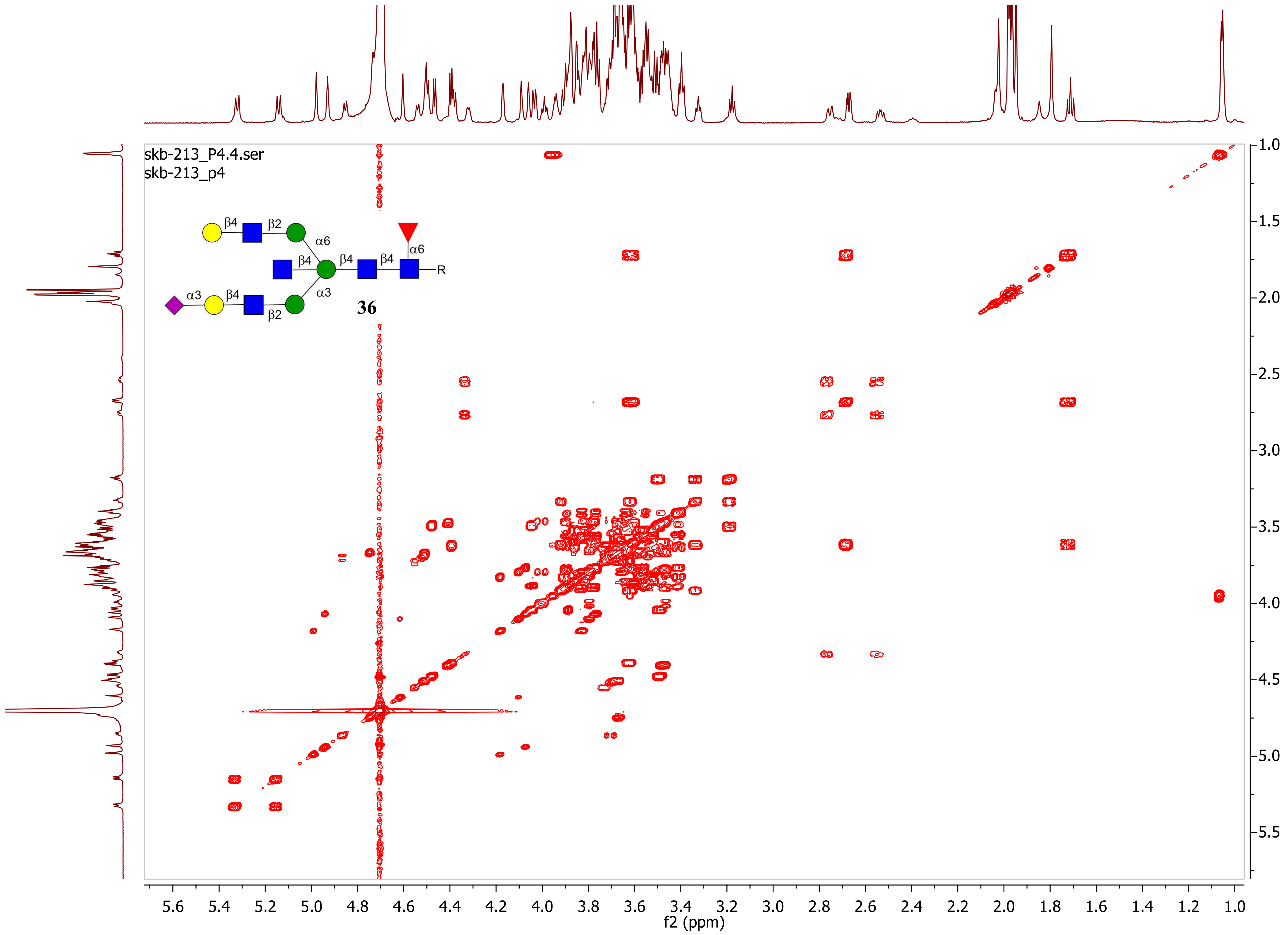

skb-213_P4.4.ser
skb-213_p4
β4
β2
α6
β4
β4
β4
α6
R
α3
β4
α3
β2
36
f2 (ppm)
f1 (ppm)

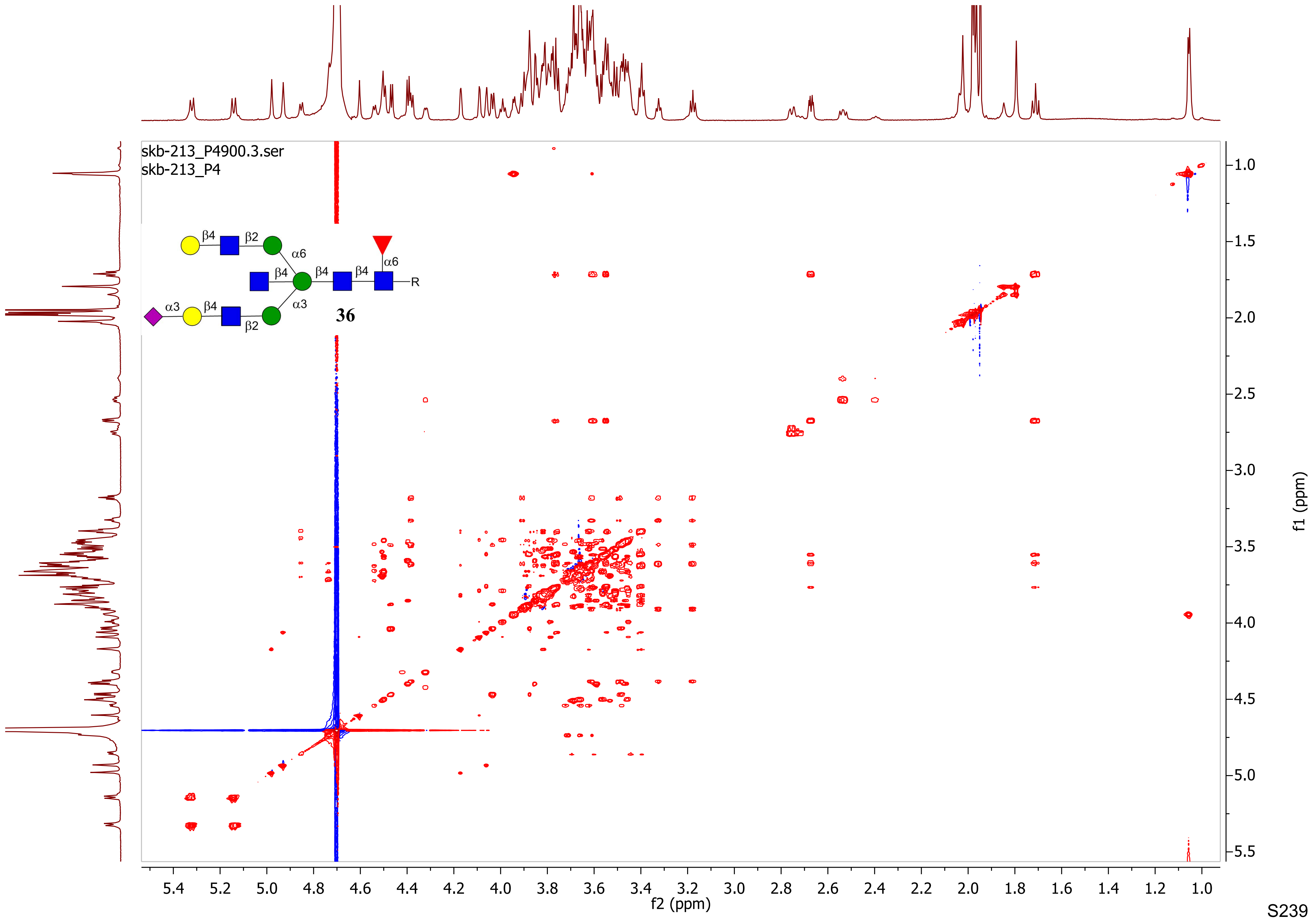
skb-213_P4900.3.ser
skb-213_P4
β4
β2
α6
β4
β4
β4
α6
R
α3
β4
α3
β2
36
f2 (ppm)
f1 (ppm)

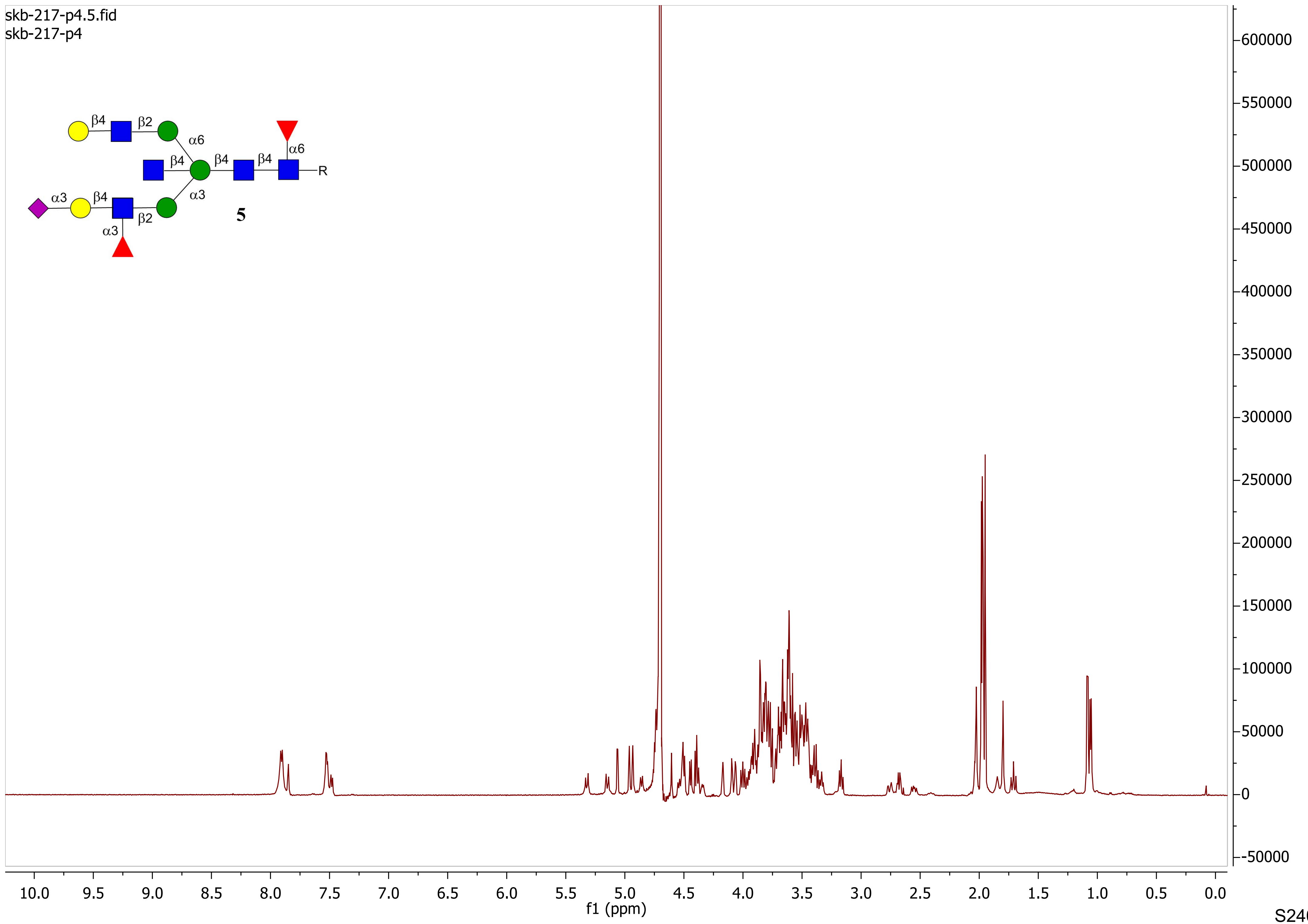
skb-217-p4.5.fid
skb-217-p4
β4
β2
α6
β4
β4
β4
α6
R
α3
β4
α3
β2
α3
5
f1 (ppm)

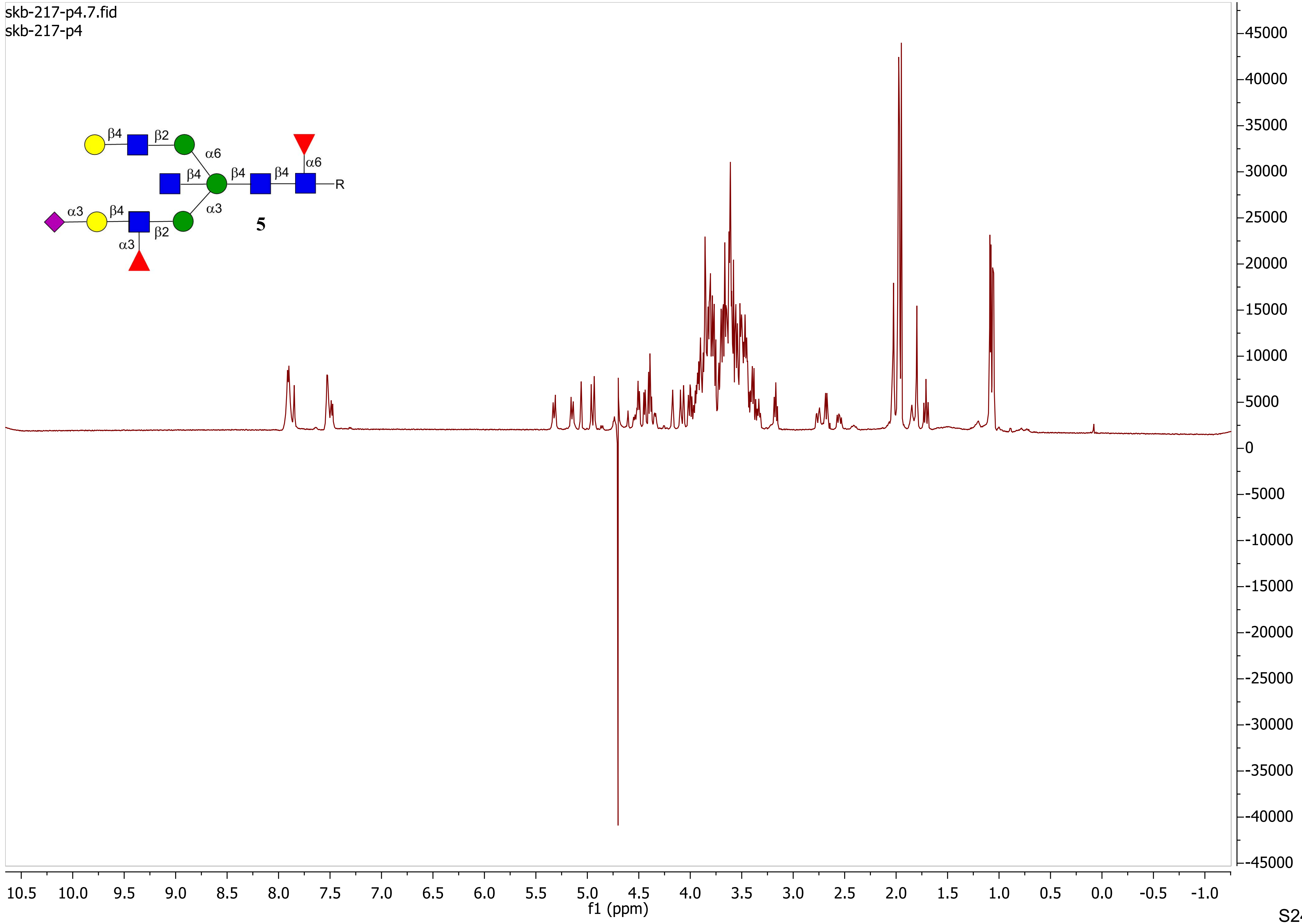

skb-217-p4.7.fid
skb-217-p4
β4
β2
α6
β4
β4
β4
α6
R
α3
β4
α3
β2
α3
5
f1 (ppm)

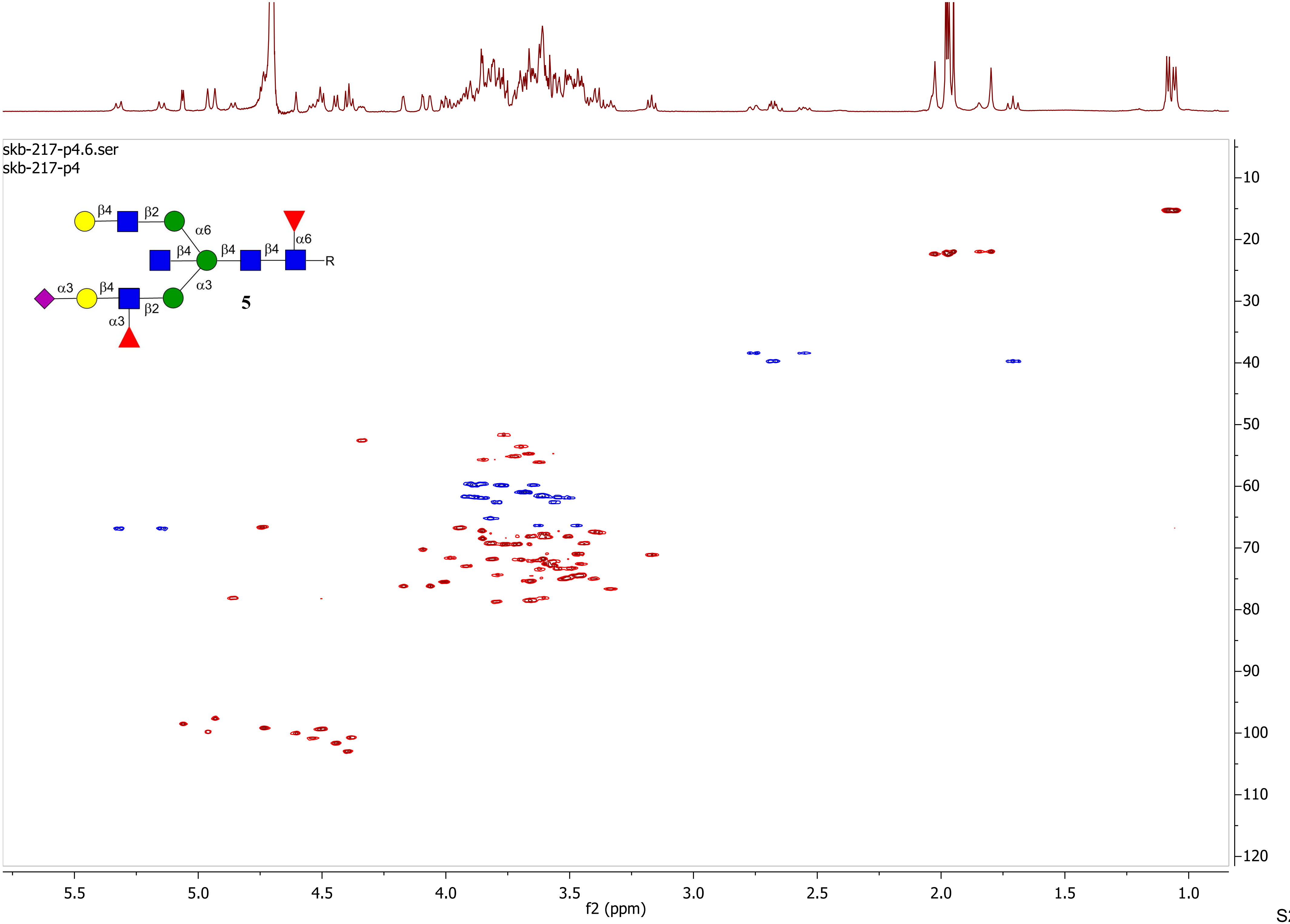
skb-217-p4.6.ser
skb-217-p4
β4
β2
α6
β4
β4
β4
α6
R
α3
β4
α3
β2
α3
5
f1 (ppm)
f2 (ppm)
5.5
5.0
4.5
4.0
3.5
3.0
2.5
2.0
1.5
1.0
10
20
30
40
50
60
70
80
90
100
110
120

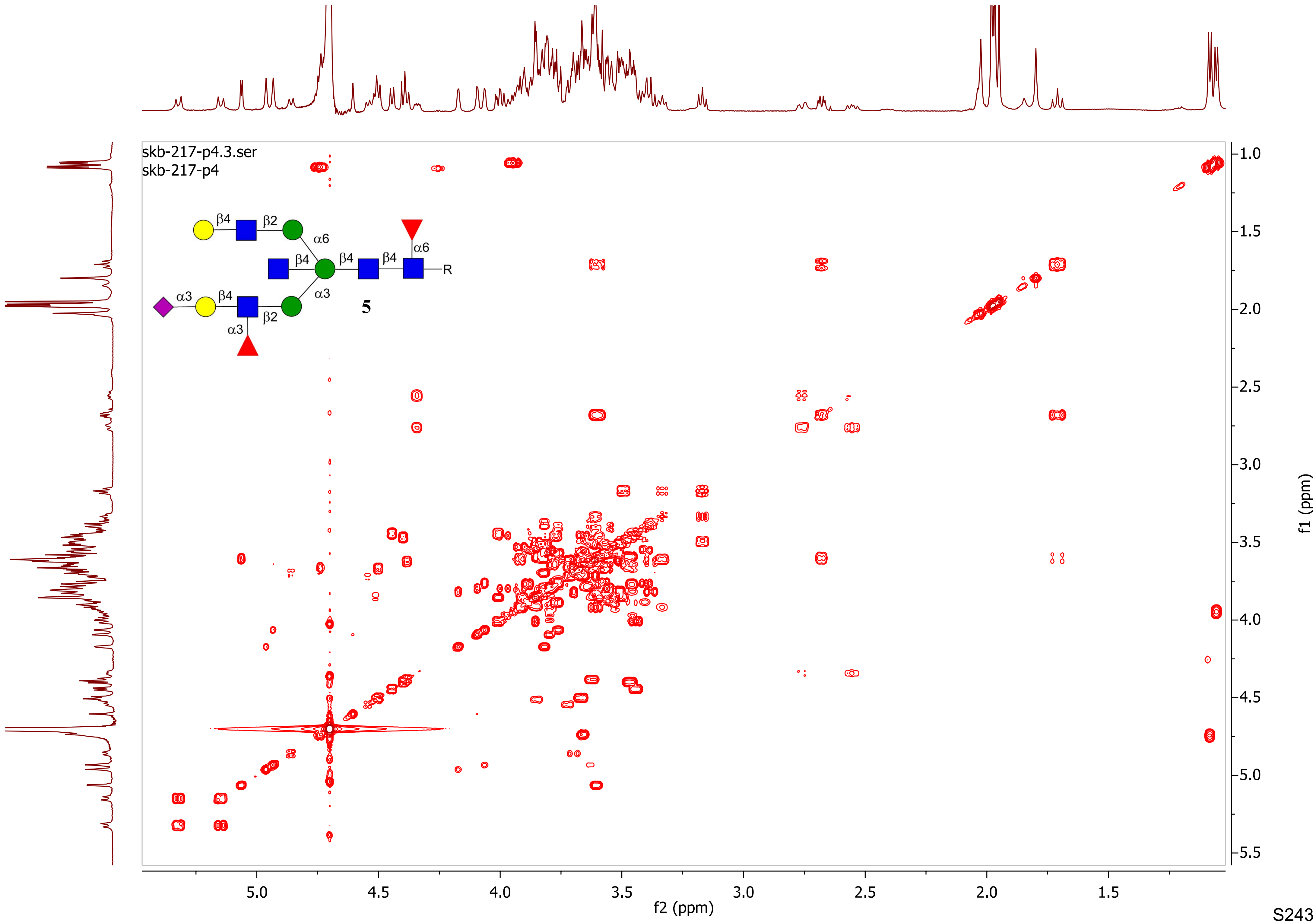
skb-217-p4.3.ser
skb-217-p4
β4
β2
α6
β4
β4
β4
α6
R
α3
β4
α3
β2
α3
5
f1 (ppm)
f2 (ppm)

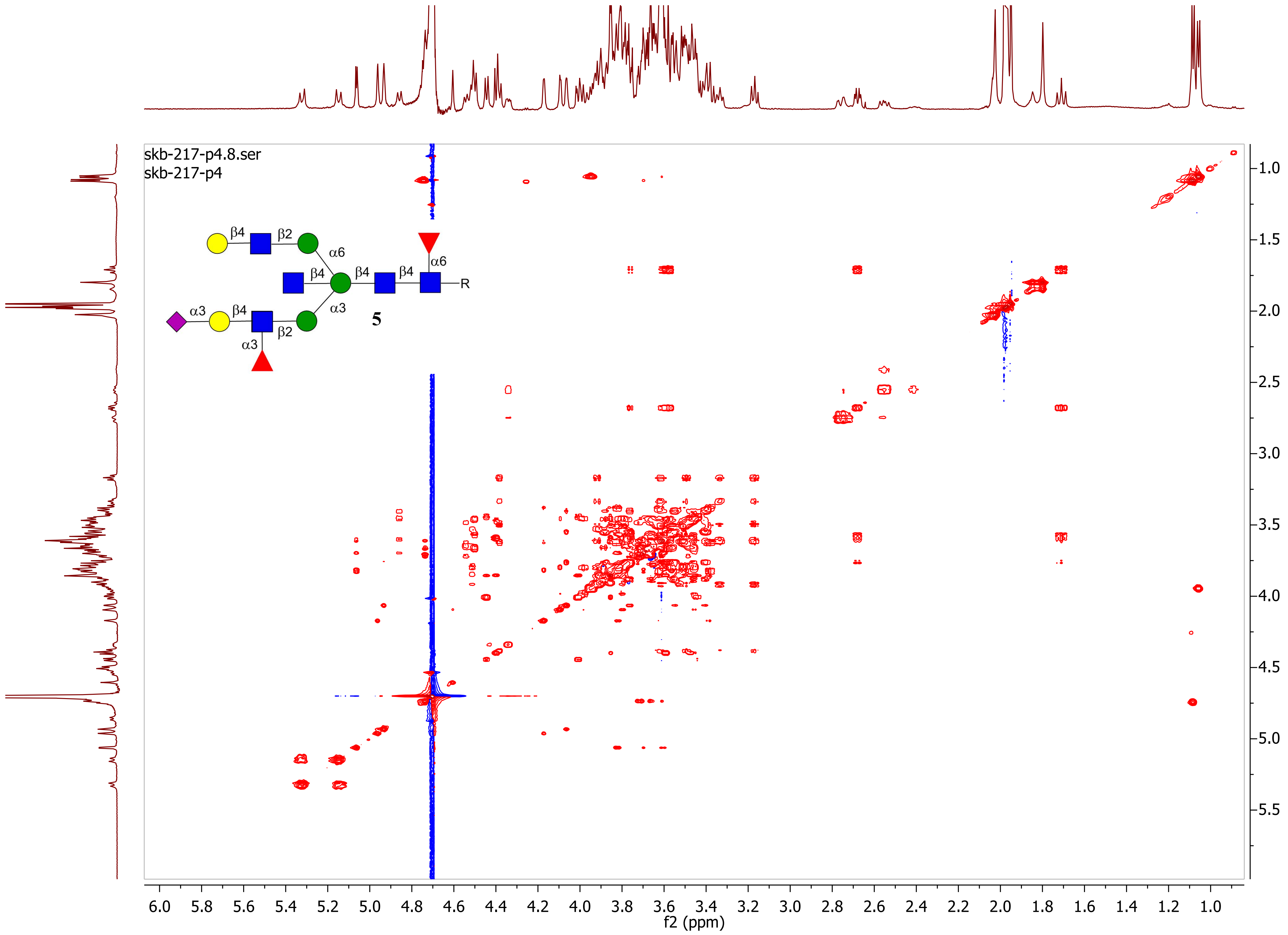

skb-217-p4.8.ser
skb-217-p4
β4
β2
α6
β4
β4
β4
α6
R
α3
β4
α3
α3
β2
5
f1 (ppm)
f2 (ppm)

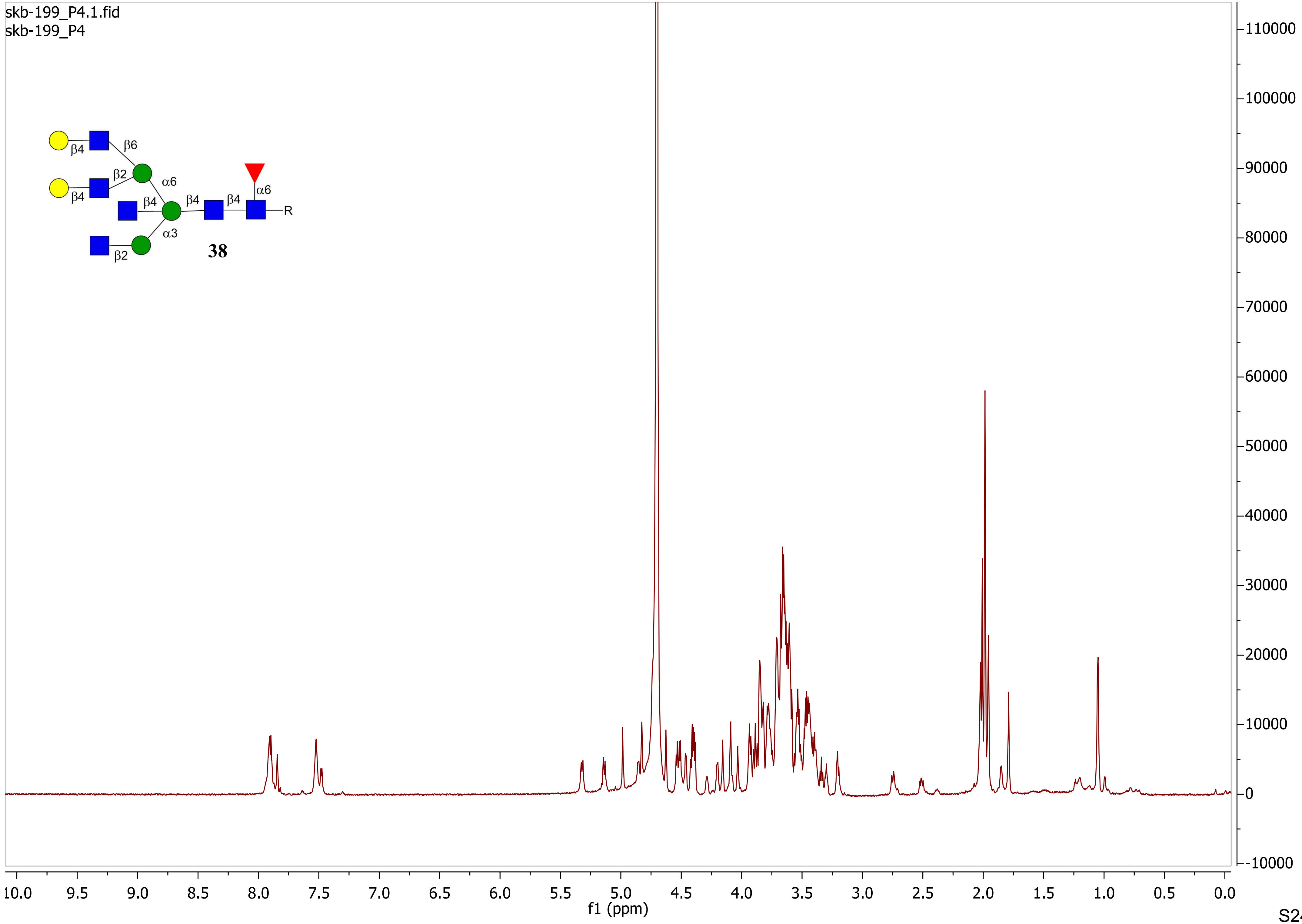
skb-199_P4.1.fid
skb-199_P4
β4
β6
β2
α6
β4
β4
β4
β4
α6
R
α3
β2
38
f1 (ppm)
10.0
9.5
9.0
8.5
8.0
7.5
7.0
6.5
6.0
5.5
5.0
4.5
4.0
3.5
3.0
2.5
2.0
1.5
1.0
0.5
0.0
110000
100000
90000
80000
70000
60000
50000
40000
30000
20000
10000
0
-10000

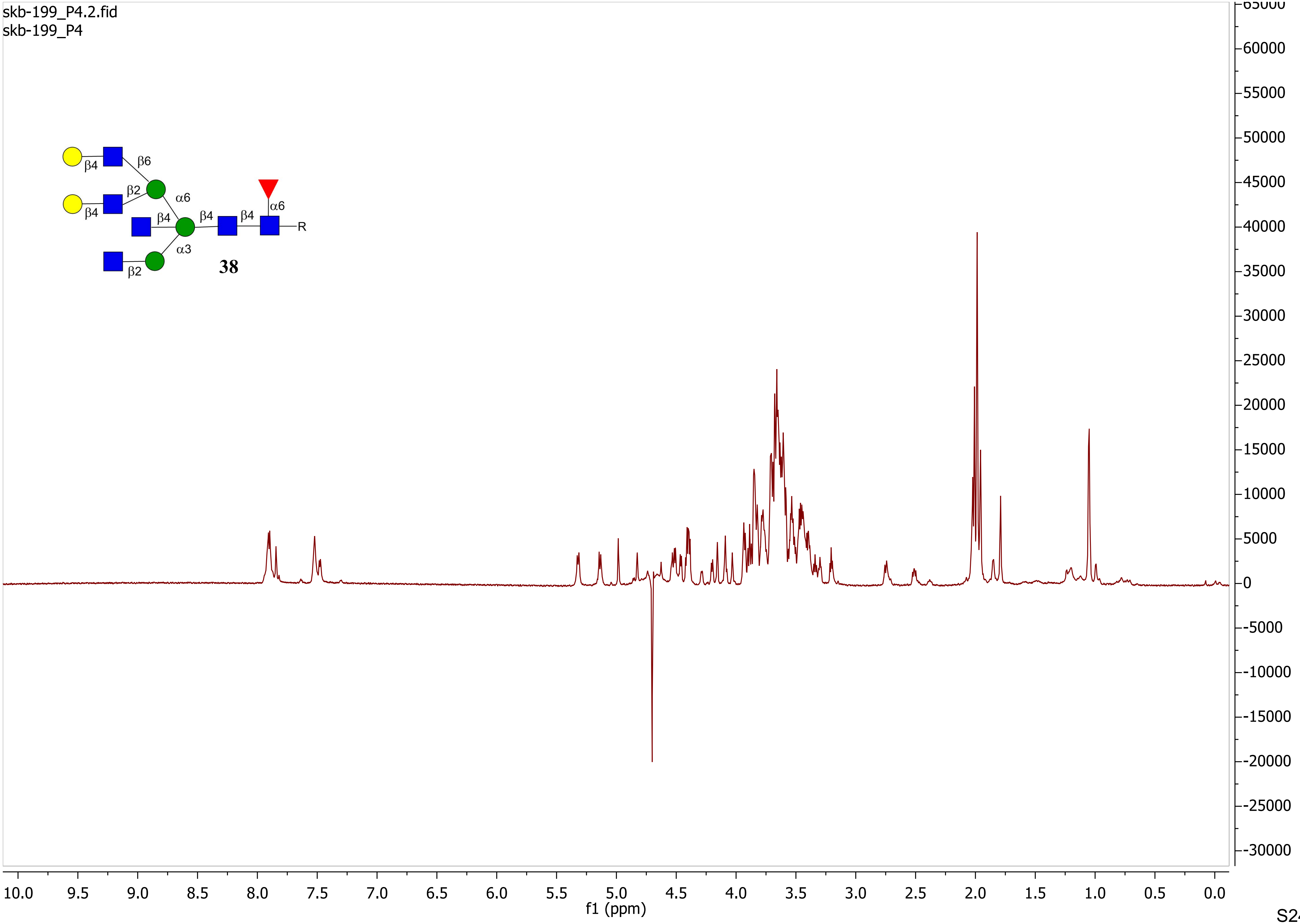
skb-199_P4.2.fid
skb-199_P4
β4
β6
β2
α6
β4
β4
β4
β4
α6
R
α3
β2
38
f1 (ppm)

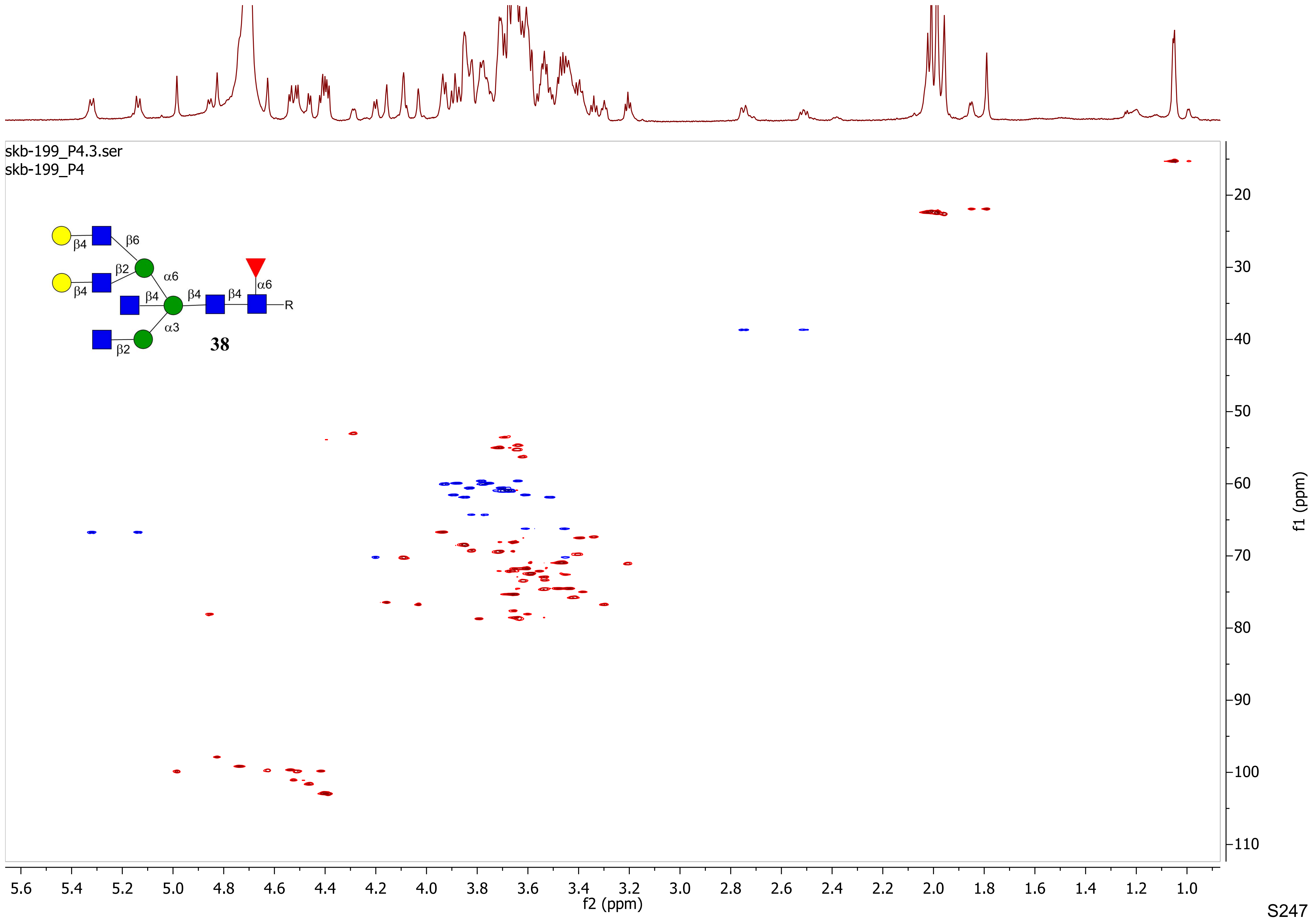
skb-199_P4.3.ser
skb-199_P4
β4
β6
β2
β4
α6
β4
β4
β4
α6
R
α3
β2
38
f2 (ppm)
f1 (ppm)

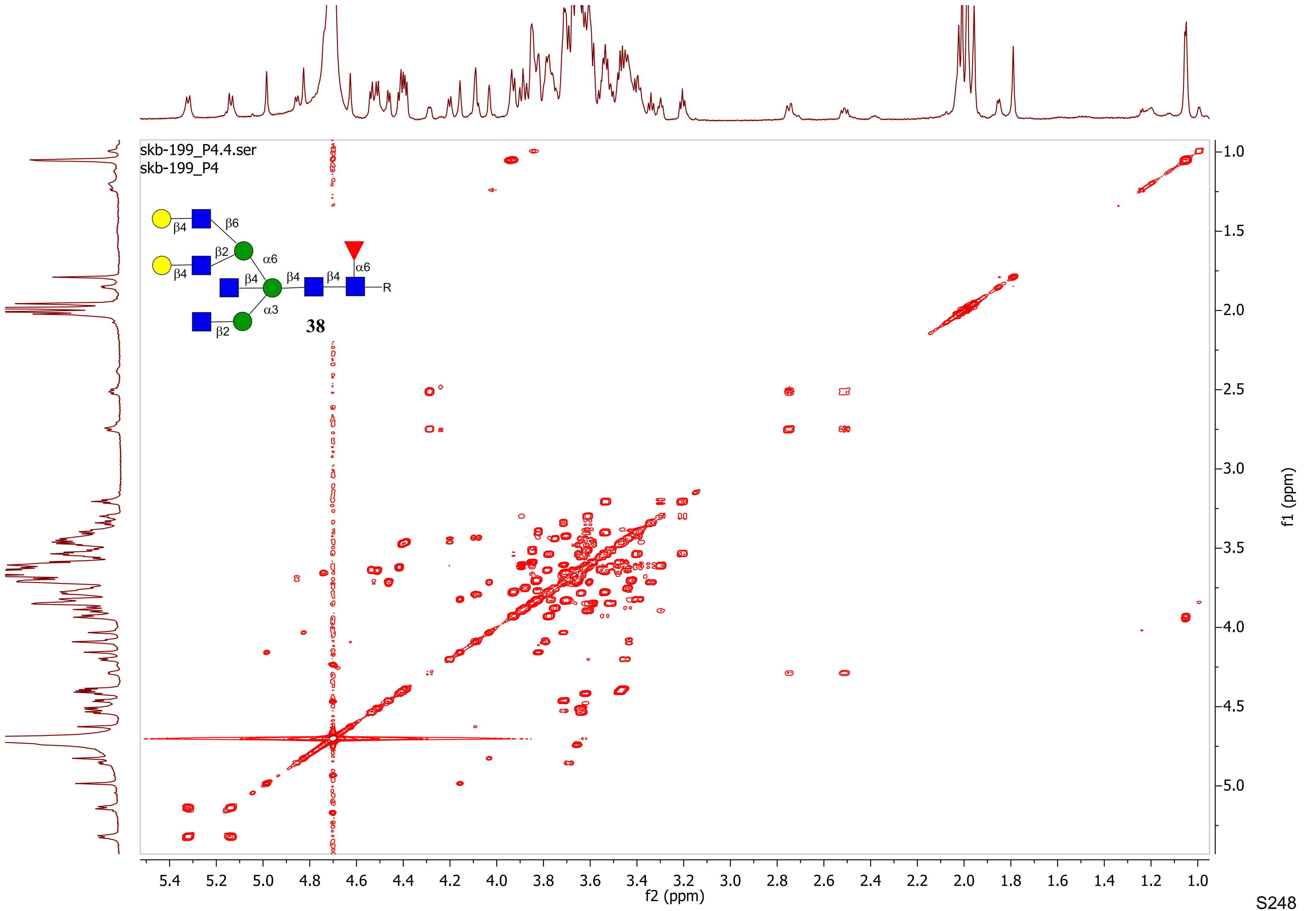
skb-199_P4.4.ser
skb-199_P4
β4
β6
β2
α6
β4
β4
β4
β4
α6
R
α3
β2
38
f2 (ppm)
f1 (ppm)

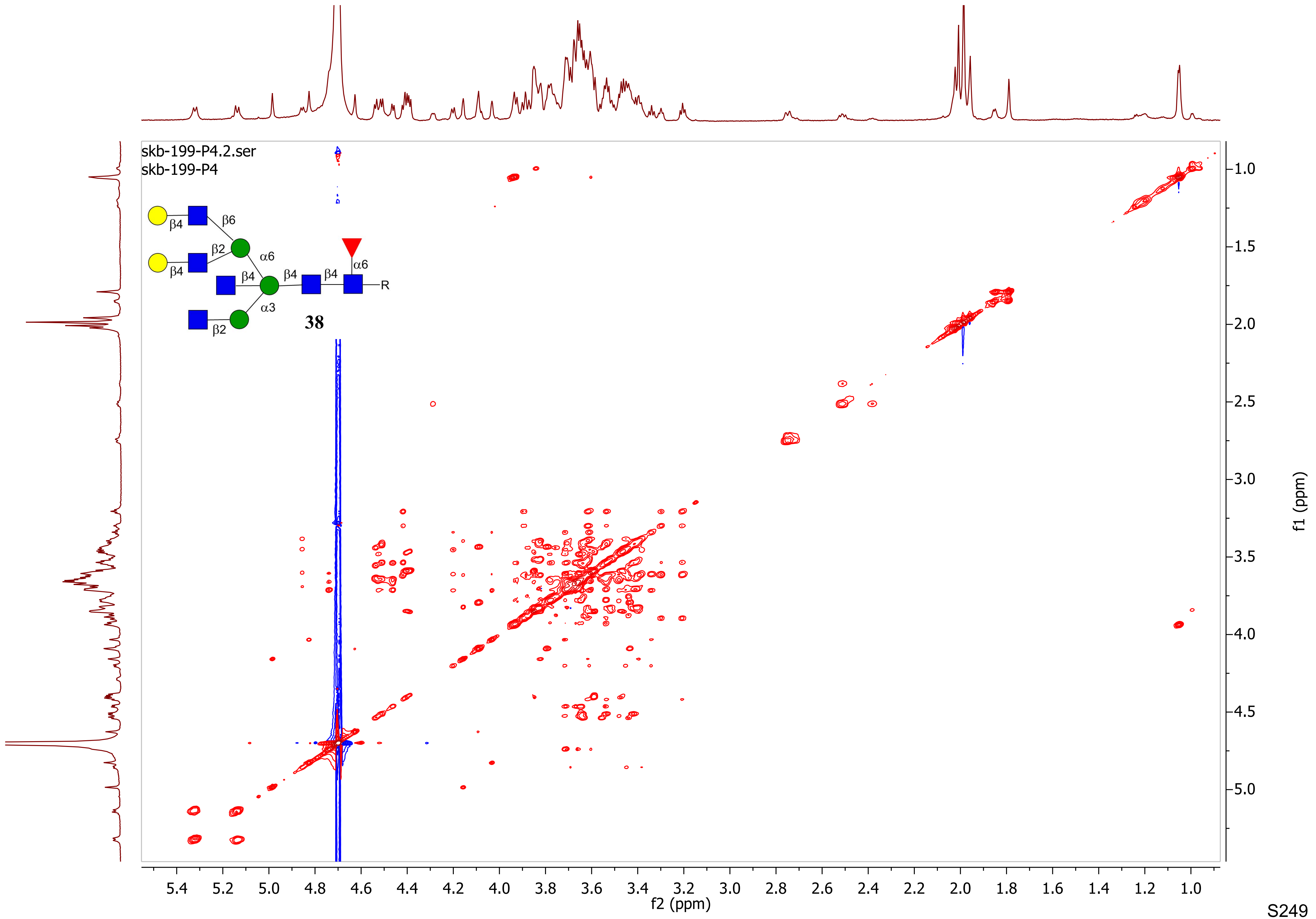
skb-199-P4.2.ser
skb-199-P4
β4
β6
β2
α6
β4
β4
β4
α6
R
β4
α3
β2
38
f2 (ppm)
f1 (ppm)

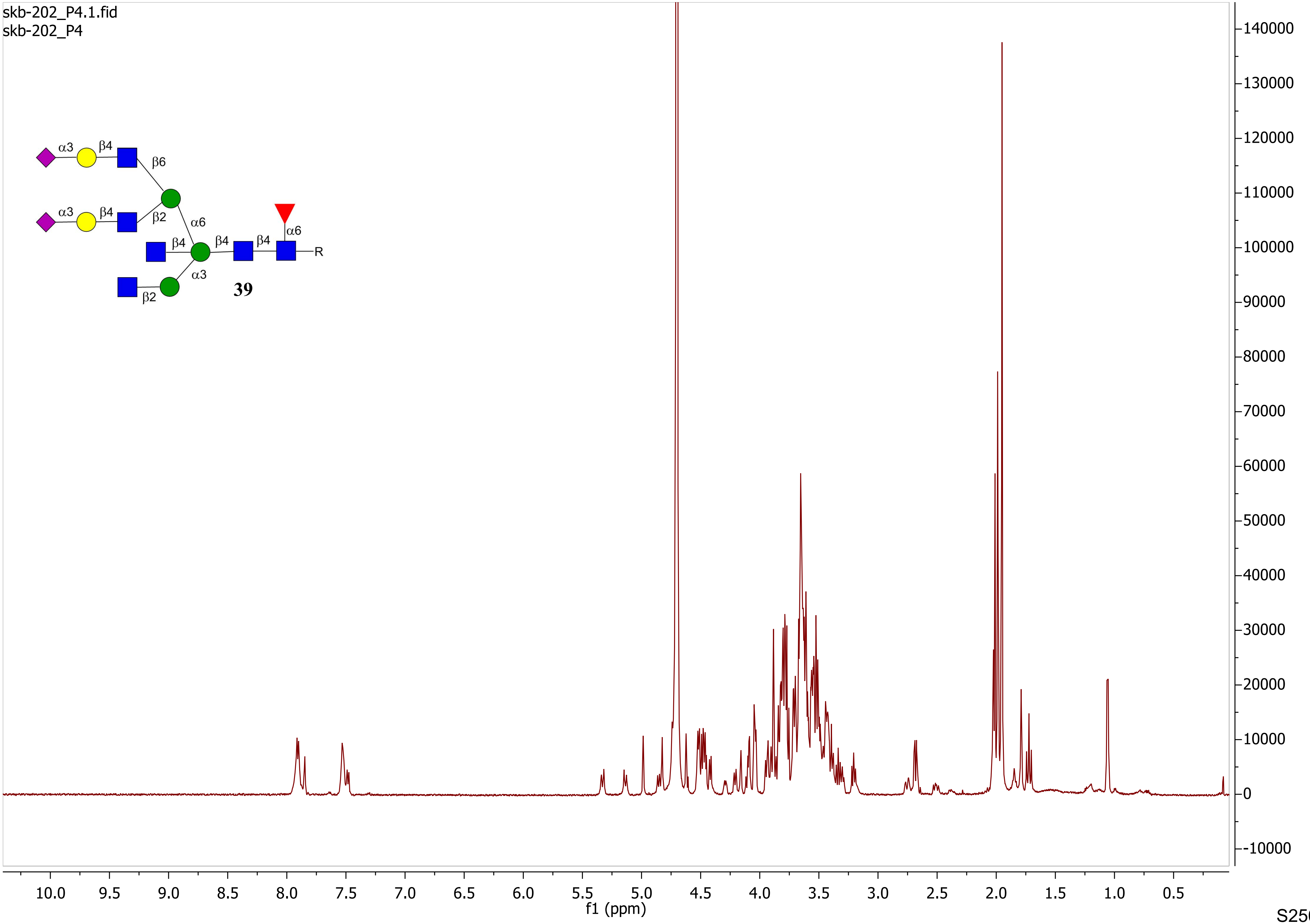

skb-202_P4.1.fid
skb-202_P4
α3
β4
β6
α3
β4
β2
α6
β4
β4
β4
α6
R
α3
β2
39
10.0
9.5
9.0
8.5
8.0
7.5
7.0
6.5
6.0
5.5
5.0
4.5
4.0
3.5
3.0
2.5
2.0
1.5
1.0
0.5
f1 (ppm)
140000
130000
120000
110000
100000
90000
80000
70000
60000
50000
40000
30000
20000
10000
0
-10000

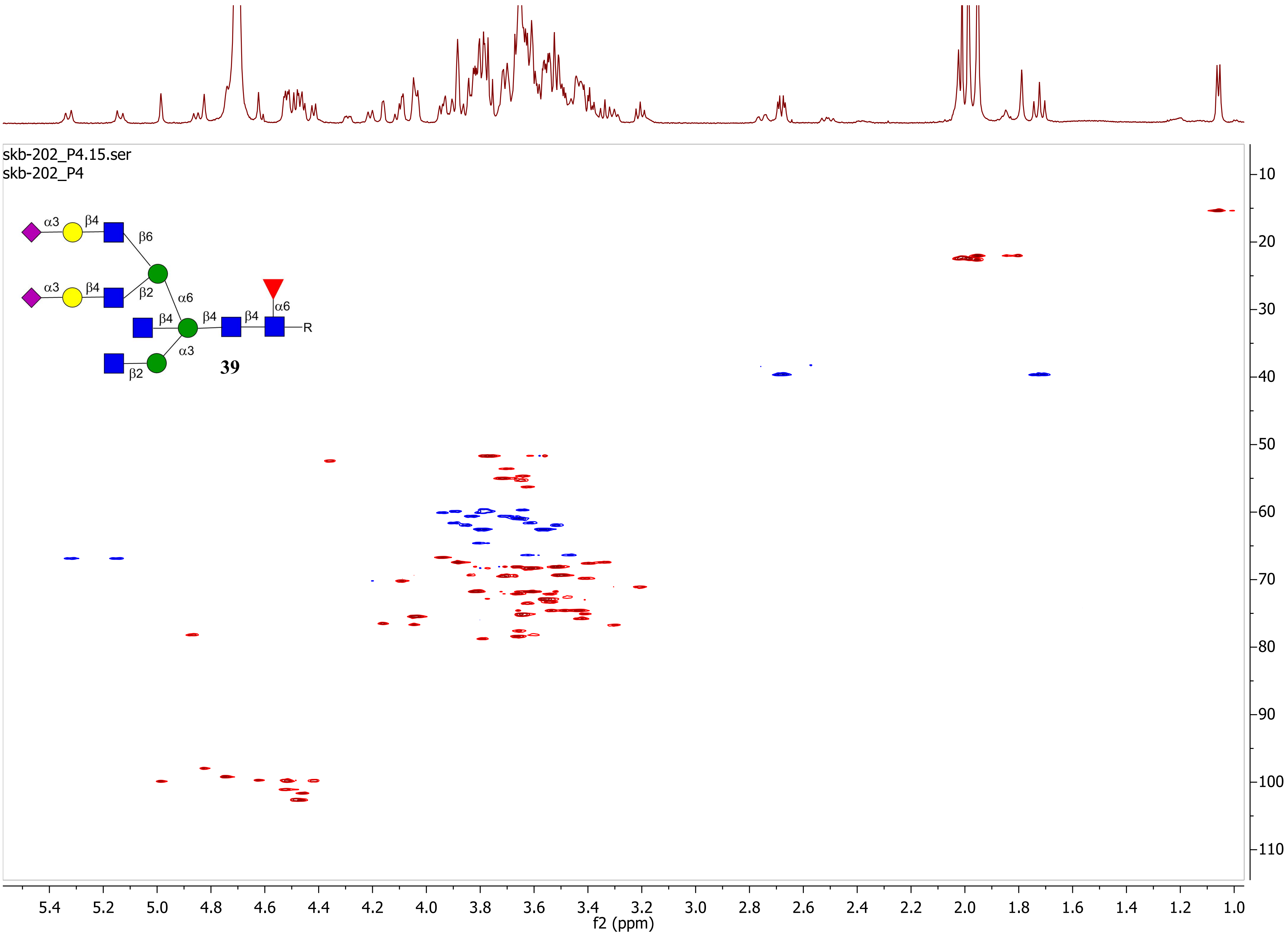

skb-202_P4.15.ser
skb-202_P4
α3
β4
β6
α3
β4
β2
α6
β4
β4
β4
α6
R
α3
β2
39
f2 (ppm)
f1 (ppm)

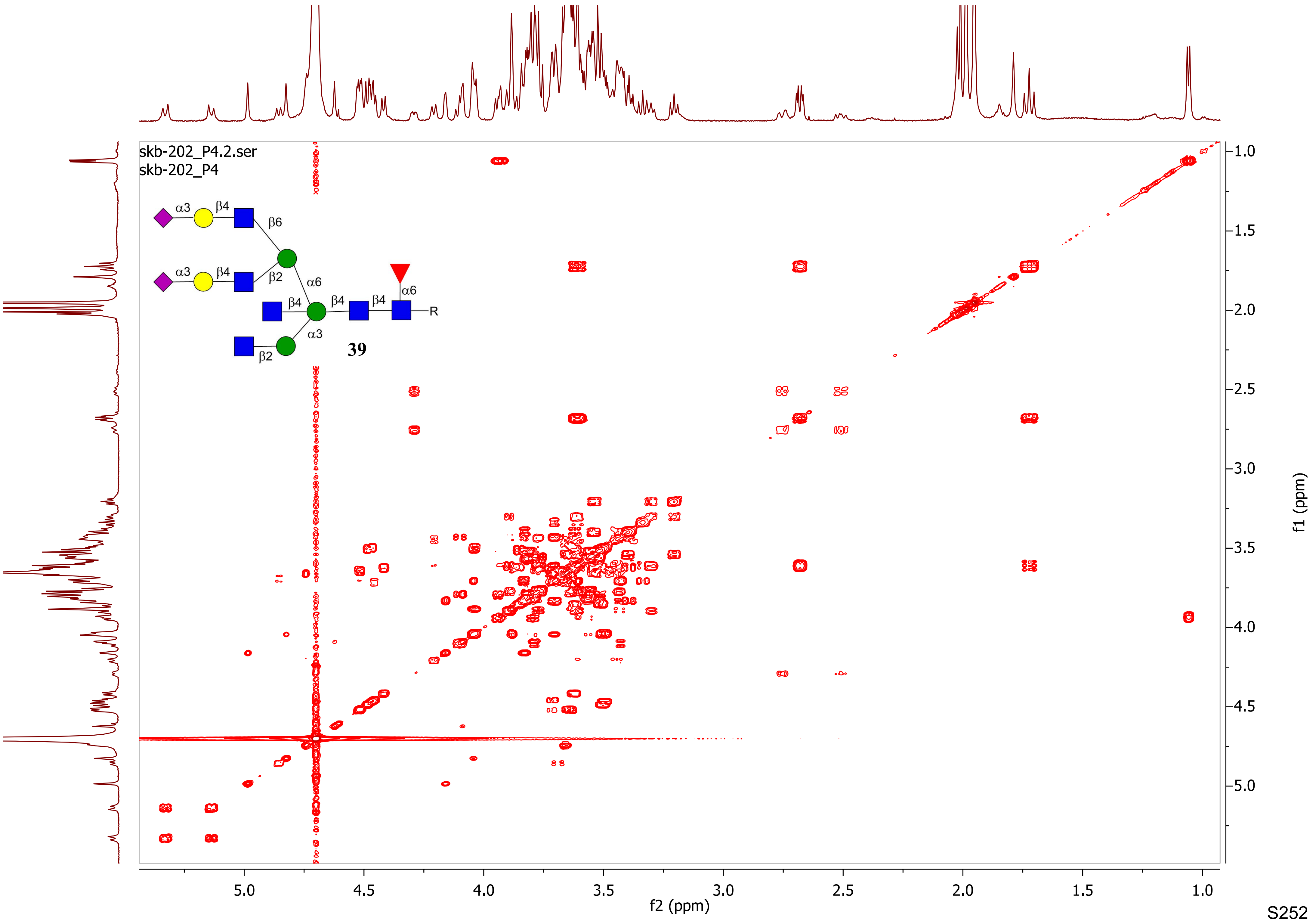
skb-202_P4.2.ser
skb-202_P4
α3
β4
β6
α3
β4
β2
α6
β4
β4
β4
α6
R
α3
β2
39
f2 (ppm)
f1 (ppm)
5.0
4.5
4.0
3.5
3.0
2.5
2.0
1.5
1.0

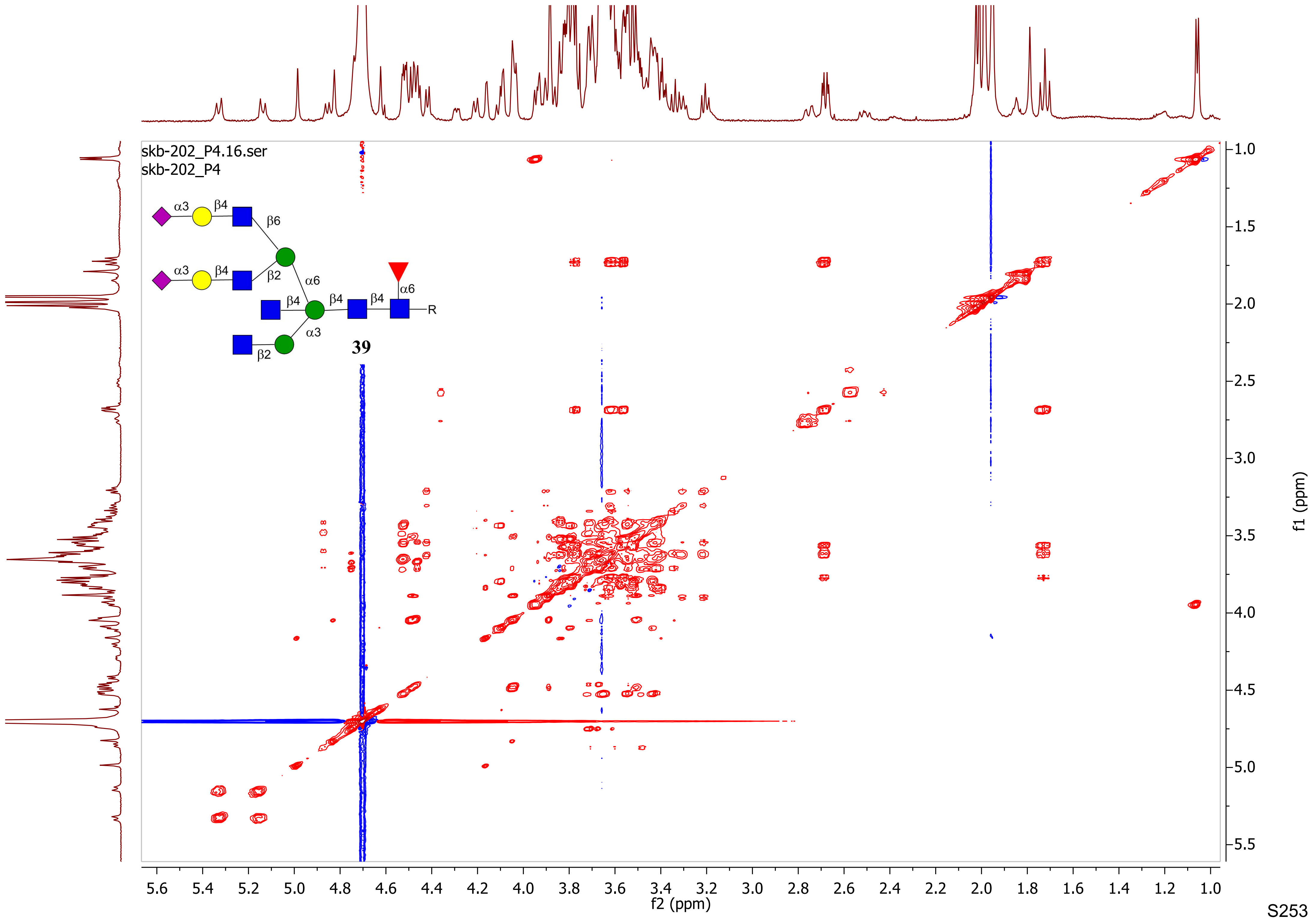
skb-202_P4.16.ser
skb-202_P4
α3
β4
β6
β2
α6
β4
β4
β4
α6
R
α3
β2
39
f2 (ppm)
f1 (ppm)

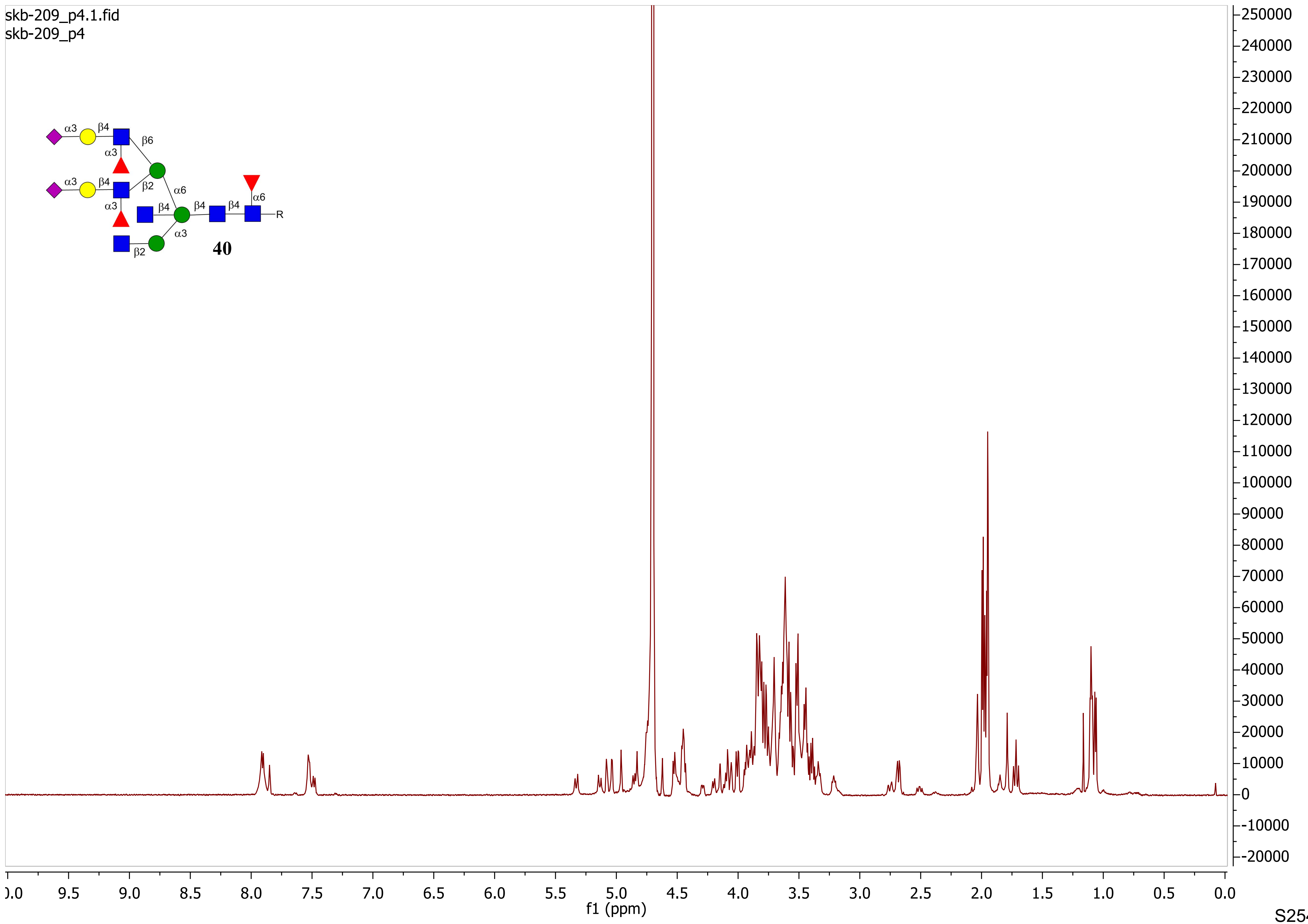

skb-209_p4.1.fid
skb-209_p4
α3
β4
β6
α3
β2
α6
α6
β4
β4
β4
α3
α3
β2
R
40
f1 (ppm)

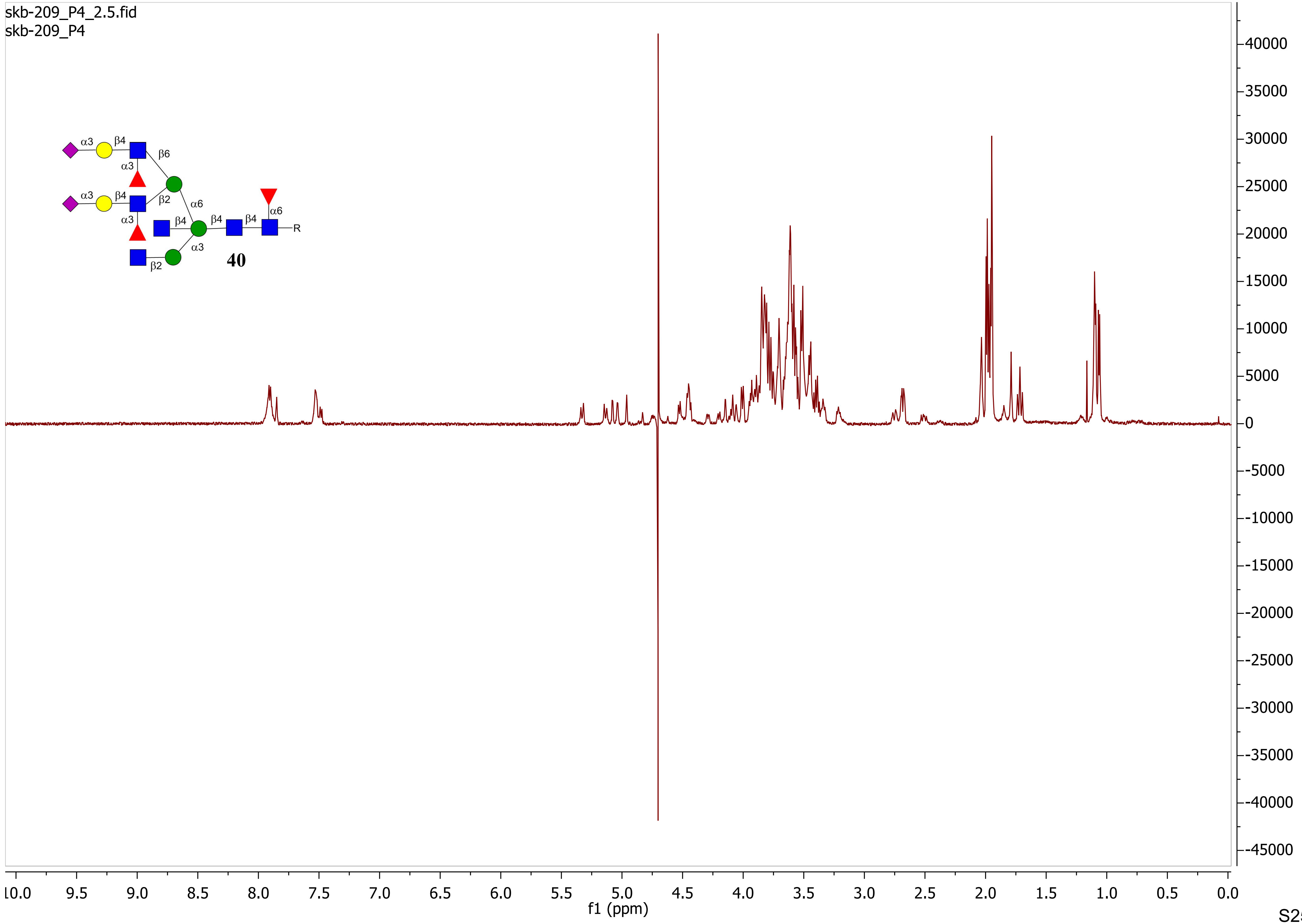
skb-209_P4_2.5.fid
skb-209_P4
α3
β4
β6
α3
α3
β4
β2
α6
α3
β4
β4
β4
α6
R
α3
β2
40
40000
35000
30000
25000
20000
15000
10000
5000
0
-5000
-10000
-15000
-20000
-25000
-30000
-35000
-40000
-45000
10.0
9.5
9.0
8.5
8.0
7.5
7.0
6.5
6.0
5.5
5.0
4.5
4.0
3.5
3.0
2.5
2.0
1.5
1.0
0.5
0.0
f1 (ppm)

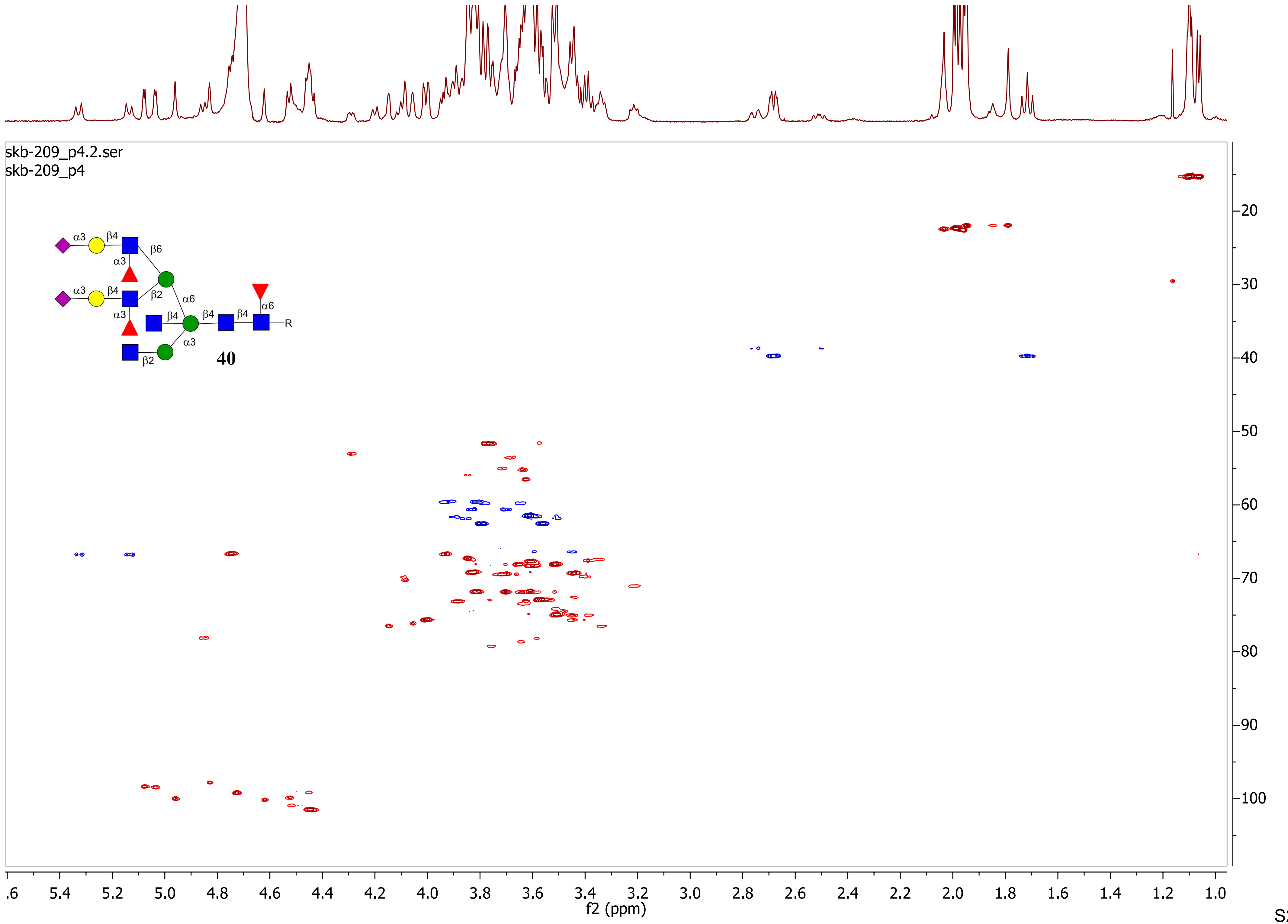
skb-209_p4.2.ser
skb-209_p4
α3
β4
β6
α3
α3
β4
β2
α6
α6
β4
β4
β4
α3
α3
R
β2
40
f2 (ppm)
f1 (ppm)

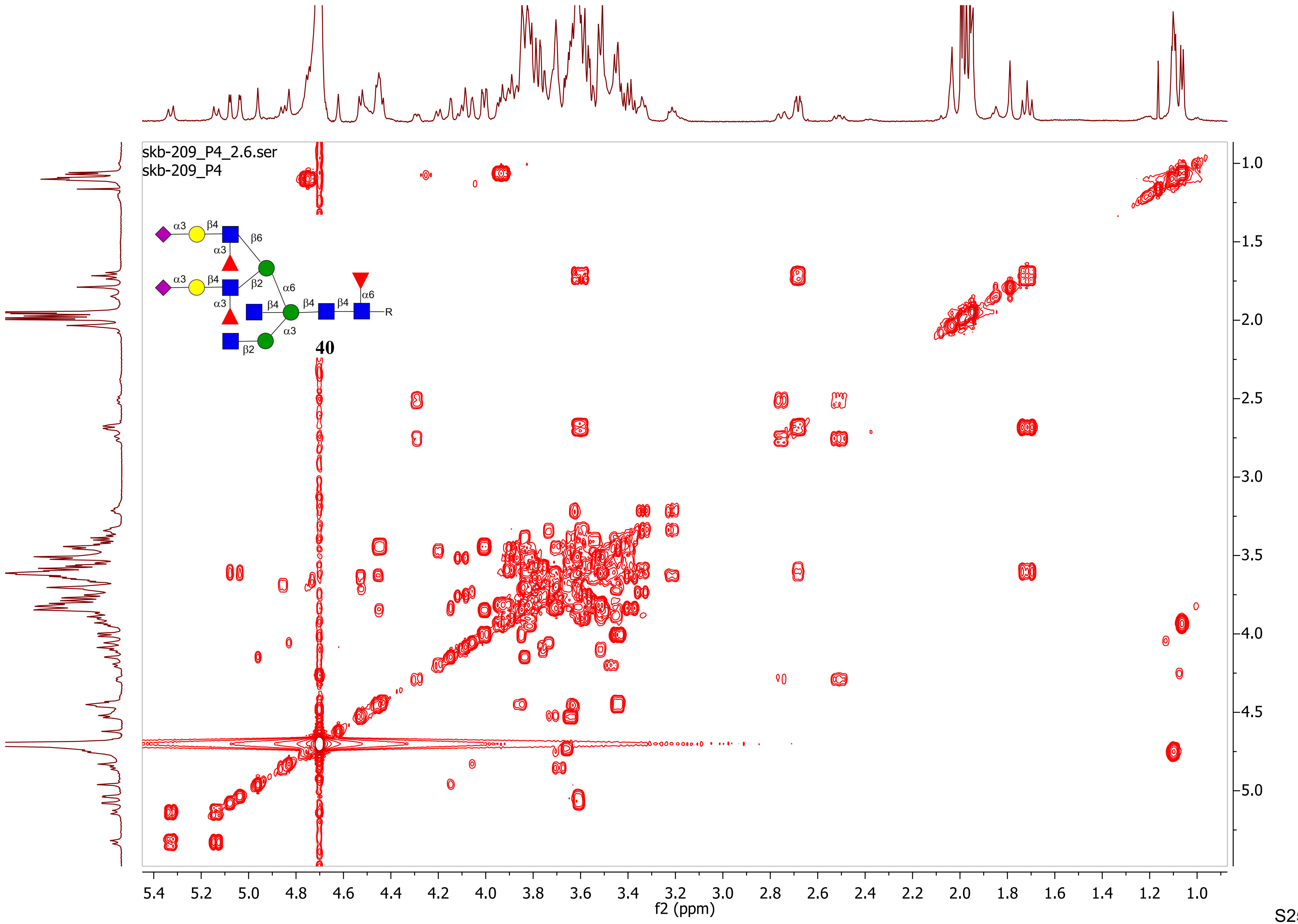

skb-209_P4_2.6.ser
skb-209_P4
40
R
f2 (ppm)
f1 (ppm)

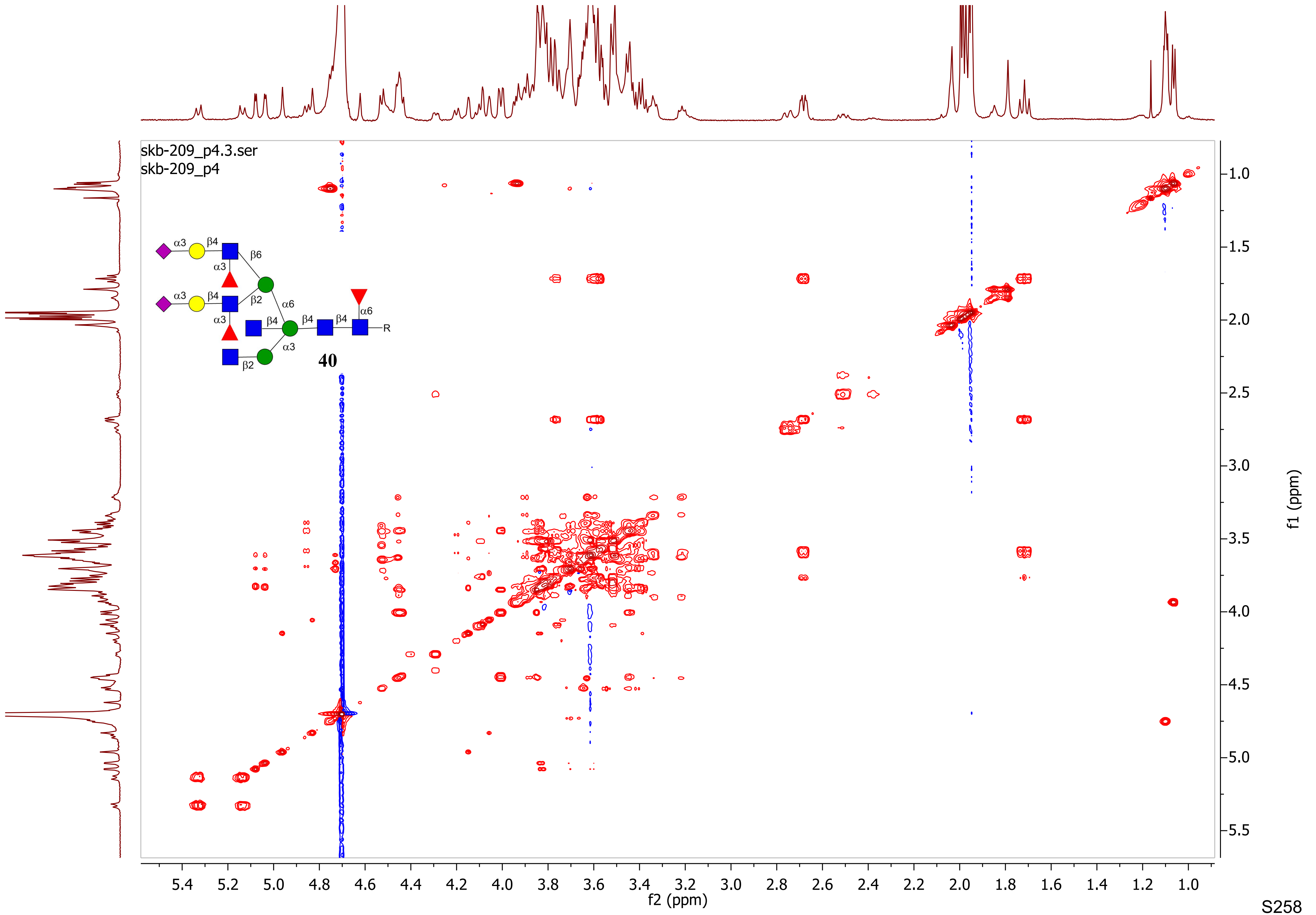

skb-209_p4.3.ser
skb-209_p4
α3
β4
β6
α3
α3
β4
β2
α6
α3
β4
β4
β4
α6
R
α3
β2
40
f2 (ppm)
f1 (ppm)

skb-210_P4.1.fid
skb-210_P4

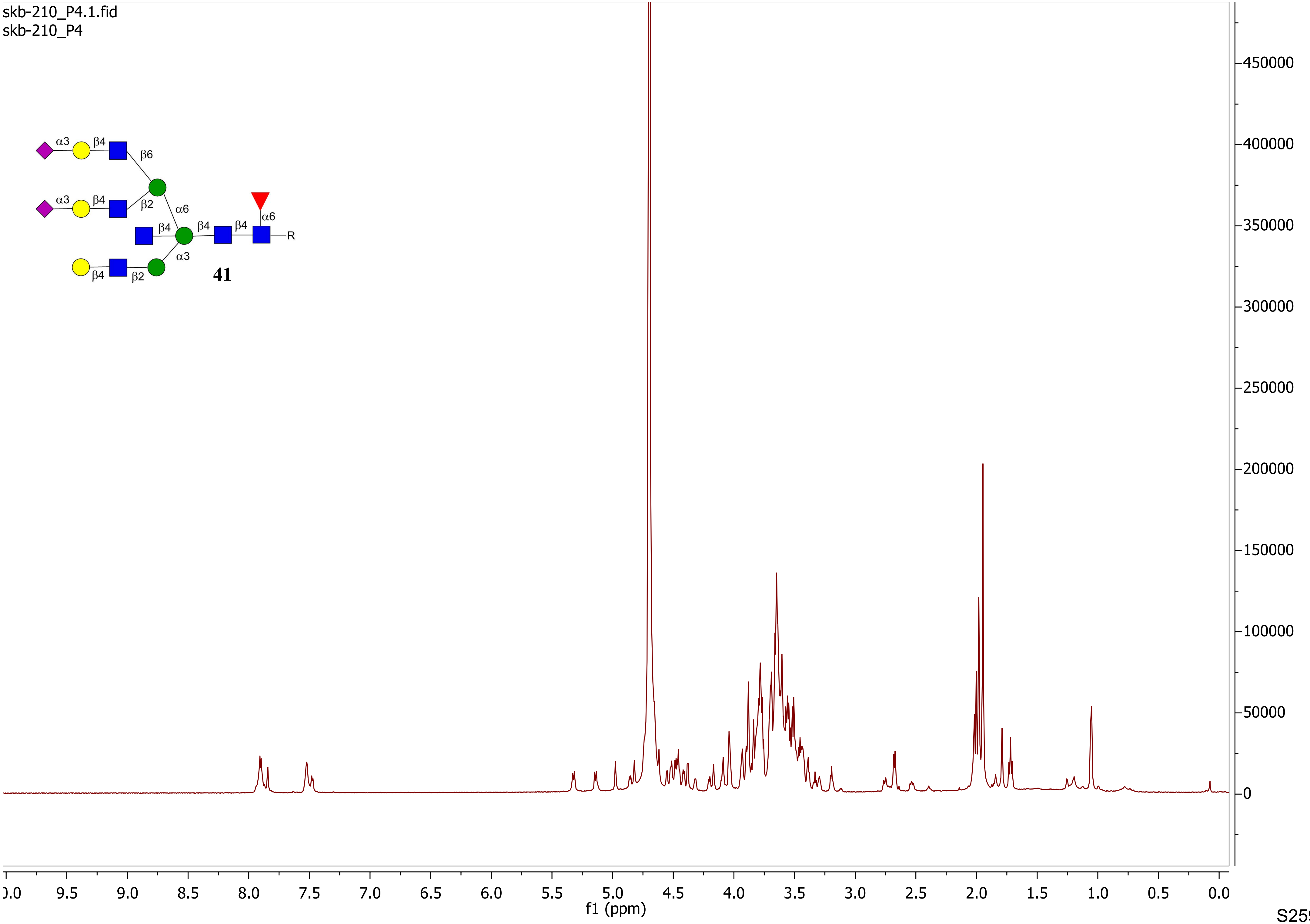

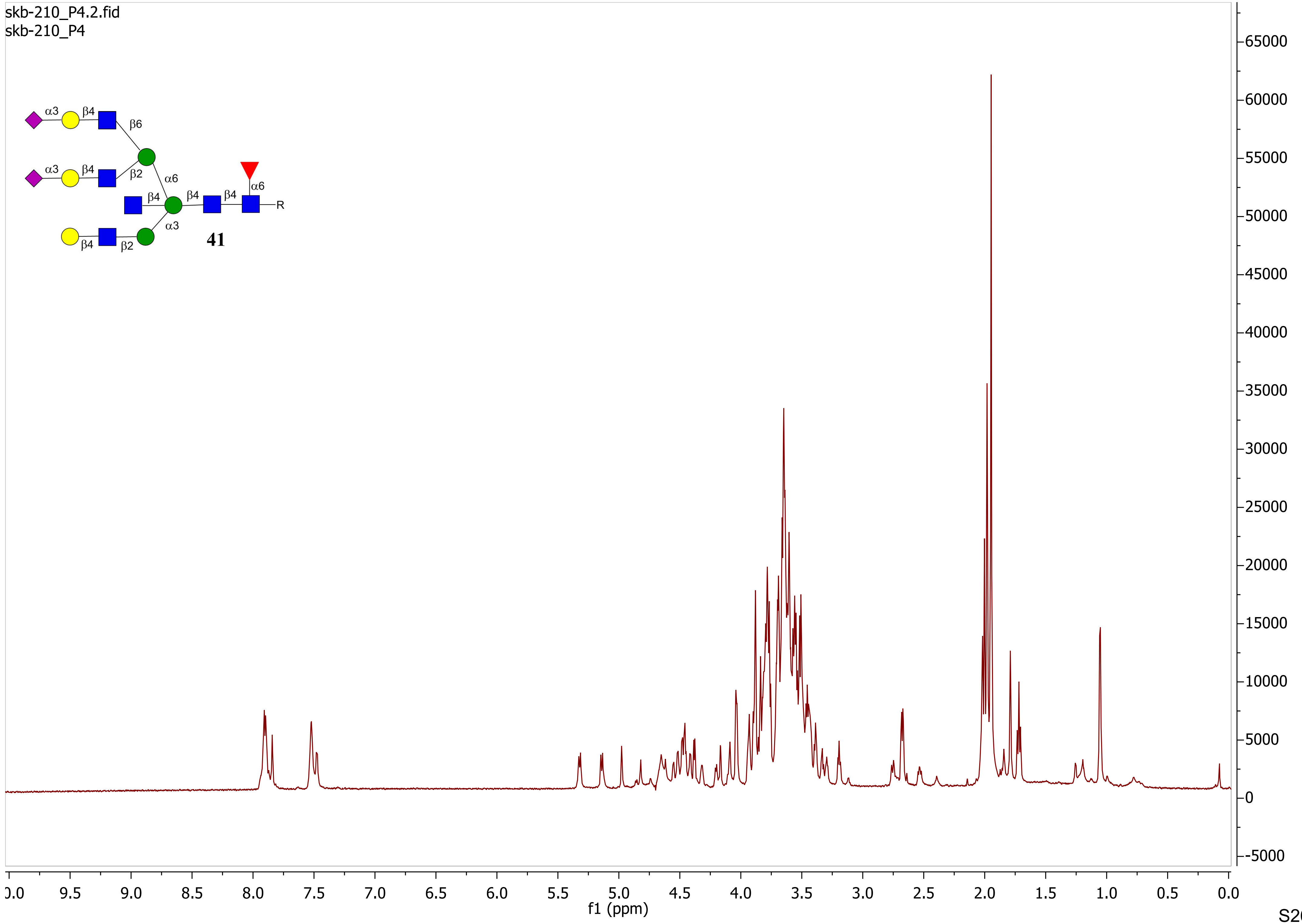
skb-210_P4.2.fid
skb-210_P4
α3
β4
β6
β2
α6
β4
β4
β4
α6
R
α3
β4
β2
41
65000
60000
55000
50000
45000
40000
35000
30000
25000
20000
15000
10000
5000
0
-5000
9.5
9.0
8.5
8.0
7.5
7.0
6.5
6.0
5.5
5.0
4.5
4.0
3.5
3.0
2.5
2.0
1.5
1.0
0.5
0.0
f1 (ppm)

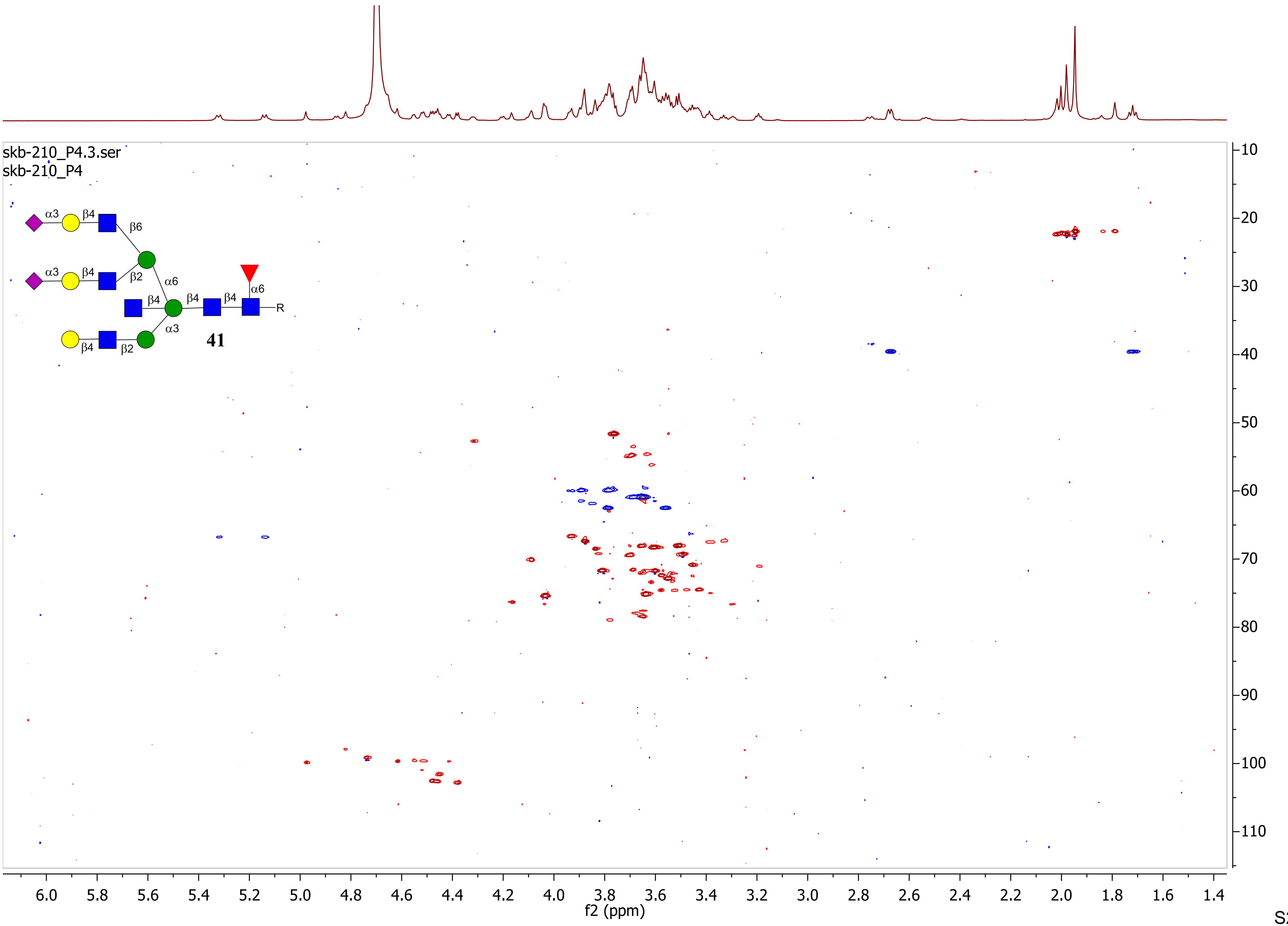
skb-210_P4.3.ser
skb-210_P4
α3
β4
β6
α3
β4
β2
α6
β4
β4
β4
α6
R
α3
β4
β2
41
f2 (ppm)
f1 (ppm)

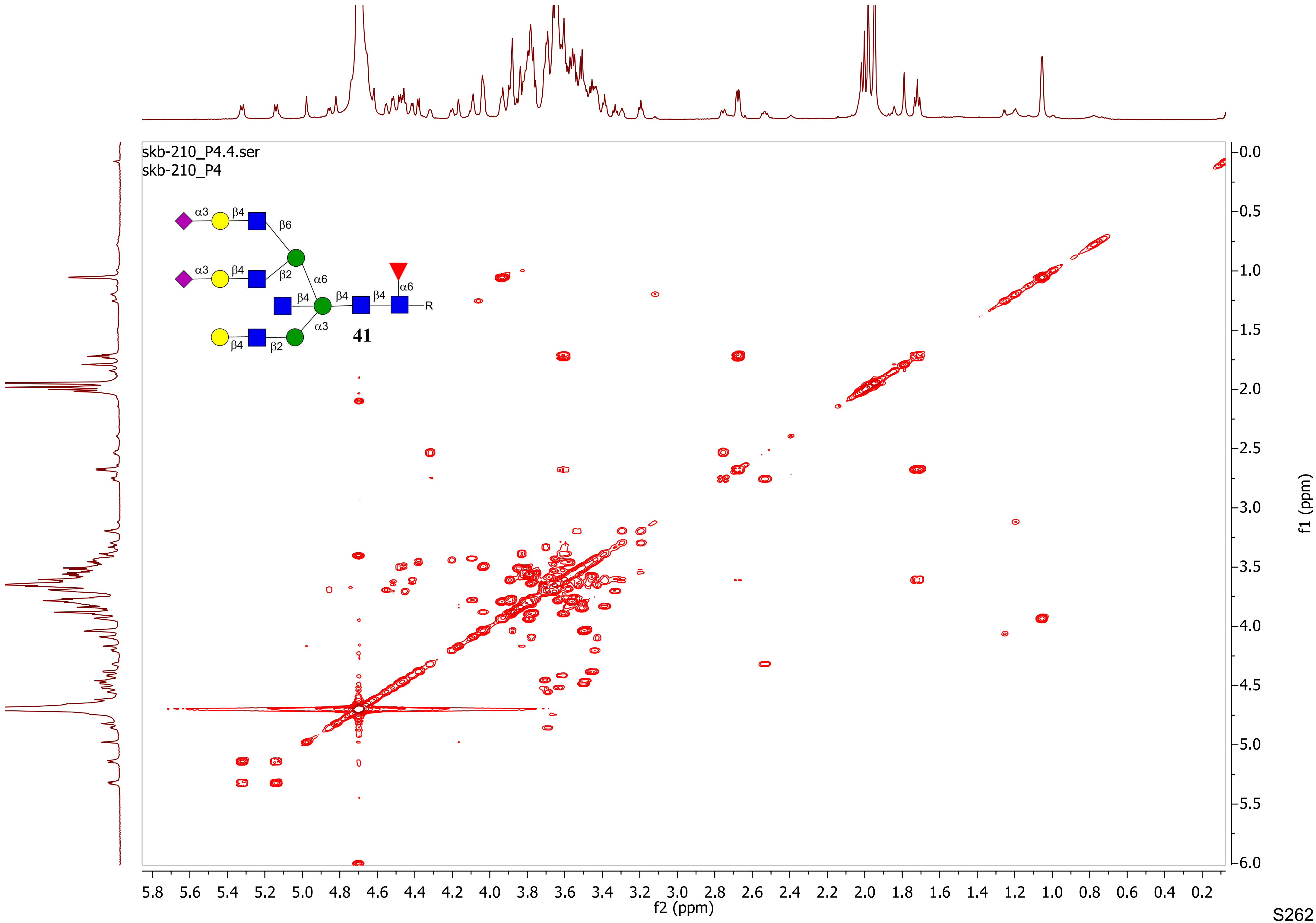
skb-210_P4.4.ser
skb-210_P4
α3
β4
β6
β2
α6
β4
β4
β4
α6
R
α3
β4
β2
41
f2 (ppm)
f1 (ppm)
5.8
5.6
5.4
5.2
5.0
4.8
4.6
4.4
4.2
4.0
3.8
3.6
3.4
3.2
3.0
2.8
2.6
2.4
2.2
2.0
1.8
1.6
1.4
1.2
1.0
0.8
0.6
0.4
0.2
0.0
0.5
1.0
1.5
2.0
2.5
3.0
3.5
4.0
4.5
5.0
5.5
6.0

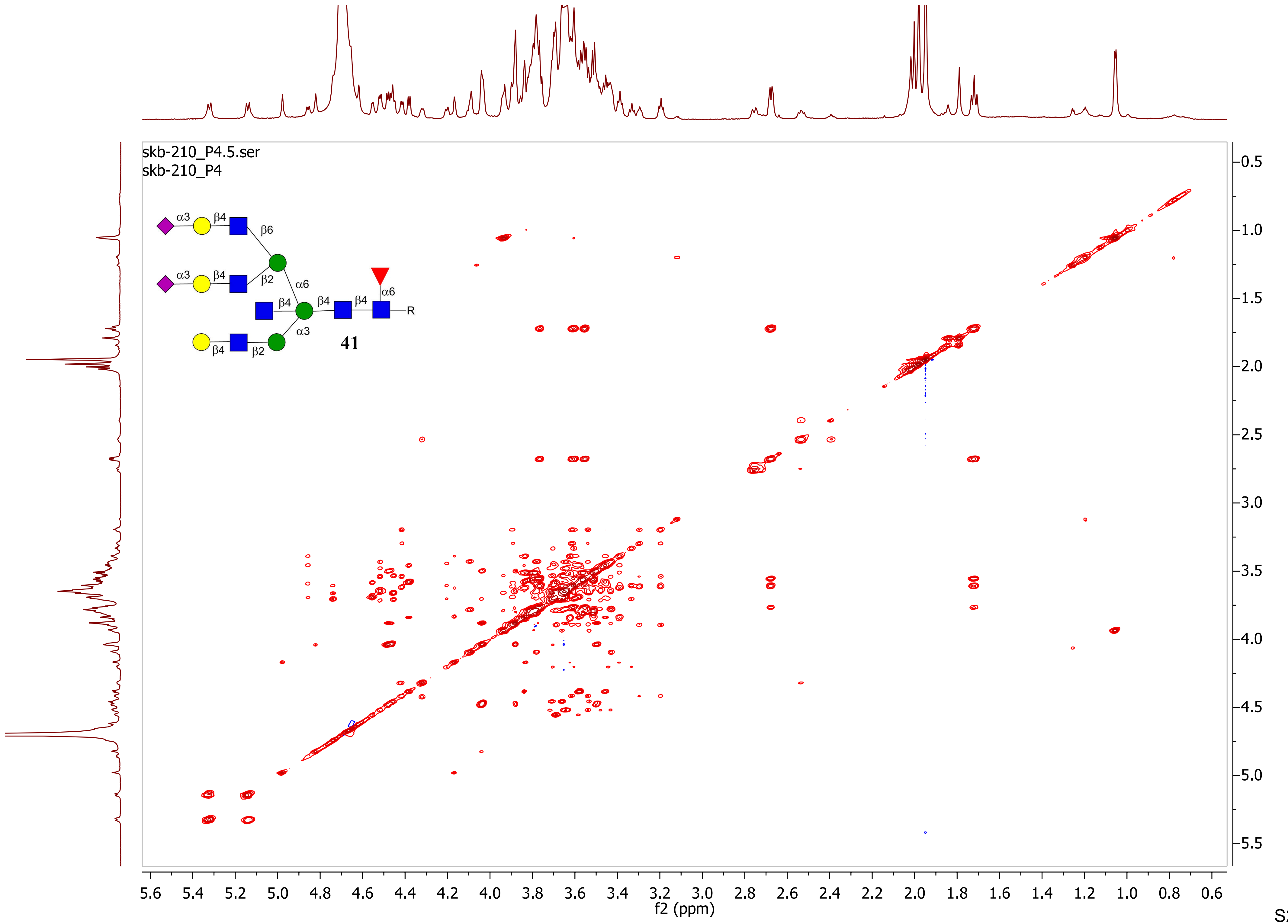
skb-210_P4.5.ser
skb-210_P4
α3
β4
β6
β2
α6
α3
β4
β4
β4
β4
α6
R
β4
β2
α3
41
f2 (ppm)
f1 (ppm)
5.6 5.4 5.2 5.0 4.8 4.6 4.4 4.2 4.0 3.8 3.6 3.4 3.2 3.0 2.8 2.6 2.4 2.2 2.0 1.8 1.6 1.4 1.2 1.0 0.8 0.6
0.5 1.0 1.5 2.0 2.5 3.0 3.5 4.0 4.5 5.0 5.5

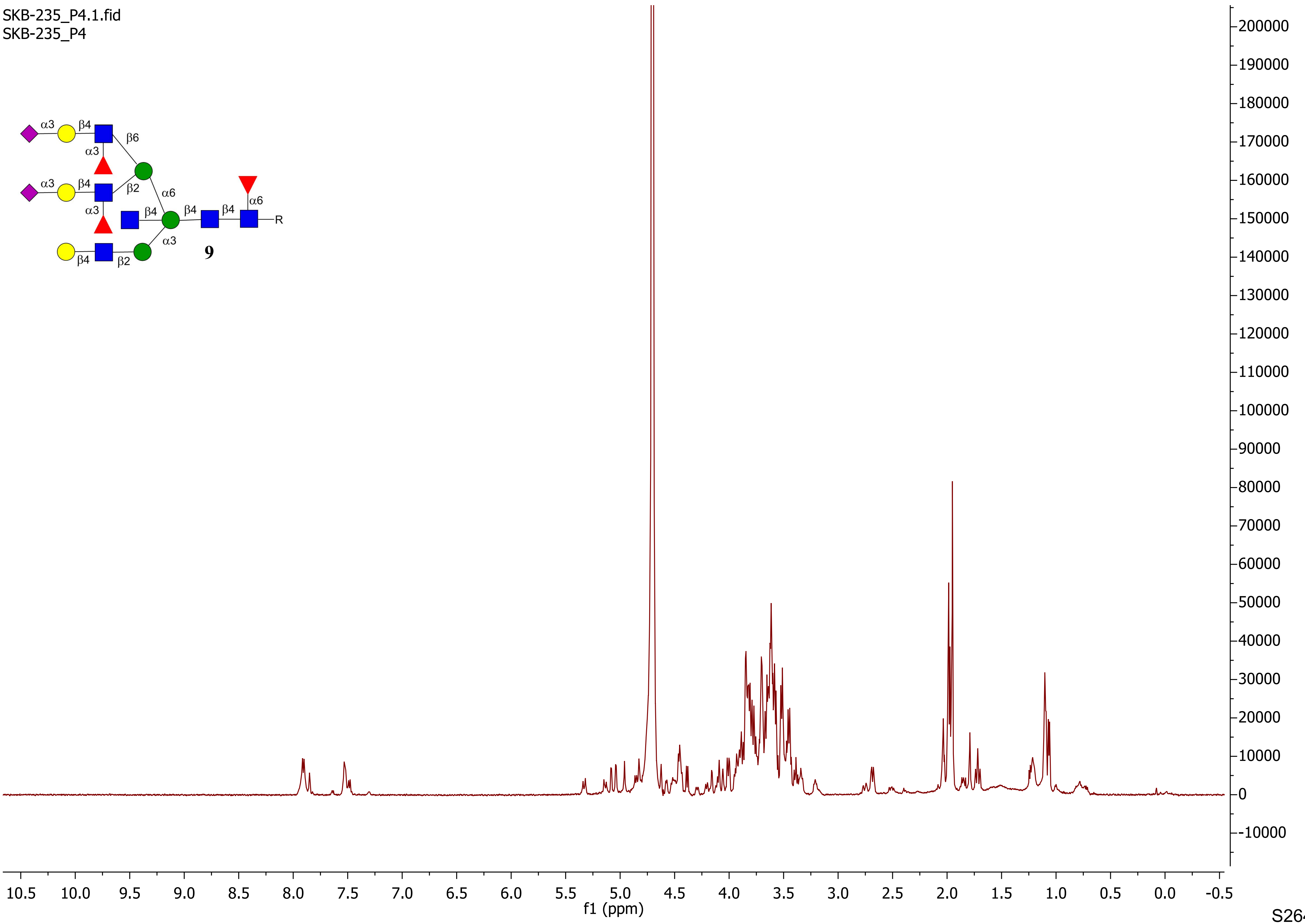
SKB-235_P4.1.fid
SKB-235_P4
α3
β4
β6
α3
α3
β4
β2
α6
α3
β4
β4
β4
α6
R
α3
β4
β2
9
200000
190000
180000
170000
160000
150000
140000
130000
120000
110000
100000
90000
80000
70000
60000
50000
40000
30000
20000
10000
0
-10000
10.5 10.0 9.5 9.0 8.5 8.0 7.5 7.0 6.5 6.0 5.5 5.0 4.5 4.0 3.5 3.0 2.5 2.0 1.5 1.0 0.5 0.0 -0.5
f1 (ppm)

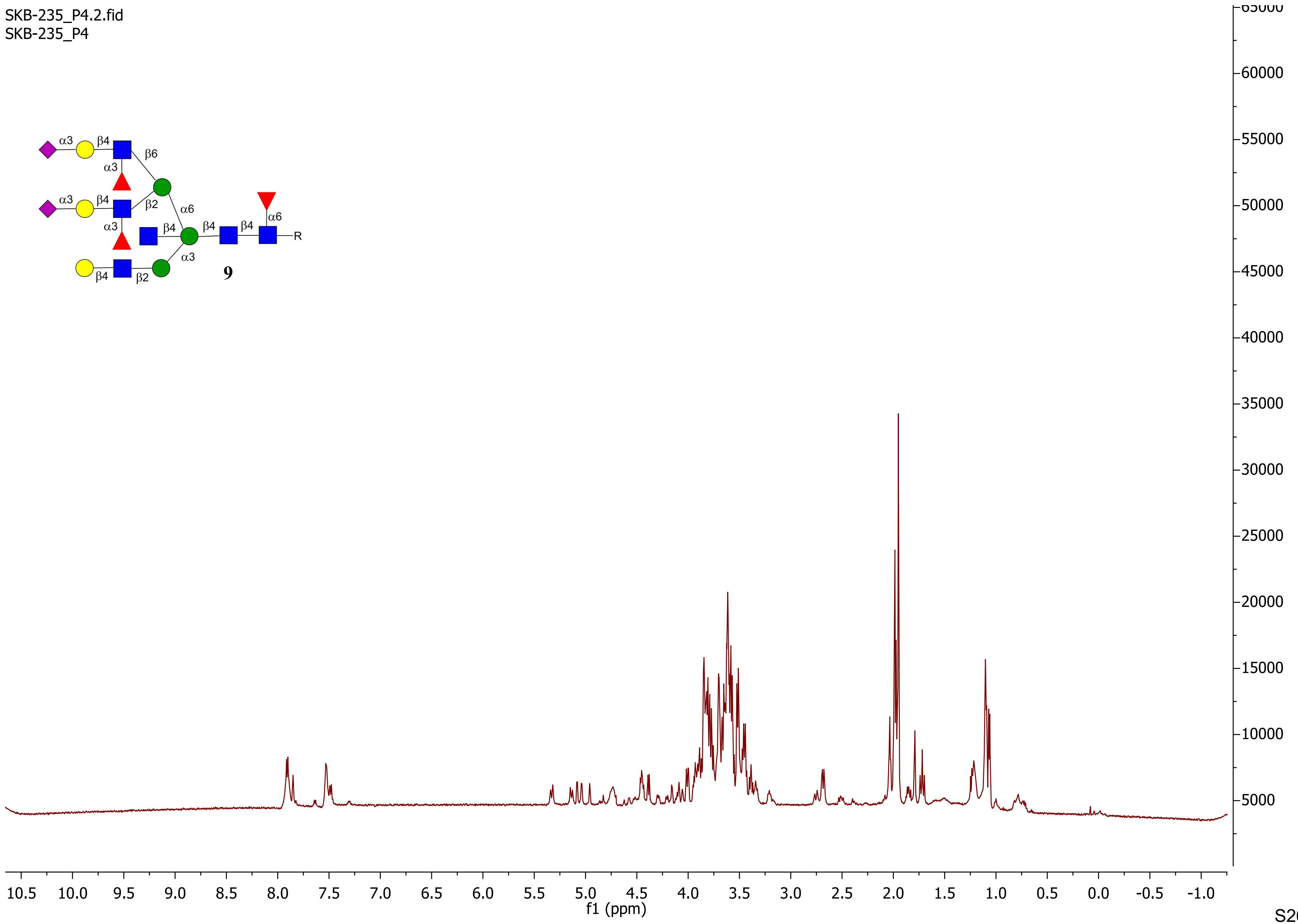
SKB-235_P4.2.fid
SKB-235_P4
α3
β4
β6
α3
α3
β4
β2
α6
α3
β4
β4
β4
α6
R
α3
β4
β2
9
65000
60000
55000
50000
45000
40000
35000
30000
25000
20000
15000
10000
5000
10.5 10.0 9.5 9.0 8.5 8.0 7.5 7.0 6.5 6.0 5.5 5.0 4.5 4.0 3.5 3.0 2.5 2.0 1.5 1.0 0.5 0.0 -0.5 -1.0
f1 (ppm)

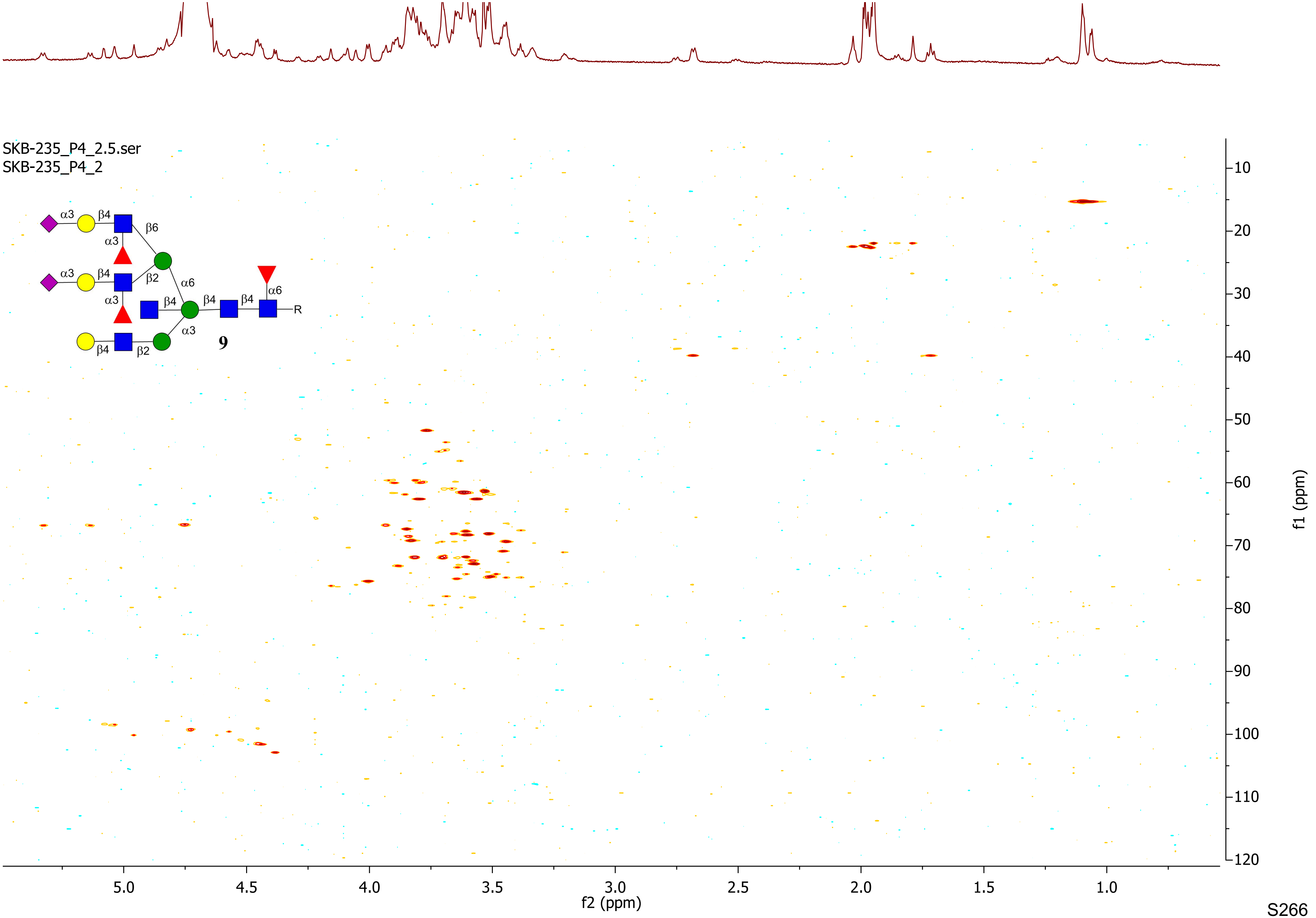

SKB-235_P4_2.5.ser
SKB-235_P4_2
α3
β4
β6
α3
α3
β4
β2
α6
α3
β4
β4
β4
α6
R
α3
β4
β2
9
f1 (ppm)
f2 (ppm)

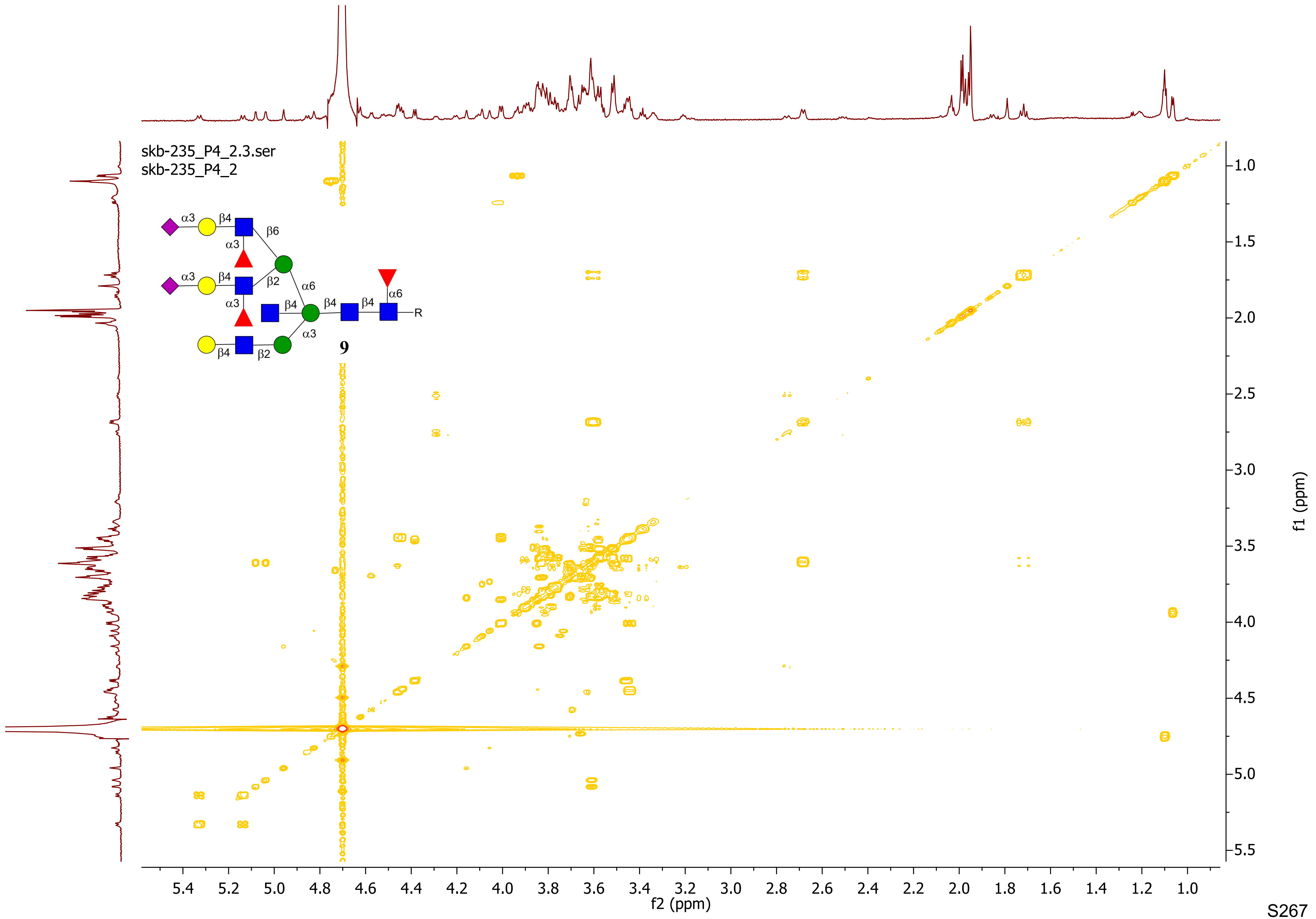
skb-235_P4_2.3.ser
skb-235_P4_2
α3
β4
β6
α3
α3
β4
β2
α6
α3
β4
β4
β4
α6
R
α3
β4
β2
9
f2 (ppm)
f1 (ppm)

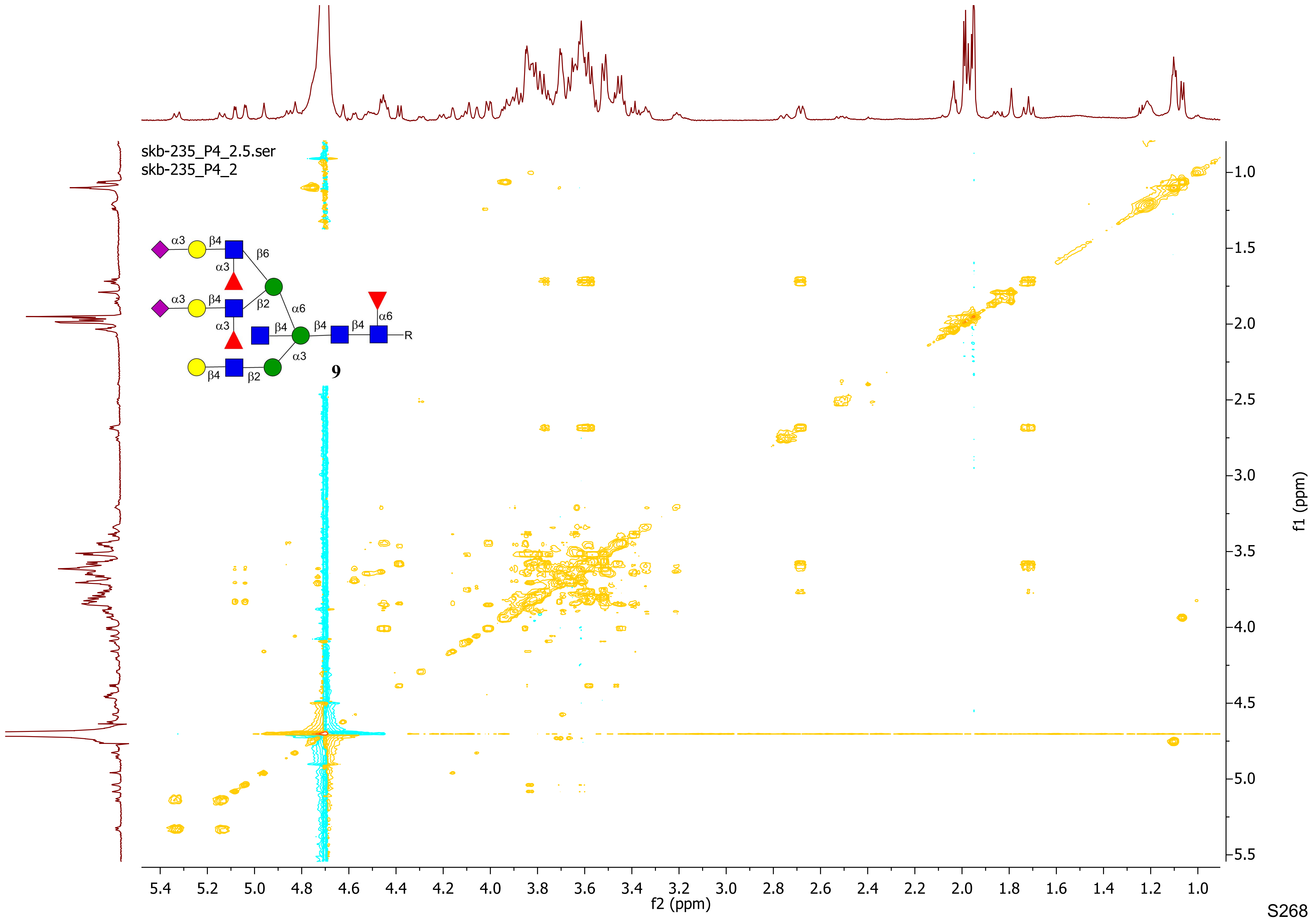
skb-235_P4_2.5.ser
skb-235_P4_2
α3
β4
β6
α3
β2
α6
β4
β4
β4
α6
R
α3
β4
β2
9
f2 (ppm)
f1 (ppm)

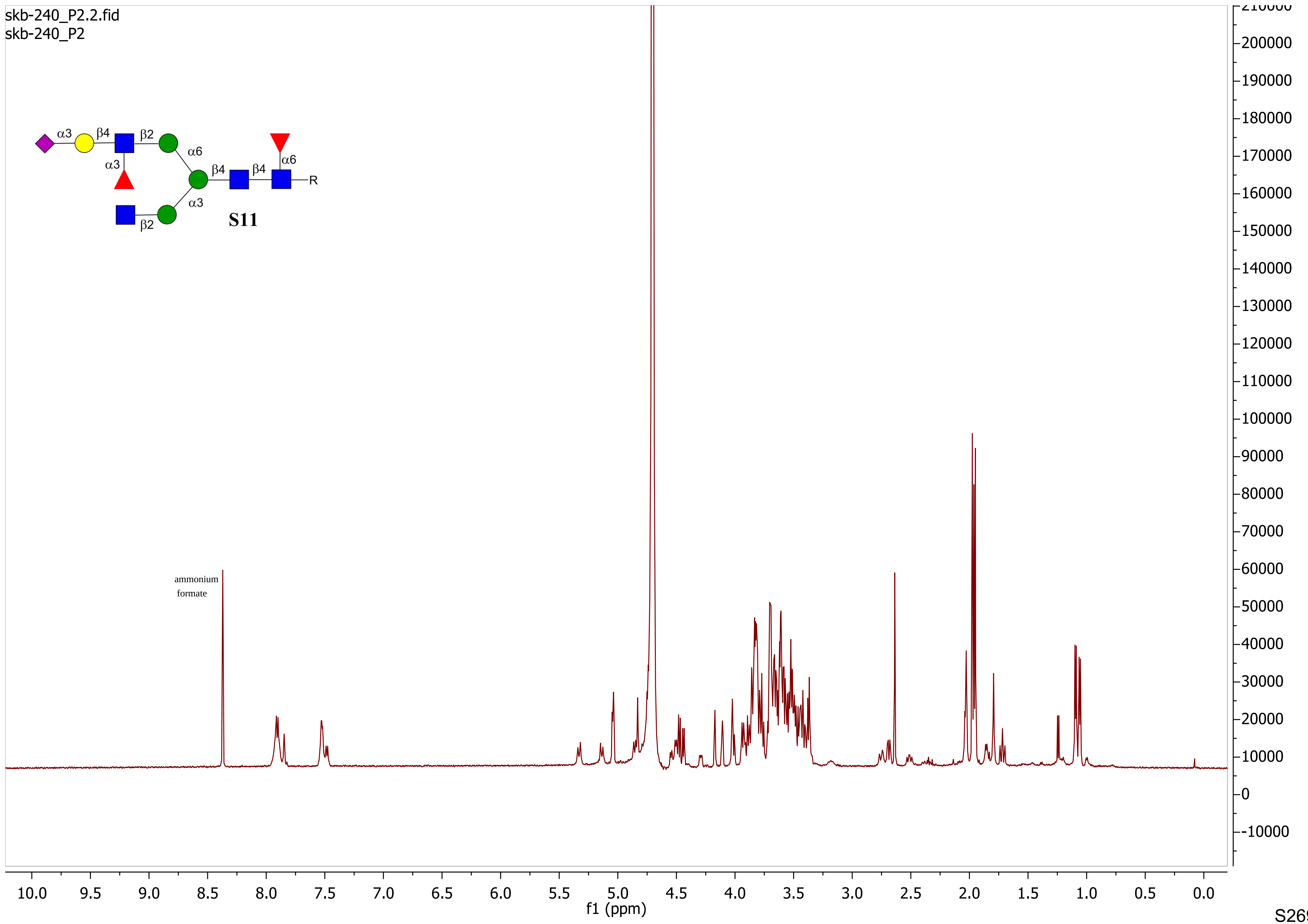
skb-240_P2.2.fid
skb-240_P2
α3 β4 β2 α6 β4 β4 α6 α3 α3 β2
R
S11
ammonium formate
f1 (ppm)

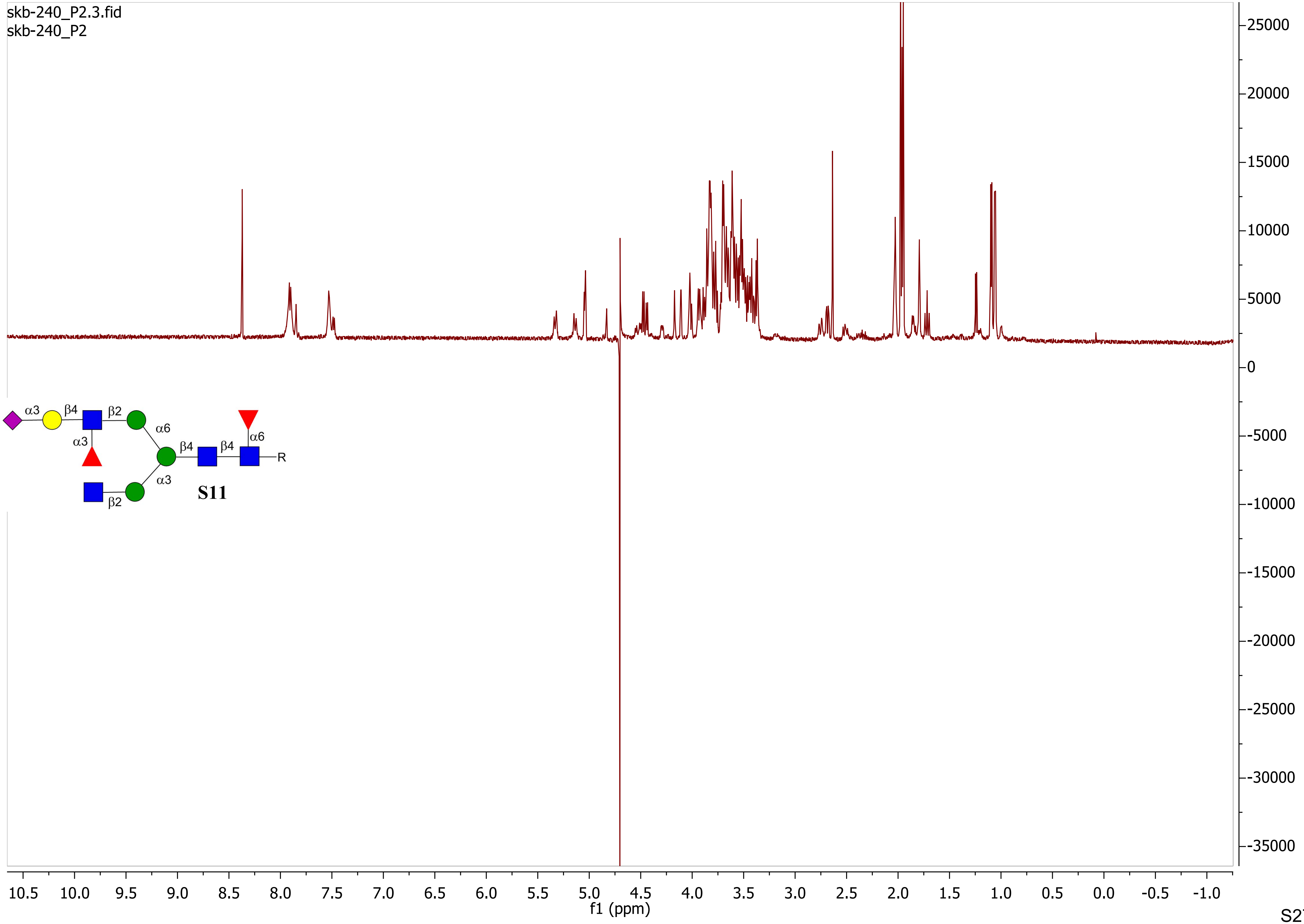

skb-240_P2.3.fid
skb-240_P2
α3
β4
β2
α6
α3
β4
β4
α6
R
α3
β2
S11
f1 (ppm)

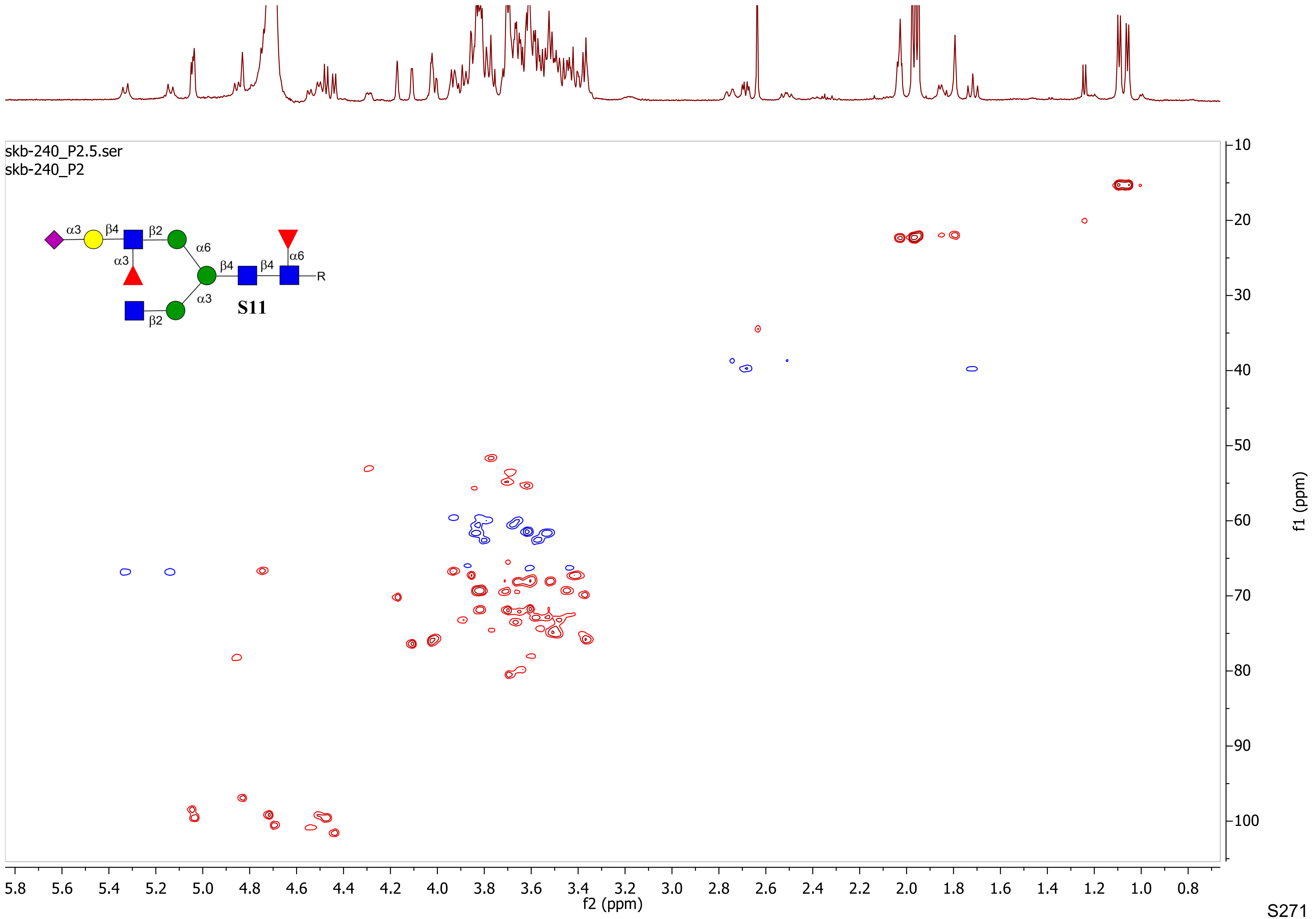

skb-240_P2.5.ser
skb-240_P2
α3
β4
β2
α6
α3
β4
β4
α6
R
α3
β2
S11
f1 (ppm)
f2 (ppm)

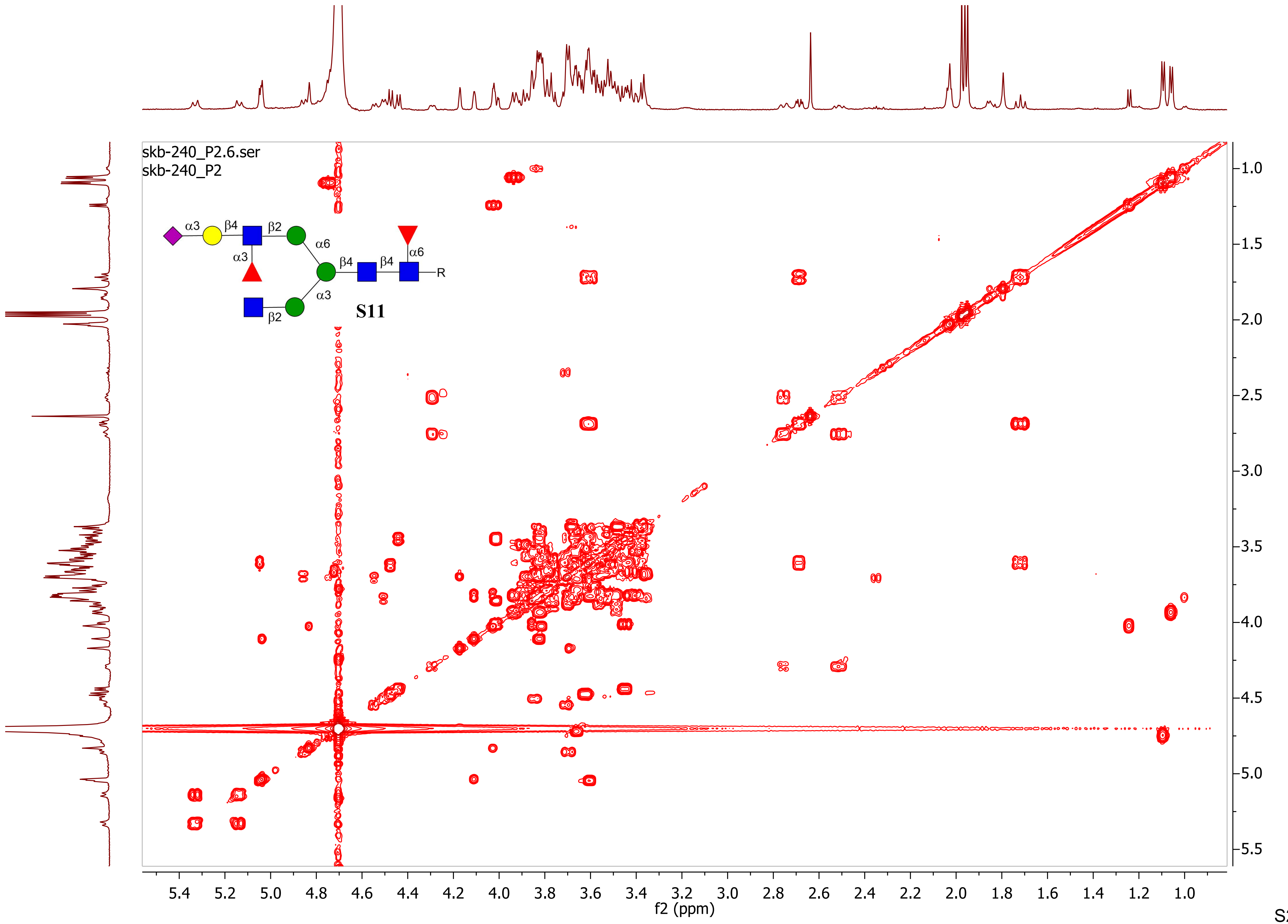

skb-240_P2.6.ser
skb-240_P2
α3
β4
β2
α6
α3
β4
β4
α6
R
α3
β2
S11
5.4
5.2
5.0
4.8
4.6
4.4
4.2
4.0
3.8
3.6
3.4
3.2
3.0
2.8
2.6
2.4
2.2
2.0
1.8
1.6
1.4
1.2
1.0
f2 (ppm)
1.0
1.5
2.0
2.5
3.0
3.5
4.0
4.5
5.0
5.5
f1 (ppm)

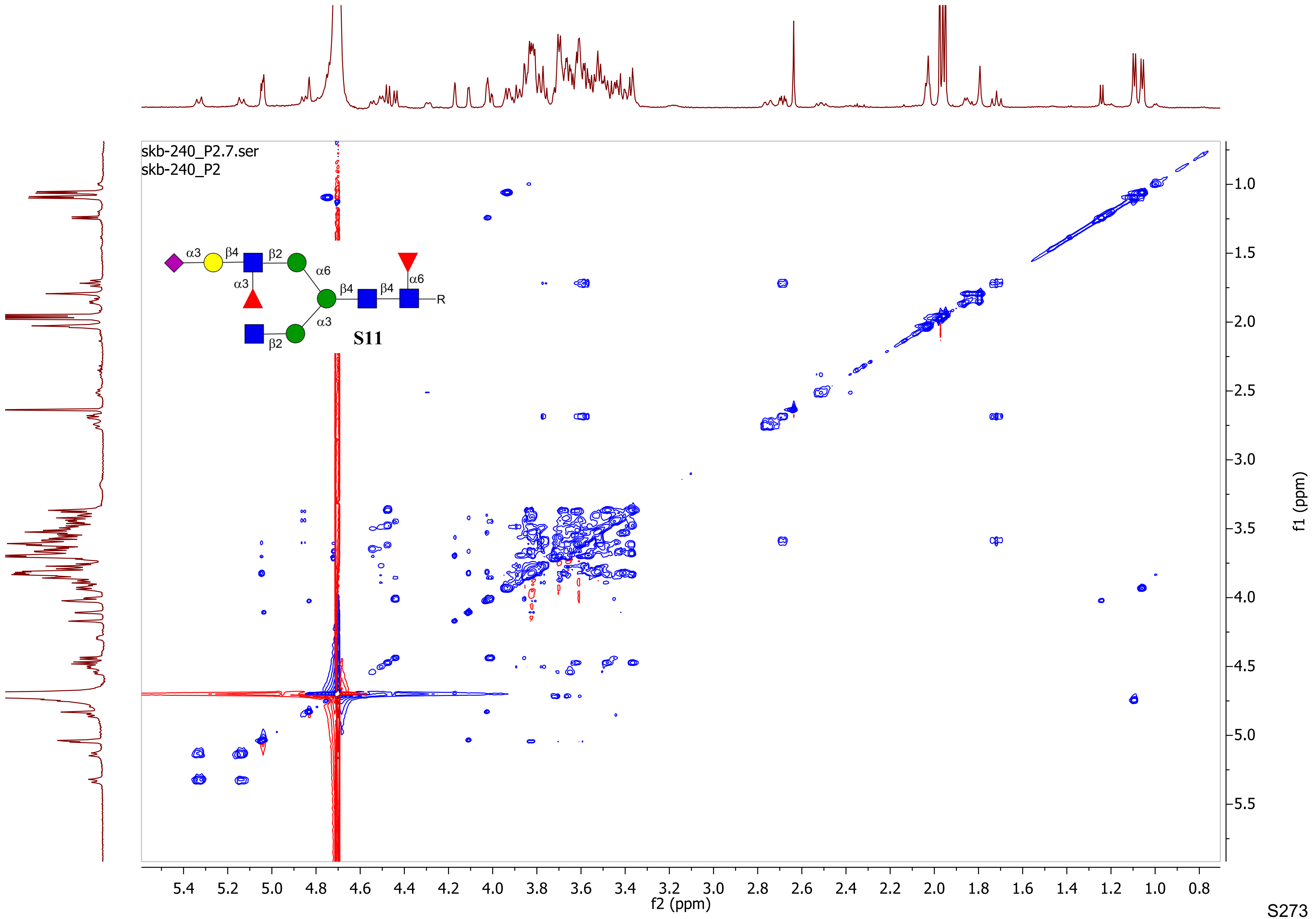

skb-240_P2.7.ser
skb-240_P2
α3
β4
β2
α6
α3
β4
β4
α6
R
α3
β2
S11
f2 (ppm)
f1 (ppm)

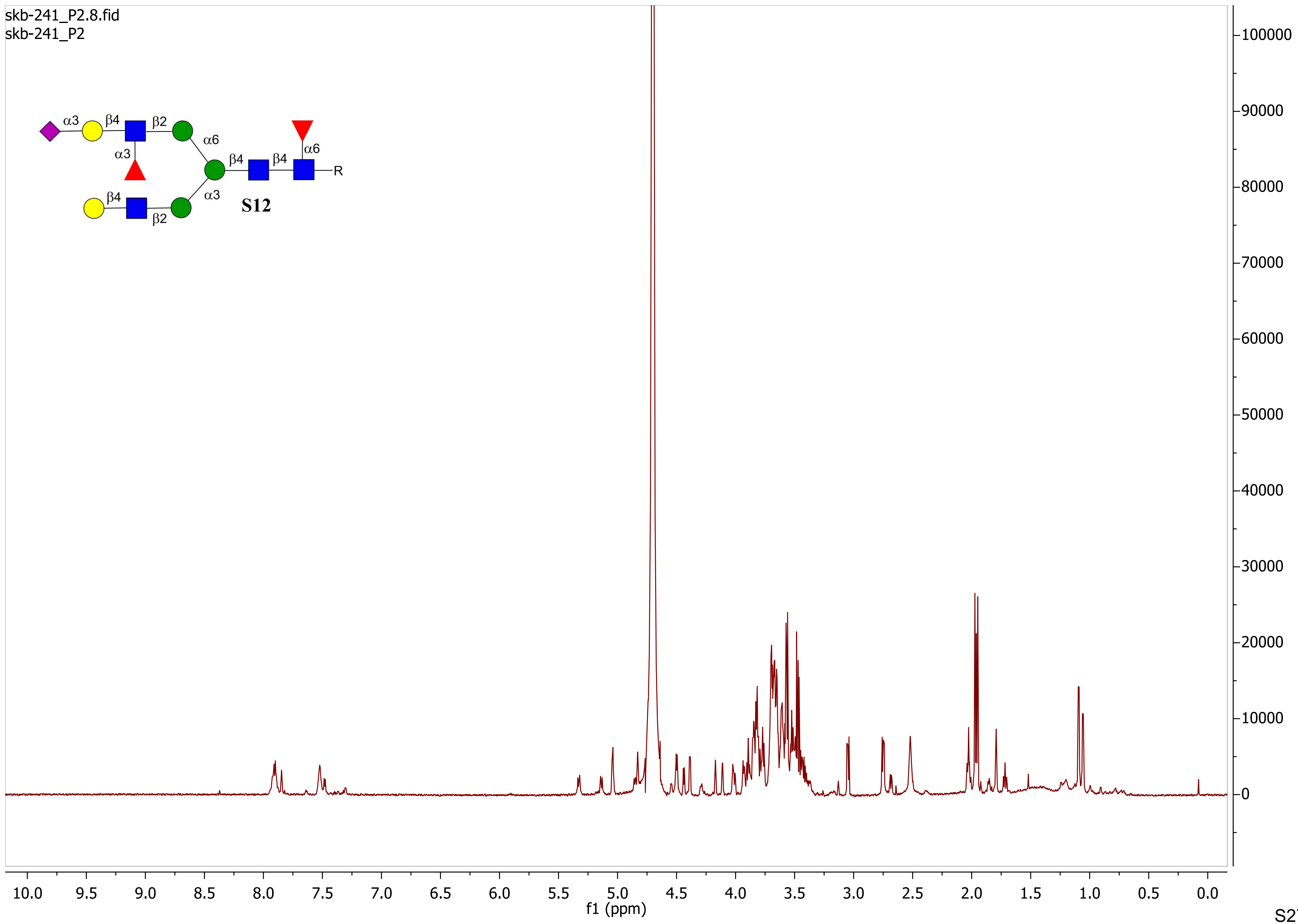
skb-241_P2.8.fid
skb-241_P2
α3
β4
β2
α6
α3
β4
β4
α6
R
β4
α3
β2
S12
100000
90000
80000
70000
60000
50000
40000
30000
20000
10000
0
10.0 9.5 9.0 8.5 8.0 7.5 7.0 6.5 6.0 5.5 5.0 4.5 4.0 3.5 3.0 2.5 2.0 1.5 1.0 0.5 0.0
f1 (ppm)

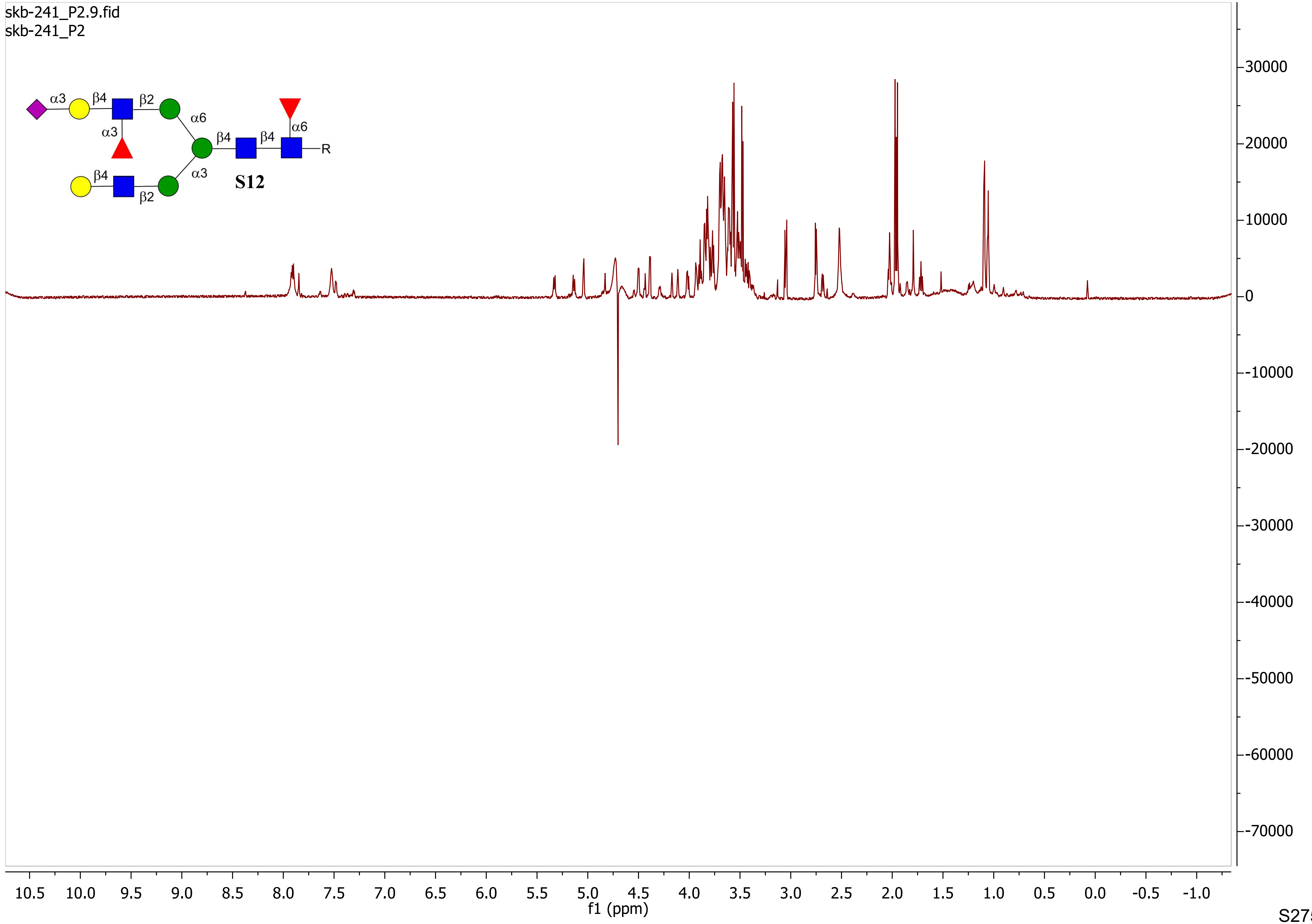
skb-241_P2.9.fid
skb-241_P2
α3
β4
β2
α6
α3
β4
β4
α6
R
β4
α3
β2
S12
f1 (ppm)

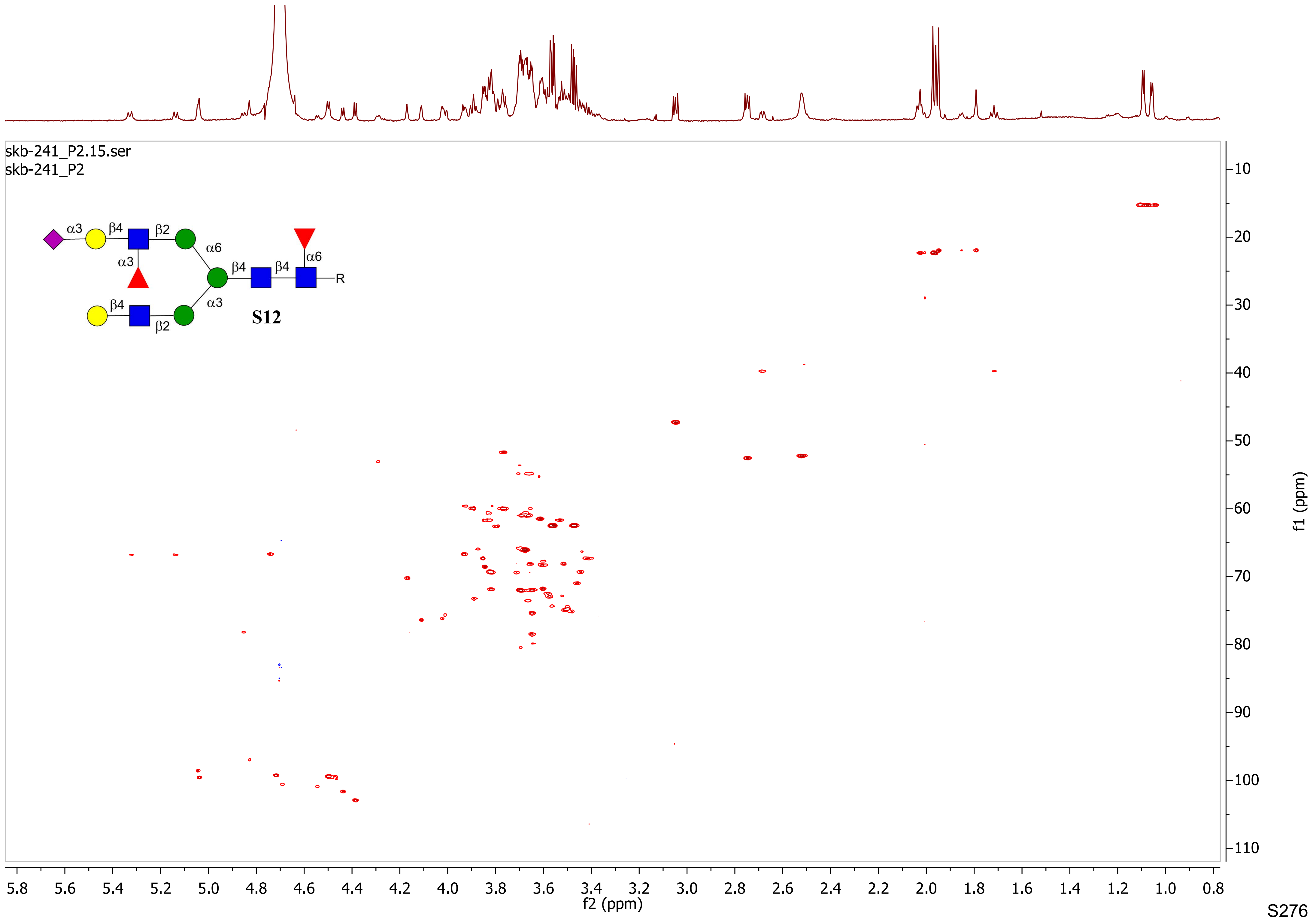
skb-241_P2.15.ser
skb-241_P2
α3
β4
β2
α6
α3
β4
β4
α6
R
β4
β2
α3
S12
f2 (ppm)
f1 (ppm)
5.8 5.6 5.4 5.2 5.0 4.8 4.6 4.4 4.2 4.0 3.8 3.6 3.4 3.2 3.0 2.8 2.6 2.4 2.2 2.0 1.8 1.6 1.4 1.2 1.0 0.8
10 20 30 40 50 60 70 80 90 100 110

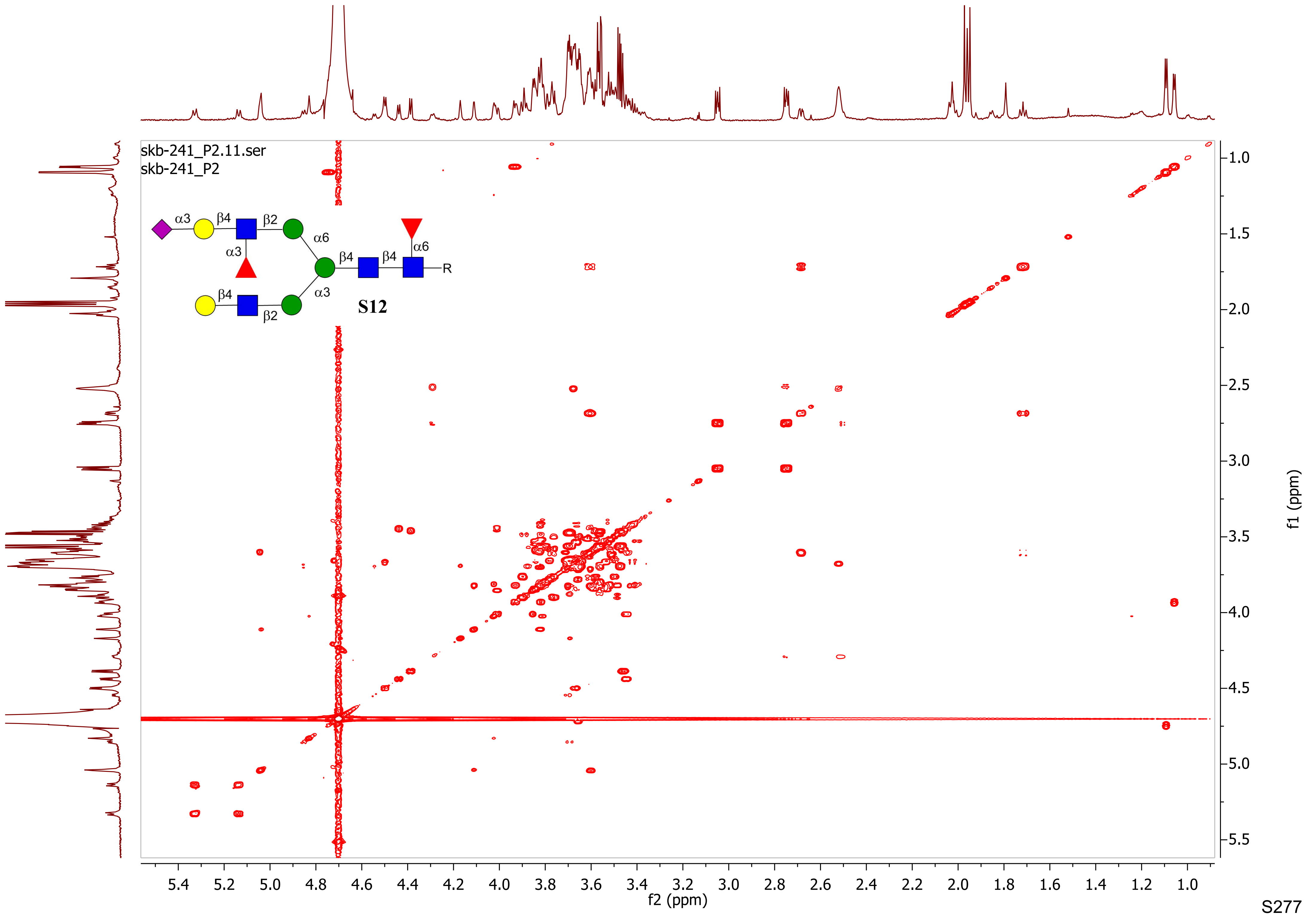
skb-241_P2.11.ser
skb-241_P2
α3
β4
β2
α6
α3
β4
β4
α6
R
α3
β4
β2
S12
f2 (ppm)
f1 (ppm)

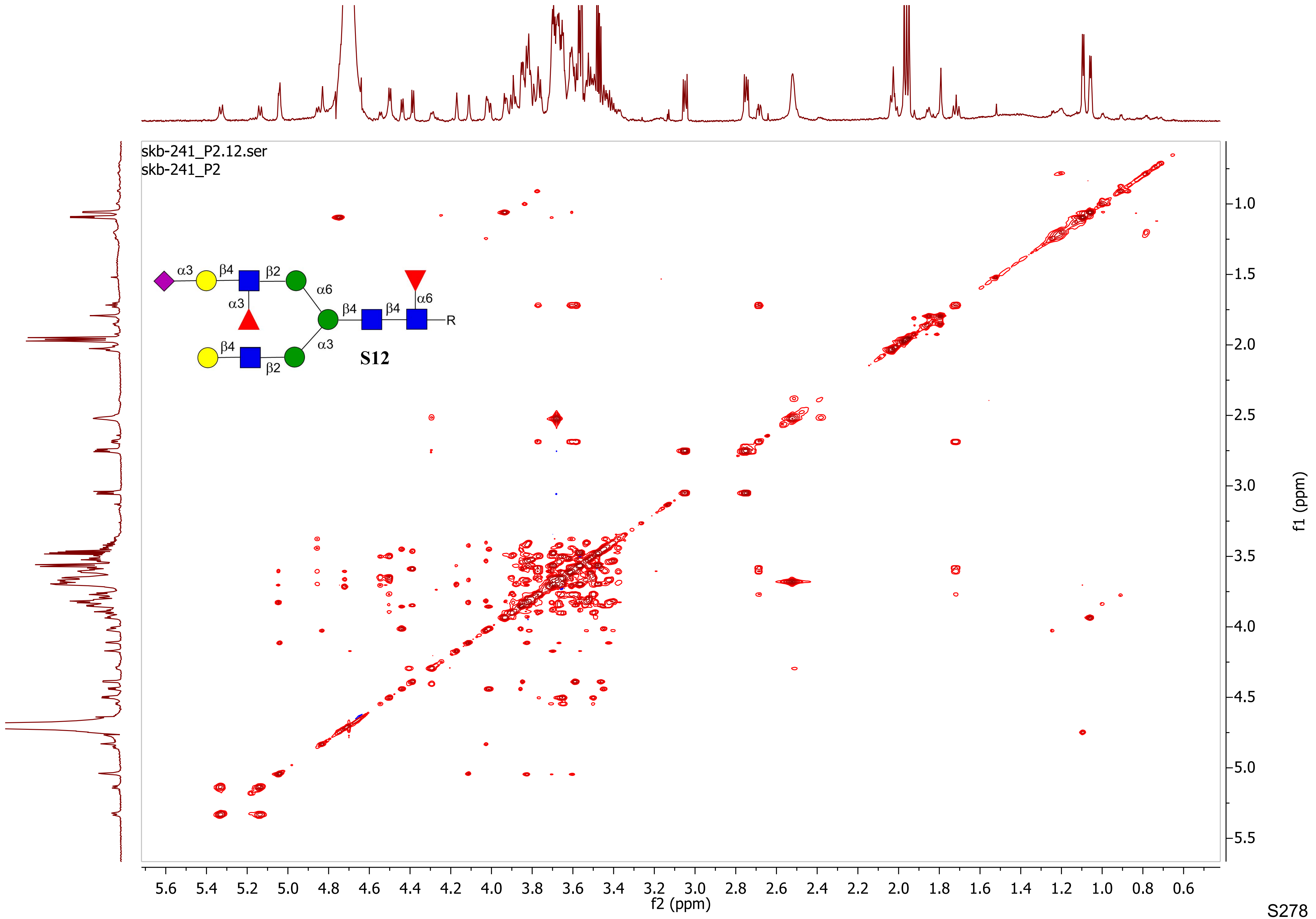

skb-241_P2.12.ser
skb-241_P2
α3
β4
β2
α6
α3
β4
β4
α6
R
β4
α3
β2
S12
5.6
5.4
5.2
5.0
4.8
4.6
4.4
4.2
4.0
3.8
3.6
3.4
3.2
3.0
2.8
2.6
2.4
2.2
2.0
1.8
1.6
1.4
1.2
1.0
0.8
0.6
f2 (ppm)
1.0
1.5
2.0
2.5
3.0
3.5
4.0
4.5
5.0
5.5
f1 (ppm)

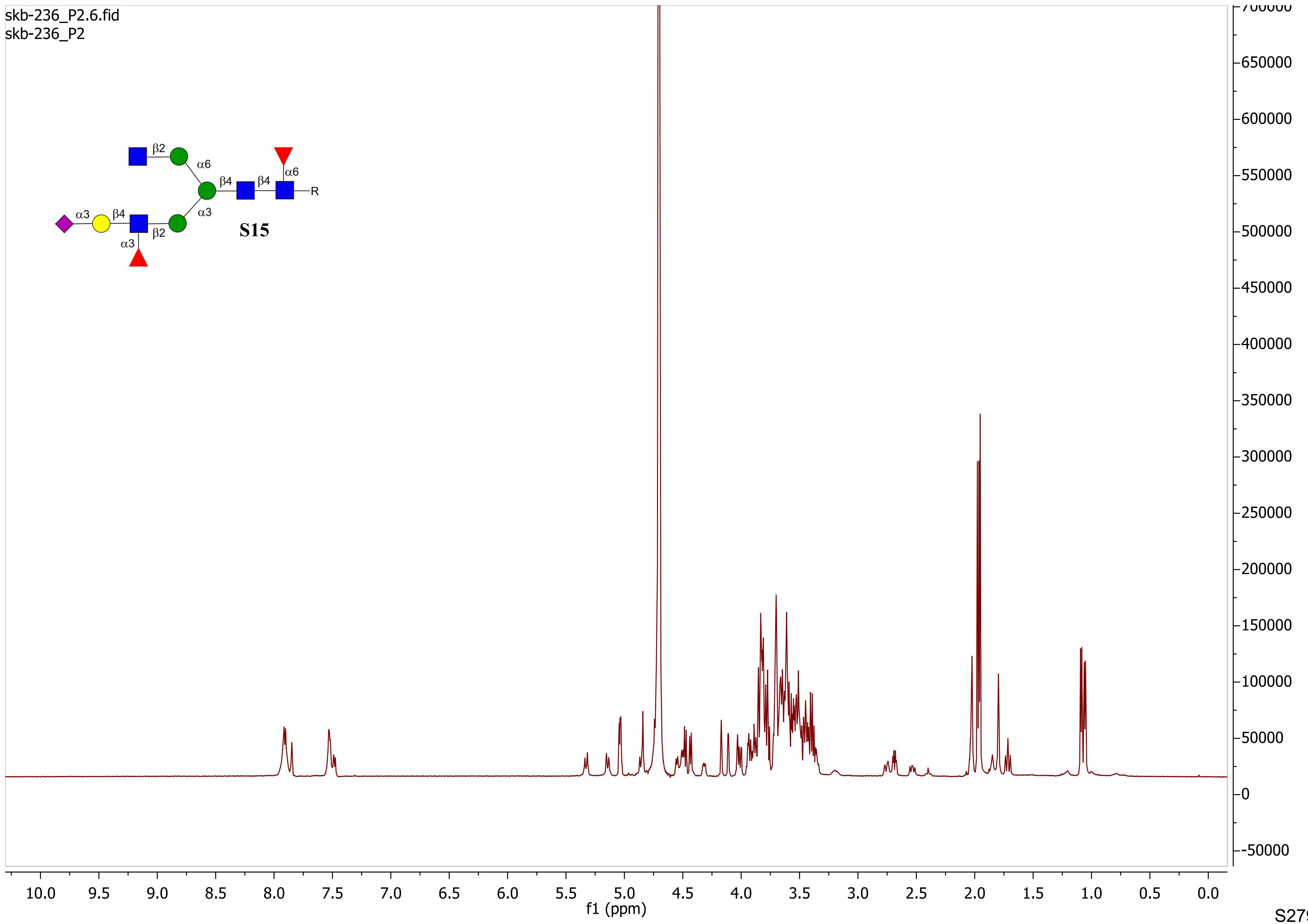
skb-236_P2.6.fid
skb-236_P2
β2
α6
β4
β4
α6
R
α3
β4
α3
β2
α3
S15
f1 (ppm)

skb-236_P2.7.fid
skb-236_P2

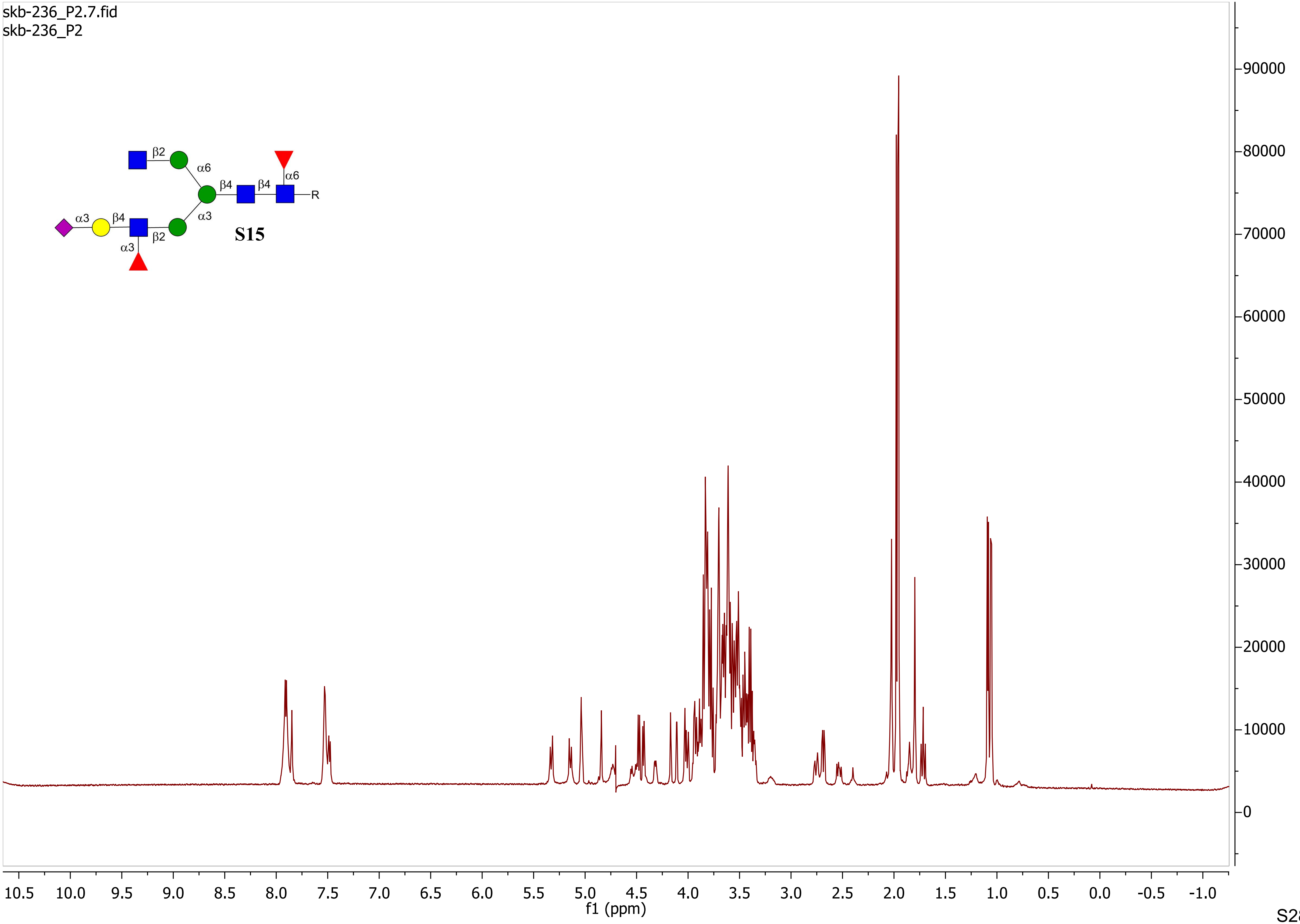

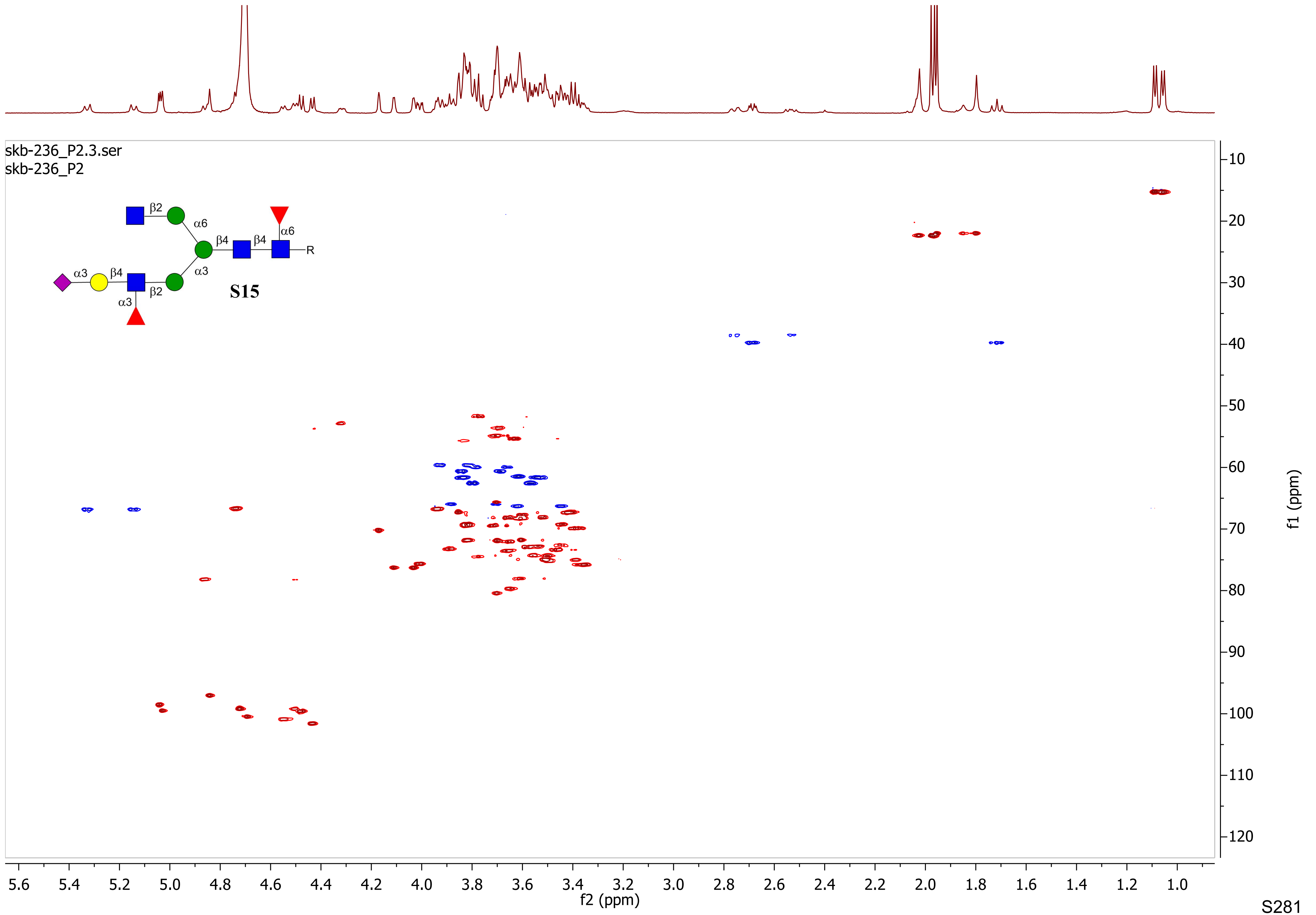
skb-236_P2.3.ser
skb-236_P2
β2
α6
β4
β4
α6
R
α3
β4
α3
β2
α3
S15
f2 (ppm)
f1 (ppm)

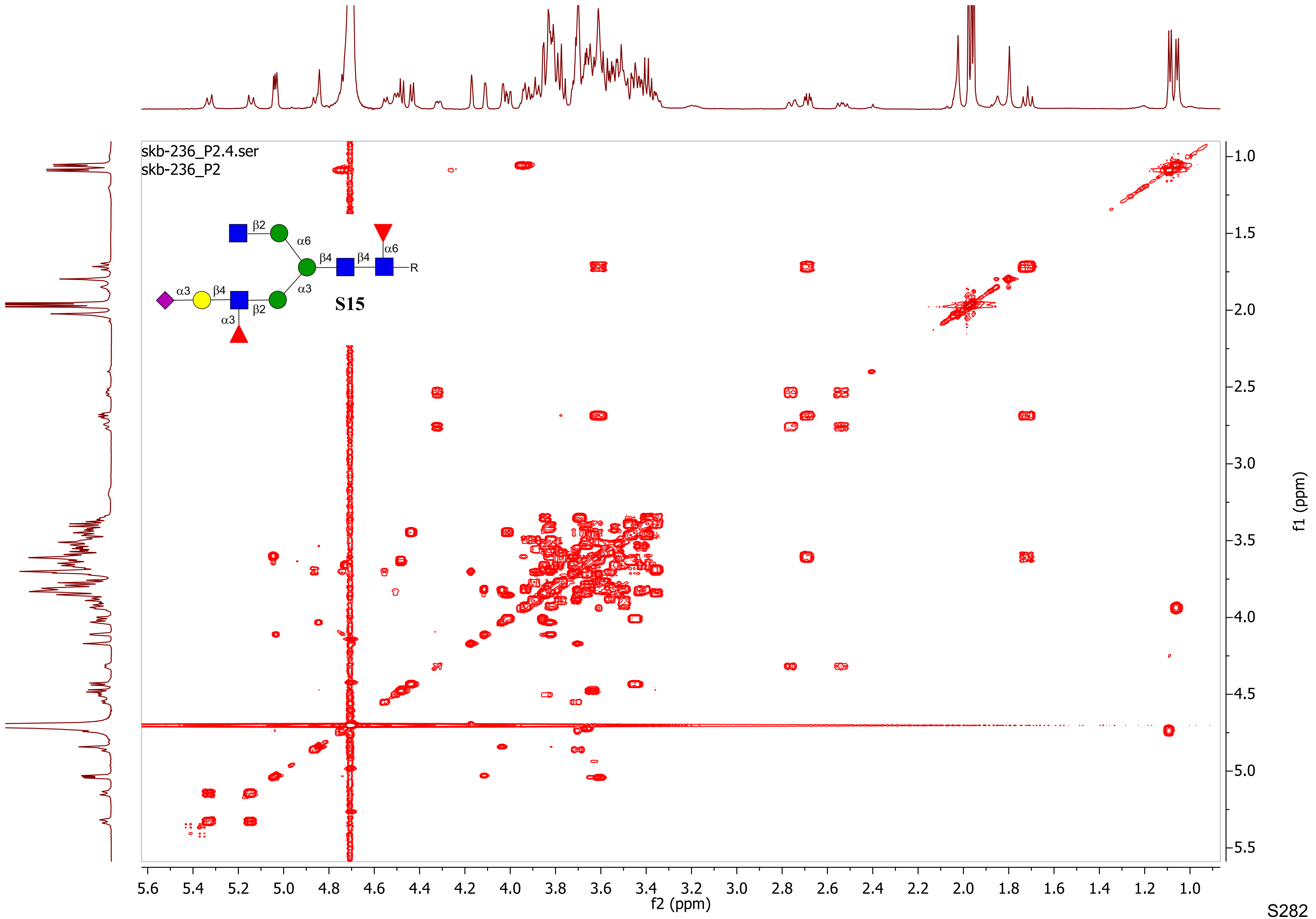

skb-236_P2.4.ser
skb-236_P2
β2
α6
β4
β4
α6
R
α3
β4
α3
β2
α3
S15
f2 (ppm)
f1 (ppm)

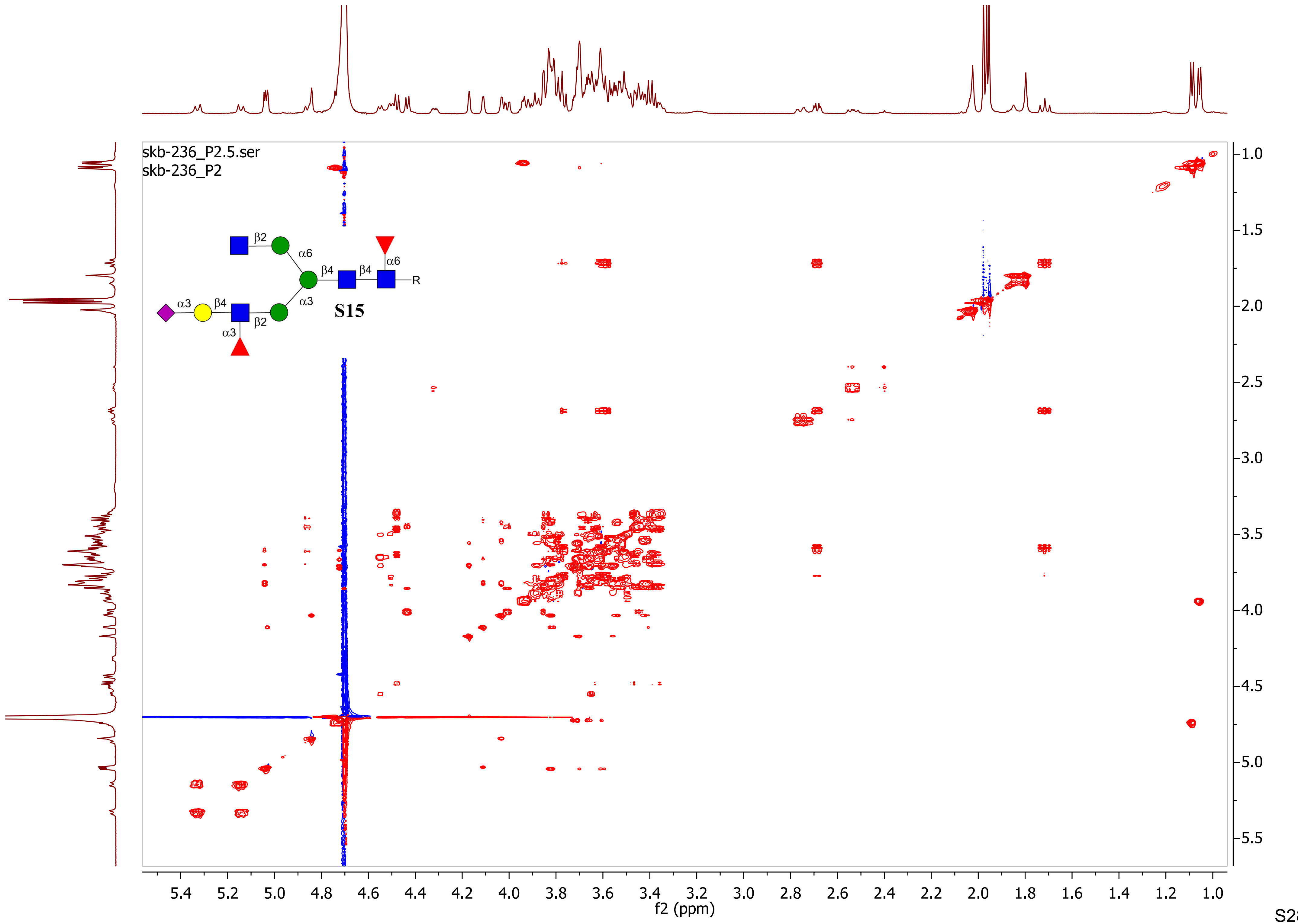
skb-236_P2.5.ser
skb-236_P2
β2
α6
β4
β4
α6
R
α3
β4
α3
β2
α3
S15
f2 (ppm)
f1 (ppm)

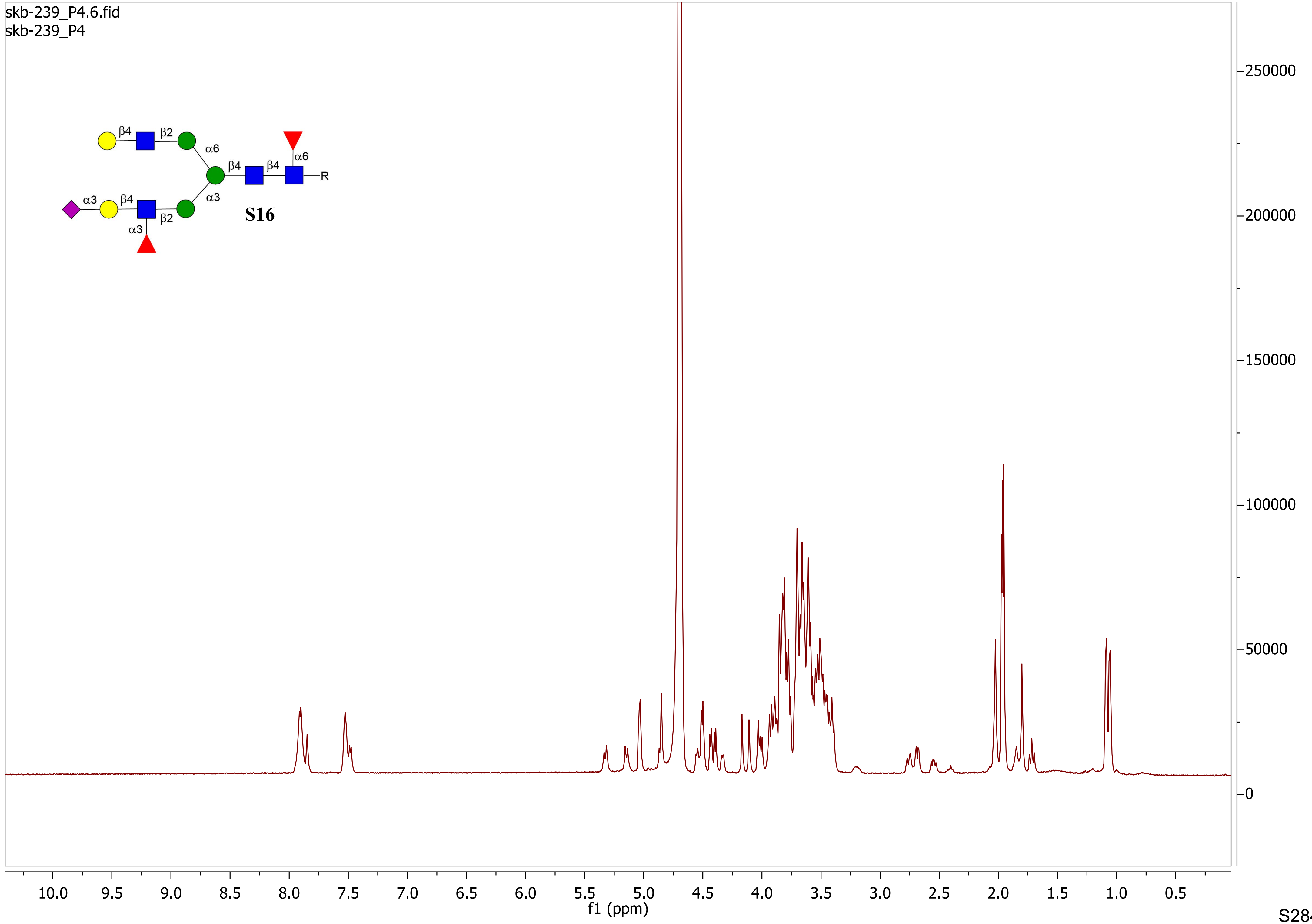
skb-239_P4.6.fid
skb-239_P4
β4
β2
α6
β4
β4
α6
R
α3
β4
α3
β2
α3
S16
250000
200000
150000
100000
50000
0
10.0
9.5
9.0
8.5
8.0
7.5
7.0
6.5
6.0
5.5
5.0
4.5
4.0
3.5
3.0
2.5
2.0
1.5
1.0
0.5
f1 (ppm)

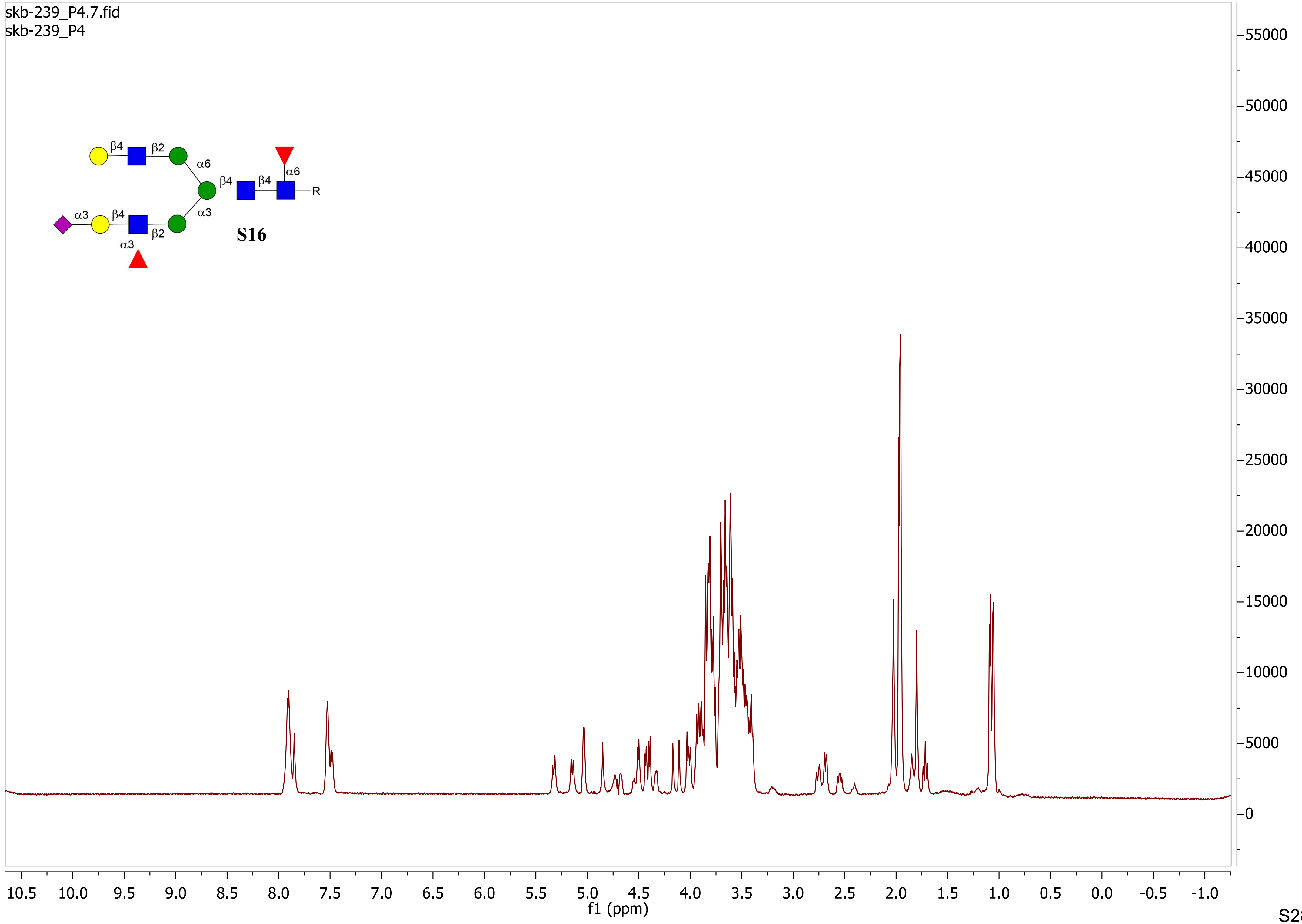
skb-239_P4.7.fid
skb-239_P4
β4
β2
α6
β4
β4
α6
R
α3
β4
α3
β2
α3
S16
f1 (ppm)

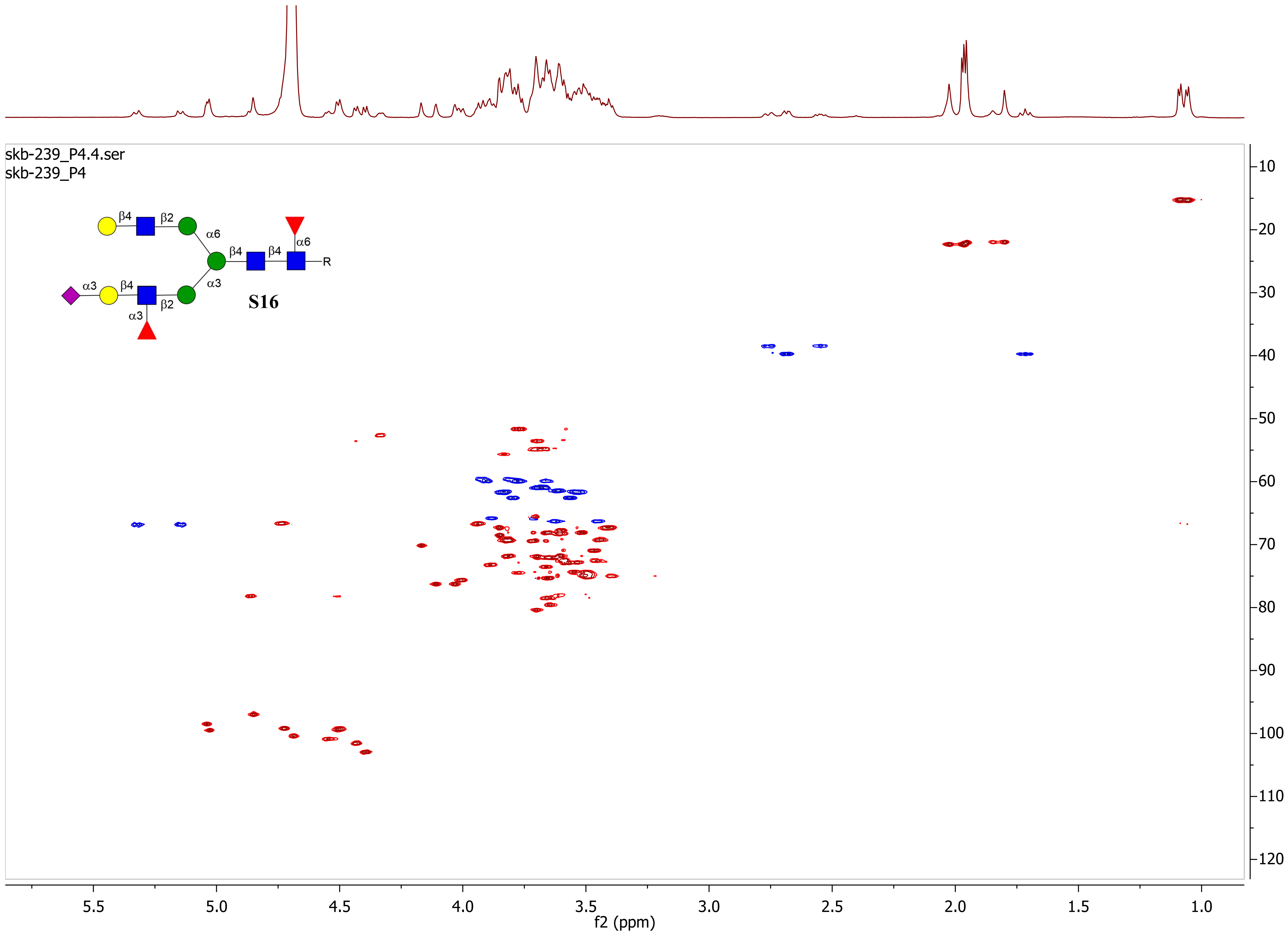
skb-239_P4.4.ser
skb-239_P4
β4
β2
α6
β4
β4
α6
R
α3
β4
α3
β2
α3
S16
f1 (ppm)
f2 (ppm)

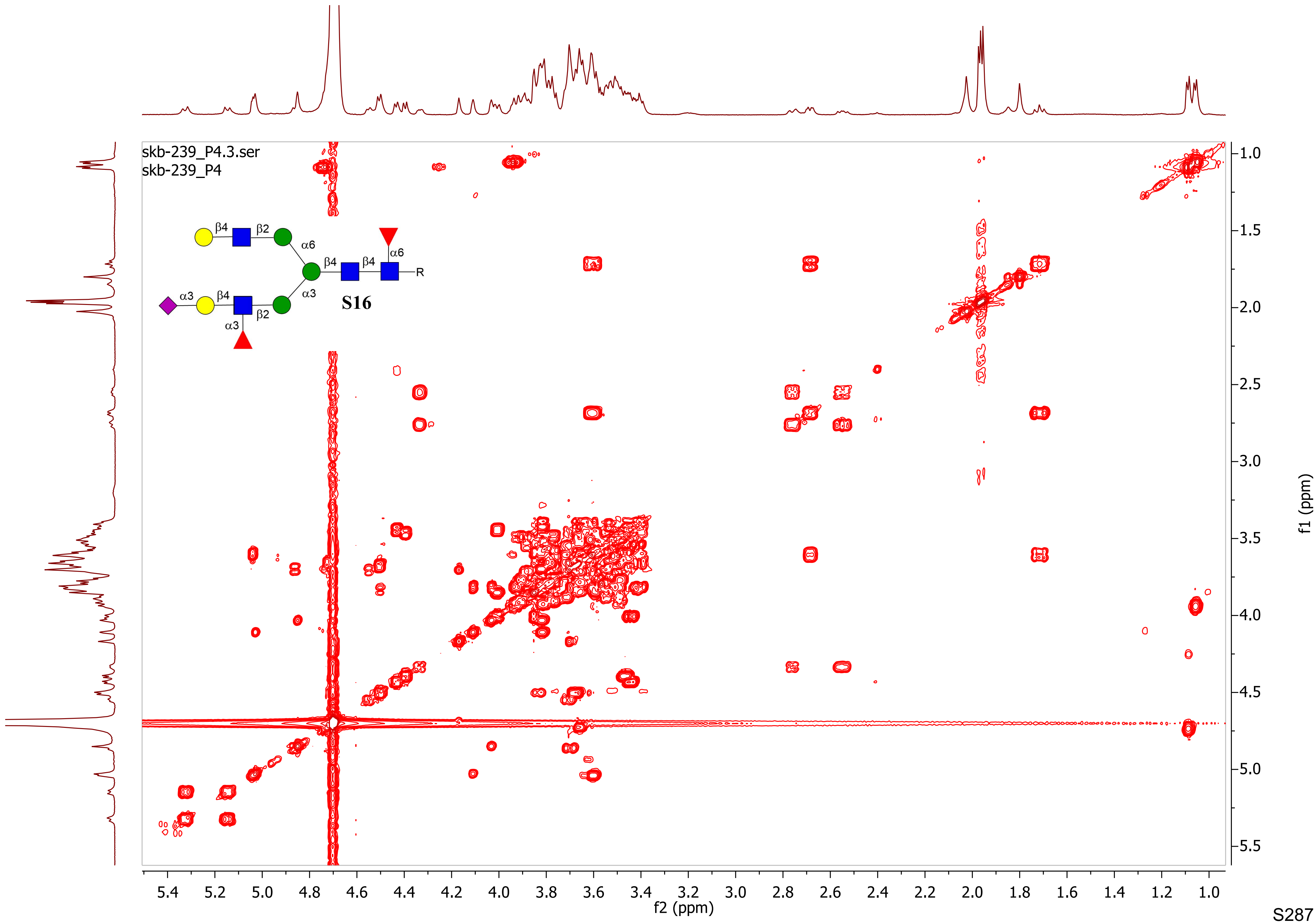
skb-239_P4.3.ser
skb-239_P4
β4
β2
α6
β4
β4
α6
R
α3
β4
α3
β2
α3
S16
f1 (ppm)
f2 (ppm)

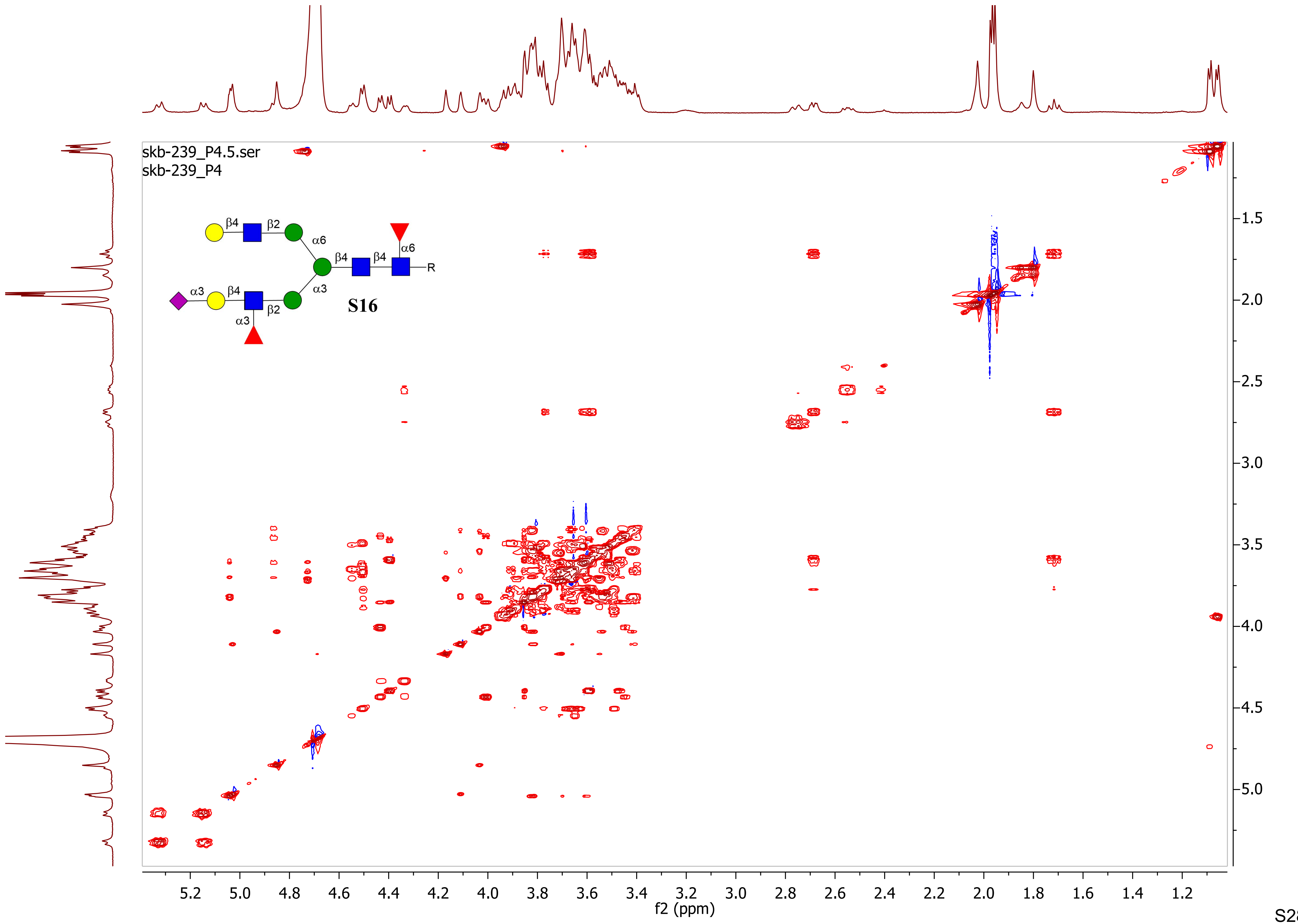

skb-239_P4.5.ser
skb-239_P4
β4
β2
α6
β4
β4
α6
R
α3
β4
α3
β2
α3
S16
f2 (ppm)
f1 (ppm)
